# Diel changes in the expression of a marker gene and candidate genes for intracellular amorphous CaCO_3_ biomineralization in *Microcystis*

**DOI:** 10.1101/2024.07.07.602159

**Authors:** Apolline Bruley, Juliette Gaëtan, Muriel Gugger, Claire Pancrace, Maxime Millet, Geoffroy Gaschignard, Manuela Dezi, Jean-François Humbert, Julie Leloup, Fériel Skouri-Panet, Isabelle Callebaut, Karim Benzerara, Elodie Duprat

## Abstract

Phylogenetically diverse cyanobacteria biomineralize intracellular amorphous calcium carbonate (iACC) inclusions. This includes several genotypes of the *Microcystis* genus, a potentially toxic, bloom-forming cyanobacterium found worldwide in freshwater ecosystems. While we ignore the biological function of iACC and the molecular mechanisms driving their formation, this process may impact local geochemical cycles and/or be used for bioremediation strategies. Recently, a marker gene of this biomineralization pathway, named *ccyA*, was discovered. However, the function of the calcyanin protein encoded by *ccyA* remains unknown. Here, based on an RNA-Seq approach, we assess the expression of the *ccyA* gene in *Microcystis aeruginosa* PCC 7806 during a 24 h day/night cycle. The *ccyA* gene shows a clear day/night expression pattern with maximum transcript abundances during the second half of the night. This is consistent with the assumption that iACC biomineralization is related with photosynthesis and may therefore follow a day/night cycle as well. Moreover, several genes directly co-localized upstream and downstream of *ccyA*, on the same DNA strand show a similar expression pattern, including a *cax* gene encoding a calcium/proton exchanger and a gene encoding a protein with a domain also present in the N-terminal region of calcyanins in many iACC-forming cyanobacteria. This suggests that they all could be part of an operon, and may play a concerted role in iACC formation. Last, several other genes involved in carbon concentrating mechanisms and calcium transport show an expression pattern similar to that of *ccyA*. Overall, this study provides a list of candidate genes that may be involved in the biomineralization of iACC by cyanobacteria and whose role could be, in the future, analyzed by biochemistry and genetics approaches.

## Introduction

Biomineralization is a process through which some living organisms form mineral phases. It is found in all the domains of life (Boskey, 2003). In some cases, biomineralization is genetically controlled, *i.e.* it is tightly controlled by specific genes, as documented in diverse eukaryotic organisms (Brownlee et al., 2015; Gilbert et al., 2022; Knoll, 2003; Marin & Luquet, 2004; Monteiro et al., 2016). For example, key genes encoding, *e.g.*, carbonic anhydrases (Chen et al., 2018; Le Roy et al., 2014), Ca²⁺, H⁺ and HCO_3_^-^ transporters (Mackinder et al., 2011), and/or molecules inhibiting mineralization (Marin et al., 2000, 1996), play pivotal roles in the control of calcium carbonate (CaCO_3_) precipitation by eukaryotic organisms. However, there are much fewer documented cases of genetically controlled biomineralization by prokaryotes (Cosmidis & Benzerara, 2022; Görgen et al., 2020). For a long time, the only well documented case was the intracellular formation of magnetites (Fe_3_O_4_) and/or greigites (Fe_3_S_4_) by magnetotactic bacteria (Monteil et al., 2018). This process is regulated by a suite of more than 40 genes involved in the control of the size, shape and crystallochemistry of the mineral byproducts (Liu et al., 2023; Taoka et al., 2023; Uebe & Schüler, 2016). More recently, the formation of intracellular amorphous calcium carbonates (iACC) by several species of cyanobacteria has been evidenced as an additional case of genetically controlled biomineralization in bacteria (Benzerara et al., 2014; Couradeau et al., 2012). Before this discovery, cyanobacteria were usually thought to induce the precipitation of CaCO_3_ as an indirect by-product of photosynthesis, with no apparent involvement of specific genes (Altermann et al., 2006). Cyanobacteria biomineralizing iACC are cosmopolitan (Ragon et al., 2014) and can sequester alkaline earth elements such as Ca, Sr, Ba and Ra, including their radioactive isotopes (Blondeau et al., 2018a; Cam et al., 2016; Mehta et al., 2019). As a result, these bacteria may be interesting candidates for bioremediating some radioactive pollutions (Mehta et al., 2022a). Morever, the biomineralization of iACC has been evidenced in phylogenetically diverse strains within the phylum of Cyanobacteria as well as in other bacterial phyla (Liu et al., 2021; Monteil et al., 2021). It is also found in several strains (but not all) of the well-studied *Microcystis* genus (Gaëtan et al., 2023). This cyanobacterium is particularly interesting since it can form blooms, with some genotypes being toxic for human and wildlife (Harke et al., 2016), causing numerous environmental issues in eutrophic freshwater ecosystems (Dick et al., 2021). Whether the formation of iACC by some *Microcystis* strains may impact their ecological success under certain conditions remains to be deciphered. Some putative biological functions have been suggested for iACC such as intracellular storage of inorganic carbon, intracellular pH buffering or their use as a ballast modifying cell buoyancy (Cosmidis & Benzerara, 2022). About the later hypothesis, we note that buoyancy is not just controlled by cell density but also by additional parameters such as EPS adhering to the cells, enhancing cell aggregation. For example, Gu et al. (2020) showed that Ca induces EPS production by *M. aeruginosa*, which can increase buoyancy through the increased size of cell aggregates. The relative contribution of these opposing parameters on the buoyancy should be assessed in the future. Moreover, we can speculate that the fulfilment of these functions may involve some kind of homeostasis which could require both iACC precipitation and dissolution processes, although iACC dissolution has not been evidenced yet in *Cyanobacteriota*. Last, the formation of iACC, a concentrated intracellular sink of Ca, by the large biomasses of *Microcystis* during blooms may impact the local geochemical cycle of Ca in these environments, although future studies will have to further document this possibility (Gaëtan et al., 2023). Indeed, the fraction of a natural *Microcystis* bloom with Ca-concentrating capabilities remains to be determined. Moreover, the dependence of Ca-sequestration by cells on extracellular Ca concentrations needs to be specified. While Ca concentrations may be lower in many lakes than in the BG11 medium (less than 50% of the lakes with a pH higher than 7.4 in the database investigated by Weyhenmeyer et al. 2009), we note, however, that concentrations in lakes vs laboratory cultures cannot be simply compared. Indeed, cultures in the laboratory of iACC-forming cyanobacteria have been conducted so far under batch conditions with a Ca supply limited by the volume of the culture, a condition that may be very different from those encountered in an open system such as a lake.

Despite the potential environmental importance of this process, the molecular mechanisms driving the biomineralization of iACC in cyanobacteria remain poorly known. It has been shown that it costs some energy to the cells (Cam et al., 2018; De Wever et al., 2019). Moreover, a genetic control has been proposed (Benzerara et al., 2022). Indeed, based on a comparative genomics approach, one family of genes, named *ccyA*, was found in phylogenetically diverse iACC-forming cyanobacteria and was absent from cyanobacteria not biomineralizing iACC. The *ccyA* gene was found in about one third of all publicly released genomes of *Microcystis i.e.*, 93 out of 282 genomes assemblies available at the time of publication (Gaëtan et al., 2023). The protein encoded by *ccyA*, called calcyanin, lacks detectable full-length homologs with a known function in public databases. Therefore, its function remains unknown. Several clues suggest that calcyanin may play a role in calcium homeostasis (Benzerara et al., 2023, 2022). However, since the expression of the *ccyA* gene in mutants of cyanobacteria not forming iACC did not result in the formation of iACC, this biomineralization process likely involves additional genes (Benzerara et al., 2022). For example, some of these genes may play a role in the fact that iACC inclusions form within an intracellular compartment, the envelope of which remains of unknown chemical composition (Blondeau et al., 2018b). Such a compartmentalization may serve to achieve chemical conditions allowing the precipitation and the stabilization of ACC. Additional examples of molecular processes that may favor iACC precipitation are carbon-concentrating mechanisms (CCMs) (Li et al., 2016; Görgen et al., 2021), which have been abundantly described in cyanobacteria and involve diverse genes and multiple mechanisms (Badger & Price, 2003; Kupriyanova et al., 2023). Last, calcium transport genes may also be involved in the accumulation of high amounts of calcium within iACC (De Wever et al., 2019). Consistently, the presence/absence of the *ccyA* gene in genomes was correlated with the presence/absence of genes involved in the transport and homeostasis of calcium and inorganic carbon (Benzerara et al., 2022). Moreover, in some genomes, *ccyA* was even colocalized with such genes.

Overall, a first step to gaining a better understanding about the molecular mechanisms involved in iACC biomineralization is to look at the expression of the *ccyA* gene together with other genes, including those neighboring *ccyA* in the genome and those mentioned above. The diel (day/night) cycle is an important parameter driving large variations of gene expression in cyanobacteria (Stöckel et al., 2008; Welkie et al., 2019; Zinser et al., 2009). For example, the diel cycle influences the expression of numerous genes involved in key cellular processes in *Microcystis aeruginosa* such as photosynthesis, carbon fixation, nitrogen metabolism, and circadian rhythms as shown, for example, by Huang et al. (2004) in *M. aeruginosa* PCC 7820. Similarly, the genes involved in a circadian cycle were studied in *M. aeruginosa* PCC 7806 (Straub et al., 2011) under a 24- hour light/dark cycle. They found that while the *kaiA* gene showed no significant variations during the cycle, the transcription patterns of the *kaiB* and *kaiC* genes, as well as the *sasA* gene encoding the two-component sensor histidine kinase, a KaiC-interacting protein, exhibited significant changes. These findings suggested that light is not the sole factor triggering the transcription of genes involved in photosynthesis and respiration; instead, their transcription may also be regulated by an endogenous circadian clock. Therefore, we have investigated the expression of the whole genome of *Microcystis aeruginosa* PCC 7806, which contains the *ccyA* gene and forms iACC under laboratory culture conditions, over a 24 h day/night cycle. In particular, we assessed whether the expression of genes either neighboring *ccyA* in the genome and/or involved in CCM and calcium transport might be correlated with that of *ccyA*.

## Methods

### Cultures and sampling

The study was conducted using the cyanobacterial strain *Microcystis aeruginosa* PCC 7806 available at the Pasteur culture of cyanobacteria (PCC) collection at the Institut Pasteur. The culture, sampling, nucleic acid extraction, and RNA sequencing were carried out at the the Institut Pasteur in 2017. The photosynthetic photon flux density provided during the day (50 μmol photons.m^-2^.s^-1^; cool white OSRAM L 18/640) was measured using a LICOR LI-185B quantum/radiometer/photometer equipped with a LICOR LI-193SB spherical sensor. A preculture of *M. aeruginosa* strain PCC 7806 was grown at 22 °C in 40 mL of the BG11 growth medium (Rippka et al., 1979) amended with 10 mM NaHCO_3_ under a 13/11 h day/night cycle. A 13-hour light period corresponds to what is observed in September at midlatitudes in the Northern hemisphere, a month during which *Microcystis* proliferates in numerous ecosystems.Then, the culture was transferred into three separate flasks (named A, B and C). The triplicate cultures of 40 mL were progressively transferred to a final set of three cultures with a volume of 1250 mL each, in fresh BG11 medium supplemented with 20 mM NaHCO_3_ and synchronized in an INFORS incubator with 1% CO_2_ and agitation at 50-100 rpm under the same day/night cycle and temperature conditions. Synchronization involved three consecutive transfers, aiming for an initial optical density (OD) at 750 nm of 0.05 and growing for a month. Sampling was performed during the mid-exponential phase (OD of 0.6-0.7) for each biological replicate, over a period of 24 h at eight time points: (i) 7 am, 60 min before the transition from night to day (t1_N); (ii) 9 am, 60 min after the transition from night to day (t2_D); (iii) 12 am (t3-D); (iv) 4 pm (t4_D); (v) 8 pm, 1 h before the transition from day to night (t5_D); (vi) 10 pm, 60 min after the transition from day to night (t6_N); (vii) 2 am, *i.e.* the middle of the night (t7_N), and (viii) in the end of the night period (t8_N), 24 h after the first sampling point. The sampling procedure involved the removal of 150 mL of culture, which were centrifuged, rinsed with sterile water, and centrifuged again. The resulting pellets were flash-frozen in liquid nitrogen and stored at -80 °C until further processing. To minimize batch effects, the samples were randomized before subsequent processing. We note that relatively high phosphate concentrations such as those found in BG11 (∼180 μmol.L^-1^) can inhibit the precipitation of extracellular carbonates and/or induce that of Ca-phosphates as shown by, *e.g.*, Rivadeneyra et al. (2006; 2010). However, by contrast, it has been shown that this does not prevent at all the precipitation of intracellular carbonates, which has been experimentally studied in BG11 before (Cam et al., 2017; De Wever et al., 2019).

### Nucleic acid extraction

RNA extraction was carried out using a Trizol reagent (Life Technologies) and the NucleoSpin miRNA kit (Macherey-Nagel). Firstly, 1 mL of Trizol preheated at 65 °C was added to the frozen pellets and incubated at 65 °C for 12 min with vortexing three times. After centrifugation (10 min at 4 °C and 12,000 rcf), the upper phase was transferred to a new tube and mixed with 500 µL of Trizol preheated at 65 °C. RNA extraction was achieved by adding 400 µL of chloroform, and incubating samples for 3 min at room temperature before centrifugation (10 min at 4°C and 12,000 rcf). The upper phase was again transferred to a new tube for extraction with 400 µL of Trizol and 400 µL of chloroform with similar incubation and centrifugation conditions. Then, the upper phase obtained from this step was further processed using the NucleoSpin miRNA kit according to the manufacturer’s instructions. The final purified RNA was eluted in 20 µL of RNase/DNase-free sterile water. The absence of genomic DNA contamination was confirmed by PCR targeting the 16S rRNA gene using specific primers. The quality and quantity of RNA were assessed using a Nanodrop spectrophotometer (ThermoFisher Scientific) and a Bioanalyzer (Agilent) with RNA Nano chips.

### Library Preparation and RNA Sequencing

Library preparation was conducted using the TruSeq Stranded mRNA sample preparation kit (Illumina, San Diego, California) following the manufacturer’s instructions. Ribosomal RNA (rRNA) depletion was performed using the Ribo-Zero rRNA removal kit (bacteria, #MRZB12424, Illumina) with 5 µg of total RNA. The rRNA-depleted RNA was then fragmented using divalent ions at 94°C for 8 min. The resulting fragmented RNA samples were reverse-transcribed using random primers, followed by complementary-strand synthesis to generate double-stranded cDNA fragments. There was no need for an end repair step in this process. A 3’-end adenine was added, and specific Illumina adapters were ligated to the cDNA fragments. The ligated products were amplified by PCR. The quality of the resulting oriented libraries was assessed using Bioanalyzer DNA1000 Chips (Agilent, # 5067-1504) and quantified using spectrofluorimetry (Quant-iT™ High-Sensitivity DNA Assay Kit, #Q33120, Invitrogen). Sequencing was performed on the Illumina HiSeq2500 platform of the Institut Pasteur, generating single-end 65 bp reads with strand specificity. Each lane of sequencing accommodated a mixture of 18 multiplexed samples.

### Read processing

Raw RNA-seq reads (*available online, see section Data, scripts, code, and supplementary information availability*) were cleaned from adapter sequences and low-quality sequences using an in-house program (https://github.com/baj12/clean_ngs). Only sequences of at least 25 nt in length were considered for further analysis. Transcript abundance was quantified from the cleaned read dataset using the Salmon software (Patro et al., 2017). Pseudo-mapping of reads was done considering the coding sequences (CDS, including pseudogenes) of the complete RefSeq genome assembly of *M. aeruginosa* PCC 7806 as a reference (GCF_002095975.1). For each replicate, the estimated transcript abundances were normalized by Salmon according to the transcript size, genome size (number of CDS) and sample size (number of reads).

### Differential expression analysis

Total abundance data were further processed using the DiCoExpress R-script-based tool (Lambert et al., 2020) in order to detect genes expressed differentially between different time points. First, normalization was achieved using Trimmed Mean of the M-value (Robinson and Oshlack, 2010). During this procedure, 220 low-count transcripts were discarded, resulting in a total of 4,440 genes with normalized counts. Principal component analysis (PCA) was performed on this normalized dataset using the R packages FactoMineR and Factoshiny (Lê et al., 2008). Then, the differential expression between time points was tested using a generalized linear model. For comparison between each time point, statistics of differential gene expression are provided as Fold Change (FC) and p-values.

### Gene functional annotation

In addition to the annotations provided by the NCBI prokaryotic genome annotation pipeline for the genome assembly of *M. aeruginosa* PCC 7806 (https://www.ncbi.nlm.nih.gov/datasets/genome/GCF_002095975.1), protein sequences translated from CDS were processed using DeepNOG protein orthologous groups assignment (Feldbauer et al., 2021) as implemented in the Genovi tool (Cumsille et al., 2023). An orthologous group with a COG accession number is assigned to each gene, as well as a confidence score ranging within [0, 1]. The stringent threshold was set at 0.8. Orthologs of genes involved in carbon-concentrating mechanisms (CCM) and calcium transport were specifically detected as bidirectional best BLASTp hits using protein sequence queries described in (Tang et al., 2022) (15 genes from *Synechocystis* sp. PCC 6803, and the *ecaA* gene from *Anabaena* PCC 7120) and (De Wever et al., 2019) (13 genes from *Chroococcidiopsis thermalis* PCC 7203), respectively. The following thresholds were used: E-value cut-off of 1E-6, ≥30% identity and 70% coverage.

### Sequence and structure analysis of unknown proteins

General information on the three domains of unknown function (DUF) encoded by genes located in the genomic neighborhood of the *ccyA* gene was extracted from their respective InterPro entries (Paysan-Lafosse et al., 2023; DUF5132: IPR033456; DUF454: IPR00740; DUF1269: IPR009200). AlphaFold2 (AF2) 3D structure models of the three DUF-containing proteins and two hypothetical proteins neighboring *ccyA* were build using ColabFold (Mirdita et al., 2022). Foldseek (van Kempen et al., 2023) was used to search for structural similarities against the Protein Data Bank (PDB) and AlphaFold Database (AFDB). 3D structures were manipulated using Chimera (Pettersen et al., 2004). Hydrophobic cluster analysis (HCA) (Callebaut et al., 1997) was used to analyze protein secondary structure features.

## Results

### Overall expression pattern

A principal component analysis (PCA) was performed on the transcript abundances of the 4,440 genes, for each replicate at each sampling time (Figure 1). All the data points were projected onto a two-dimensional plane defined by the first two principal components, which accounted for 64.69% of the total variance (Figure 1). Among the triplicates (i.e. three independent cultures for each time-point) only minor variations were observed, at least along axis 1, and their values were clearly separated from the samplings at other time points. Furthermore, the resulting plot exhibited a clear separation along axis 1 between night (N) and day (D) points.

**Figure 1.**
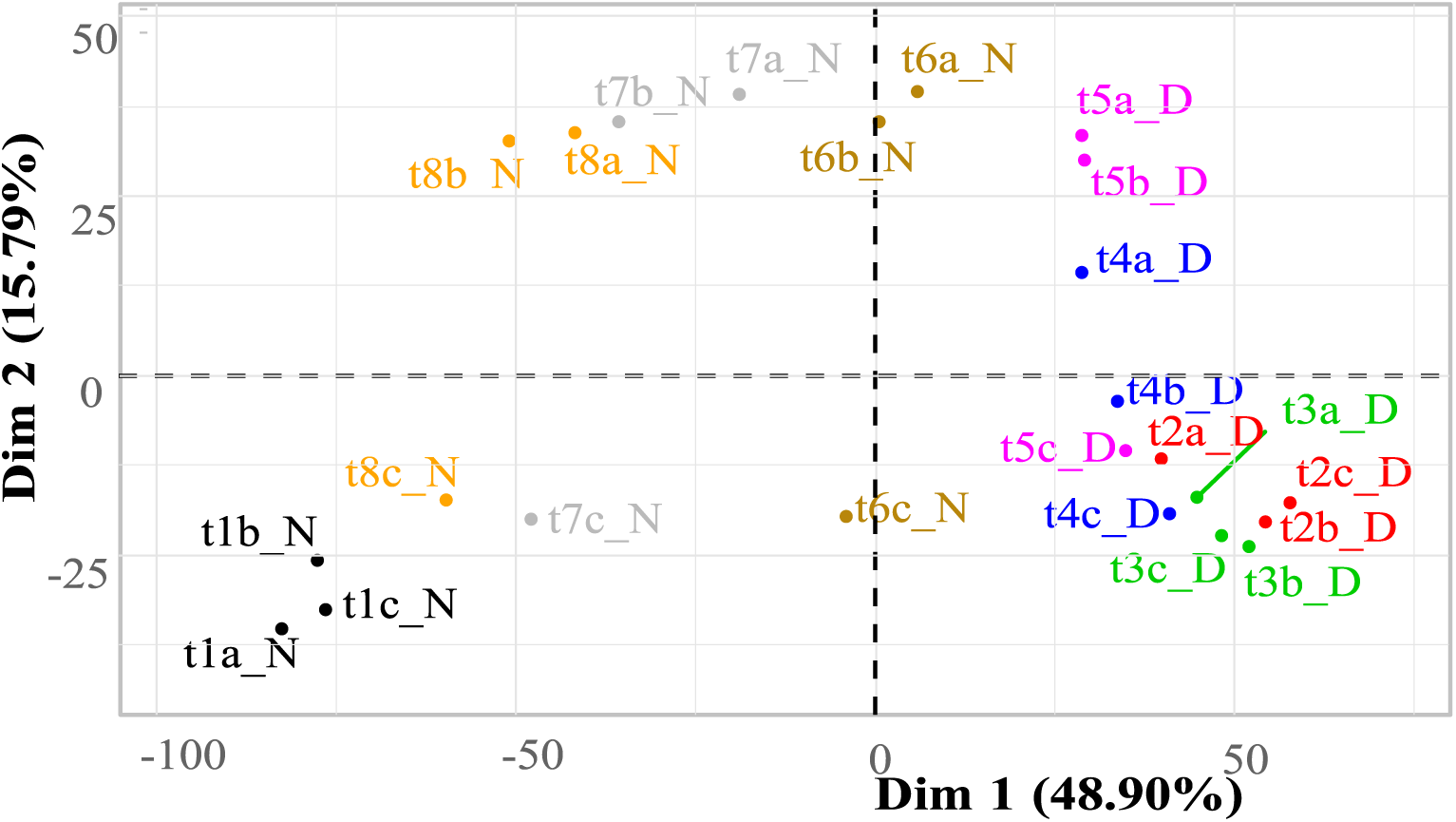
Principal component analysis of the transcript abundances of 4440 *M. aeruginosa* genes during a day/night cycle. Abundances are expressed as log base 2 of the normalized counts (see Methods section for normalization procedure), for each replicate (a-c) at each sampling time (t1-t8). The 24 samples are projected onto the first two principal components (Dim. 1 and Dim. 2), explaining 48.9% and 15.8% of the variance of the dataset, respectively. Samples/replicates collected at different times over the day/night cycle appear with a different color code. The “_N” and “_D” codes in the sample names refer to night and day, respectively.

### Expression of the *ccyA* gene

In order to analyze the expression profile of the *ccyA* gene during the 24 h day/night cycle, we plotted the normalized expression of *ccyA* in all replicates at the different time points (Figure 2). A clear day/night pattern can be observed in the expression of *ccyA*, with maximum transcript abundances during the second half of the night (t7_N and t8_N, Table 1) and a minimum transcript abundance at 12 am during daytime (t3_D, Table 1). Pairwise comparisons of *ccyA* transcript abundance at different time points show that the expression of *ccyA* at daytime points (t2_D to t5_D) did not significantly differ between each other, but were significantly different (p-value < 0.01) from some of the nightime points, t7_N and t8_N, respectively (Table S1). The 10 pm nighttime point (t6_N) did not significantly differ from any others except from the 12 am daytime point (t3_D; p-value < 0.05) and can therefore be considered as a transition point between day and night. When compared with the mean expression of the 4,440 genes of the whole transcriptome, the mean expression of the *ccyA* gene was significantly higher (p-value < 0.05) during the second half of the night, *i.e.* at the two-time points 2 am (t7_N) and 7 am (t8_N) (Table 1).

**Figure 2.**
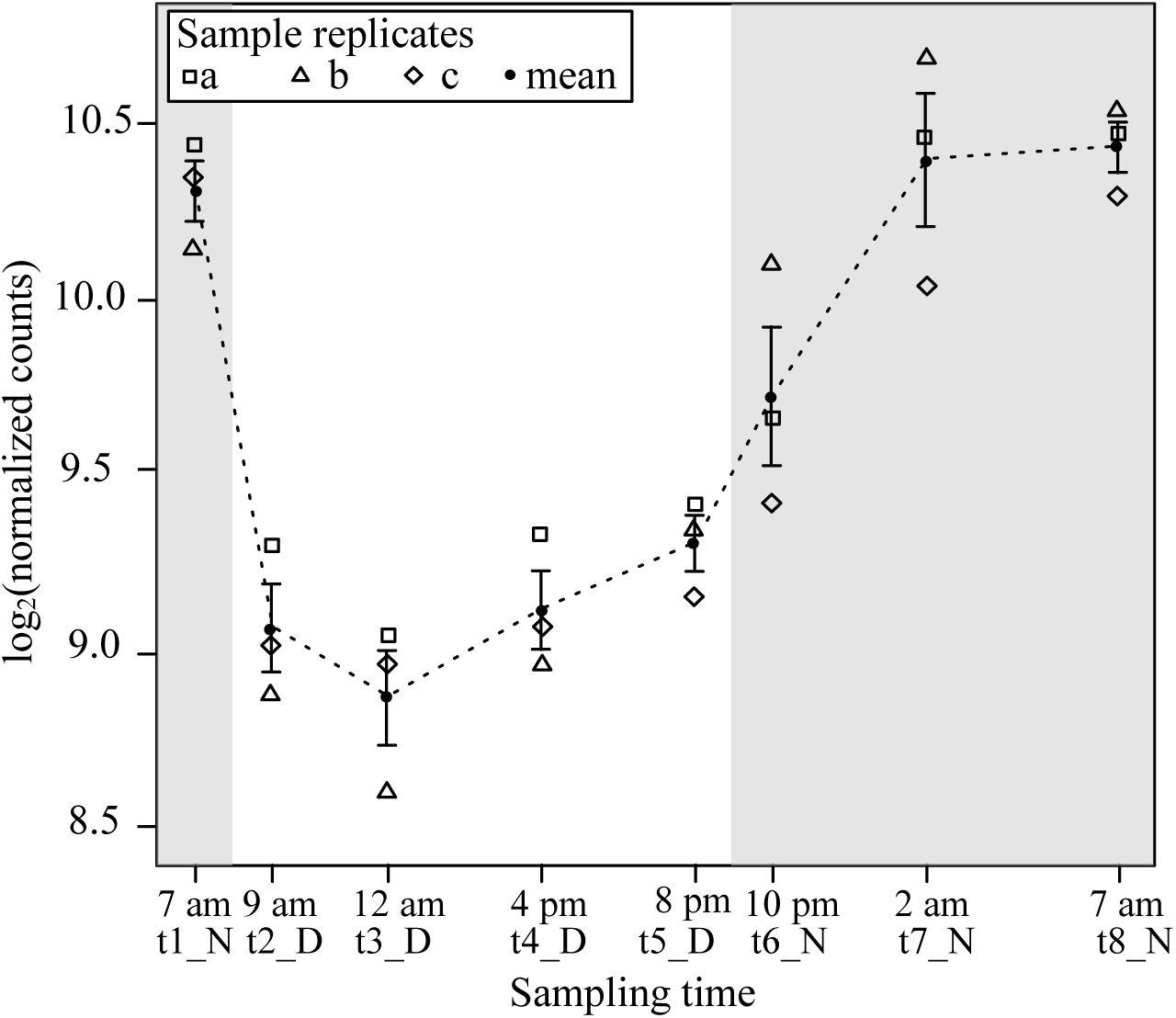
Time course of the abundance of *ccyA* transcripts during a day/night cycle. Abundances are expressed as log base 2 of the normalized counts. The abundance mean for the three replicates is shown by a black filled circle. Replicates a, b and c are represented by different symbols. For each sampling time (t1-t8), error bar represents the standard error on the mean of the three replicates. Grey shaded areas outline the night periods.

**Table 1.** Comparison between the transcript abundances of *ccyA* and the whole transcriptome, for each time point. Abundances are expressed as log base 2 of the normalized counts. When means significantly differ, p-values of Wilcoxon signed rank test with continuity correction are depicted as follows: (**): 0.01 – 0.05, (*): 0.05 – 0.1. Night points are shaded in grey.

|  | t1_N(*)<br>7 am | t2_D<br>9 am | t3_D<br>12 am | t4_D<br>4 pm | t5_D(*)<br>8 pm | t6_N(*)<br>10 pm | t7_N(**)<br>2 am | t8_N(**)<br>7 am |
| --- | --- | --- | --- | --- | --- | --- | --- | --- |
| **Mean**<br>(*ccyA* gene, n=3 replicates) | 10.31 | 9.07 | 8.88 | 9.12 | 9.31 | 9.73 | 10.39 | 10.43 |
| **Standard deviation**<br>(*ccyA* gene, n=3 replicates) | 0.15 | 0.22 | 0.23 | 0.19 | 0.14 | 0.34 | 0.33 | 0.12 |
| **Mean**<br>(4440 genes, n=3 replicates) | 6.87 | 7.35 | 7.39 | 7.44 | 7.44 | 7.36 | 7.23 | 7.18 |
| **Standard deviation**<br>(4440 genes, n=3 replicates) | 3.79 | 2.45 | 2.57 | 2.67 | 2.61 | 2.87 | 3.21 | 3.33 |

### Search of differential gene expression during the day/night cycle

Subsequently, a comprehensive analysis was conducted throughout the genome to identify all genes exhibiting diel variations in their expression, using the minima and maxima of *ccyA* gene expression (t3_D and t8_N) as reference time points (Figure 3). Overall, 1,799 of the 4,440 total genes showed a significant difference in expression (at least 2-fold) between 12 am (during the day, t3_D) and 7 am (during the night, t8_N) (Figure 3 and Table S2). Among these 1,799 genes, 906 were significantly more expressed during the day (t3_D), and conversely, 893 were over-expressed during the night (t8_N).

**Figure 3.**
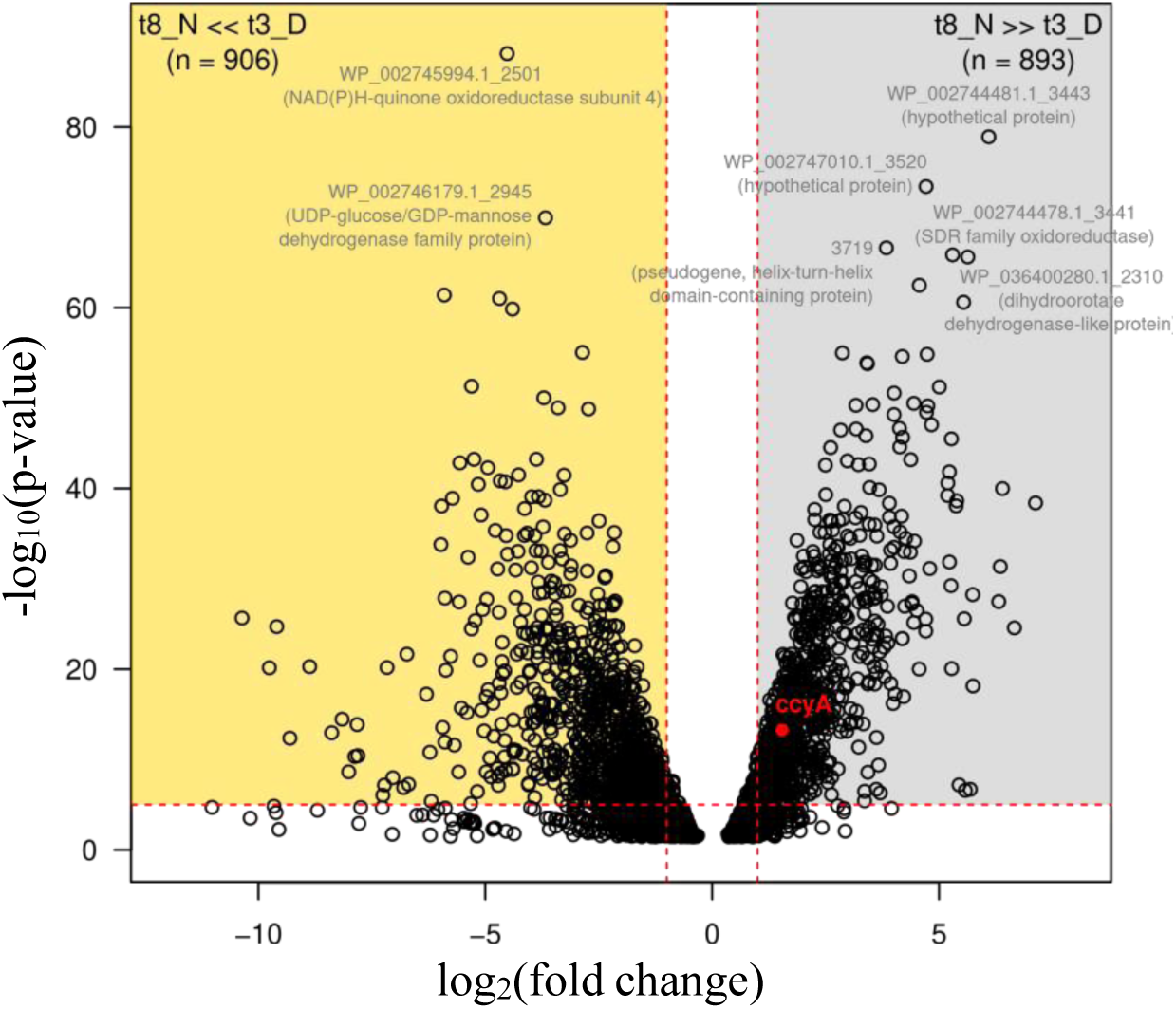
Differential expression of the whole transcriptome between sampling time t3_D (12 am, during the day period) and t8_N (7 am, end of the night period). Log base 2 of the fold change (FC) values (x-axis) are defined for 3116 CDS. CDS with a negative log2(FC) value correspond to transcripts significantly more abundant at 12 am (t3_D) compared with 7 am (t8_N) (gold area; 906 CDS). CDS with a positive log2(FC) value correspond to transcripts significantly more abundant at 7 am (t8_N) compared with 12 am (t3_D) (grey area; 893 CDS). The *ccyA* gene is indicated by a red dot. The horizontal red dashed line indicates the significance p-value threshold after Bonferroni correction (-log10(P-value)=5). The vertical red dashed lines indicate FC=2 and FC=0.5, *e.g.*, 2-fold abundance ratio between the two sampling times. RefSeq accessions and annotations are indicated for CDS with P<2.0e-66). See Suppl. Table S2 for details.

Figure 3 provides the annotations for the genes with the most substantial expression differences between these two time points (-log10(p-value)>60): genes overexpressed during the day included a NAD(P)H-quinone oxidoreductase and a protein of the UDP-glucose/GDP-mannose dehydrogenase family. The genes overexpressed at night included: two genes encoding hypothetical proteins, a pseudo-gene encoding a protein containing a helix-turn-helix domain, a gene encoding a protein from the SDR oxidoreductase family, and a gene encoding a dihydroorotate dehydrogenase-like protein.

### Genomic neighborhood of *ccyA*

Figure 4 displays the map of the 20 genes (18 genes and 2 pseudo-genes) located upstream (10 genes) and downstream (10 genes) to *ccyA*. This number of 20 genes was set arbitrarily and considered as a close neighborhood. When available, the functional annotations were extracted from the NCBI RefSeq database (Table 2). In this database, the *ccyA* gene is incorrectly annotated as a gene coding for the cell envelope biogenesis protein OmpA (Benzerara et al., 2022). Interestingly, three transposition elements were found near the *ccyA* gene: an ISL3 family and an IS1 family transposases, both pseudo-genes located at -6 and -5 genes compared to *ccyA*, and an IS1634 family transposase located at +2 genes compared with *ccyA*. The gene located directly upstream of *ccyA* was annotated as encoding a domain-of-unknown-function (DUF5132)-containing protein, and the one directly downstream a calcium/proton exchanger (*cax* gene), previously characterized in eukaryotes (Hirschi et al., 1996; Waight et al., 2013). At position -3 (upstream of *ccyA*), another DUF-containing protein (DUF454) was detected.

**Figure 4.**
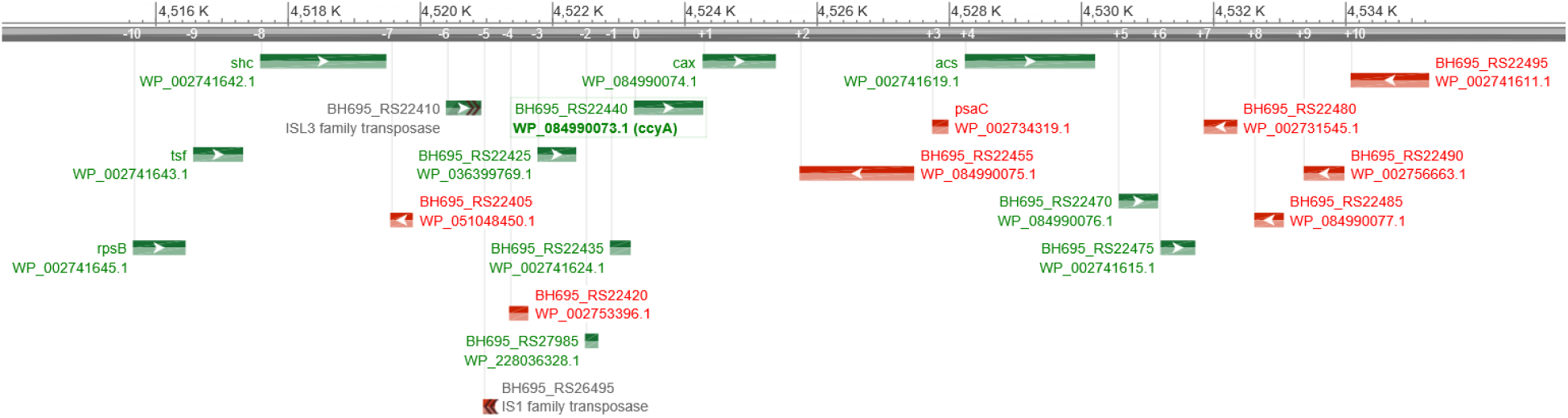
Schematic representation of the *ccyA* gene neighborhood in *M. aeruginosa* PCC 7806 complete genome assembly (NCBI genome browser display). The *ccyA* gene is outlined by a green box. RefSeq protein accession, gene name (if defined), gene orientation (red for minus; green for plus) and locus tag are indicated for the 10 CDS located upstream (-10 to -1) and the 10 CDS located downstream (+1 to +10) with respect to *ccyA*. Protein functional annotation is provided in Table 2. This genomic region spans within the nucleotide position interval [4515650, 4535259] in the NCBI genome assembly NZ_CP020771.1. CDS boundaries and features correspond to RefSeq annotations (GCF_002095975.1). Gene symbols stand for *shc*: squalene--hopene cyclase; *tsf*: translation elongation factor Ts; *rspsB*: 30S ribosomal protein S2 ; *cax*: calcium/proton exchanger; *acs*: acetate--CoA ligase; *psaC*: photosystem I iron-sulfur center protein PsaC.

**Table 2.**
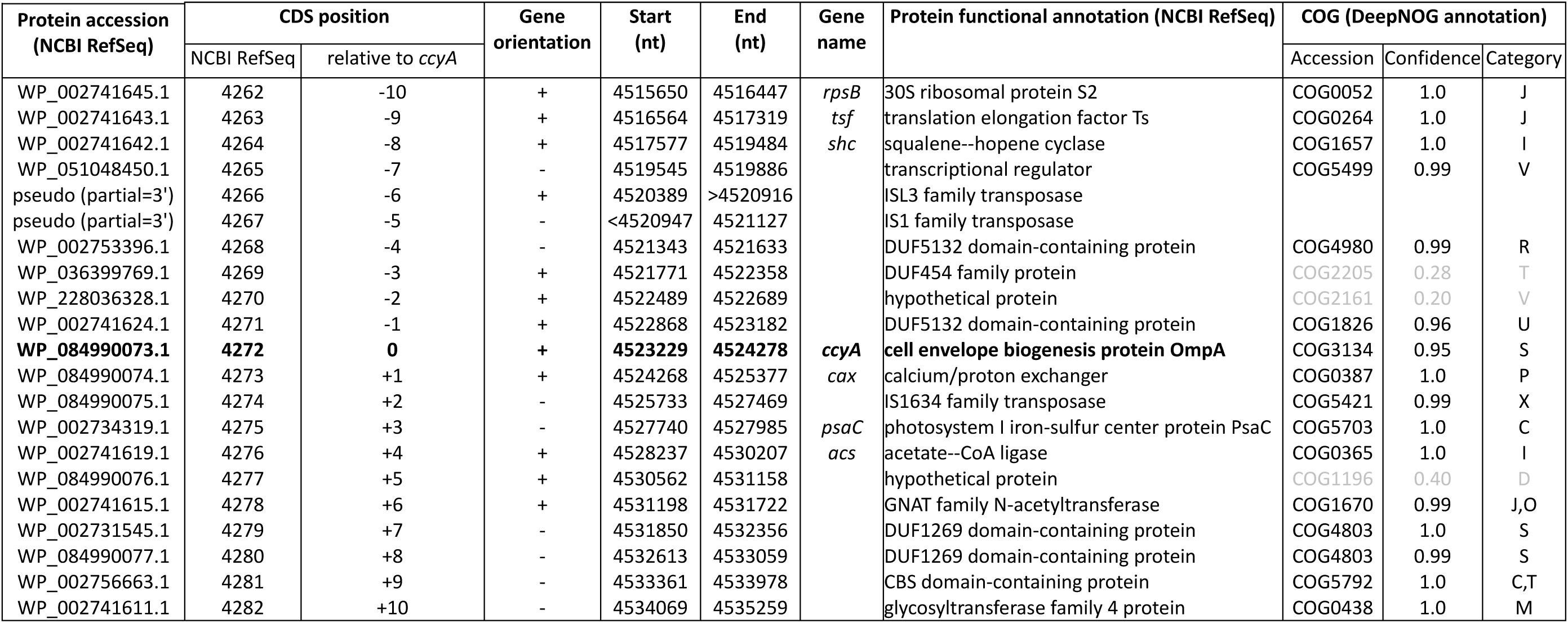
Neighborhood of the *ccyA* gene in *M. aeruginosa* PCC 7806 complete genome assembly. General features and annotations are listed for the ten CDS located upstream (-10 to -1) and the ten CDS located downstream (+1 to +10) of the position of *ccyA* (in bold, protein improperly annotated in RefSeq as OmpA). RefSeq annotations correspond to the assembly accession GCF_002095975.1 (NZ_CP020771.1).

AlphaFold 2 (AF2) 3D structure models of two hypothetical proteins, as well as five proteins containing DUFs present in the *ccyA* genomic neighborhood were analyzed (Figure 5) and compared with experimental 3D structures using Foldseek, in order to highlight possible structural and functional relationships. (i) Analysis of the DUF454 family protein (WP_036399769.1; position -3 relatively to *ccyA*) indicates that its C-terminal end matches the C-terminal end of the DUF454 IPR profile (inner membrane protein YbaN from *E.coli*). Meanwhile, its N-terminal part matches the Pfam HMA_2 profile (PF19991), corresponding to a CoBaHMA domain. CoBaHMA (after Conserved Basic residues HMA) is the name of a recently identified family of domains belonging to the HMA superfamily (Gaschignard et al., 2024). Within this superfamily, the CoBaHMA domain family shows distinct features such as the presence of an additional strand β0 at the N-terminus of the domain and the presence of a charged patch on one side of the β-sheet. The AF2 3D structure model also consistently highlights this two-domain architecture, with the C-terminal, DUF454- like domain folding as a helical hairpin, while the CoBaHMA domain possesses the basic conserved amino acids characteristic of this family (Figure 5-A). (ii) The DUF5132-containing protein (WP_002741624.1; position -1 relatively to *ccyA*) is covered by the DUF5132 profile (aa 48-89) over a little less than half of its length and contains a predicted transmembrane segment in its N-terminal part (aa 12-32). The AF2 3D structure model, characterized overall by good pLDDT values, indicates a putative assembly, in a hairpin-like conformation, of the helical transmembrane segment and another alpha-helix preceding a long, soluble helix matching the DUF5132 profile (Figure 5-B). (iii) A second member of this family, with similar structural features is also present in the *ccyA* neighborhood (WP_002753396.1; position -4 relatively to *ccyA*). No significant relationship with any known 3D structures could be detected by FoldSeek. (iv) A hypothetical protein (WP_228036328.1; position -2 relatively to *ccyA*) corresponds to a truncated part of a hypothetical protein referenced under RefSeq WP_271988195.1, which is also composed of a CoBaHMA domain sharing the basic signature of this family (Figure 5-C). (v) Another hypothetical protein (WP_084990076.1; position +5 relatively to *ccyA*) is predicted as a long coiled-coil (data not shown). (vi) and (vii) Finally, two DUF1269-containing proteins (WP_002731545.1 and WP_084990077.1; genes at +7 and +8 relatively to *ccyA*), corresponding to two isoforms differing by their C-terminal end, match the DUF1269 IPR profile over their whole lengths and include a long glycine-zipper motif embedded in a soluble domain, which based on Foldseek searches, corresponds to a ferredoxin fold, as found for instance in the copper tolerance CutA1 protein (pdb 1V6H, Bagautdinov, 2014) and in the nickel-responsive transcription factor NikR (pdb 2BJ3, Chivers and Tahirov 2003) (Figure 5-D). This glycine-zipper motif is similar, albeit more hydrophobic, to the GlyZip motif in the C-ter domain of calcyanins, composed of two parts separated by a proline preceded by a small amino acid (here a serine, while it is a glycine in calcyanins) (Figure 5-D). Overall, this analysis highlighted some features in the neighborhood of the *ccya* gene, which were not captured by automatic annotations, including motifs that are found in calcyanins of all cyanobacteria (GlyZip and hairpin-like helices) as well as calcyanins of some species (CoBaHMA domains).

**Figure 5.**
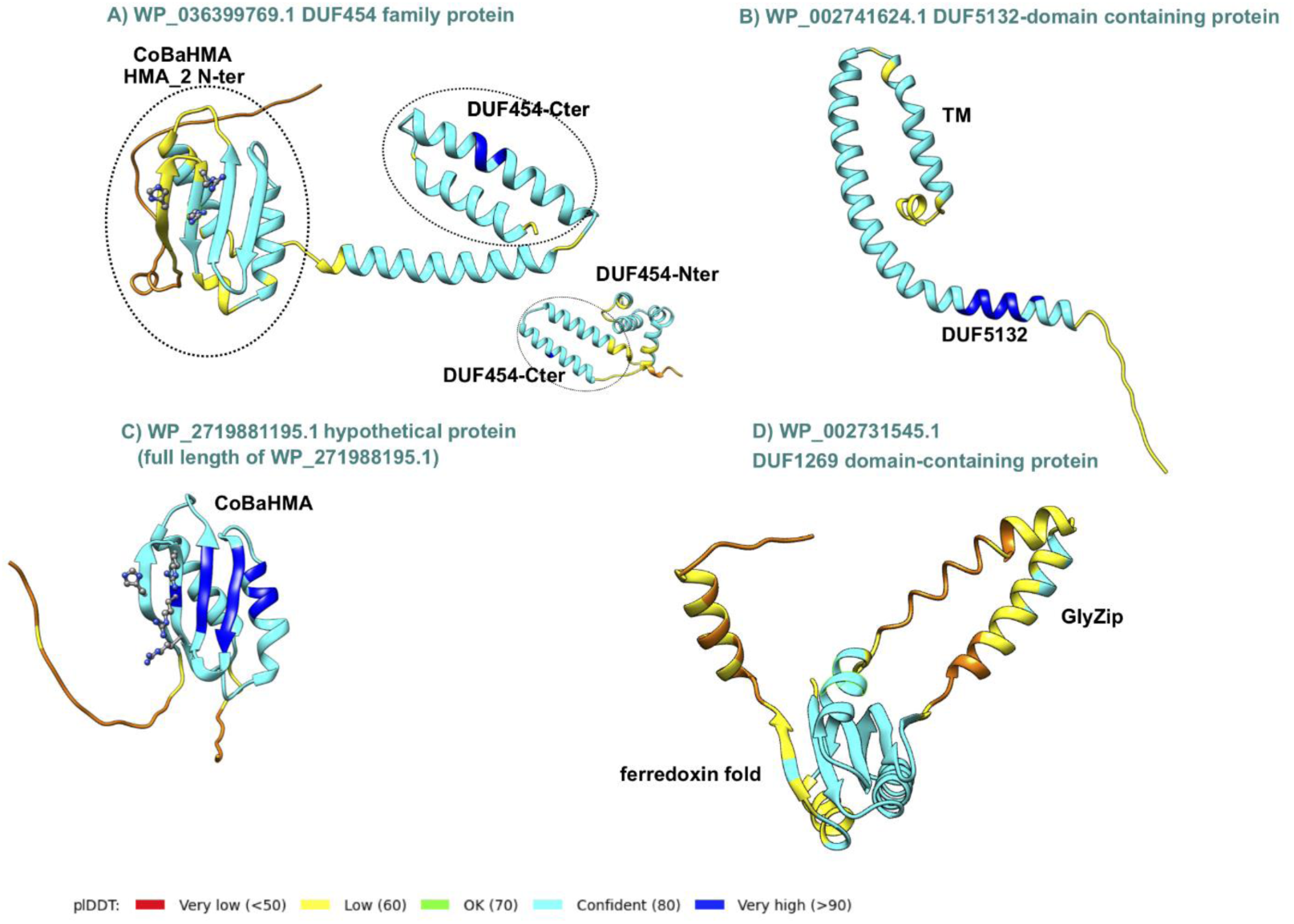
AlphaFold2 3D structure models of some proteins encoded by the genomic neighbors of *ccya.* The 3D structure models are represented as ribbons colored according to the plDDT values (color code given at bottom). In panels A and C, the basic amino acids constituting the signature of the CoBaHMA family are shown as balls and sticks. (A) The CoBaHMA_DUF454-Cter architecture (encoding gene in position -3 relatively to *ccyA*) is compared with a protein entirely covered by the DUF454 IPR profile (inner membrane protein YbaN (YABN_ECOLI, UniProt P0AAR5); insert). (B) DUF5132 protein (encoding gene in position -1 relatively to *ccyA*). TM stands for TransMembrane. (C) Hypothetical protein (encoding gene in position -2 relatively to *ccyA*) with a typical CoBaHMA domain. (D) DUF1269 domain-containing protein (encoding gene in position +7 relatively to *ccyA*) containing a GlyZip motif, predicted to fold as a helical hairpin (see HCA plot and sequence alignment in Figure S1).

#### Correlated expression of other genes with the *ccyA* gene

The expression patterns of genes exhibiting a significant difference in their circadian expression between daytime (t3_D, 12 am) and nighttime (t8_N, 7 am) were compared with that of *ccyA* to figure out which ones may have a maximum expression at the same time (Figure 6; Table S3). We note that such gene expression variations over a day/night cycle are expected in an autotrophic organisms and at least some correlations are likely not causal. However, their identification offers one possibility to better understand which other pathways may be activated at the same time as *ccyA* is expressed. As a result, two subsets were identified: on the one hand, 773 genes showed a highly positive correlation with the expression pattern of *ccyA* (with a Pearson correlation coefficient rho > 0.75, this threshold being arbitrarily fixed; grey subset in Figure 6; see also Figure S2 and Table S3); on the other hand, 706 displayed a negative correlation with the expression pattern of *ccyA* (rho < -0.75, yellow subset in Figure 6; see also Figure S2 and Table S3). Figure 6 displays the significance level (p-value) of the correlations between the expression profiles of these 1479 genes and that of *ccyA*, and provides annotation for the genes with the most significant correlation.

**Figure 6.**
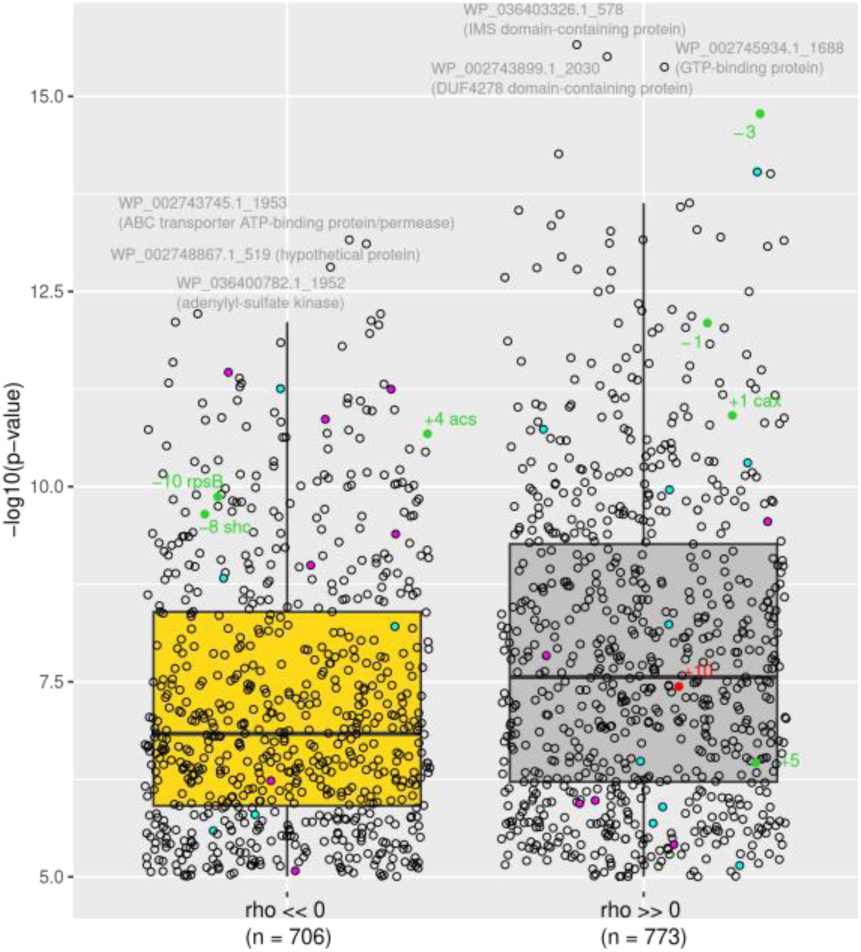
CDS with a day/night expression pattern strongly (anti-)correlated with that of *ccyA*. The Pearson correlation coefficient (rho) is highly negative for 706 CDS (left). The Pearson correlation coefficient is highly positive for 773 CDS (right). The distribution of correlation coefficients over the 4440 CDS is shown in Figure S2. RefSeq accessions and annotations are indicated for the 3 CDS with the strongest negative and positive correlations (see Table S3 for details). CDS belonging to the carbon concentrating mechanism or the calcium transport system are represented as magenta and cyan dots, respectively. CDS located in the [-10, +10] neighborhood of *ccyA* (Table 2 and Figure 4) are represented by green and red dots (for positive and negative DNA strands, respectively). The *cax* gene located at a genomic position of +1 from the *ccyA* gene also belongs to the calcium transport system and appears in green (see Table S4 for details).

The genes for which the expression patterns were the most significantly positively correlated with that of the *ccyA* gene were: (i) an intermembrane mitochondrial space (IMS) domain-containing protein, (ii) a GTP-binding protein, and (iii) a DUF-containing protein (DUF4278). The genes the most significantly anticorrelated with *ccyA* expression profile were annotated as (i) an ABC transporter ATP-binding protein/permease, (ii) an adenylyl-sulfate kinase, and (iii) a hypothetical protein.

Among the 20 neighboring genes of *ccyA*, eight showed an expression profile either positively or negatively correlated with that of *ccyA* (see Figure S3 for a time plot of expression of all 20 genes).

Interestingly, the expression profile of the two genes adjacent to *ccyA*, located on the same DNA strand, *i.e.* encoding for a calcium/proton exchanger (CAX) and DUF5132 (Table 2), displayed a strong positive correlation with the expression profile of *ccyA* (Figure 6 and 7). The expression profile of the gene coding for a DUF454 family protein (position -3 relatively to *ccyA*; Table 2), exhibited one of the strongest positive correlations with the expression profile of *ccyA* (Figure 6), with a nearly identical expression profile (Figure 7). Finally, the expression profiles of two other genes, located at +10 and +5 genes with respect to *ccyA* showed a weaker but significant positive correlation with the expression profile of *ccyA*. The gene at +10 was annotated as a glycosyltransferase family 4 protein. The gene at +5 codes for a hypothetical protein predicted as a long coiled-coil (Table 2).

**Figure 7.**
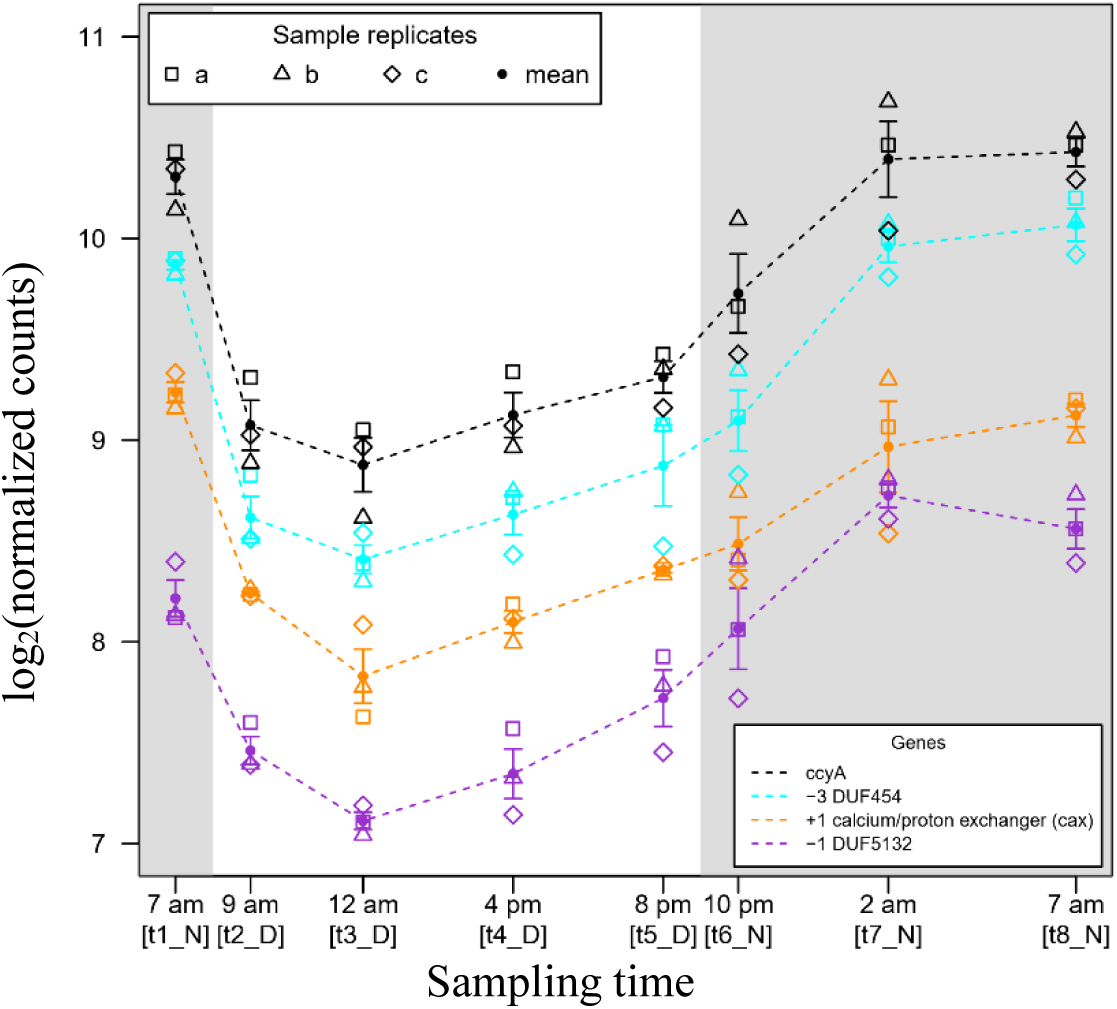
Time course of the abundances of three genomic neighbors of the *ccyA* gene during a day/night cycle. The CDS located at positions -3 (blue), -1 (purple) and +1 (orange) correspond to the *ccyA* neighbors with transcript abundance profiles the most correlated to *ccyA* transcript abundance profile: Pearson correlation coefficient of 0.97 (p-value=1.7e-15), 0.95 (p-value=8.0e-13), 0.94 (p-value=1.2e-11), respectively. See Methods section for more details about how counts were normalized.

Moreover, proteins involved in calcium transport and in CCM were searched in the genome of PCC 7806 (Table S4). In total, 24 genes involved in CCM and/or their homologs and 24 genes involved in calcium passive or active transport and their homologs were identified. The expression patterns of seven out of the 24 genes involved in CCM were correlated negatively with the expression pattern of *ccyA*, whereas the expression patterns of five of CCM genes were positively correlated with the expression pattern of *ccyA* (Figure 6, in magenta; see Figure S4 for a time plot of the expression of all CCM genes). Among the 24 genes involved in calcium transport, the expression patterns of five of them were negatively correlated with that of *ccyA*, while the expression patterns of ten of them were positively correlated with that of *ccyA* (Figure 6, in cyan except the *cax* gene in green; see Figure S5 for a plot of time expression of all “Ca genes”).

## Discussion

The identification of a marker gene of iACC biomineralisation in cyanobacteria was a first step towards understanding the molecular mechanisms controlling the formation of these biomineral phases (Benzerara et al., 2022). However, up to now, only the presence/absence of the gene and the structural features of the encoded protein have been investigated (Benzerara et al., 2022; Gaëtan et al., 2023; Gaschignard et al., 2024). Here, we explored whether this marker gene is transcribed in an organism capable of forming iACC, how this may be influenced by day/night cycles and how the expression of neighboring genes compares.

Several tools can be used for assessing the transcription of a large number of genes within a genome, including DNA microarray and high-throughput RNA sequencing (RNAseq) (Wang et al., 2009). Microarrays are limited by the use of probes: the number of studied genes remains limited by the available knowledge of the targeted genome. By contrast, RNAseq allows targeting all the mRNAs in a cell and has the advantage of measuring the level of transcriptional activity in the organisms (Mantione et al., 2014). Moreover, RNAseq can achieve higher resolution (*i.e.* higher accuracy in the estimation of gene expression) than microarrays. Because this technique does not suffer from microarray-based limitations such as background noise and saturation, it can also achieve a lower detection limit, and thus detect very low gene expression (Zhao et al., 2014). Straub et al. (2011) already studied the expression of the whole genome of *M. aeruginosa* PCC 7806 over a 24-h day/night cycle, using DNA microarray. Using their publicly available dataset, we determined that the expression level of the *ccyA* gene (MIC_2500) was not significantly different from the background noise. Here, the use of RNA-seq on the same organism enabled the detection and analysis of the *ccyA* gene expression dynamics. Indeed, the expression level of *ccyA* in our dataset was above the mean expression of the 4,440 genes of *Microcystis* at all time and this was statistically significant for several time points during the night. We can thus conclude that the *ccyA* gene is clearly expressed in an iACC-forming strain.

In line with the study by Straub et al. (2011) on the dynamics of gene expression in *M. aeruginosa* PCC 7806, we found, as expected, that the day/night cycle is an important factor driving variations in global gene expression in PCC 7806. Cyanobacteria have a highly defined circadian clock, and this day/night gene expression pattern has been demonstrated in several strains (Pattanayak & Rust, 2014; Stöckel et al., 2008; Zinser et al., 2009). Straub et al. (2011) showed that the metabolism of *M. aeruginosa* undergoes significant changes between the light and dark periods. Specifically, during the light period, processes including carbon uptake, photosynthesis, and the reductive pentose phosphate pathway result in glycogen synthesis. Conversely, during the dark period, glycogen degradation, the oxidative pentose phosphate pathway, the tricarboxylic acid (TCA) branched pathway, and ammonium uptake promote amino acid biosynthesis. Furthermore, the biosynthesis of secondary metabolites, such as microcystins, aeruginosin, and cyanopeptolin, predominantly occurs during the light period.

A clear day-night expression pattern was also observed for the *ccyA* gene: the expression was significantly higher than the average genome expression from the middle (2 am, t7_N) to the end of the night (7 am, t8_N). In all other time points of the 24-h cycle, the expression level of this gene did not statistically differ from the average expression of all genes. The gene is therefore overtranscribed during the second half of the night. While mRNA abundance provides valuable information about gene expression, it does not always directly correlate with the protein levels in the cell (Ingolia et al., 2009; Riba et al., 2019; Schwanhäusser et al., 2011). Several factors, including translation, initiation and elongation rate (Riba et al., 2019), post-translational modifications, protein stability, and degradation rates (Christiano et al., 2014; Gancedo et al., 1982) may impact the relationship between mRNA abundance and protein content. As such, it is difficult to infer the abundance of calcyanin at a given time from the transcript expression level only. However, one can speculate that the protein may be more produced during the second part of the night, and remain at an optimal abundance during the day when it may most function. In the future, direct measurements of the protein abundance should be performed using proteomics in order to quantify the calcyanin content in cells. Last, Straub et al. (2011) mentioned that the metabolism of *M. aeruginosa* strongly changes between the light and the dark period. For example, during the latter, amino acid biosynthesis is promoted by glycogen degradation, the oxidative pentose phosphate pathway, the TCA branched pathway and ammonium uptake. This illustrates that seeing *ccyA* expression at its strongest at the end of the dark period is not particularly surprising energetically-wise.

Interestingly, we showed that a Ca^2+^/H^+^ antiporter (*cax* gene) and two DUF-containing proteins were not only directly co-localized upstream and downstream of *ccyA*, on the same DNA brand but they were also co-expressed with *ccyA*. This suggests that those three genes could be part of a functioning unit of DNA that could be transcribed together into a single mRNA strand, *i.e.* an operon, and may play a concerted role in iACC formation.

The process of iACC formation necessitates the accumulation of calcium ions at concentrations higher than the cytosolic concentrations typically measured in cells (Cam et al., 2015). Therefore, it may be suggested that the protein encoded by the *cax* gene is involved in the build-up of these accumulations. However, the functioning of CAX proteins raises questions. Indeed, the single Ca^2+^/H^+^ exchanger found in *Synechocystis* PCC 6803 has been shown to be located in the plasma membrane and to catalyze Ca^2+^ efflux out of the cells (Waditee et al., 2004). By contrast, it has been suggested that CAX proteins in plants may have different locations and efflux Ca^2+^ from the cytoplasm either to the extracellular space or to intracellular vacuoles (Shigaki et al., 2006). Investigating further the similarities/dissimilarities of the CAX protein of *M. aeruginosa* PCC 7806 with already known CAX proteins and locating these transporters within *M. aeruginosa* PCC 7806 cells would therefore be crucial to better understand their potential involvement in iACC formation.

The gene encoding a DUF454 protein that co-localizes (position -3 relatively to *ccyA*) and co-expresses with *ccya* is particularly interesting. Indeed, we show here that this DUF454 protein bears a CoBaHMA (after Conserved Basic residues HMA) domain (Gaschignard et al., 2024). Strikingly, the calcyanins of many cyanobacteria are similarly composed of a CoBaHMA domain in their N-terminal region and a C-terminal three-fold repeat of a long glycine zipper motif conserved in all iACC-forming cyanobacteria (Benzerara et al 2022). This is not the case for the calcyanins of *Microcystis,* which does not contain a CoBaHMA domain but instead displays an unknown domain in their N-terminal region. Since the discovery of the calcyanin family, it has been surprising to observe such a broad diversity in their N-terminal domains despite an expected common function. Here, although we still ignore the function of calcyanin, the discovery of a CoBaHMA-encoding gene neighboring the *ccyA* gene suggests that *Microcystis* calcyanin may work in concert with other proteins, including a DUF454 protein and resulting in a protein complex with a possibly similar architecture in *Microcystis* and other iACC-forming cyanobacteria. Moreover, the presence in the close genomic neighborhood of *ccyA* of genes encoding for another protein of the CoBaHMA family (-2 position relative to *ccyA*), as well as proteins possessing a GlyZip-like motif (+7 and +8 positions relative to *ccyA*) relatively similar to the GlyZip motif in the C-ter domain of calcyanins additionally suggests the possibility of complementary roles of these proteins for a function to identify.

Moreover, building upon the approach followed by De Wever et al. (2019), we conducted a comprehensive search for annotated genes involved in CCM and calcium transport, as well as their homologous counterparts in the genome of *M. aeruginosa* PCC 7806. Firstly, a greater number of genes associated with CCM or their homologs appeared to be expressed preferentially during the day instead of night (nine *vs* four, respectively), compared with genes associated with calcium transport (three *vs* nine, respectively) (Table S4). Similarly, Straub et al. (2011) showed that the transcript abundance of CCM genes showed a peak in expression just after the night/day transition and then a rapid decrease after the day/night transition. The elevated concentration of CO_2_ in carboxysomes resulting from the CCM is a way to improve carbon fixation (Price and Howitt, 2011). The expression of CCM genes during the day is therefore expected. Yet, among the 24 genes identified as involved or potentially involved in CCM, 5 had an expression pattern positively correlated with that of *ccyA*, *i.e.* maximum at night (Table S4). Notably, two of these genes (*cmpC* and *cmpD*) encode two subunits of the inorganic carbon transporter BCT1 involved in the cyanobacterial CCM (Price & Howitt, 2011). BCT1 is a high-affinity but low-flux HCO_3_^-^ transporter. It is composed of four subunits, and among them, CmpC and CmpD are extrinsic cytoplasmic proteins that possess binding sites for ATP (Price & Howitt, 2011). It has been shown that the expression of these genes in *M. aeruginosa* PCC 7806 depends on the ambient pCO_2_ (Sandrini et al., 2015). Notably, under high pCO_2_ conditions (1450 ppm in their experiments), the expression of high-affinity bicarbonate uptake systems (notably *cmpA*, *cmpB*, and *cmpC* genes) is down-regulated, and cells shift towards CO_2_ and low-affinity bicarbonate uptake compared with cells grown under low pCO_2_ conditions (200 ppm) that the authors considered as C_i_-limited. For the production of the transcriptomes that we studied here, pCO_2_ was not regulated nor monitored, and the change of expression in CCM-involved genes such as *cmpD* and *cmpC* could be due to changes in pCO_2_ conditions during day/night transitions, even if there are expected to be lower than in the Sandrini et al. (2015)’s study. Moreover, we note that cmpA and cmpB were more expressed during the day. Therefore, the potential variation of the BCT1 activity over a diel cycle remains to be ascertained. In future studies, exploring whether the co-expression of the five genes involved in CCM with *ccyA* may play a role in iACC biomineralization would be interesting, and should include a special focus on the pCO_2_ variations during the experiments. Last, Walter et al. (2016) demonstrated that in *Anabaena* sp. PCC 7120, the concentration of extracellular Ca^2+^ can also impact the expression of genes, including those involved in bicarbonate uptake. While we do not expect a dramatic change of the extracellular Ca^2+^ concentration over a 24-hour period based on previous studies (Cam et al., 2018; De Wever et al., 2019), this parameter should also be considered in future studies. In contrast to the bicarbonate transporters and their homologs, the number of genes possibly linked with calcium transport (24 genes) and having an expression pattern positively correlated with that of *ccyA* (10 genes) was higher than genes with an expression pattern negatively correlated with *ccyA* (5 genes) (Table S4). The transport of Ca^2+^ ions into the cytosol is commonly considered as a passive process facilitated by channels that exhibit low ionic specificity (Domínguez et al., 2015). Here, only one out of the ten genes whose expression was positively correlated with that of *ccyA* was identified as a passive transporter (hB1-1). The other nine genes include the *cax* gene, neighboring *ccyA*, two *apnhaP* genes (Na^+^/H^+^ antiporter acting as a Ca^2+^/H^+^ antiporter at alkaline pH as shown by Waditee et al. (2001)) and three of their homologs, a UPF0016 gene (putative Ca^2+^/H^+^ transporter) and a Ca^2+^-ATPase and one of their homologs. They could thus also be candidate genes contributing to the intracellular accumulation of calcium with the same question arising about the polarity of the transport as for *cax* (towards the exterior of the cells or inside compartments containing iACC).

In a comparative genomics approach performed at the scale of the whole *Cyanobacteriota* phylum, *ccyA* was the only gene identified as shared by iACC+ strains and absent in iACC- strains (Benzerara et al., 2022). However, some genes from both phenotypes (iACC- and iACC+), and therefore not highlighted by the above-mentioned comparative genomics approach, could also be involved in iACC formation. Indeed, these genes may be recruited in iACC+ strains for iACC formation, whereas they would have a different function in iACC- strains. Moreover, there could also be genes specific to *Microcystis* involved in iACC formation, with other non-homologous genes but of similar functions involved in other cyanobacterial genera. The present study is a step forward towards identifying possible partners in iACC formation. However, only future studies specifically targeting these genes will be able to provide definitive answers about their implications in the iACC formation process. Furthermore, investigations into the comprehensive response of *Microcystis* to fluctuations in calcium and/or pCO_2_ levels could provide valuable insights into the molecular mechanisms underlying the biomineralization of iACC. These studies should encompass monitoring of the physicochemical parameters of the environment and the quantification of iACC formation. Notably, certain semi-quantitative techniques, such as Fourier transform infrared spectroscopy (FTIR) (Mehta et al., 2022b), scanning transmission x-ray microscopy (STXM) (Benzerara et al., 2023) and X-ray absorption near edge structure (XANES) spectroscopy (Mehta et al., 2023), now offer the capability to detect cyanobacterial iACC and assess their relative abundance. In the future, studying control strains not hosting the *ccyA* gene might be an additional interesting perspective. However, for this purpose, one cannot consider any *ccyA*- strain as a robust control since its genome may differ from the one of PCC 7806 in several genes. A more reliable control would consist in a yet-unavailable mutant of PCC 7806 with a desactivated/deleted *ccyA* gene. Last, one may speculate that cells may sometimes, also direct iACC dissolution to regulate their abundance. While this process has been shown in *Achromatium* (e.g., Yang et al., 2019), it still needs to be unambiguously evidenced in *Cyanobacteriota* before assessing which genes might be involved.

Finally, it can be noted that the correlation between the *ccyA* gene and genes involved in calcium transport and the CCM is not restricted to *Microcystis*. For example, the combined presence of homologs of a Ca^2+^/H^+^ antiporter gene and the Na^+^-dependent bicarbonate *bicA* gene was shown to be significantly correlated with that of *ccyA* in the *Cyanobacteriota* phylum (Benzerara et al., 2022). Moreover, the *ccyA* gene was co-localized with a Ca^2+^/H^+^ antiporter gene in several other cyanobacteria (*C. fritschii* PCC 9212 and PCC 6912, and *Fischerella* sp. NIES-4106), mirroring the situation observed in *M. aeruginosa* PCC 7806. Overall, this offers some target genes for future genetic studies to better depict the molecular mechanisms of iACC biomineralization.

## Supporting information

Readme_Supporting information

## Acknowledgements

We thank Varet H, Sismeiro O, Dillies MA, Legendre R and Coppe JY who run the RNA platform of the Institut Pasteur and provided us with the processed trancriptomics data. We also thank two reviewers for comments that significantly improved the manuscript.

## Data, scripts, code, and supplementary information availability

Raw transcriptomics data are available online on NCBI, Gene expression Omnibus, GEO accession number: GSE255450. Supplementary information, supplementary data (output data from Salmon, DICOEXPRESS and DeepNOG analyses as well as gene expression correlation) and supplementary figures and tables are available online on bioRxiv, BIORXIV/2024/602159 - Version 3, https://doi.org/10.1101/2024.07.07.602159. Links to the tools used in this study are noted in the Material and methods section.

## References

1. Raw transcriptomics data are available on NCBI, Gene expression Omnibus, GEO accession number: GSE255450. https://www.ncbi.nlm.nih.gov/geo/query/acc.cgi?acc=GSE255450

2. Supplementary information and supplementary data, figures and tables can be found on bioRxiv, at the following address: 10.1101/2024.07.07.602159

3. Altermann, W., Kazmierczak, J., Oren, A., Wright, D.T., 2006. Cyanobacterial calcification and its rock-building potential during 3.5 billion years of Earth history. Geobiology 4, 147–166. 10.1111/j.1472-4669.2006.00076.x

4. Badger, M.R., Price, G.D., 2003. CO_2_ concentrating mechanisms in cyanobacteria: molecular components, their diversity and evolution. Journal of Experimental Botany 54, 609–622. 10.1093/jxb/erg076

5. Benzerara, K., Duprat, E., Bitard-Feildel, T., Caumes, G., Cassier-Chauvat, C., Chauvat, F., Dezi, M., Diop, S.I., Gaschignard, G., Görgen, S., Gugger, M., López-García, P., Millet, M., Skouri-Panet, F., Moreira, D., Callebaut, I., 2022. A new gene family diagnostic for intracellular biomineralization of amorphous Ca carbonates by cyanobacteria. Genome Biology and Evolution 14, evac026. 10.1093/gbe/evac026

6. Benzerara, K., Görgen, S., Athar, K.M., Chauvat, F., March, K., Menguy, N., Mehta, N., Skouri-Panet, F., Swaraj, S., Travert, C., Cassier-Chauvat, C., Duprat, E., 2023. Quantitative mapping of calcium cell reservoirs in cyanobacteria at the submicrometer scale. Journal of Electron Spectroscopy and Related Phenomena 267, 147369. 10.1016/j.elspec.2023.147369

7. Benzerara, K., Skouri-Panet, F., Li, J., Férard, C., Gugger, M., Laurent, T., Couradeau, E., Ragon, M., Cosmidis, J., Menguy, N., Margaret-Oliver, I., Tavera, R., López-García, P., Moreira, D., 2014. Intracellular Ca-carbonate biomineralization is widespread in cyanobacteria. Proceedings of the National Academy of Sciences 111, 10933–10938. 10.1073/pnas.1403510111

8. Blondeau, M., Benzerara, K., Ferard, C., Guigner, J.-M., Poinsot, M., Coutaud, M., Tharaud, M., Cordier, L., Skouri-Panet, F., 2018a. Impact of the cyanobacterium *Gloeomargarita lithophora* on the geochemical cycles of Sr and Ba. Chemical Geology 483, 88–97. 10.1016/j.chemgeo.2018.02.029

9. Blondeau, M., Sachse, M., Boulogne, C., Gillet, C., Guigner, J.-M., Skouri-Panet, F., Poinsot, M., Ferard, C., Miot, J., Benzerara, K., 2018b. Amorphous calcium carbonate granules form within an intracellular compartment in calcifying cyanobacteria. Front. Microbiol. 9, 1768. 10.3389/fmicb.2018.01768

10. Boskey, A.L., 2003. Biomineralization: an overview. Connective Tissue Research 44, 5–9. 10.1080/03008200390152007

11. Brownlee, C., Wheeler, G.L., Taylor, A.R., 2015. Coccolithophore biomineralization: New questions, new answers. Seminars in Cell & Developmental Biology, 46, 11–16. 10.1016/j.semcdb.2015.10.027

12. Callebaut, I., Labesse, G., Durand, P., Poupon, A., Canard, L., Chomilier, J., Henrissat, B., Mornon, J.P., 1997. Deciphering protein sequence information through hydrophobic cluster analysis (HCA): current status and perspectives. Cell Mol Life Sci, 53, 621–645. 10.1007/s000180050082

13. Cam, N., Benzerara, K., Georgelin, T., Jaber, M., Lambert, J.-F., Poinsot, M., Skouri-Panet, F., Cordier, L., 2016. Selective uptake of alkaline earth metals by cyanobacteria forming intracellular carbonates. Environ. Sci. Technol. 50, 11654–11662. 10.1021/acs.est.6b02872

14. Cam, N., Benzerara, K., Georgelin, T., Jaber, M., Lambert, J.-F., Poinsot, M., Skouri-Panet, F., Moreira, D., Lopez-Garcia, P., Raimbault, E., Cordier, L., Jezequel, D., 2018. Cyanobacterial formation of intracellular Ca-carbonates in undersaturated solutions. Geobiology 16, 49–61. 10.1111/gbi.12261

15. Cam, N., Georgelin, T., Jaber, M., Lambert, J.-F., Benzerara, K., 2015. In vitro synthesis of amorphous Mg-, Ca-, Sr- and Ba-carbonates: What do we learn about intracellular calcification by cyanobacteria? Geochimica et Cosmochimica Acta 161, 36–49. 10.1016/j.gca.2015.04.003

16. Chen, S., Gagnon, A.C., Adkins, J.F., 2018. Carbonic anhydrase, coral calcification and a new model of stable isotope vital effects. Geochimica et Cosmochimica Acta 236, 179–197. 10.1016/j.gca.2018.02.032

17. Christiano, R., Nagaraj, N., Fröhlich, F., Walther, T.C., 2014. Global proteome turnover analyses of the yeasts *S. cerevisiae* and *S. pombe*. Cell Reports 9, 1959–1965. 10.1016/j.celrep.2014.10.065

18. Cosmidis, J., Benzerara, K., 2022. Why do microbes make minerals? Comptes Rendus. Géoscience 354, 1–39. 10.5802/crgeos.107

19. Couradeau, E., Benzerara, K., Gerard, E., Moreira, D., Bernard, S., Brown, G.E., Lopez-Garcia, P., 2012. An early-branching microbialite cyanobacterium forms intracellular carbonates. Science 336, 459–462. 10.1126/science.1216171

20. Cumsille, A., Durán, R.E., Rodríguez-Delherbe, A., Saona-Urmeneta, V., Cámara, B., Seeger, M., Araya, M., Jara, N., Buil-Aranda, C., 2023. GenoVi, an open-source automated circular genome visualizer for bacteria and archaea. PLOS Computational Biology 19, e1010998. 10.1371/journal.pcbi.1010998

21. De Wever, A., Benzerara, K., Coutaud, M., Caumes, G., Poinsot, M., Skouri-Panet, F., Laurent, T., Duprat, E., Gugger, M., 2019. Evidence of high Ca uptake by cyanobacteria forming intracellular CaCO_3_ and impact on their growth. Geobiology 17, 676–690. 10.1111/gbi.12358

22. Dick, G.J., Duhaime, M.B., Evans, J.T., Errera, R.M., Godwin, C.M., Kharbush, J.J., Nitschky, H.S., Powers, M.A., Vanderploeg, H.A., Schmidt, K.C., Smith, D.J., Yancey, C.E., Zwiers, C.C., Denef, V.J., 2021. The genetic and ecophysiological diversity of *Microcystis*. Environmental Microbiology 23, 7278–7313. 10.1111/1462-2920.15615

23. Domínguez, D.C., Guragain, M., Patrauchan, M., 2015. Calcium binding proteins and calcium signaling in prokaryotes. *Cell Calcium*, Evolution of Calcium Signaling 57, 151–165. 10.1016/j.ceca.2014.12.006

24. Feldbauer, R., Gosch, L., Lüftinger, L., Hyden, P., Flexer, A., Rattei, T., 2021. DeepNOG: fast and accurate protein orthologous group assignment. Bioinformatics 36, 5304–5312. 10.1093/bioinformatics/btaa1051

25. Gaëtan, J., Halary, S., Millet, M., Bernard, C., Duval, C., Hamlaoui, S., Hecquet, A., Gugger, M., Marie, B., Mehta, N., Moreira, D., Skouri-Panet, F., Travert, C., Duprat, E., Leloup, J., Benzerara, K., 2023. Widespread formation of intracellular calcium carbonates by the bloom-forming cyanobacterium *Microcystis*. Environmental Microbiology 25, 751–765. 10.1111/1462-2920.16322

26. Gancedo, J.M., López, S., Ballesteros, F., 1982. Calculation of half-lives of proteins in vivo. Heterogeneity in the rate of degradation of yeast proteins. Mol Cell Biochem 43, 89–95. 10.1007/BF00423096

27. Gaschignard, G., Millet, M., Bruley, A., Benzerara, K., Dezi, M., Skouri-Panet, F., Duprat, E., Callebaut, I., 2024. AlphaFold2-guided description of CoBaHMA, a novel family of bacterial domains within the heavy-metal-associated superfamily. Proteins, 1-19. 10.1002/prot.26668

28. Gilbert, P.U.P.A., Bergmann, K.D., Boekelheide, N., Tambutté, S., Mass, T., Marin, F., Adkins, J.F., Erez, J., Gilbert, B., Knutson, V., Cantine, M., Hernández, J.O., Knoll, A.H., 2022. Biomineralization: Integrating mechanism and evolutionary history. Science Advances 8, eabl9653. 10.1126/sciadv.abl9653

29. Görgen, S., Benzerara, K., Skouri-Panet, F., Gugger, M., Chauvat, F., Cassier-Chauvat, C., 2020. The diversity of molecular mechanisms of carbonate biomineralization by bacteria. Discov Mater 1, 2. 10.1007/s43939-020-00001-9

30. Gu, P., Li, Q., Zhang, W., Zheng, Z., Luo, X., 2020. Effects of different metal ions (Ca, Cu, Pb, Cd) on formation of cyanobacterial blooms. Ecotoxicol. Environ. Saf. 189, 109976. 10.1016/j.ecoenv.2019.109976

31. Harke, M.J., Steffen, M.M., Gobler, C.J., Otten, T.G., Wilhelm, S.W., Wood, S.A., Paerl, H.W., 2016. A review of the global ecology, genomics, and biogeography of the toxic cyanobacterium, *Microcystis* spp. Harmful Algae 54, 4–20. 10.1016/j.hal.2015.12.007

32. Hirschi, K.D., Zhen, R.G., Cunningham, K.W., Rea, P.A., Fink, G.R., 1996. CAX1, an H^+^/Ca^2+^ antiporter from *Arabidopsis*. Proceedings of the National Academy of Sciences 93, 8782–8786. 10.1073/pnas.93.16.8782

33. Huang, J., Wang, J., Xu, H., 2014. The circadian rhythms of photosynthesis, ATP content and cell division in *Microcystis aeruginosa* PCC7820. Acta Physiol Plant 36, 3315–3323. 10.1007/s11738-014-1699-1

34. Ingolia, N.T., Ghaemmaghami, S., Newman, J.R.S., Weissman, J.S., 2009. Genome-wide analysis in vivo of translation with nucleotide resolution using ribosome profiling. Science 324, 218–223. 10.1126/science.1168978

35. Knoll, A.H., 2003. Biomineralization and evolutionary history. Reviews in Mineralogy and Geochemistry 54, 329–356. 10.2113/0540329

36. Kupriyanova, E.V., Pronina, N.A., Los, D.A., 2023. Adapting from low to high: an update to CO_2_- concentrating mechanisms of cyanobacteria and microalgae. Plants 12, 1569. doi: 10.3390/plants12071569.

37. Lambert, I., Paysant-Le Roux, C., Colella, S., Martin-Magniette, M.-L., 2020. DiCoExpress: a tool to process multifactorial RNAseq experiments from quality controls to co-expression analysis through differential analysis based on contrasts inside GLM models. Plant Methods 16, 68. 10.1186/s13007-020-00611-7

38. Le Roy, N., Jackson, D.J., Marie, B., Ramos-Silva, P., Marin, F., 2014. The evolution of metazoan α- carbonic anhydrases and their roles in calcium carbonate biomineralization. Frontiers in Zoology 11, 75. 10.1186/s12983-014-0075-8

39. Lê, S., Josse, J., Husson, F., 2008. FactoMineR: an R package for multivariate analysis. Journal of Statistical Software 25, 1–18. 10.18637/jss.v025.i01

40. Li, J., Margaret Oliver, I., Cam, N., Boudier, T., Blondeau, M., Leroy, E., Cosmidis, J., Skouri-Panet, F., Guigner, J.-M., Férard, C., Poinsot, M., Moreira, D., Lopez-Garcia, P., Cassier-Chauvat, C., Chauvat, F., Benzerara, K., 2016. Biomineralization patterns of intracellular carbonatogenesis in cyanobacteria: molecular hypotheses. Minerals 6, 10. 10.3390/min6010010

41. Liu, P., Liu, Y., Ren, X., Zhang, Z., Zhao, X., Roberts, A.P., Pan, Y., Li, J., 2021. A novel magnetotactic alphaproteobacterium producing intracellular magnetite and calcium-bearing minerals. Applied and Environmental Microbiology 87, e0155621. 10.1128/AEM.01556-21

42. Liu, P., Zheng, Y., Zhang, R., Bai, J., Zhu, K., Benzerara, K., Menguy, N., Zhao, X., Roberts, A.P., Pan, Y., Li, J., 2023. Key gene networks that control magnetosome biomineralization in magnetotactic bacteria. National Science Review 10, nwac238. 10.1093/nsr/nwac238

43. Mackinder, L., Wheeler, G., Schroeder, D., von Dassow, P., Riebesell, U., Brownlee, C., 2011. Expression of biomineralization-related ion transport genes in *Emiliania huxleyi*. Environmental Microbiology 13, 3250–3265. 10.1111/j.1462-2920.2011.02561.x

44. Mantione, K.J., Kream, R.M., Kuzelova, H., Ptacek, R., Raboch, J., Samuel, J.M., Stefano, G.B., 2014. Comparing bioinformatic gene expression profiling methods: microarray and RNA-Seq. Med Sci Monit Basic Res 20, 138–141. 10.12659/MSMBR.892101

45. Marin, F., Corstjens, P., de Gaulejac, B., de Vrind-De Jong, E., Westbroek, P., 2000. Mucins and molluscan calcification: molecular characterization of mucoperlin, a novel mucin-like protein from the nacreous shell layer of the fan mussel *Pinna nobilis* (Bivalvia, pteriomorphia). Journal of Biological Chemistry 275, 20667–20675. 10.1074/jbc.M003006200

46. Marin, F., Luquet, G., 2004. Molluscan shell proteins. Comptes Rendus Palevol 3, 469–492. 10.1016/j.crpv.2004.07.009

47. Marin, F., Smith, M., Isa, Y., Muyzer, G., Westbroek, P., 1996. Skeletal matrices, muci, and the origin of invertebrate calcification. Proceedings of the National Academy of Sciences 93, 1554–1559. 10.1073/pnas.93.4.1554

48. Mehta, N., Benzerara, K., Kocar, B.D., Chapon, V., 2019. Sequestration of radionuclides Radium-226 and Strontium-90 by cyanobacteria forming intracellular calcium carbonates. Environ. Sci. Technol. 53, 12639–12647. 10.1021/acs.est.9b03982

49. Mehta, N., Bougoure, J., Kocar, B.D., Duprat, E., Benzerara, K., 2022a. Cyanobacteria accumulate radium (^226^Ra) within intracellular amorphous calcium carbonate inclusions. ACS EST Water 2, 616–623. 10.1021/acsestwater.1c00473

50. Mehta, N., Gaëtan, J., Giura, P., Azaïs, T., Benzerara, K., 2022b. Detection of biogenic amorphous calcium carbonate (ACC) formed by bacteria using FTIR spectroscopy. Spectrochimica Acta Part A: Molecular and Biomolecular Spectroscopy 278, 121262. 10.1016/j.saa.2022.121262

51. Mehta, N., Vantelon, D., Gaëtan, J., Fernandez-Martinez, A., Delbes, L., Travert, C., Benzerara, K., 2023. Calcium speciation and coordination environment in intracellular amorphous calcium carbonate (ACC) formed by cyanobacteria. Chemical Geology 641, 121765. 10.1016/j.chemgeo.2023.121765

52. Mirdita, M., Schütze, K., Moriwaki, Y., Heo, L., Ovchinnikov, S., Steinegger, M., 2022. ColabFold: making protein folding accessible to all. Nat Methods 19, 679–682. 10.1038/s41592-022-01488-1

53. Monteil, C.L., Benzerara, K., Menguy, N., Bidaud, C.C., Michot-Achdjian, E., Bolzoni, R., Mathon, F.P., Coutaud, M., Alonso, B., Garau, C., Jézéquel, D., Viollier, E., Ginet, N., Floriani, M., Swaraj, S., Sachse, M., Busigny, V., Duprat, E., Guyot, F., Lefevre, C.T., 2021. Intracellular amorphous Ca-carbonate and magnetite biomineralization by a magnetotactic bacterium affiliated to the Alphaproteobacteria. ISME J 15, 1–18. 10.1038/s41396-020-00747-3

54. Monteil, C.L., Perrière, G., Menguy, N., Ginet, N., Alonso, B., Waisbord, N., Cruveiller, S., Pignol, D., Lefèvre, C.T., 2018. Genomic study of a novel magnetotactic Alphaproteobacteria uncovers the multiple ancestry of magnetotaxis. Environmental Microbiology 20, 4415–4430. 10.1111/1462-2920.14364

55. Monteiro, F.M., Bach, L.T., Brownlee, C., Bown, P., Rickaby, R.E.M., Poulton, A.J., Tyrrell, T., Beaufort, L., Dutkiewicz, S., Gibbs, S., Gutowska, M.A., Lee, R., Riebesell, U., Young, J., Ridgwell, A., 2016. Why marine phytoplankton calcify. Science Advances 2, e1501822. 10.1126/sciadv.1501822

56. Patro, R., Duggal, G., Love, M.I., Irizarry, R.A., Kingsford, C., 2017. Salmon provides fast and bias-aware quantification of transcript expression. Nat Methods 14, 417–419. 10.1038/nmeth.4197

57. Pattanayak, G., Rust, M.J., 2014. The cyanobacterial clock and metabolism. Current Opinion in Microbiology 18, 90–95. 10.1016/j.mib.2014.02.010

58. Paysan-Lafosse, T., Blum, M., Chuguransky, S., Grego, T., Pinto, B.L., Salazar, G.A., Bileschi, M.L., Bork, P., Bridge, A., Colwell, L., Gough, J., Haft, D.H., Letunić, I., Marchler-Bauer, A., Mi, H., Natale, D.A., Orengo, C.A., Pandurangan, A.P., Rivoire, C., Sigrist, C.J.A., Sillitoe, I., Thanki, N., Thomas, P.D., Tosatto, S.C.E., Wu, C.H., Bateman, A., 2023. InterPro in 2022. Nucleic Acids Research 51, D418–D427. 10.1093/nar/gkac993

59. Pettersen, E.F., Goddard, T.D., Huang, C.C., Couch, G.S., Greenblatt, D.M., Meng, E.C., Ferrin, T.E., 2004. UCSF Chimera-a visualization system for exploratory research and analysis. J Comput Chem 25, 1605–1612. 10.1002/jcc.20084

60. Price, G.D., Howitt, S.M., 2011. The cyanobacterial bicarbonate transporter *BicA*: its physiological role and the implications of structural similarities with human SLC26 transporters. Biochem. Cell Biol. 89, 178–188. 10.1139/O10-136

61. Ragon, M., Benzerara, K., Moreira, D., Tavera, R., Lopez-Garcia, P., 2014. 16S rDNA-based analysis reveals cosmopolitan occurrence but limited diversity of two cyanobacterial lineages with contrasted patterns of intracellular carbonate mineralization. Front. Microbiol. 5, 331. 10.3389/fmicb.2014.00331

62. Riba, A., Di Nanni, N., Mittal, N., Arhné, E., Schmidt, A., Zavolan, M., 2019. Protein synthesis rates and ribosome occupancies reveal determinants of translation elongation rates. Proceedings of the National Academy of Sciences 116, 15023–15032. 10.1073/pnas.1817299116

63. Rippka, R., Deruelles, J., Waterbury, J.B., Herdman, M., Stanier, R.Y., 1979. Generic assignments, strain histories and properties of pure cultures of cyanobacteria. Microbiology 111, 1–61. 10.1099/00221287-111-1-1

64. Rivadeneyra, M.A., Martín-Algarra, A., Sánchez-Navas, A., Martíin-Ramos, D. 2006. Carbonate and phosphate precipitation by Chromohalobacter marismortui, Geomicrobiology Journal, 23, 89–101. 10.1080/01490450500533882

65. Rivadeneyra, M.A., Martín-Algarra, A., Sánchez-Román, M., Sánchez-Navas, A., Martín-Ramos, J.D., 2010. Amorphous Ca-phosphate precursors for Ca-carbonate biominerals mediated by *Chromohalobacter marismortui*. ISME J, 4, 922–932. 10.1038/ismej.2010.17

66. Robinson, M.D., Oshlack, A., 2010. A scaling normalization method for differential expression analysis of RNA-seq data. Genome Biology 11, R25. 10.1186/gb-2010-11-3-r25

67. Sandrini, G., Cunsolo, S., Schuurmans, J.M., Matthijs, H.C.P., Huisman, J., 2015. Changes in gene expression, cell physiology and toxicity of the harmful cyanobacterium *Microcystis aeruginosa* at elevated CO_2_. Frontiers in Microbiology 6, 401. 10.3389/fmicb.2015.00401

68. Schwanhäusser, B., Busse, D., Li, N., Dittmar, G., Schuchhardt, J., Wolf, J., Chen, W., Selbach, M., 2011. Global quantification of mammalian gene expression control. Nature 473, 337–342. 10.1038/nature10098

69. Shigaki, T., Rees, I., Nakhleh, L., Hirschi, K.D., 2006. Identification of three distinct phylogenetic groups of CAX cation/proton antiporters. J Mol Evol 63, 815–825. 10.1007/s00239-006-0048-4

70. Stöckel, J., Welsh, E.A., Liberton, M., Kunnvakkam, R., Aurora, R., Pakrasi, H.B., 2008. Global transcriptomic analysis of *Cyanothece* 51142 reveals robust diurnal oscillation of central metabolic processes. Proceedings of the National Academy of Sciences 105, 6156–6161. 10.1073/pnas.0711068105

71. Straub, C., Quillardet, P., Vergalli, J., Tandeau de Marsac, N., Humbert, J.-F., 2011. A day in the life of *Microcystis aeruginosa* strain PCC 7806 as revealed by a transcriptomic analysis. PLOS ONE 6, e16208. 10.1371/journal.pone.0016208

72. Tang, J., Zhou, H., Yao, D., Riaz, S., You, D., Klepacz-Smółka, A., Daroch, M., 2022. Comparative genomic analysis revealed distinct molecular components and organization of CO_2_- concentrating mechanism in thermophilic cyanobacteria. Frontiers in Microbiology 13, 876272. 10.3389/fmicb.2022.876272

73. Taoka, A., Eguchi, Y., Shimoshige, R., Fukumori, Y., 2023. Recent advances in studies on magnetosome-associated proteins composing the bacterial geomagnetic sensor organelle. Microbiology and Immunology 67, 228–238. 10.1111/1348-0421.13062

74. Uebe, R., Schüler, D., 2016. Magnetosome biogenesis in magnetotactic bacteria. Nat Rev Microbiol 14, 621–637. 10.1038/nrmicro.2016.99

75. van Kempen, M., Kim, S.S., Tumescheit, C., Mirdita, M., Lee, J., Gilchrist, C.L.M., Söding, J., Steinegger, M., 2024. Fast and accurate protein structure search with Foldseek. Nat Biotechnol 42, 243–246. 10.1038/s41587-023-01773-0

76. Waditee, R., Hibino, T., Tanaka, Y., Nakamura, T., Incharoensakdi, A., Takabe, T., 2001. Halotolerant cyanobacterium *Aphanothece halophytica* contains an Na(+)/H(+) antiporter, homologous to eukaryotic ones, with novel ion specificity affected by C-terminal tail. J Biol Chem. 276, 36931–36938. 10.1074/jbc.M10365020

77. Waditee, R., Hossain, G.S., Tanaka, Y., Nakamura, T., Shikata, M., Takano, J., Takabe, Tetsuko, Takabe, Teruhiro, 2004. Isolation and functional characterization of Ca^2+^/H^+^ antiporters from Cyanobacteria. J. Biol. Chem. 279, 4330–4338. 10.1074/jbc.M310282200

78. Waight, A.B., Pedersen, B.P., Schlessinger, A., Bonomi, M., Chau, B.H., Roe-Zurz, Z., Risenmay, A.J., Sali, A., Stroud, R.M., 2013. Structural basis for alternating access of a eukaryotic calcium/proton exchanger. Nature 499, 107–110. 10.1038/nature12233

79. Wang, Z., Gerstein, M., Snyder, M., 2009. RNA-Seq: a revolutionary tool for transcriptomics. Nat Rev Genet 10, 57–63. 10.1038/nrg2484

80. Welkie, D.G., Rubin, B.E., Diamond, S., Hood, R.D., Savage, D.F., Golden, S.S., 2019. A Hard Day’s Night: cyanobacteria in diel cycles. Trends in Microbiology 27, 231–242. 10.1016/j.tim.2018.11.002

81. Weyhenmeyer, G.A., Hartmann, J., Hessen, D.O., Kopáček, J., Hejzlar, J., Jacquet, S., Hamilton, S.K., Verburg, P., Leach, T.H., Schmid, M., Flaim, G., Nõges, T., Nõges, P., Wentzky, V.C., Rogora, M., Rusak, J.A., Kosten, S., Paterson, A.M., Teubner, L., Higgins, S.N., Lawrence, G., Kangur, K., Kokorite, I., Cerasino, L., Funk, C., Harvey, R., Moatar, F., de Wit, H.A., Zechmeister, T., 2019. Widespread diminishing anthropogenic effects on calcium in freshwaters. Sci Rep 9, 10450. 10.1038/s41598-019-46838-w

82. Yang, T., Teske, A., Ambrose, W., Salman-Carvalho, V., Bagnell, R., & Nielsen, L. P., 2019. Intracellular calcite and sulfur dynamics of *Achromatium* cells observed in a lab-based enrichment and aerobic incubation experiment. Antonie van Leeuwenhoek, 112, 263-274. 10.1007/s10482-018-1153-2

83. Zhao, S., Fung-Leung, W.-P., Bittner, A., Ngo, K., Liu, X., 2014. Comparison of RNA-Seq and microarray in transcriptome profiling of activated T cells. PLOS ONE 9, e78644. 10.1371/journal.pone.0078644

84. Zinser, E.R., Lindell, D., Johnson, Z.I., Futschik, M.E., Steglich, C., Coleman, M.L., Wright, M.A., Rector, T., Steen, R., McNulty, N., Thompson, L.R., Chisholm, S.W., 2009. Choreography of the transcriptome, photophysiology, and cell cycle of a minimal photoautotroph, *Prochlorococcus*. PLOS ONE 4, e5135. 10.1371/journal.pone.0005135

