## Supplementary material for "Diel changes in the expression of a marker gene and candidate genes for intracellular amorphous CaCO_3_ biomineralization in *Microcystis*": Readme_Supporting information

This file describes the supplementary data, files and tables

**CONTENTS**

1. **Supplementary Data**  **(Separate files)**

**Supplementary Data 1:** “DICOEXPRESS_input” folder containing 2 files used as inputs for DICOEXPRESS differential expression analyses. COUNTS.csv file: for each replicate, the estimated transcript abundances as normalized by Salmon according to the transcript size, genome size (number of CDS) and sample size (number of reads); these data are described as total abundance data in the manuscript. TARGET.csv file: list of sample replicate names and labels. More information about DiCoExpress input and output file description can be found in ***Lambert et al. (2019). DiCoExpress: a workspace to process multifactorial RNAseq experiments from quality controls to co-expression analysis through differential analysis based on contrasts inside GLM models. 10.21203/rs.2.19732/v1.***

**Supplementary Data 2**: DICOEXPRESS_output folder containing two subfolders:

a) the “QualityControl_Normalization” subfolder, containing information about quality control and normalization results. During this procedure, 220 low-count transcripts were discarded (list in the PCC7806SL_Low_count_genes file), resulting in a total of 4,440 genes with normalized counts (file PCC7806SL_NormCounts_log2), mean and standard deviation (PCC7806SL_NormCounts_log2_Mean_SD) for each time point.

b) the “DiffAnalysis” subfolder, containing data about the differential expression between time points as tested using a generalized linear model. For comparison between each time point, statistics of differentially expressed genes (DEGs) are provided as Fold Change (FC) and p-values. For each time point pair, the PCC7806_[t*x*-t*xx*]_LRT_BH files provide for all normalized genes the LogFoldChange (logFC), logCountsperMillion (logCPM), Likelihood ratio (LR), p-value and False Discovery Rate (FDR). The PCC7806_[t*x*-t*xx*]_Id_DEG files provide the list of DEGs identifiers. The PCC7806_[t*x*-t*xx*]_DEG.BH files provide for all DEGs the logFC, logCPM, LR, p-values and FDR. The PCC7806_[t*x*-t*xx*]_plotSmear pdf files show a plot the log of the ratio of expression levels for each gene between two experimental groups (the log fold-change) against the overall average expression level for each gene across the two groups (the log-concentration). The DEGs are plotted in red. The PCC7806_[tx-txx]_Top50_Clustering pdf files contain a hierarchical clustering for the top 50 DEGs. The PCC7806_[tx-txx]_Top50_Profile pdf files contain single gene profiles for the top 50 among the DEGs.

**Supplementary Data 3**: “Pearson_pairwise_correlations_4439-profiles_vs_ccyA” folder; Pearson pairwise correlation tests between the temporal expression patterns of each *M. aeruginosa* PCC 7806 gene and the temporal expression pattern of *ccyA.* For each gene, the gene ID, its annotation, the estimated correlation coefficient (rho) and the p-value of the test are provided.

**Supplementary Data 4**: “DeepNOG_annotations” folder; functional annotation of the *M. aeruginosa* PCC 7806 genes (DeepNOG protein orthologous groups assignment; see Methods section in the main manuscript for details).

2. **Supplementary Tables (Separate files)**

**Supplementary Table 1**: One .xlsx file. Pairwise comparisons of *ccyA* transcript abundance at different time points. Abundances are expressed as log base 2 of the normalized counts. Each time point is represented by n=3 replicates. The p-values of two-side pairwise t-tests are indicated and highlighted when significant (* and ** for p-values between 0.01 and 0.05, and below 0.01, respectively).

**Supplementary Table 2**: Two .csv files. File #1: list of the 893 genes plotted in Figure 3 with an expression significantly higher during the day at t3_D (12 am) than during the night at t8_N (7 am) [n893-gray_CDSoverexpressed_at_night.csv]. file #2: list of the 906 genes with an expression significantly higher at t8_N (7 am) than at t3_D (12 am) [file n906-gold_CDSoverexpressed_day]. In both files, access codes, annotations and COG predictions of the CDS are provided as well as the log of the fold change value (FC), the gene expression in count-per-million (CPM), the likelihood ratio (LR), the p-value (PValue), the false discovery rate (FDR).

**Supplementary Table 3**: Two .csv files. File #1: list of the 773 genes whose expression profiles are highly correlated (rho>>0) with that of the *ccyA* gene [n773-gray_CDS_correlated-ccyA.csv]. These genes are plotted in Figure 6 in the grey shaded area. File #2: list of the 706 genes whose expression profiles are highly anticorrelated (rho<<0) with that of the *ccyA* gene [n706-gold_CDS_anticorrelated-ccyA.csv]. These genes are plotted in Figure 6 in the golden shaded area. In both files, access codes, annotations and COG predictions of the CDS are provided as well as the value of the Pearson correlation coefficient (rho.pearson.), the p-value (P.value).

**Supplementary Table 4**: One .xlsx file with two tabs. Tab 1: list of the 24 genes involved in CCM identified in the genome of PCC 7806SL, their NCBI and COG annotations and expression parameters. The fold change (FC) and p-value between night (t8_N) and day (t3_D) are provided (also plotted in figure 3). High positive fold changes from night to day (t3_D>>t8_N) are highlighted in yellow. High negative fold changed are highlighted in grey. Moreover, the Pearson correlation coefficient (rho_ccyA) and the p-value for the correlation between the expression profile of these genes in a day/night cycle and that of *ccyA* are also provided. High correlations between CCM and *ccyA* genes are highlighted in grey; high anticorrelation between CCM and *ccyA* genes are highlighted in yellow. Tab 2: list of the 24 genes involved in Ca transportation identified in the genome of PCC 7806SL, their NCBI and COG annotations and expression parameters. For the fold change (FC) and the Pearson correlation coefficient (rho_ccyA) the yellow/grey color codes are similar as for the CCM genes in tab 1.

3. **Supplementary Figures (Separate files)**

**Supplementary Figure 1:** Hydrophobic cluster analysis (HCA) bidimensional plot of a protein sequence segment from DUF1269 (encoding gene in position +7 relatively to *ccyA*; see Figure 5D for AF2 structure prediction) shown to illustrate the topological features of the GlyZip motif (glycine highlighted in yellow), predicted to fold as a helical hairpin. The sequence is shown on a duplicated alpha-helical net, in which strong hydrophobic amino acids are contoured, forming hydrophobic clusters (see Callebaut et al., 1997 for more details about the HCA method). The GlyZip sequence of the DUF1269 domain-containing protein (top row) is aligned against the first GlyZip motif of the *Microcystis* calcyanin (bottom row).

**Supplementary Figure 2**: Histogram showing the distribution of the Pearson correlation coefficient (rho) between the expression pattern of *ccyA* and that of all other 4439 genes in the transcriptome. The part of the distribution where rho < -0.75 is coloured in yellow (genes with an expression pattern anticorrelated with that of *ccyA*). The part of the distribution where rho > 0.75 is coloured in grey (genes with an expression pattern correlated with that of *ccyA*).

**Supplementary Figure 3**: Comparison of the abundance profiles of transcripts of genes neighboring *ccyA* (solid line) with that of *ccyA* transcripts (dashed line) during a day/night cycle. One page per gene neighboring *ccyA*, total of 20 pages. Abundances are expressed as log base 2 of the normalized counts. Replicates a, b and c and the mean of the abundance are represented by different symbols. For each sampling time (t1-t8), error bar represents the standard error on the mean of the three replicates. Grey shaded areas outline the night periods. On top of the graph, it is indicated when rho <<0 (rho<-0.75), rho>>0 (rho>0.75) and nothing otherwise. The 0.75 treshold was chosen arbitrarily.

**Supplementary Figure 4**: Comparison of the abundance profiles of transcripts of genes involved in the CCM (solid line) with that of *ccyA* transcripts (dashed line) during a day/night cycle. One page per gene involved in the CCM, total of 24 pages. Abundances are expressed as log base 2 of the normalized counts. Replicates a, b and c and the mean of the abundance are represented by different symbols. For each sampling time (t1-t8), error bar represents the standard error on the mean of the three replicates. Grey shaded areas outline the night periods. On top of the graph, it is indicated when rho <<0 (rho<-0.75), rho>>0 (rho>0.75) and nothing otherwise. The 0.75 treshold was chosen arbitrarily.

**Supplementary Figure 5:** Comparison of the abundance profiles of transcripts of genes involved in Ca transport (solid line) with that of *ccyA* transcripts (dashed line) during a day/night cycle. One page per gene involved in the Ca transport, total of 24 pages. Abundances are expressed as log base 2 of the normalized counts. Replicates a, b and c and the mean of the abundance are represented by different symbols. For each sampling time (t1-t8), error bar represents the standard error on the mean of the three replicates. Grey shaded areas outline the night periods. On top of the graph, it is indicated when rho <<0 (rho<-0.75), rho>>0 (rho>0.75) and nothing otherwise. The 0.75 treshold was chosen arbitrarily.
