## Supplementary_Data for "Diel changes in the expression of a marker gene and candidate genes for intracellular amorphous CaCO_3_ biomineralization in *Microcystis*": PCC7806_[t1-t3]_Top50_Profile.pdf

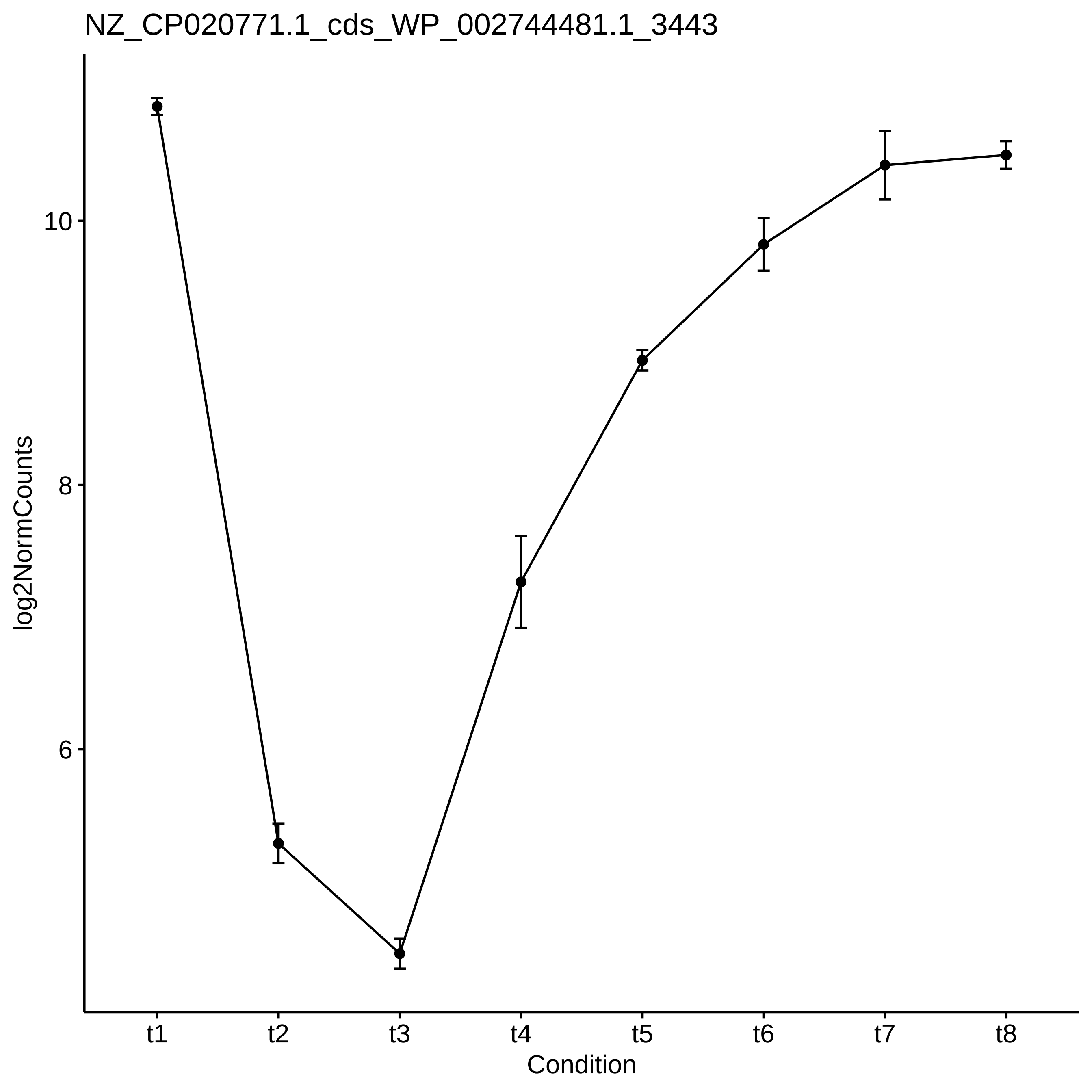
NZ_CP020771.1_cds_WP_002744481.1_3443
log2NormCounts
10
8
6
t1
t2
t3
t4
t5
t6
t7
t8
Condition

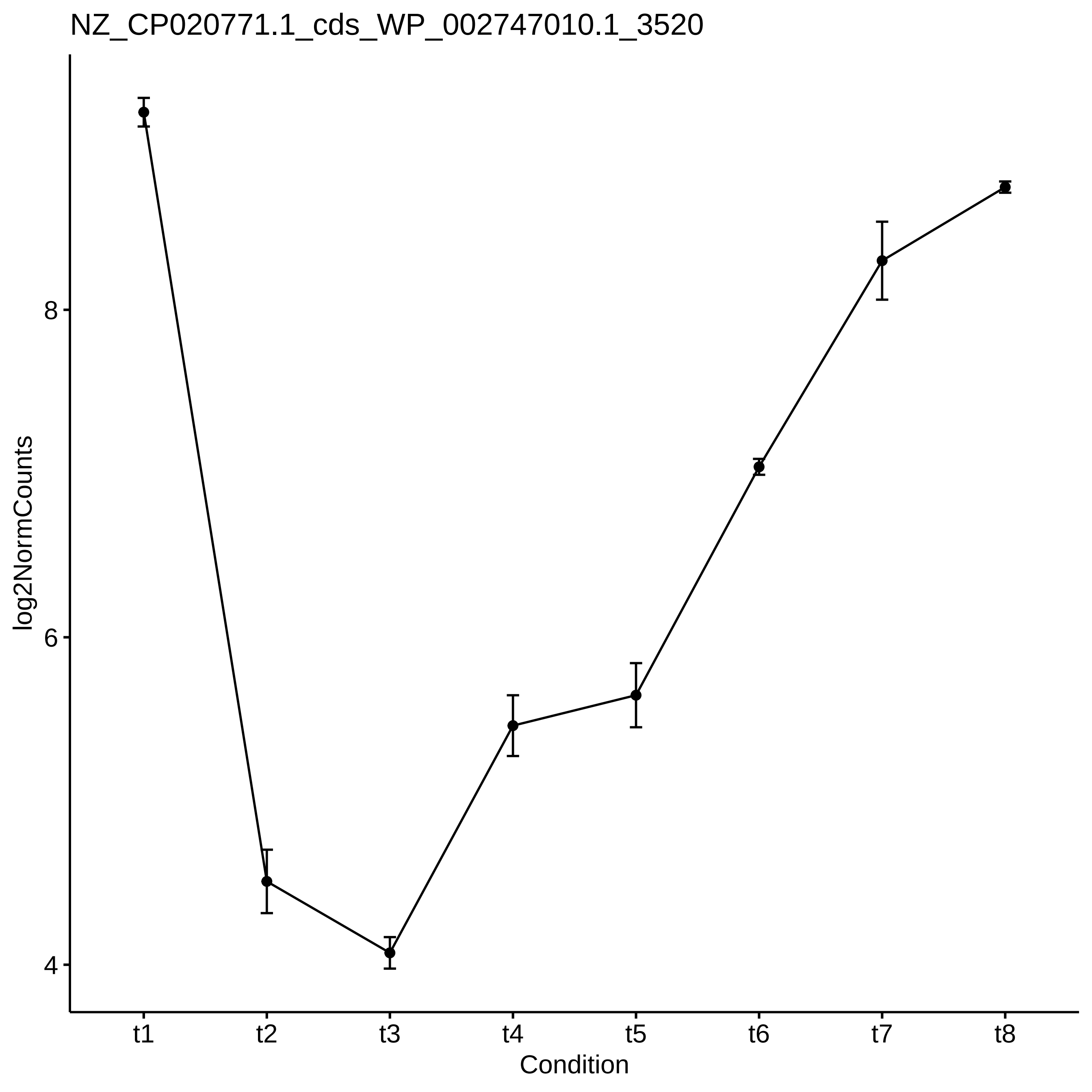
NZ_CP020771.1_cds_WP_002747010.1_3520
log2NormCounts
8
6
4
t1
t2
t3
t4
t5
t6
t7
t8
Condition

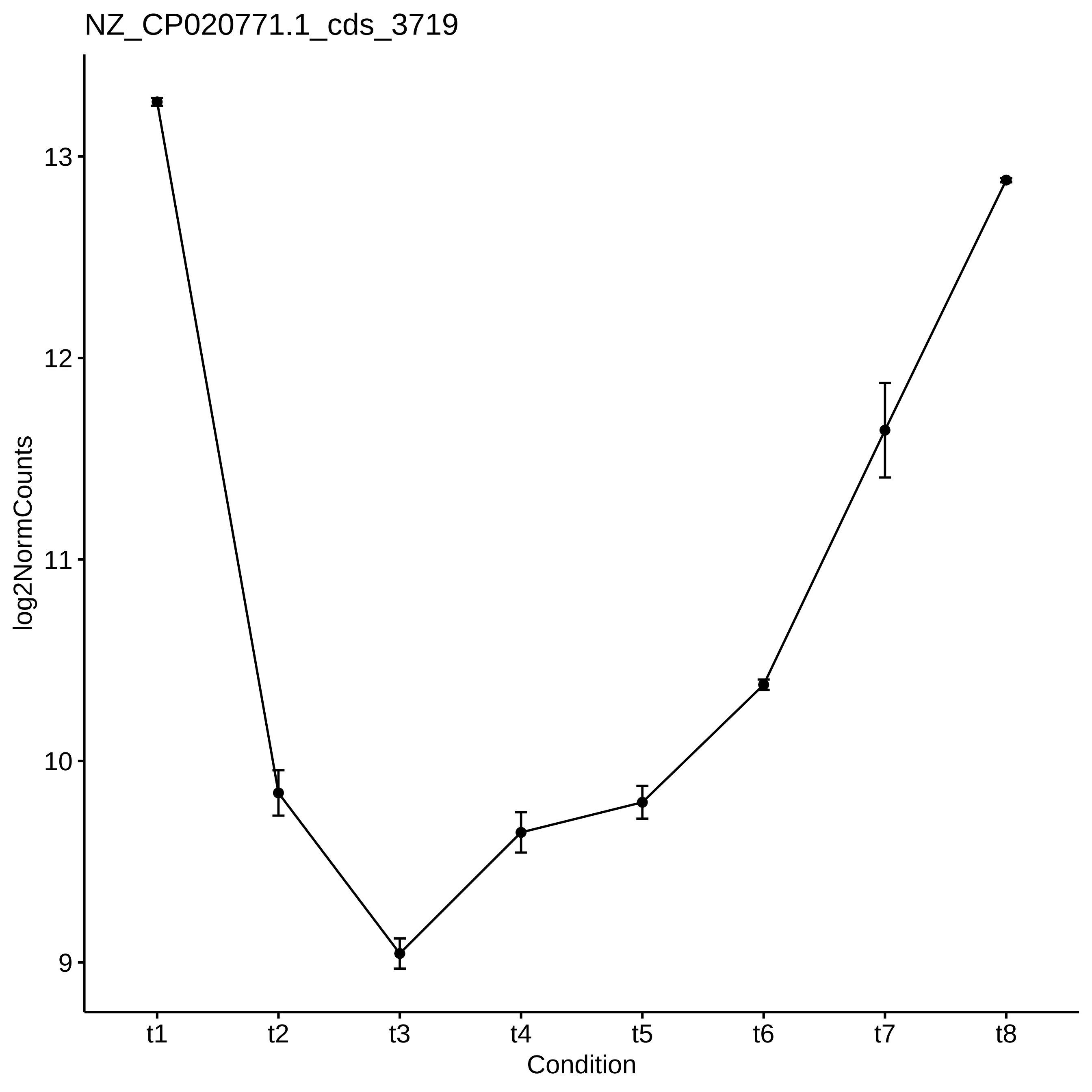
NZ_CP020771.1_cds_3719
log2NormCounts
13
12
11
10
9
t1
t2
t3
t4
t5
t6
t7
t8
Condition

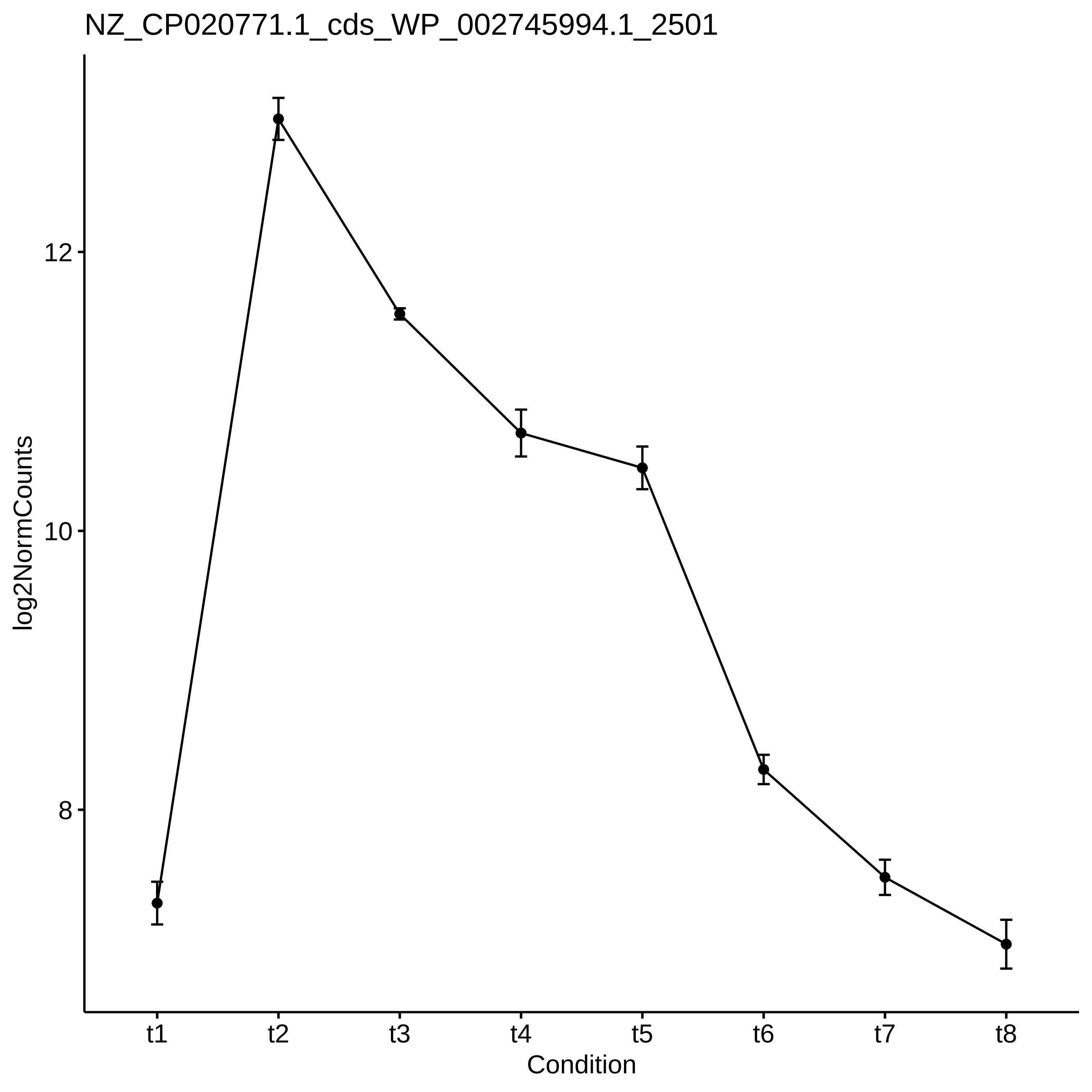
NZ_CP020771.1_cds_WP_002745994.1_2501
12
10
8
log2NormCounts
t1
t2
t3
t4
t5
t6
t7
t8
Condition

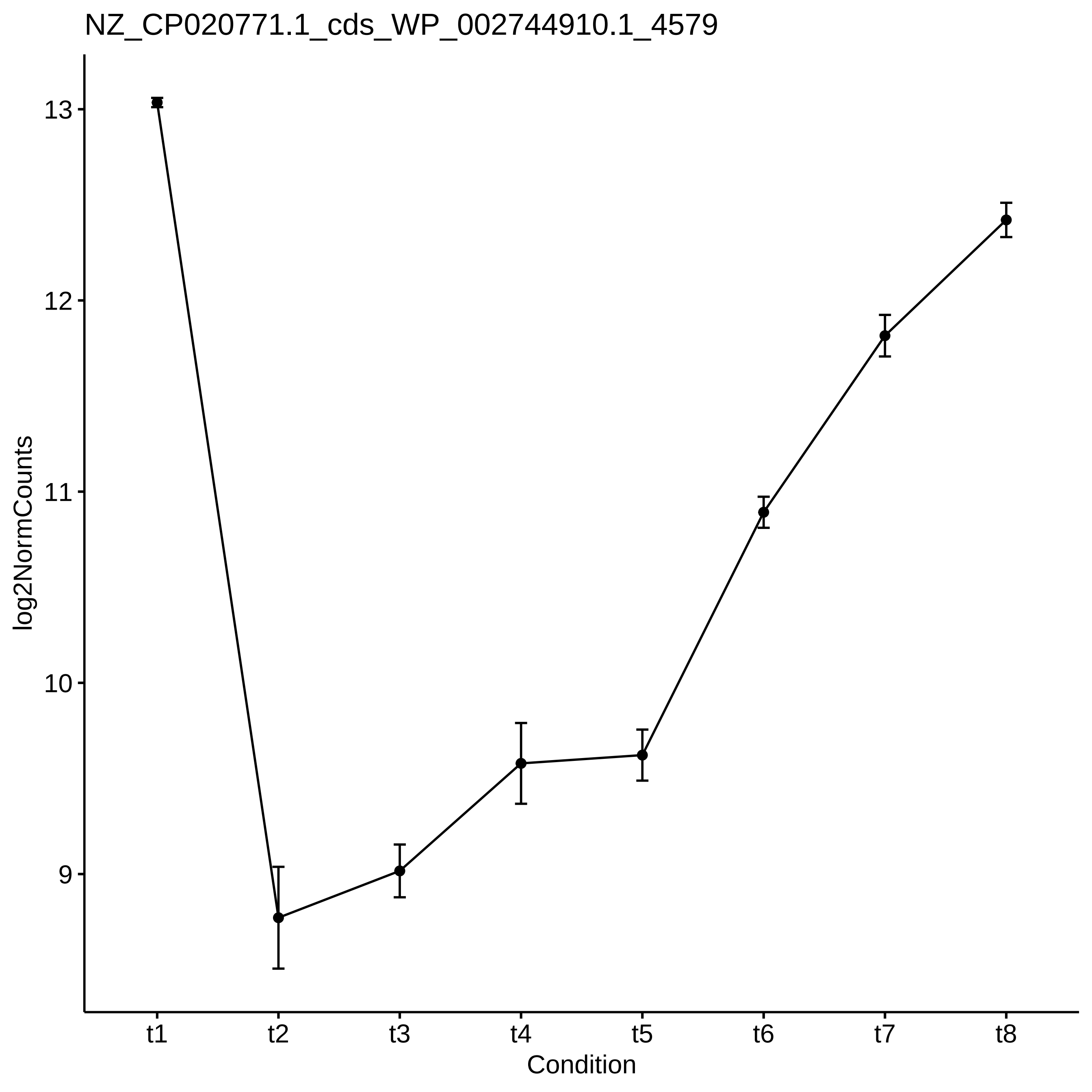
NZ_CP020771.1_cds_WP_002744910.1_4579
log2NormCounts
13
12
11
10
9
t1
t2
t3
t4
t5
t6
t7
t8
Condition

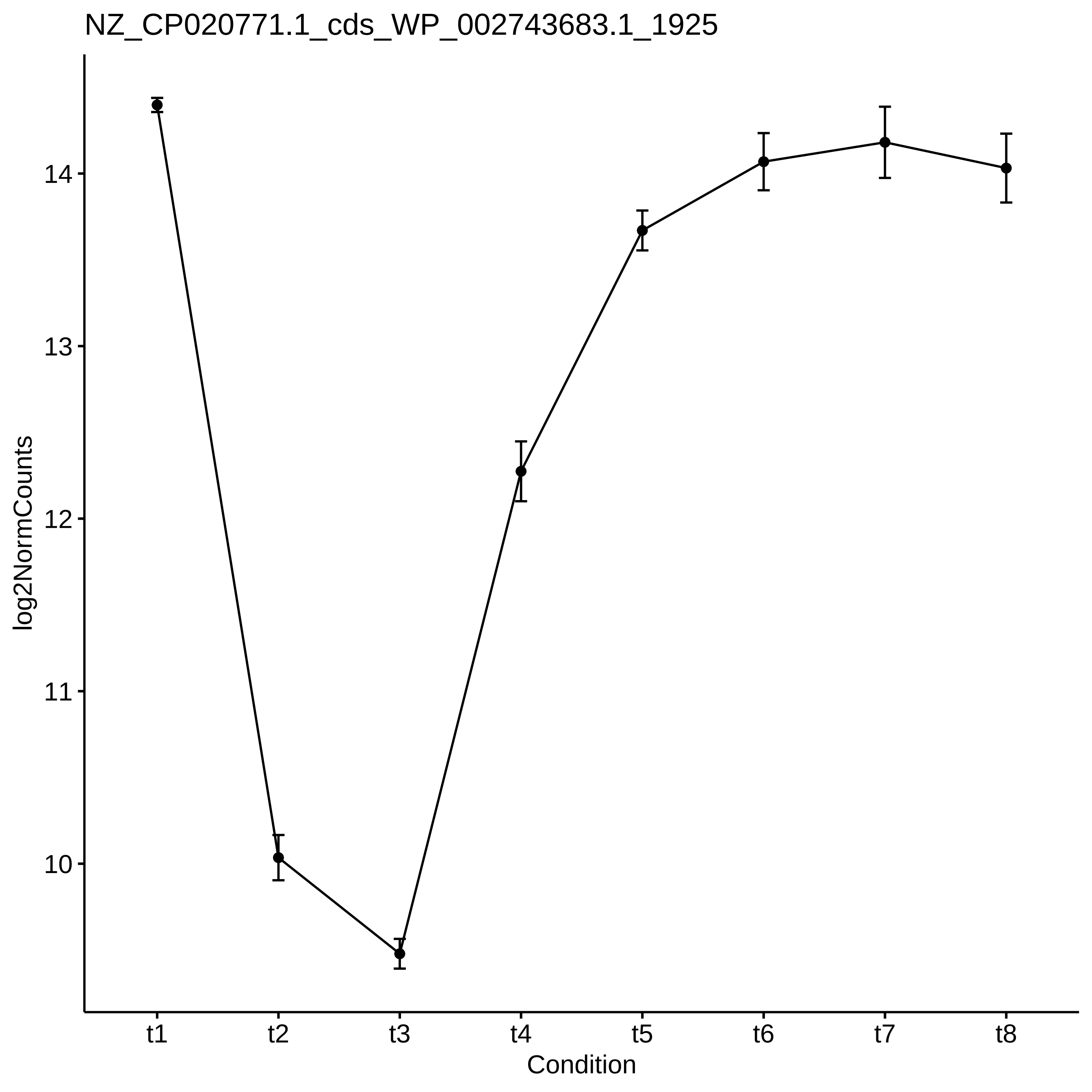

NZ_CP020771.1_cds_WP_002743683.1_1925
log2NormCounts
14
13
12
11
10
t1
t2
t3
t4
t5
t6
t7
t8
Condition

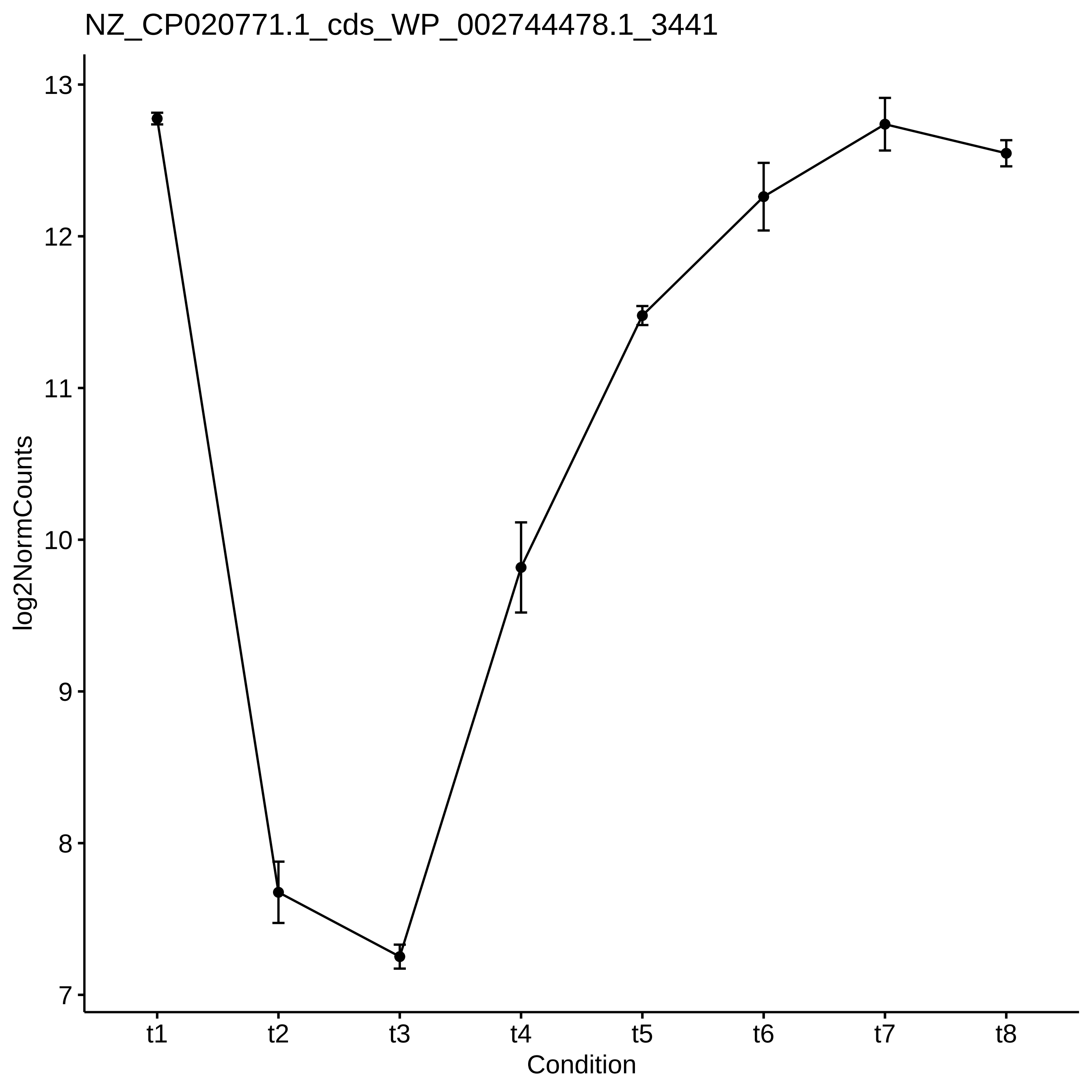

NZ_CP020771.1_cds_WP_002744478.1_3441
log2NormCounts
13
12
11
10
9
8
7
t1
t2
t3
t4
t5
t6
t7
t8
Condition

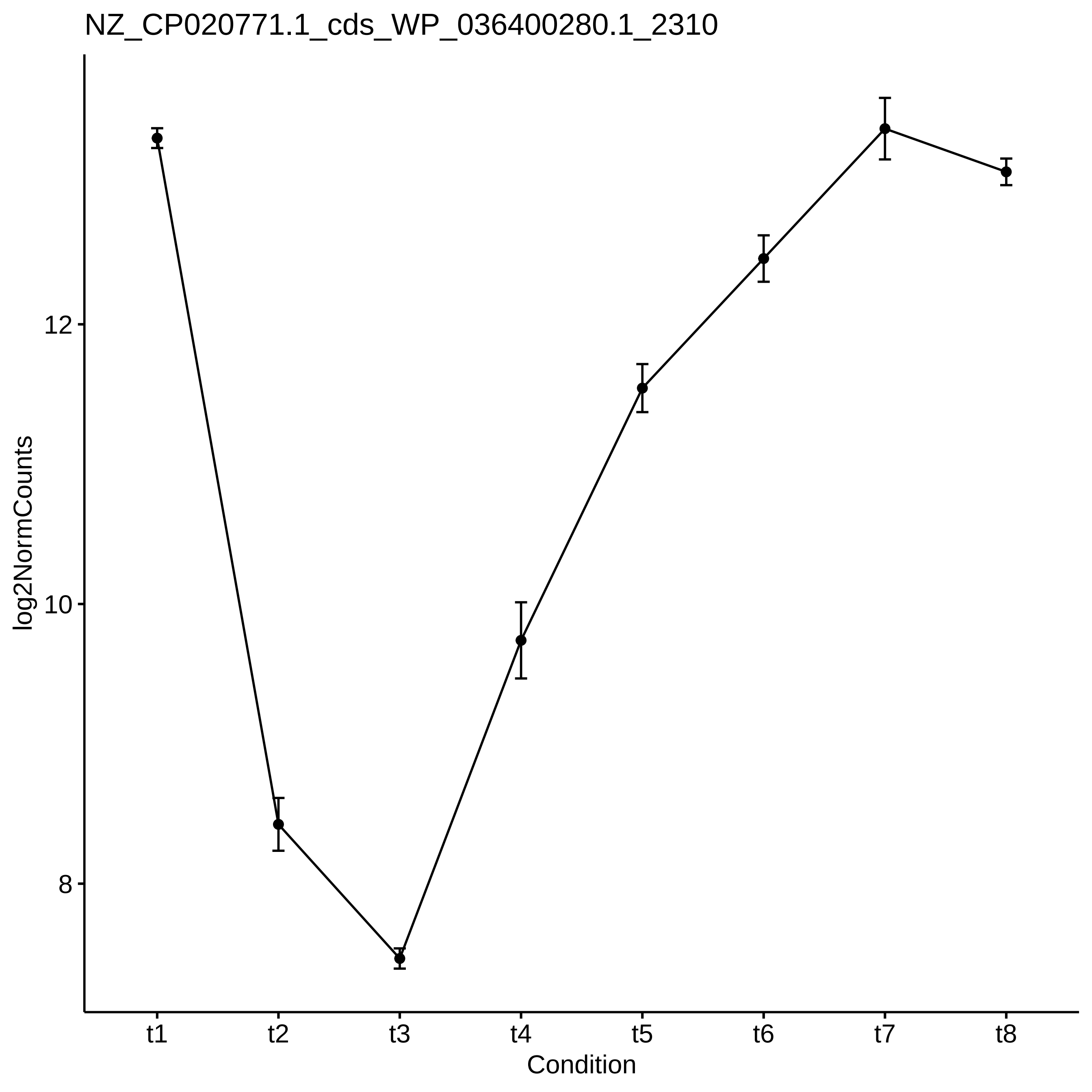
NZ_CP020771.1_cds_WP_036400280.1_2310
log2NormCounts
12
10
8
t1
t2
t3
t4
t5
t6
t7
t8
Condition

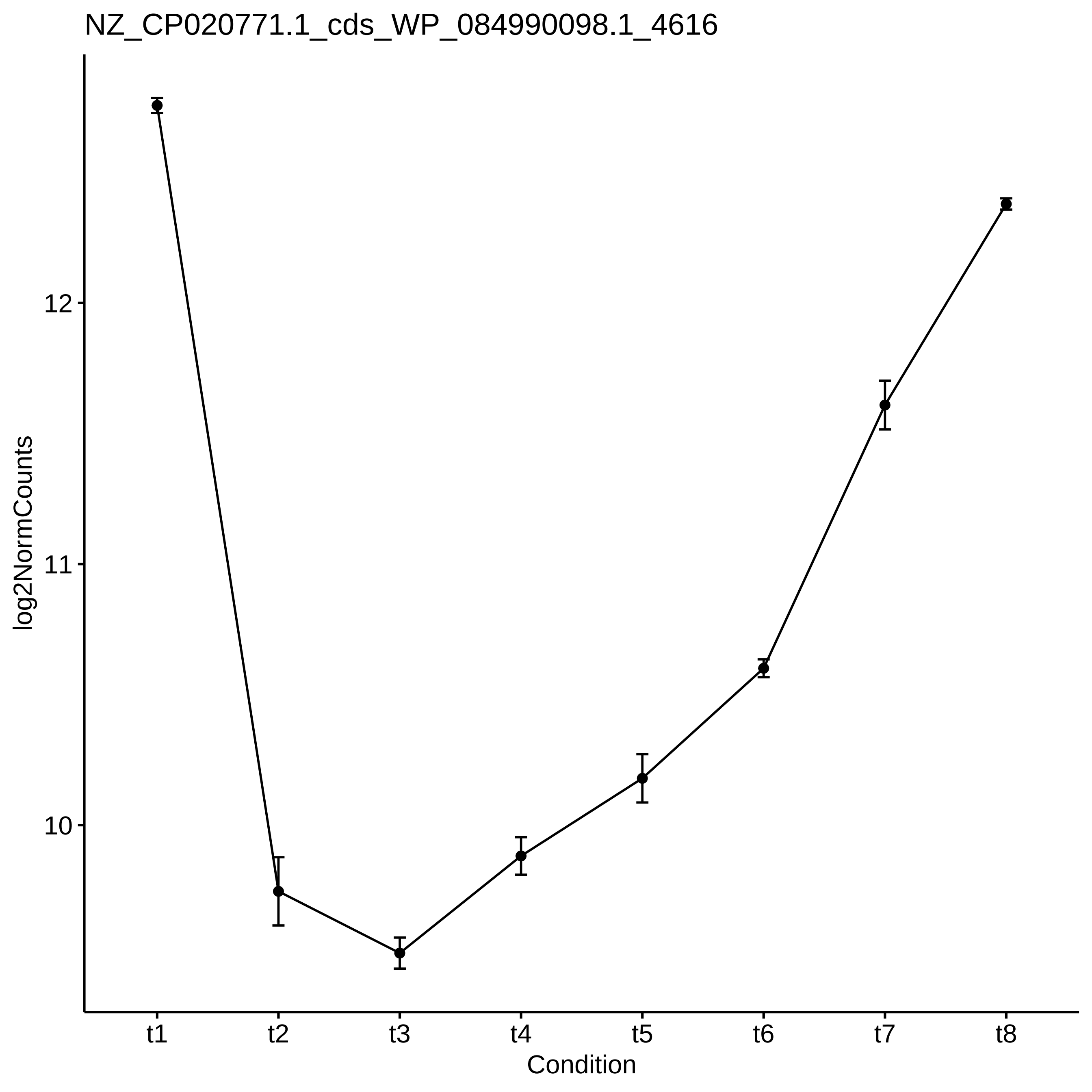
NZ_CP020771.1_cds_WP_084990098.1_4616
log2NormCounts
12
11
10
t1
t2
t3
t4
t5
t6
t7
t8
Condition

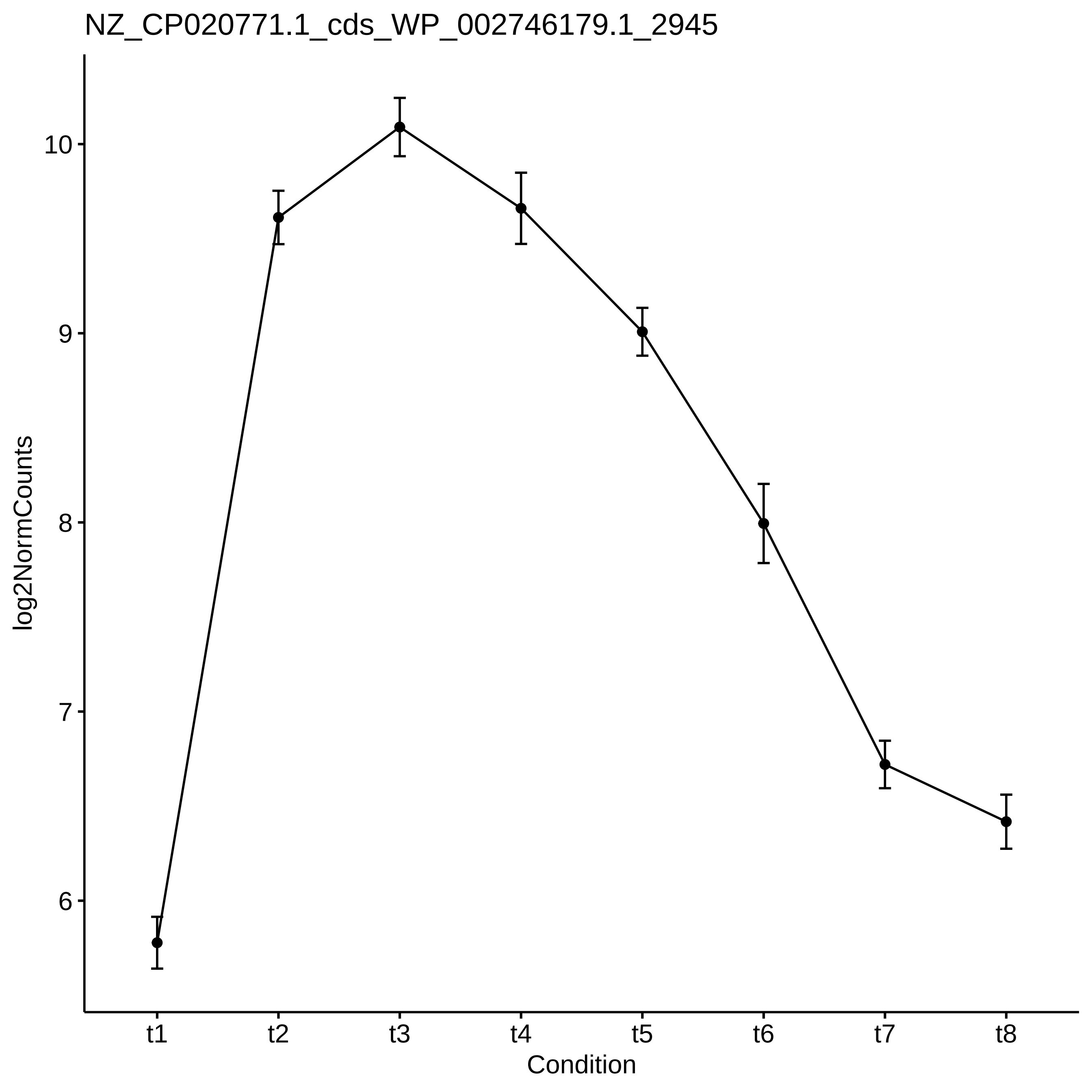
NZ_CP020771.1_cds_WP_002746179.1_2945
log2NormCounts
10
9
8
7
6
t1
t2
t3
t4
t5
t6
t7
t8
Condition

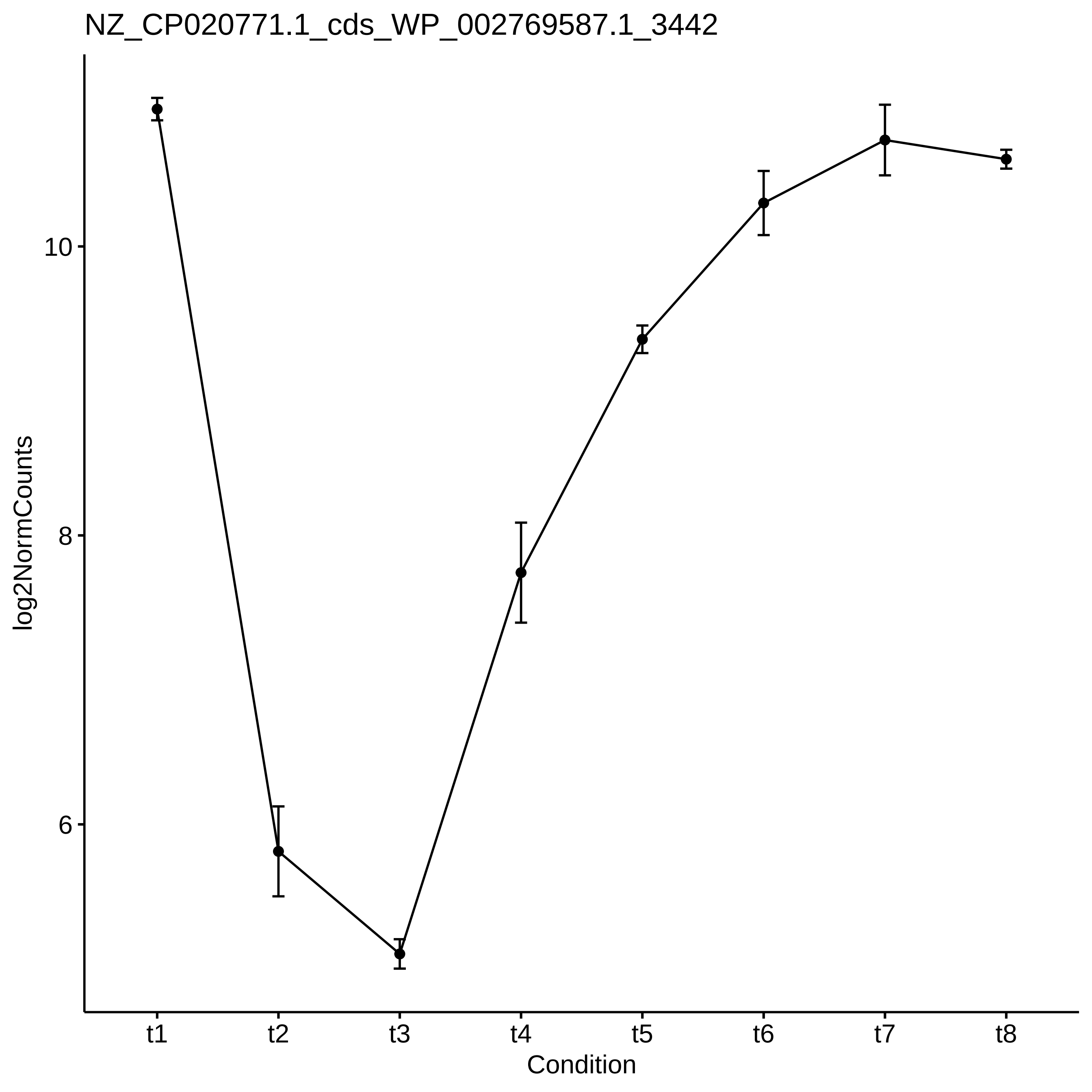
NZ_CP020771.1_cds_WP_002769587.1_3442
log2NormCounts
10
8
6
t1
t2
t3
t4
t5
t6
t7
t8
Condition

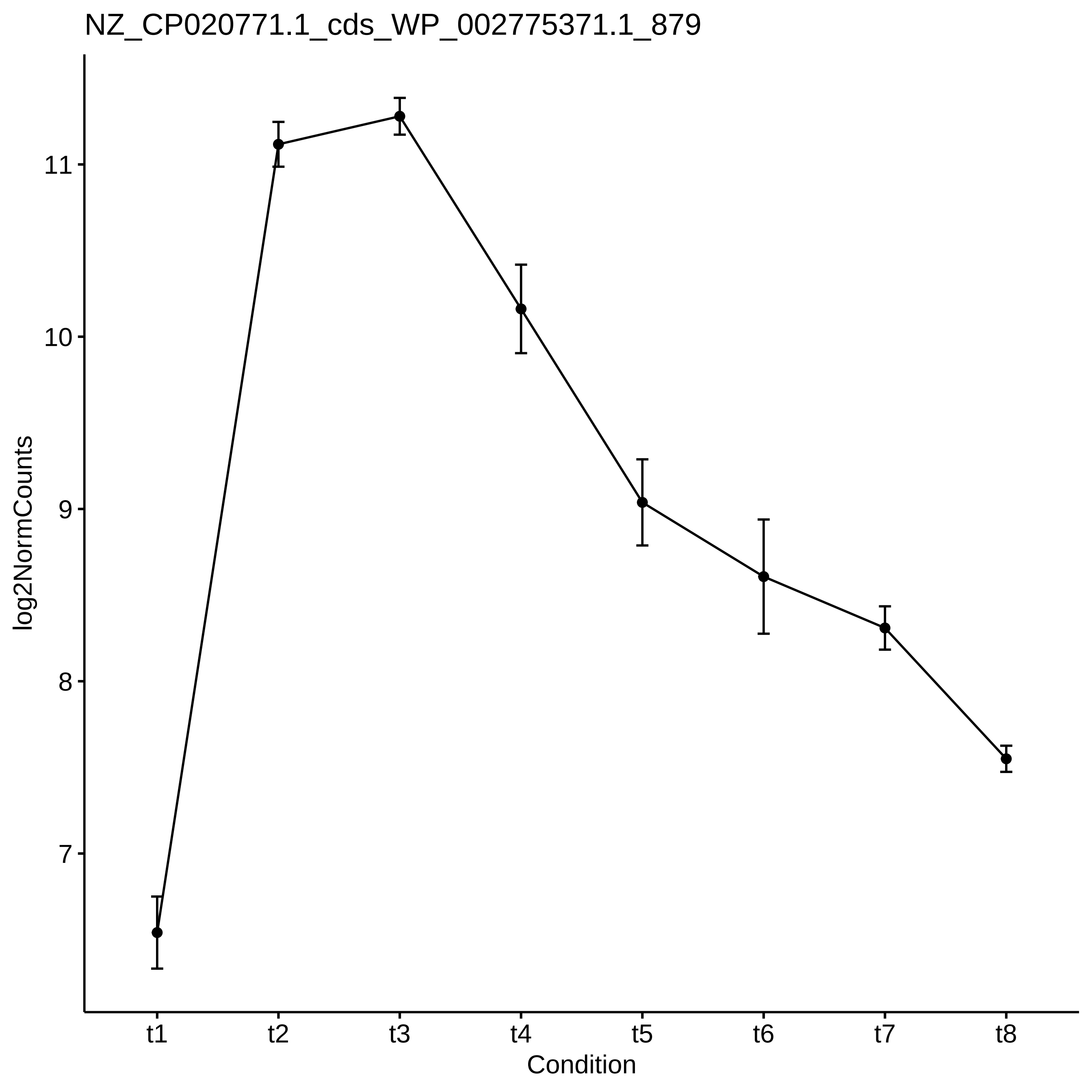
NZ_CP020771.1_cds_WP_002775371.1_879
log2NormCounts
11
10
9
8
7
t1
t2
t3
t4
t5
t6
t7
t8
Condition

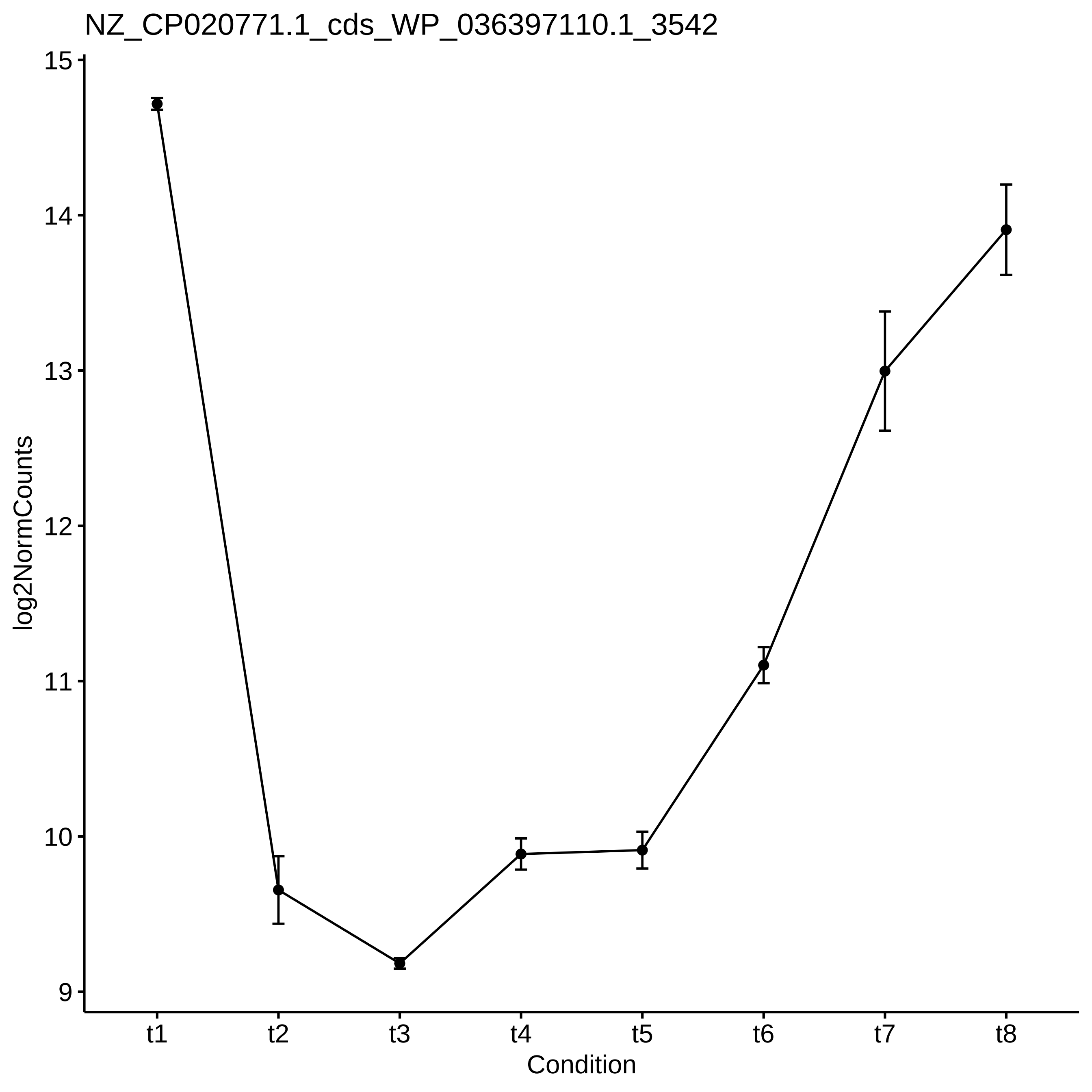

NZ_CP020771.1_cds_WP_036397110.1_3542
log2NormCounts
9
10
11
12
13
14
15
t1
t2
t3
t4
t5
t6
t7
t8
Condition

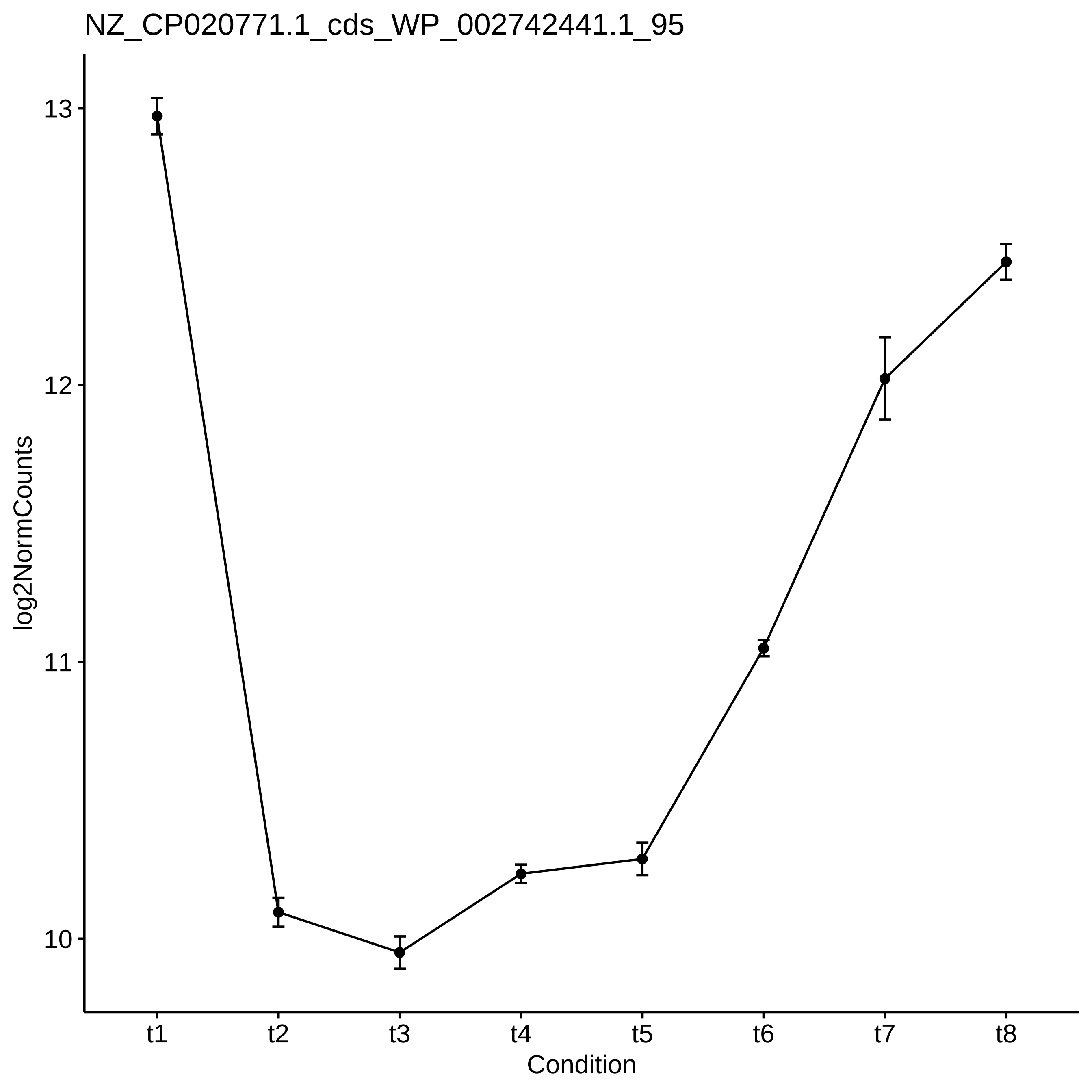
NZ_CP020771.1_cds_WP_002742441.1_95
log2NormCounts
13
12
11
10
t1
t2
t3
t4
t5
t6
t7
t8
Condition

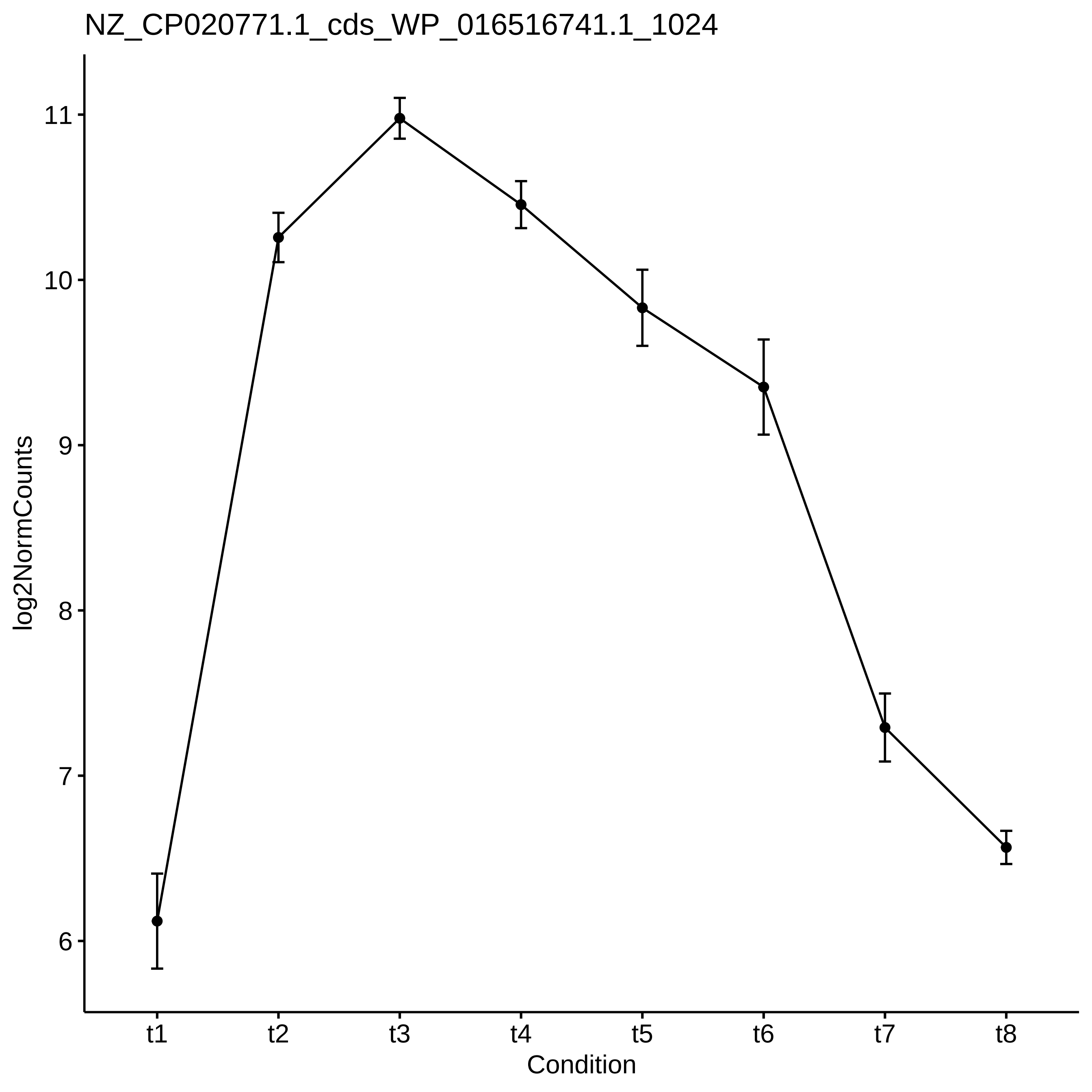

NZ_CP020771.1_cds_WP_016516741.1_1024
log2NormCounts
11
10
9
8
7
6
t1
t2
t3
t4
t5
t6
t7
t8
Condition

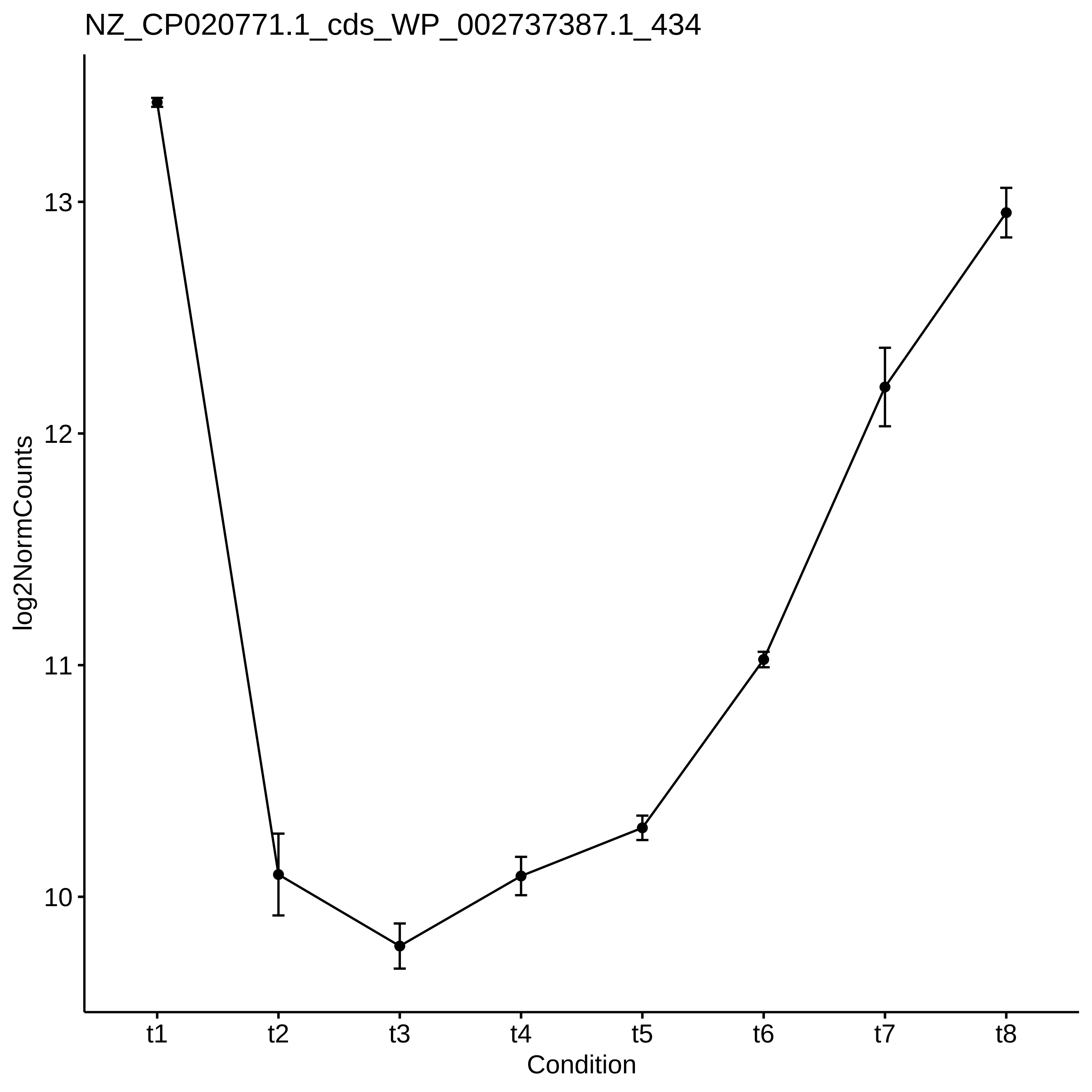
NZ_CP020771.1_cds_WP_002737387.1_434
log2NormCounts
13
12
11
10
t1
t2
t3
t4
t5
t6
t7
t8
Condition

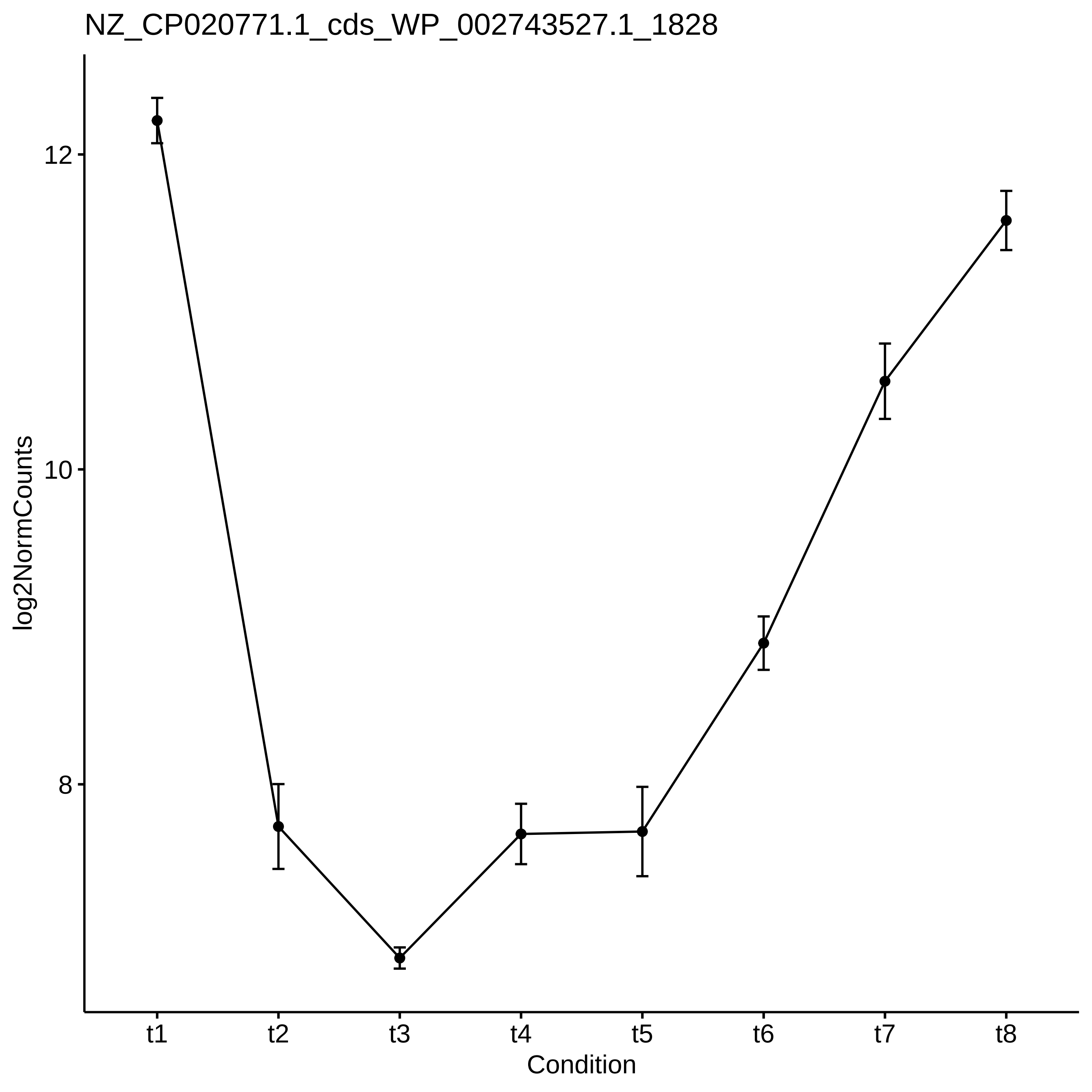
NZ_CP020771.1_cds_WP_002743527.1_1828
log2NormCounts
12
10
8
t1
t2
t3
t4
t5
t6
t7
t8
Condition

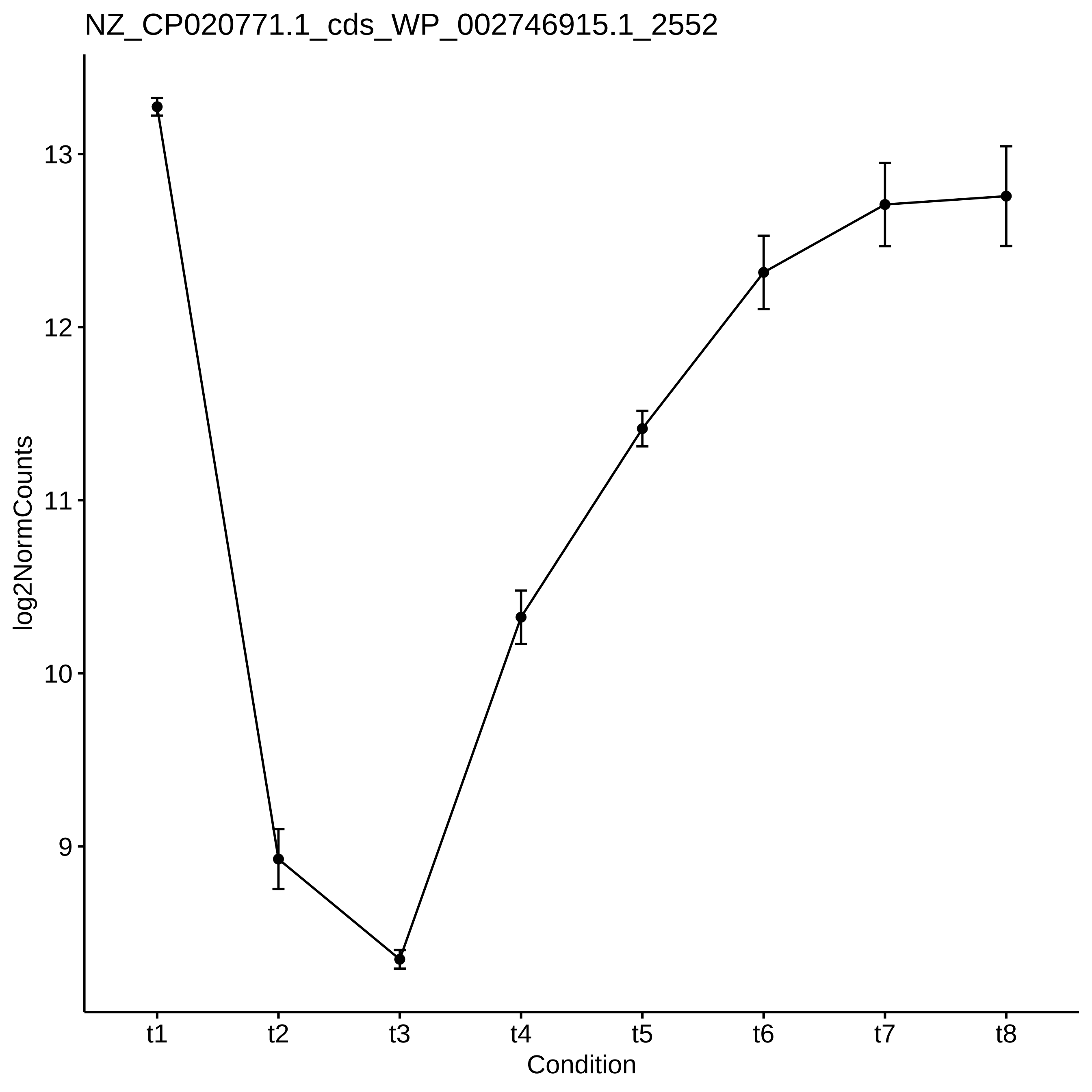
NZ_CP020771.1_cds_WP_002746915.1_2552
log2NormCounts
13
12
11
10
9
t1
t2
t3
t4
t5
t6
t7
t8
Condition

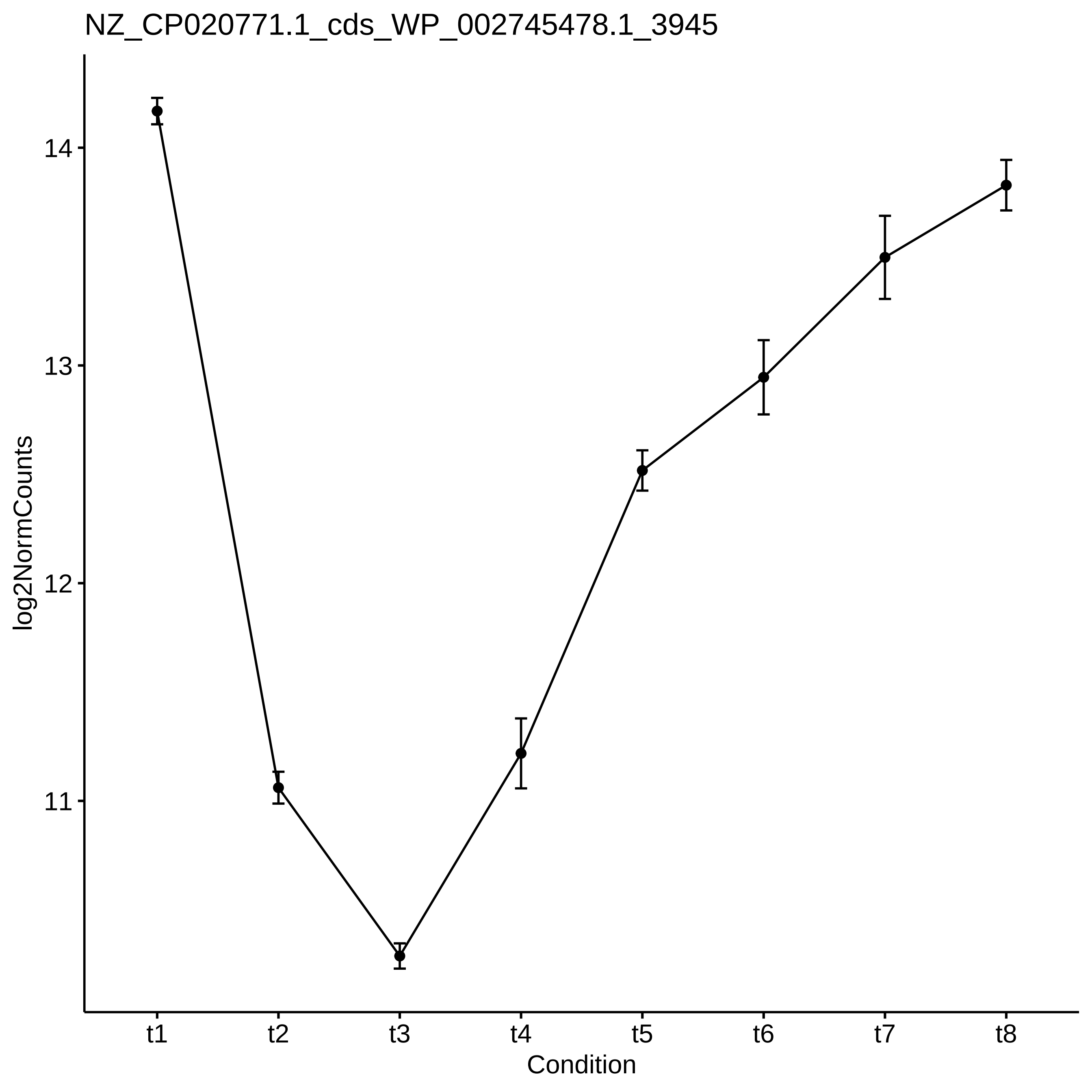

NZ_CP020771.1_cds_WP_002745478.1_3945
log2NormCounts
14
13
12
11
t1
t2
t3
t4
t5
t6
t7
t8
Condition

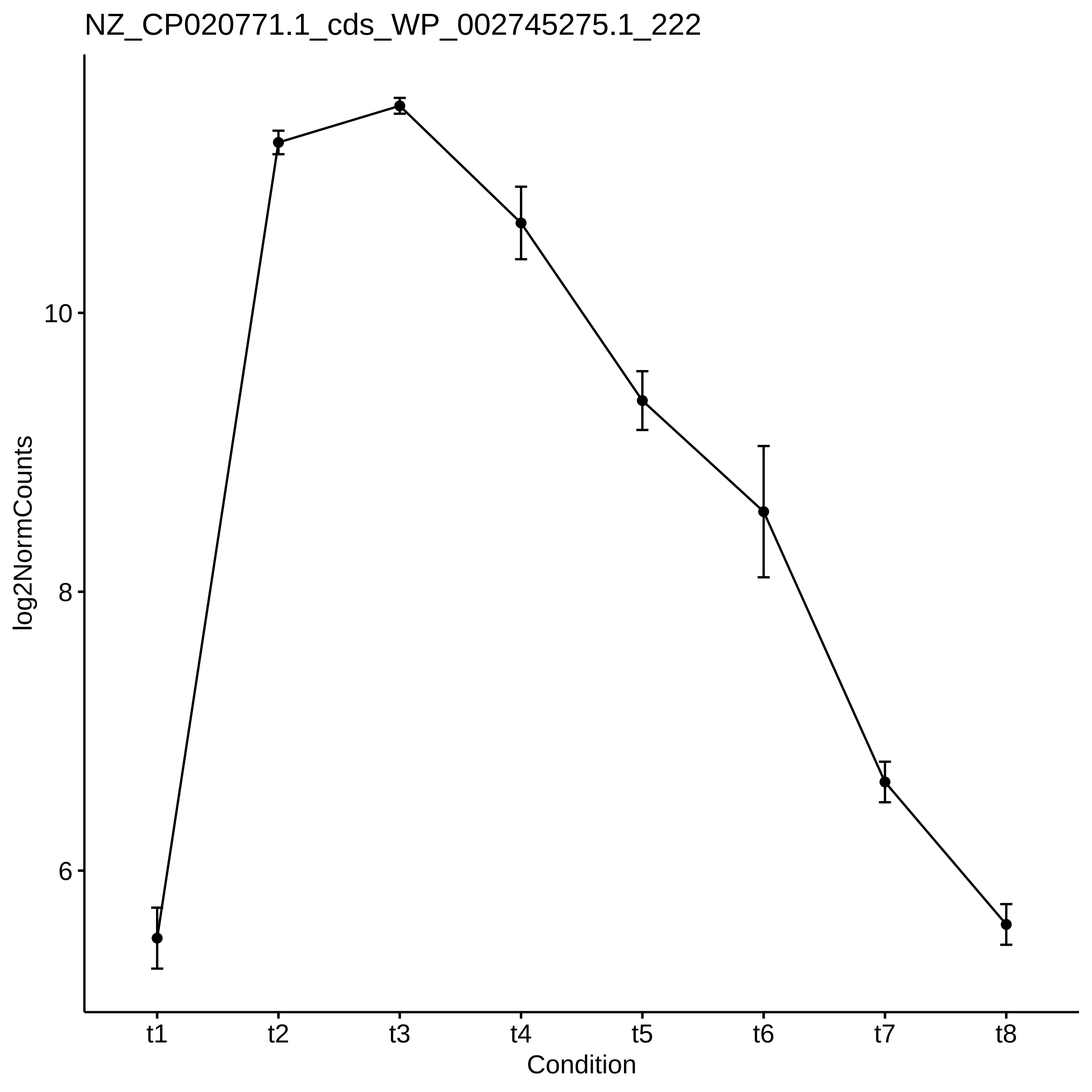
NZ_CP020771.1_cds_WP_002745275.1_222
log2NormCounts
10
8
6
t1
t2
t3
t4
t5
t6
t7
t8
Condition

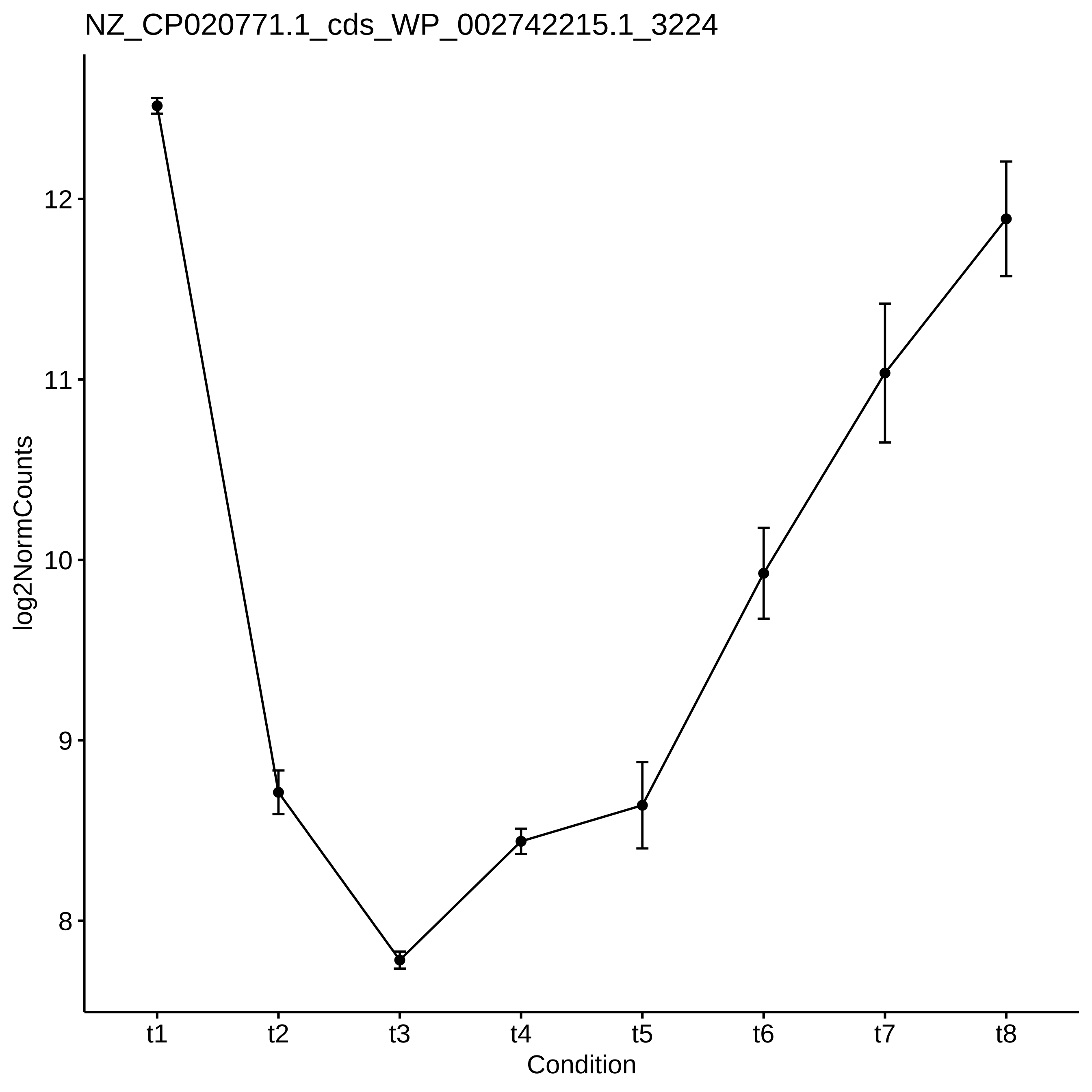

NZ_CP020771.1_cds_WP_002742215.1_3224
log2NormCounts
12
11
10
9
8
t1
t2
t3
t4
t5
t6
t7
t8
Condition

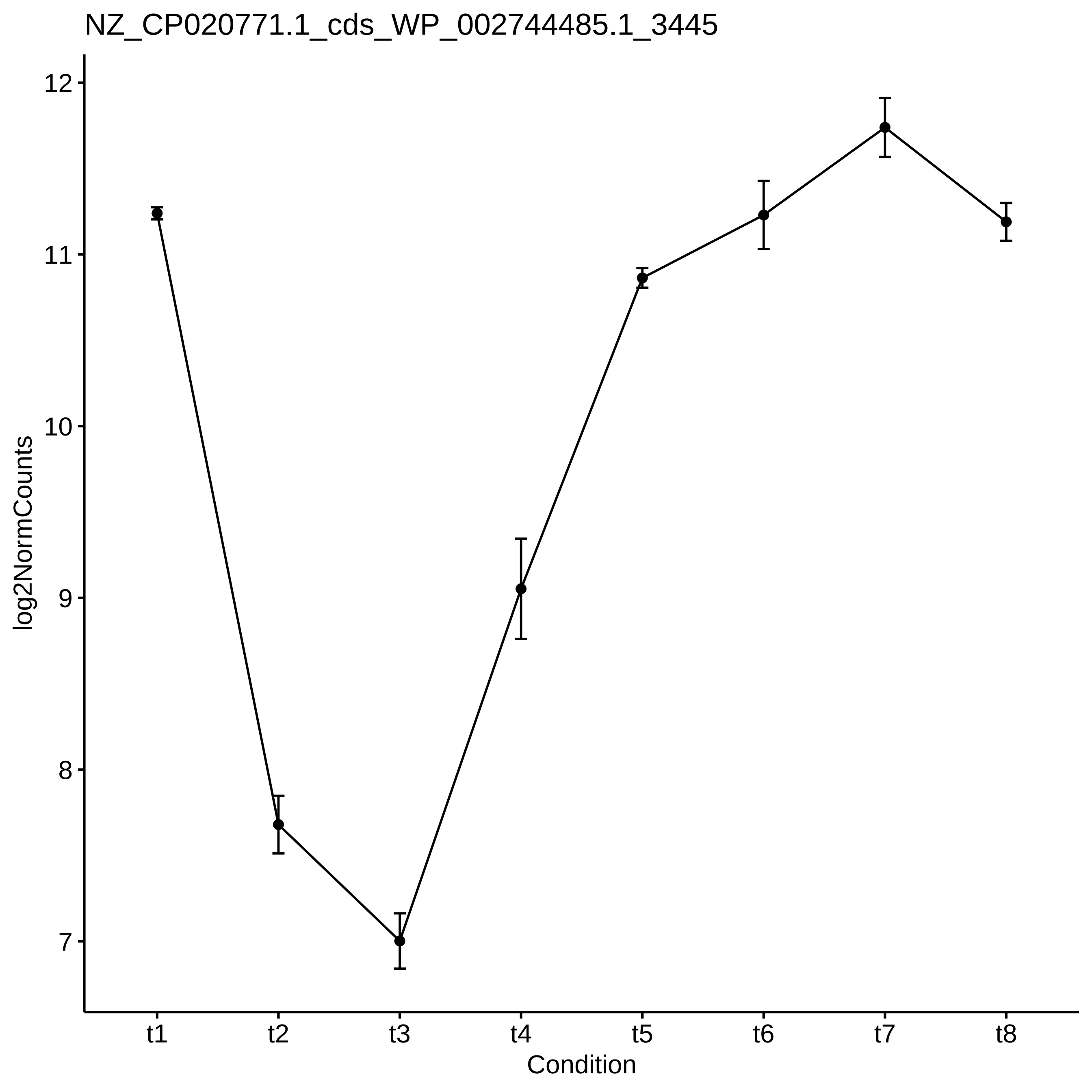
NZ_CP020771.1_cds_WP_002744485.1_3445
log2NormCounts
12
11
10
9
8
7
t1
t2
t3
t4
t5
t6
t7
t8
Condition

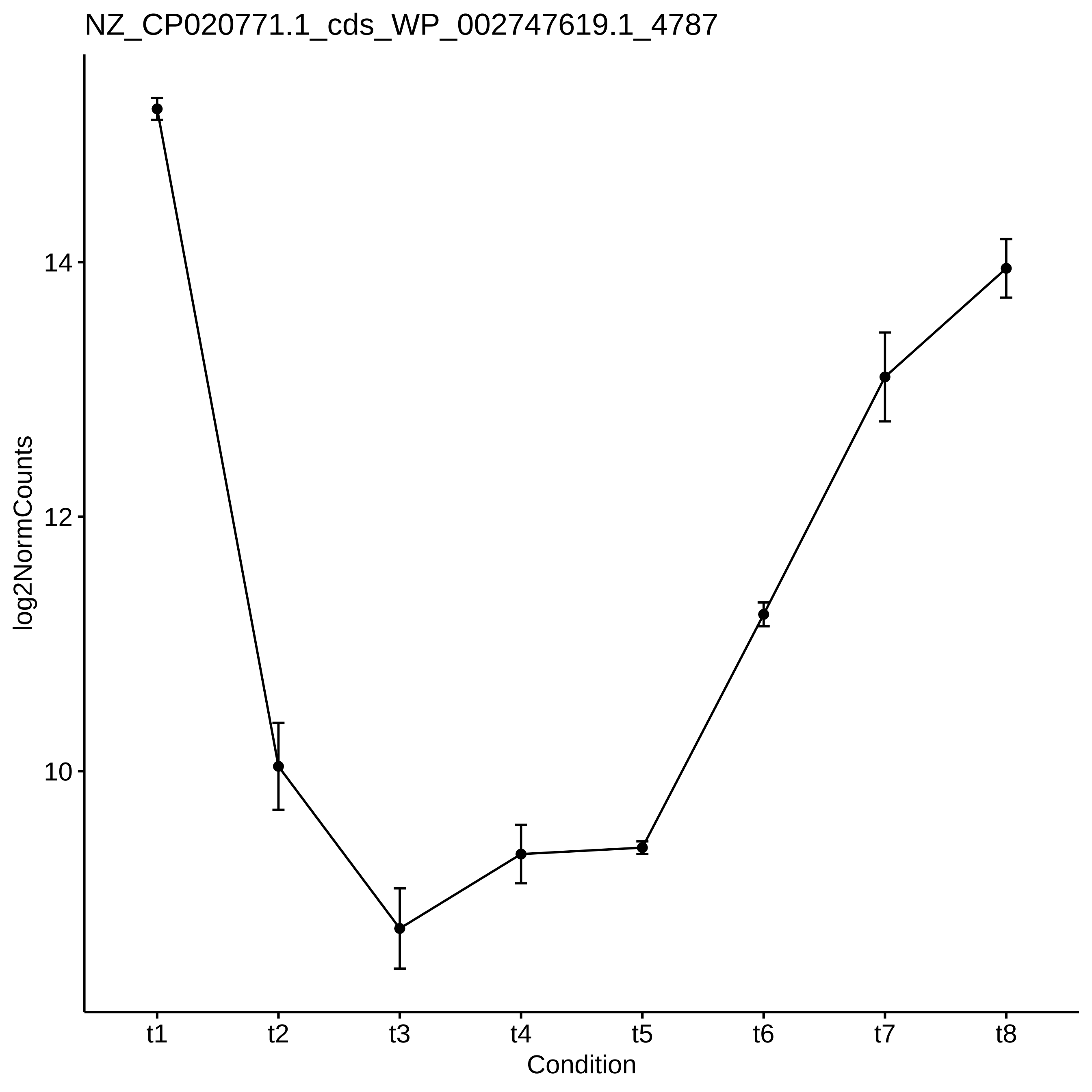
NZ_CP020771.1_cds_WP_002747619.1_4787
log2NormCounts
14
12
10
t1
t2
t3
t4
t5
t6
t7
t8
Condition

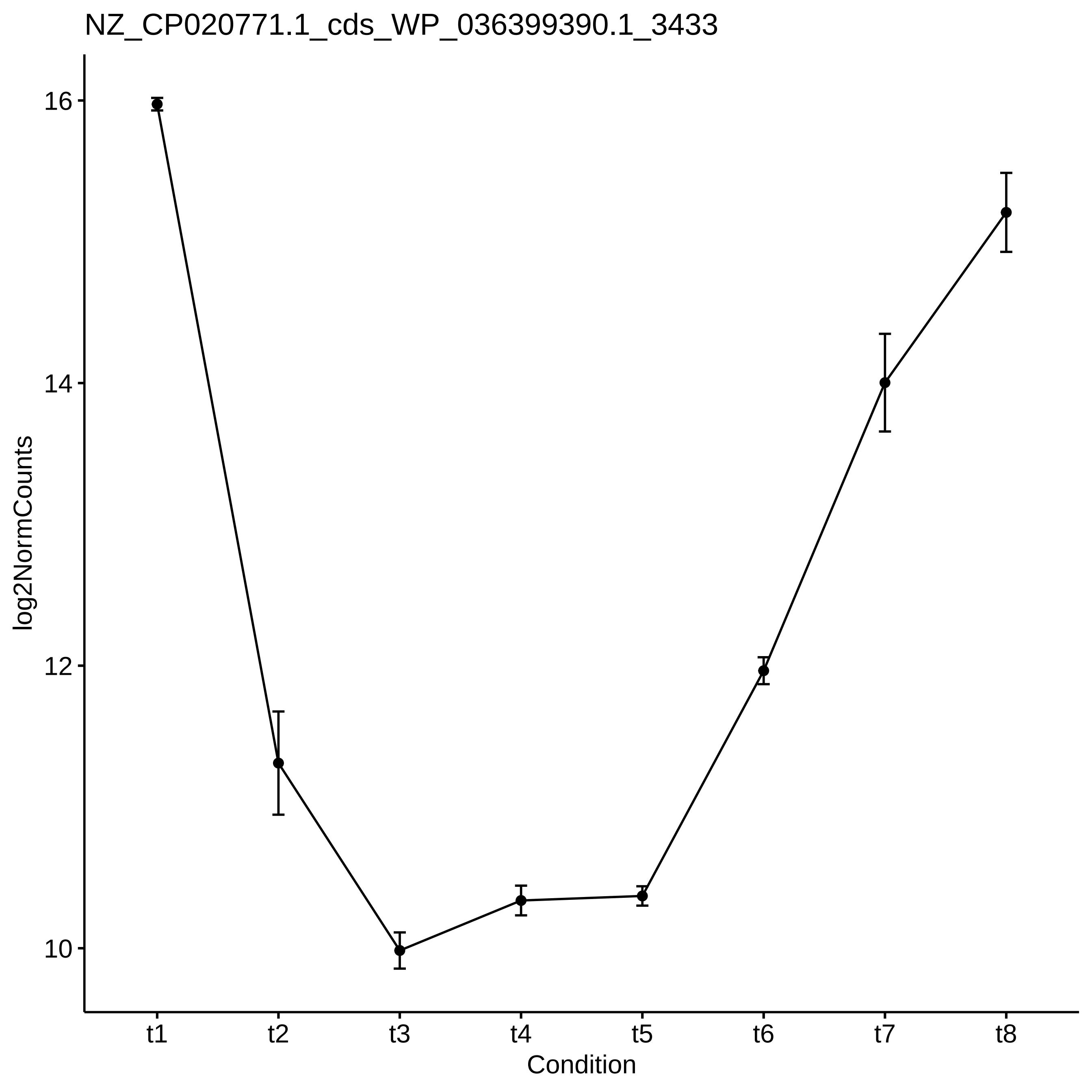
NZ_CP020771.1_cds_WP_036399390.1_3433
log2NormCounts
16
14
12
10
t1
t2
t3
t4
t5
t6
t7
t8
Condition

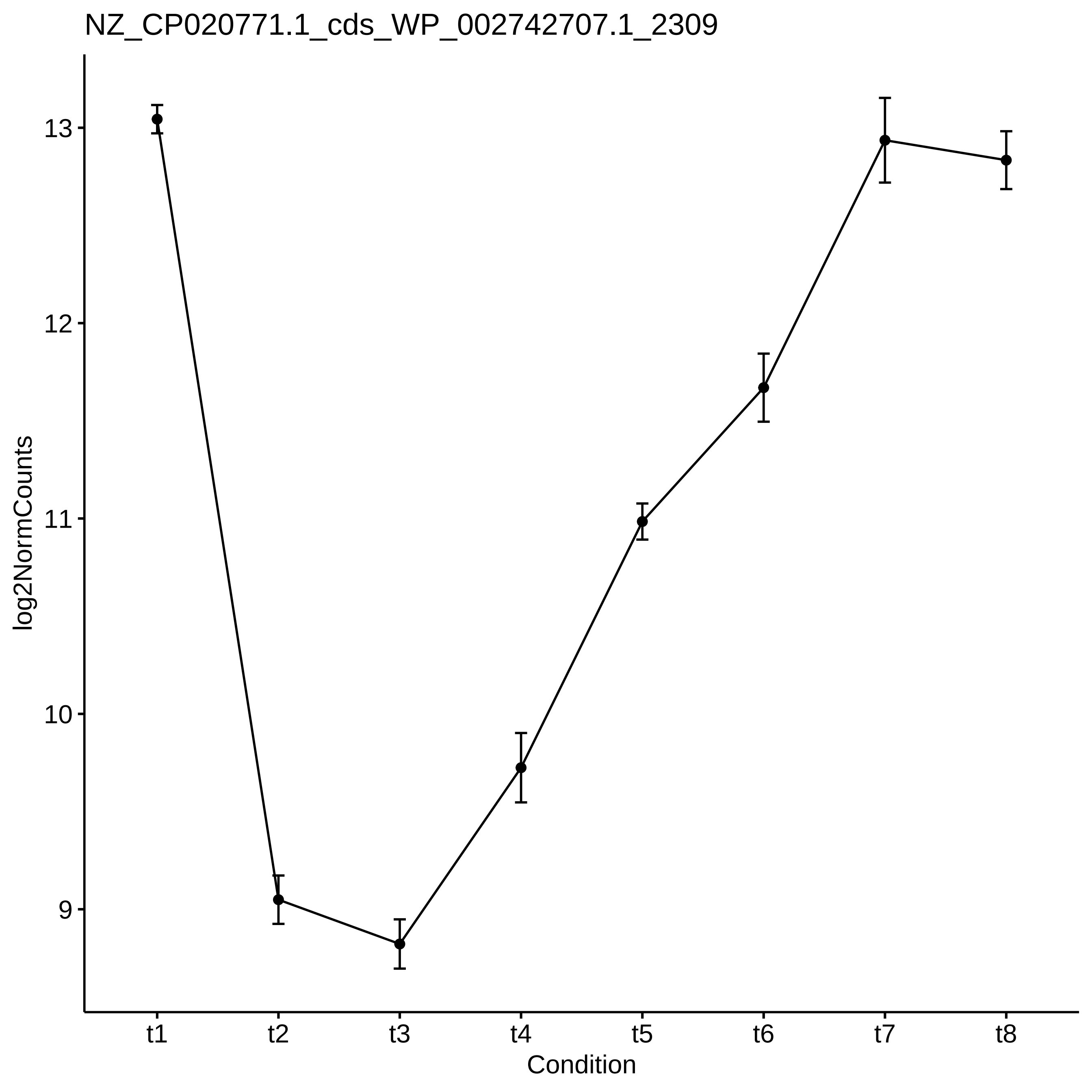
NZ_CP020771.1_cds_WP_002742707.1_2309
log2NormCounts
13
12
11
10
9
t1
t2
t3
t4
t5
t6
t7
t8
Condition

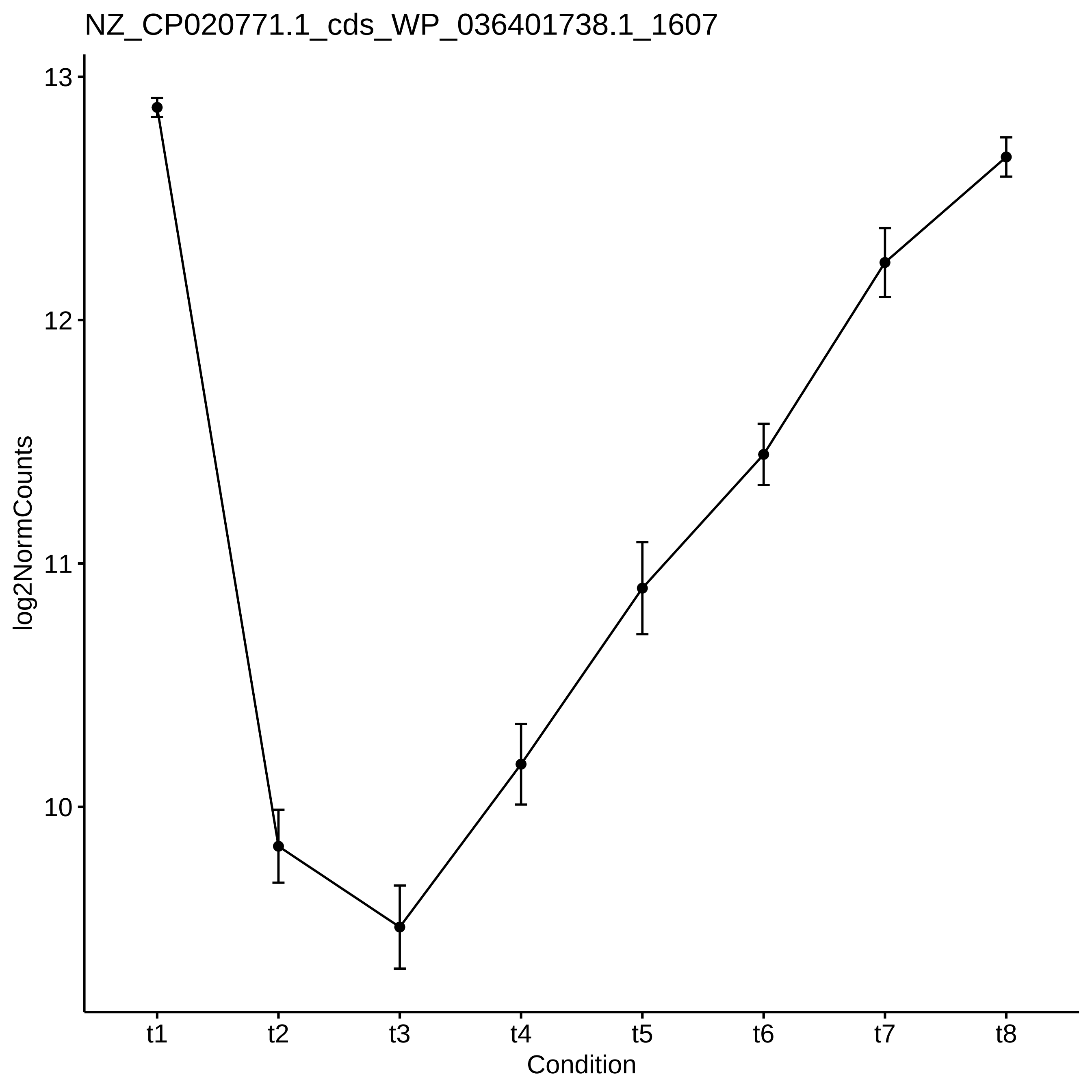
NZ_CP020771.1_cds_WP_036401738.1_1607
13
12
11
10
log2NormCounts
t1
t2
t3
t4
t5
t6
t7
t8
Condition

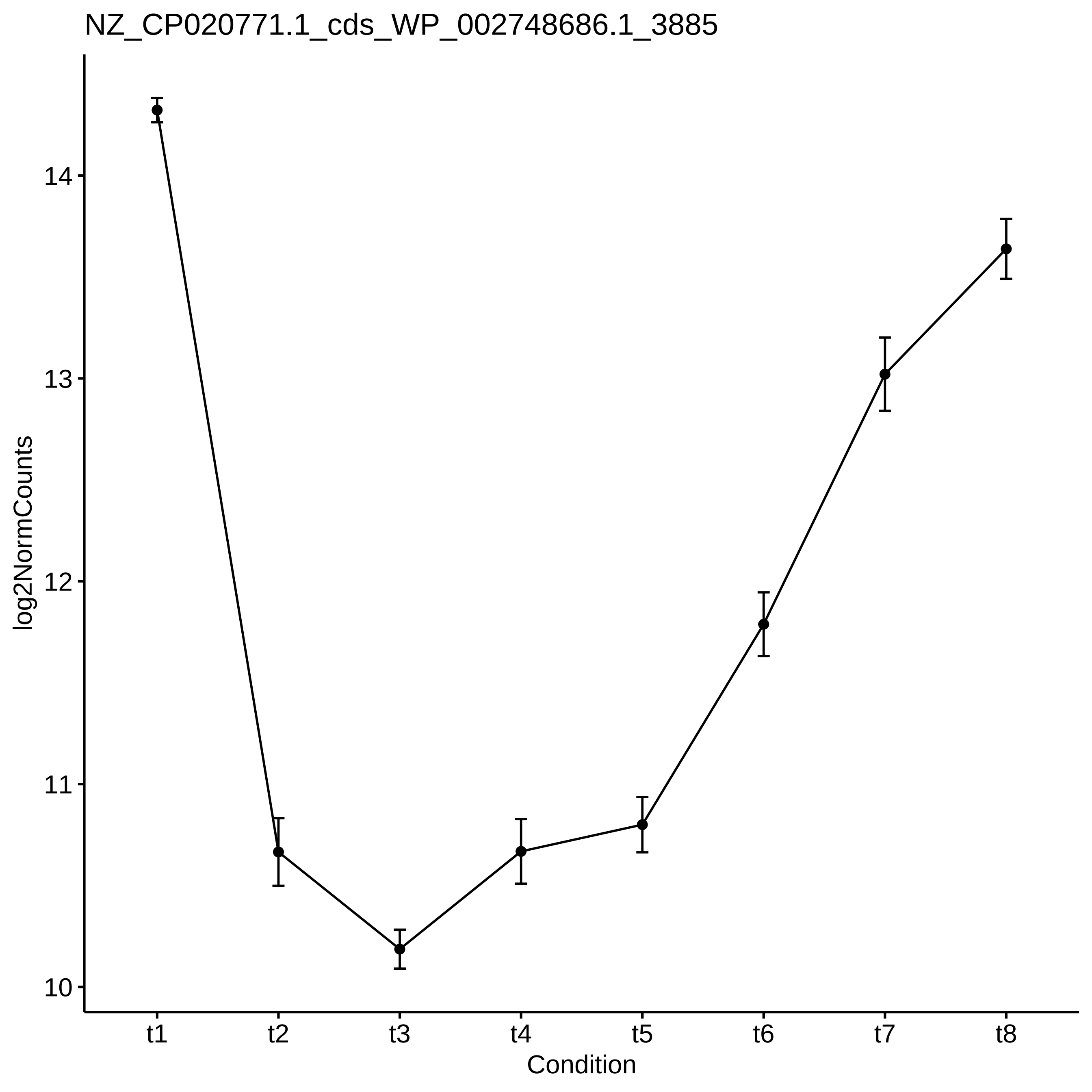
NZ_CP020771.1_cds_WP_002748686.1_3885
log2NormCounts
14
13
12
11
10
t1
t2
t3
t4
t5
t6
t7
t8
Condition

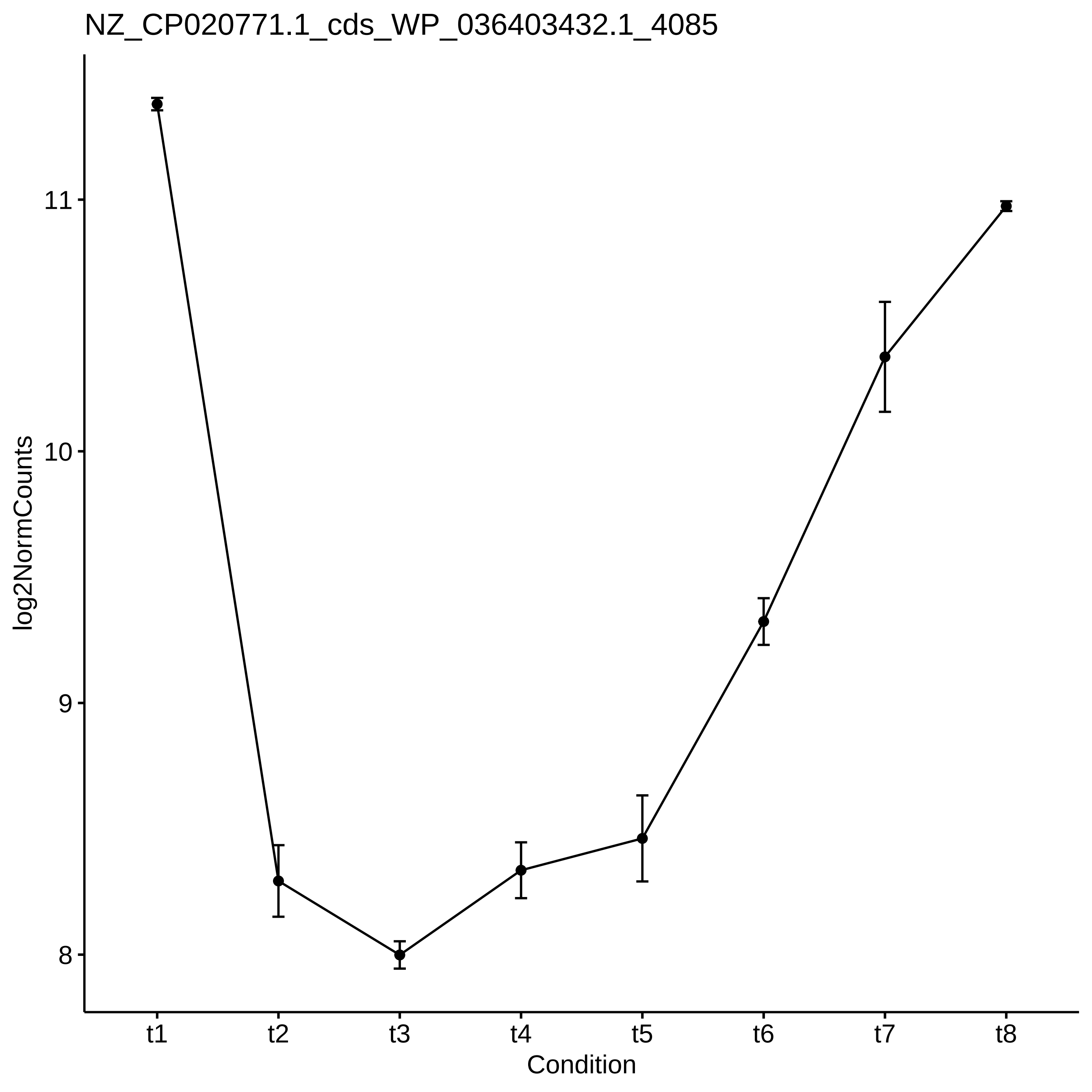
NZ_CP020771.1_cds_WP_036403432.1_4085
log2NormCounts
11
10
9
8
t1
t2
t3
t4
t5
t6
t7
t8
Condition

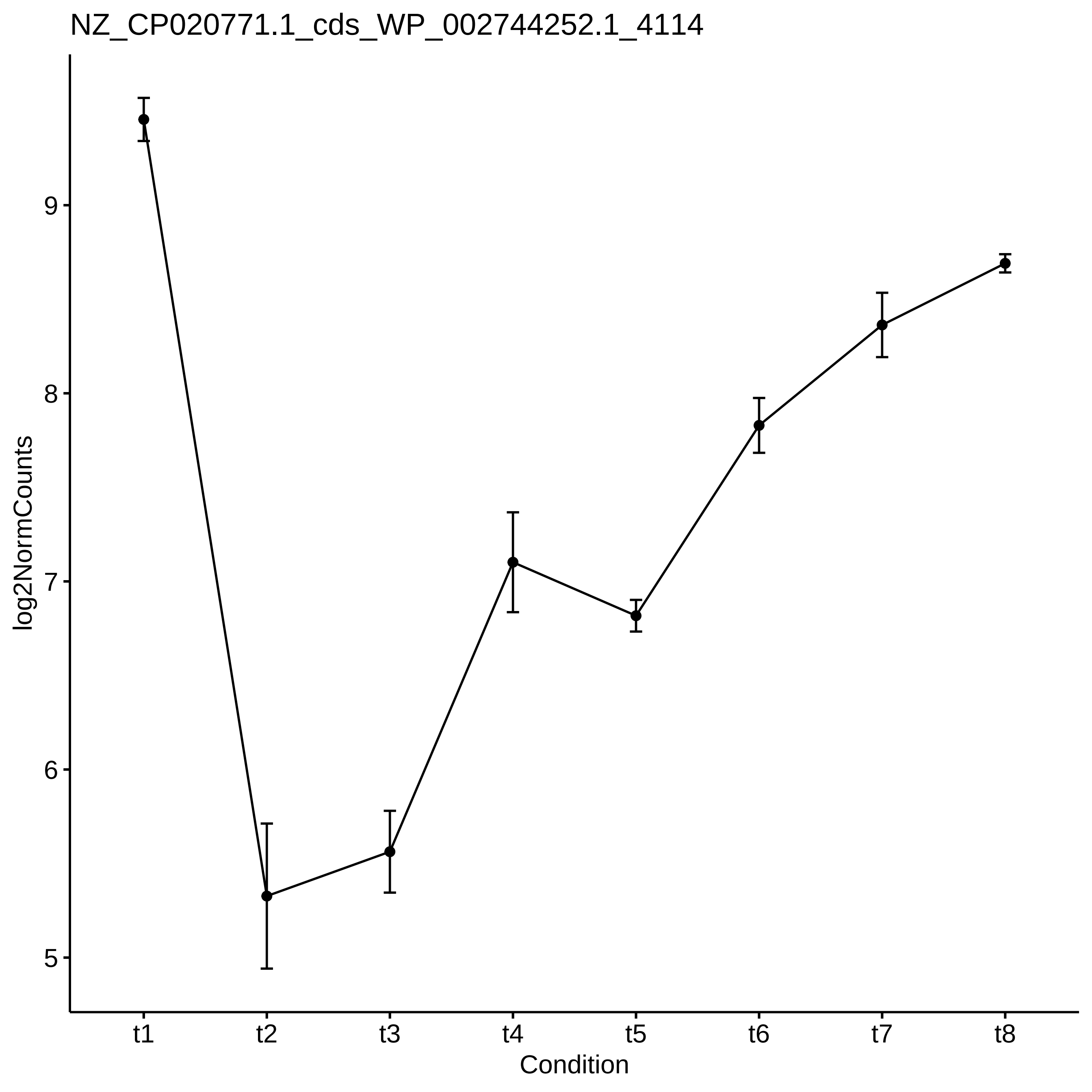
NZ_CP020771.1_cds_WP_002744252.1_4114
log2NormCounts
9
8
7
6
5
t1
t2
t3
t4
t5
t6
t7
t8
Condition

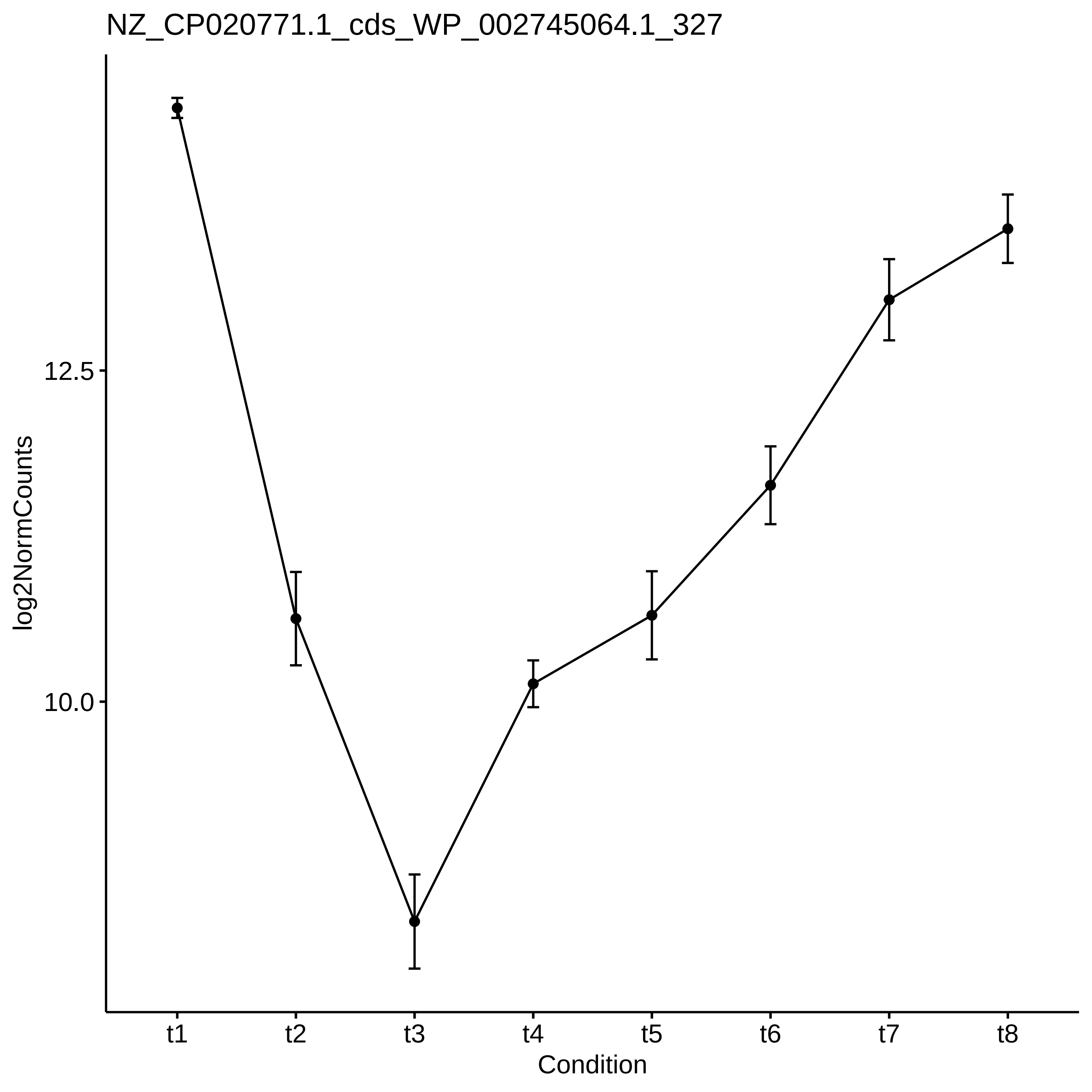
NZ_CP020771.1_cds_WP_002745064.1_327
log2NormCounts
12.5
10.0
t1
t2
t3
t4
t5
t6
t7
t8
Condition

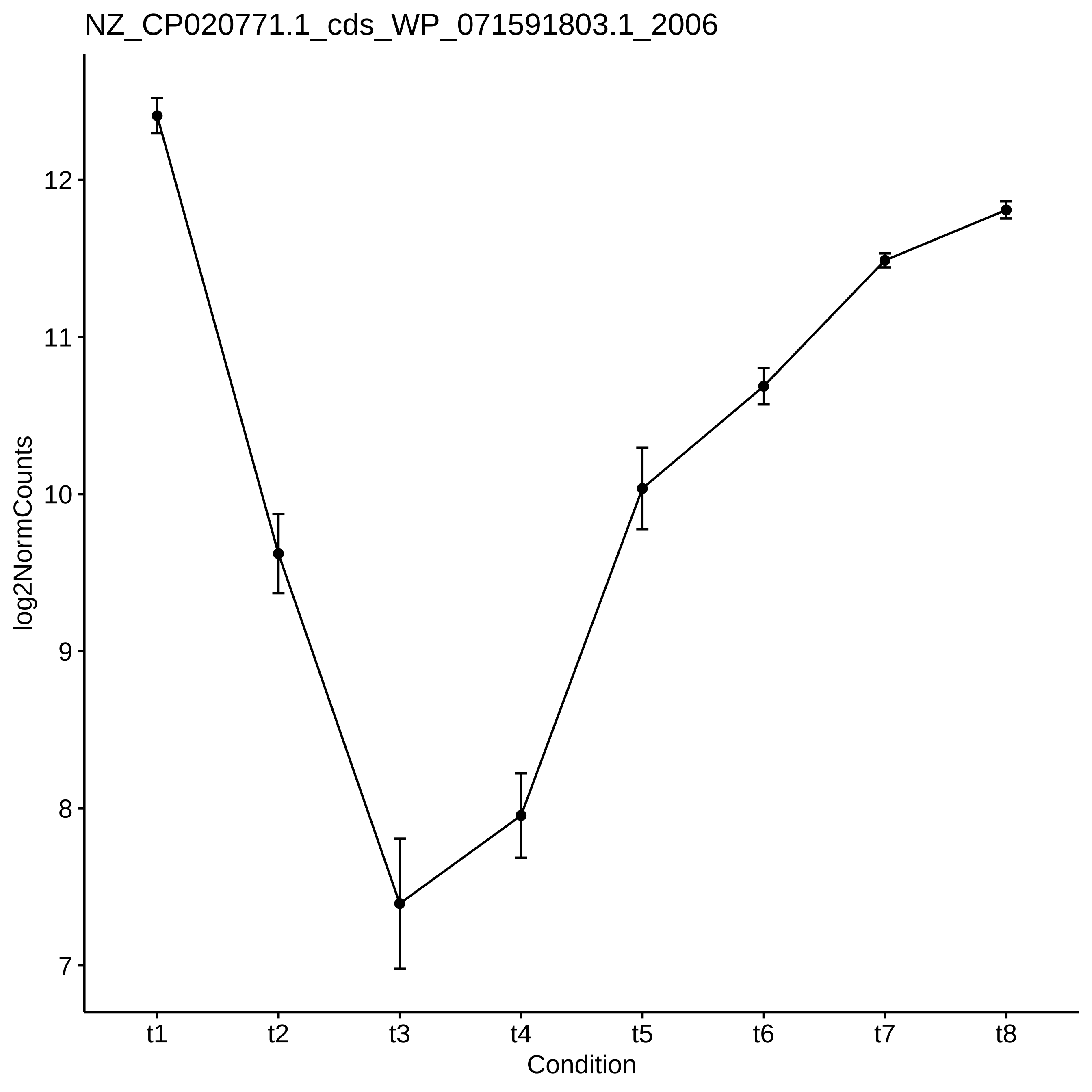
NZ_CP020771.1_cds_WP_071591803.1_2006
log2NormCounts
12
11
10
9
8
7
t1
t2
t3
t4
t5
t6
t7
t8
Condition

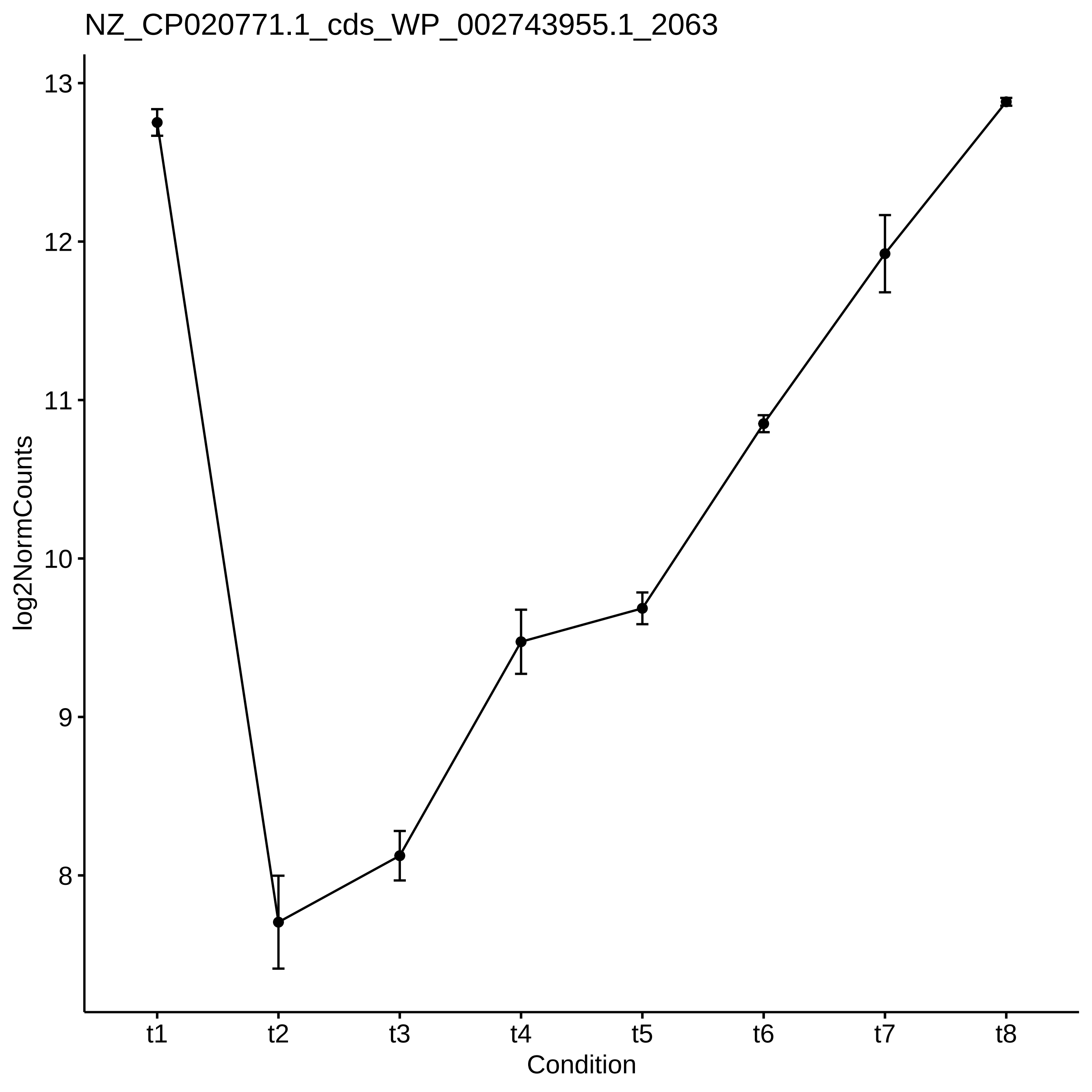
NZ_CP020771.1_cds_WP_002743955.1_2063
log2NormCounts
13
12
11
10
9
8
t1
t2
t3
t4
t5
t6
t7
t8
Condition

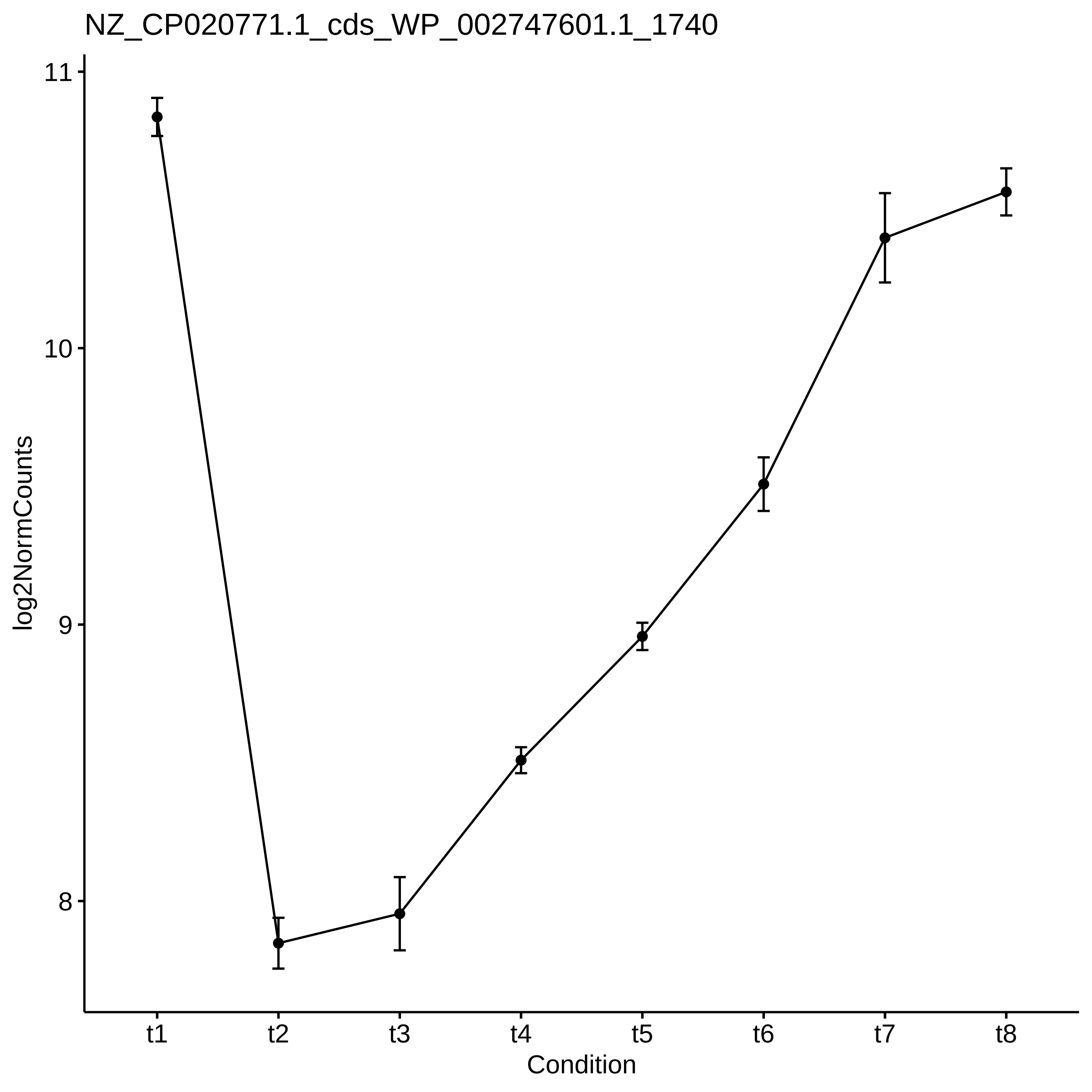
NZ_CP020771.1_cds_WP_002747601.1_1740
log2NormCounts
11
10
9
8
t1
t2
t3
t4
t5
t6
t7
t8
Condition

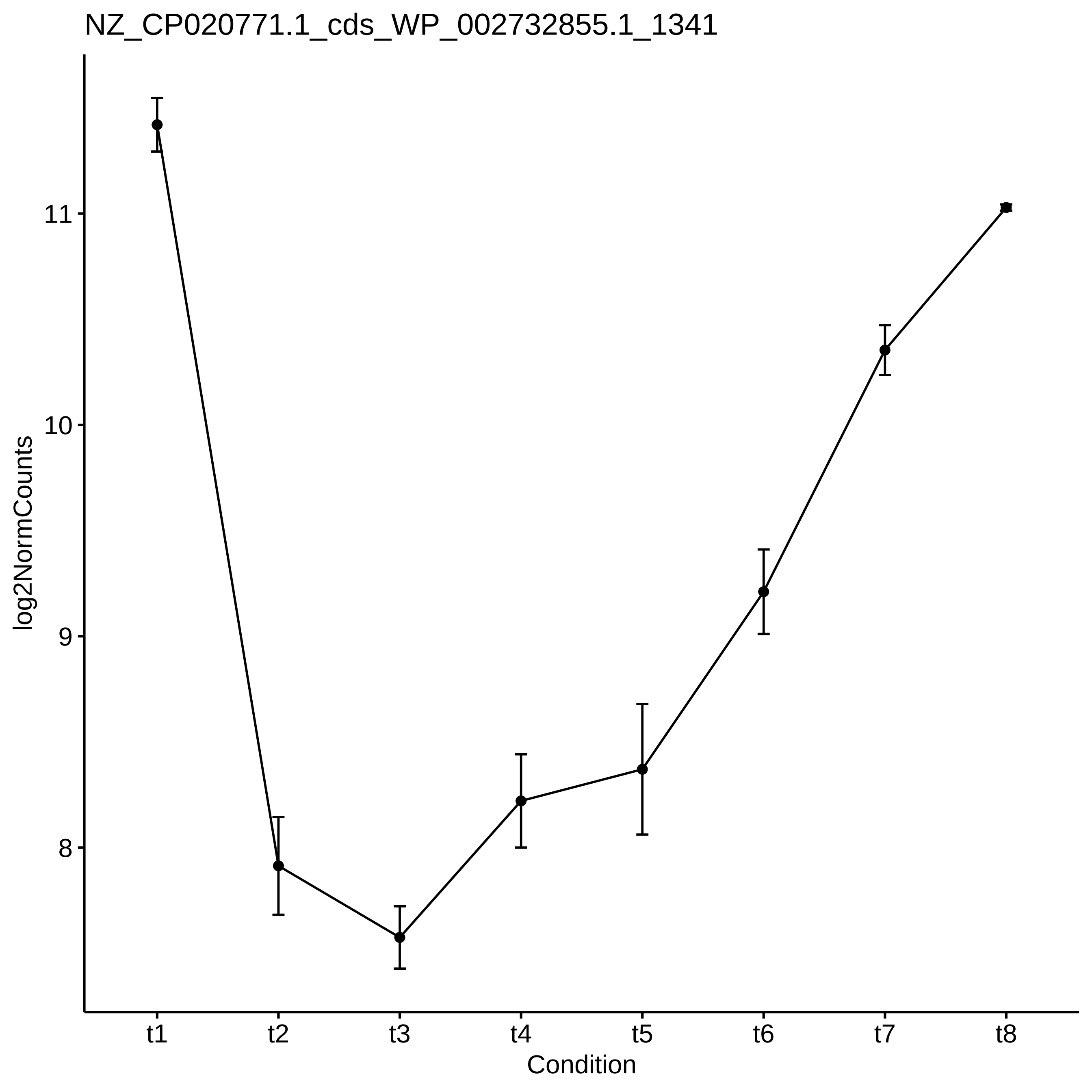

NZ_CP020771.1_cds_WP_002732855.1_1341
log2NormCounts
11
10
9
8
t1
t2
t3
t4
t5
t6
t7
t8
Condition

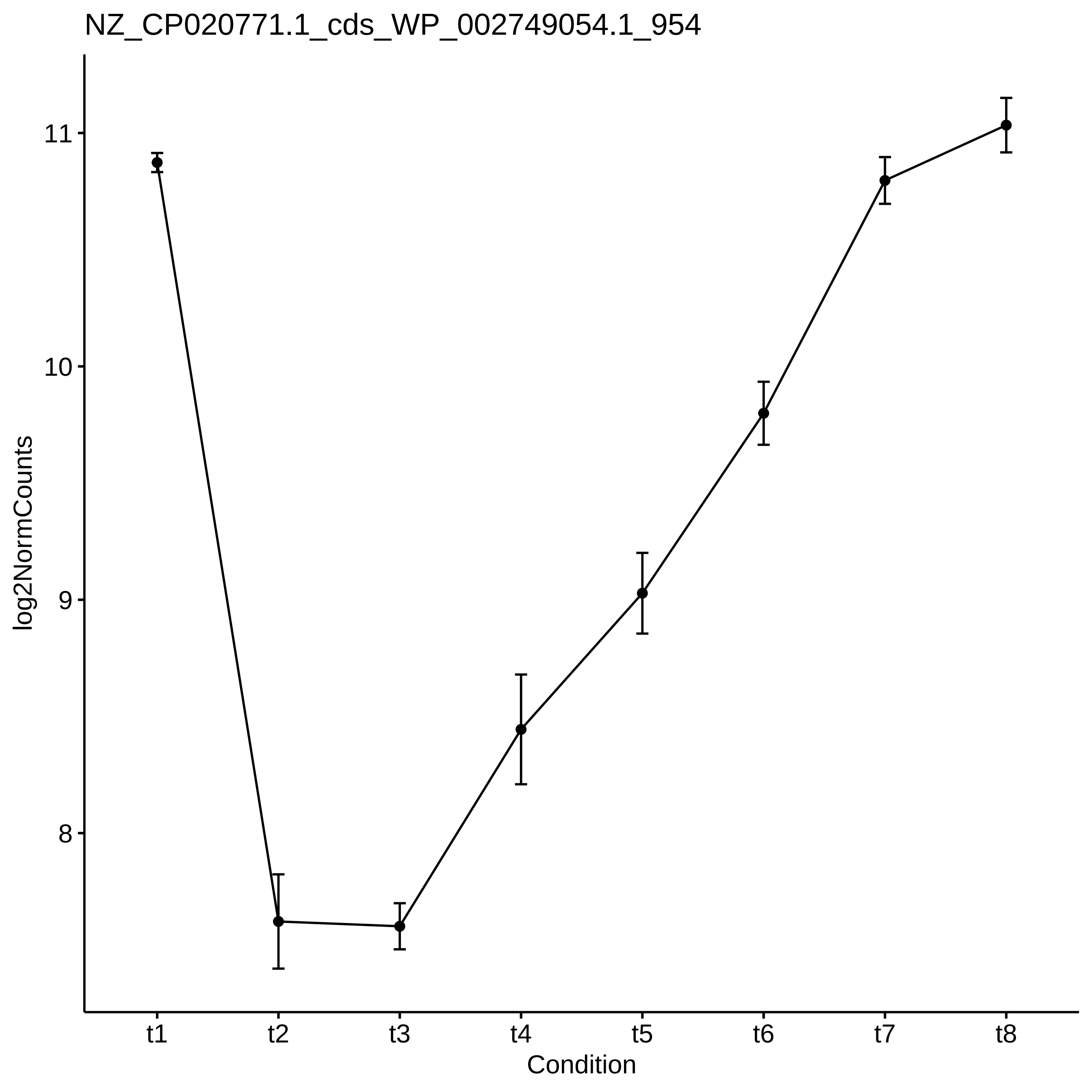
NZ_CP020771.1_cds_WP_002749054.1_954
log2NormCounts
11
10
9
8
t1
t2
t3
t4
t5
t6
t7
t8
Condition

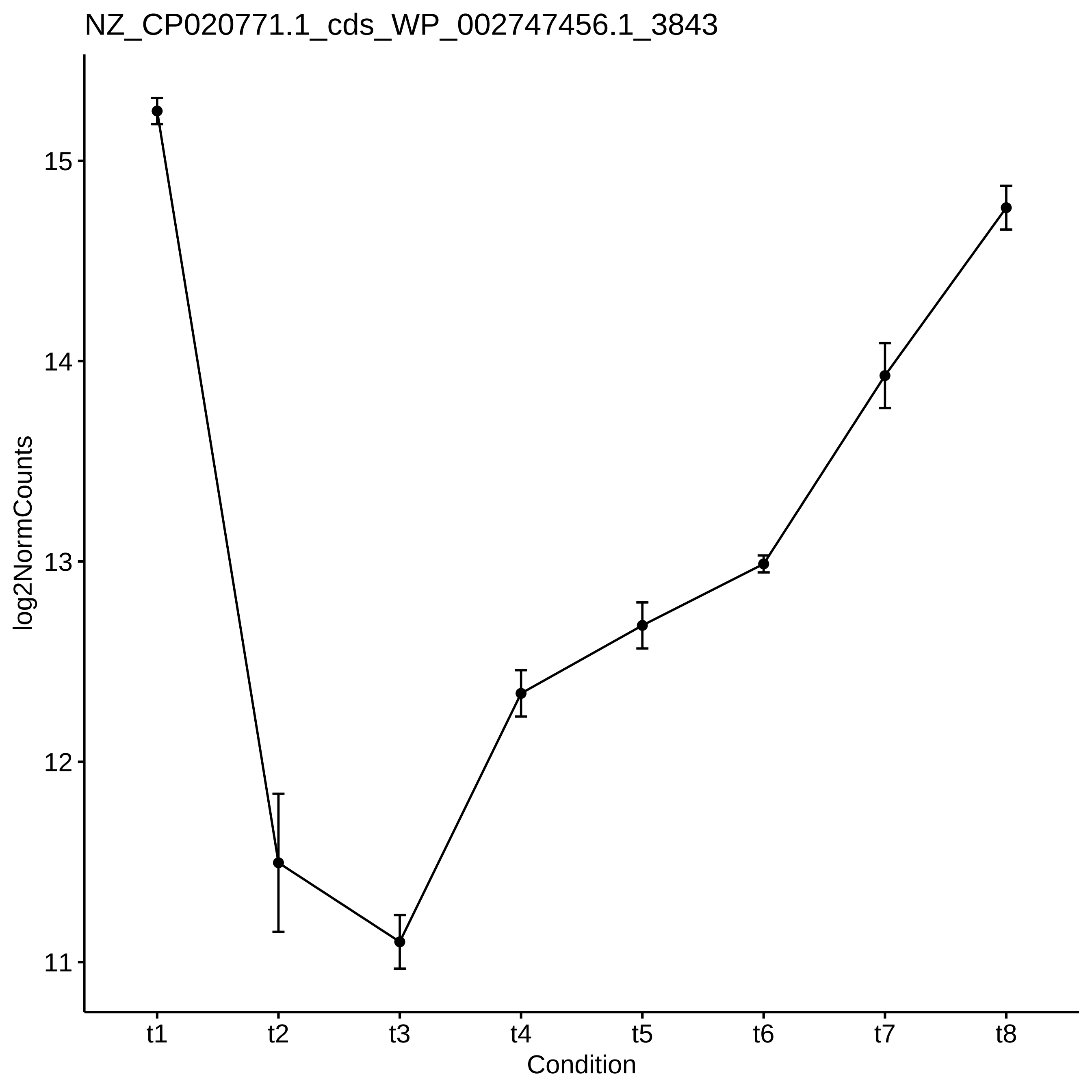

NZ_CP020771.1_cds_WP_002747456.1_3843
log2NormCounts
15
14
13
12
11
t1
t2
t3
t4
t5
t6
t7
t8
Condition

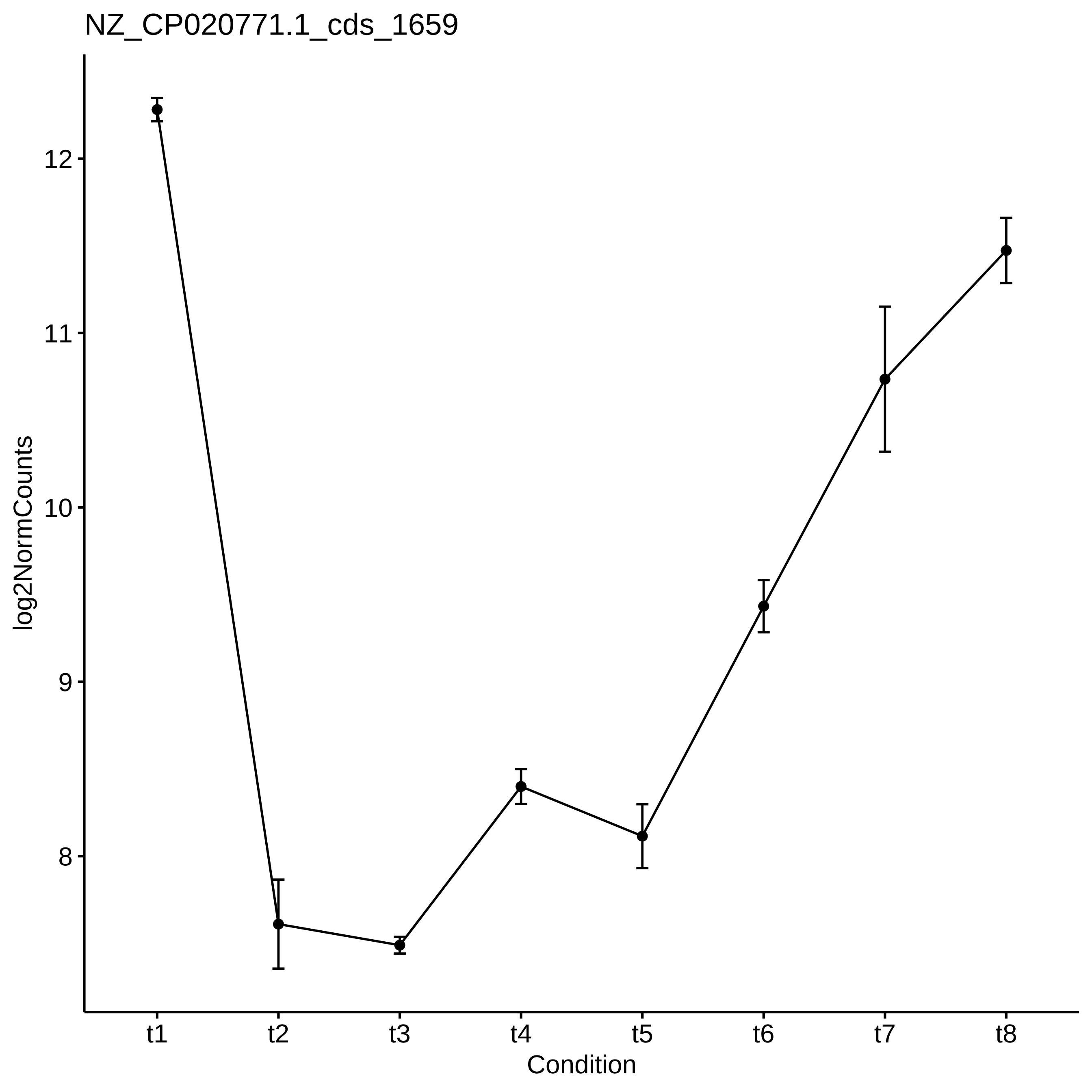
NZ_CP020771.1_cds_1659
log2NormCounts
12
11
10
9
8
t1
t2
t3
t4
t5
t6
t7
t8
Condition

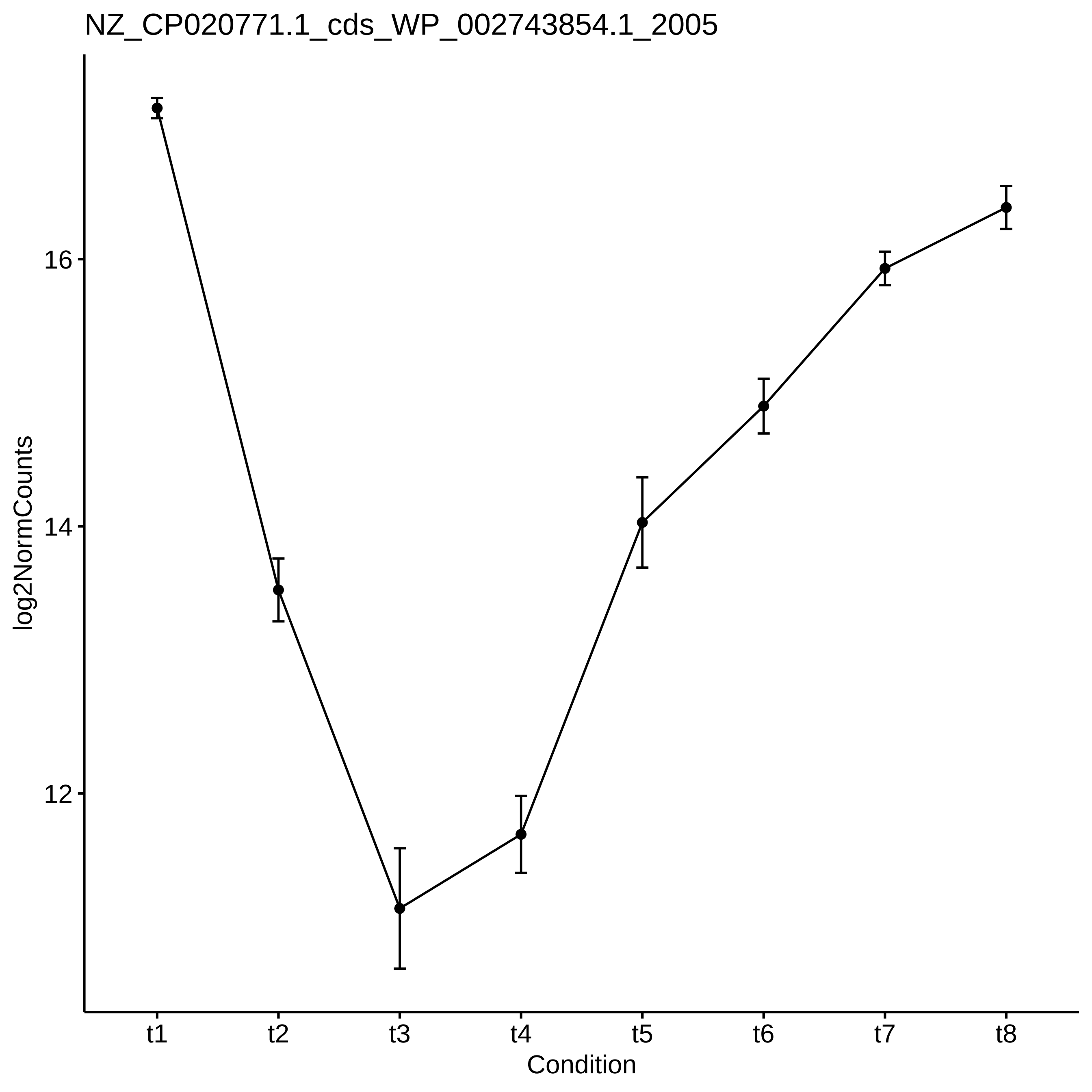

NZ_CP020771.1_cds_WP_002743854.1_2005
log2NormCounts
16
14
12
t1
t2
t3
t4
t5
t6
t7
t8
Condition

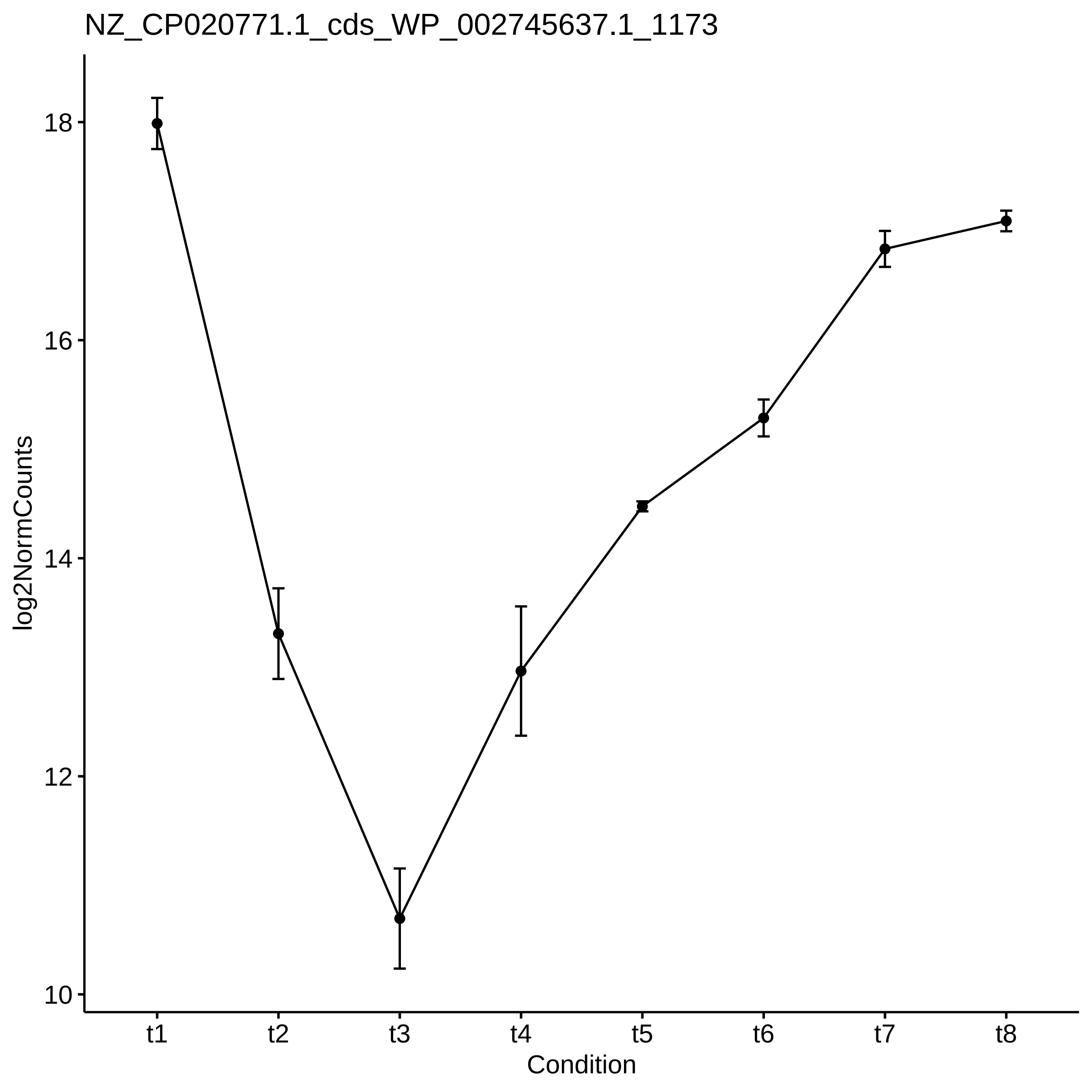
NZ_CP020771.1_cds_WP_002745637.1_1173
log2NormCounts
18
16
14
12
10
t1
t2
t3
t4
t5
t6
t7
t8
Condition

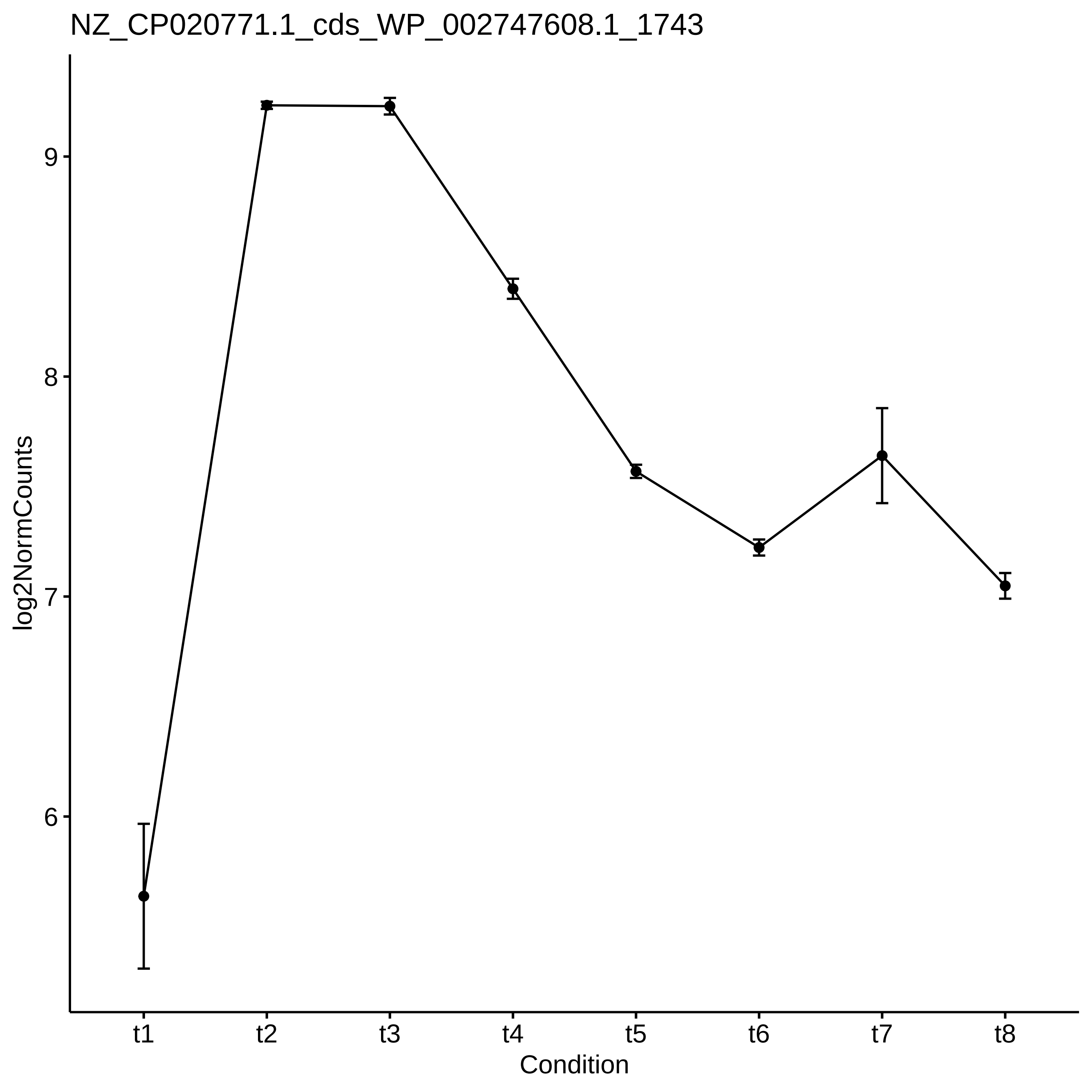
NZ_CP020771.1_cds_WP_002747608.1_1743
log2NormCounts
9
8
7
6
t1
t2
t3
t4
t5
t6
t7
t8
Condition

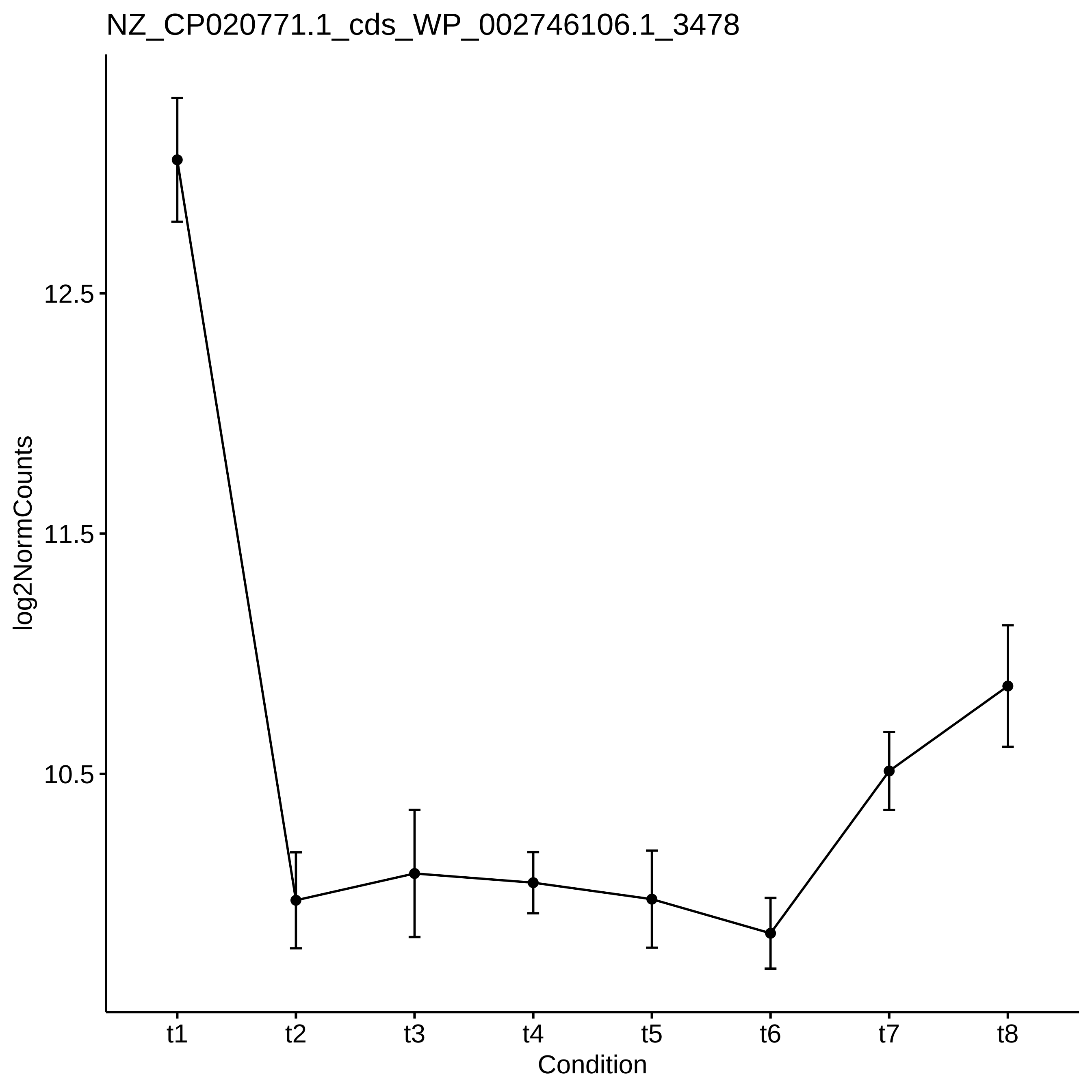
NZ_CP020771.1_cds_WP_002746106.1_3478
log2NormCounts
12.5
11.5
10.5
t1
t2
t3
t4
t5
t6
t7
t8
Condition

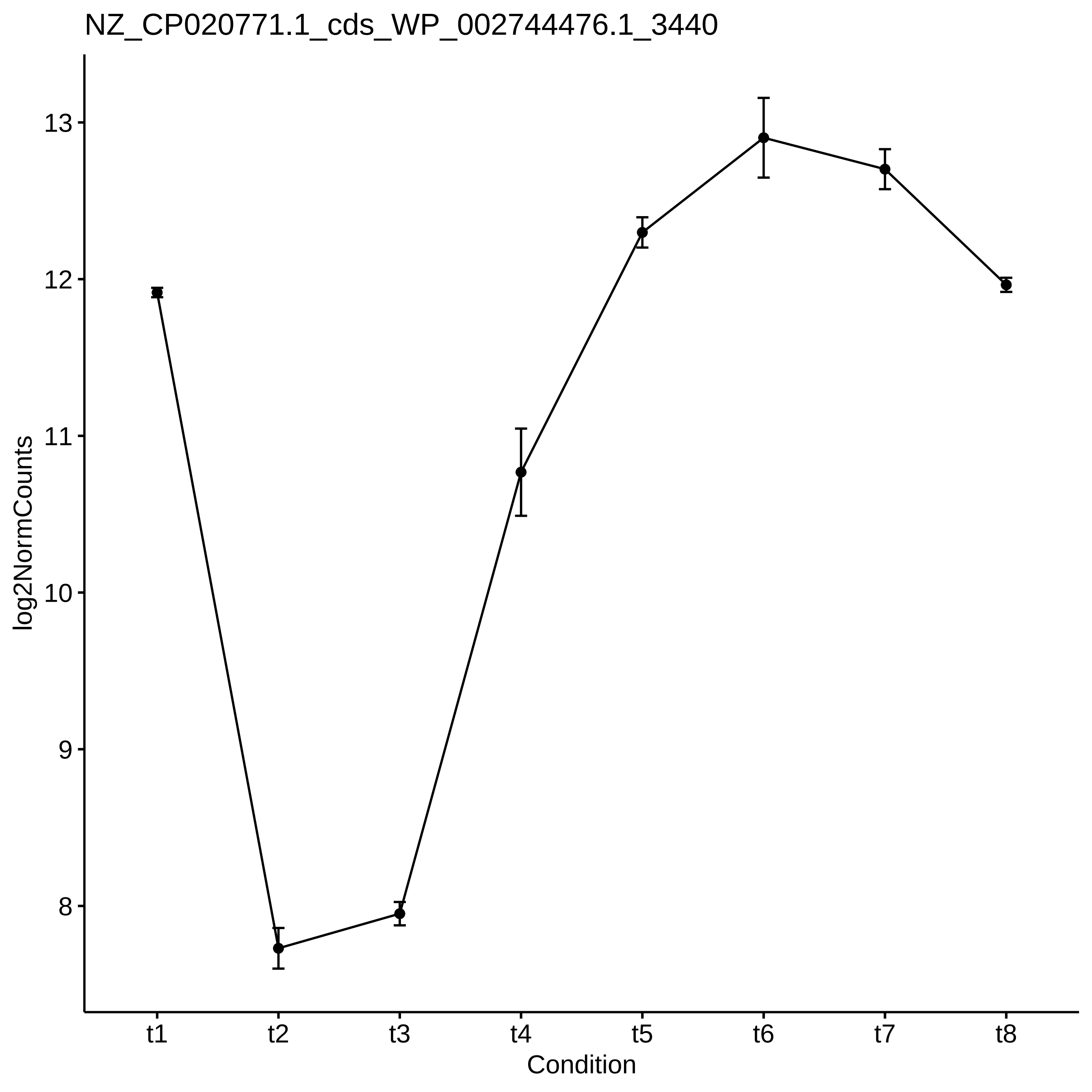
NZ_CP020771.1_cds_WP_002744476.1_3440
log2NormCounts
13
12
11
10
9
8
t1
t2
t3
t4
t5
t6
t7
t8
Condition

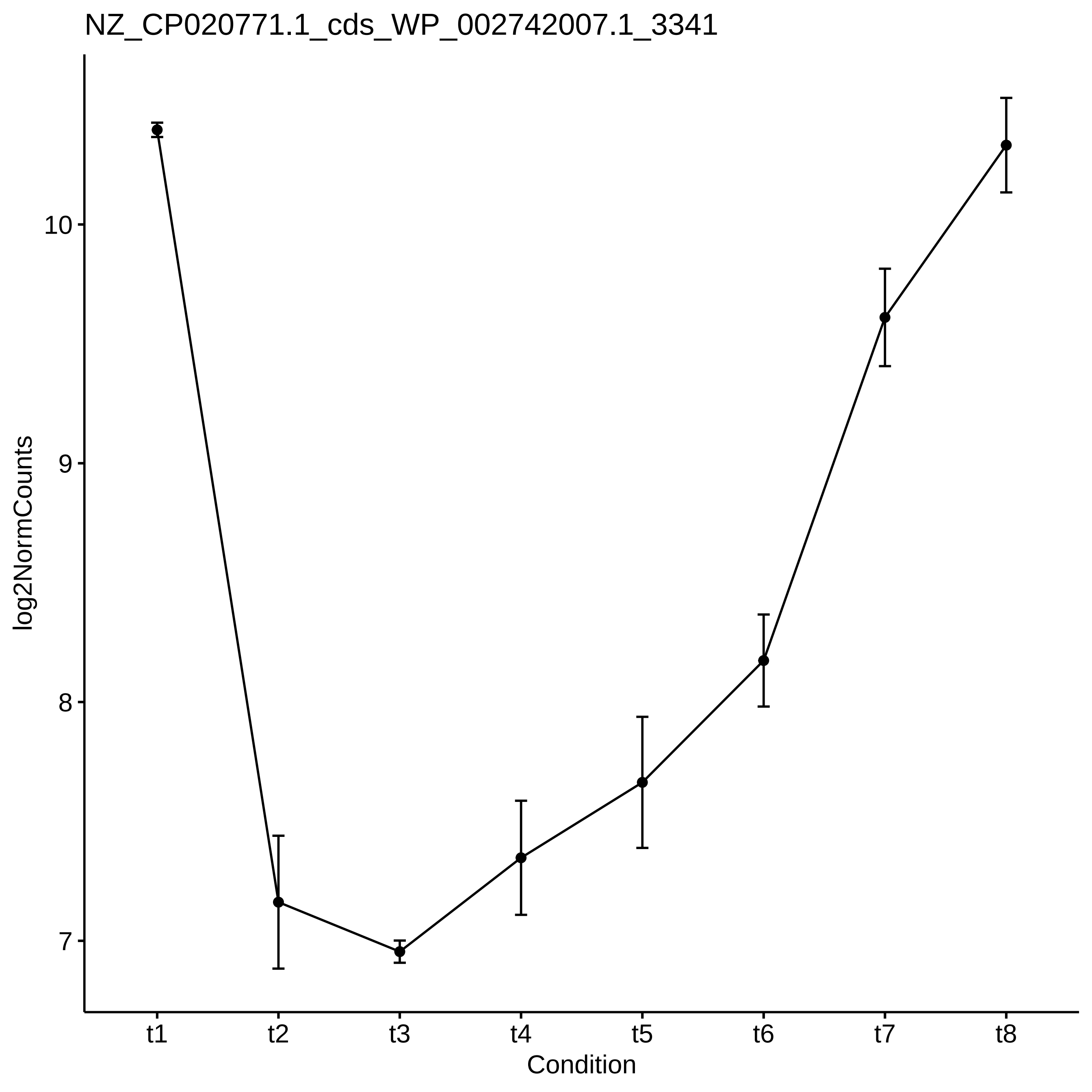
NZ_CP020771.1_cds_WP_002742007.1_3341
log2NormCounts
10
9
8
7
t1
t2
t3
t4
t5
t6
t7
t8
Condition

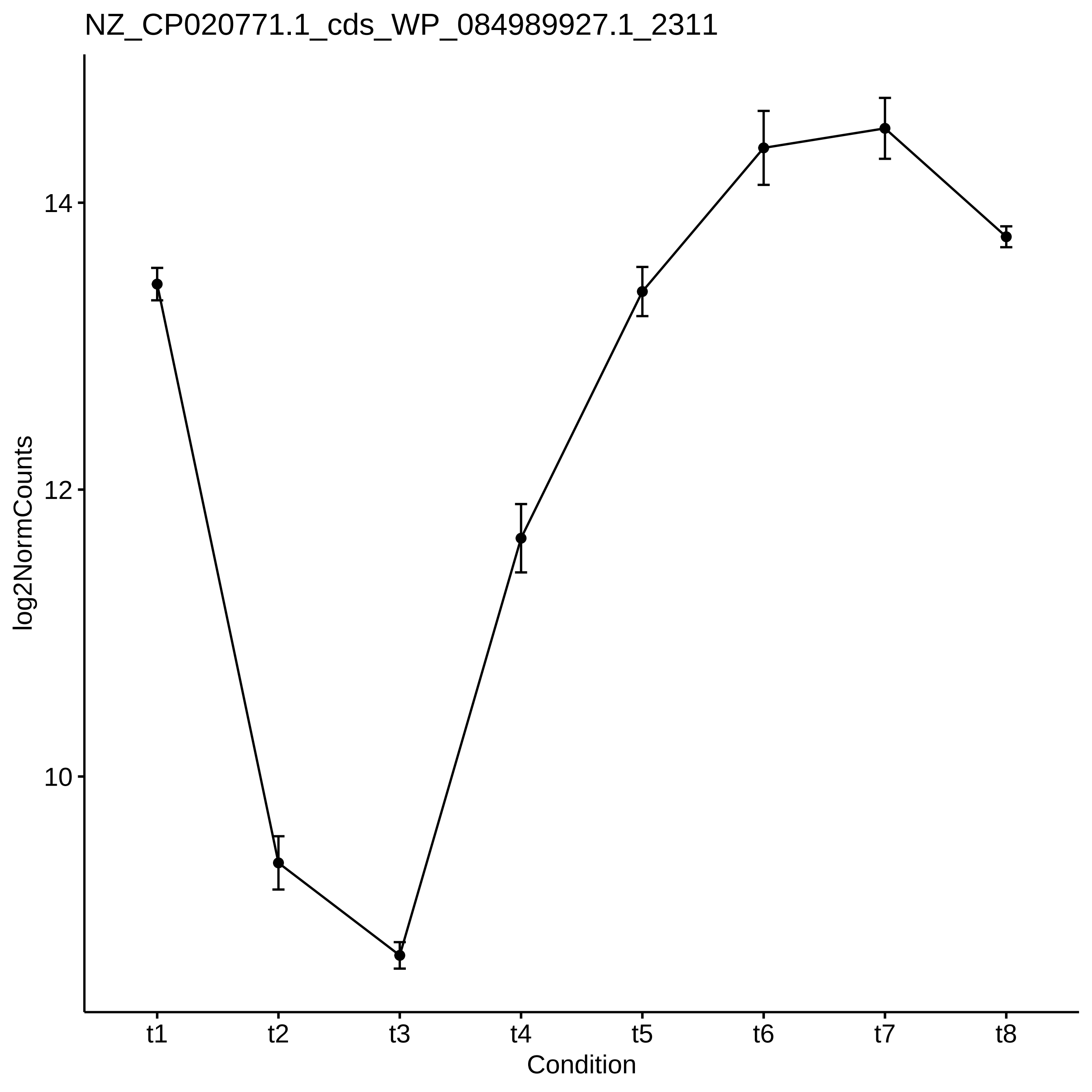
NZ_CP020771.1_cds_WP_084989927.1_2311
log2NormCounts
14
12
10
t1
t2
t3
t4
t5
t6
t7
t8
Condition

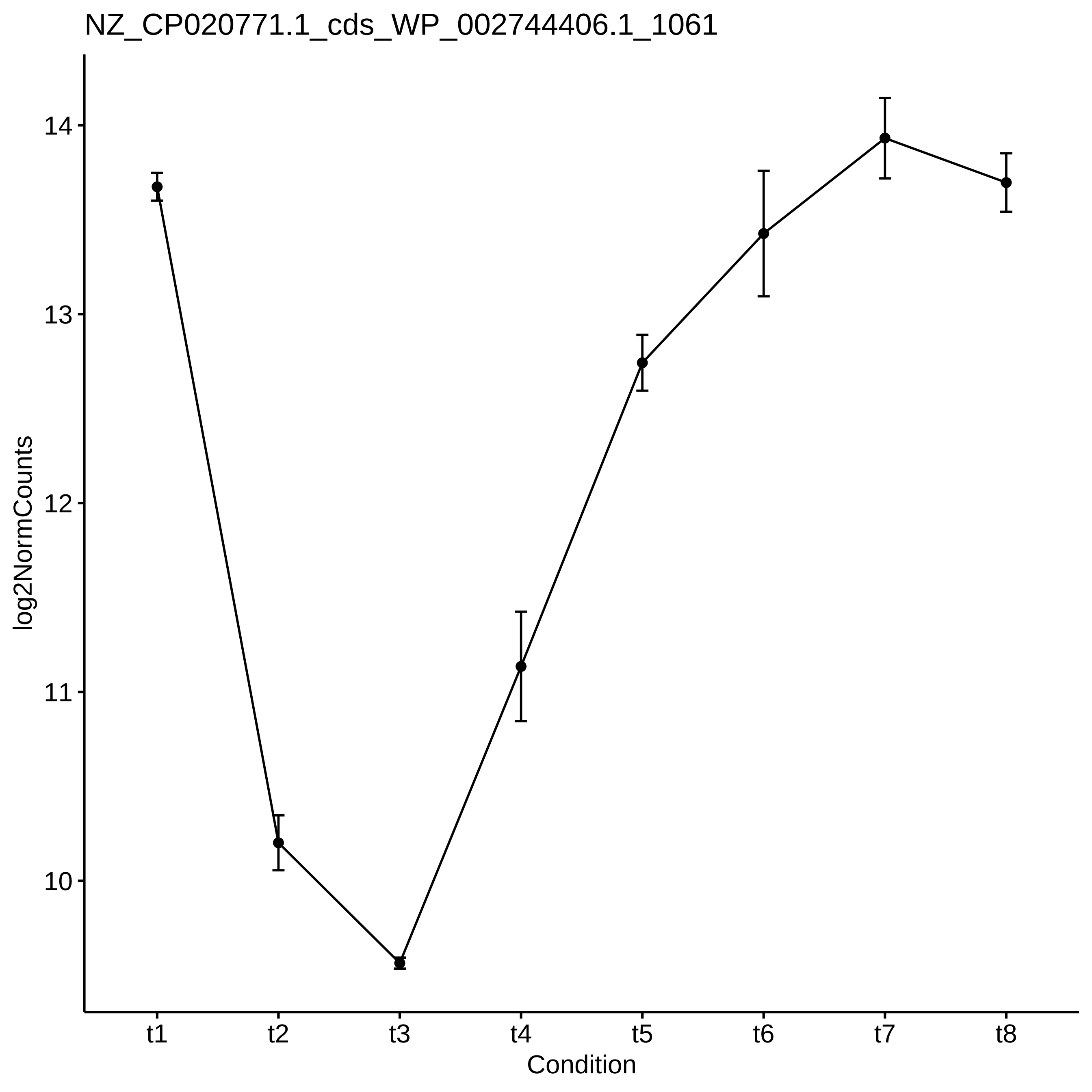
NZ_CP020771.1_cds_WP_002744406.1_1061
log2NormCounts
14
13
12
11
10
t1
t2
t3
t4
t5
t6
t7
t8
Condition

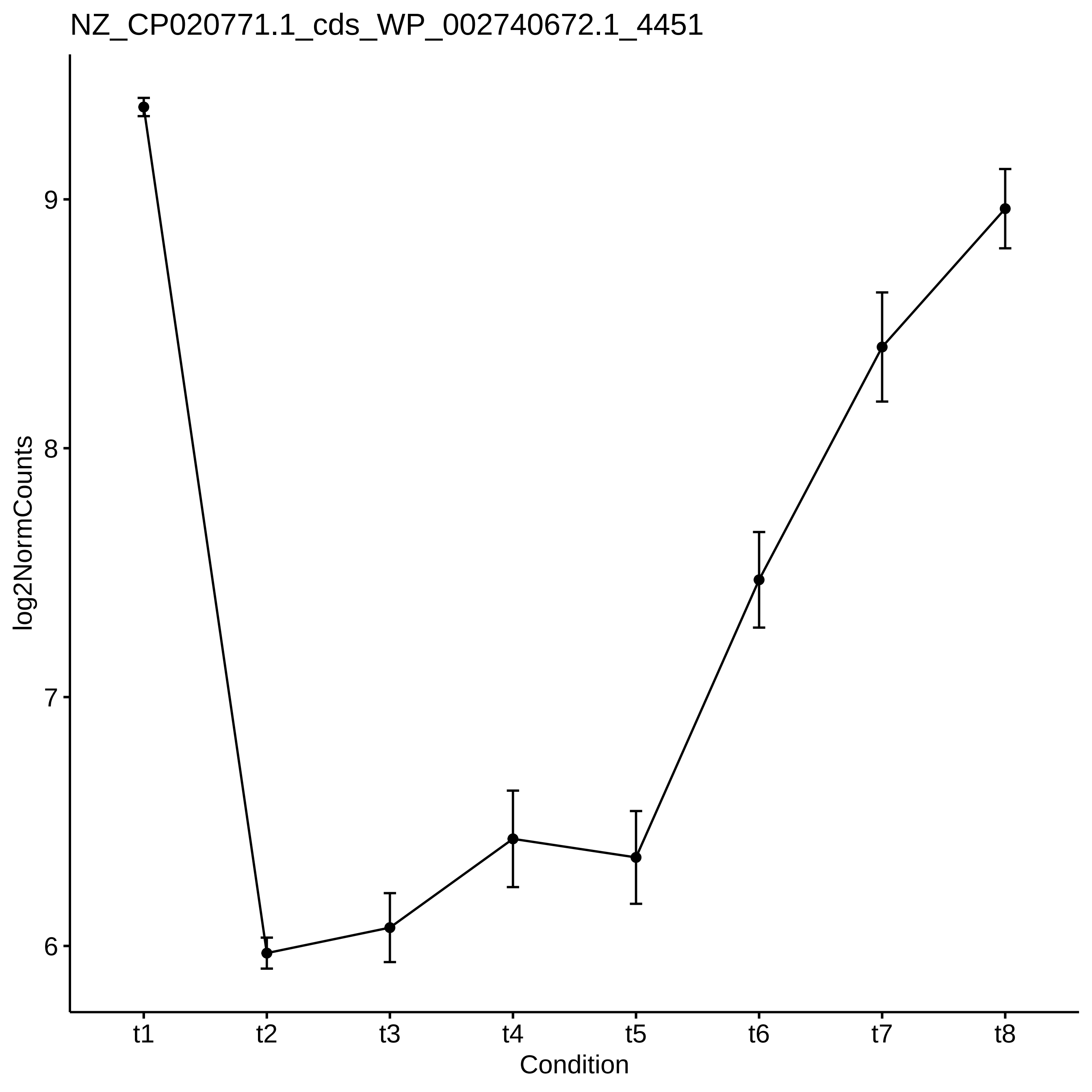
NZ_CP020771.1_cds_WP_002740672.1_4451
log2NormCounts
9
8
7
6
t1
t2
t3
t4
t5
t6
t7
t8
Condition

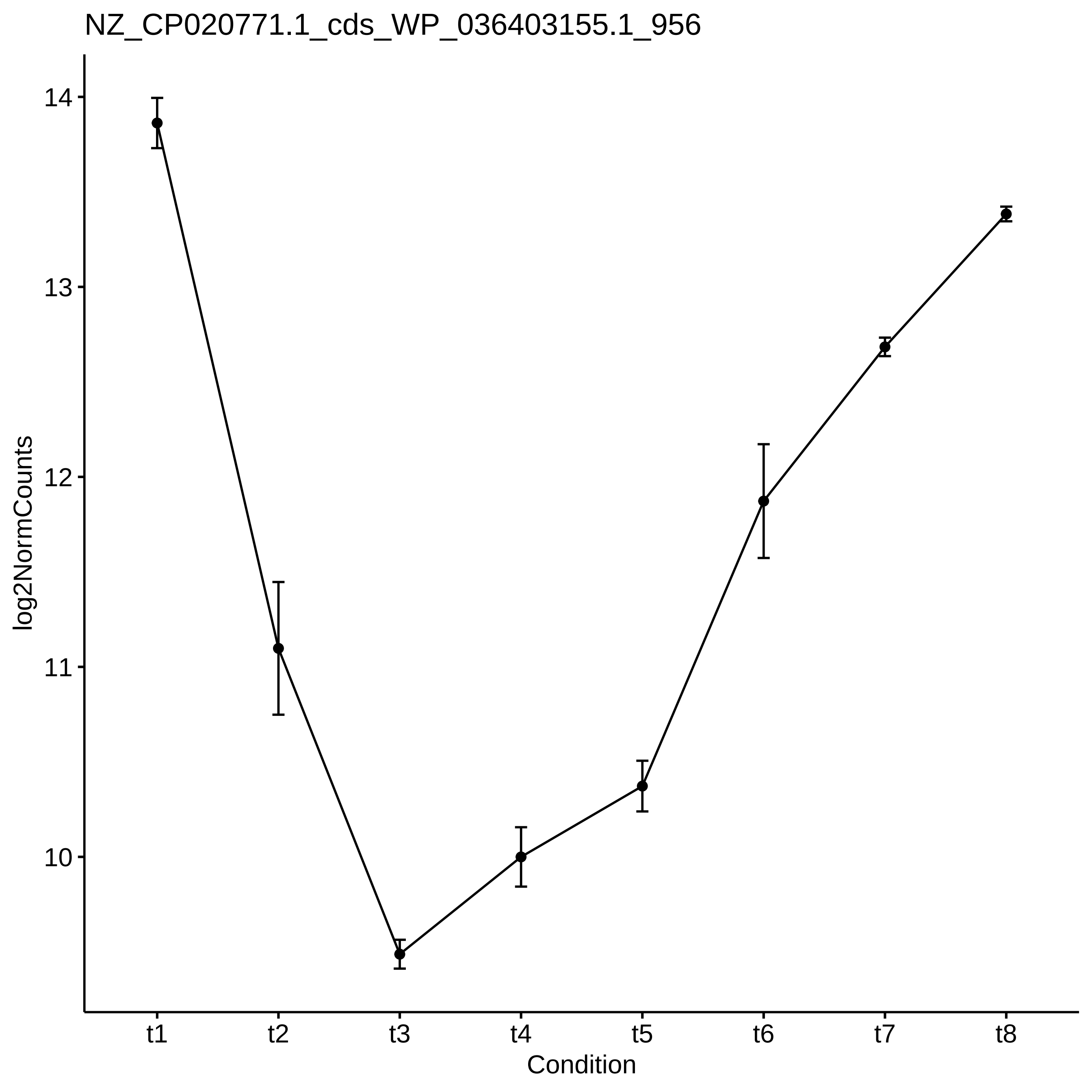
NZ_CP020771.1_cds_WP_036403155.1_956
log2NormCounts
14
13
12
11
10
t1
t2
t3
t4
t5
t6
t7
t8
Condition

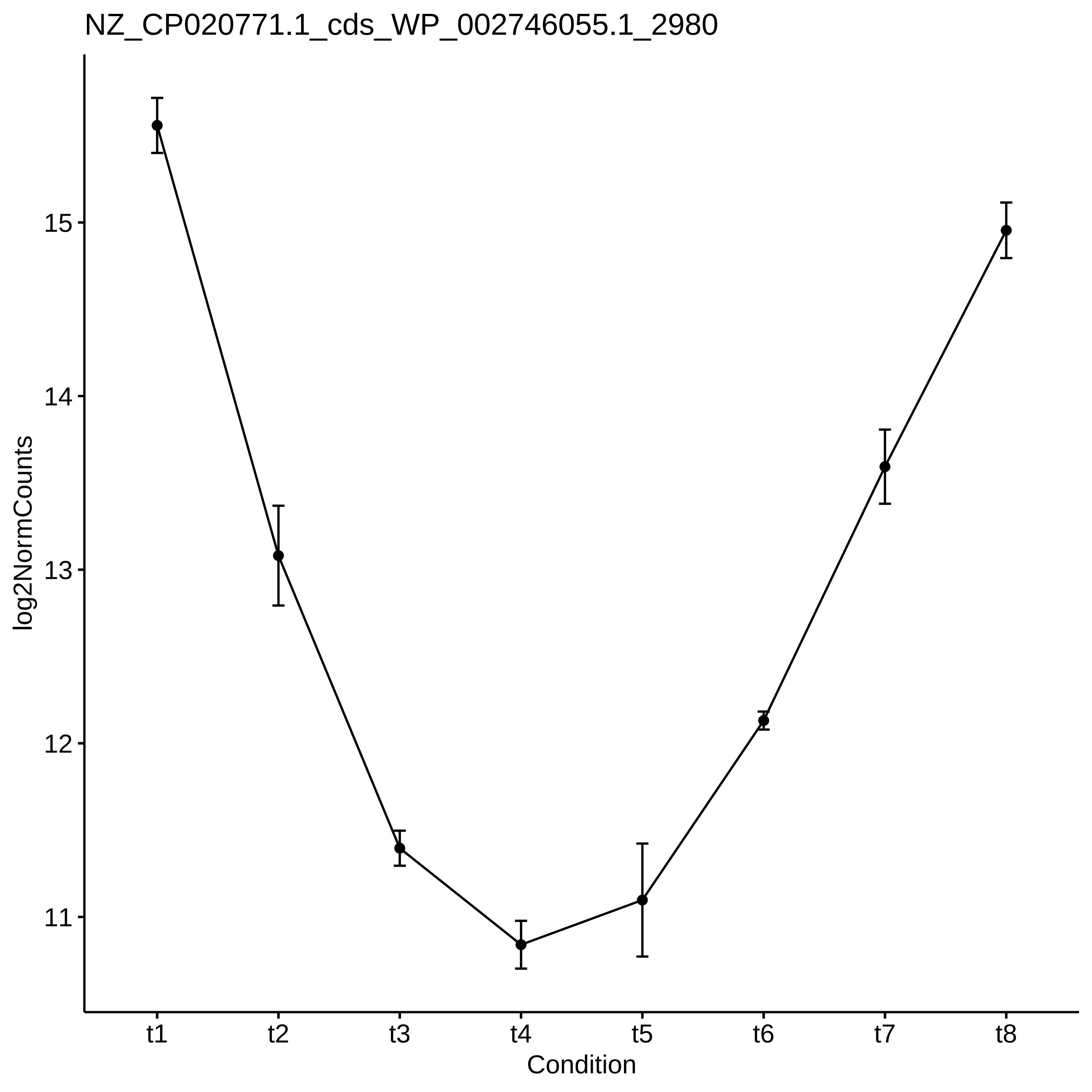
NZ_CP020771.1_cds_WP_002746055.1_2980
log2NormCounts
15
14
13
12
11
t1
t2
t3
t4
t5
t6
t7
t8
Condition

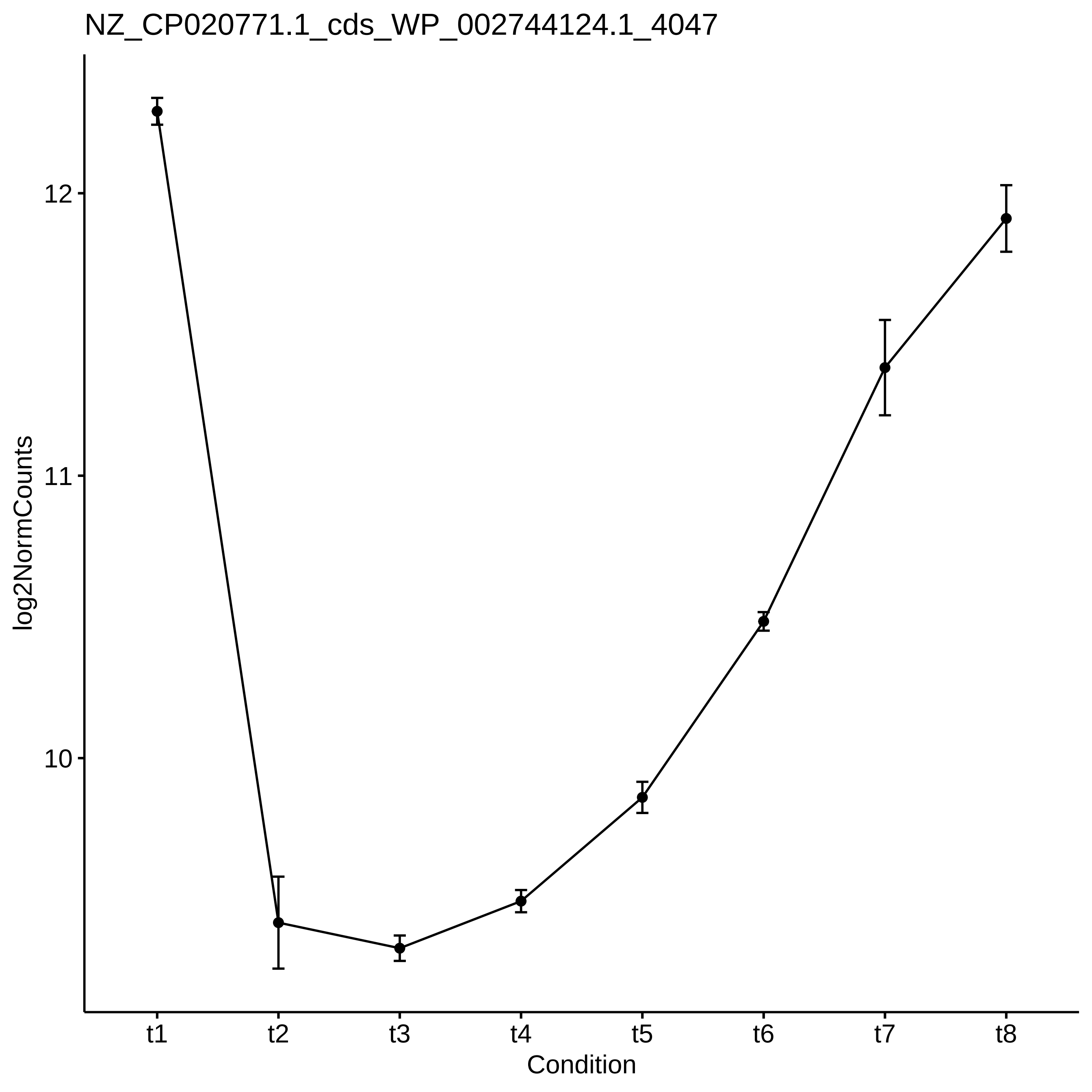

NZ_CP020771.1_cds_WP_002744124.1_4047
log2NormCounts
12
11
10
t1
t2
t3
t4
t5
t6
t7
t8
Condition

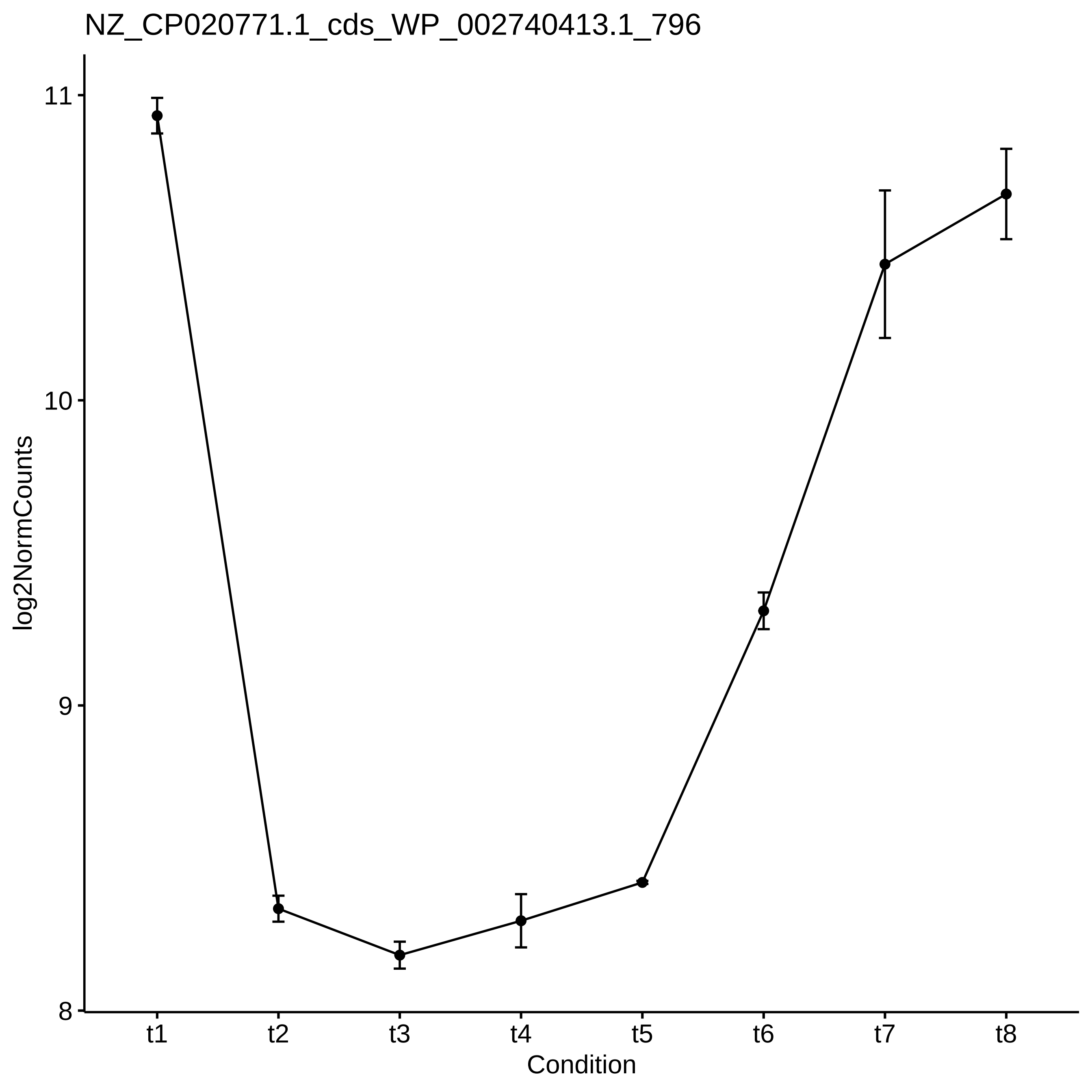
NZ_CP020771.1_cds_WP_002740413.1_796
log2NormCounts
11
10
9
8
t1
t2
t3
t4
t5
t6
t7
t8
Condition
