## Supplementary_Data for "Diel changes in the expression of a marker gene and candidate genes for intracellular amorphous CaCO_3_ biomineralization in *Microcystis*": PCC7806_[t1-t8]_Top50_Profile.pdf

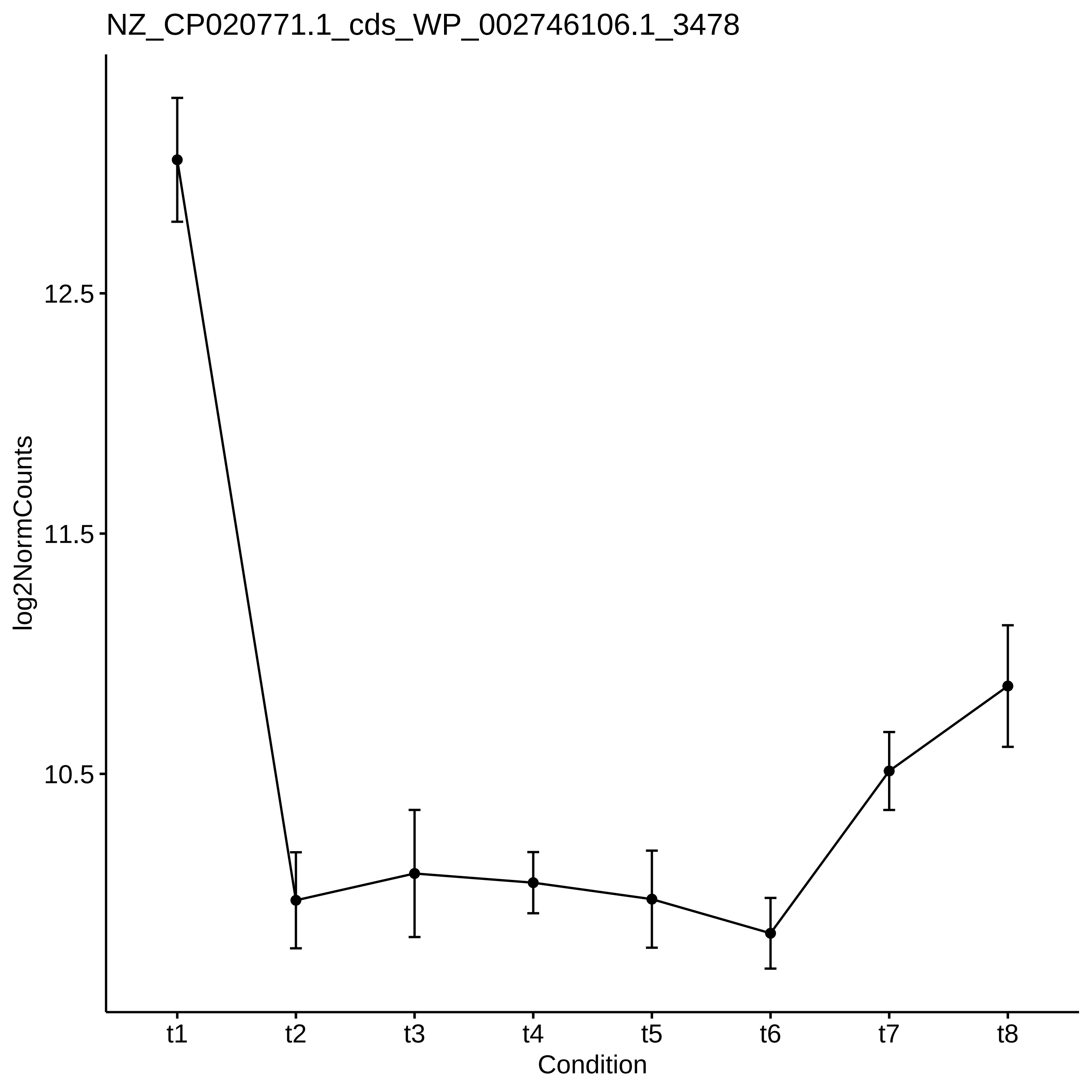
NZ_CP020771.1_cds_WP_002746106.1_3478
log2NormCounts
12.5
11.5
10.5
t1
t2
t3
t4
t5
t6
t7
t8
Condition

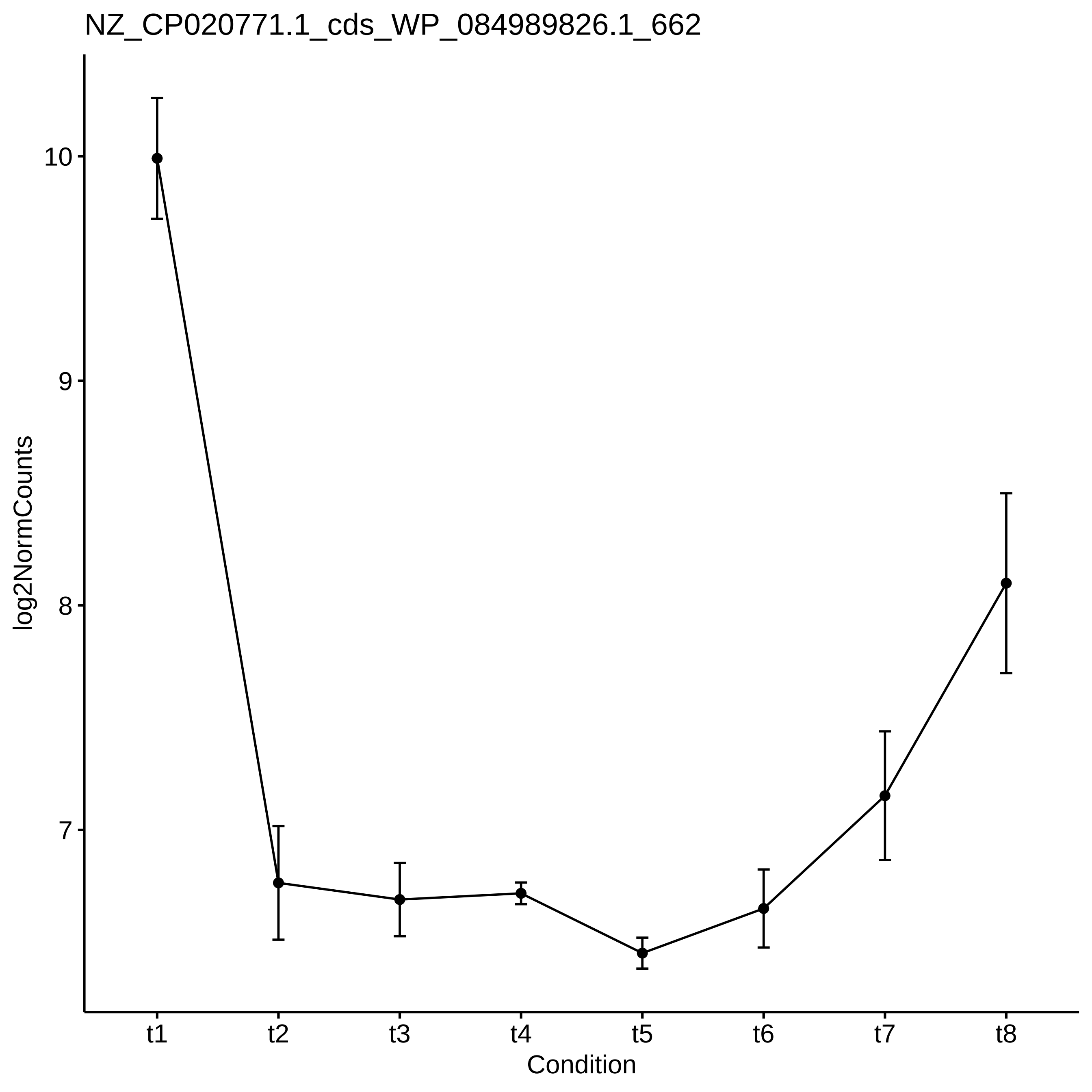
NZ_CP020771.1_cds_WP_084989826.1_662
log2NormCounts
10
9
8
7
t1
t2
t3
t4
t5
t6
t7
t8
Condition

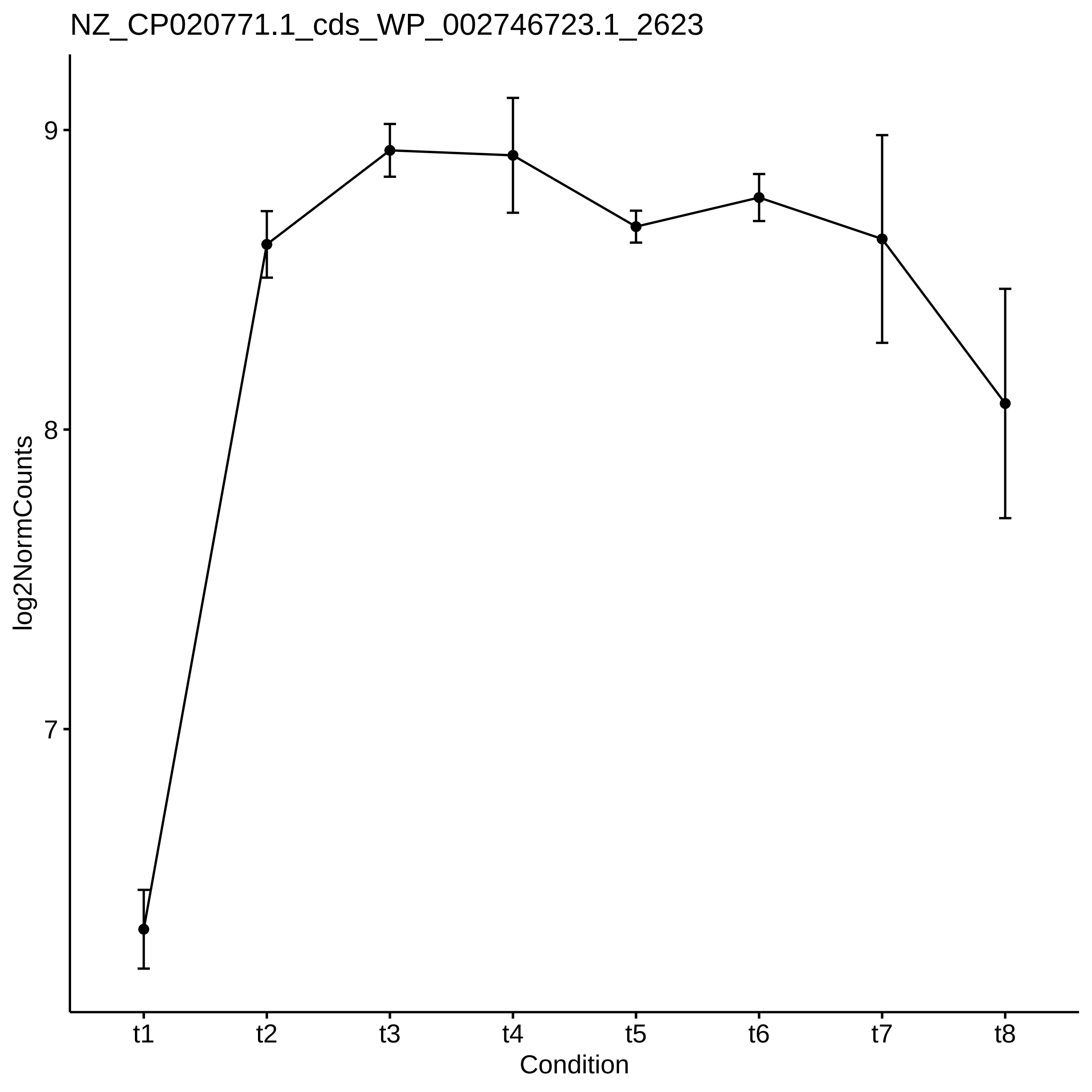
NZ_CP020771.1_cds_WP_002746723.1_2623
log2NormCounts
9
8
7
t1
t2
t3
t4
t5
t6
t7
t8
Condition

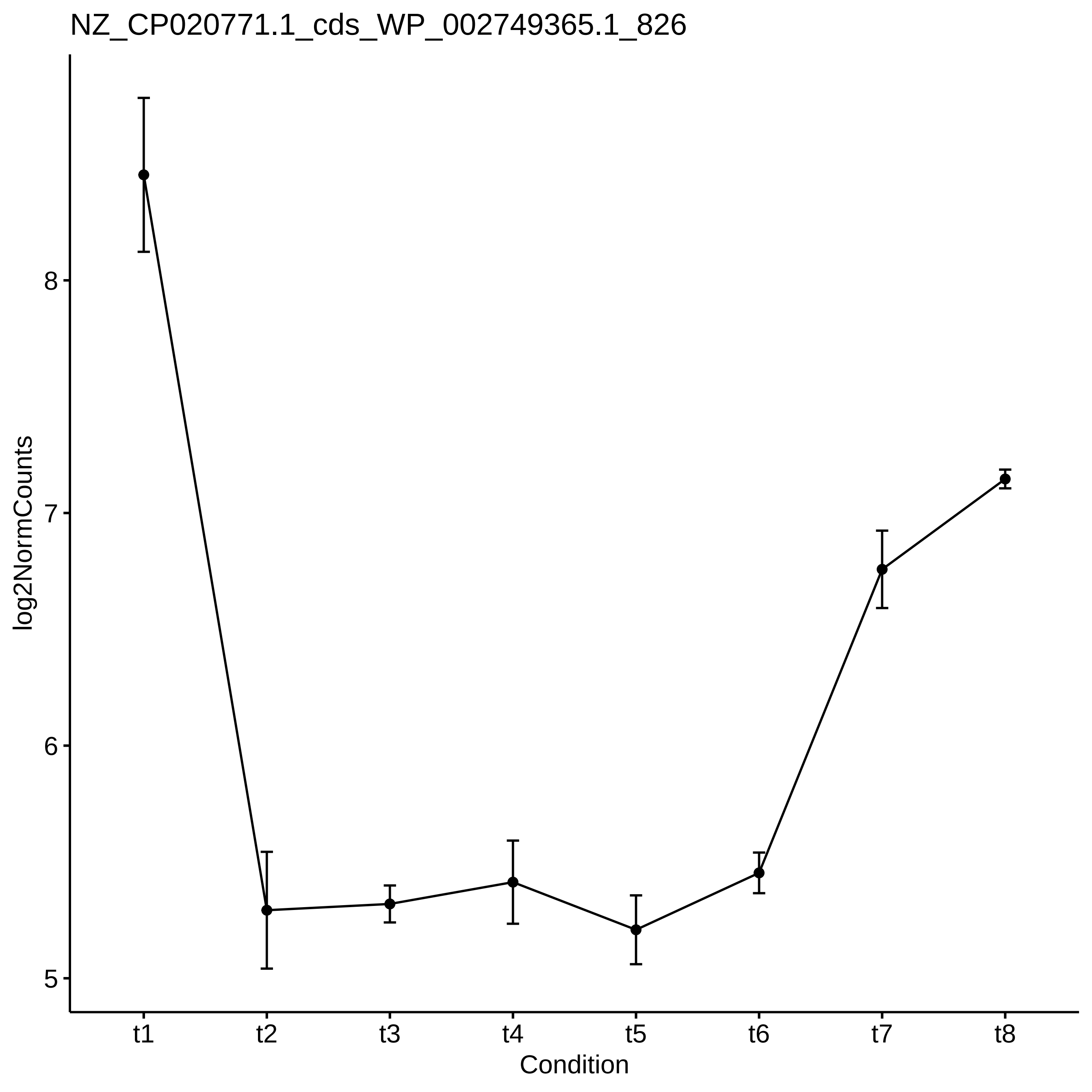

NZ_CP020771.1_cds_WP_002749365.1_826
log2NormCounts
8
7
6
5
t1
t2
t3
t4
t5
t6
t7
t8
Condition

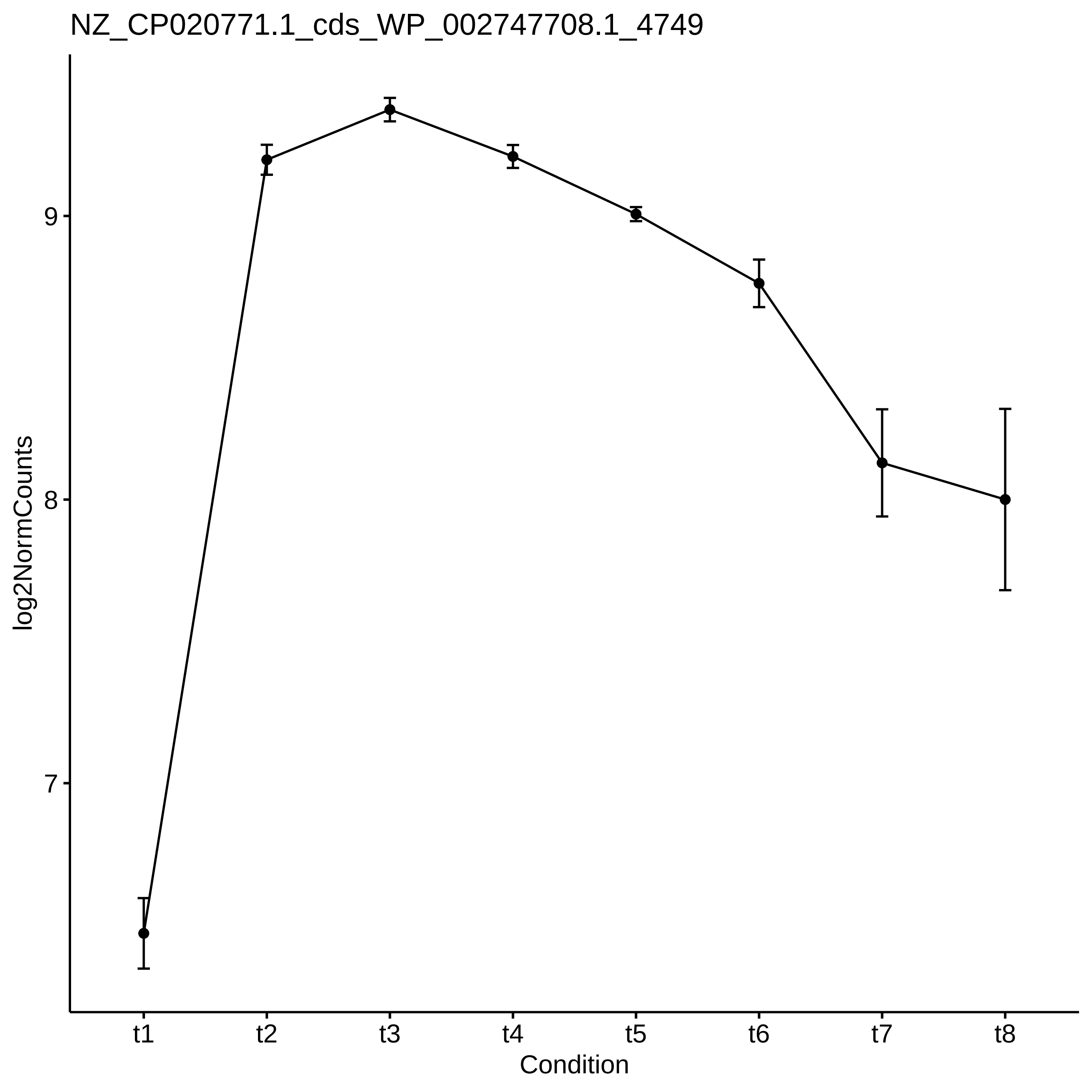

NZ_CP020771.1_cds_WP_002747708.1_4749
log2NormCounts
9
8
7
t1
t2
t3
t4
t5
t6
t7
t8
Condition

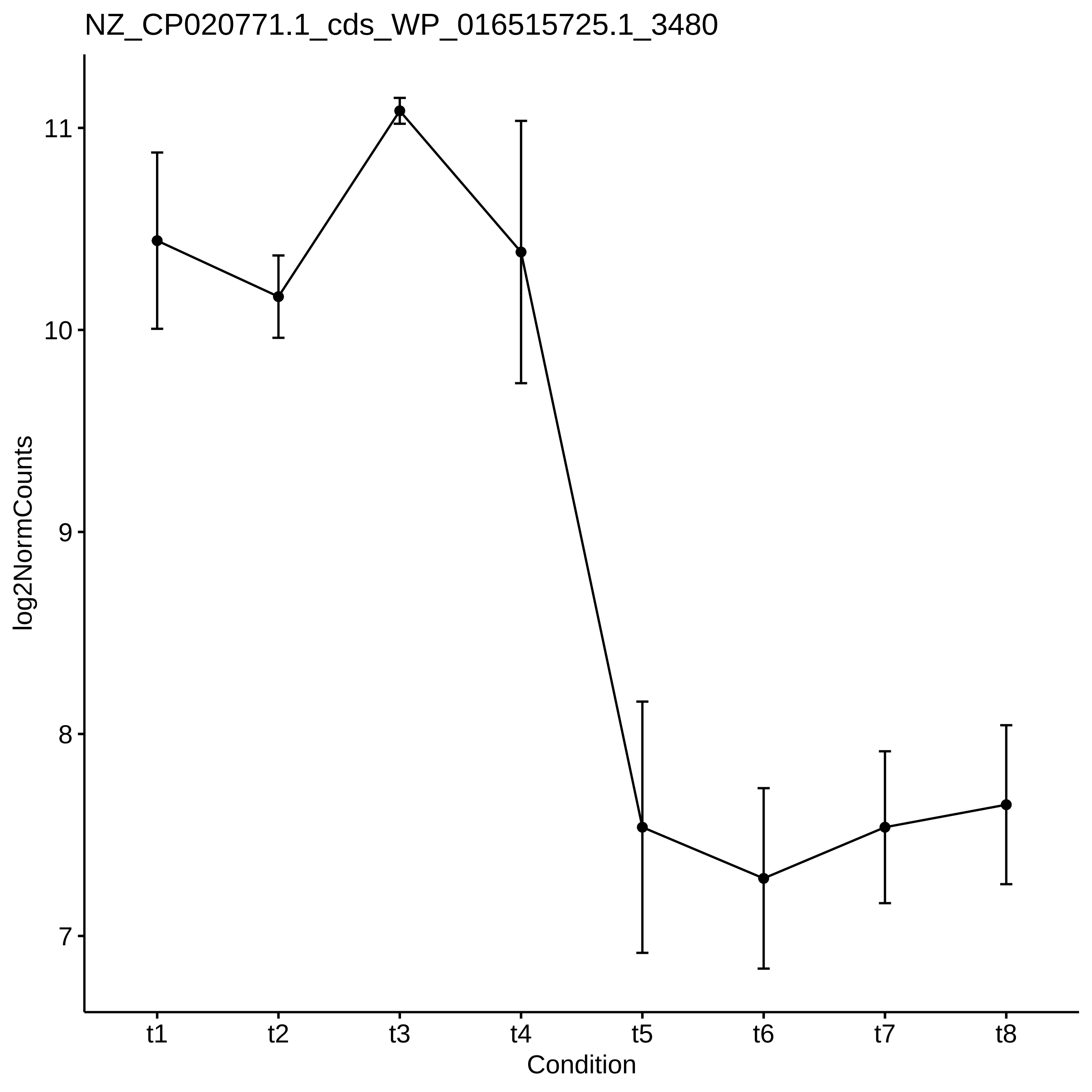

NZ_CP020771.1_cds_WP_016515725.1_3480
log2NormCounts
11
10
9
8
7
t1
t2
t3
t4
t5
t6
t7
t8
Condition

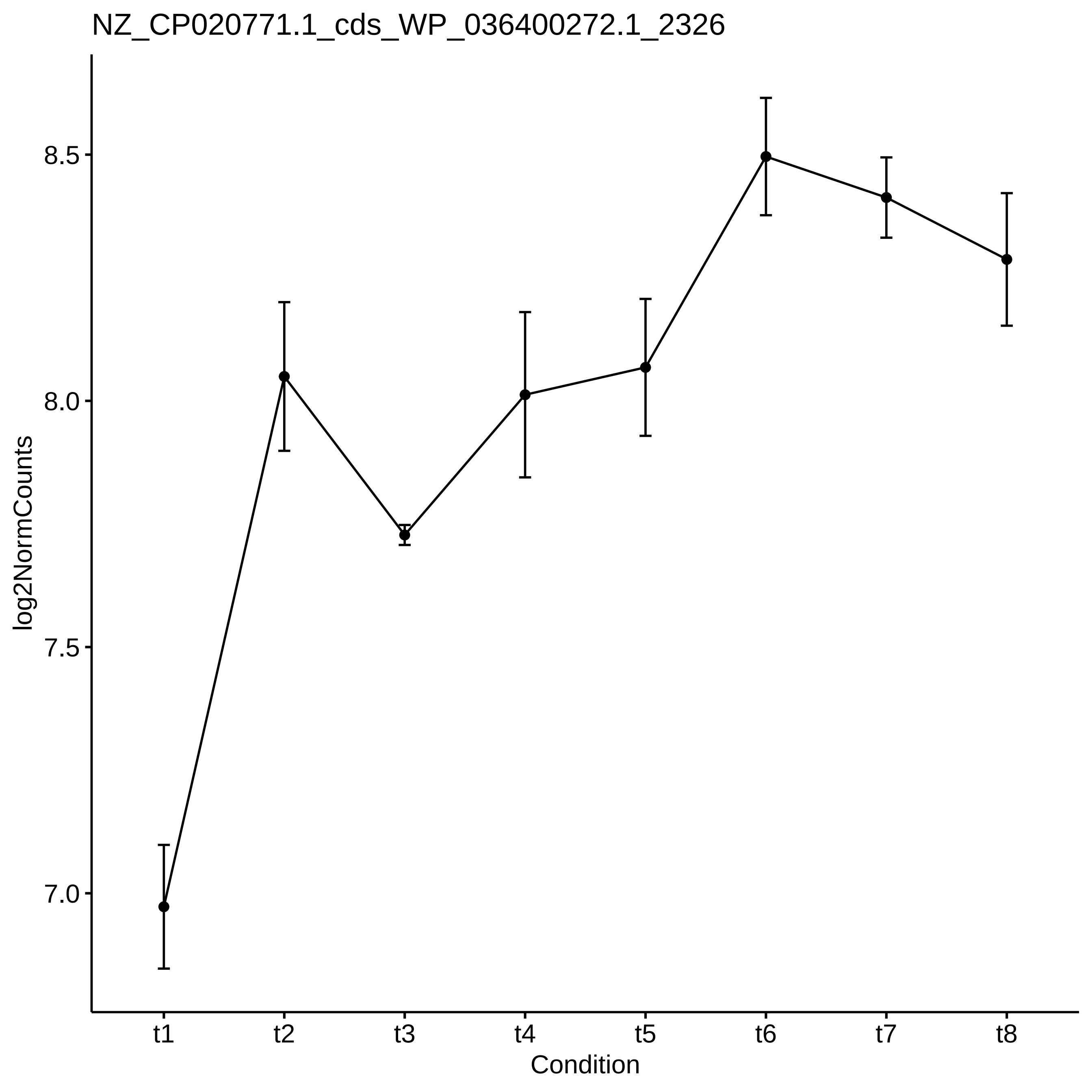
NZ_CP020771.1_cds_WP_036400272.1_2326
log2NormCounts
8.5
8.0
7.5
7.0
t1
t2
t3
t4
t5
t6
t7
t8
Condition

NZ_CP020771.1_cds_WP_002740487.1_754

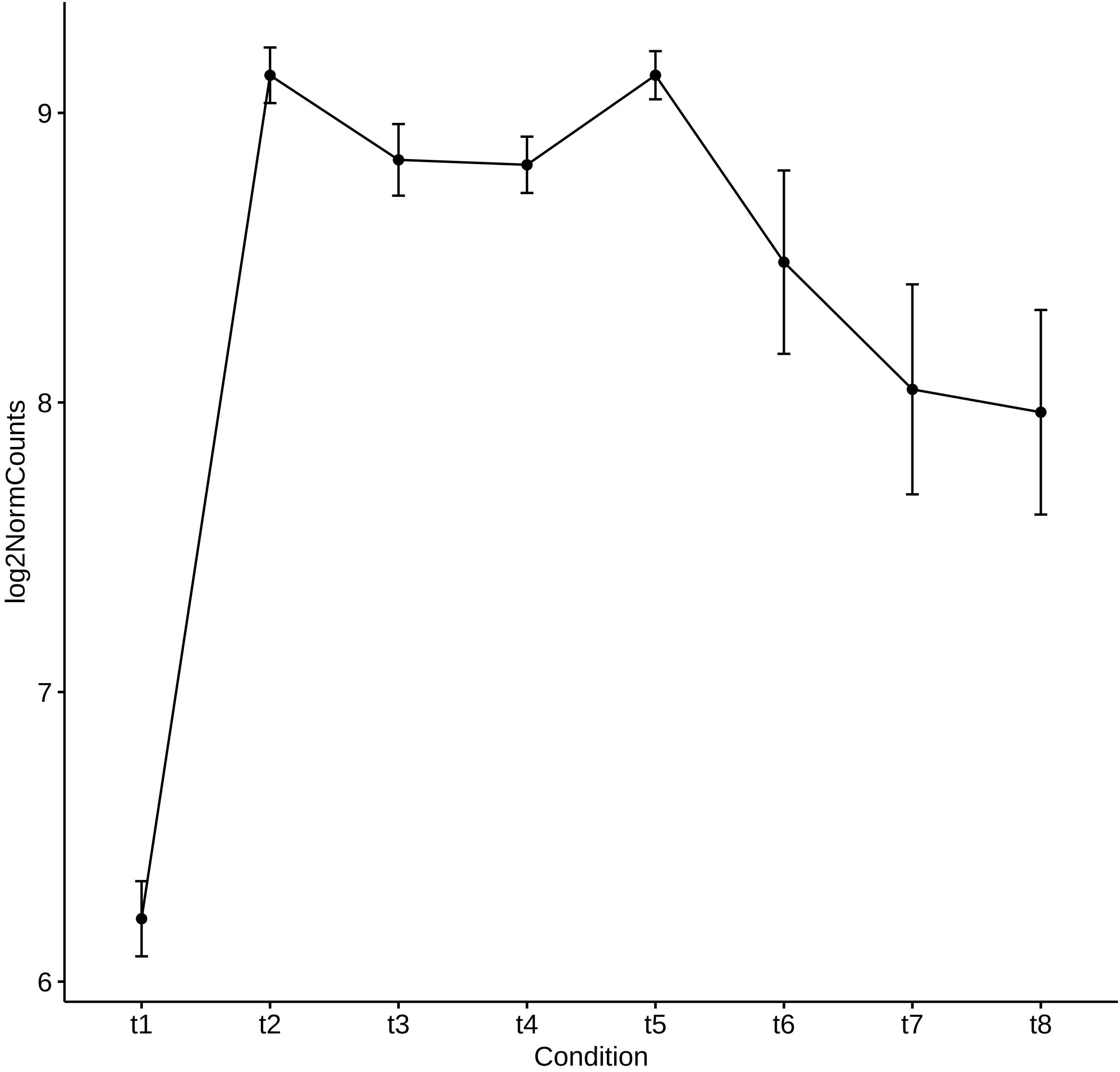

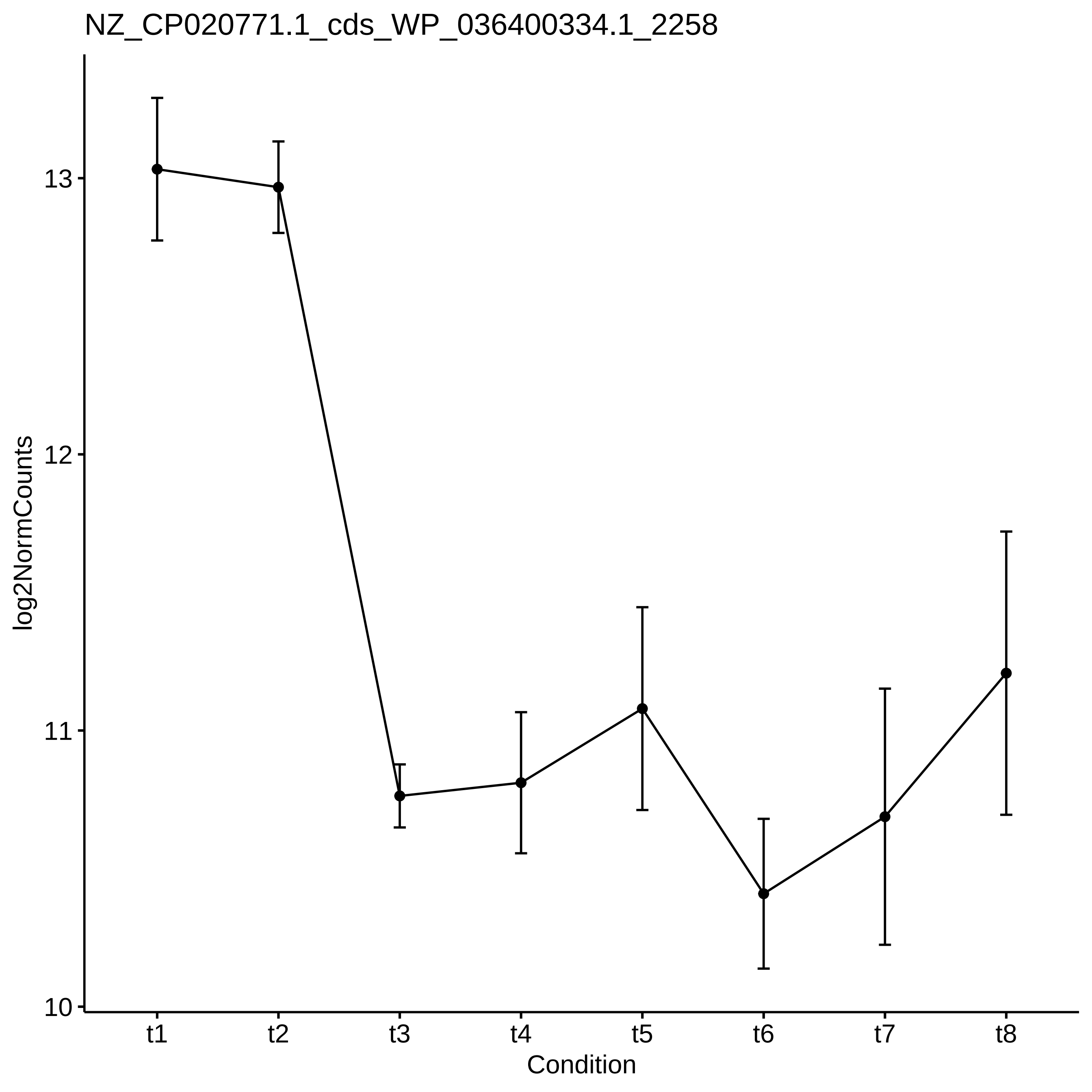

NZ_CP020771.1_cds_WP_036400334.1_2258
log2NormCounts
13
12
11
10
t1
t2
t3
t4
t5
t6
t7
t8
Condition

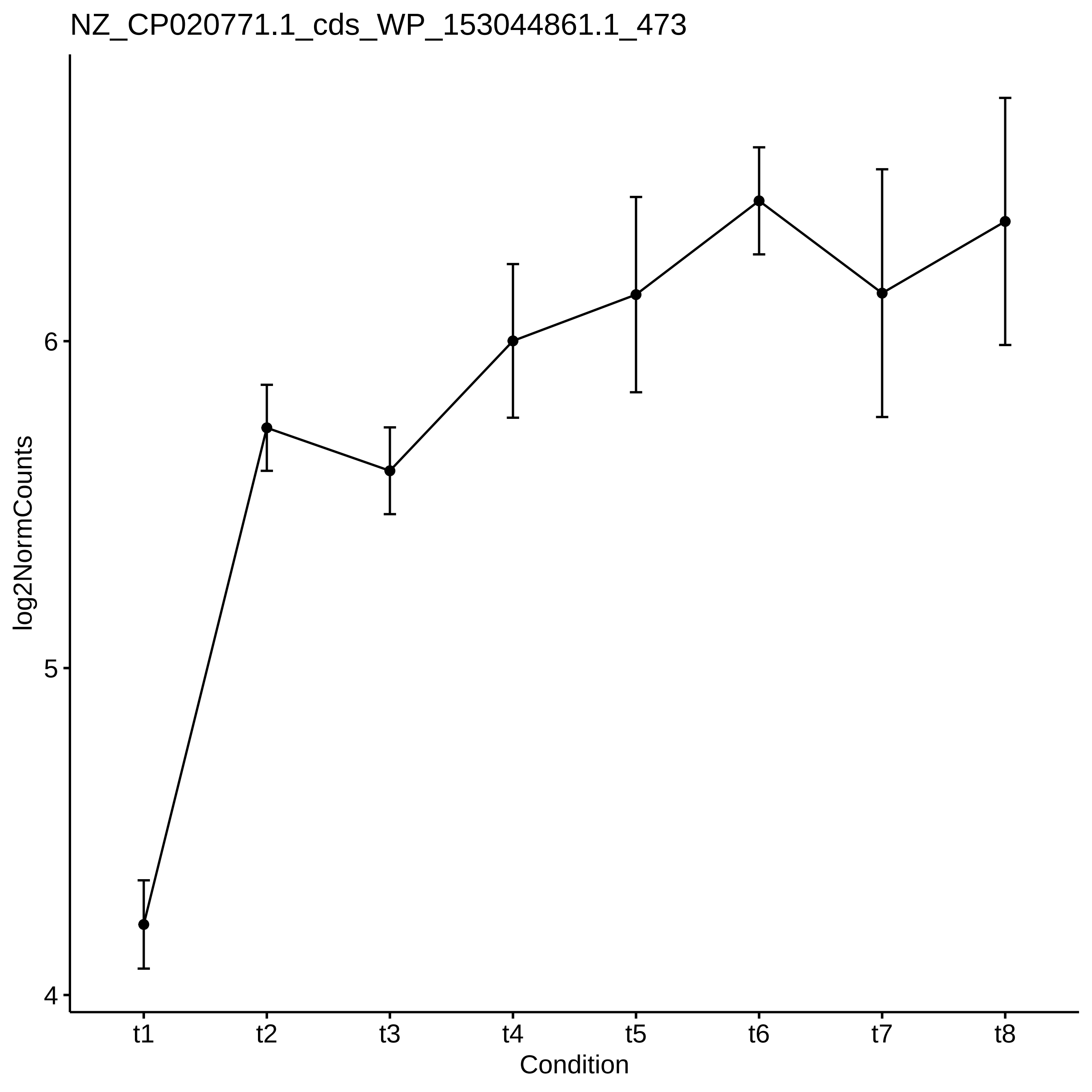
NZ_CP020771.1_cds_WP_153044861.1_473
log2NormCounts
6
5
4
t1
t2
t3
t4
t5
t6
t7
t8
Condition

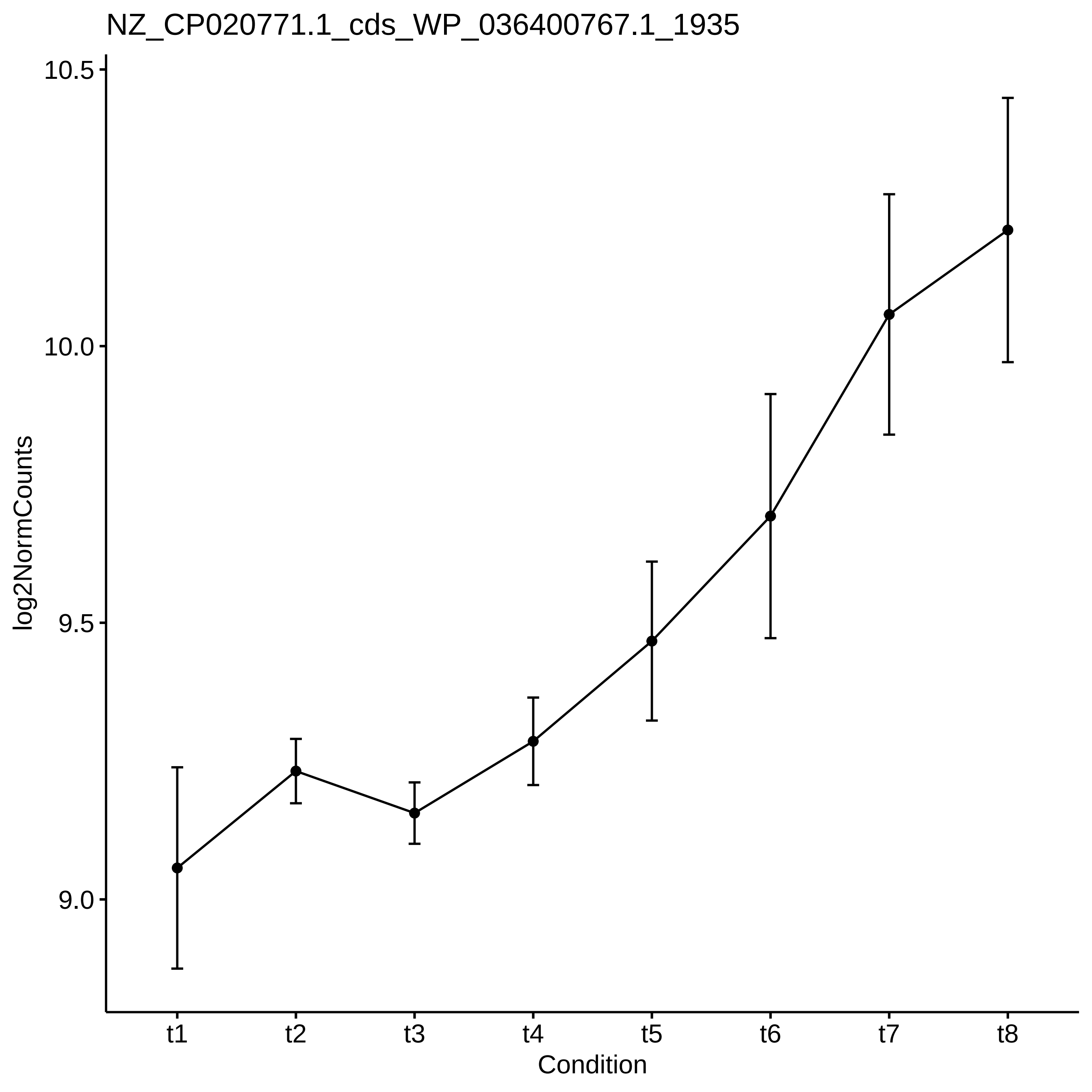

NZ_CP020771.1_cds_WP_036400767.1_1935
log2NormCounts
10.5
10.0
9.5
9.0
t1
t2
t3
t4
t5
t6
t7
t8
Condition

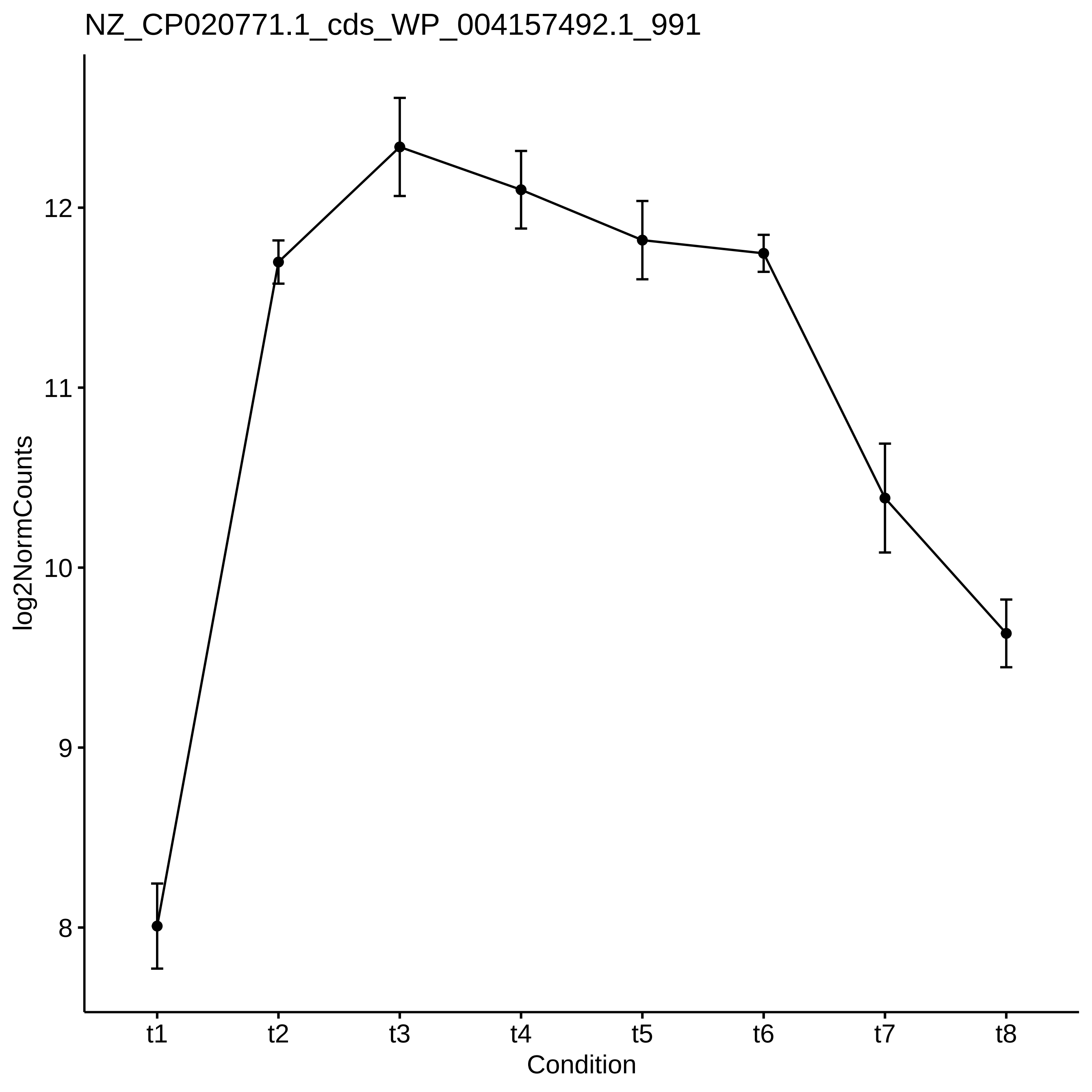
NZ_CP020771.1_cds_WP_004157492.1_991
log2NormCounts
12
11
10
9
8
t1
t2
t3
t4
t5
t6
t7
t8
Condition

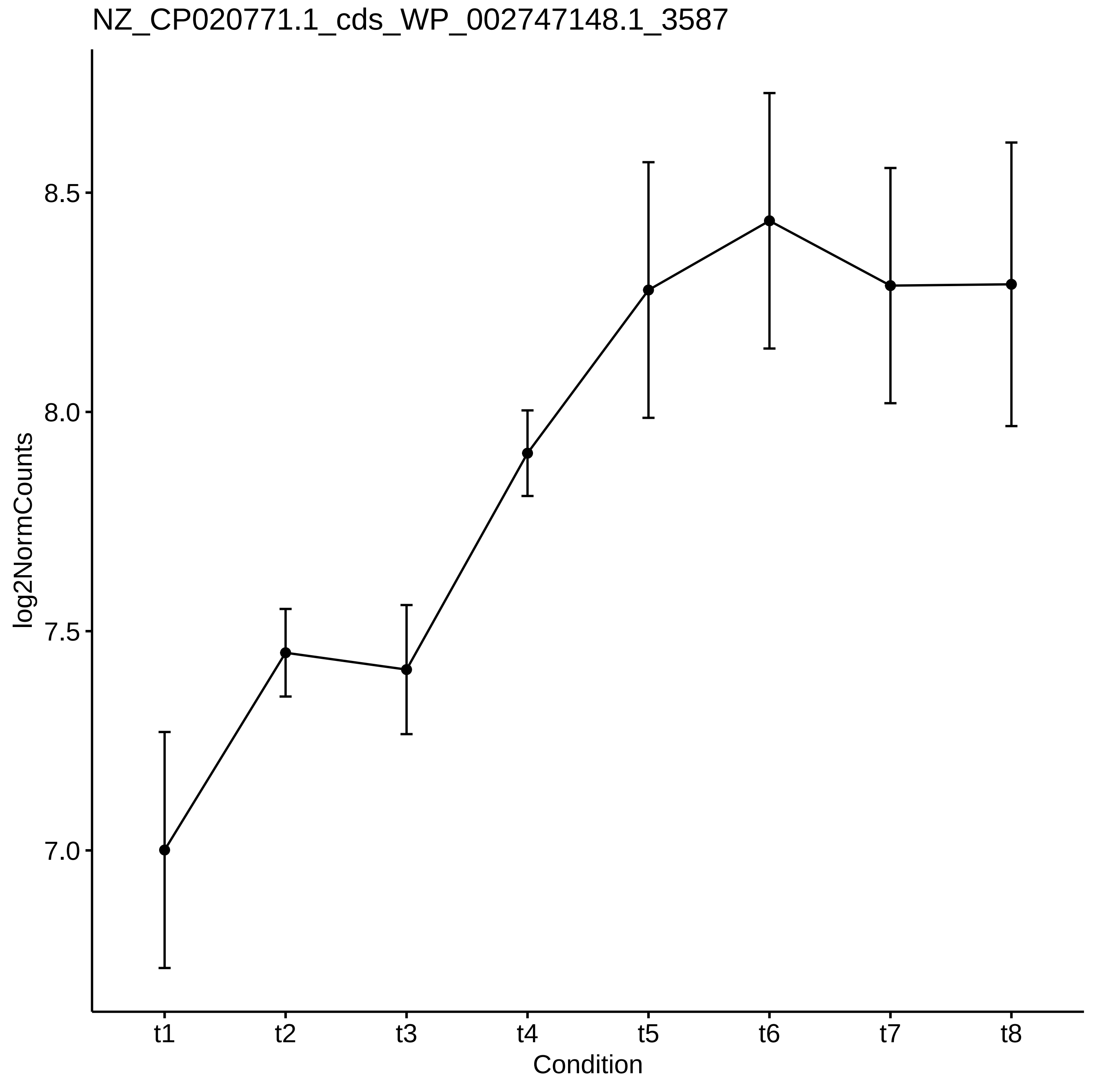

NZ_CP020771.1_cds_WP_002747148.1_3587
log2NormCounts
8.5
8.0
7.5
7.0
t1
t2
t3
t4
t5
t6
t7
t8
Condition

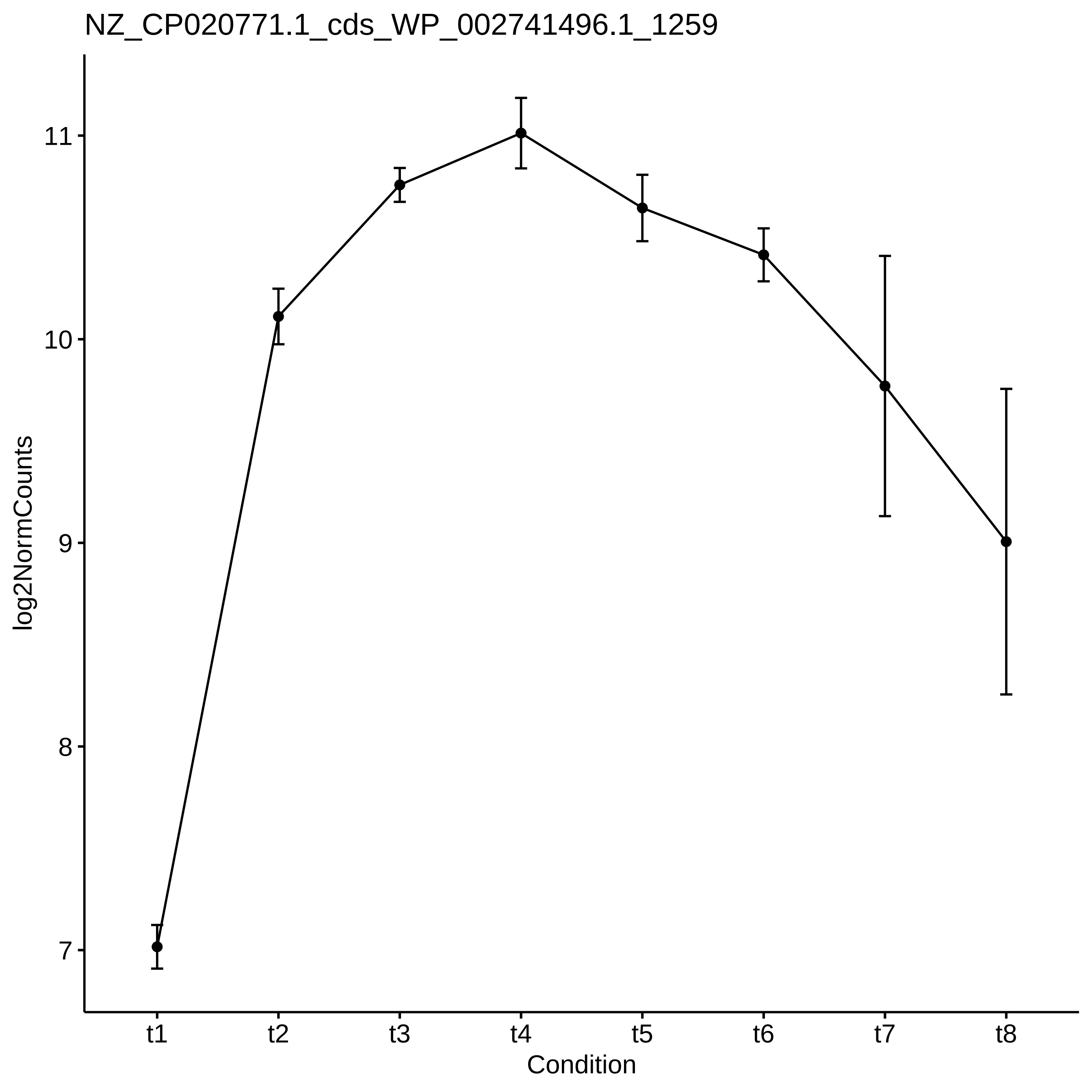
NZ_CP020771.1_cds_WP_002741496.1_1259
log2NormCounts
11
10
9
8
7
t1
t2
t3
t4
t5
t6
t7
t8
Condition

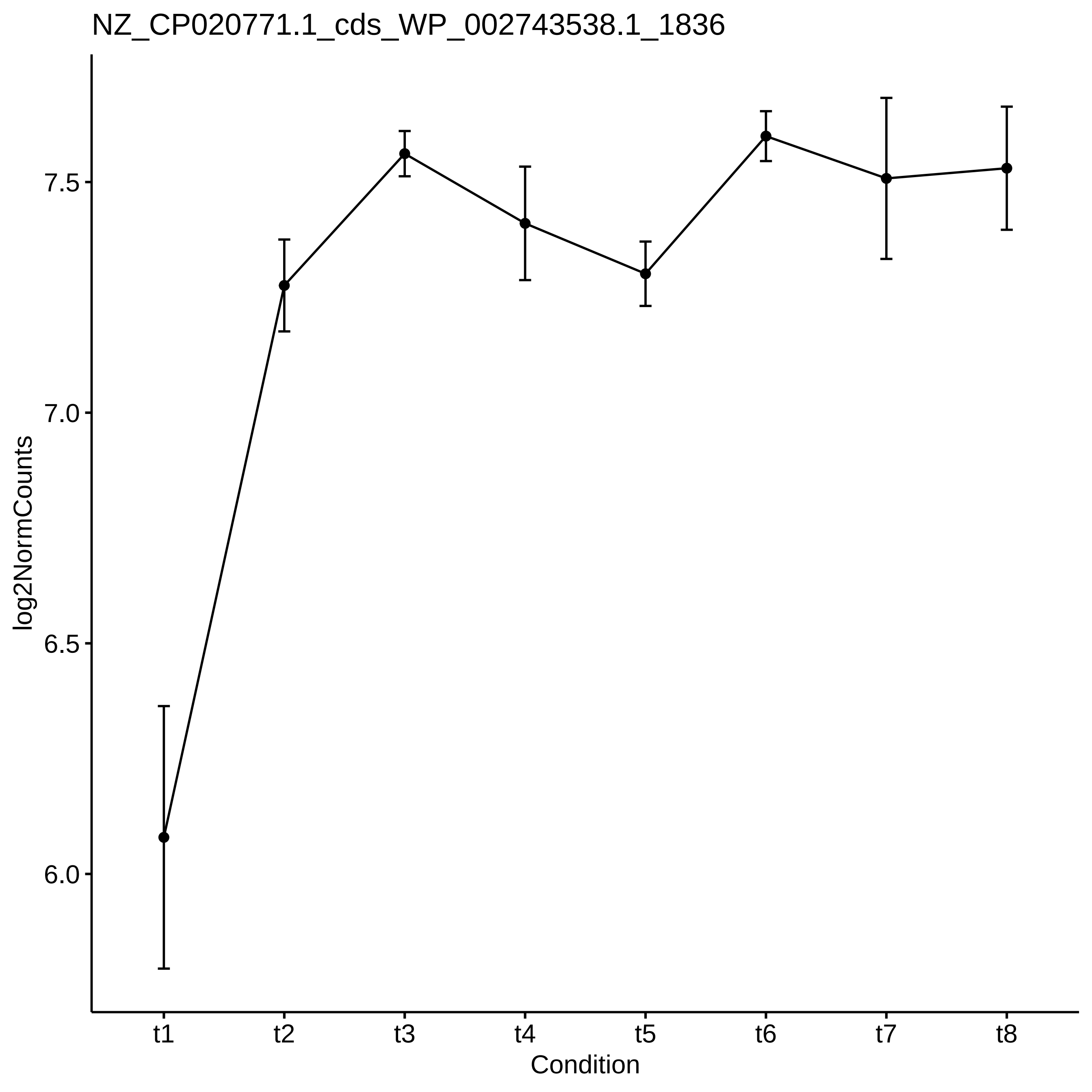

NZ_CP020771.1_cds_WP_002743538.1_1836
log2NormCounts
7.5
7.0
6.5
6.0
t1
t2
t3
t4
t5
t6
t7
t8
Condition

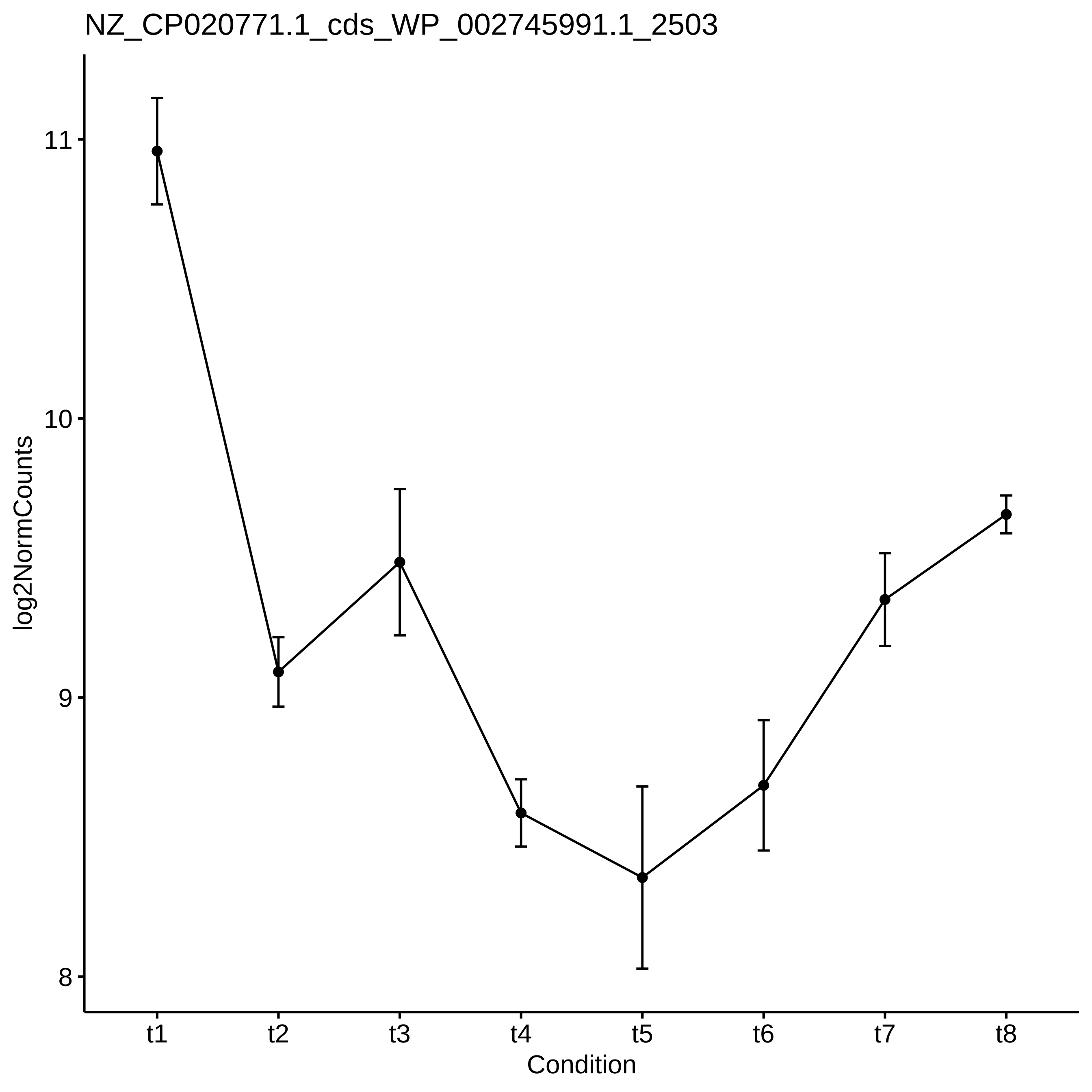

NZ_CP020771.1_cds_WP_002745991.1_2503
log2NormCounts
11
10
9
8
t1
t2
t3
t4
t5
t6
t7
t8
Condition

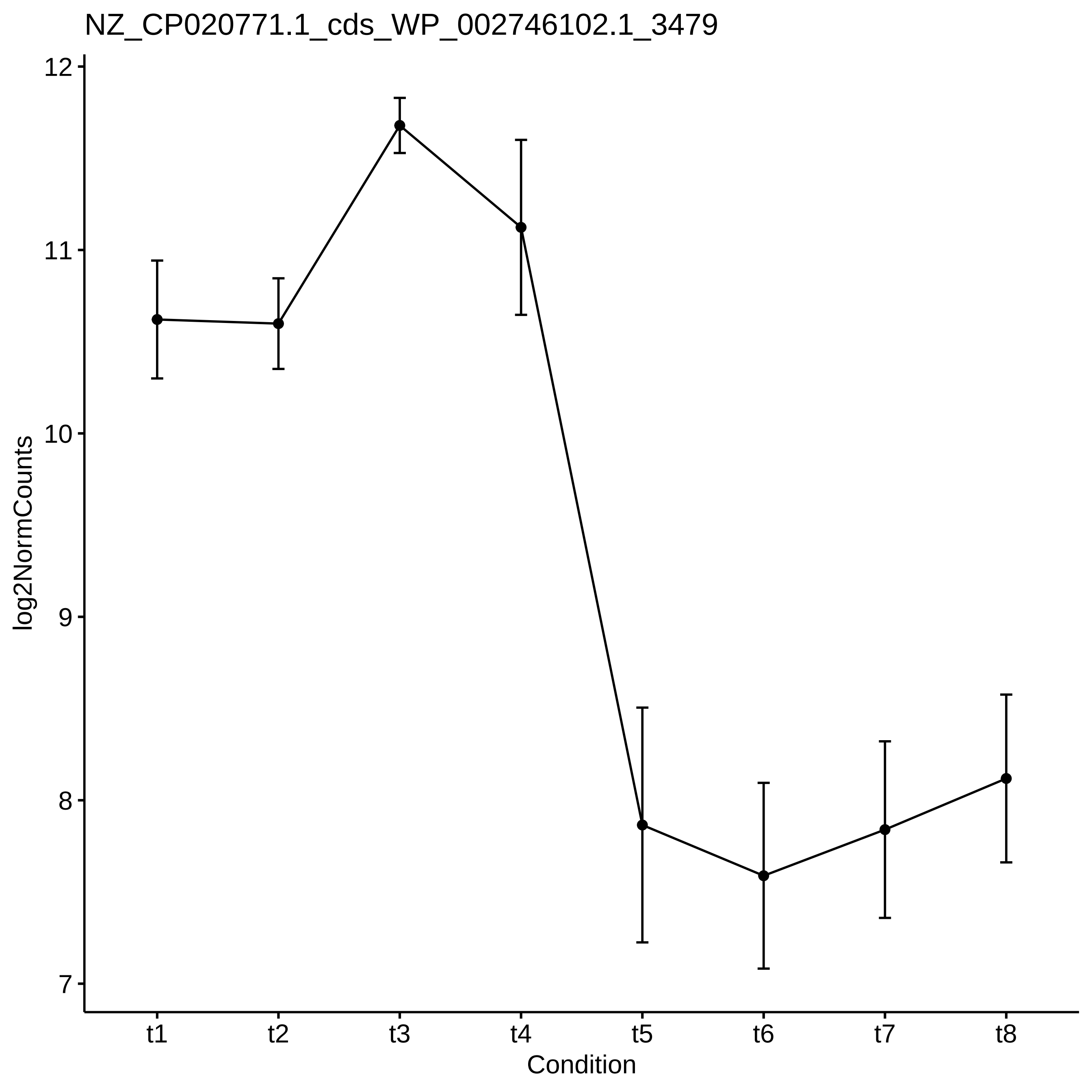
NZ_CP020771.1_cds_WP_002746102.1_3479
log2NormCounts
12
11
10
9
8
7
t1
t2
t3
t4
t5
t6
t7
t8
Condition

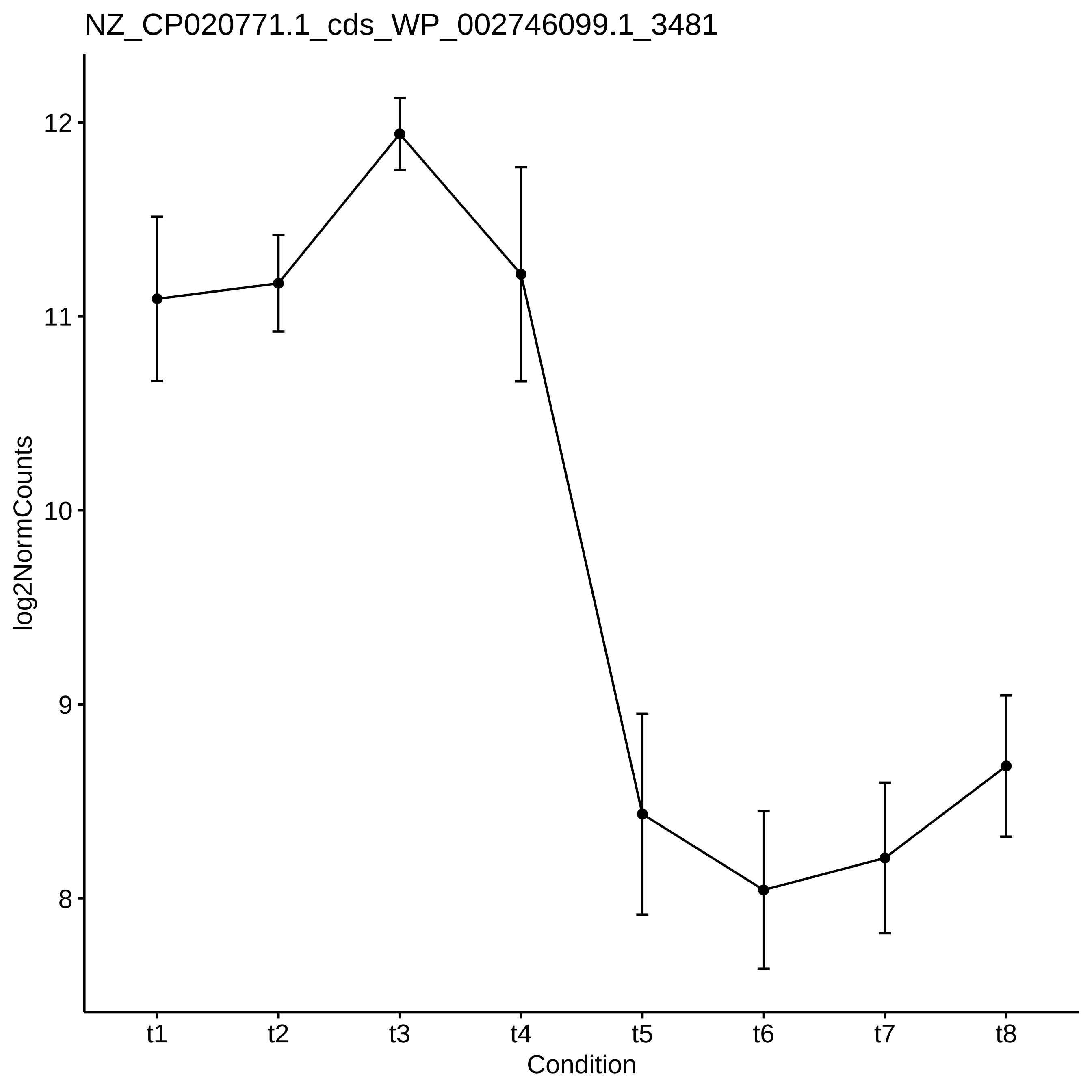

NZ_CP020771.1_cds_WP_002746099.1_3481
log2NormCounts
12
11
10
9
8
t1
t2
t3
t4
t5
t6
t7
t8
Condition

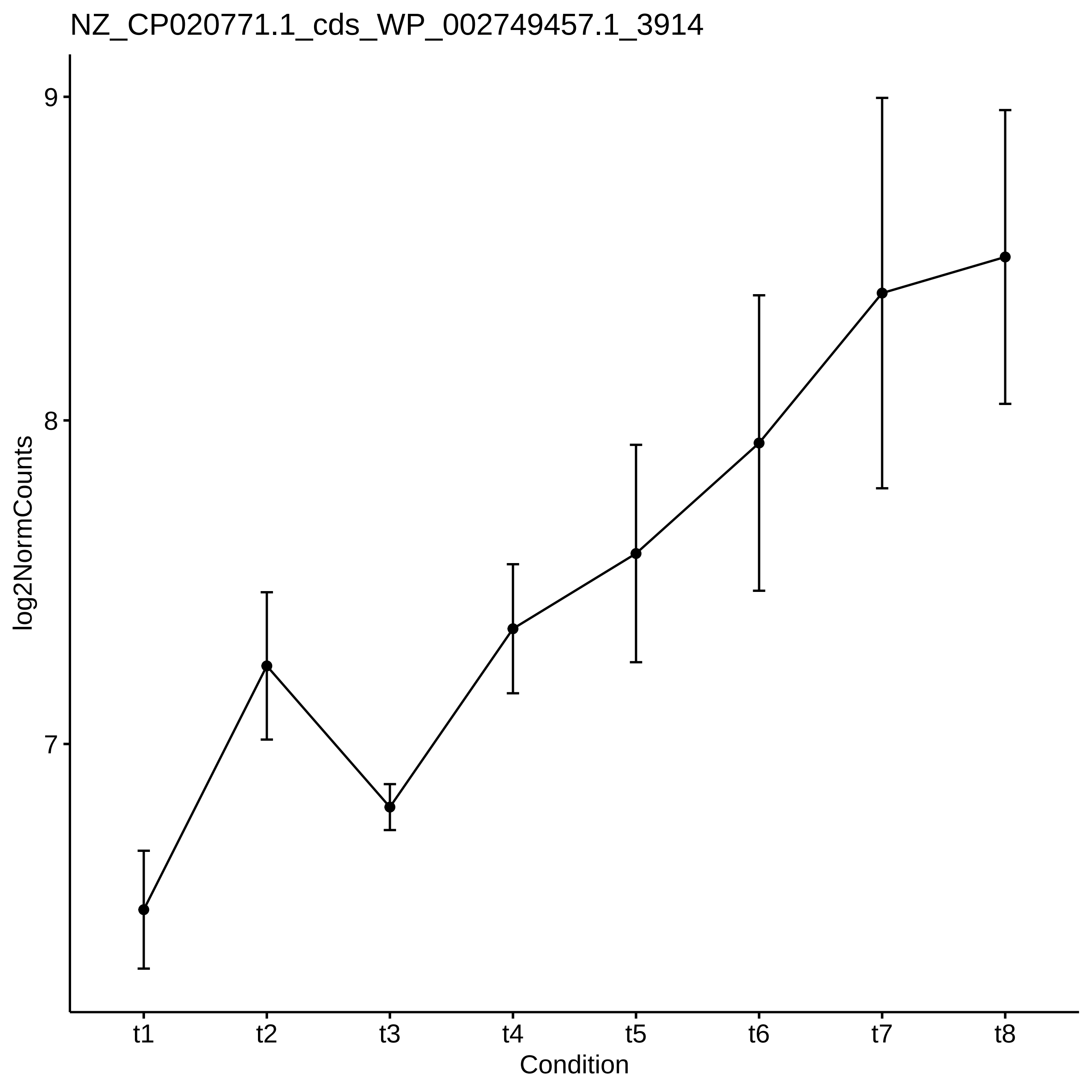

NZ_CP020771.1_cds_WP_002749457.1_3914
log2NormCounts
9
8
7
t1
t2
t3
t4
t5
t6
t7
t8
Condition

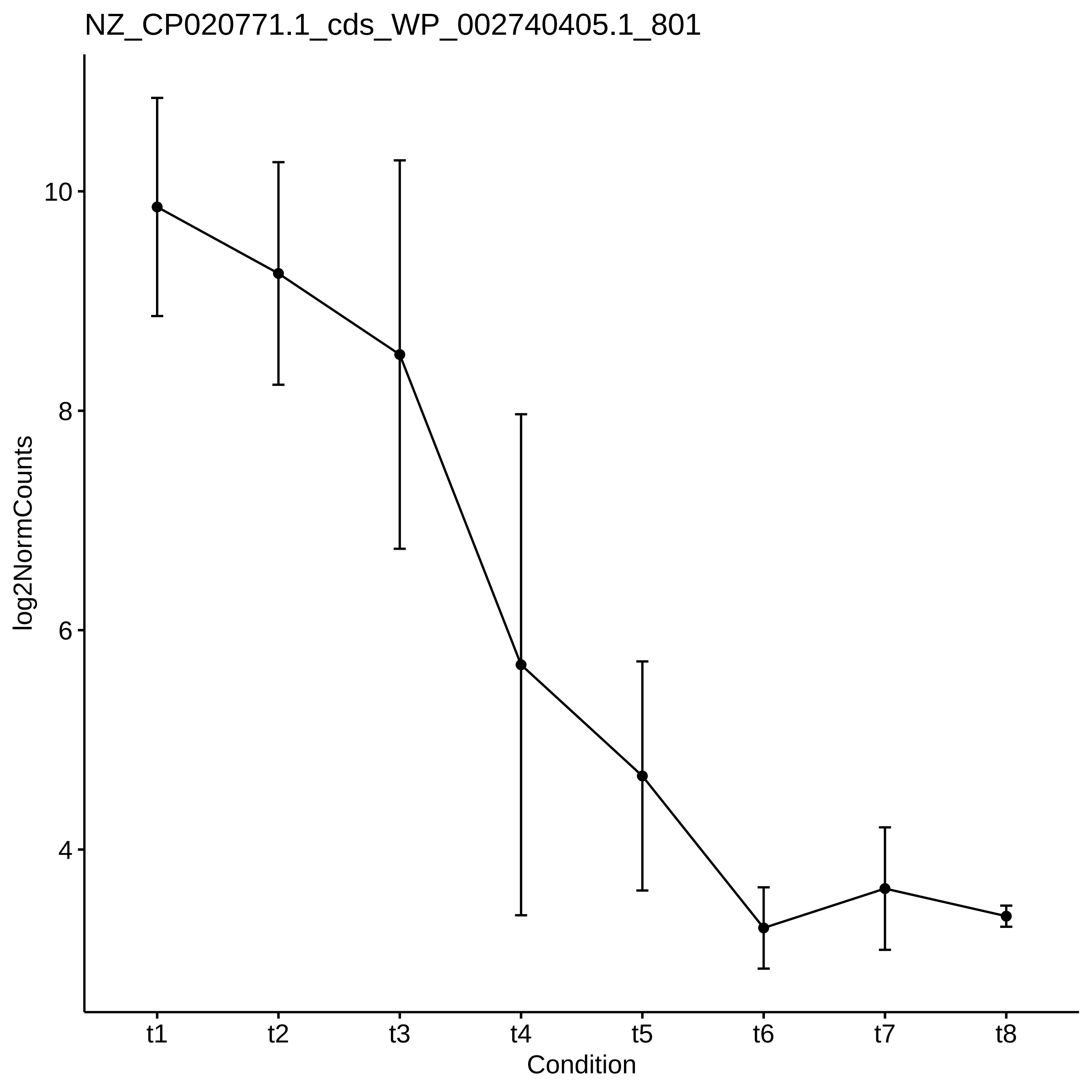

NZ_CP020771.1_cds_WP_002740405.1_801
log2NormCounts
10
8
6
4
t1
t2
t3
t4
t5
t6
t7
t8
Condition

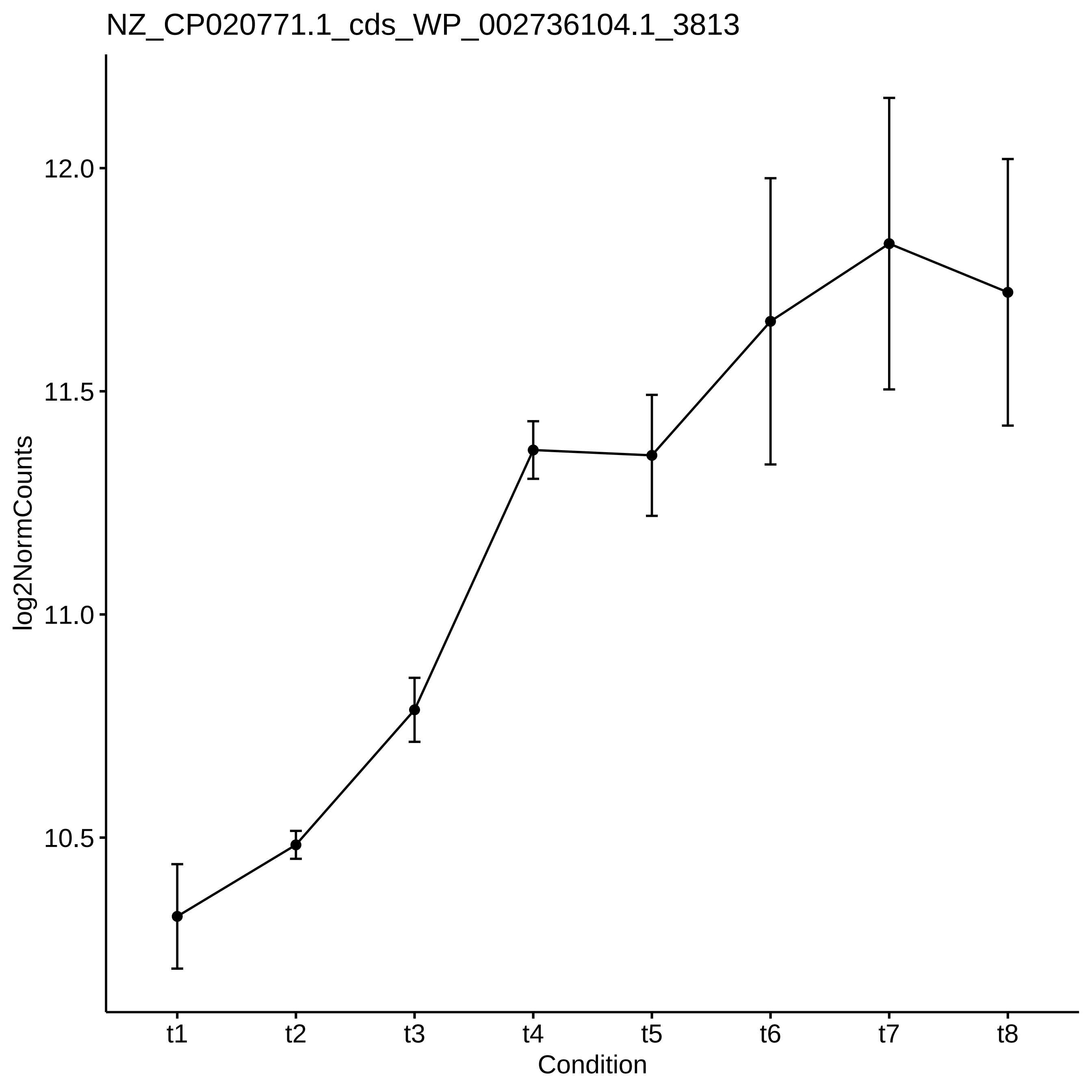
NZ_CP020771.1_cds_WP_002736104.1_3813
log2NormCounts
12.0
11.5
11.0
10.5
t1
t2
t3
t4
t5
t6
t7
t8
Condition

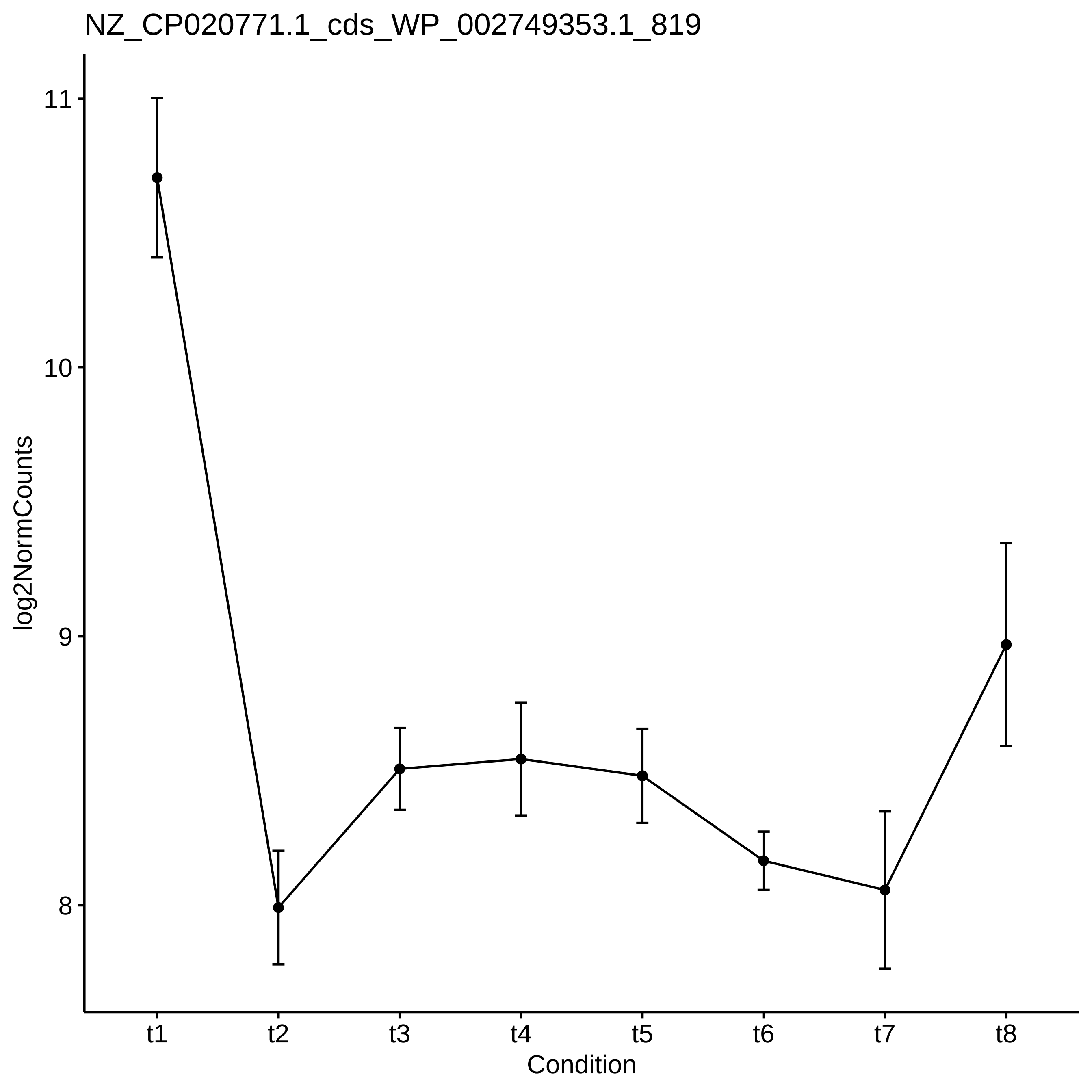
NZ_CP020771.1_cds_WP_002749353.1_819
log2NormCounts
11
10
9
8
t1
t2
t3
t4
t5
t6
t7
t8
Condition

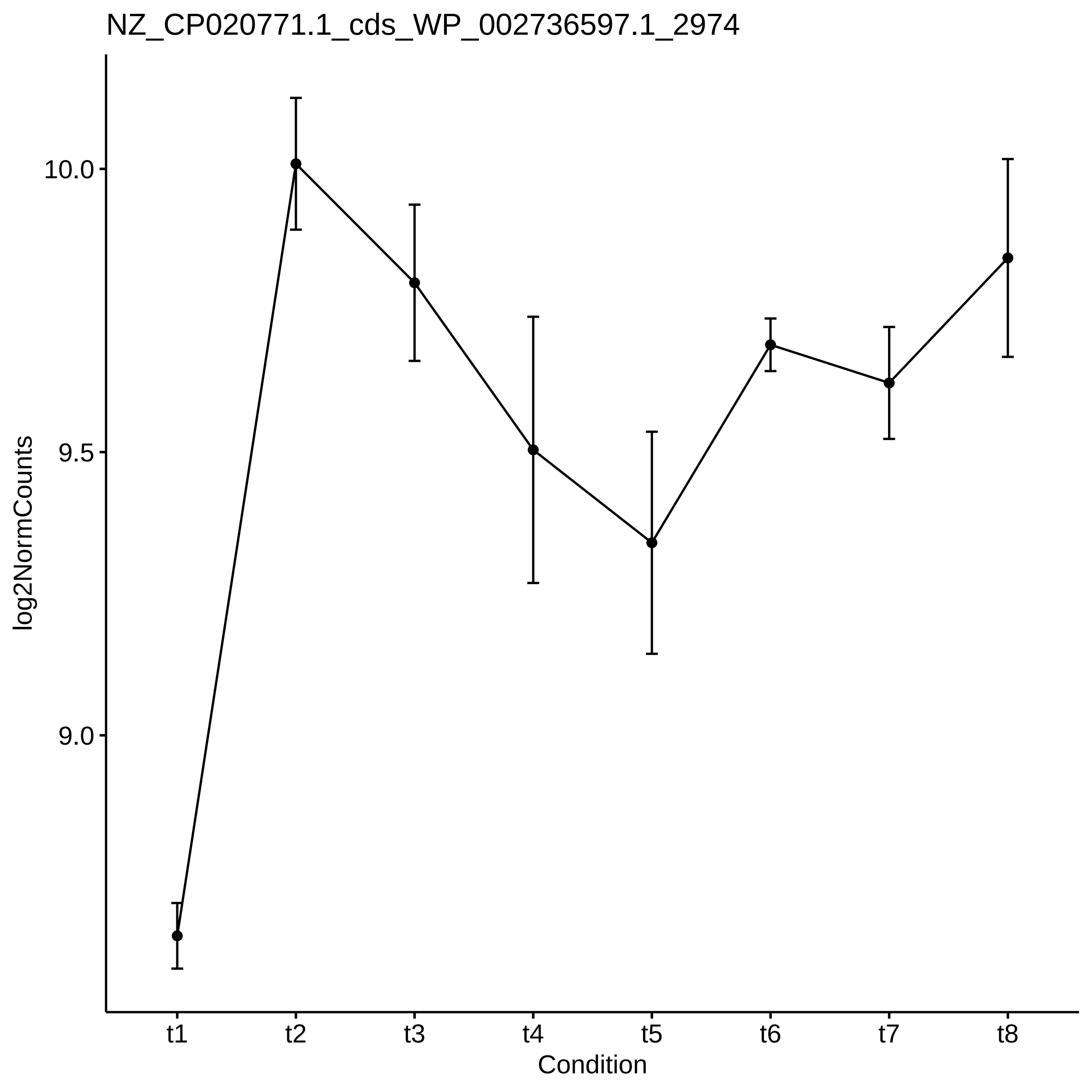

NZ_CP020771.1_cds_WP_002736597.1_2974
log2NormCounts
10.0
9.5
9.0
t1
t2
t3
t4
t5
t6
t7
t8
Condition

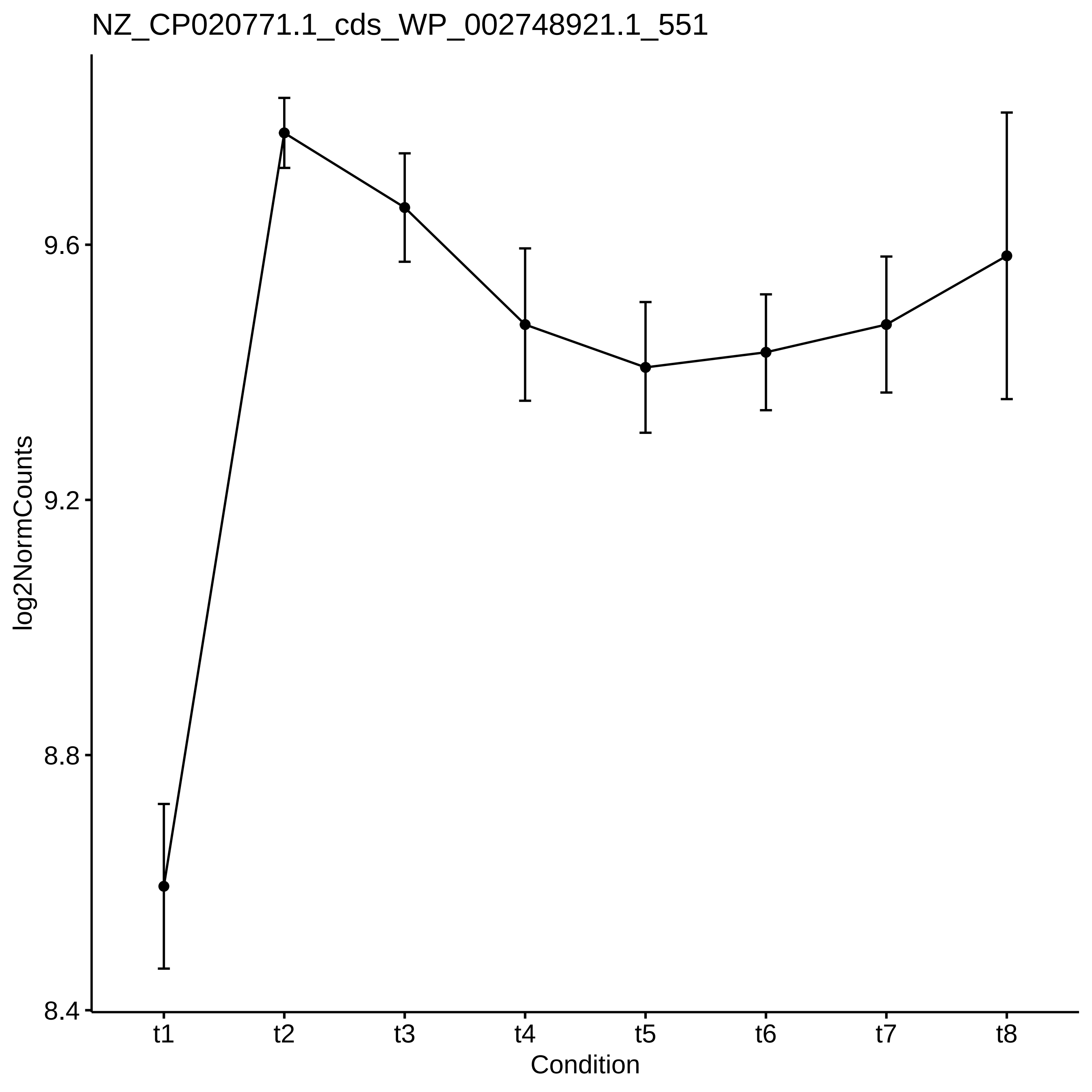

NZ_CP020771.1_cds_WP_002748921.1_551
log2NormCounts
9.6
9.2
8.8
8.4
t1
t2
t3
t4
t5
t6
t7
t8
Condition

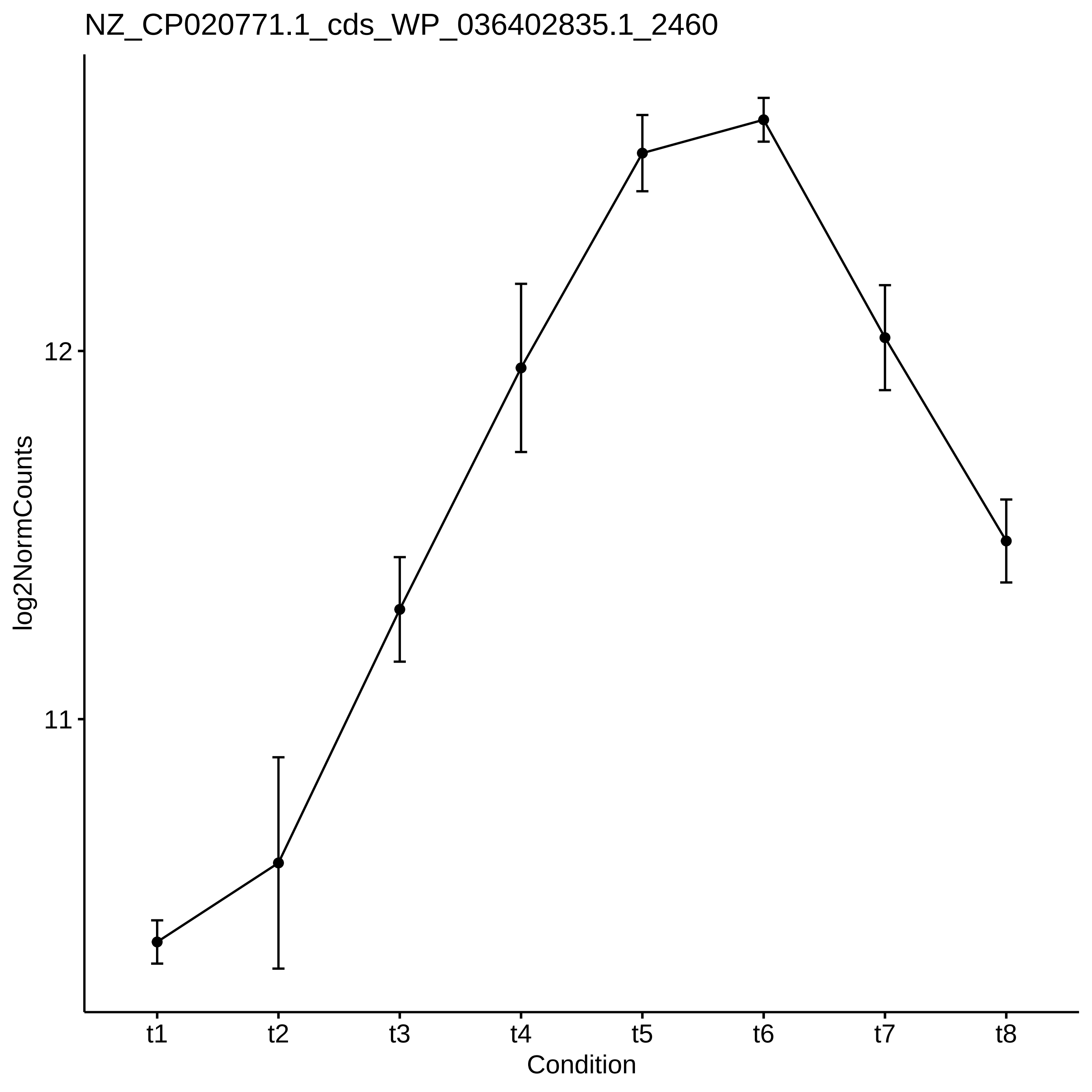

NZ_CP020771.1_cds_WP_036402835.1_2460
log2NormCounts
12
11
t1
t2
t3
t4
t5
t6
t7
t8
Condition

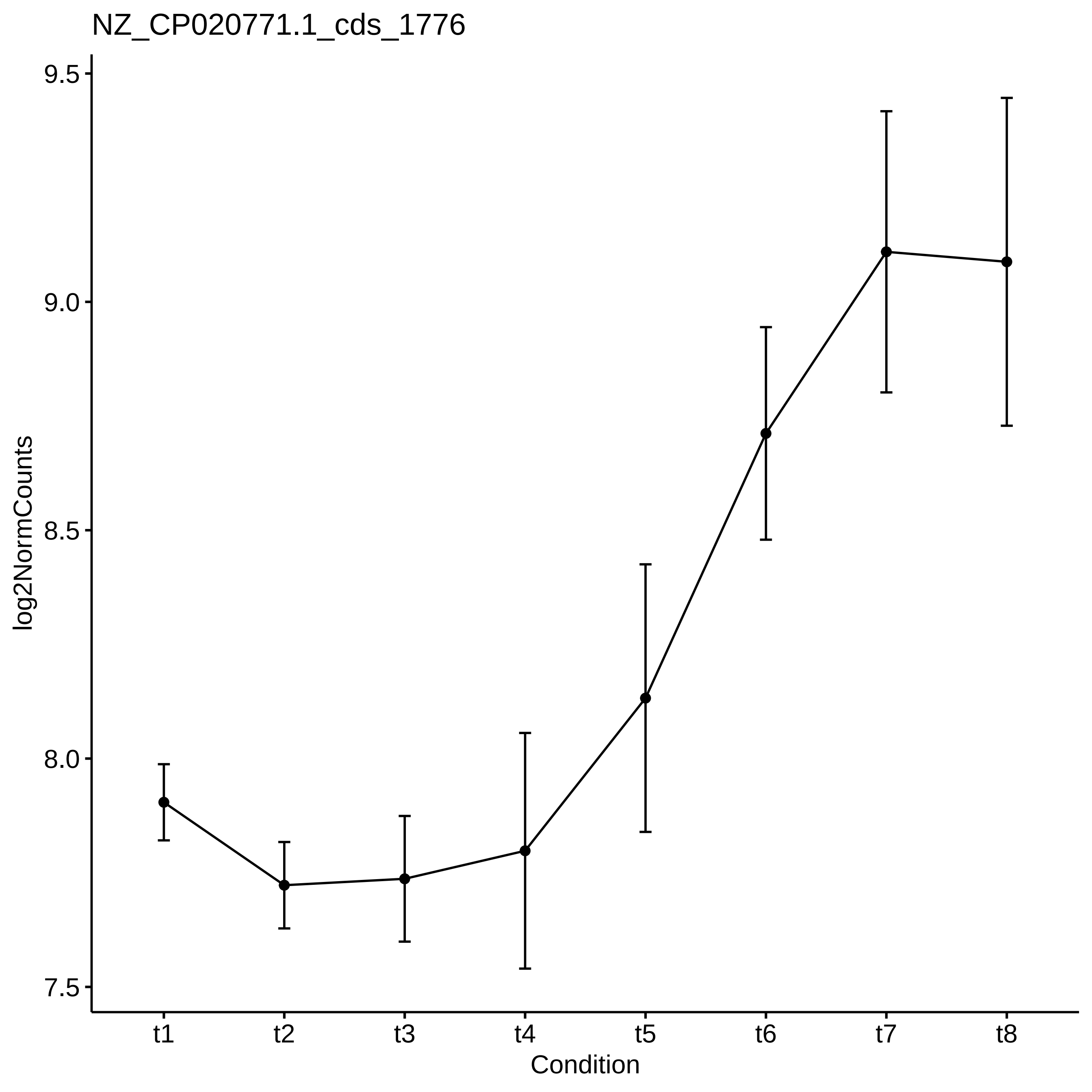
NZ_CP020771.1_cds_1776
log2NormCounts
9.5
9.0
8.5
8.0
7.5
t1
t2
t3
t4
t5
t6
t7
t8
Condition

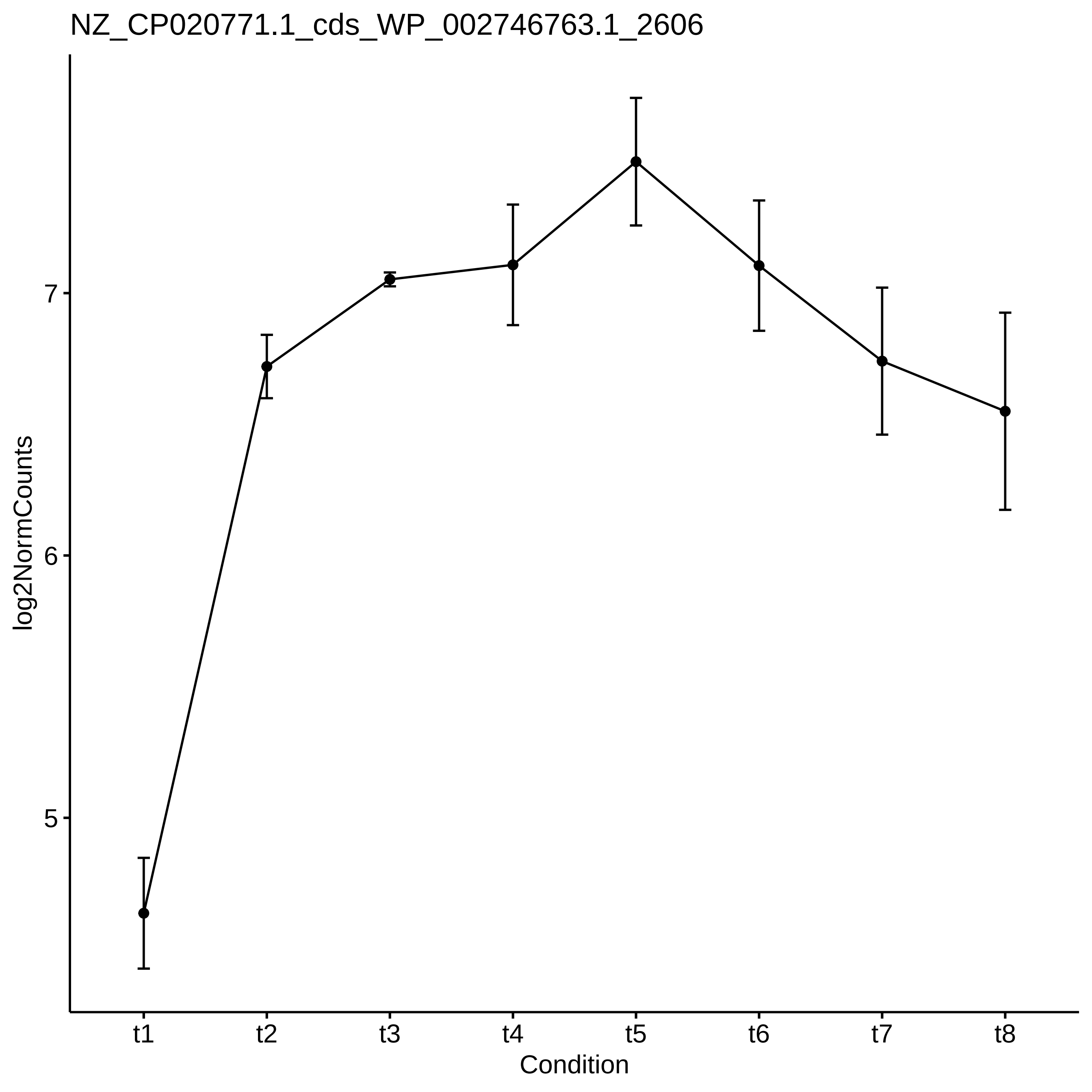
NZ_CP020771.1_cds_WP_002746763.1_2606
log2NormCounts
7
6
5
t1
t2
t3
t4
t5
t6
t7
t8
Condition

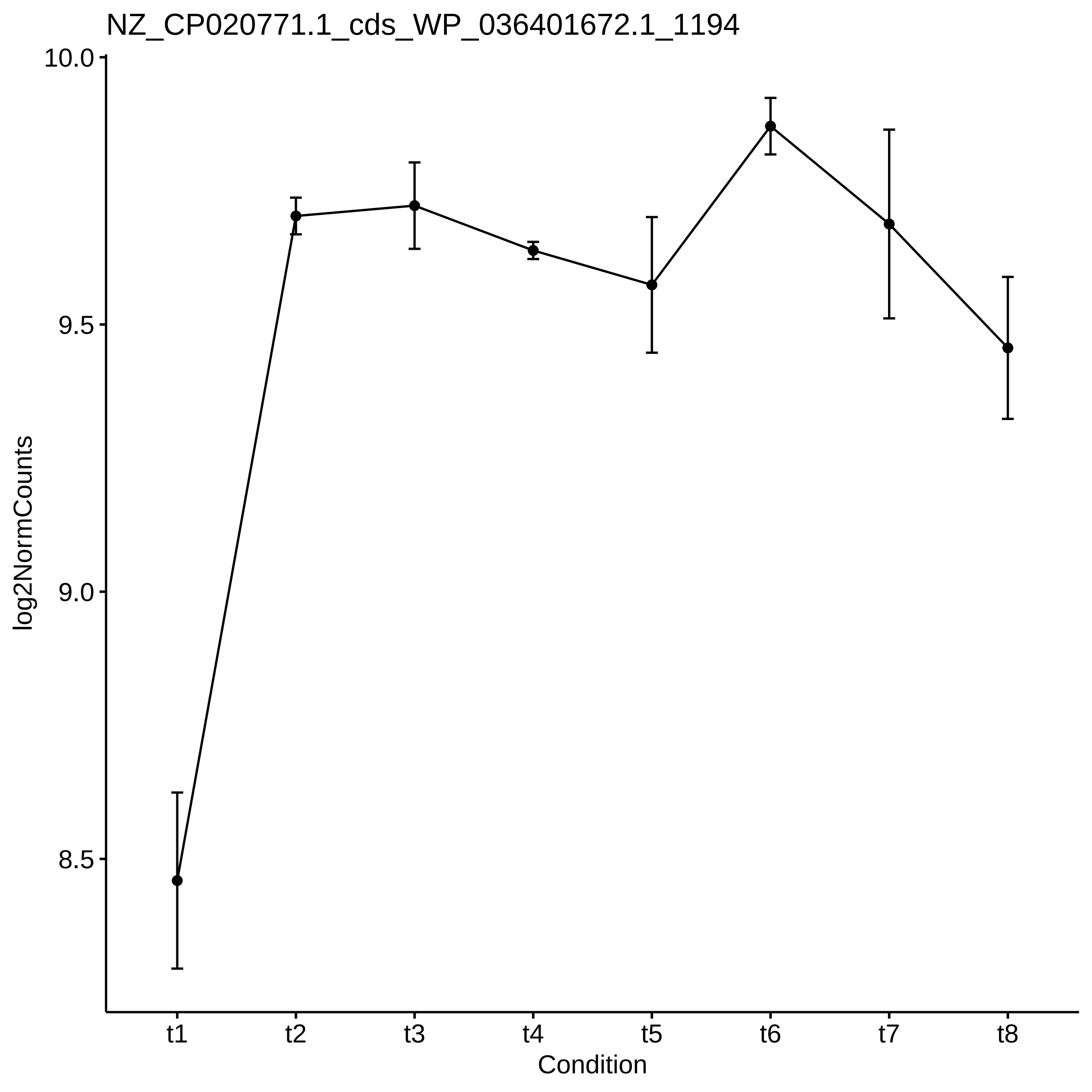
NZ_CP020771.1_cds_WP_036401672.1_1194
10.0
9.5
9.0
8.5
log2NormCounts
t1
t2
t3
t4
t5
t6
t7
t8
Condition

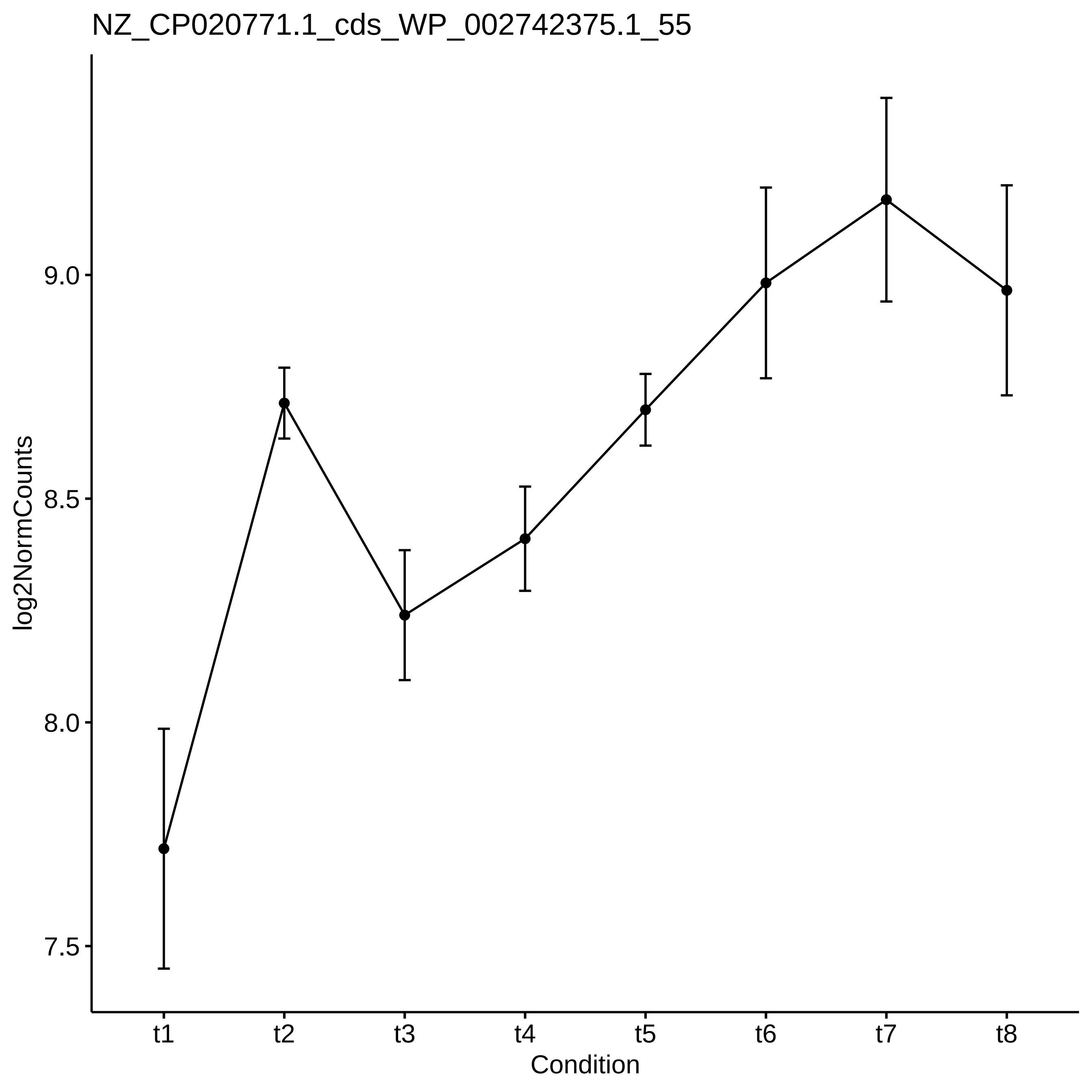

NZ_CP020771.1_cds_WP_002742375.1_55
log2NormCounts
9.0
8.5
8.0
7.5
t1
t2
t3
t4
t5
t6
t7
t8
Condition

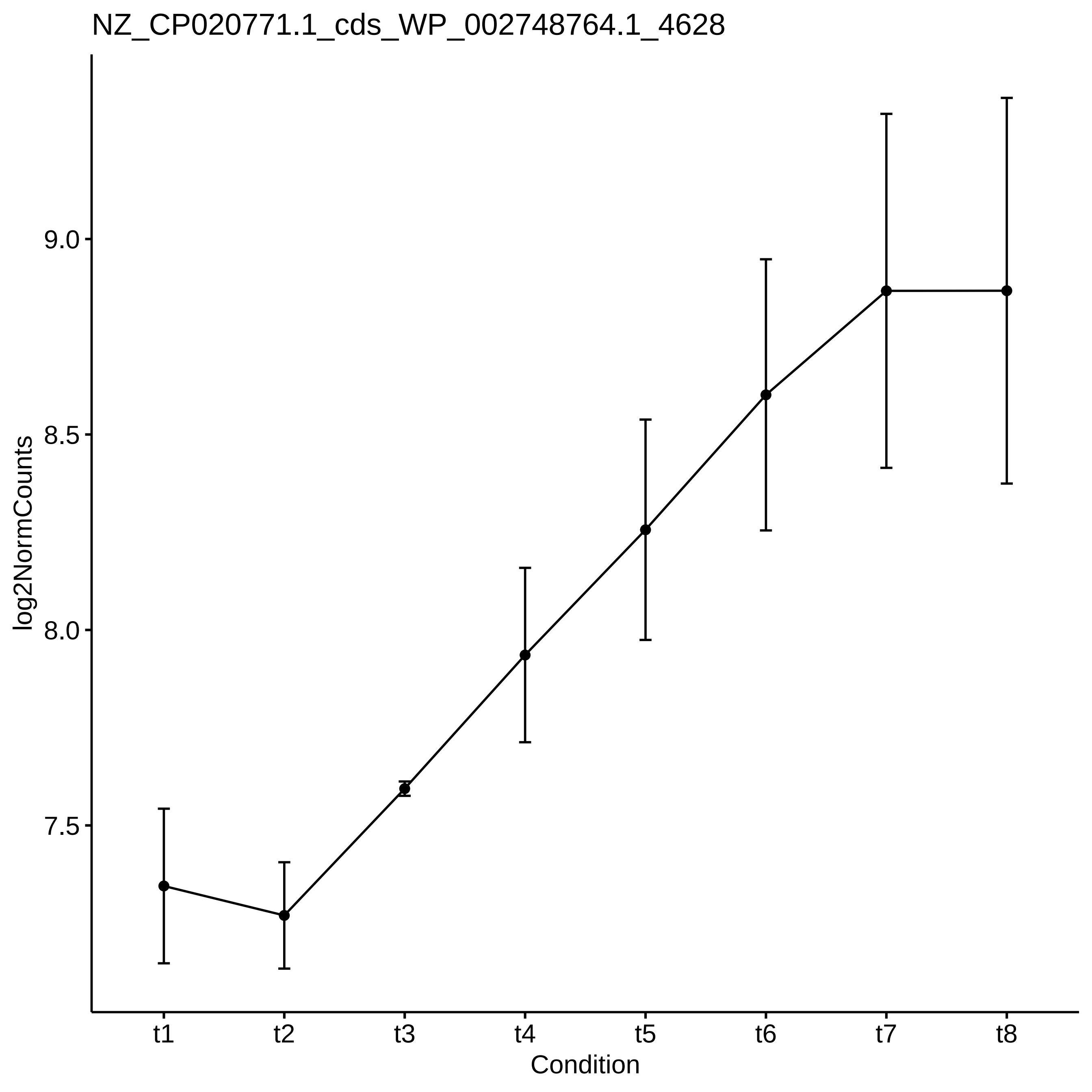
NZ_CP020771.1_cds_WP_002748764.1_4628
log2NormCounts
9.0
8.5
8.0
7.5
t1
t2
t3
t4
t5
t6
t7
t8
Condition

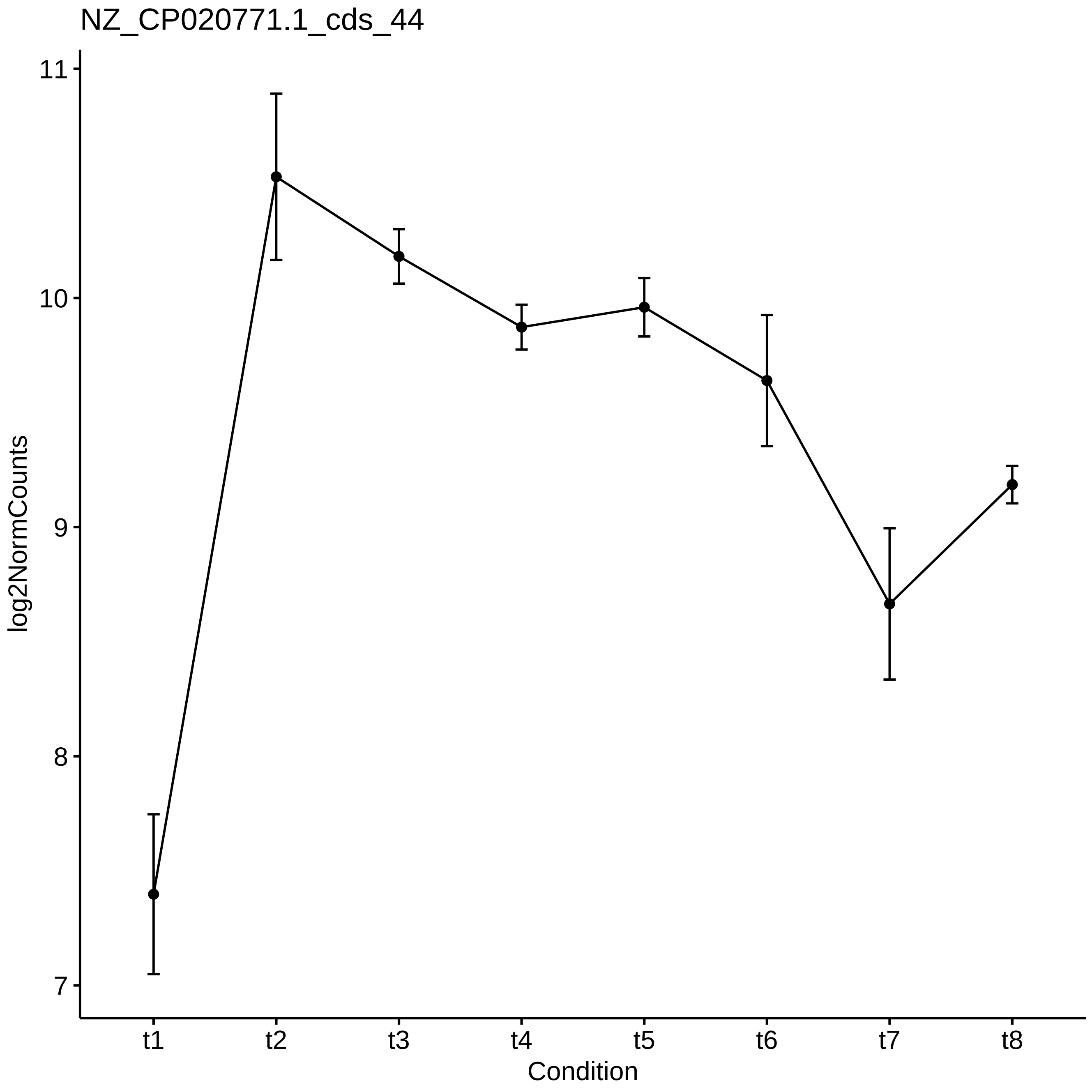

NZ_CP020771.1_cds_44
log2NormCounts
11
10
9
8
7
t1
t2
t3
t4
t5
t6
t7
t8
Condition

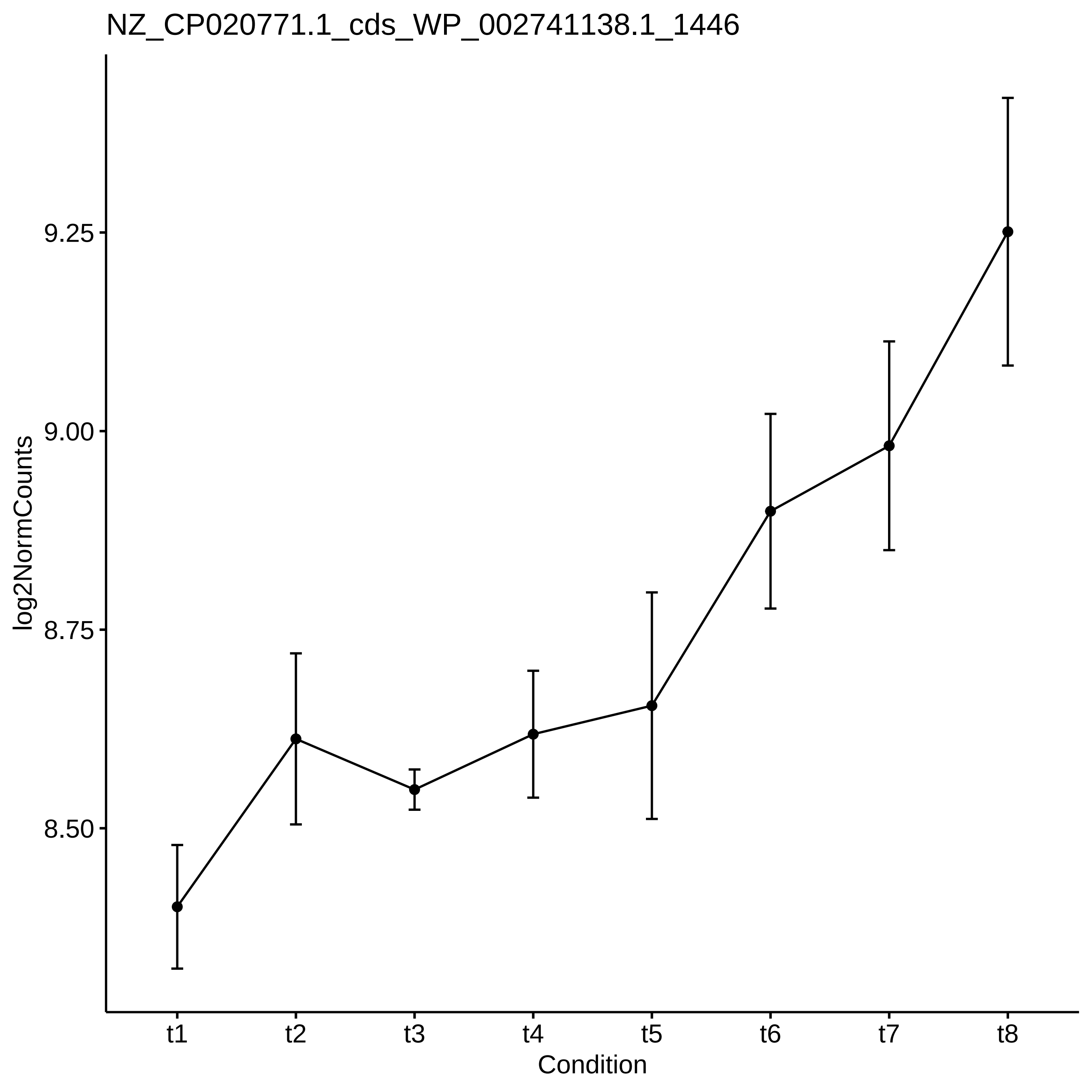
NZ_CP020771.1_cds_WP_002741138.1_1446
log2NormCounts
9.25
9.00
8.75
8.50
t1
t2
t3
t4
t5
t6
t7
t8
Condition

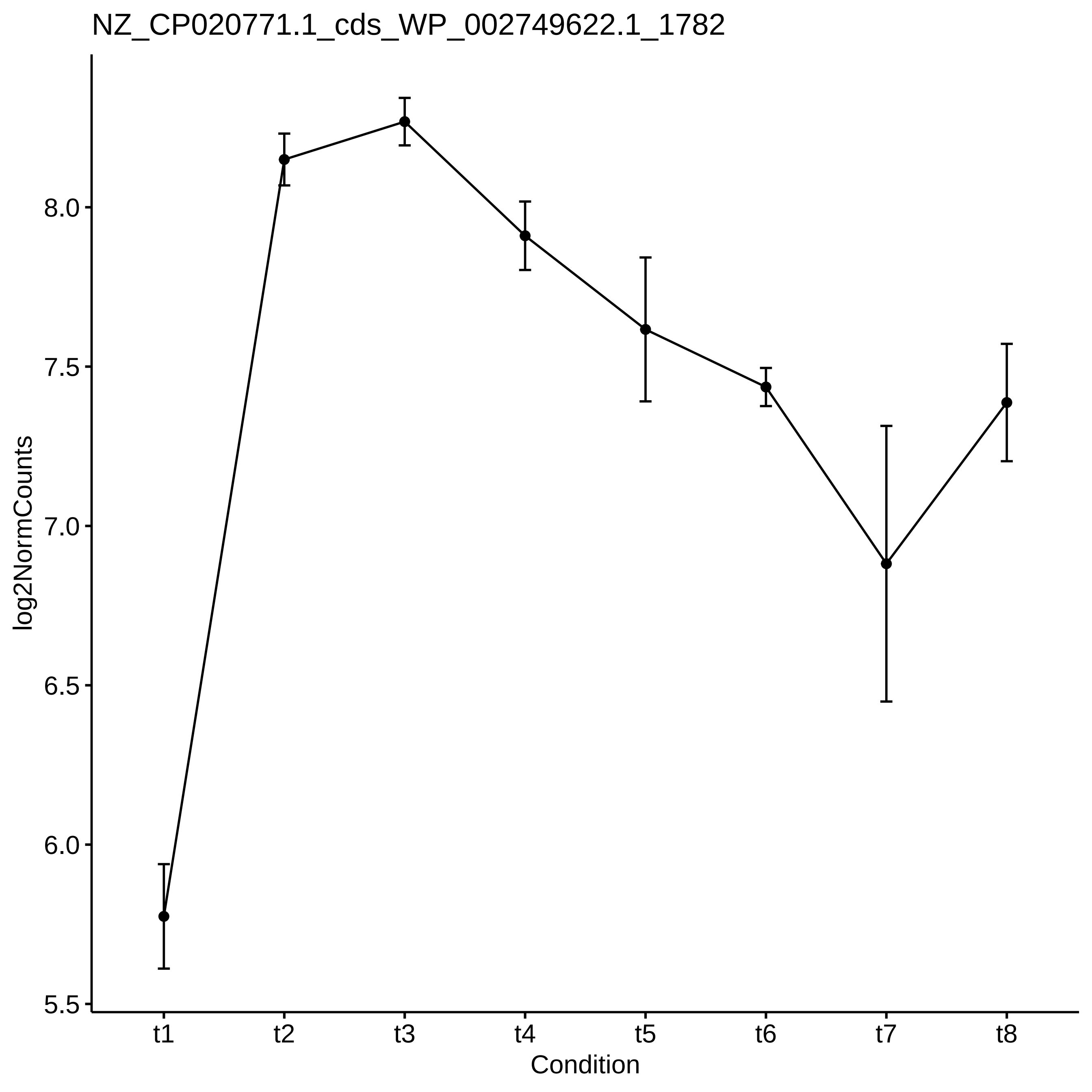

NZ_CP020771.1_cds_WP_002749622.1_1782
log2NormCounts
8.0
7.5
7.0
6.5
6.0
5.5
t1
t2
t3
t4
t5
t6
t7
t8
Condition

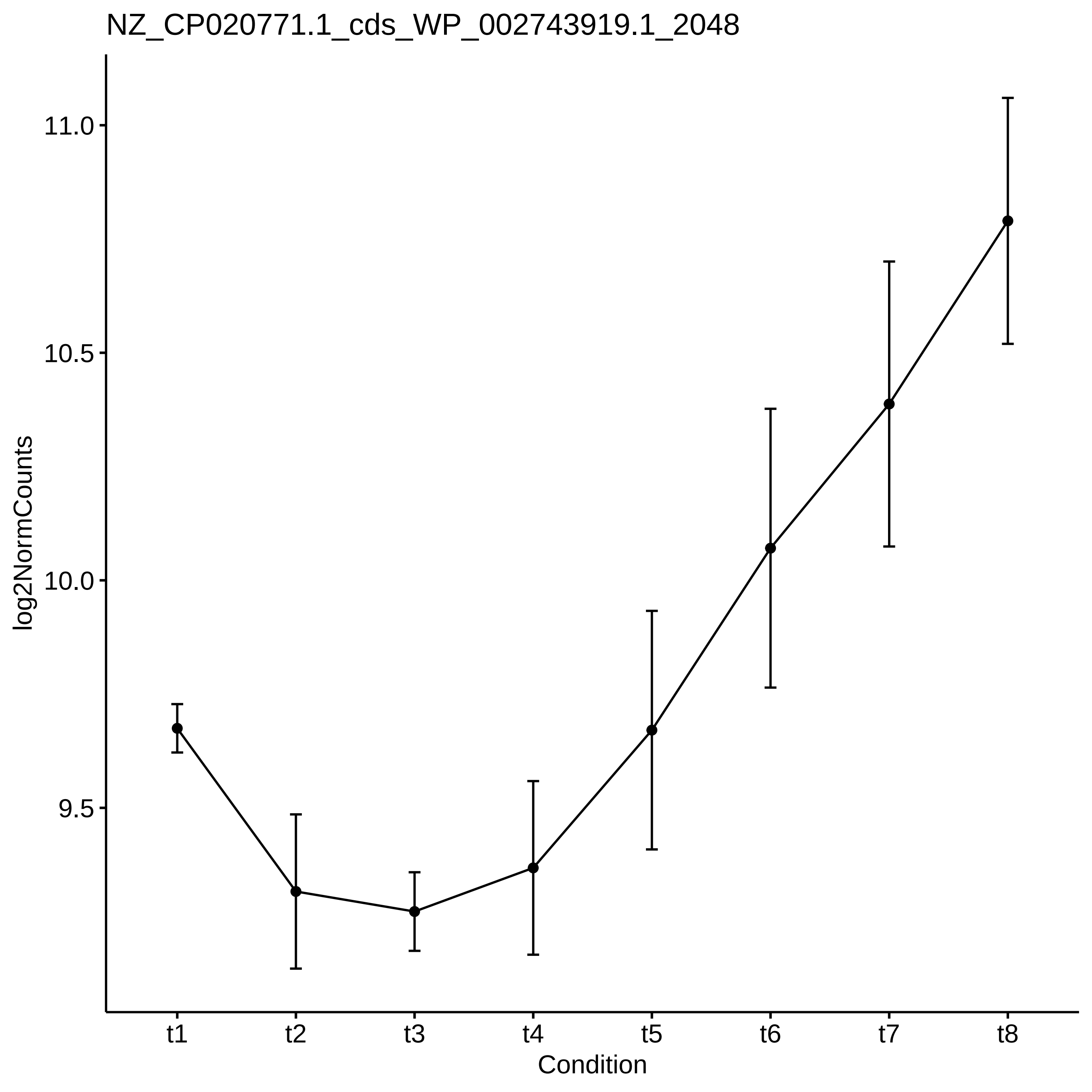

NZ_CP020771.1_cds_WP_002743919.1_2048
log2NormCounts
11.0
10.5
10.0
9.5
t1
t2
t3
t4
t5
t6
t7
t8
Condition

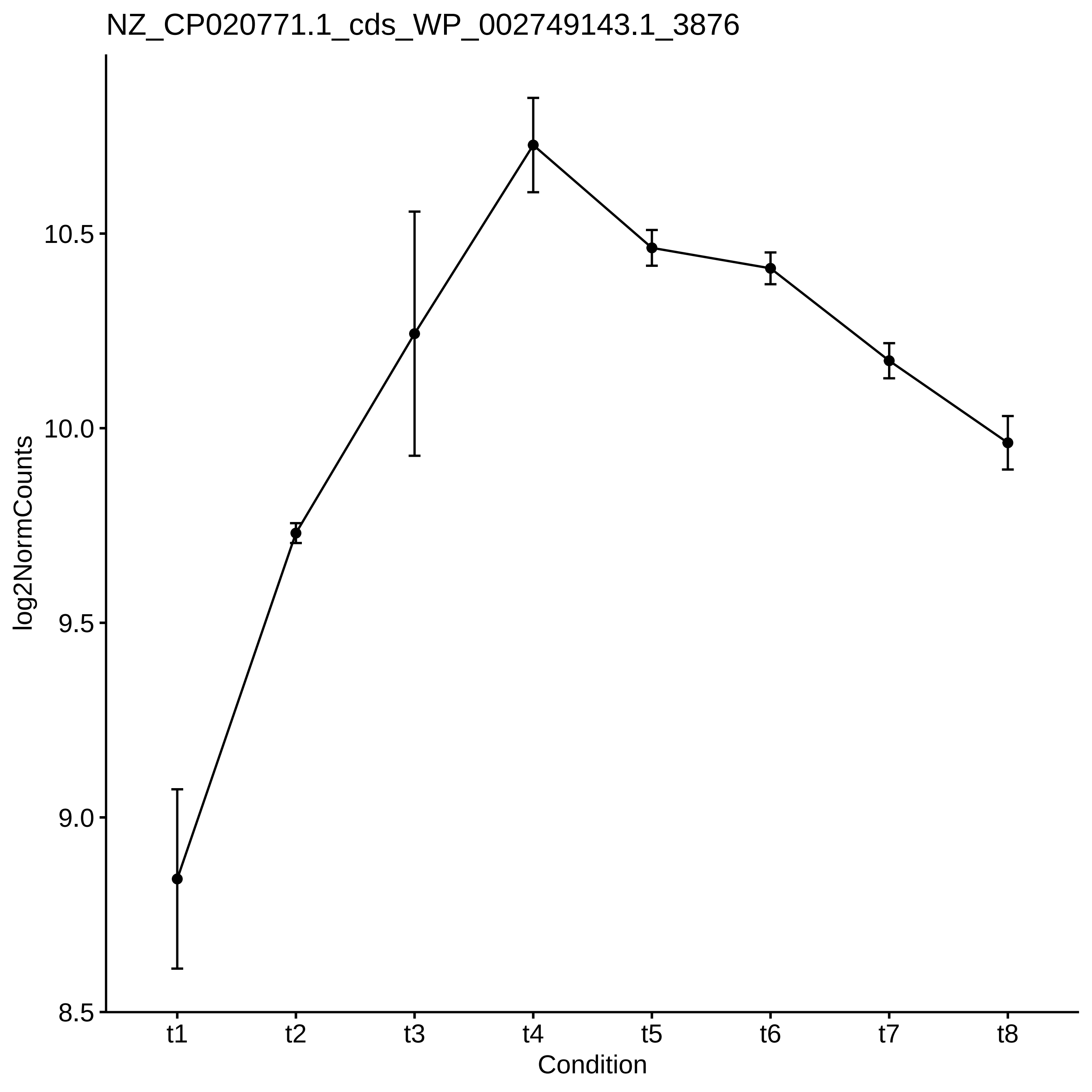

NZ_CP020771.1_cds_WP_002749143.1_3876
log2NormCounts
10.5
10.0
9.5
9.0
8.5
t1
t2
t3
t4
t5
t6
t7
t8
Condition

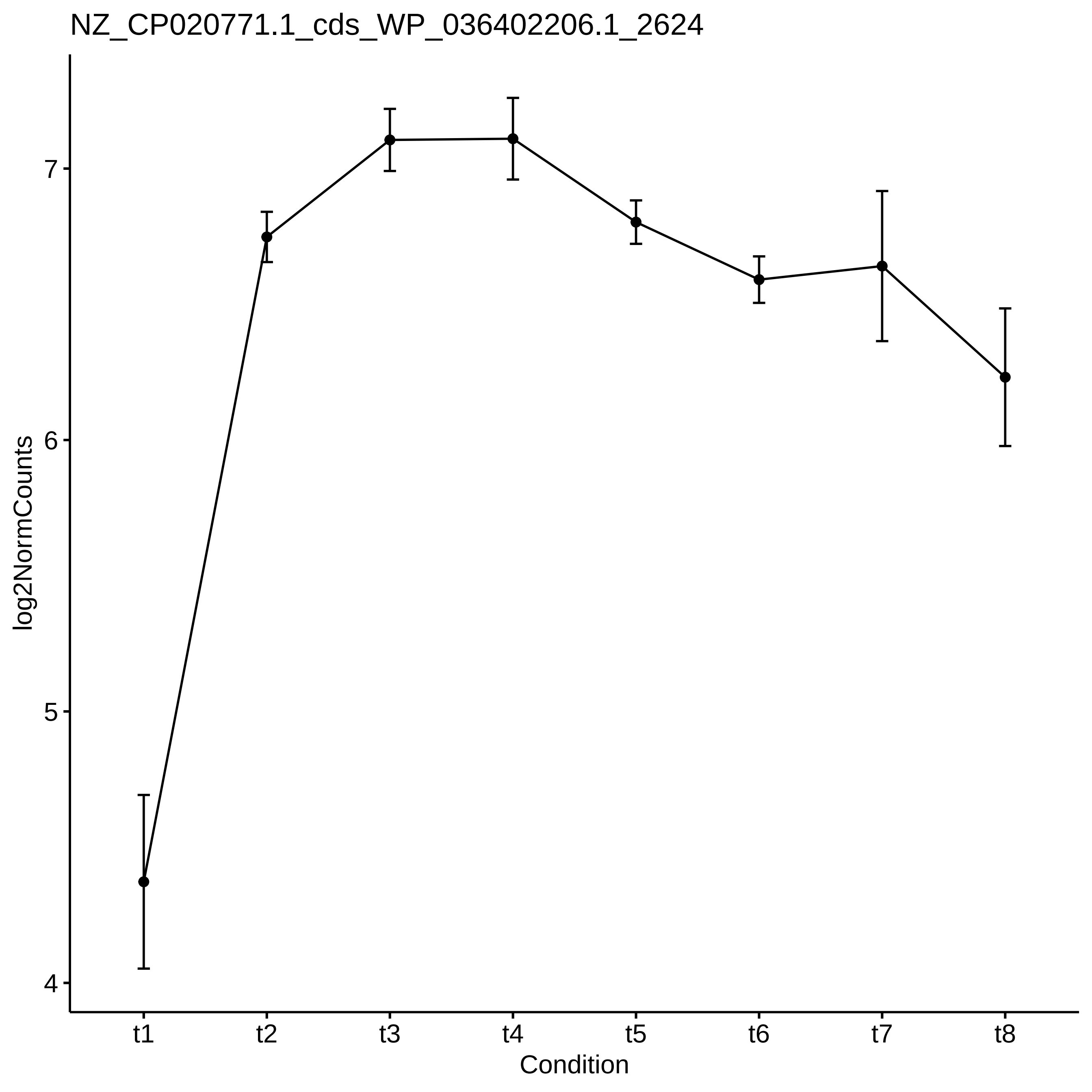

NZ_CP020771.1_cds_WP_036402206.1_2624
log2NormCounts
7
6
5
4
t1
t2
t3
t4
t5
t6
t7
t8
Condition

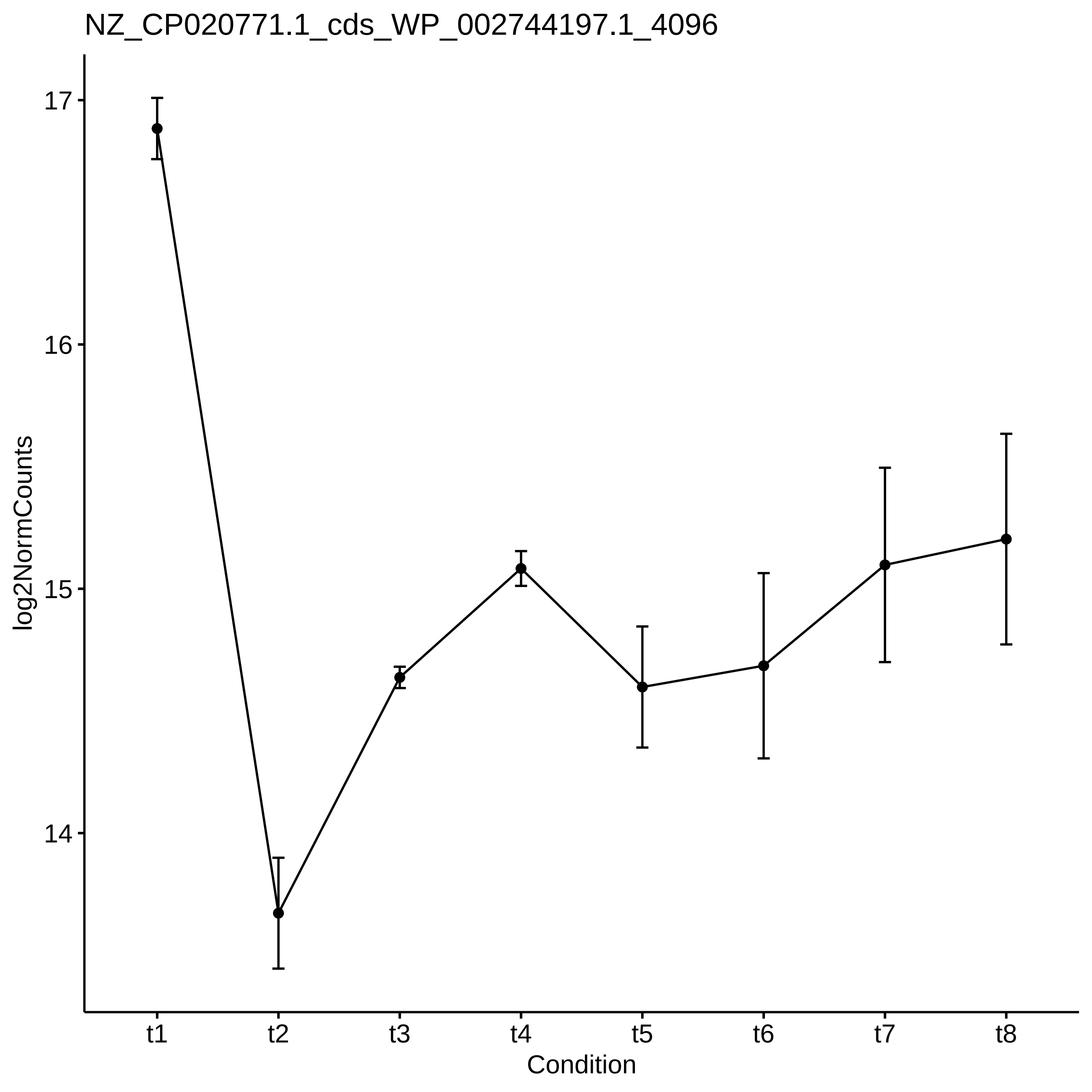

NZ_CP020771.1_cds_WP_002744197.1_4096
log2NormCounts
17
16
15
14
t1
t2
t3
t4
t5
t6
t7
t8
Condition

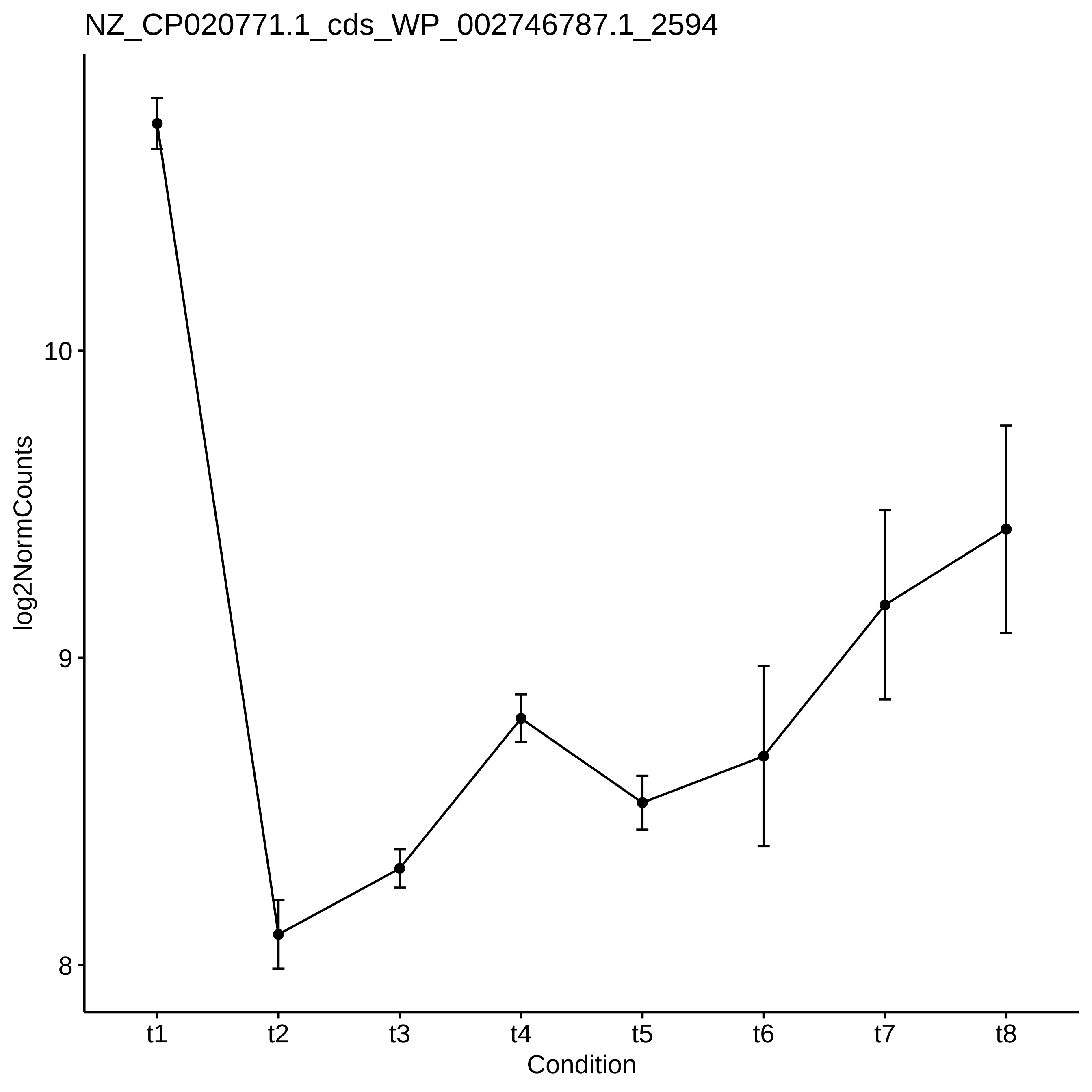
NZ_CP020771.1_cds_WP_002746787.1_2594
log2NormCounts
10
9
8
t1
t2
t3
t4
t5
t6
t7
t8
Condition

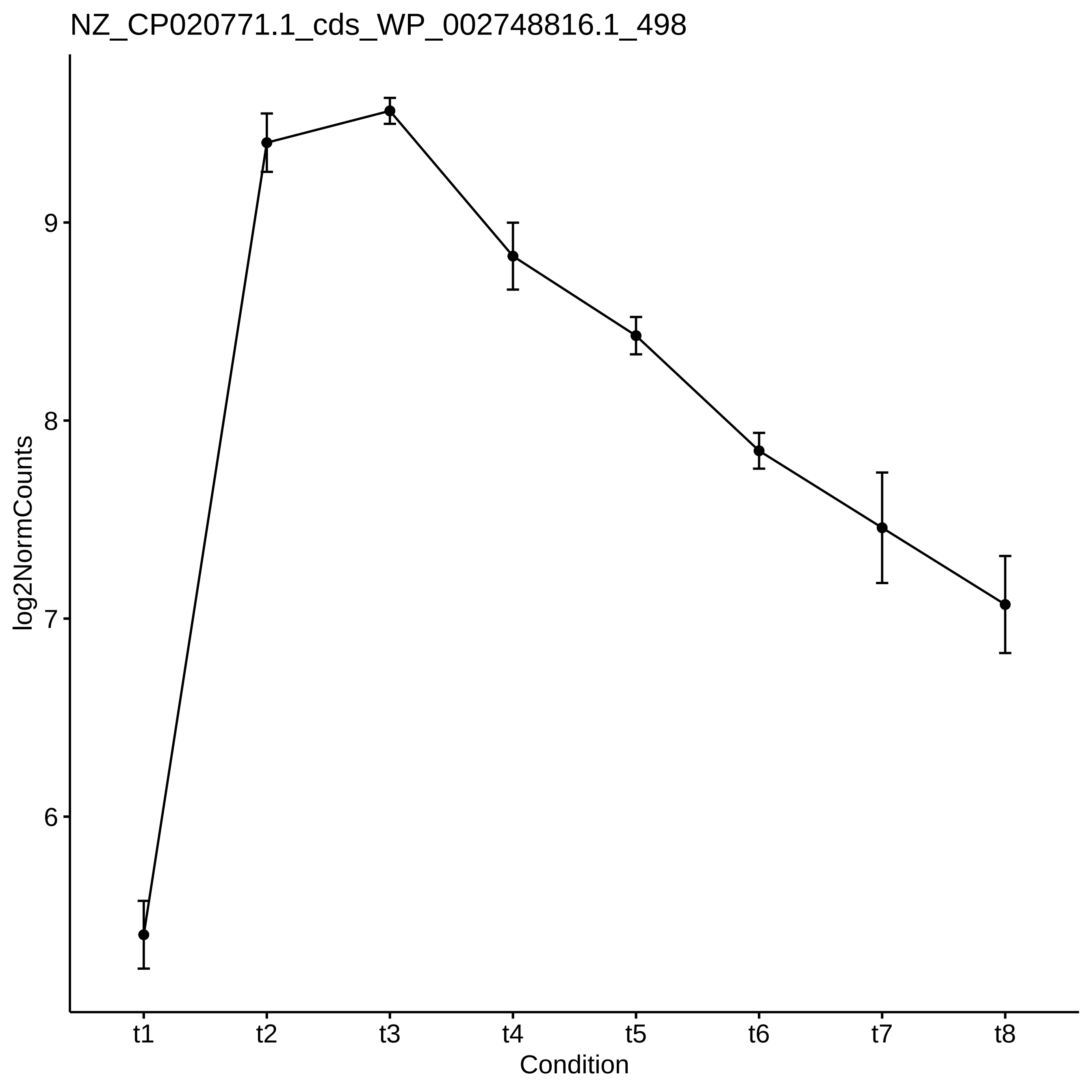

NZ_CP020771.1_cds_WP_002748816.1_498
log2NormCounts
9
8
7
6
t1
t2
t3
t4
t5
t6
t7
t8
Condition

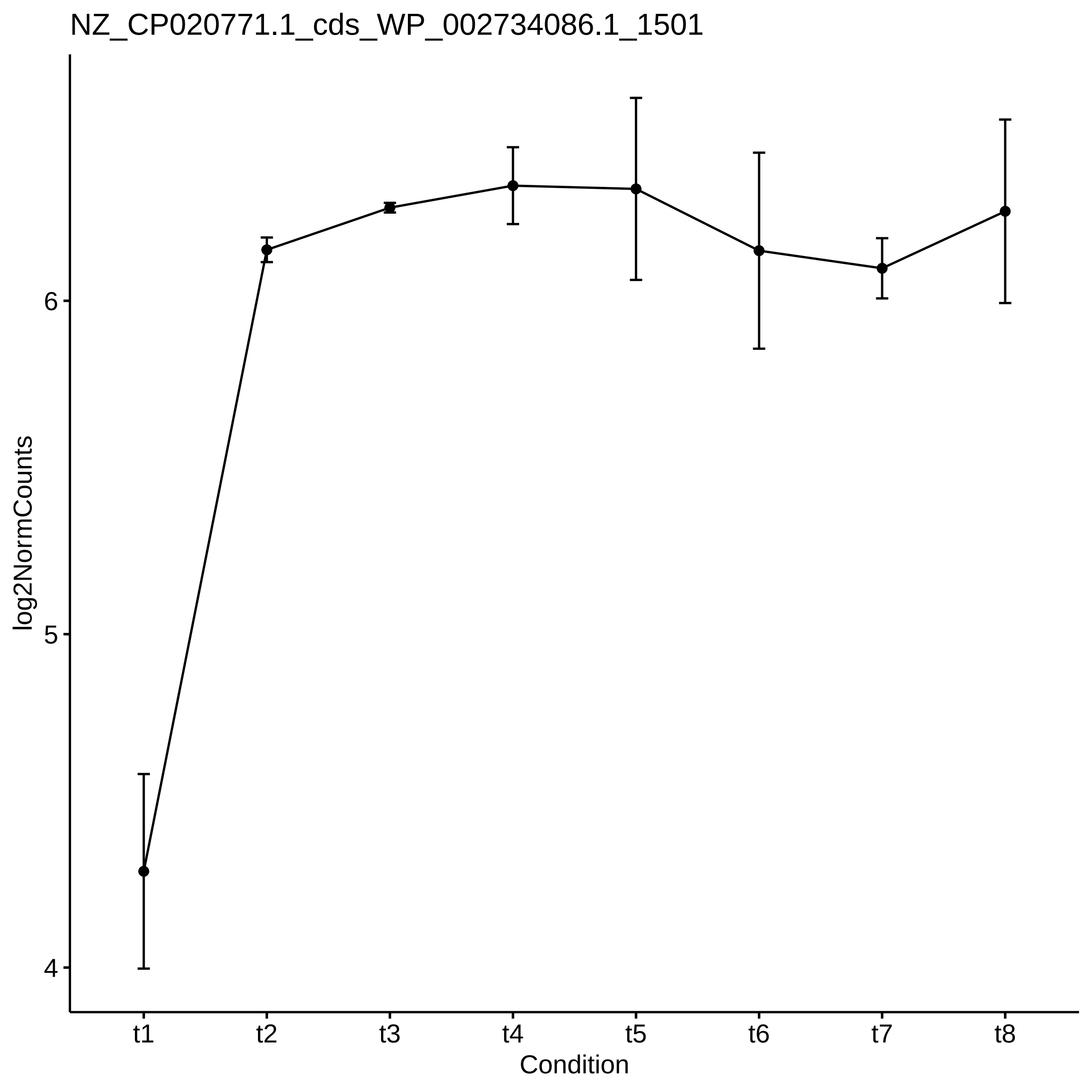

NZ_CP020771.1_cds_WP_002734086.1_1501
log2NormCounts
6
5
4
t1
t2
t3
t4
t5
t6
t7
t8
Condition

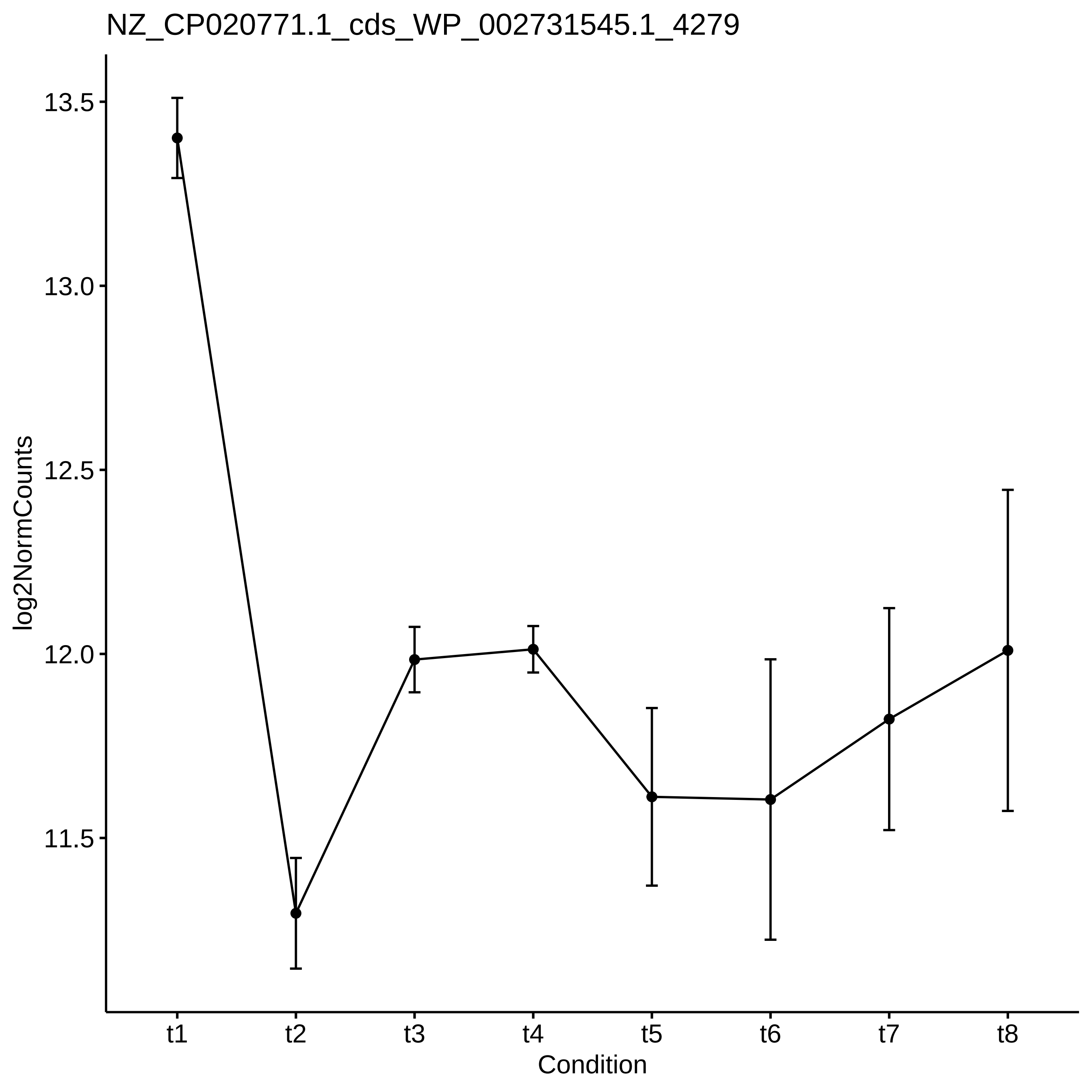
NZ_CP020771.1_cds_WP_002731545.1_4279
log2NormCounts
13.5
13.0
12.5
12.0
11.5
t1
t2
t3
t4
t5
t6
t7
t8
Condition

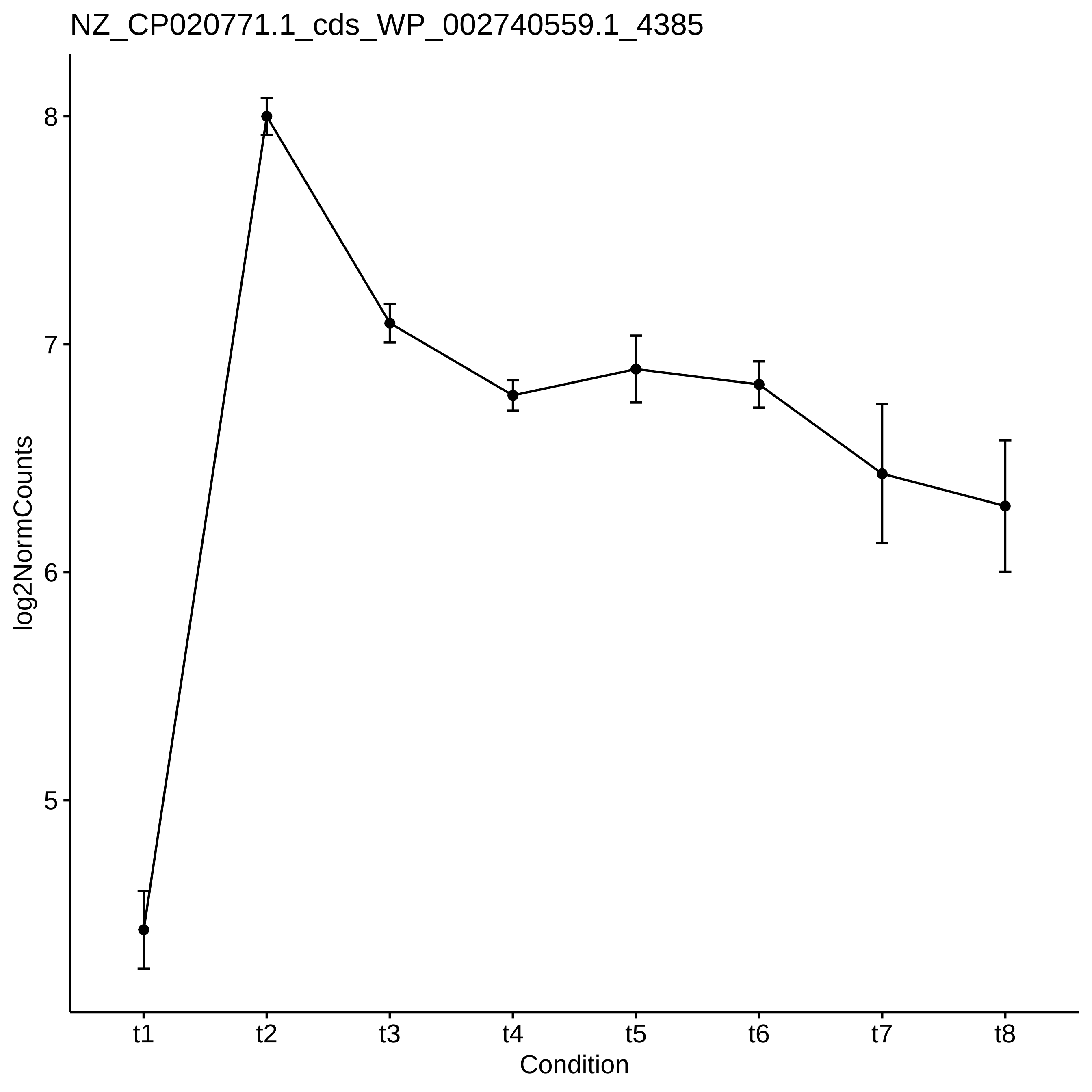

NZ_CP020771.1_cds_WP_002740559.1_4385
log2NormCounts
8
7
6
5
t1
t2
t3
t4
t5
t6
t7
t8
Condition

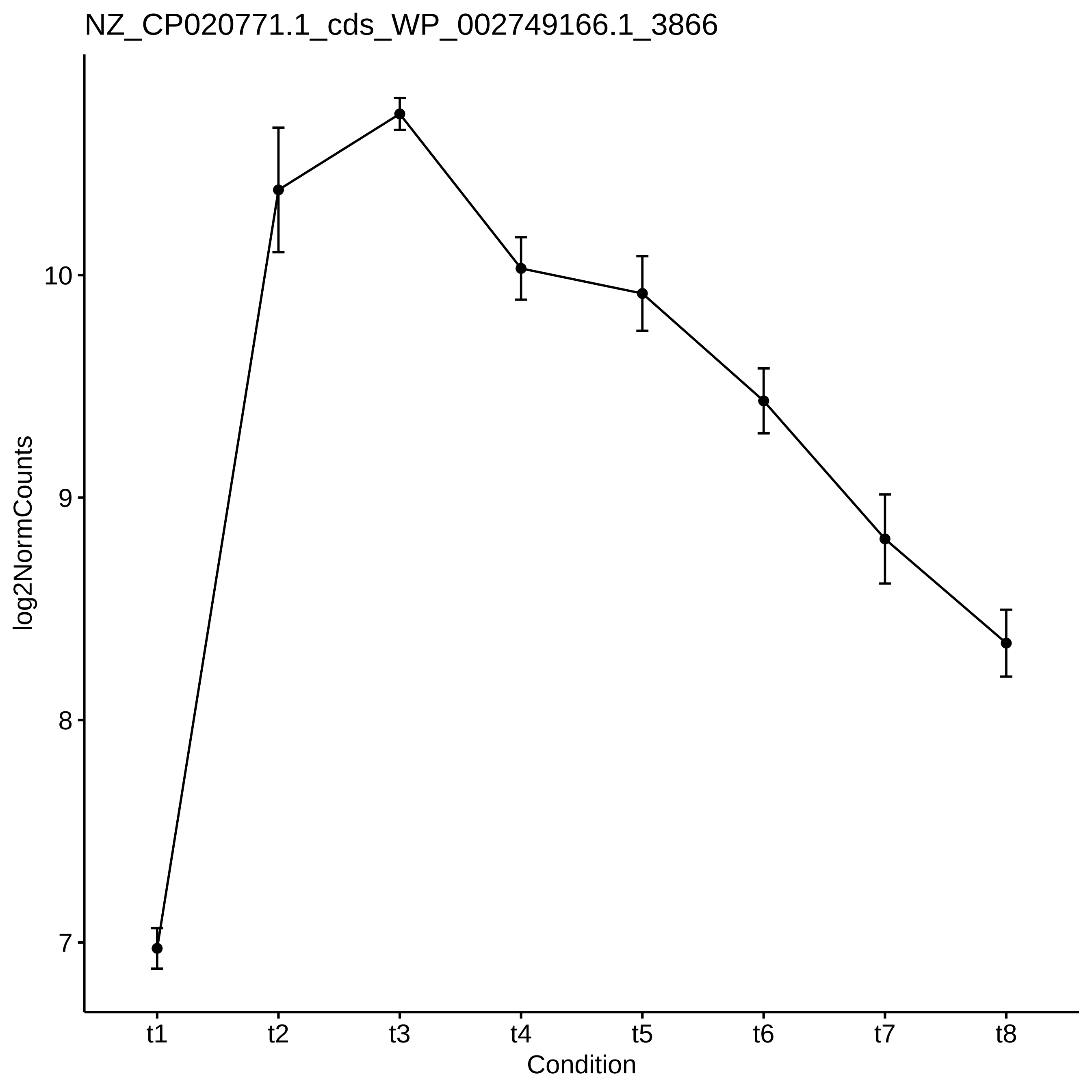
NZ_CP020771.1_cds_WP_002749166.1_3866
log2NormCounts
10
9
8
7
t1
t2
t3
t4
t5
t6
t7
t8
Condition

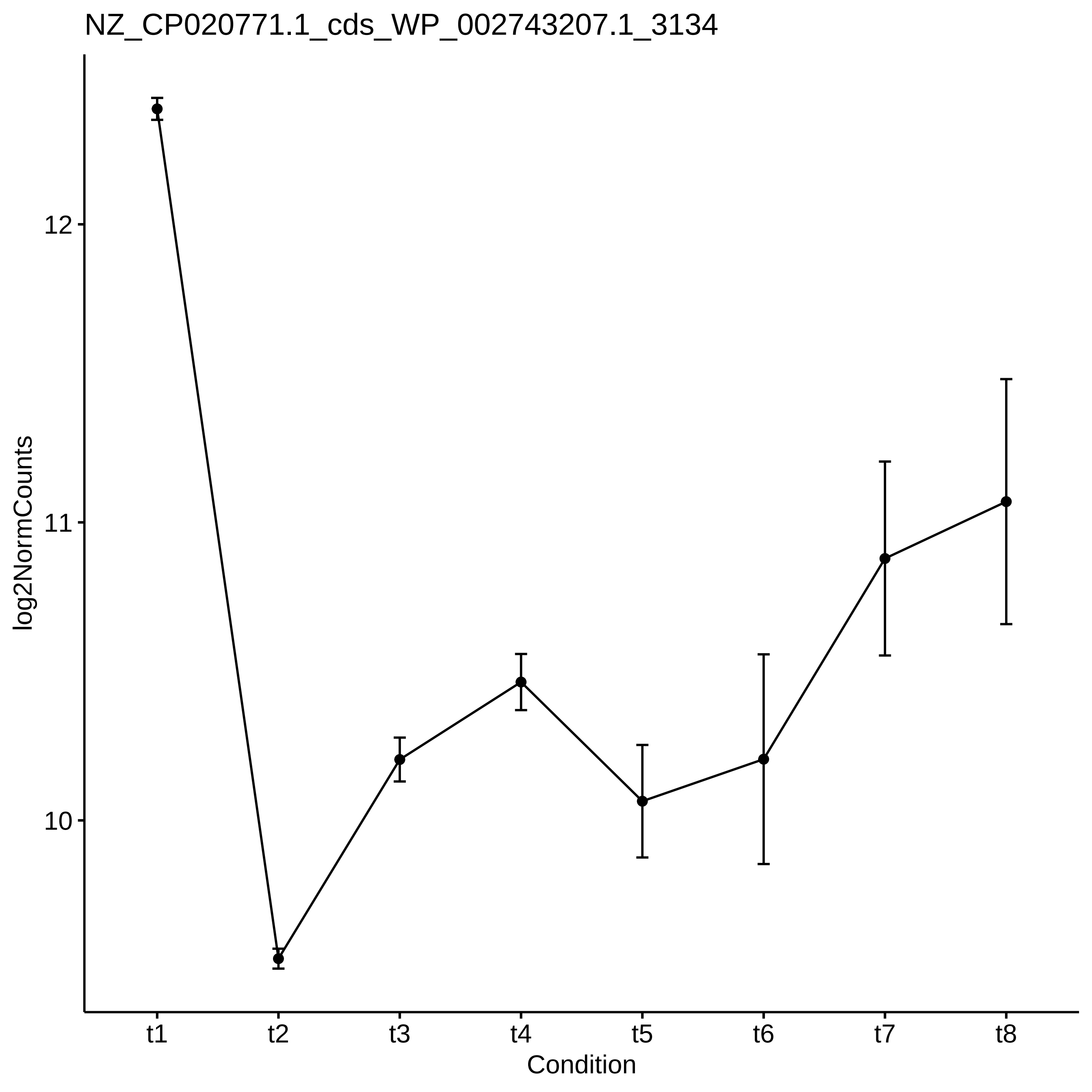

NZ_CP020771.1_cds_WP_002743207.1_3134
log2NormCounts
12
11
10
t1
t2
t3
t4
t5
t6
t7
t8
Condition

NZ_CP020771.1_cds_WP_002738148.1_544

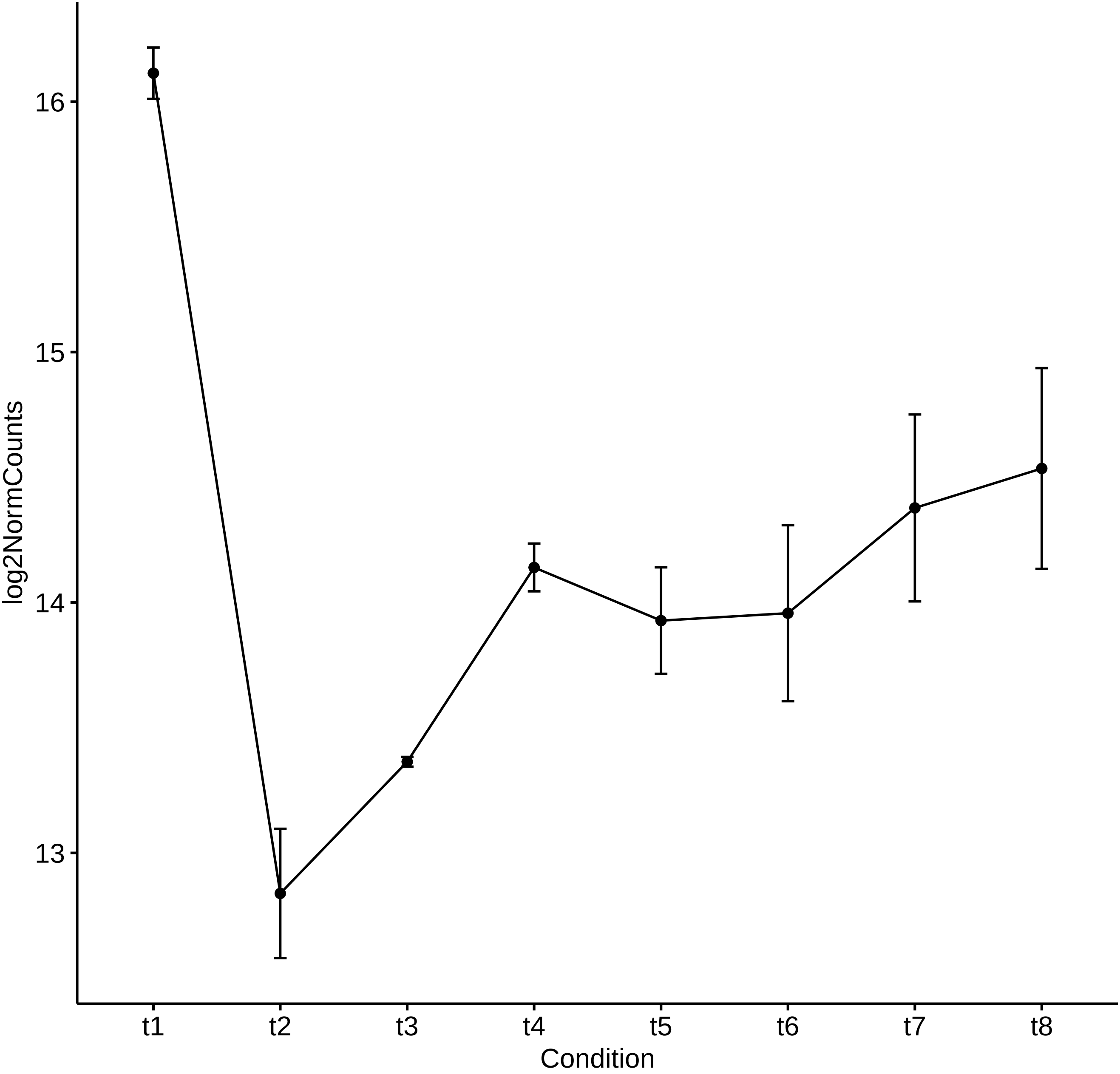

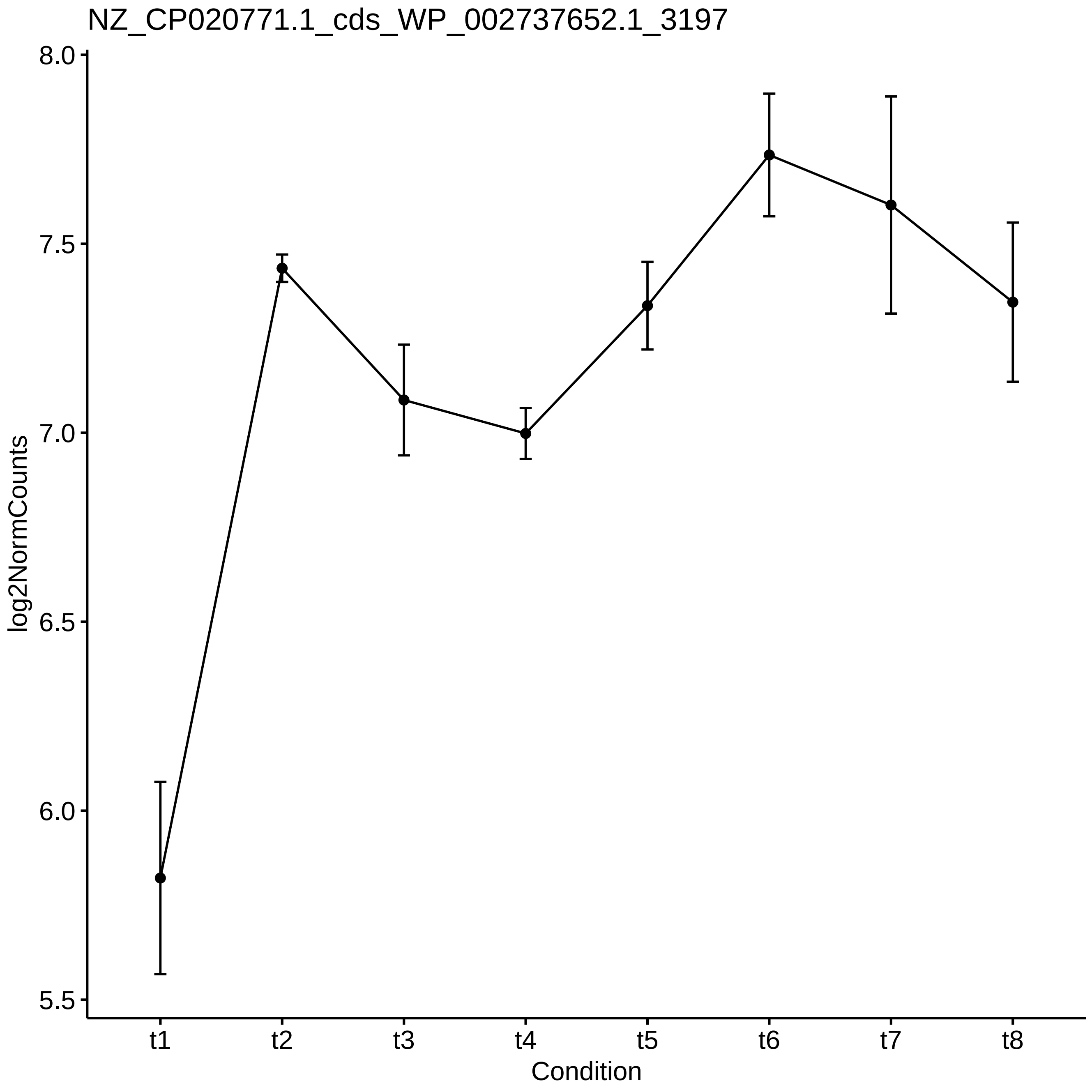

NZ_CP020771.1_cds_WP_002737652.1_3197
log2NormCounts
8.0
7.5
7.0
6.5
6.0
5.5
t1
t2
t3
t4
t5
t6
t7
t8
Condition

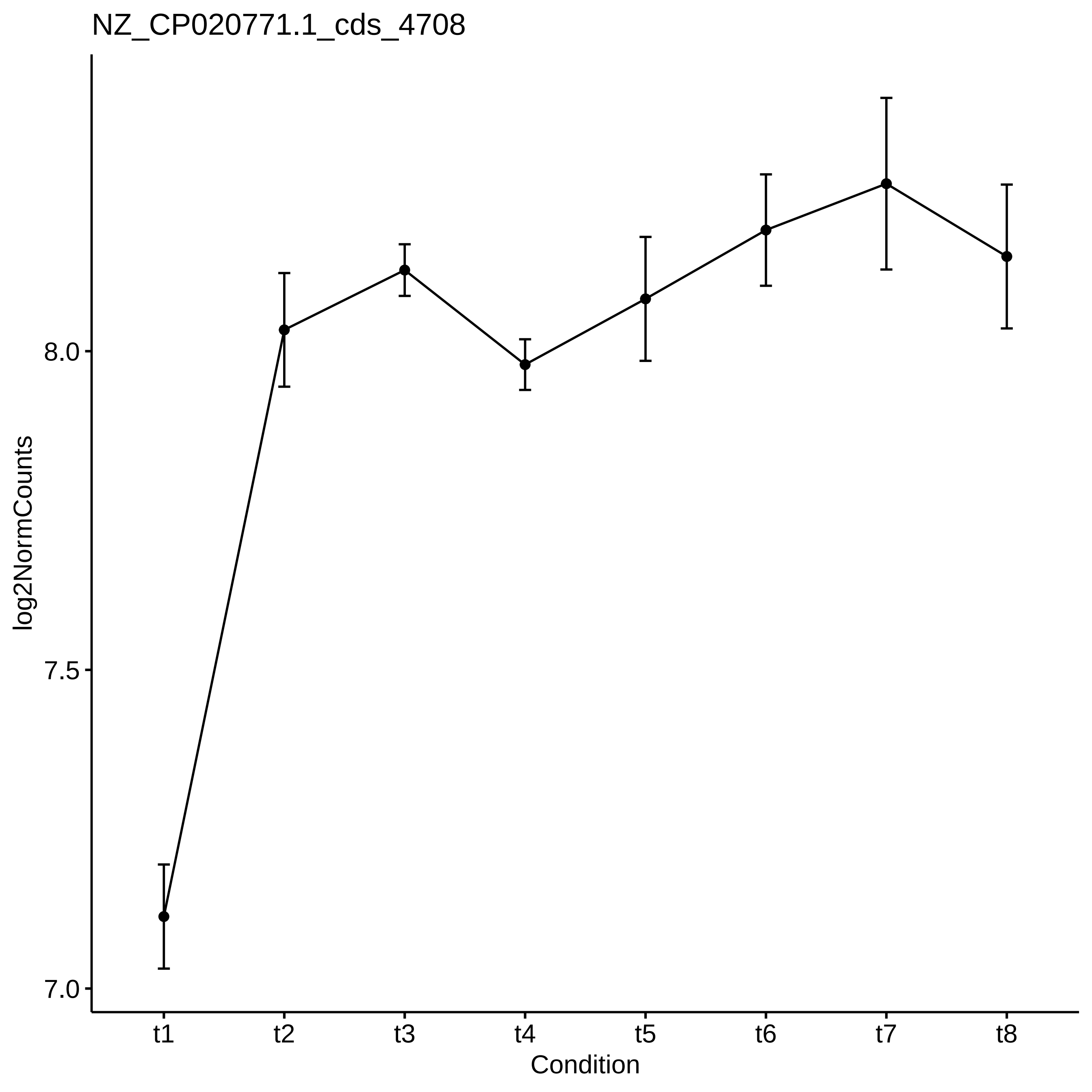

NZ_CP020771.1_cds_4708
log2NormCounts
8.0
7.5
7.0
t1
t2
t3
t4
t5
t6
t7
t8
Condition

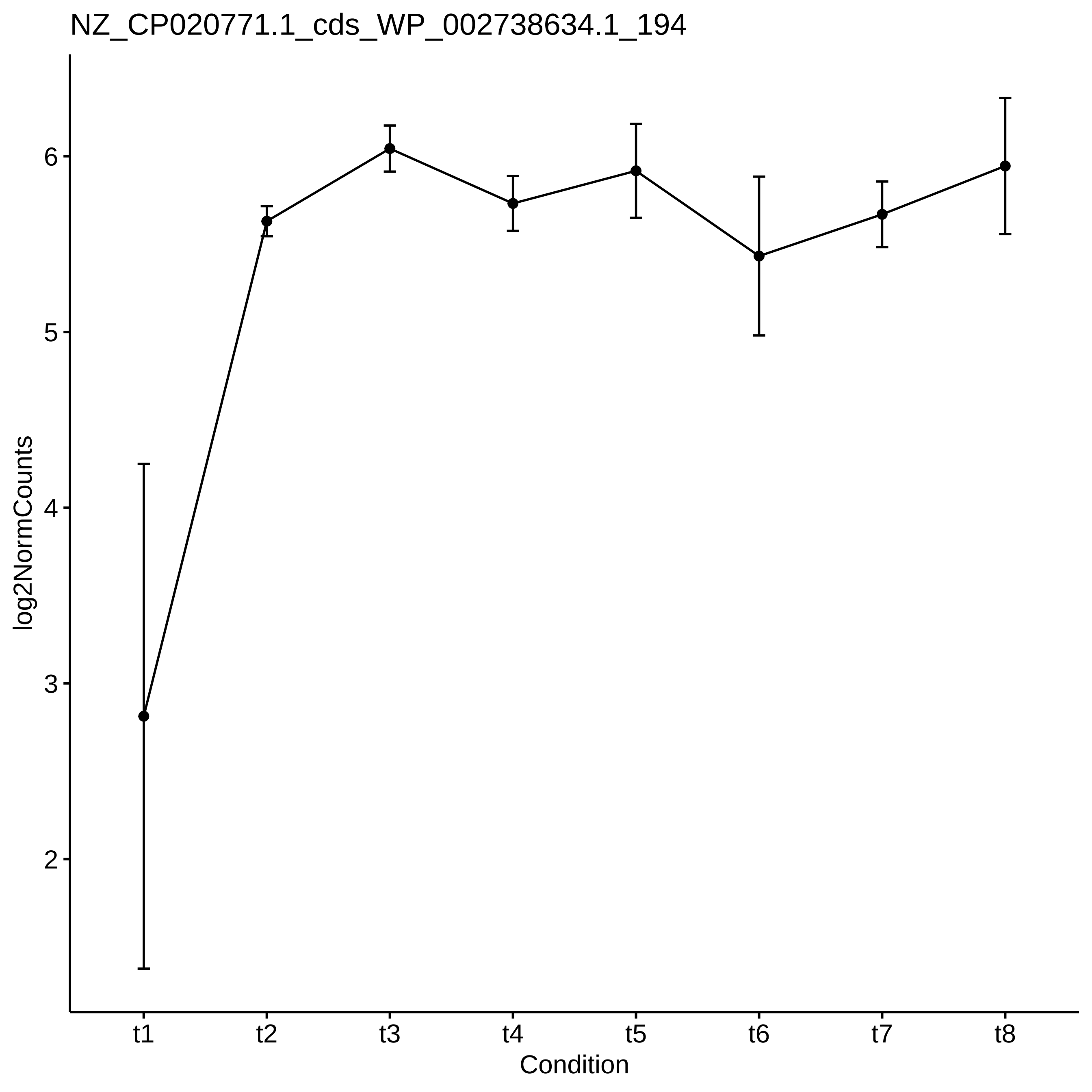
NZ_CP020771.1_cds_WP_002738634.1_194
log2NormCounts
2
3
4
5
6
t1
t2
t3
t4
t5
t6
t7
t8
Condition

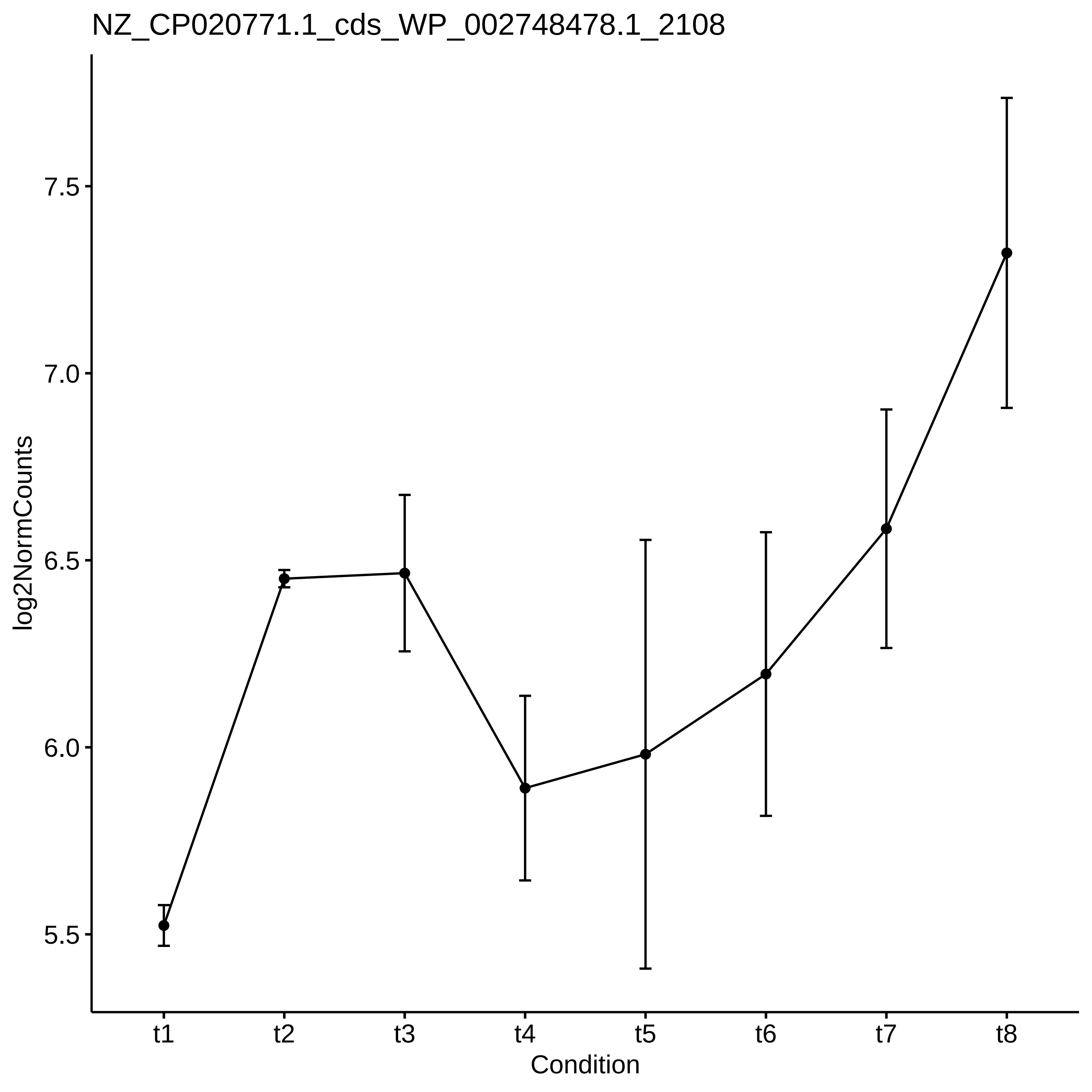

NZ_CP020771.1_cds_WP_002748478.1_2108
log2NormCounts
7.5
7.0
6.5
6.0
5.5
t1
t2
t3
t4
t5
t6
t7
t8
Condition

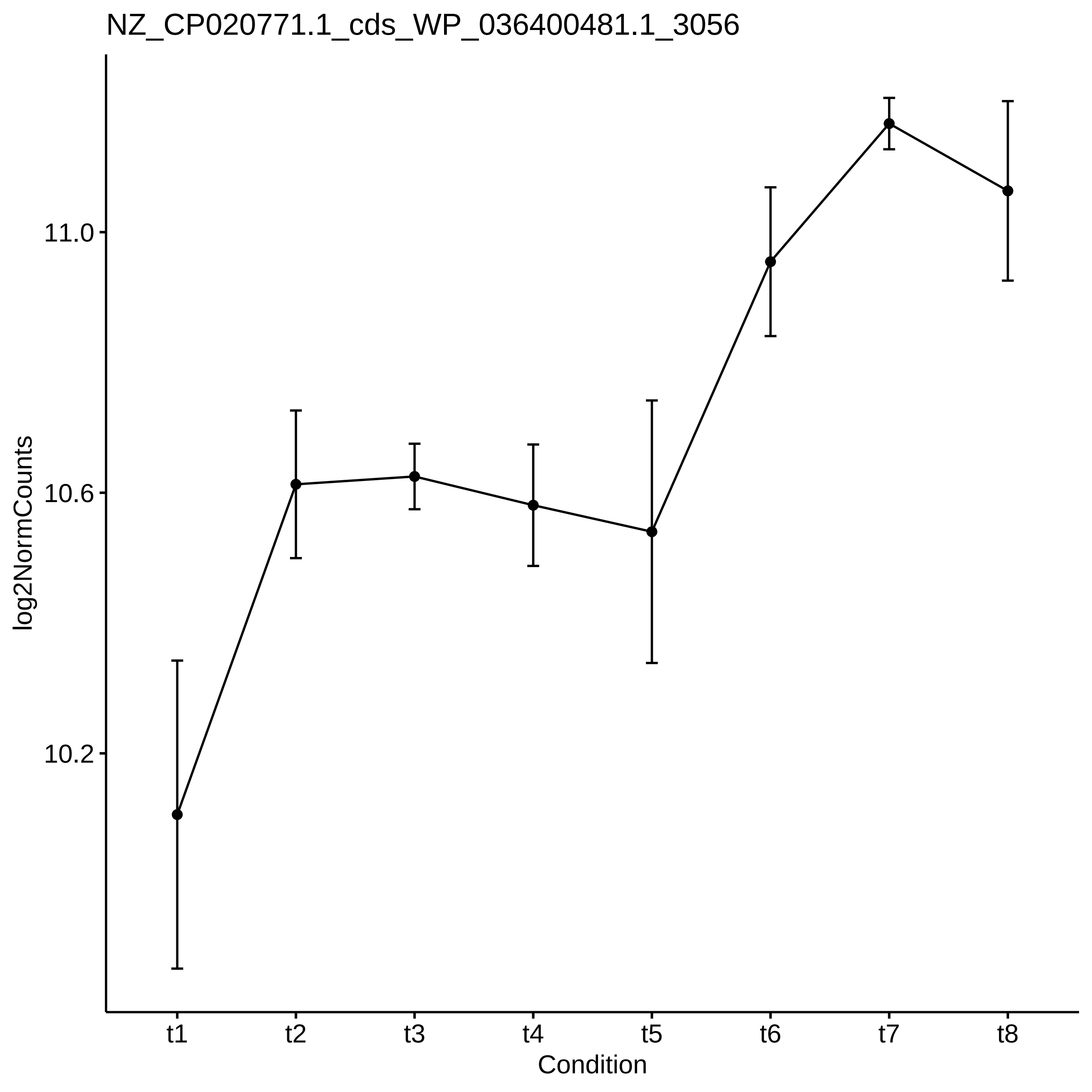
NZ_CP020771.1_cds_WP_036400481.1_3056
log2NormCounts
11.0
10.6
10.2
t1
t2
t3
t4
t5
t6
t7
t8
Condition
