## Supplementary_Data for "Diel changes in the expression of a marker gene and candidate genes for intracellular amorphous CaCO_3_ biomineralization in *Microcystis*": PCC7806_[t2-t3]_Top50_Profile.pdf

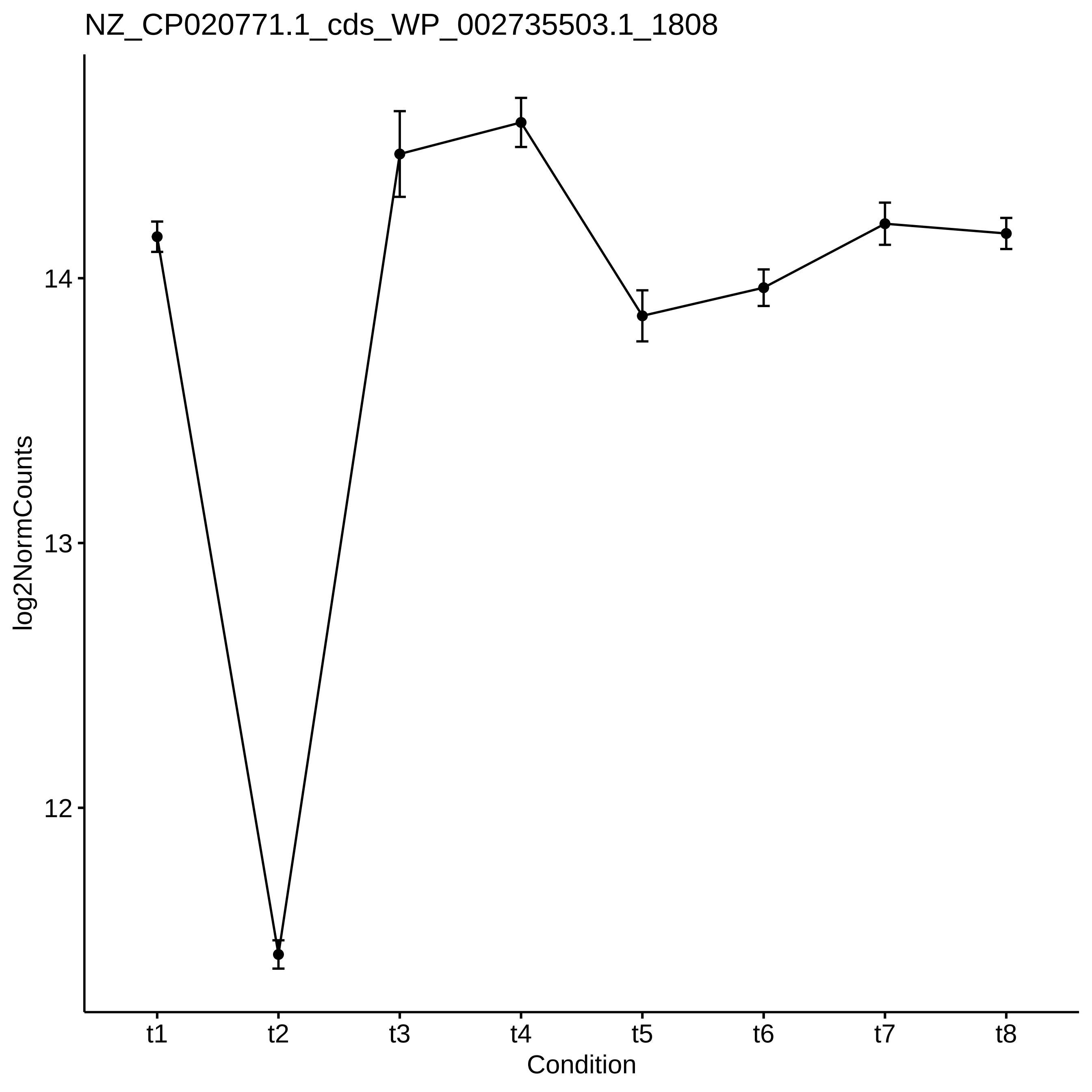
NZ_CP020771.1_cds_WP_002735503.1_1808
log2NormCounts
14
13
12
t1
t2
t3
t4
t5
t6
t7
t8
Condition

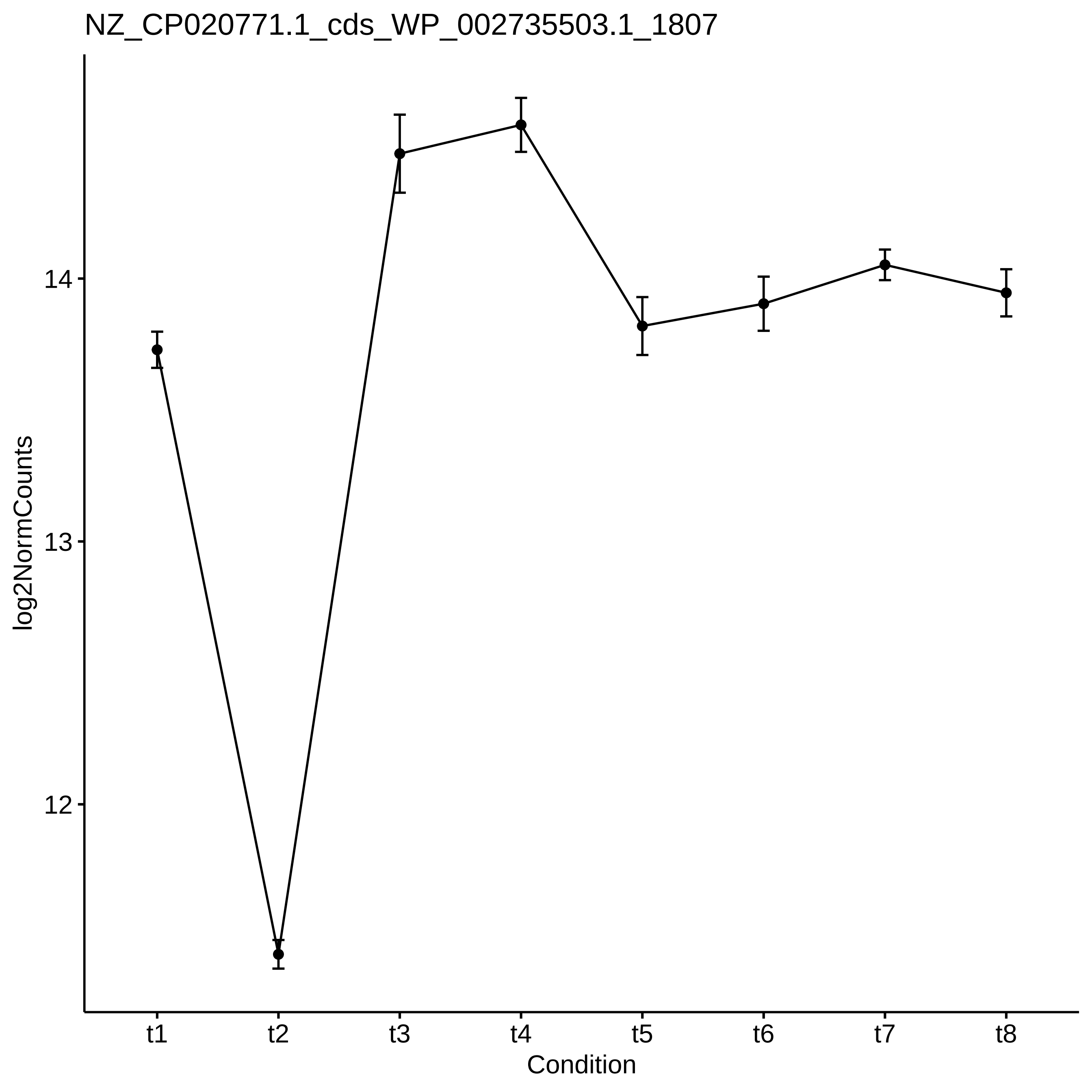
NZ_CP020771.1_cds_WP_002735503.1_1807
log2NormCounts
14
13
12
t1
t2
t3
t4
t5
t6
t7
t8
Condition

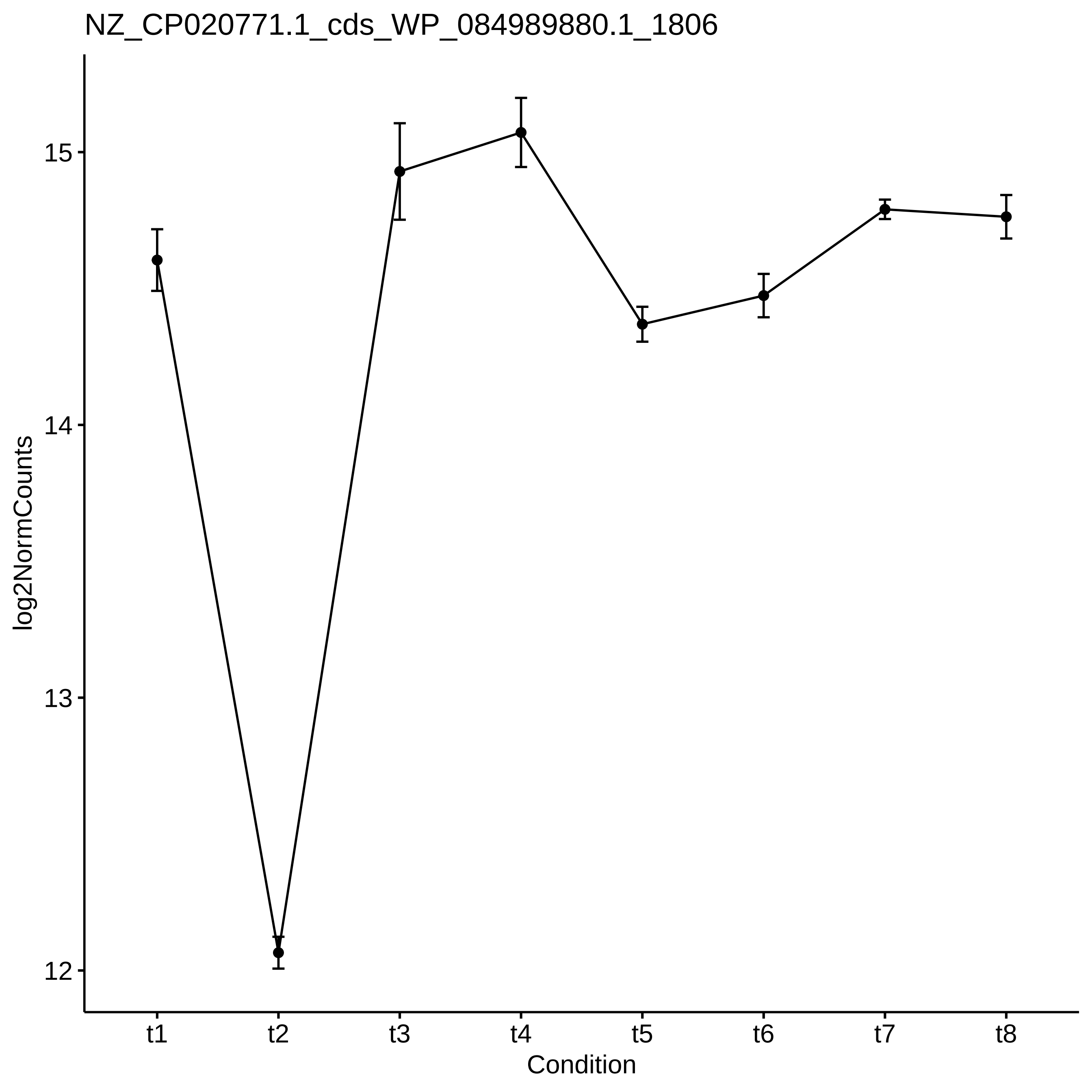

NZ_CP020771.1_cds_WP_084989880.1_1806
log2NormCounts
15
14
13
12
t1
t2
t3
t4
t5
t6
t7
t8
Condition

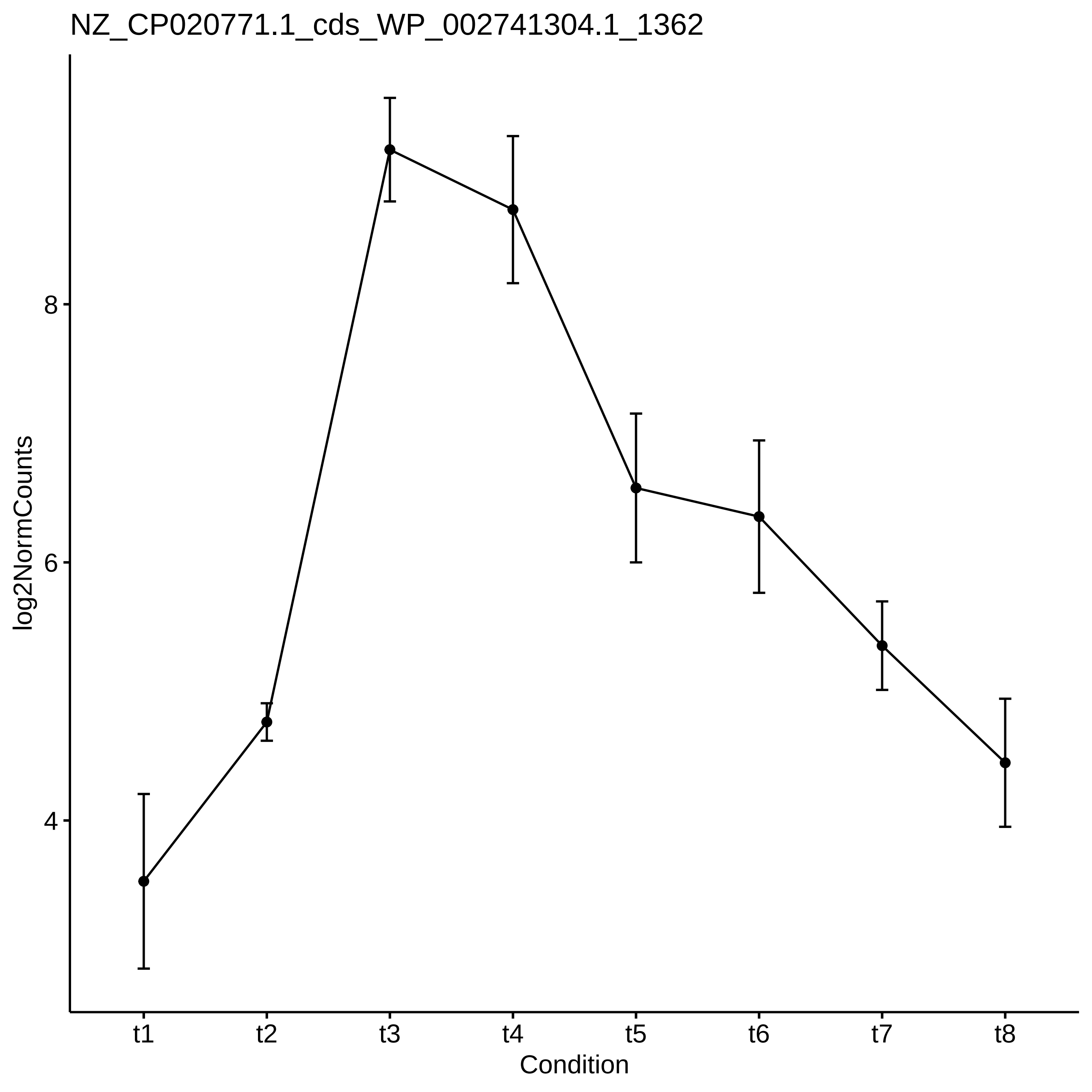
NZ_CP020771.1_cds_WP_002741304.1_1362
log2NormCounts
8
6
4
t1
t2
t3
t4
t5
t6
t7
t8
Condition

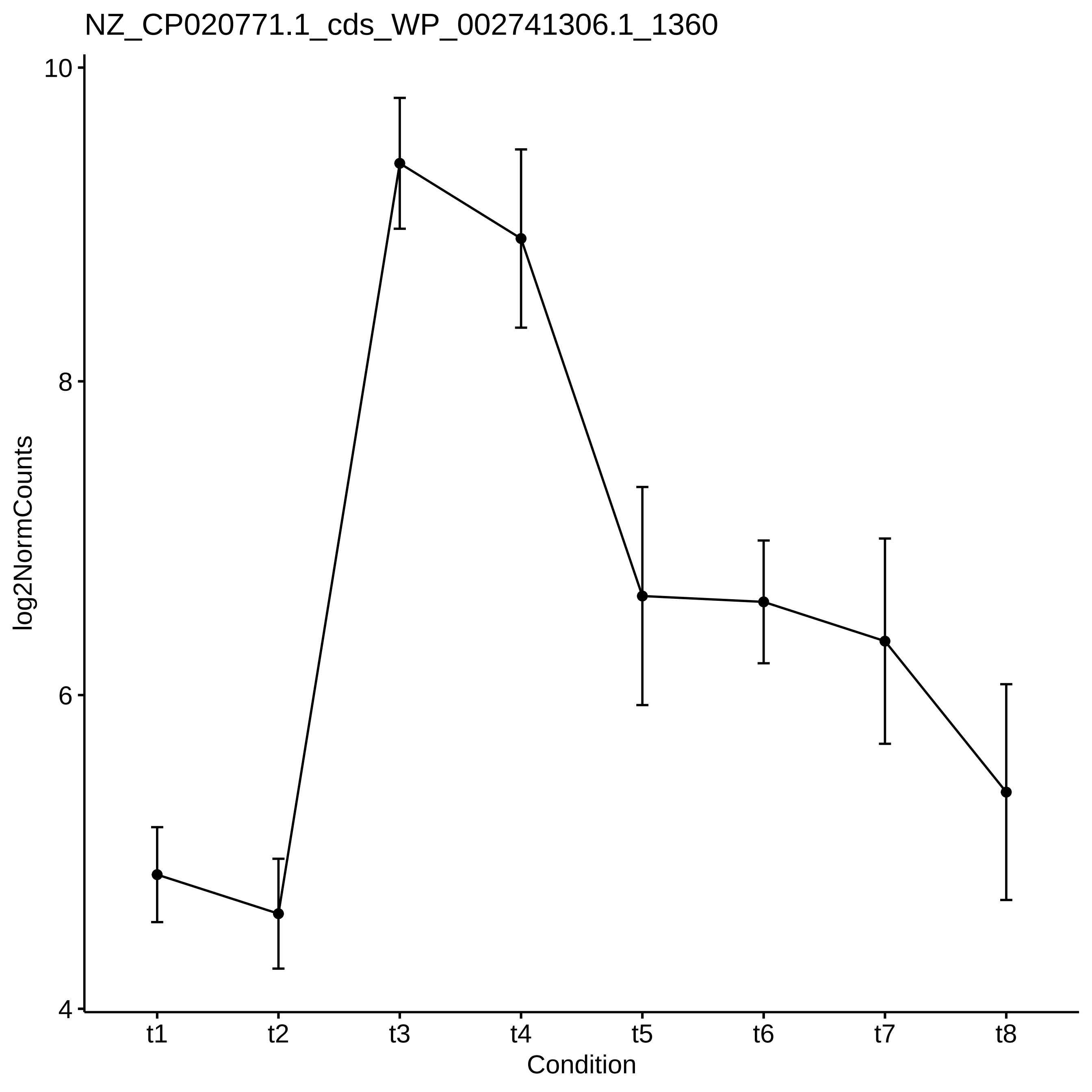
NZ_CP020771.1_cds_WP_002741306.1_1360
log2NormCounts
10
8
6
4
t1
t2
t3
t4
t5
t6
t7
t8
Condition

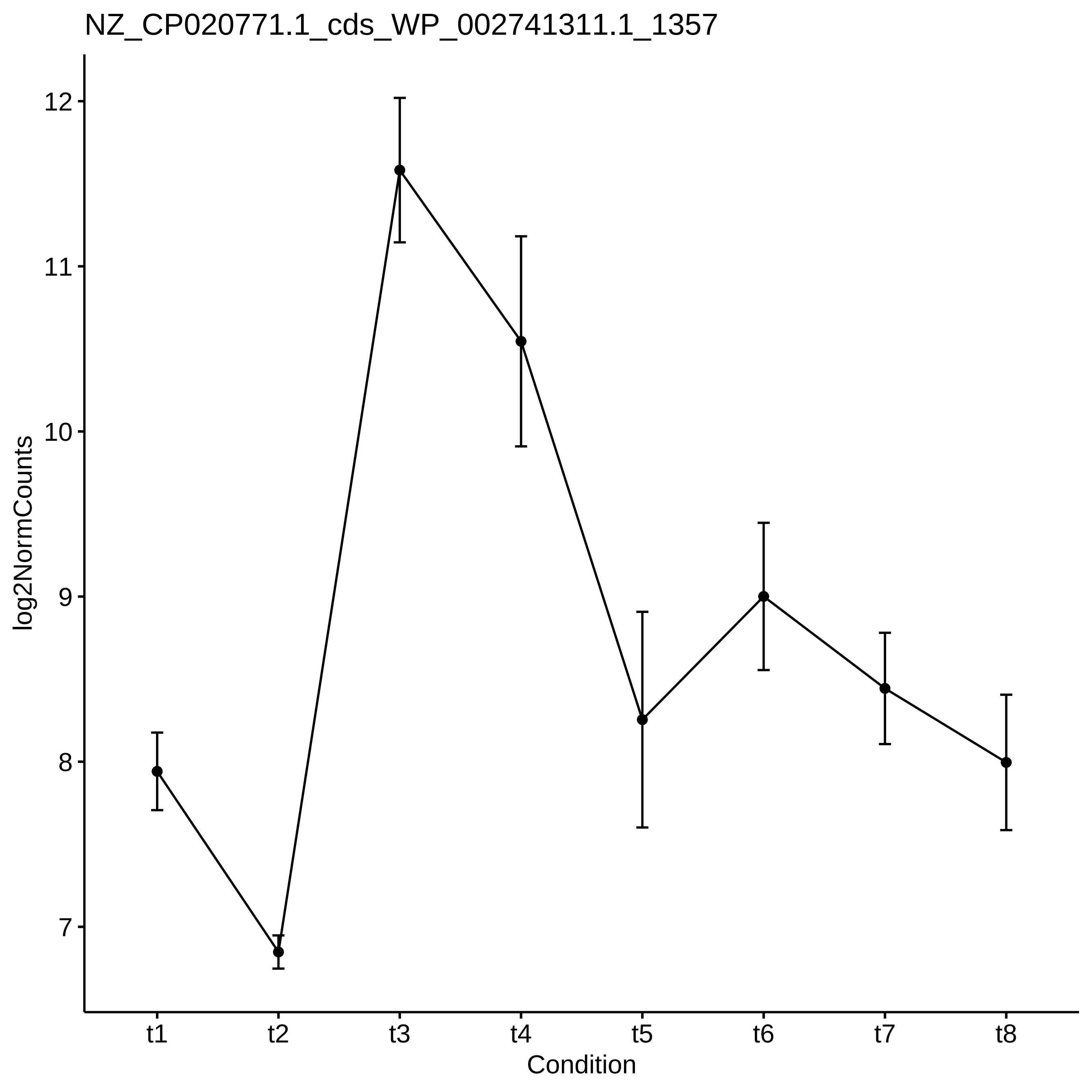
NZ_CP020771.1_cds_WP_002741311.1_1357
log2NormCounts
12
11
10
9
8
7
t1
t2
t3
t4
t5
t6
t7
t8
Condition

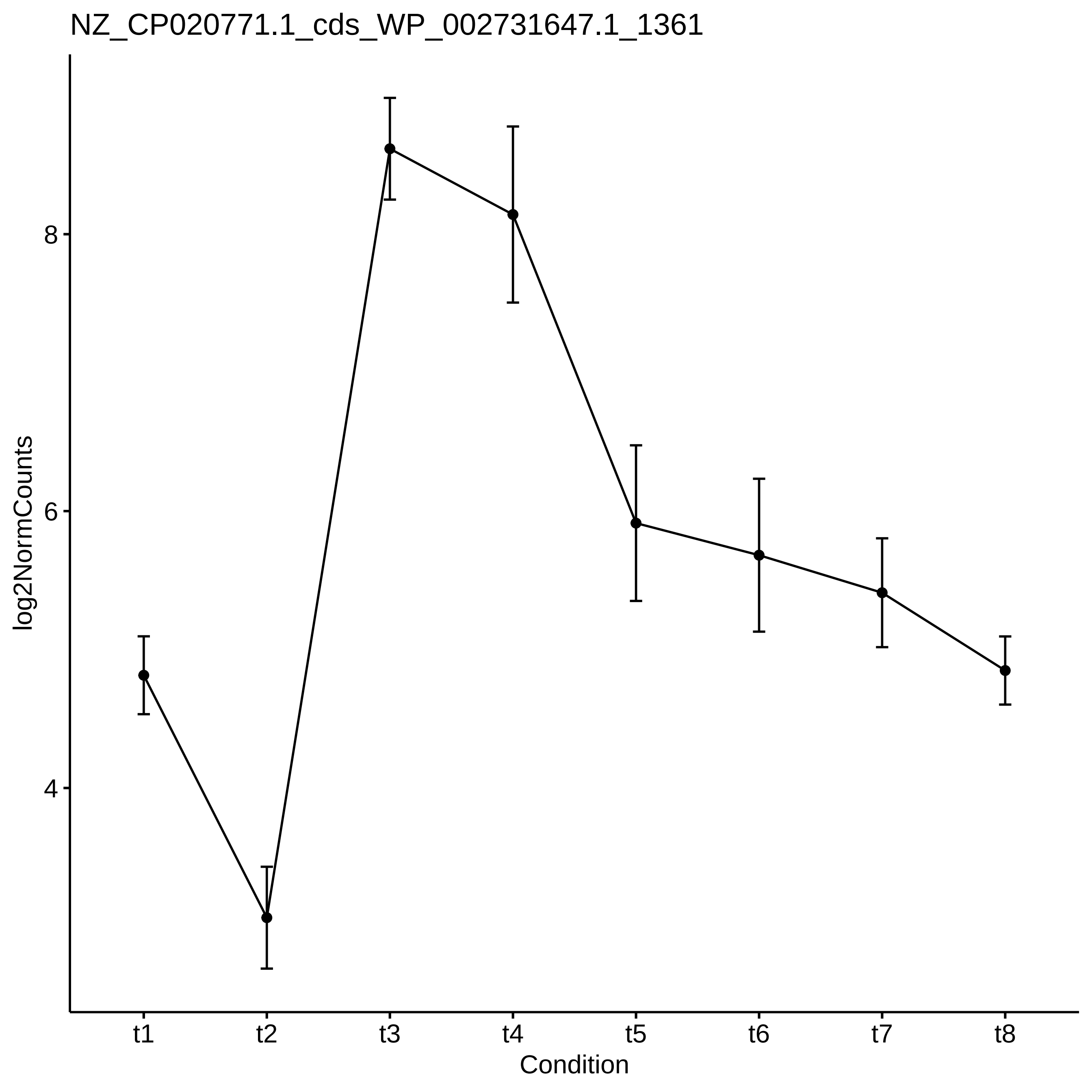
NZ_CP020771.1_cds_WP_002731647.1_1361
log2NormCounts
8
6
4
t1
t2
t3
t4
t5
t6
t7
t8
Condition

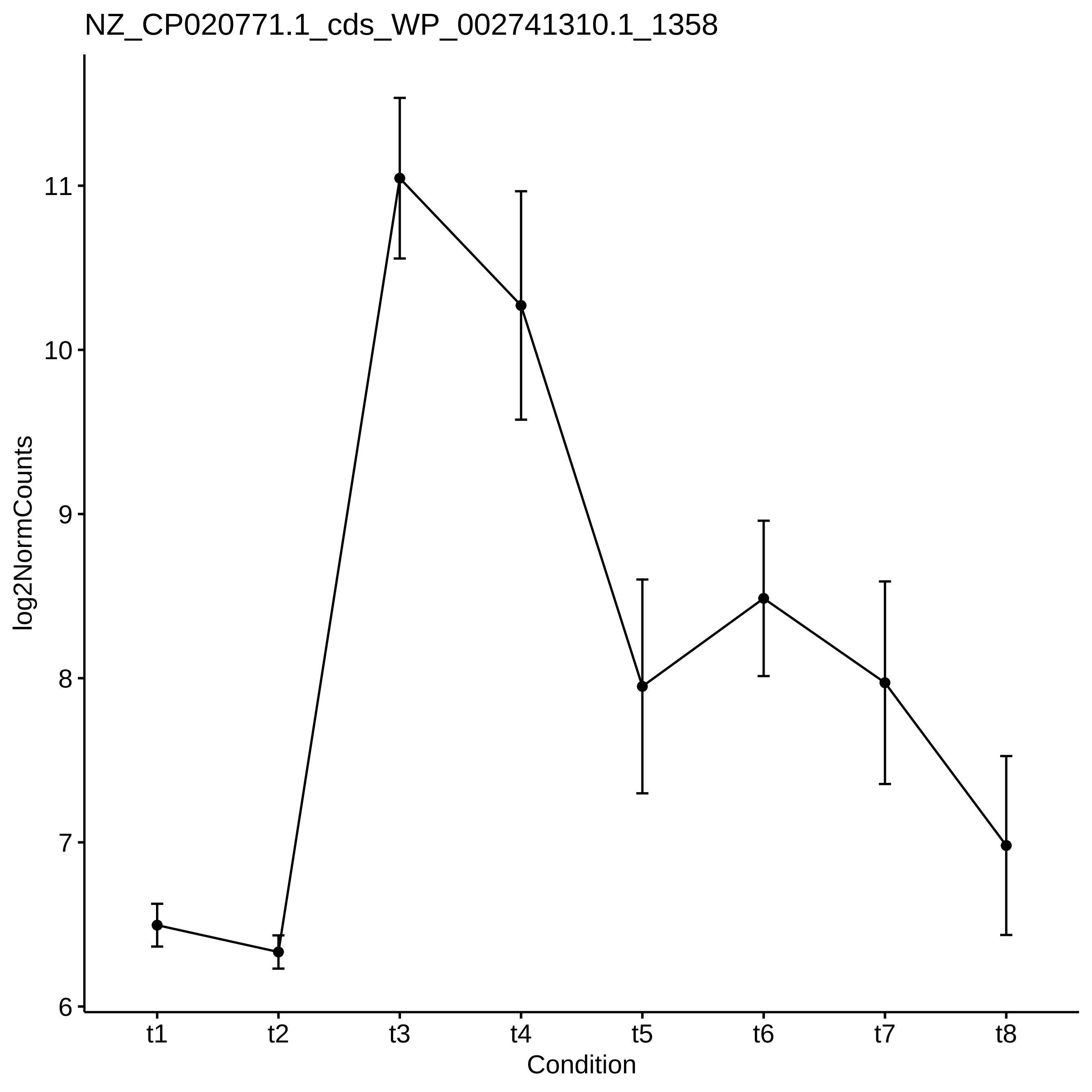
NZ_CP020771.1_cds_WP_002741310.1_1358
log2NormCounts
11
10
9
8
7
6
t1
t2
t3
t4
t5
t6
t7
t8
Condition

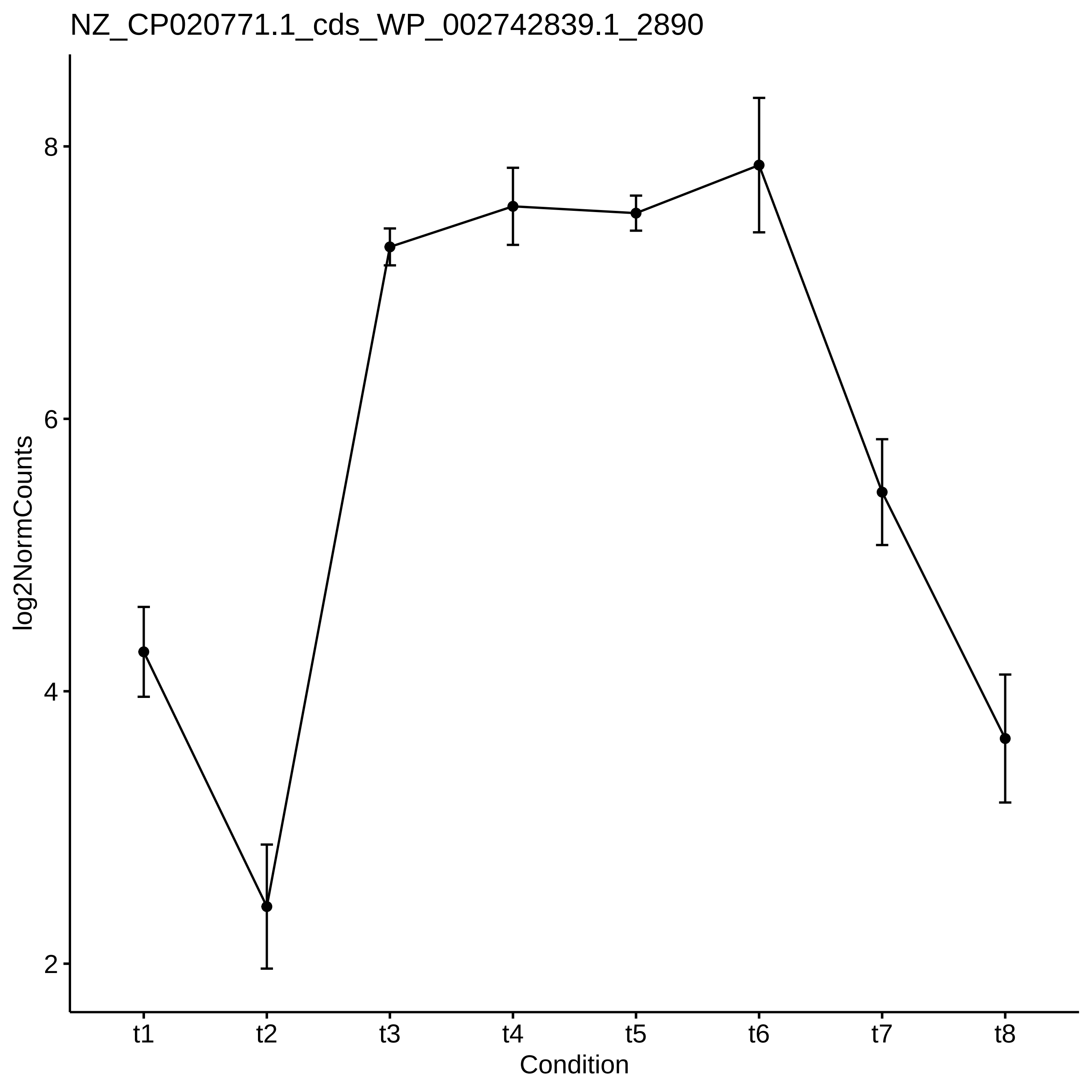
NZ_CP020771.1_cds_WP_002742839.1_2890
log2NormCounts
8
6
4
2
t1
t2
t3
t4
t5
t6
t7
t8
Condition

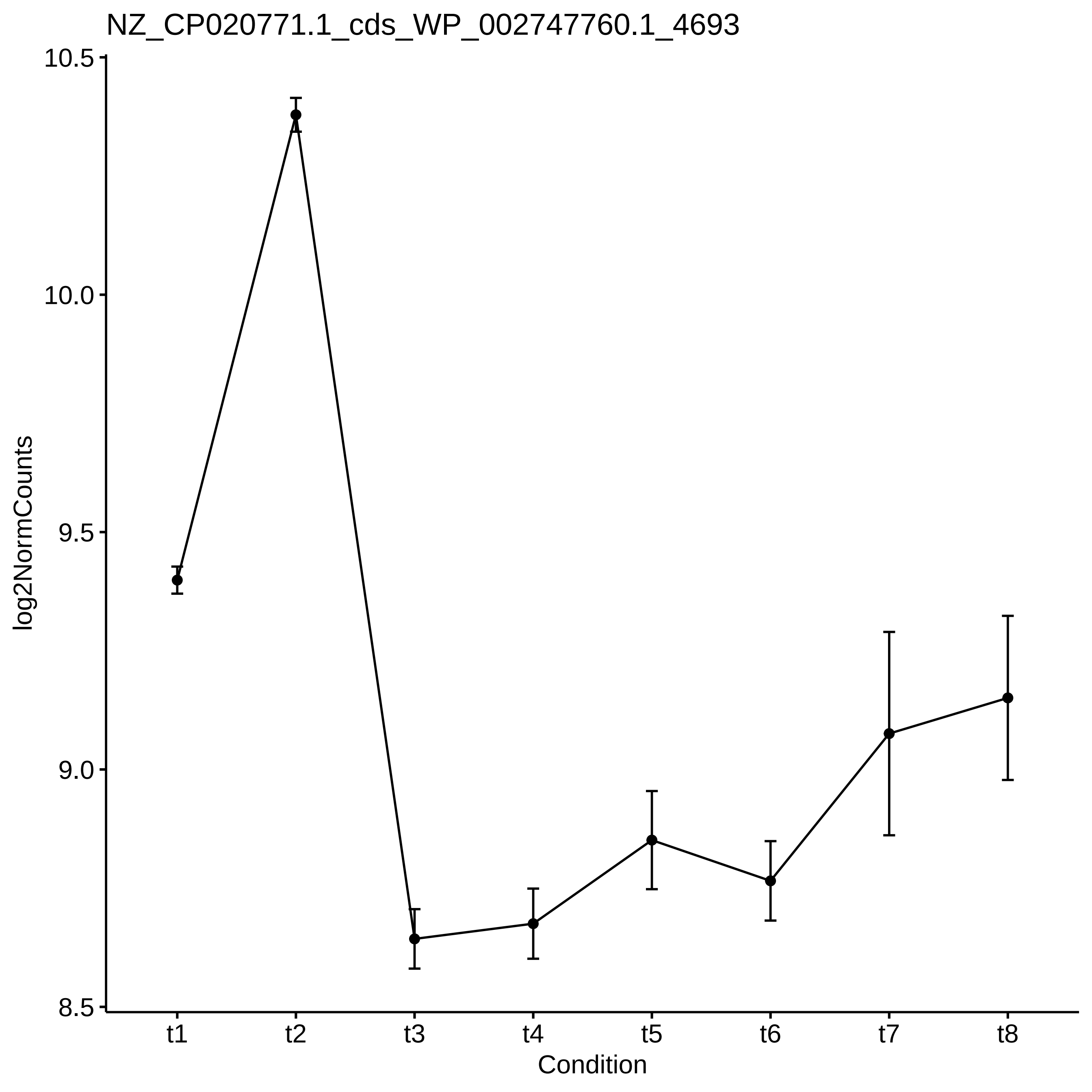

NZ_CP020771.1_cds_WP_002747760.1_4693
10.5
10.0
9.5
9.0
8.5
log2NormCounts
t1
t2
t3
t4
t5
t6
t7
t8
Condition

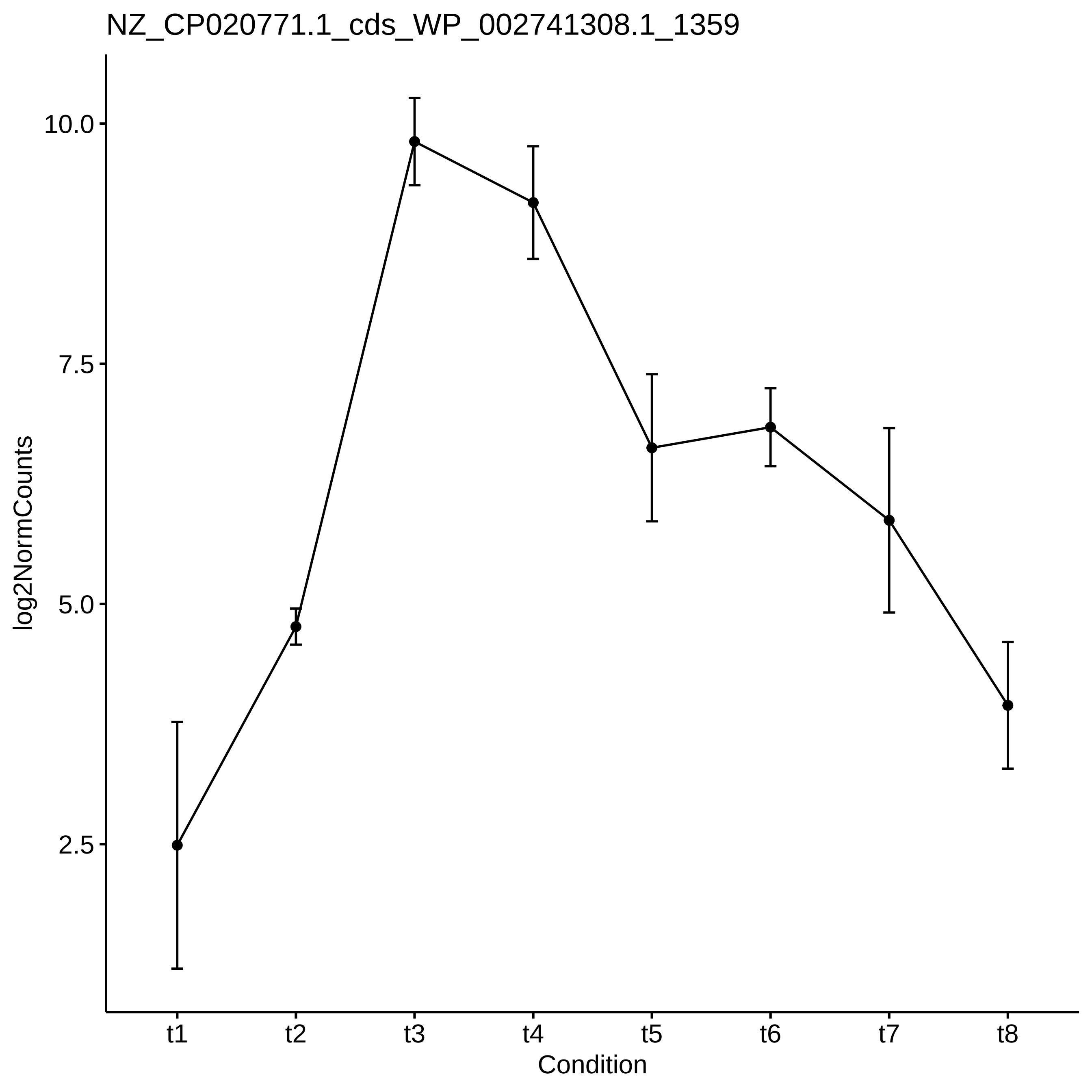
NZ_CP020771.1_cds_WP_002741308.1_1359
log2NormCounts
10.0
7.5
5.0
2.5
t1
t2
t3
t4
t5
t6
t7
t8
Condition

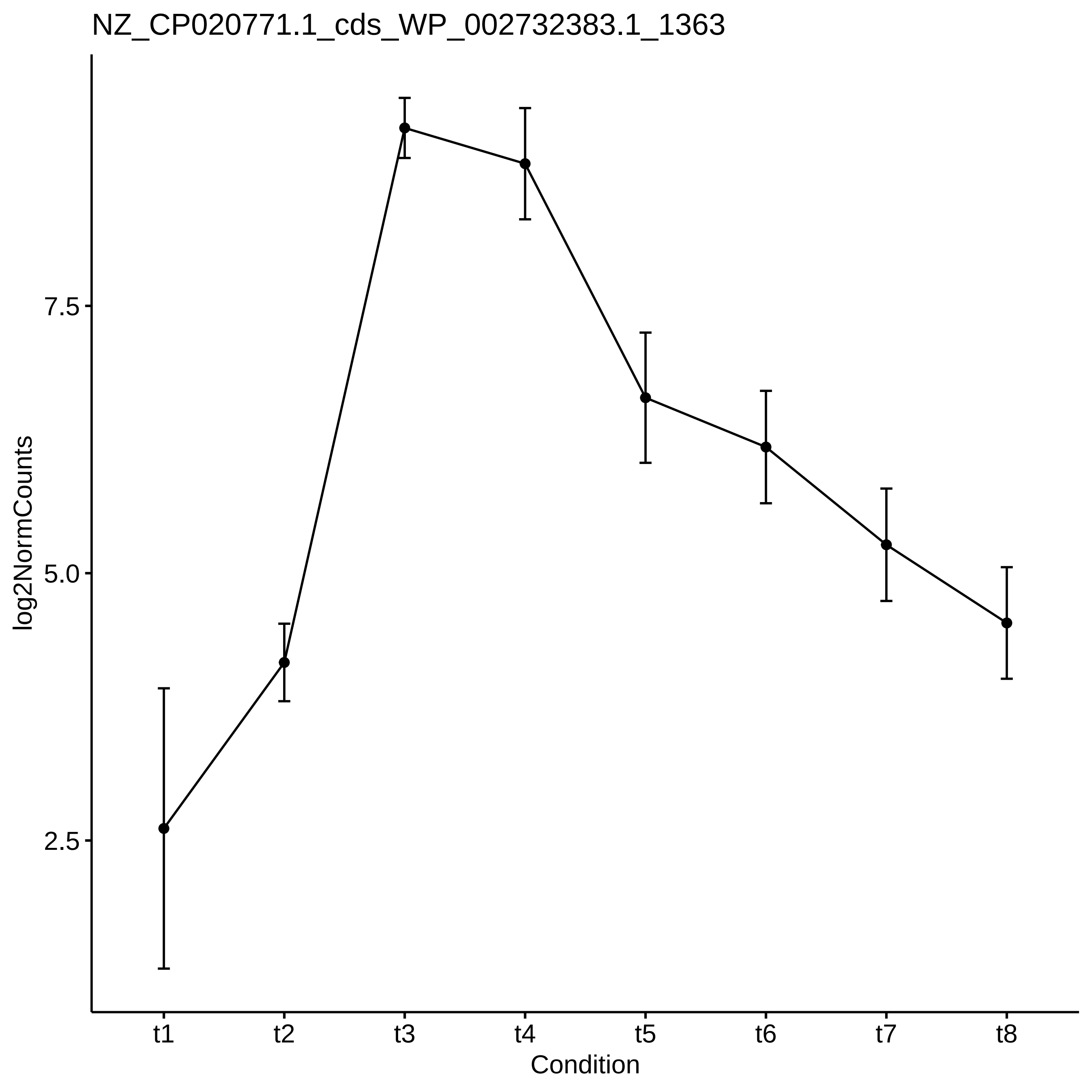

NZ_CP020771.1_cds_WP_002732383.1_1363
log2NormCounts
7.5
5.0
2.5
t1
t2
t3
t4
t5
t6
t7
t8
Condition

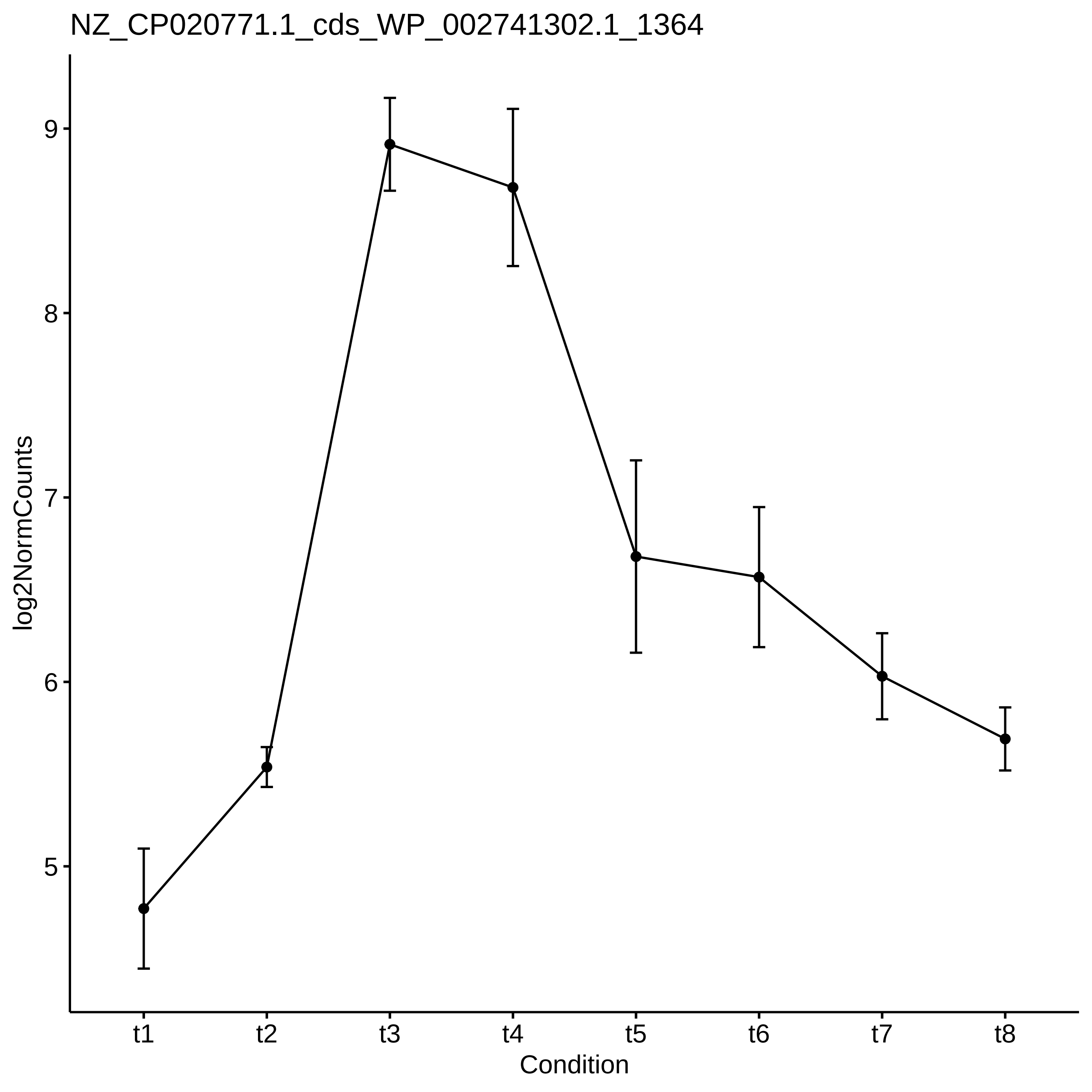
NZ_CP020771.1_cds_WP_002741302.1_1364
log2NormCounts
9
8
7
6
5
t1
t2
t3
t4
t5
t6
t7
t8
Condition

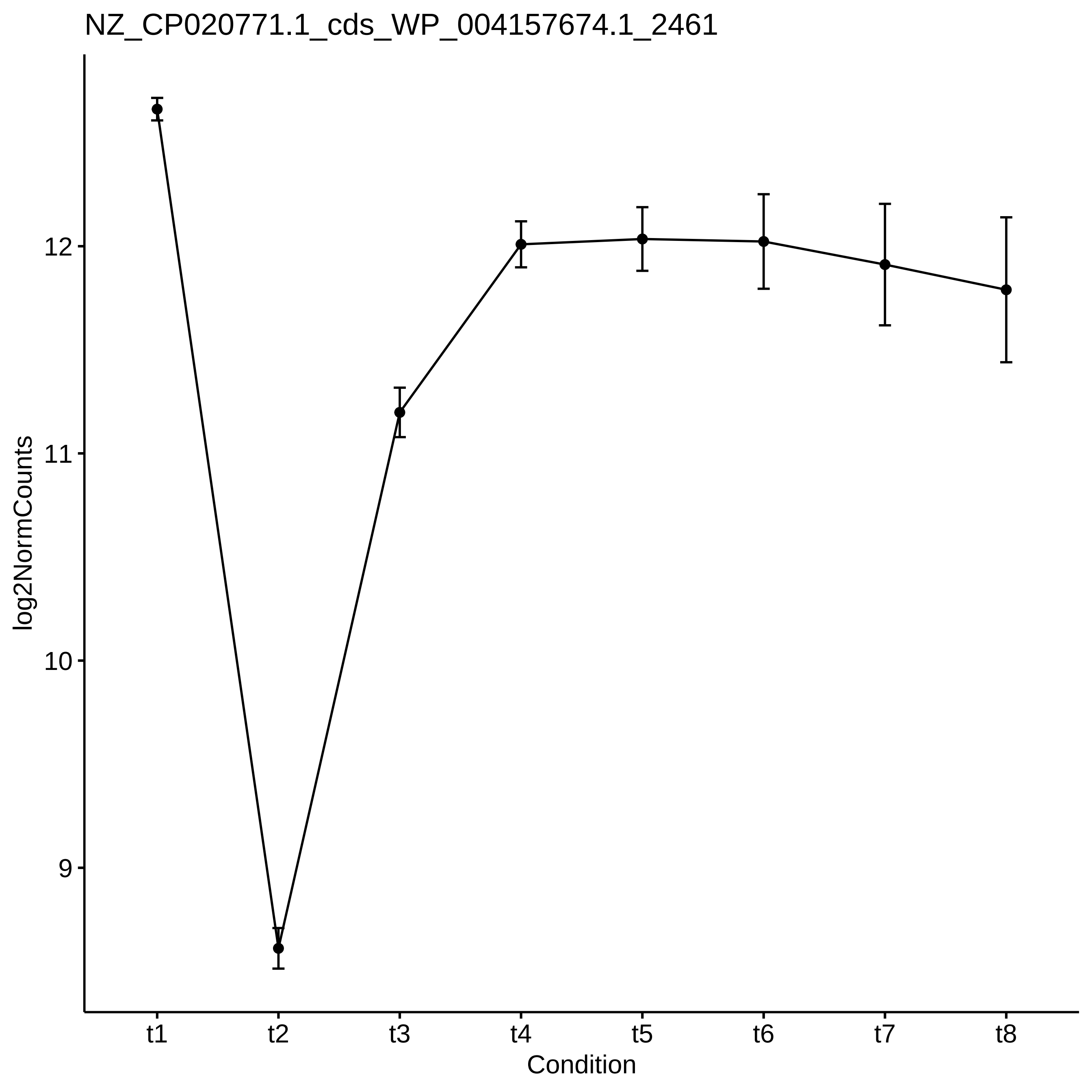
NZ_CP020771.1_cds_WP_004157674.1_2461
log2NormCounts
12
11
10
9
t1
t2
t3
t4
t5
t6
t7
t8
Condition

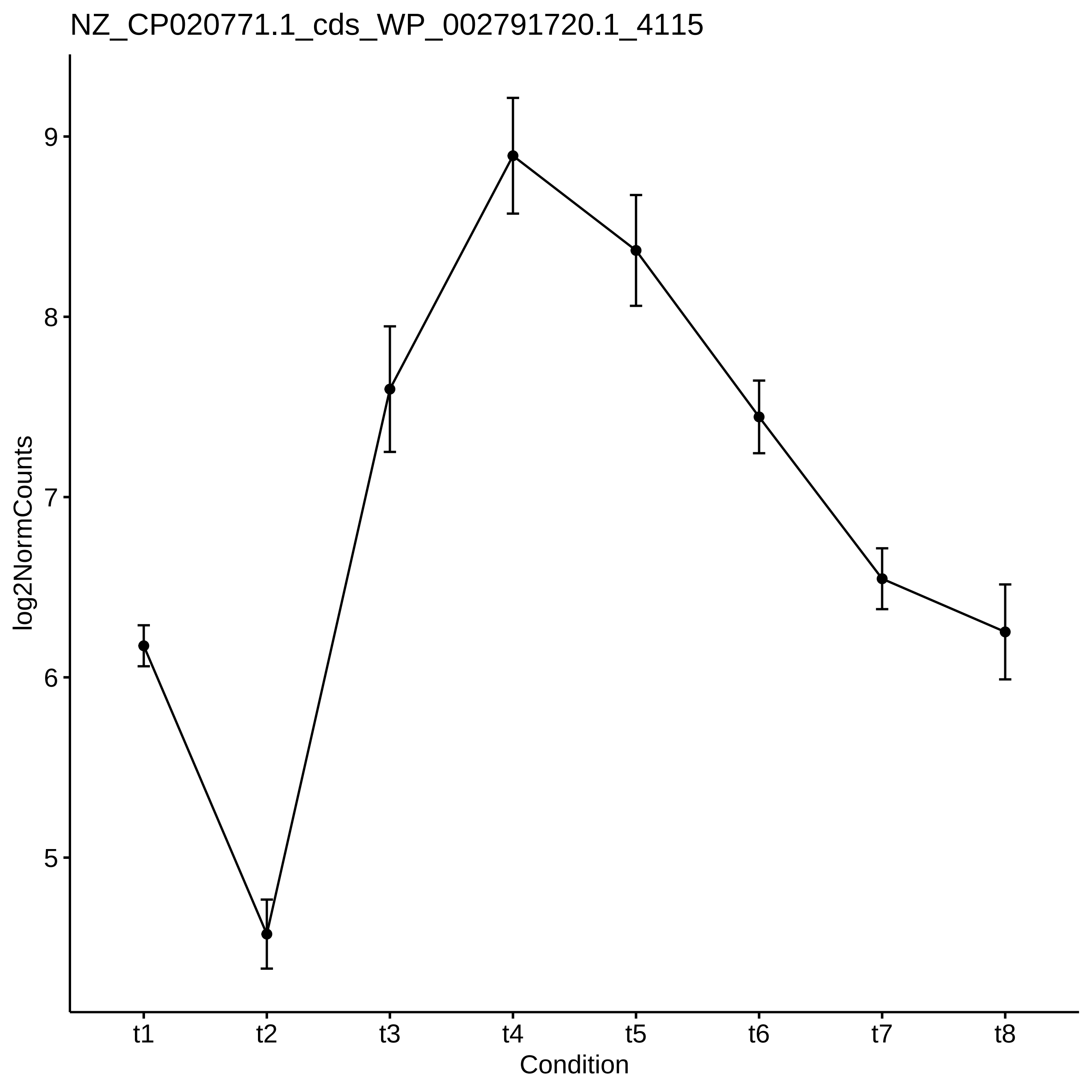
NZ_CP020771.1_cds_WP_002791720.1_4115
log2NormCounts
9
8
7
6
5
t1
t2
t3
t4
t5
t6
t7
t8
Condition

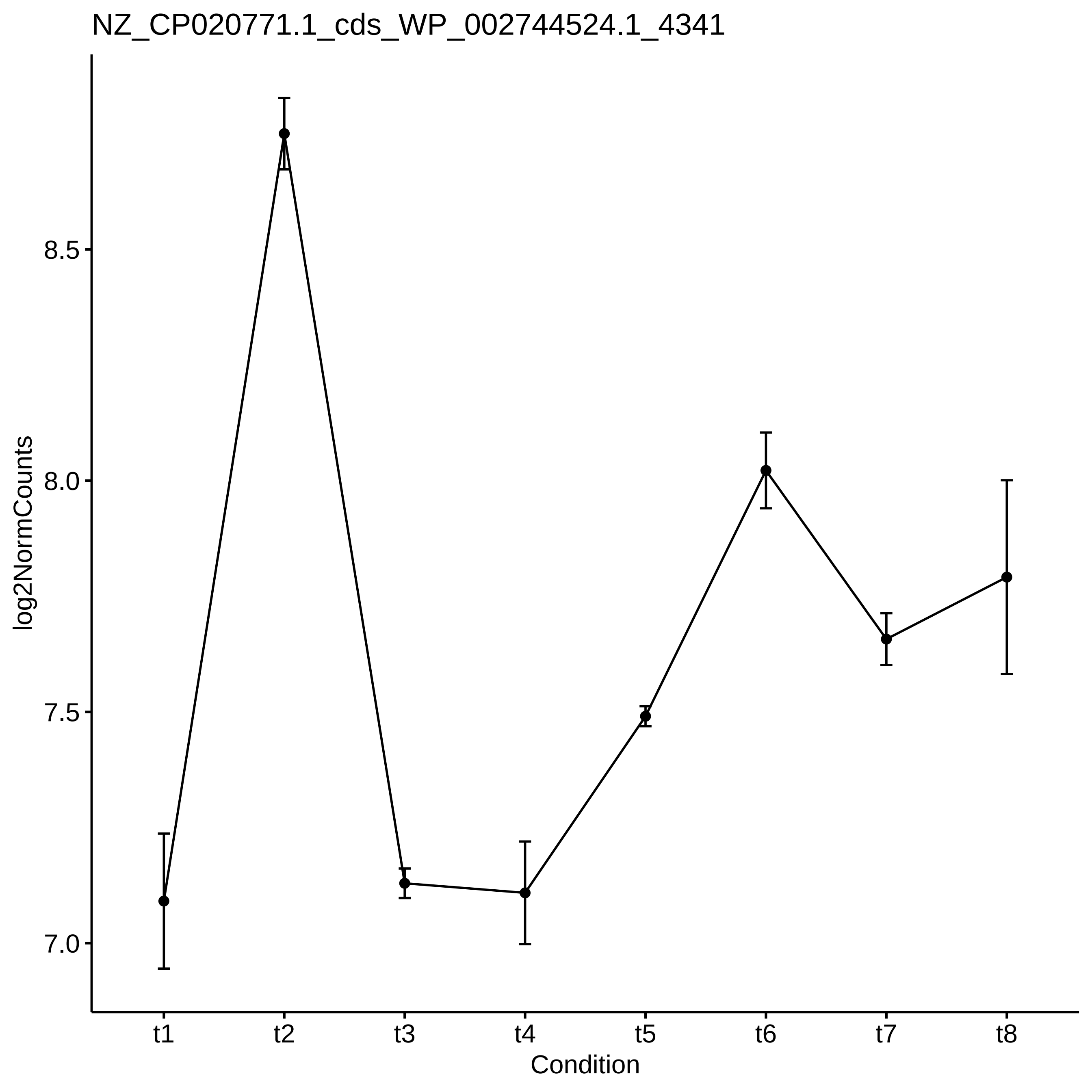

NZ_CP020771.1_cds_WP_002744524.1_4341
log2NormCounts
8.5
8.0
7.5
7.0
t1
t2
t3
t4
t5
t6
t7
t8
Condition

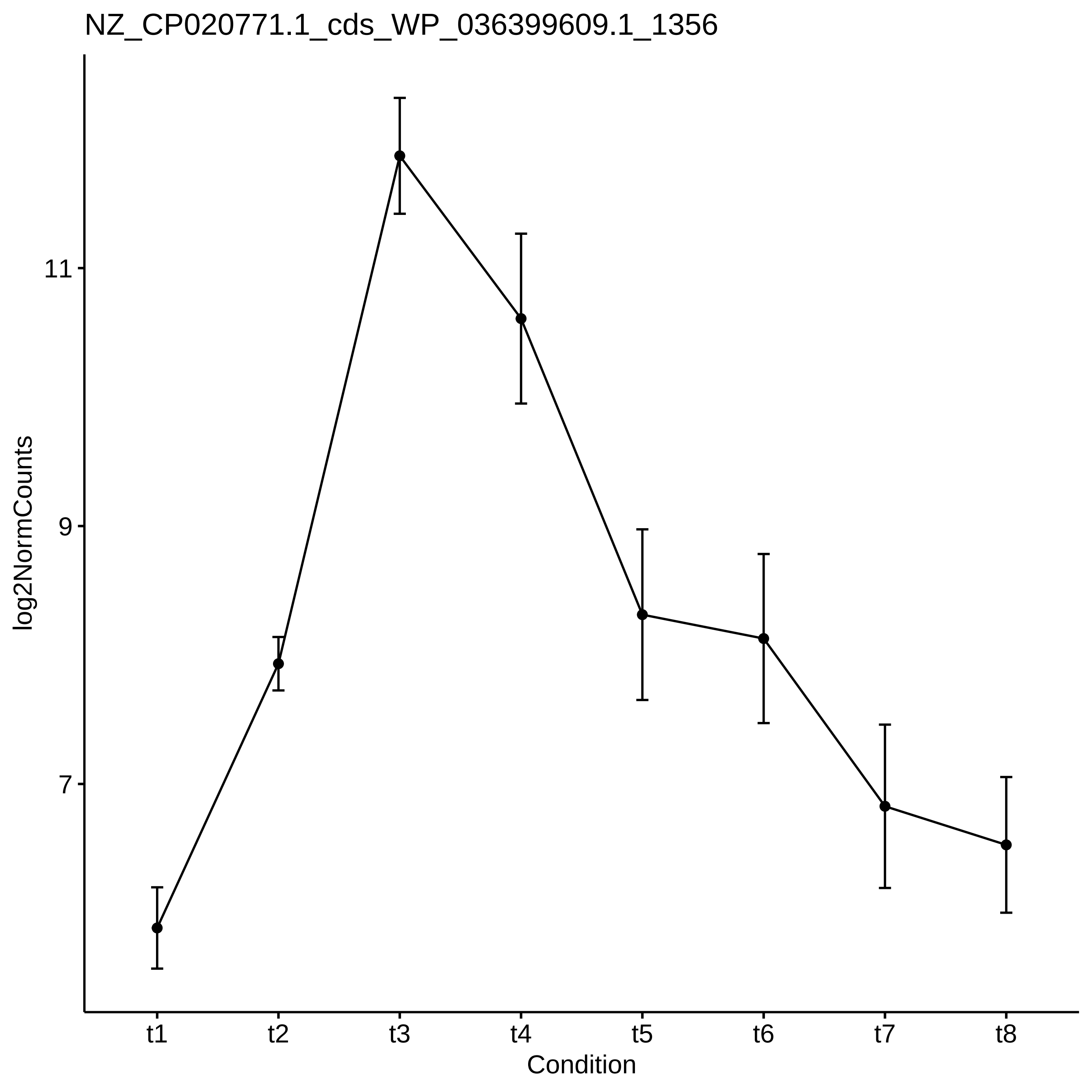
NZ_CP020771.1_cds_WP_036399609.1_1356
log2NormCounts
11
9
7
t1
t2
t3
t4
t5
t6
t7
t8
Condition

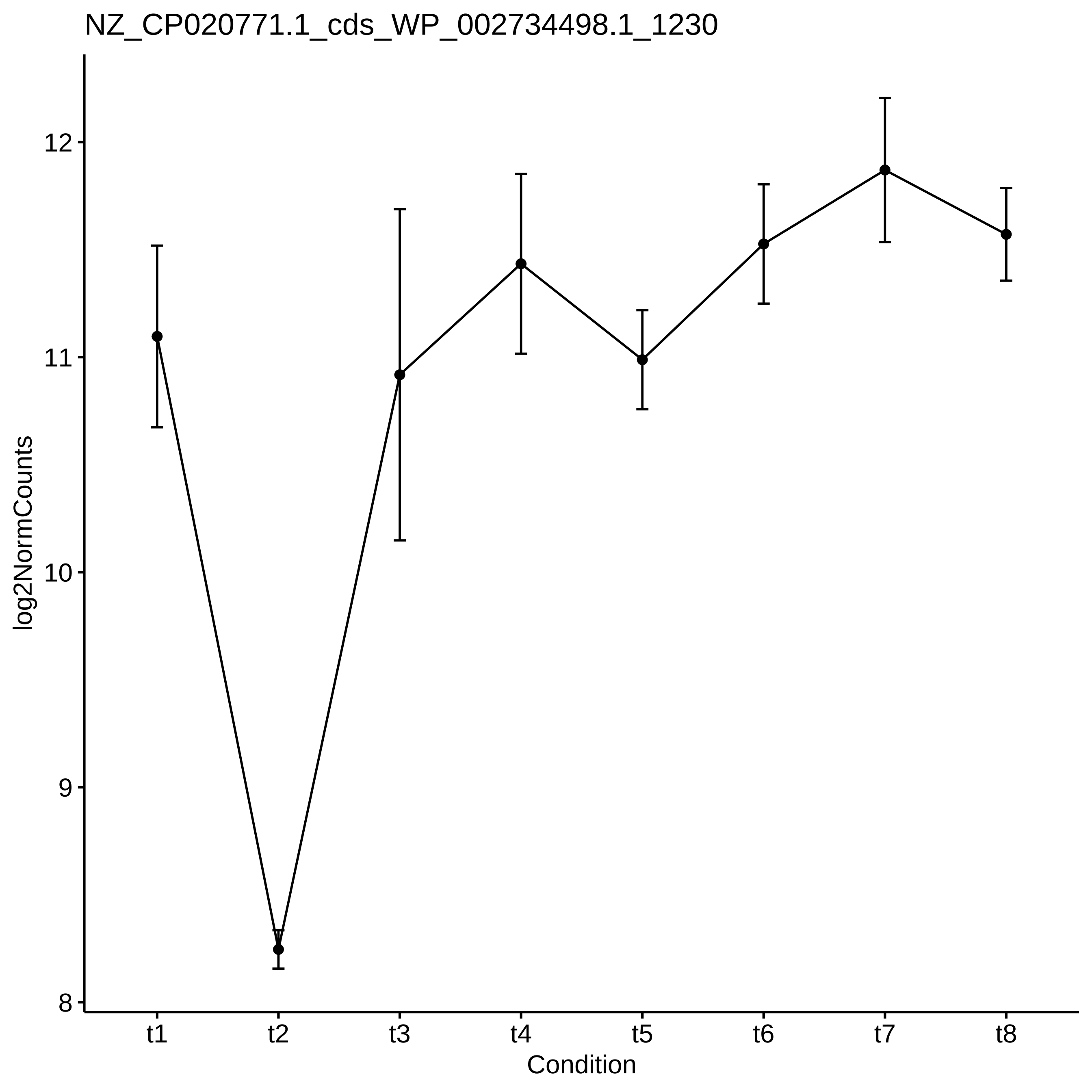
NZ_CP020771.1_cds_WP_002734498.1_1230
log2NormCounts
12
11
10
9
8
t1
t2
t3
t4
t5
t6
t7
t8
Condition

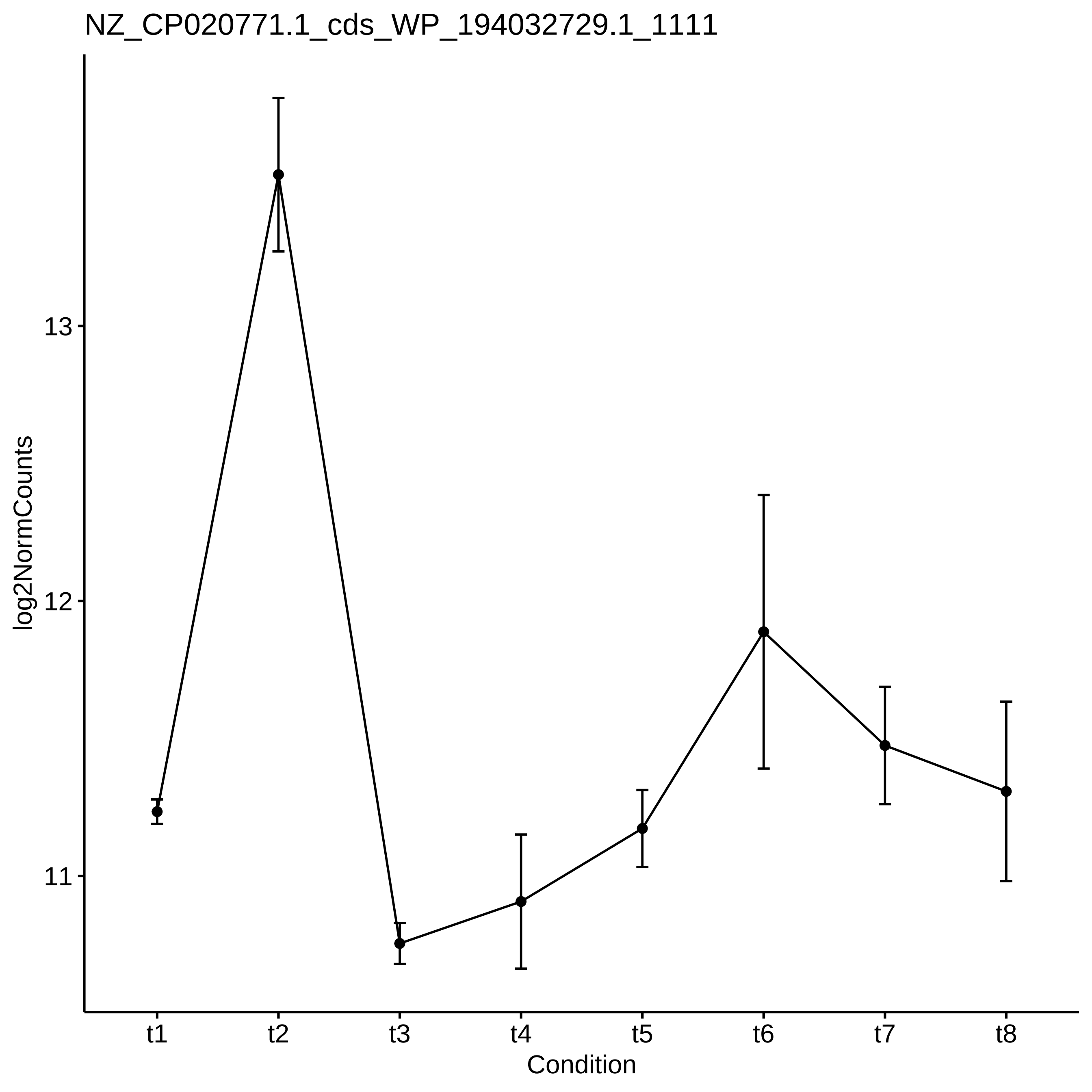
NZ_CP020771.1_cds_WP_194032729.1_1111
log2NormCounts
13
12
11
t1
t2
t3
t4
t5
t6
t7
t8
Condition

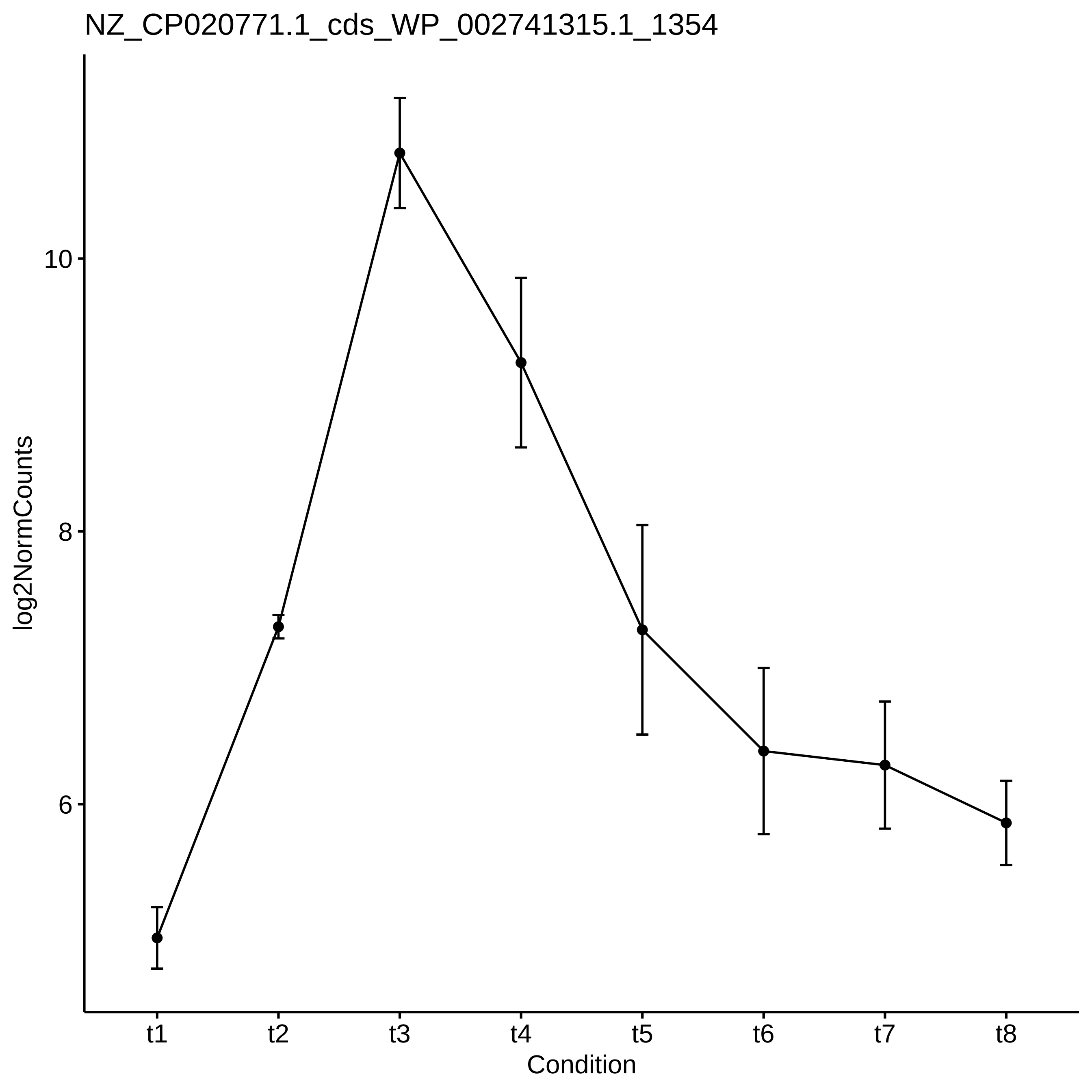
NZ_CP020771.1_cds_WP_002741315.1_1354
log2NormCounts
10
8
6
t1
t2
t3
t4
t5
t6
t7
t8
Condition

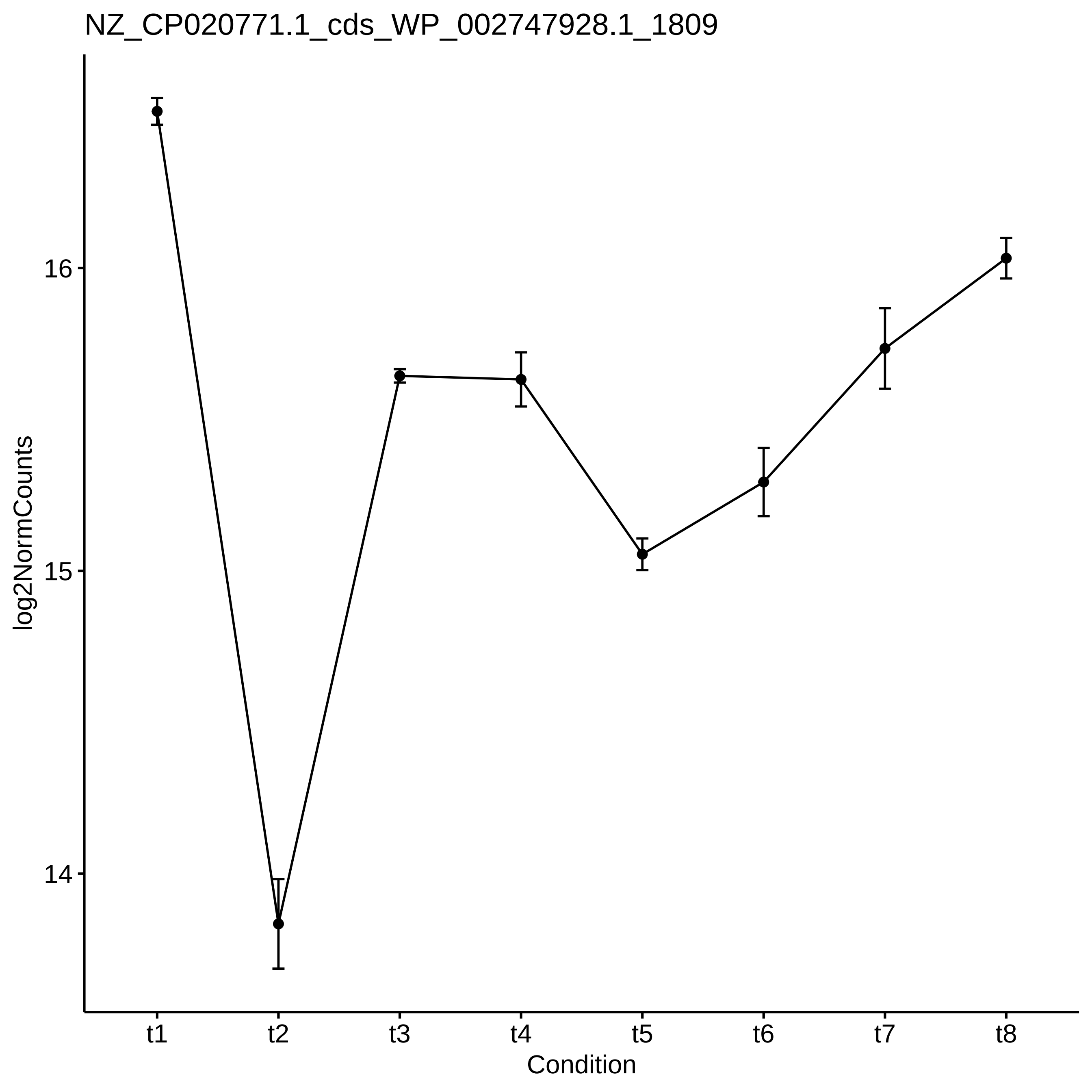
NZ_CP020771.1_cds_WP_002747928.1_1809
log2NormCounts
16
15
14
t1
t2
t3
t4
t5
t6
t7
t8
Condition

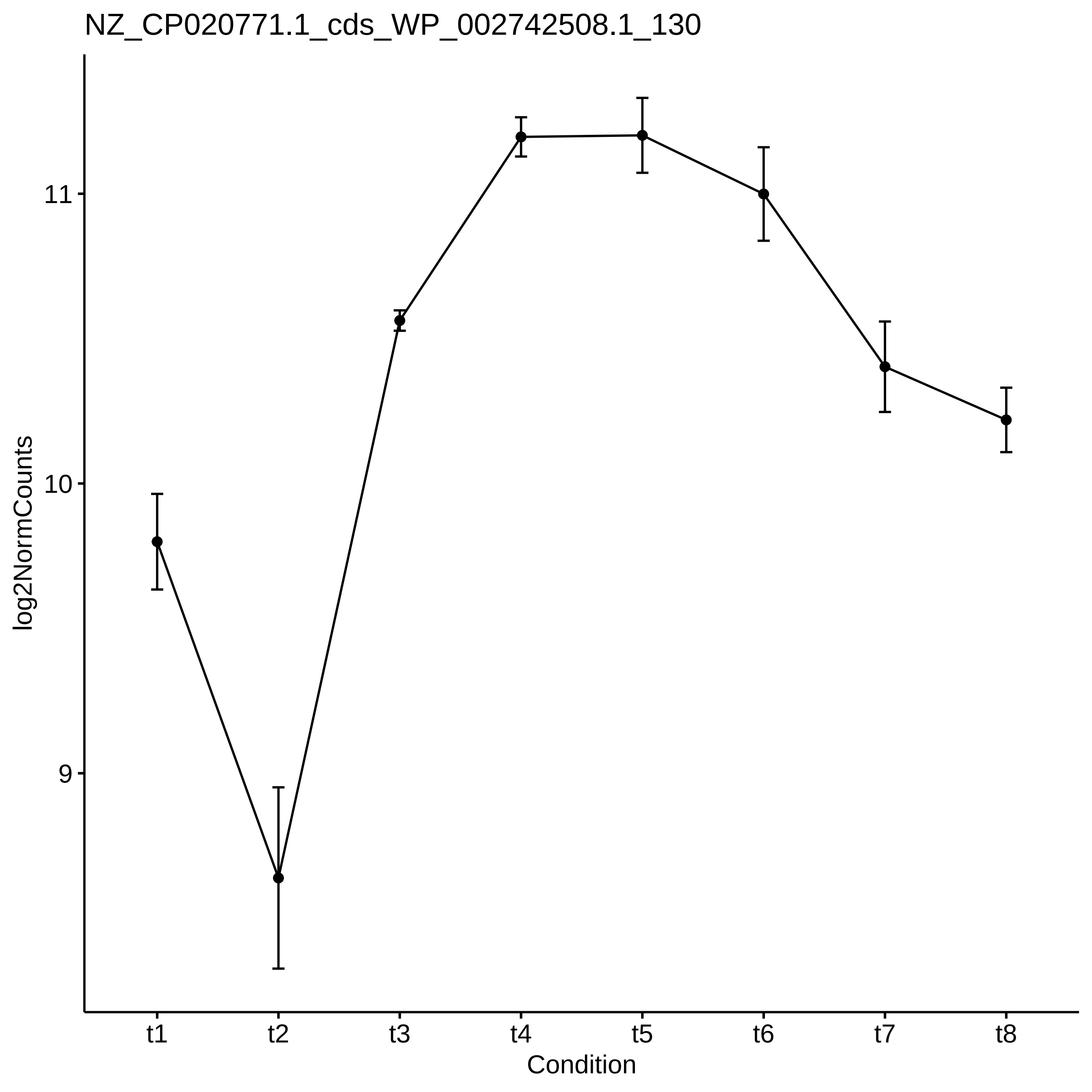

NZ_CP020771.1_cds_WP_002742508.1_130
log2NormCounts
11
10
9
t1
t2
t3
t4
t5
t6
t7
t8
Condition

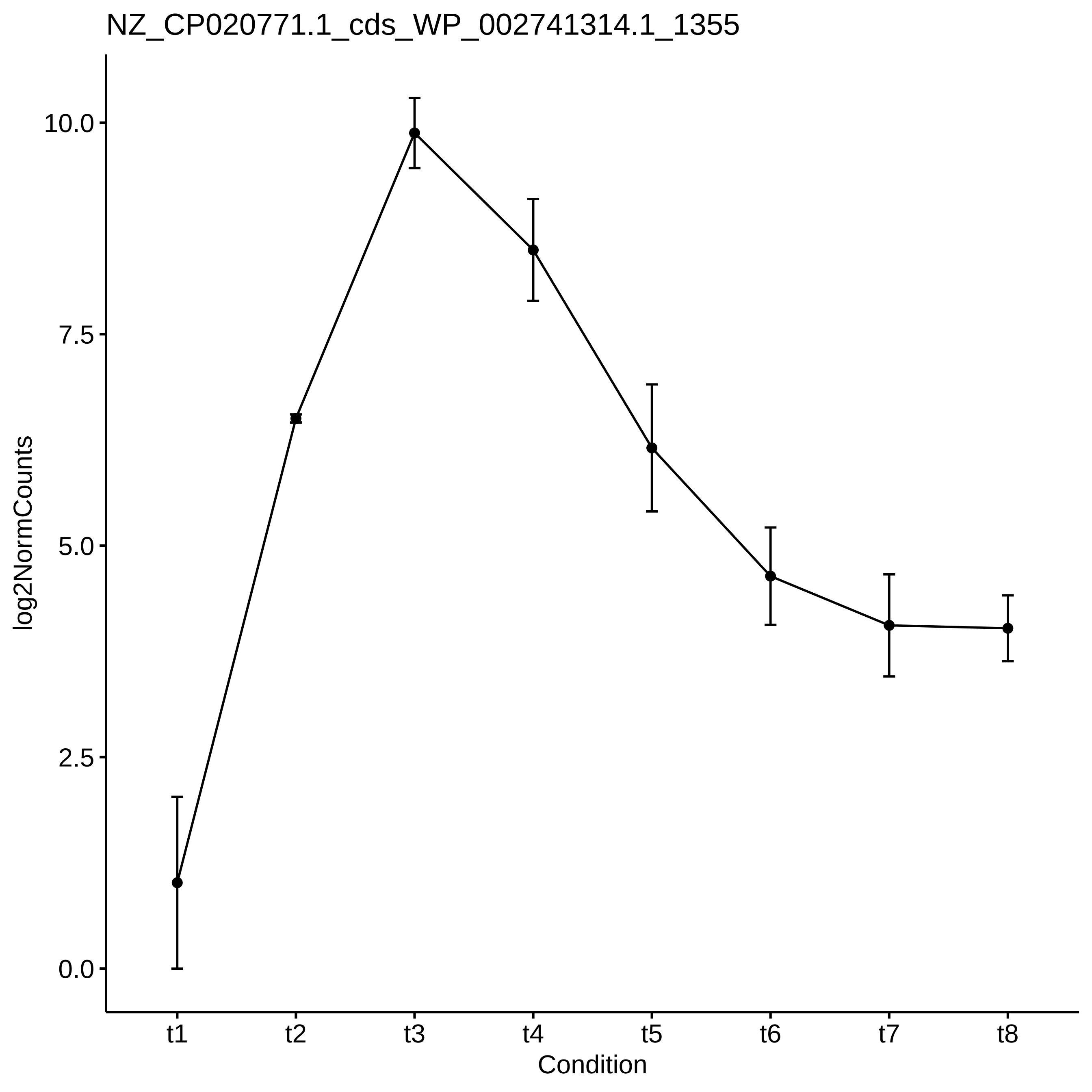
NZ_CP020771.1_cds_WP_002741314.1_1355
log2NormCounts
10.0
7.5
5.0
2.5
0.0
t1
t2
t3
t4
t5
t6
t7
t8
Condition

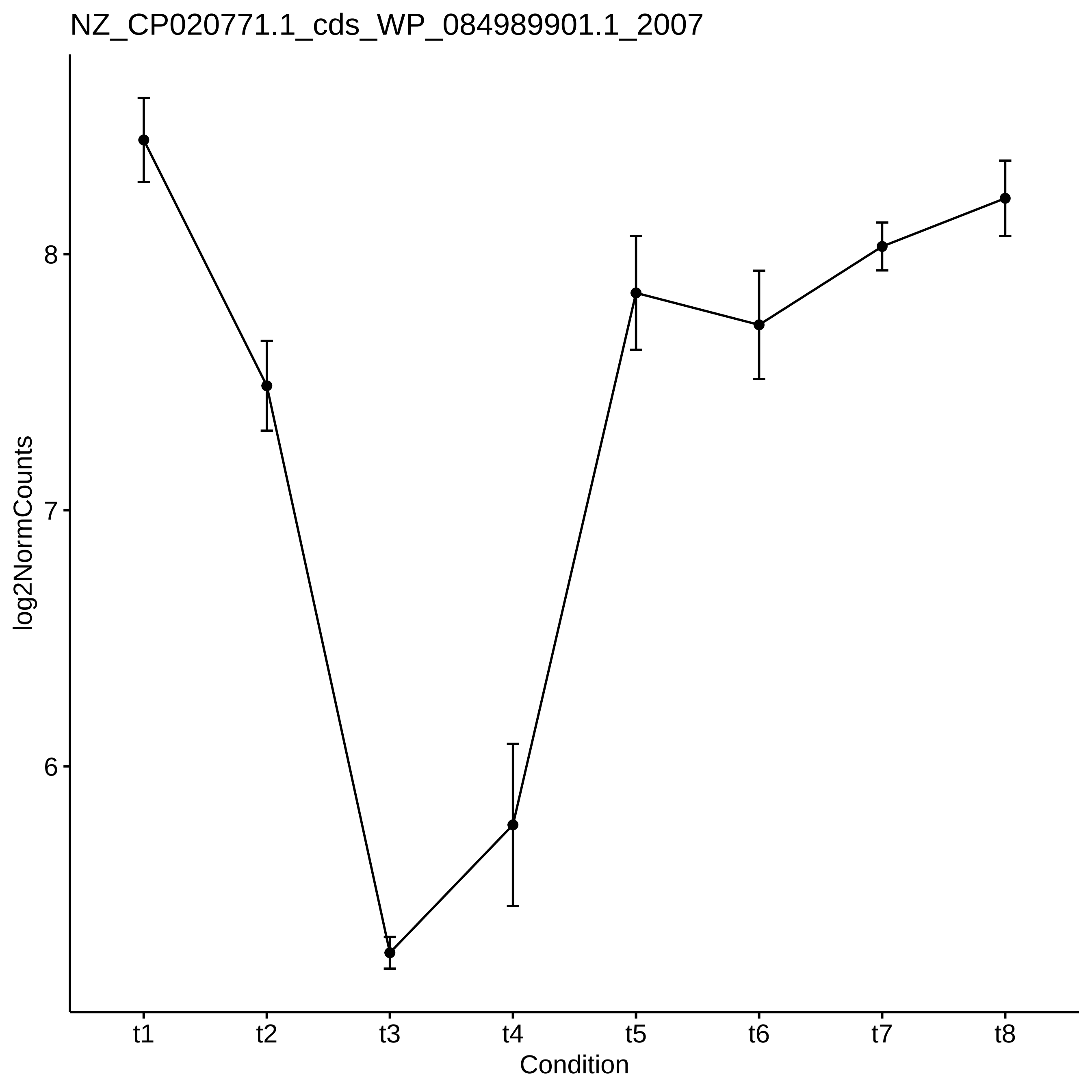
NZ_CP020771.1_cds_WP_084989901.1_2007
log2NormCounts
8
7
6
t1
t2
t3
t4
t5
t6
t7
t8
Condition

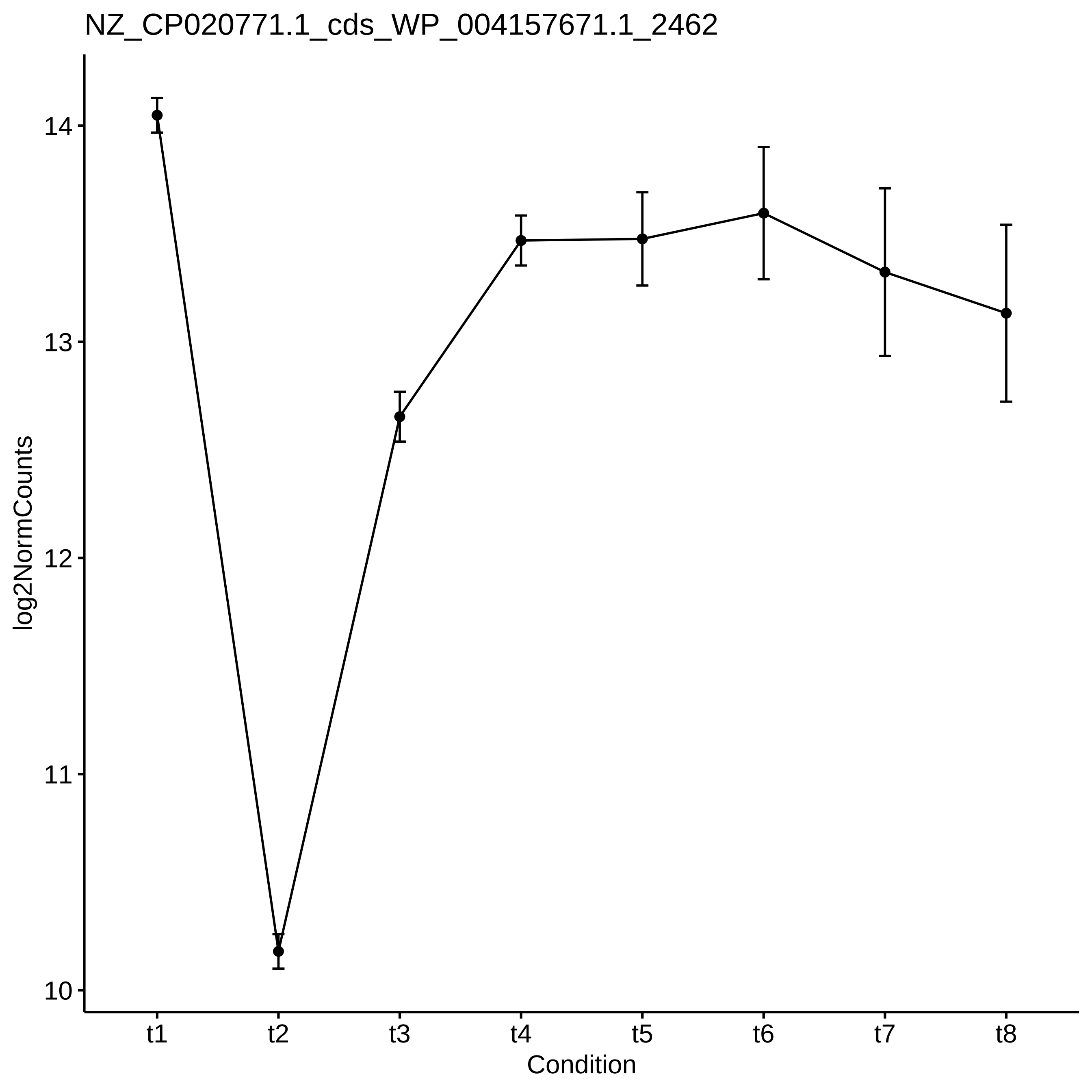
NZ_CP020771.1_cds_WP_004157671.1_2462
log2NormCounts
14
13
12
11
10
t1
t2
t3
t4
t5
t6
t7
t8
Condition

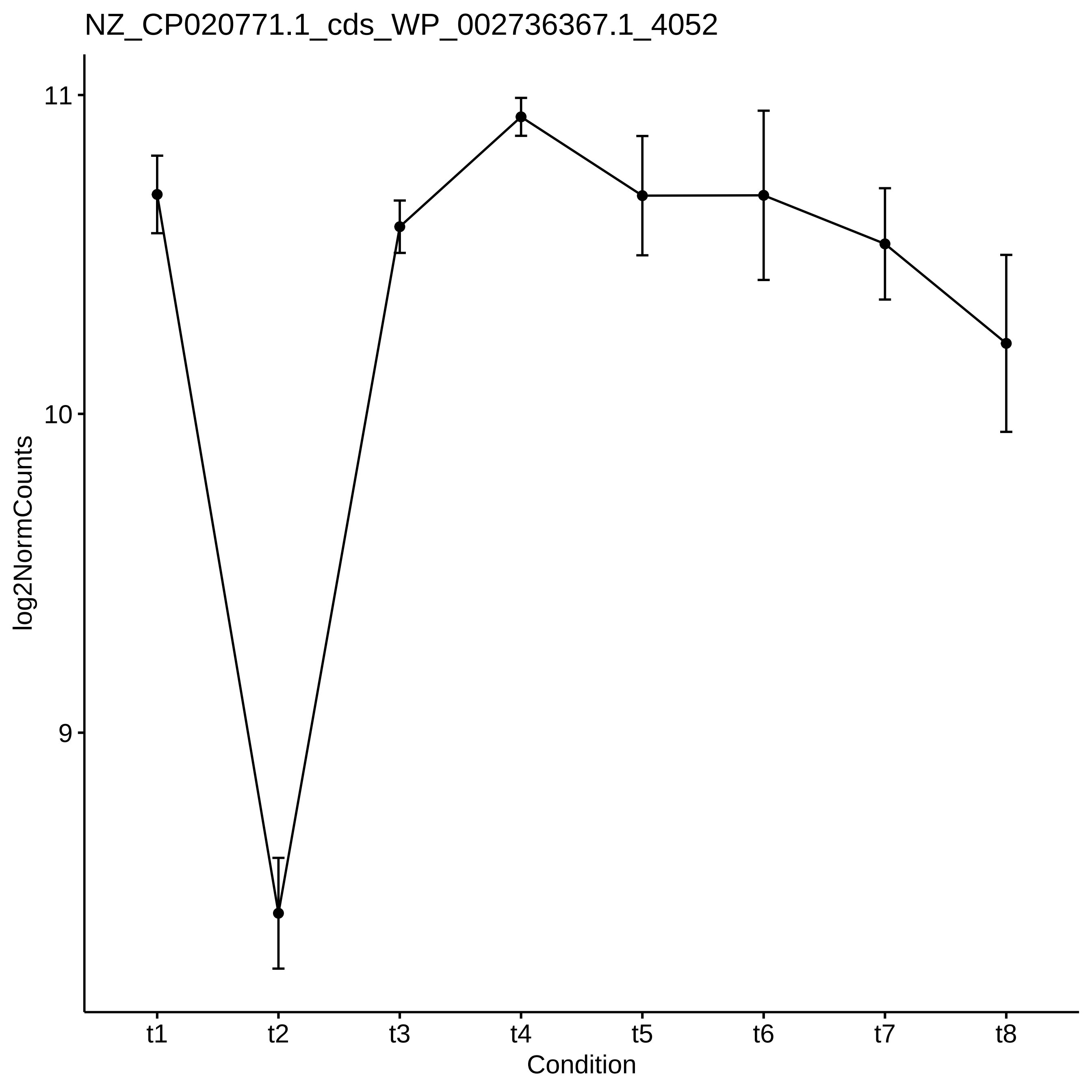
NZ_CP020771.1_cds_WP_002736367.1_4052
11
10
9
log2NormCounts
t1
t2
t3
t4
t5
t6
t7
t8
Condition

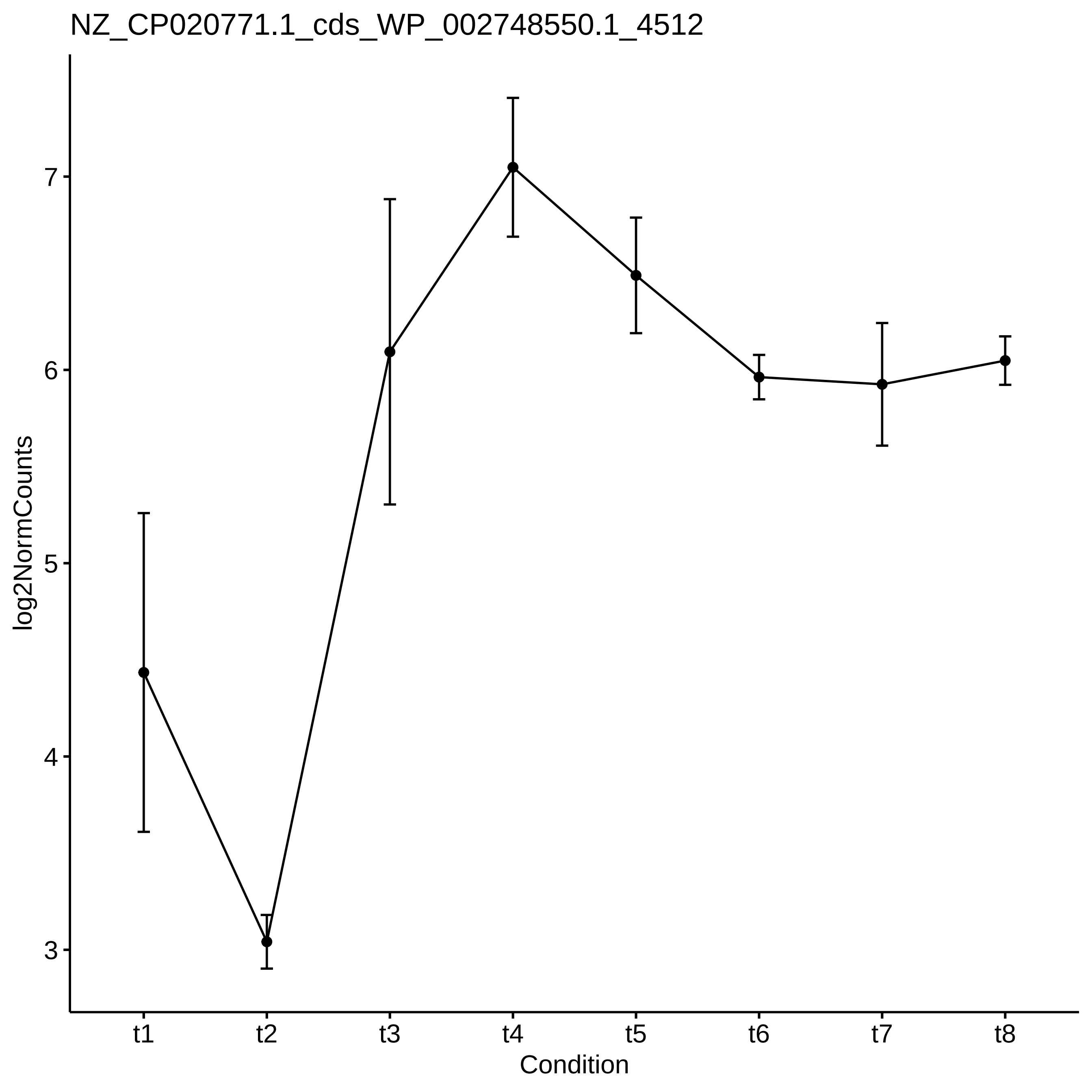

NZ_CP020771.1_cds_WP_002748550.1_4512
log2NormCounts
7
6
5
4
3
t1
t2
t3
t4
t5
t6
t7
t8
Condition

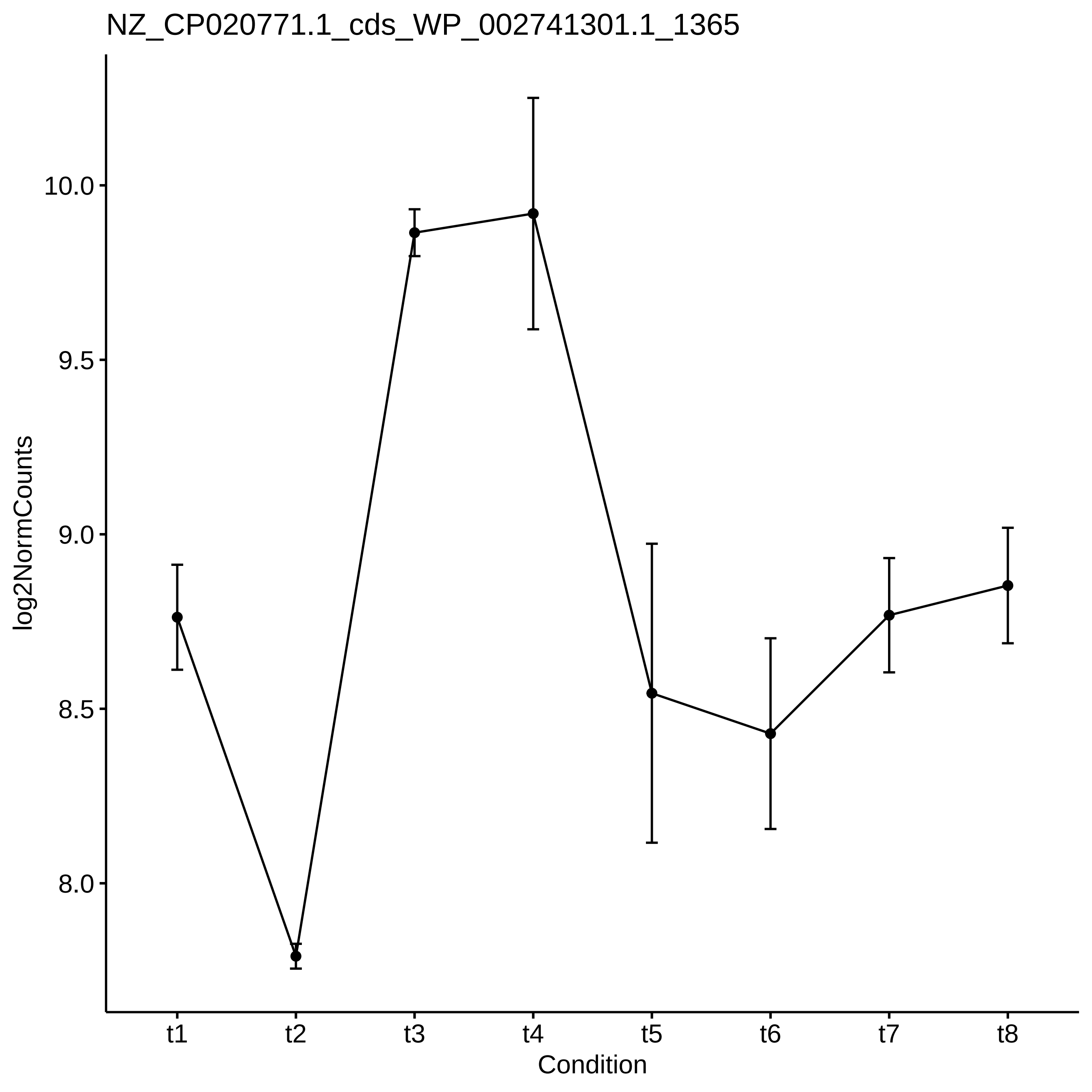
NZ_CP020771.1_cds_WP_002741301.1_1365
log2NormCounts
10.0
9.5
9.0
8.5
8.0
t1
t2
t3
t4
t5
t6
t7
t8
Condition

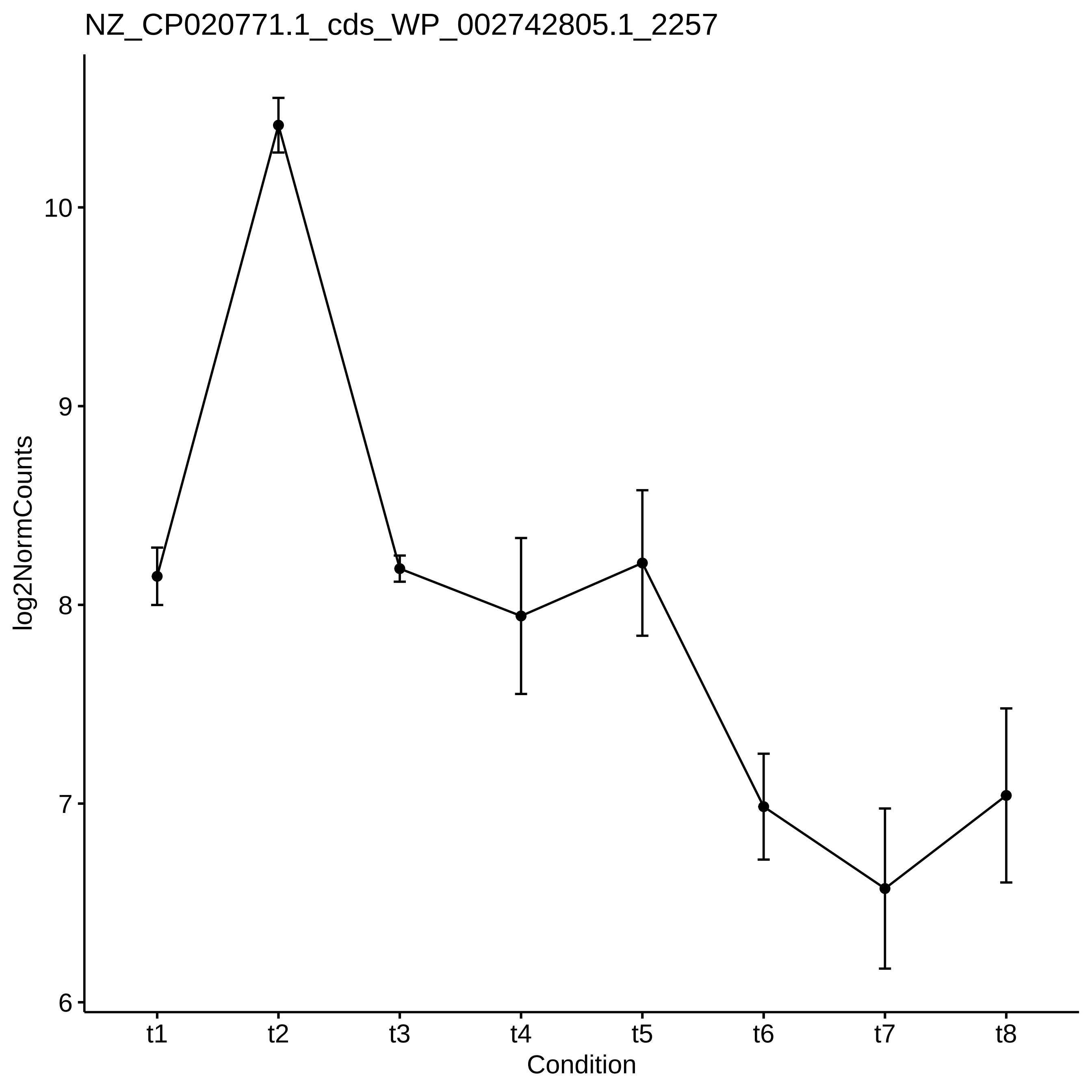
NZ_CP020771.1_cds_WP_002742805.1_2257
log2NormCounts
10
9
8
7
6
t1
t2
t3
t4
t5
t6
t7
t8
Condition

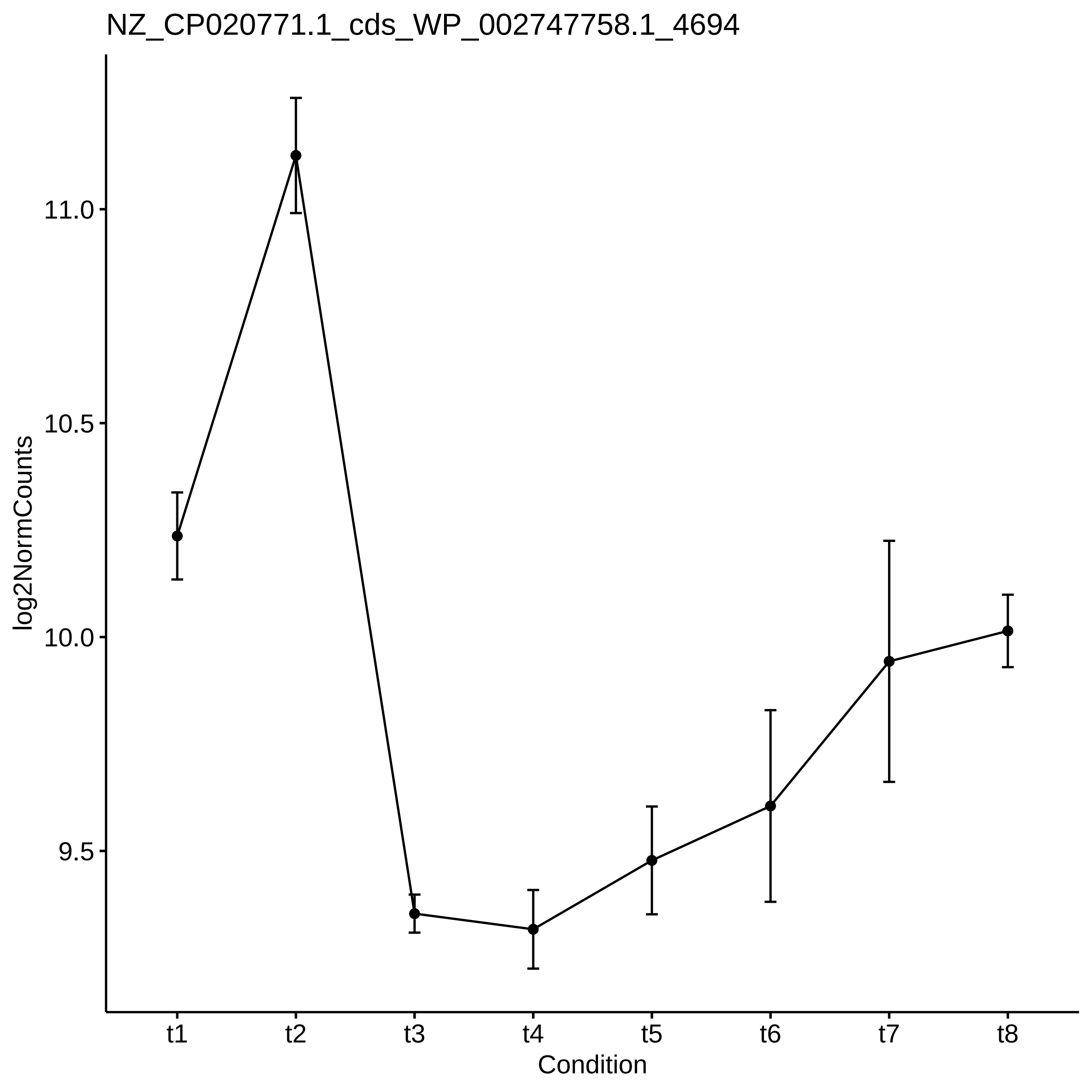
NZ_CP020771.1_cds_WP_002747758.1_4694
log2NormCounts
11.0
10.5
10.0
9.5
t1
t2
t3
t4
t5
t6
t7
t8
Condition

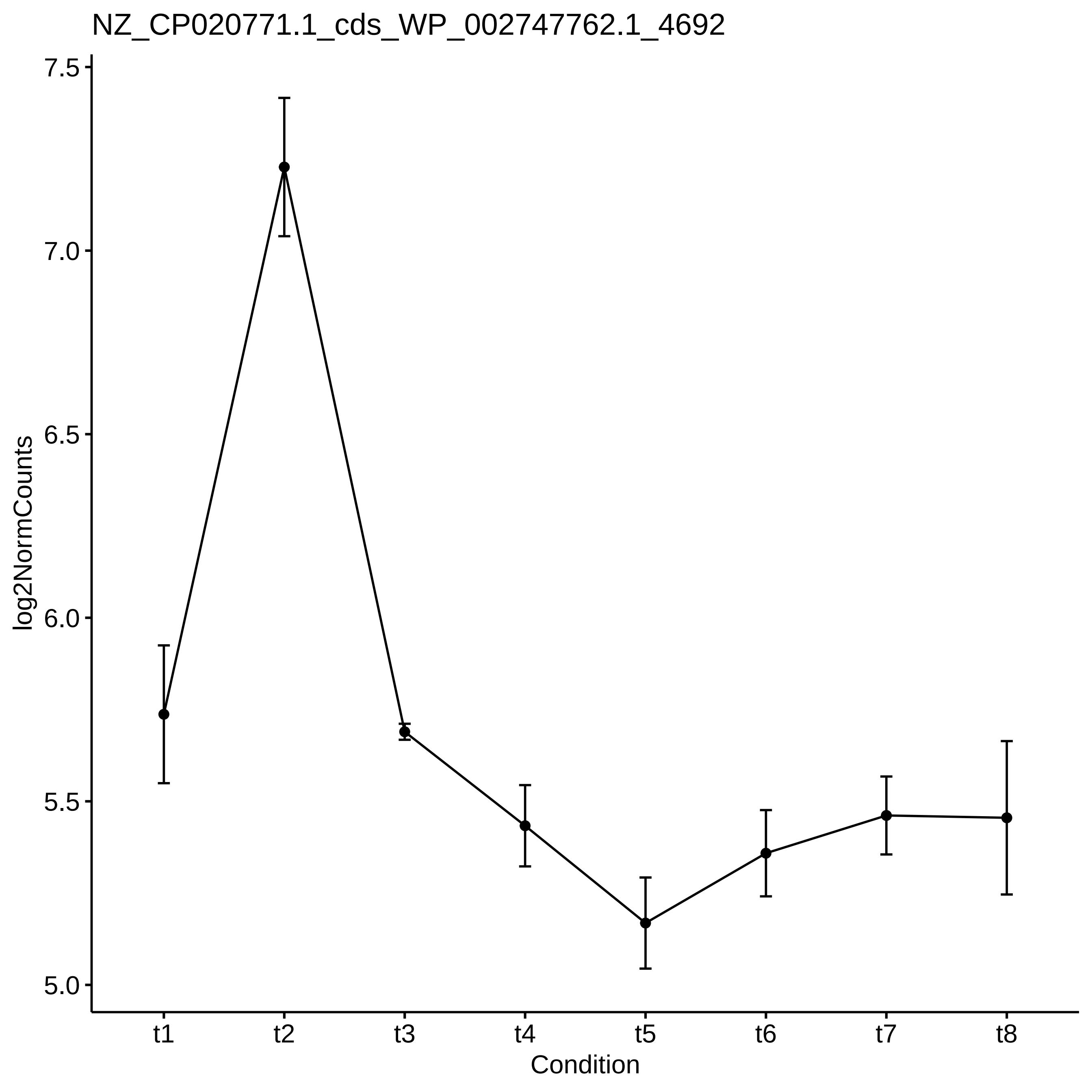
NZ_CP020771.1_cds_WP_002747762.1_4692
7.5
7.0
6.5
6.0
5.5
5.0
log2NormCounts
t1
t2
t3
t4
t5
t6
t7
t8
Condition

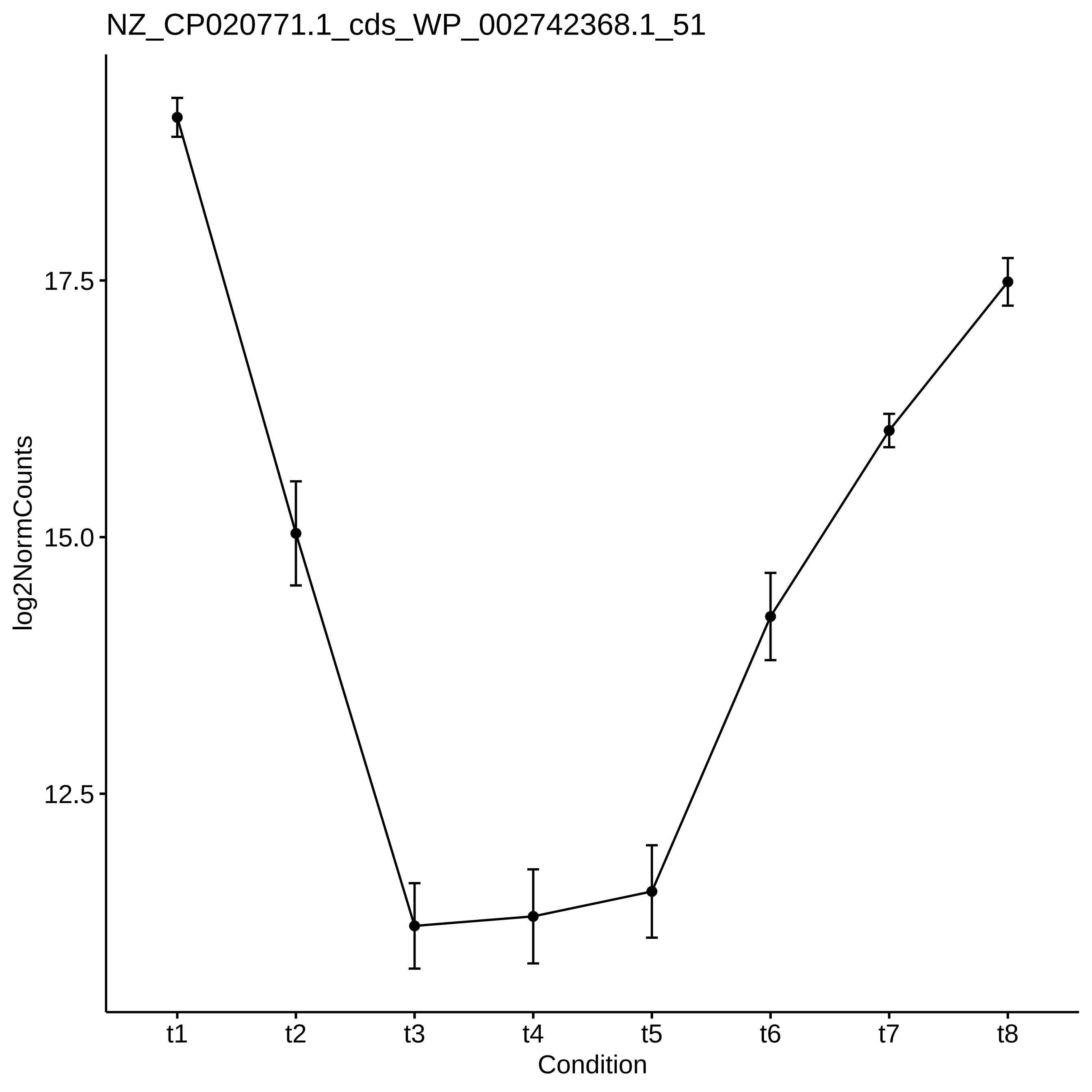
NZ_CP020771.1_cds_WP_002742368.1_51
log2NormCounts
17.5
15.0
12.5
t1
t2
t3
t4
t5
t6
t7
t8
Condition

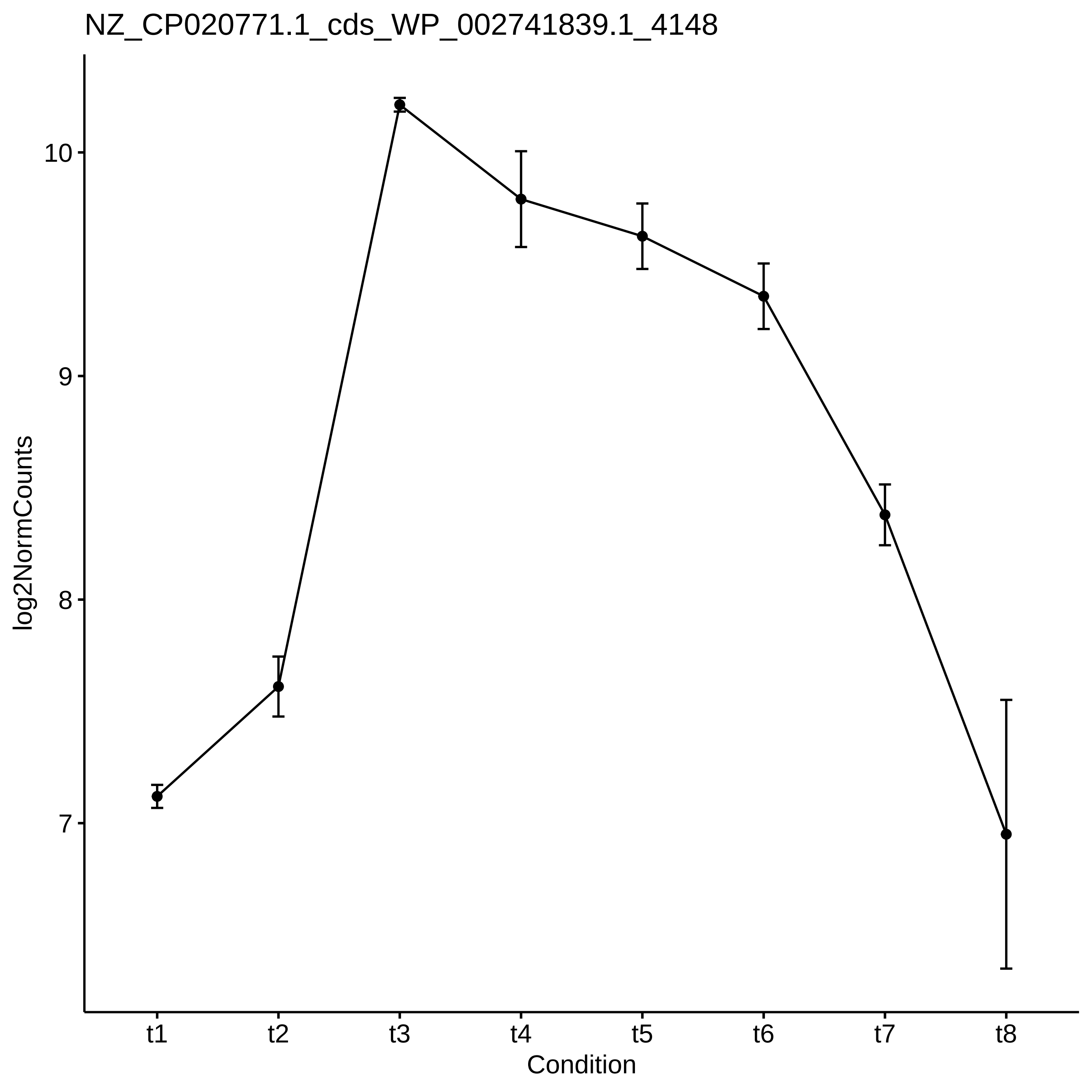

NZ_CP020771.1_cds_WP_002741839.1_4148
log2NormCounts
10
9
8
7
t1
t2
t3
t4
t5
t6
t7
t8
Condition

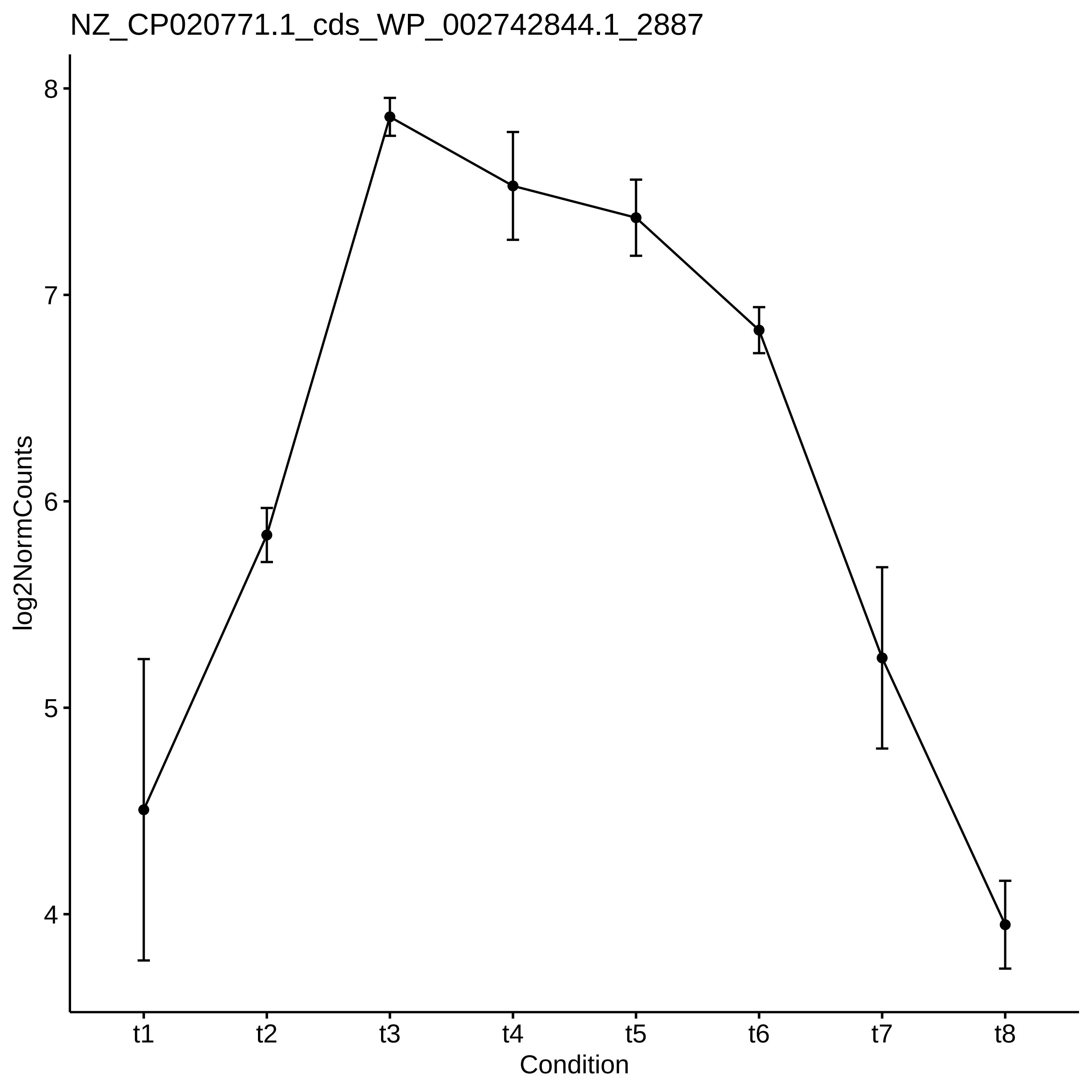
NZ_CP020771.1_cds_WP_002742844.1_2887
log2NormCounts
8
7
6
5
4
t1
t2
t3
t4
t5
t6
t7
t8
Condition

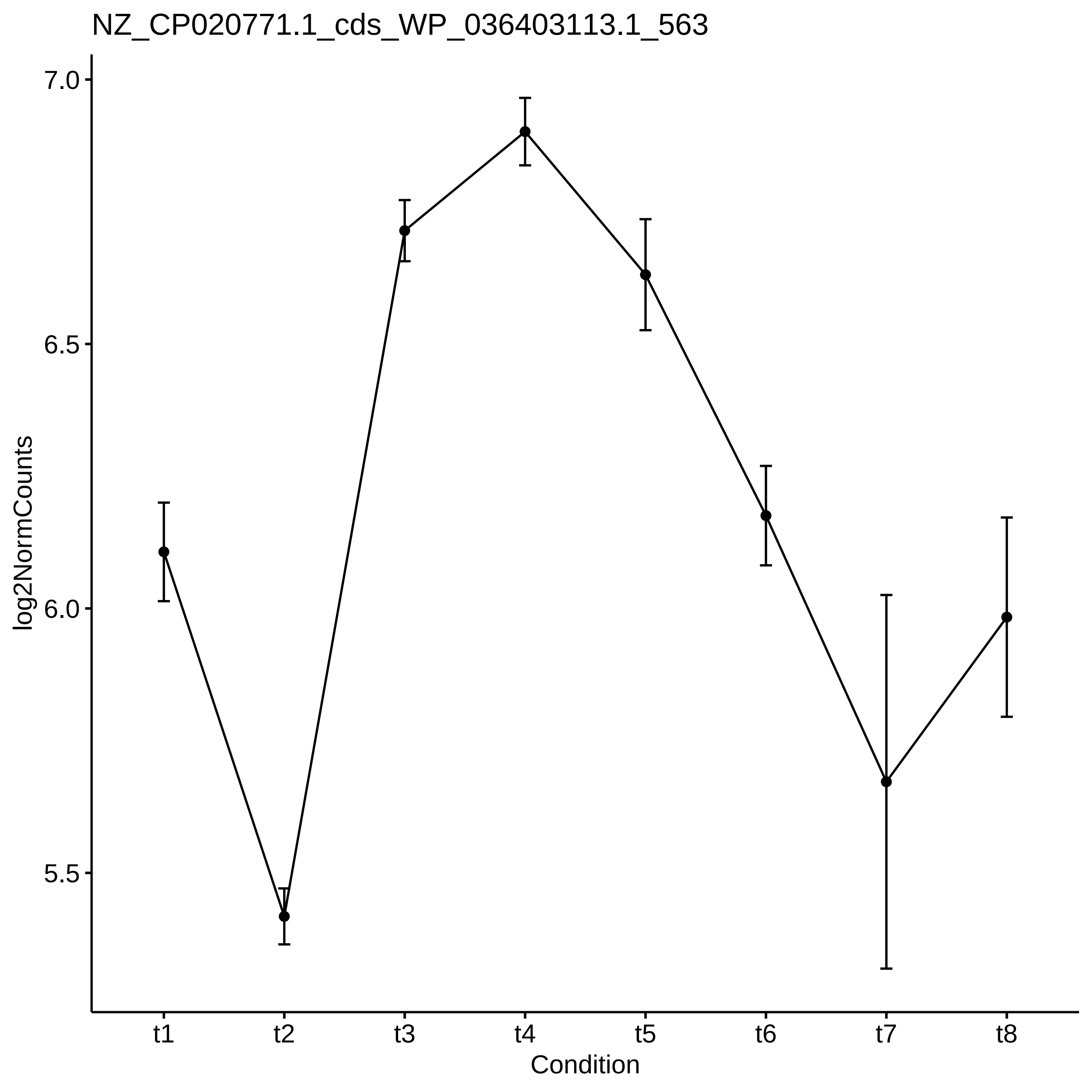
NZ_CP020771.1_cds_WP_036403113.1_563
log2NormCounts
7.0
6.5
6.0
5.5
t1
t2
t3
t4
t5
t6
t7
t8
Condition

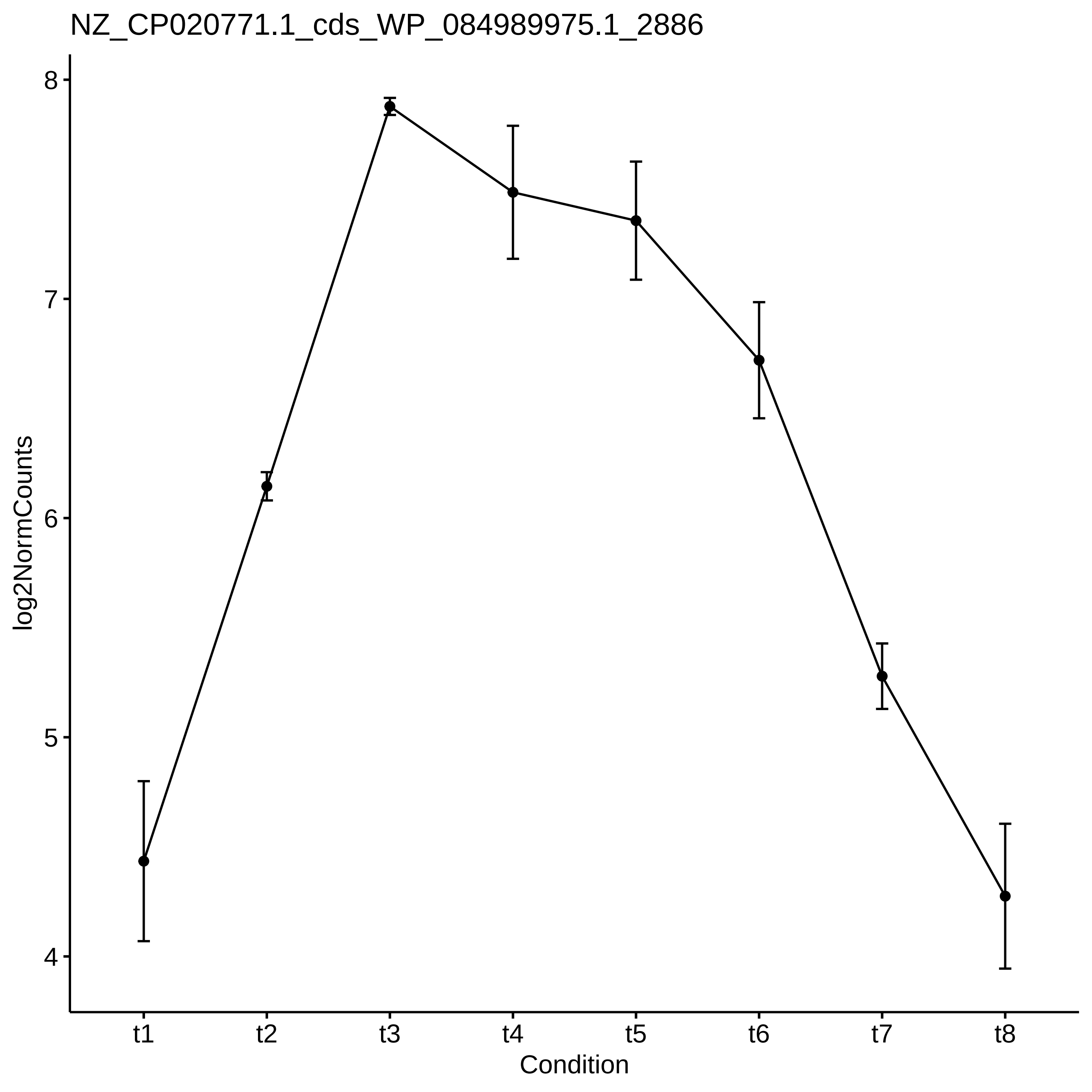

NZ_CP020771.1_cds_WP_084989975.1_2886
log2NormCounts
8
7
6
5
4
t1
t2
t3
t4
t5
t6
t7
t8
Condition

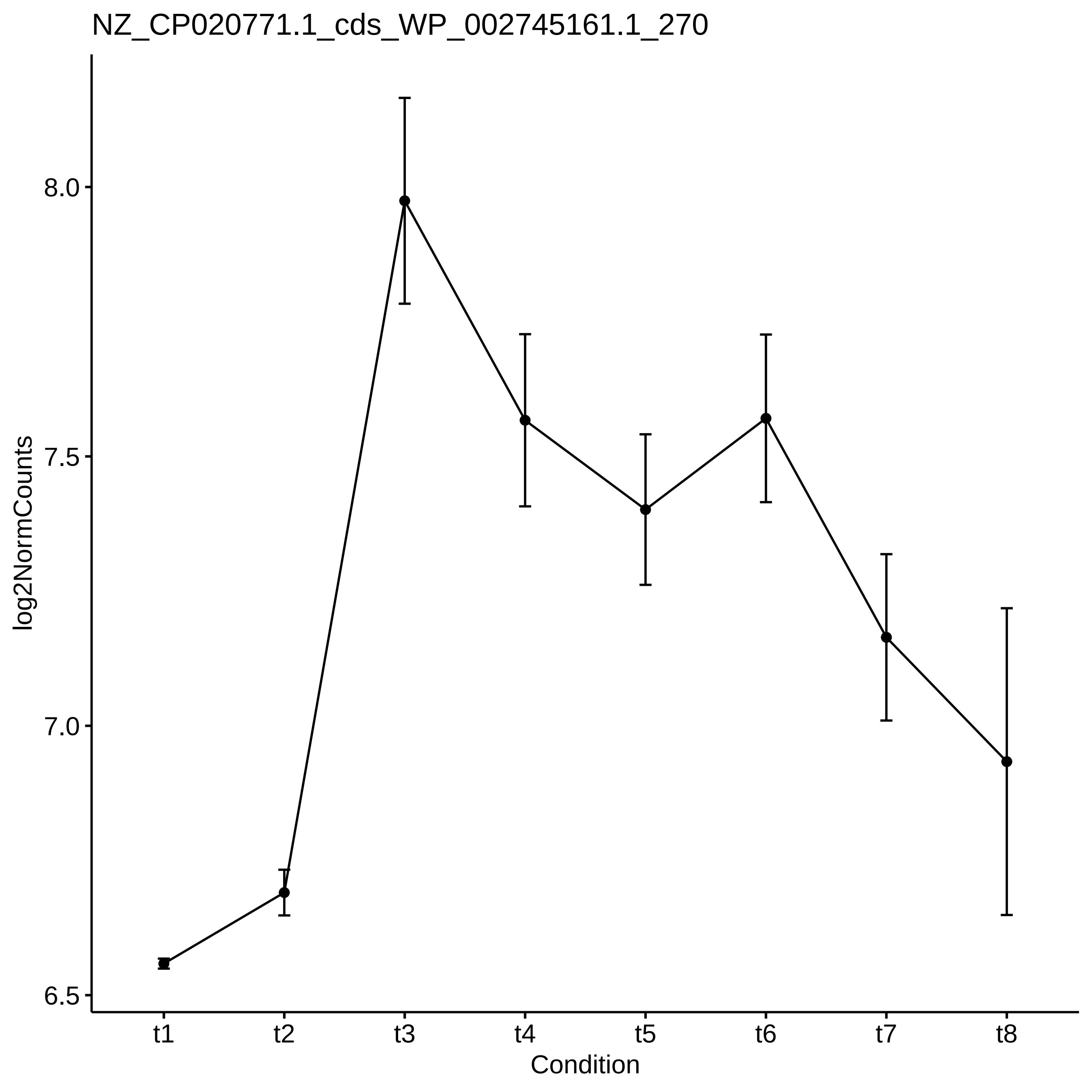
NZ_CP020771.1_cds_WP_002745161.1_270
log2NormCounts
8.0
7.5
7.0
6.5
t1
t2
t3
t4
t5
t6
t7
t8
Condition

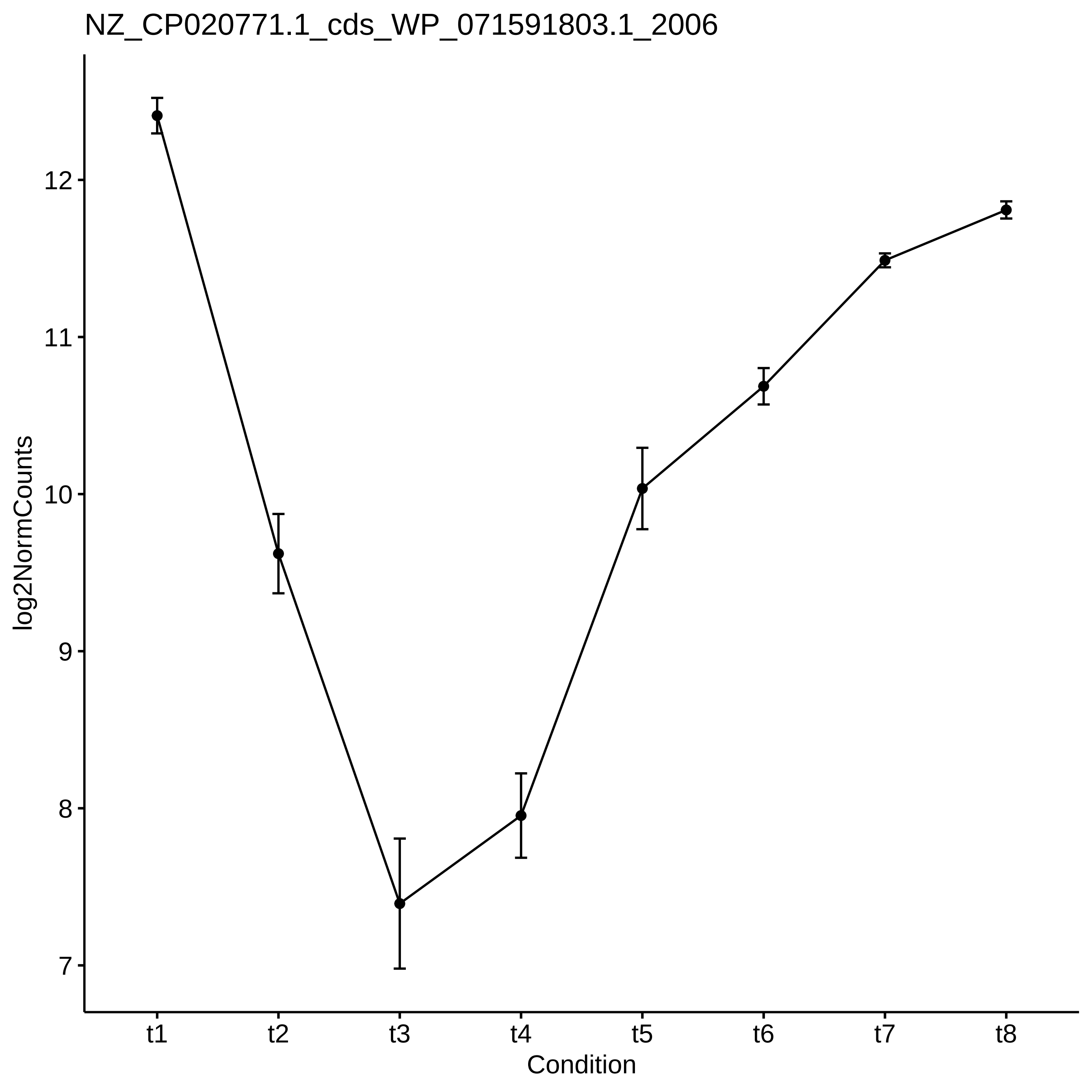
NZ_CP020771.1_cds_WP_071591803.1_2006
log2NormCounts
12
11
10
9
8
7
t1
t2
t3
t4
t5
t6
t7
t8
Condition

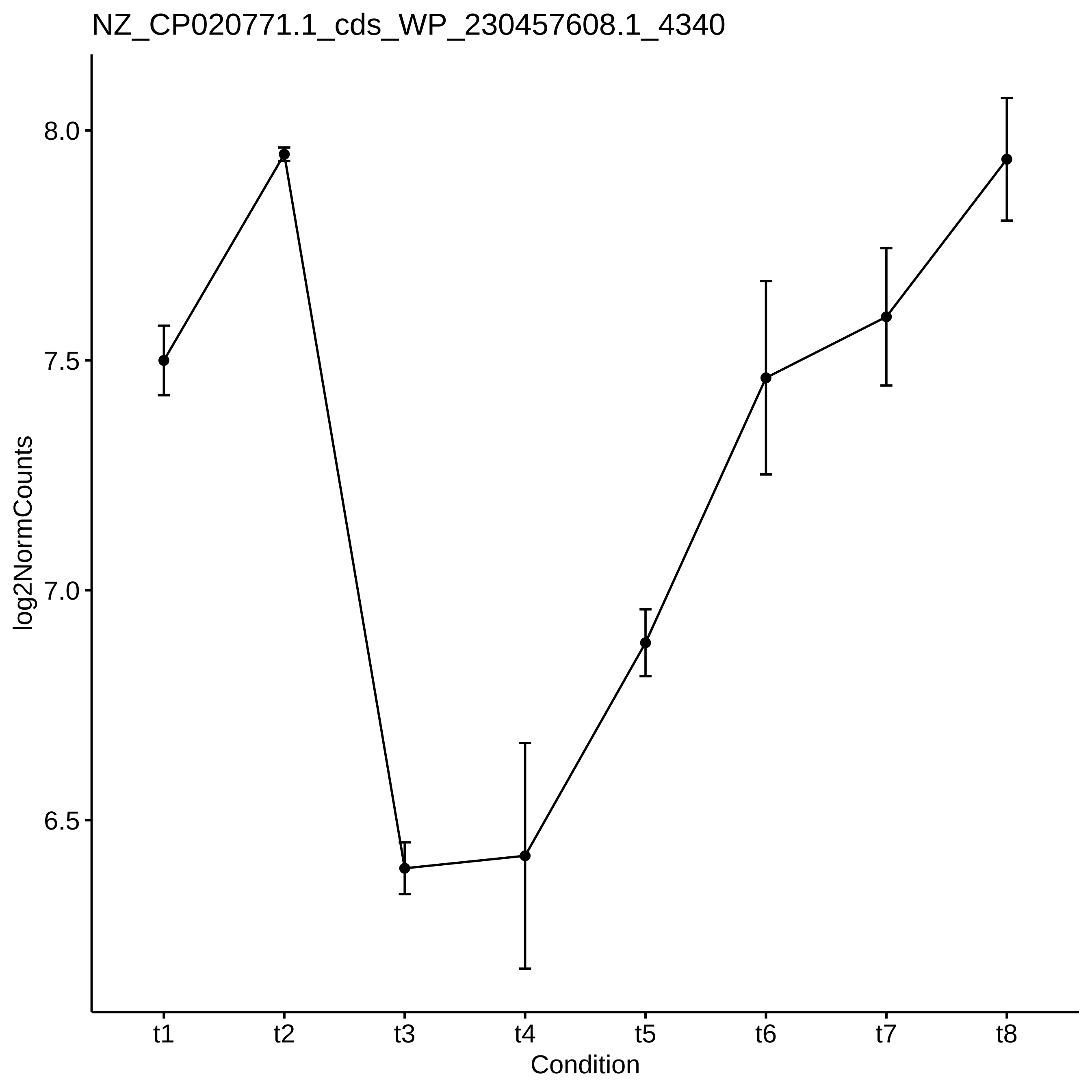
NZ_CP020771.1_cds_WP_230457608.1_4340
log2NormCounts
8.0
7.5
7.0
6.5
t1
t2
t3
t4
t5
t6
t7
t8
Condition

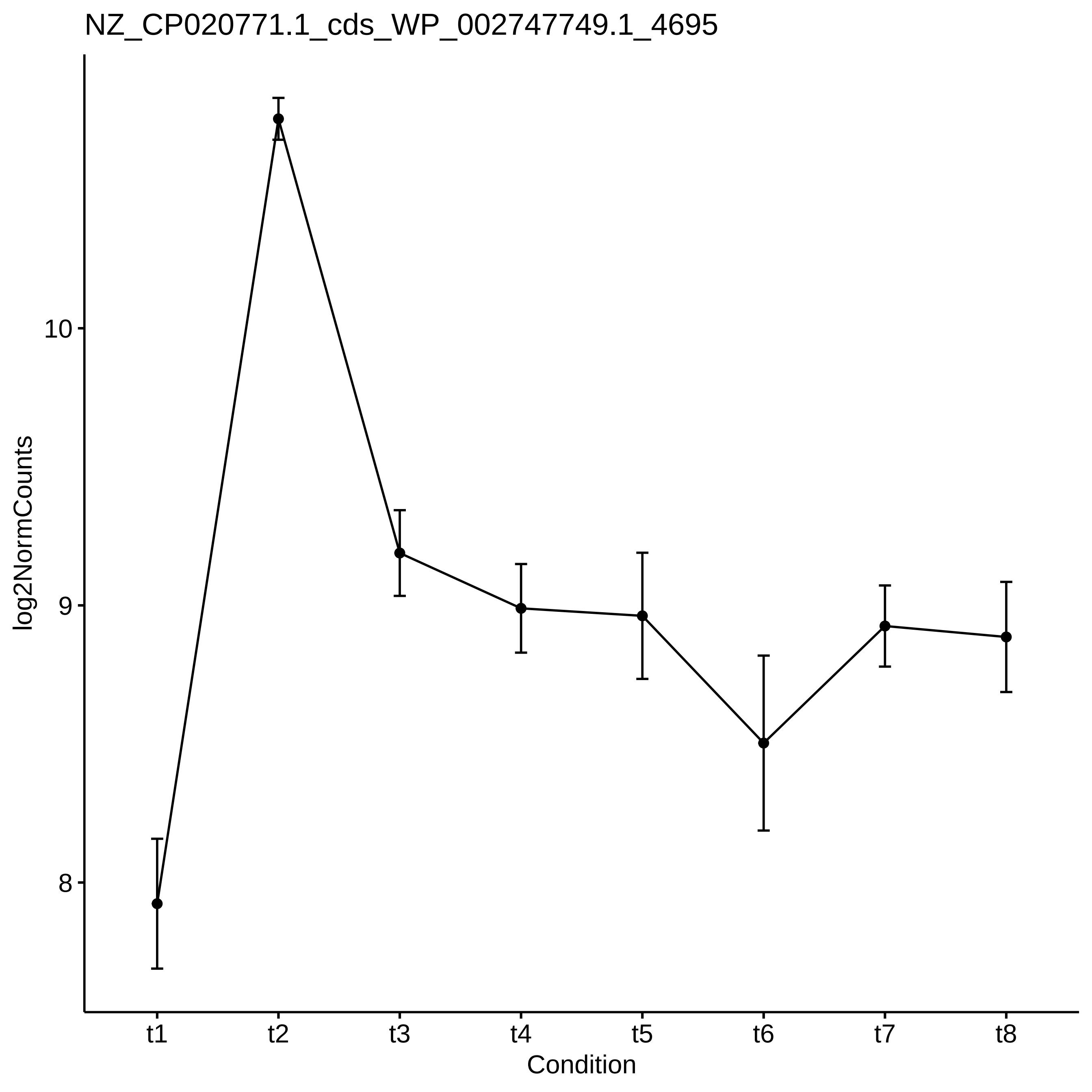

NZ_CP020771.1_cds_WP_002747749.1_4695
log2NormCounts
10
9
8
t1
t2
t3
t4
t5
t6
t7
t8
Condition

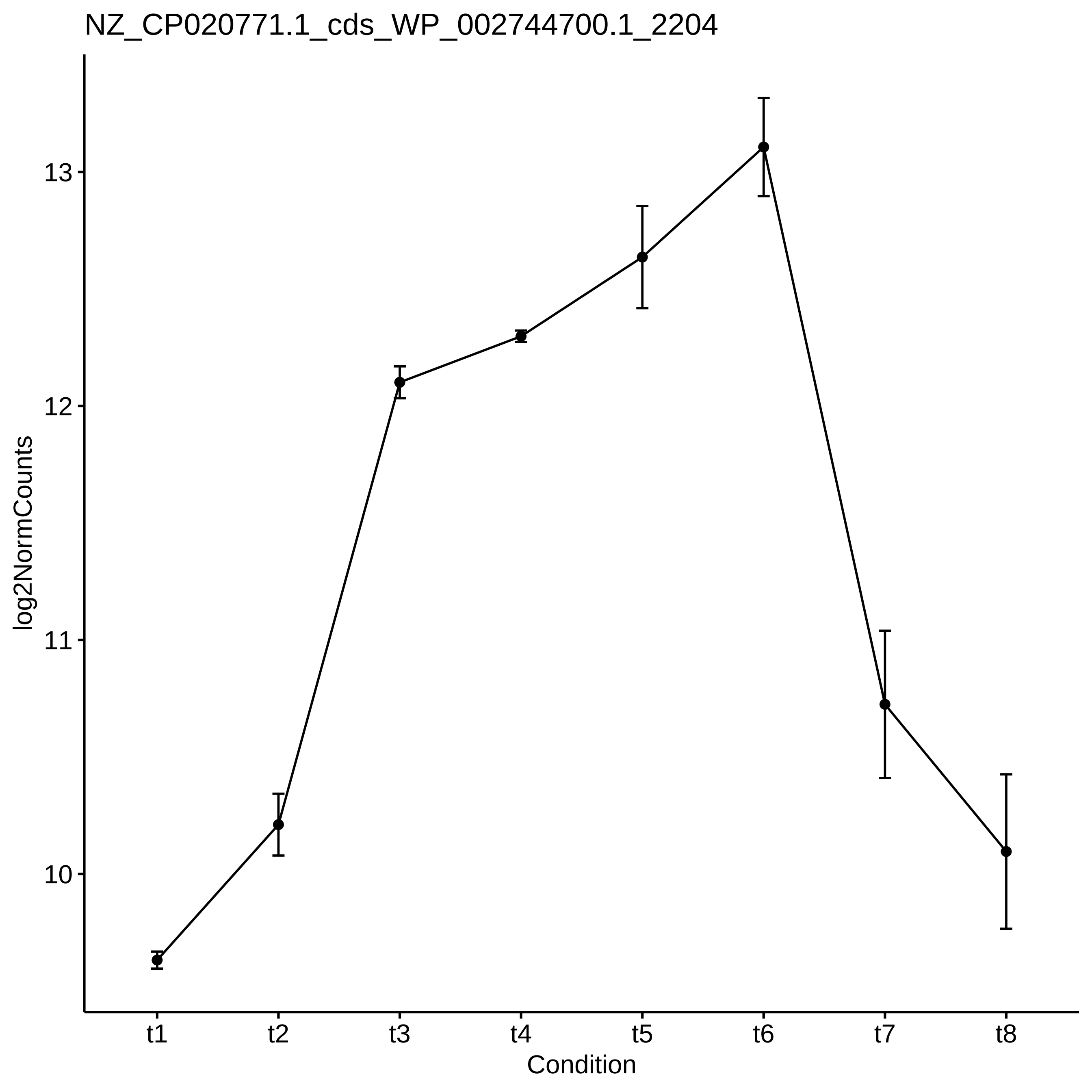
NZ_CP020771.1_cds_WP_002744700.1_2204
log2NormCounts
13
12
11
10
t1
t2
t3
t4
t5
t6
t7
t8
Condition

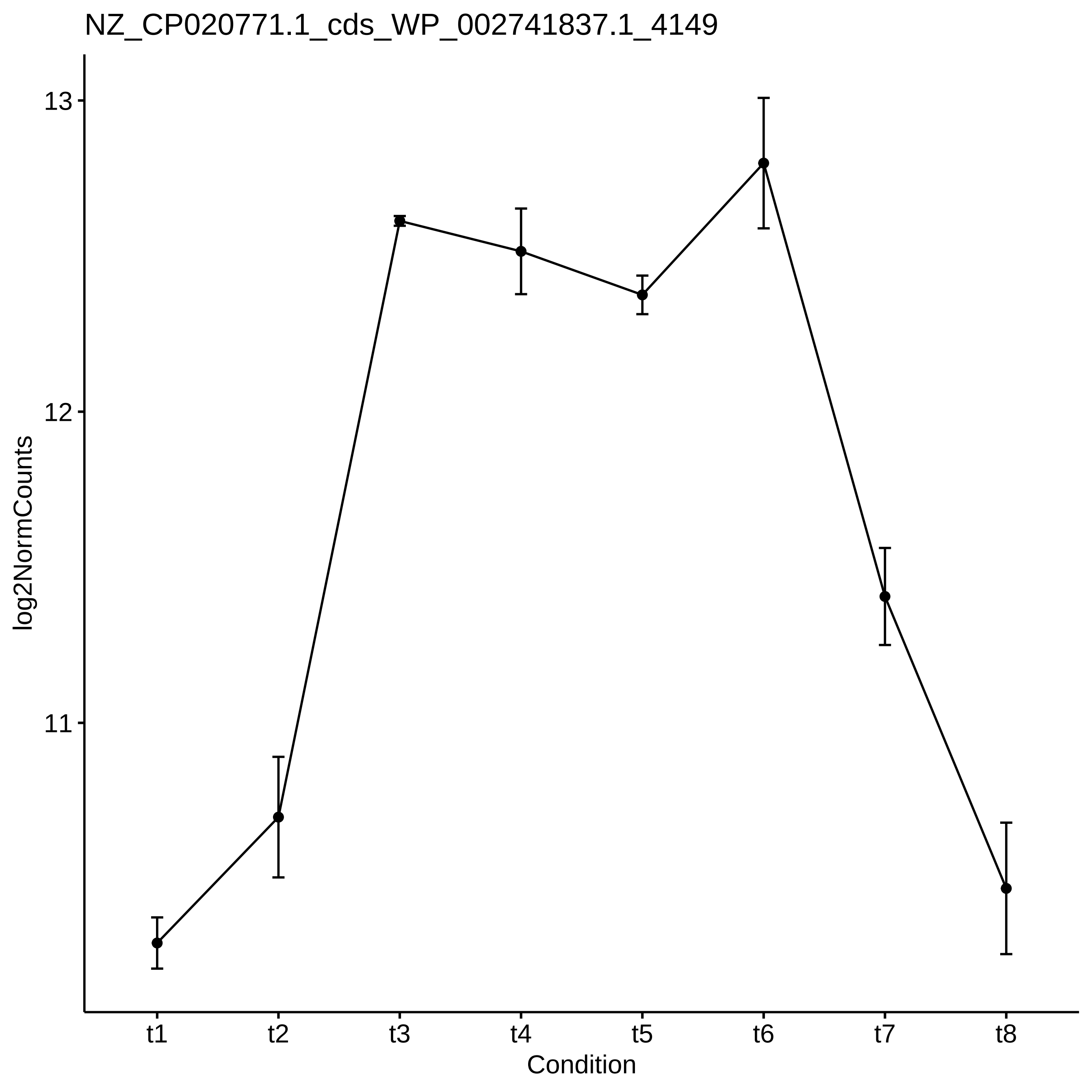
NZ_CP020771.1_cds_WP_002741837.1_4149
13
12
11
log2NormCounts
t1
t2
t3
t4
t5
t6
t7
t8
Condition

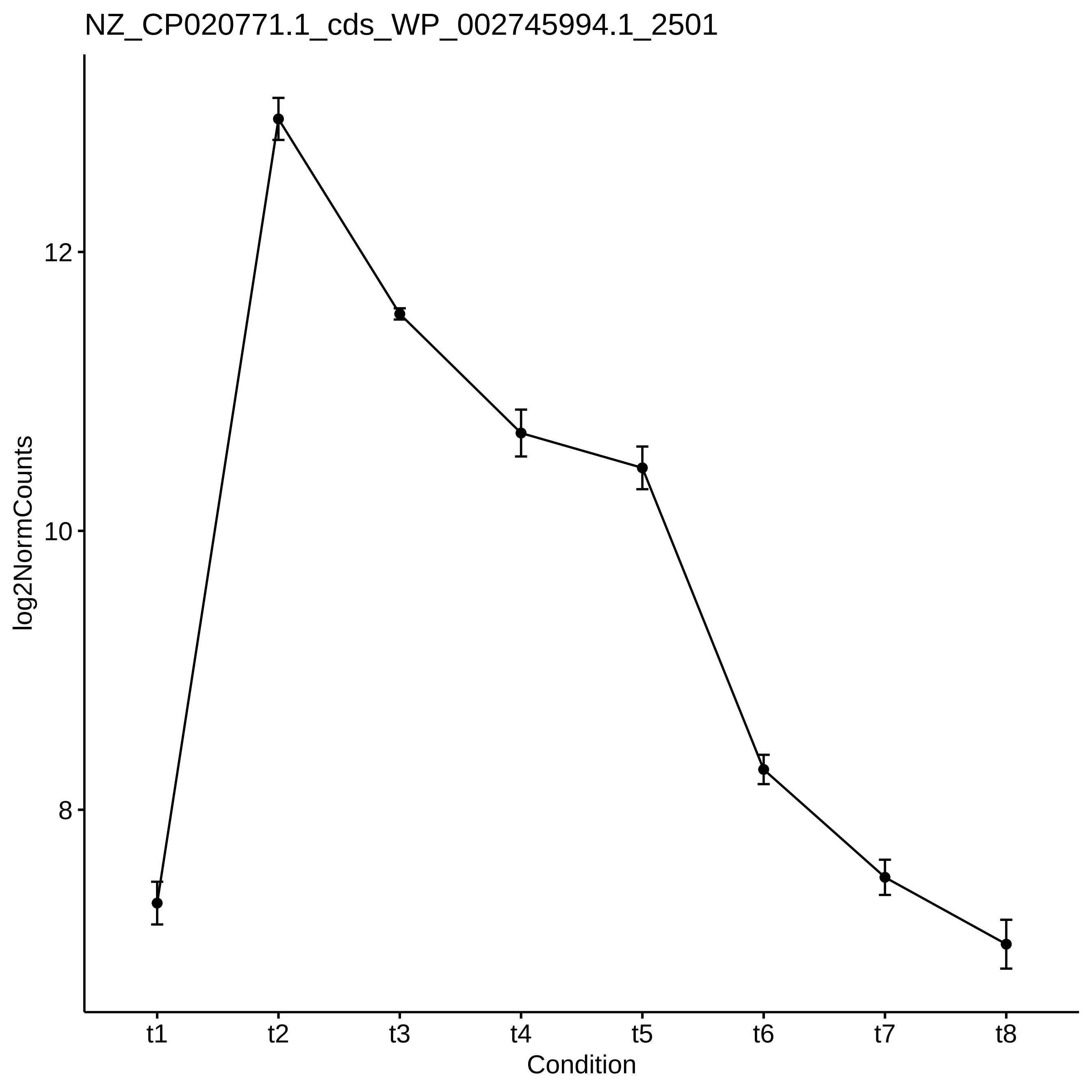
NZ_CP020771.1_cds_WP_002745994.1_2501
log2NormCounts
12
10
8
t1
t2
t3
t4
t5
t6
t7
t8
Condition

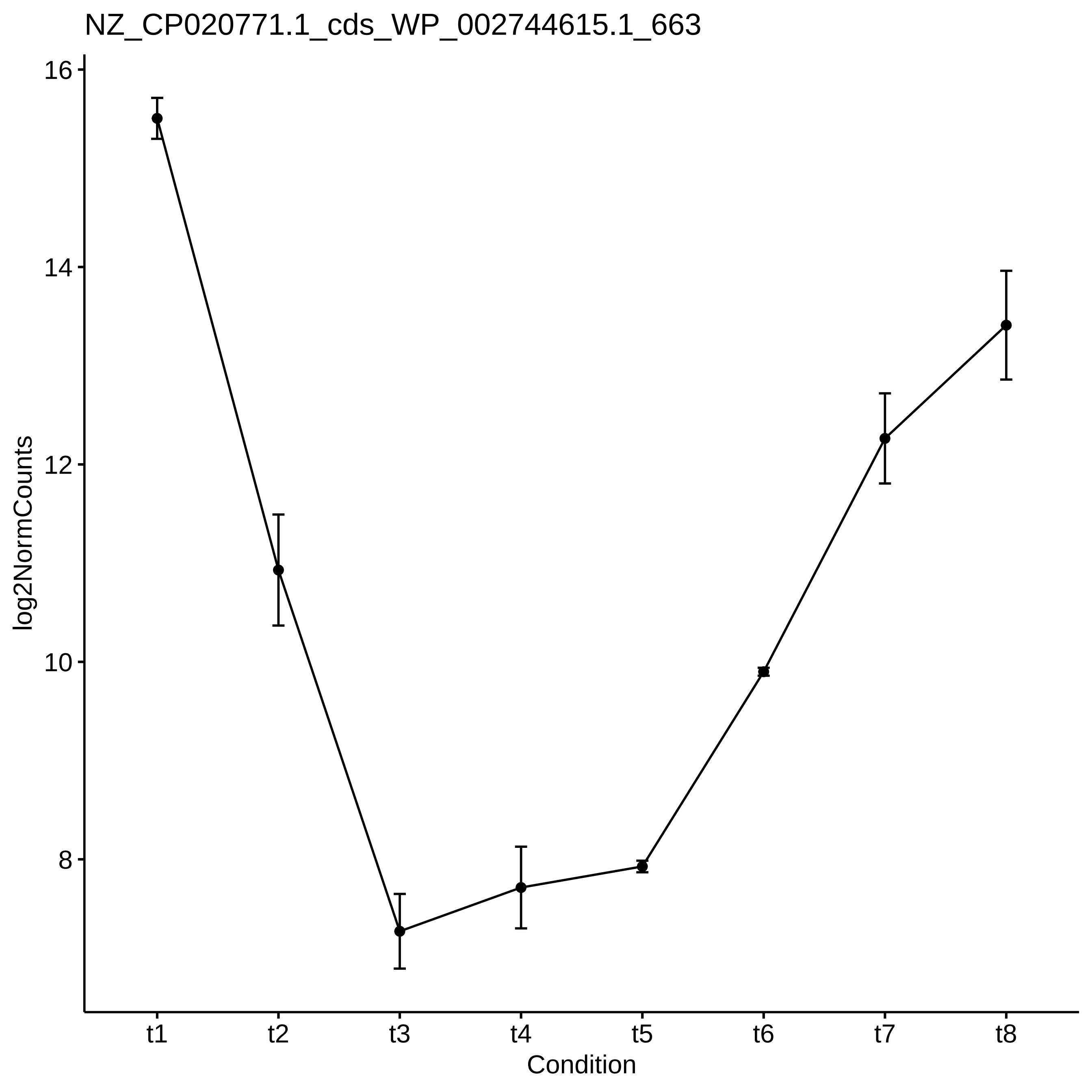
NZ_CP020771.1_cds_WP_002744615.1_663
log2NormCounts
16
14
12
10
8
t1
t2
t3
t4
t5
t6
t7
t8
Condition

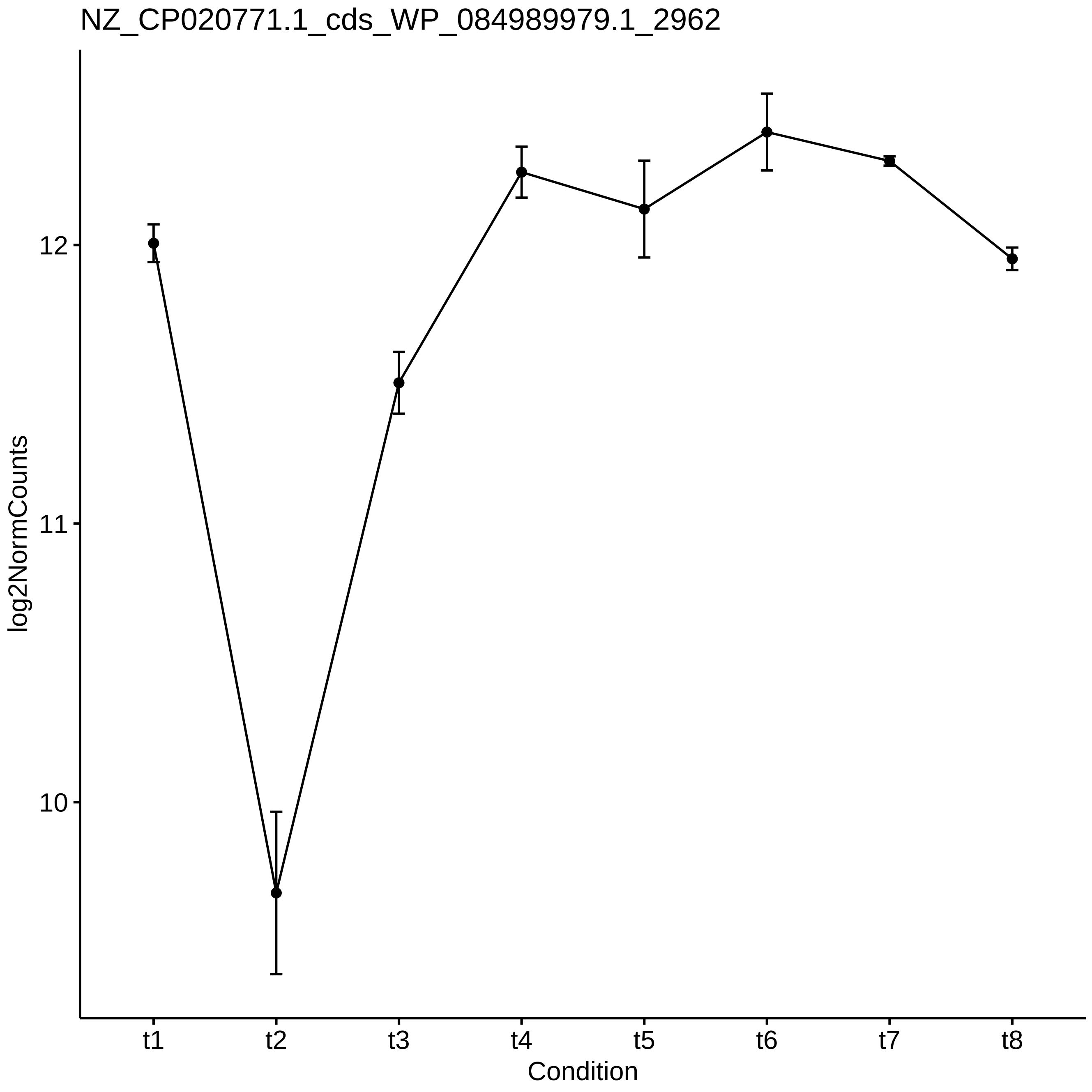
NZ_CP020771.1_cds_WP_084989979.1_2962
log2NormCounts
12
11
10
t1
t2
t3
t4
t5
t6
t7
t8
Condition

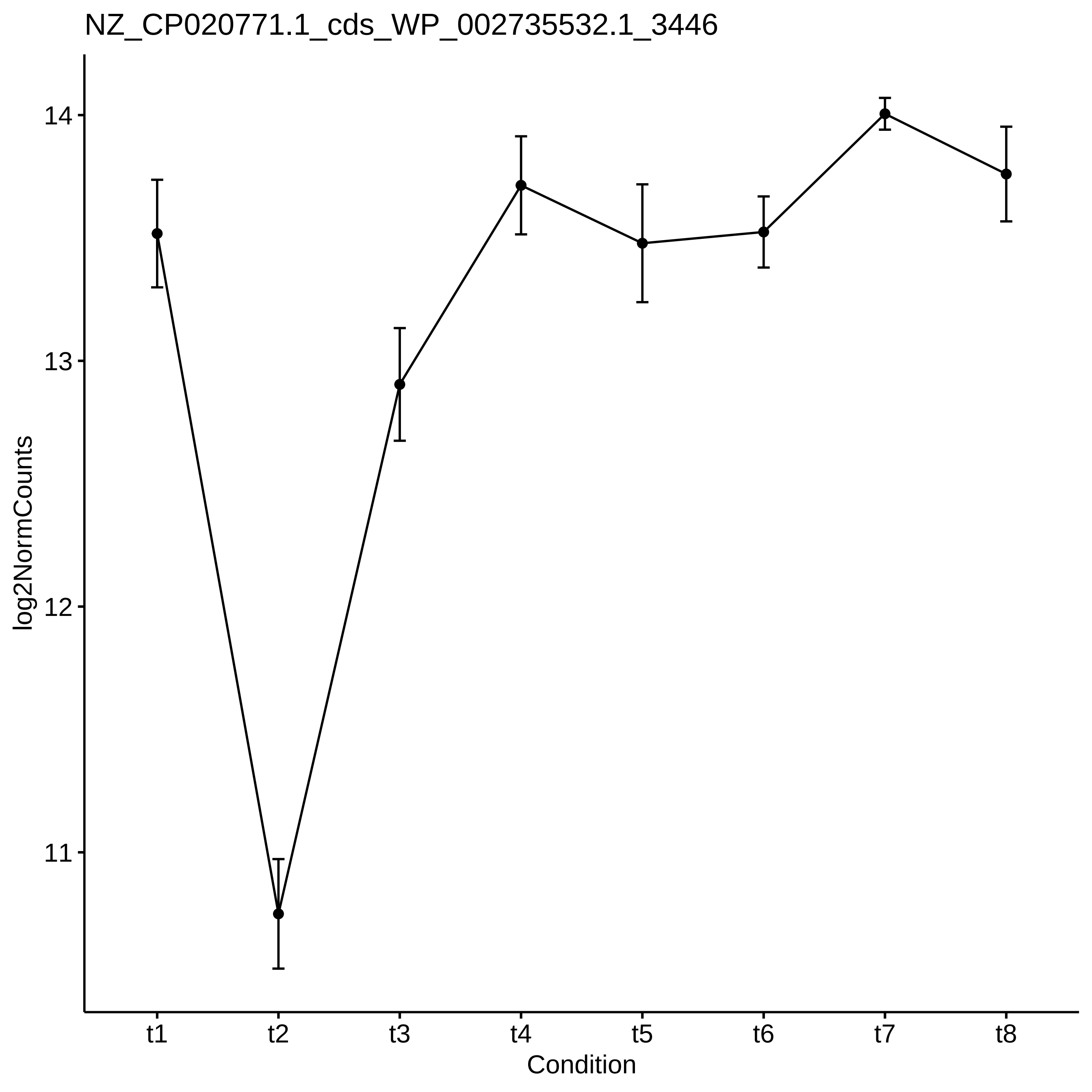
NZ_CP020771.1_cds_WP_002735532.1_3446
log2NormCounts
14
13
12
11
t1
t2
t3
t4
t5
t6
t7
t8
Condition

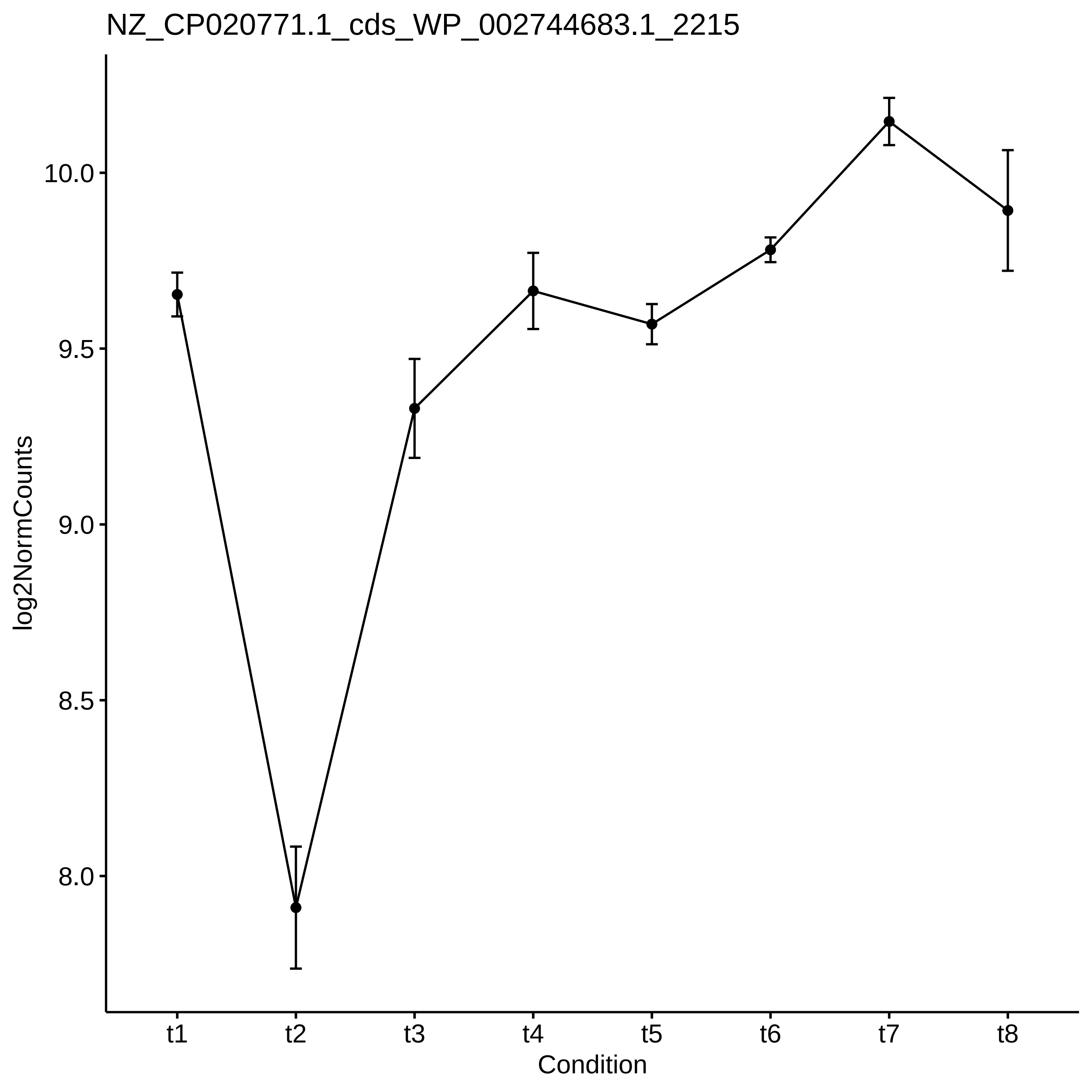
NZ_CP020771.1_cds_WP_002744683.1_2215
log2NormCounts
10.0
9.5
9.0
8.5
8.0
t1
t2
t3
t4
t5
t6
t7
t8
Condition

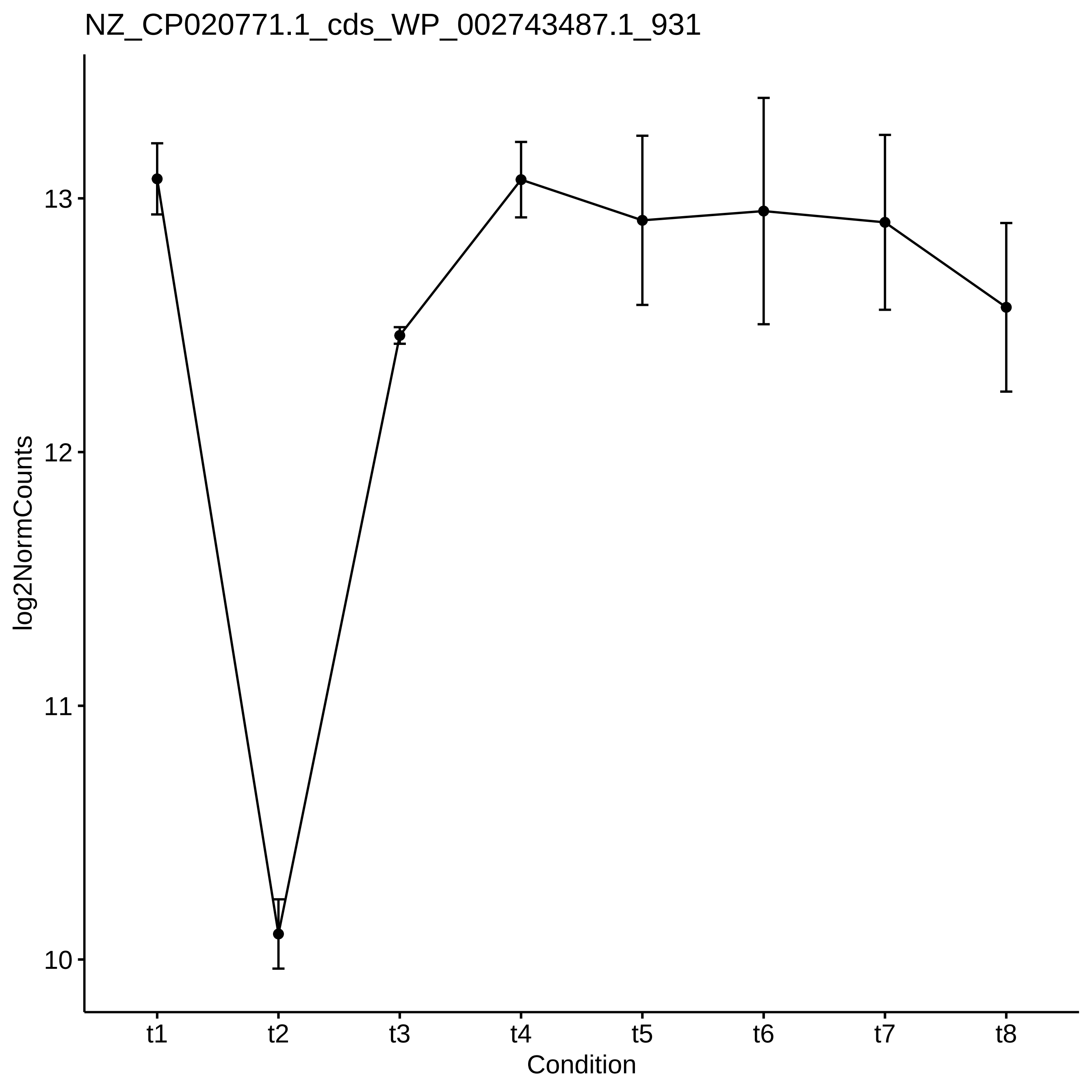
NZ_CP020771.1_cds_WP_002743487.1_931
log2NormCounts
13
12
11
10
t1
t2
t3
t4
t5
t6
t7
t8
Condition

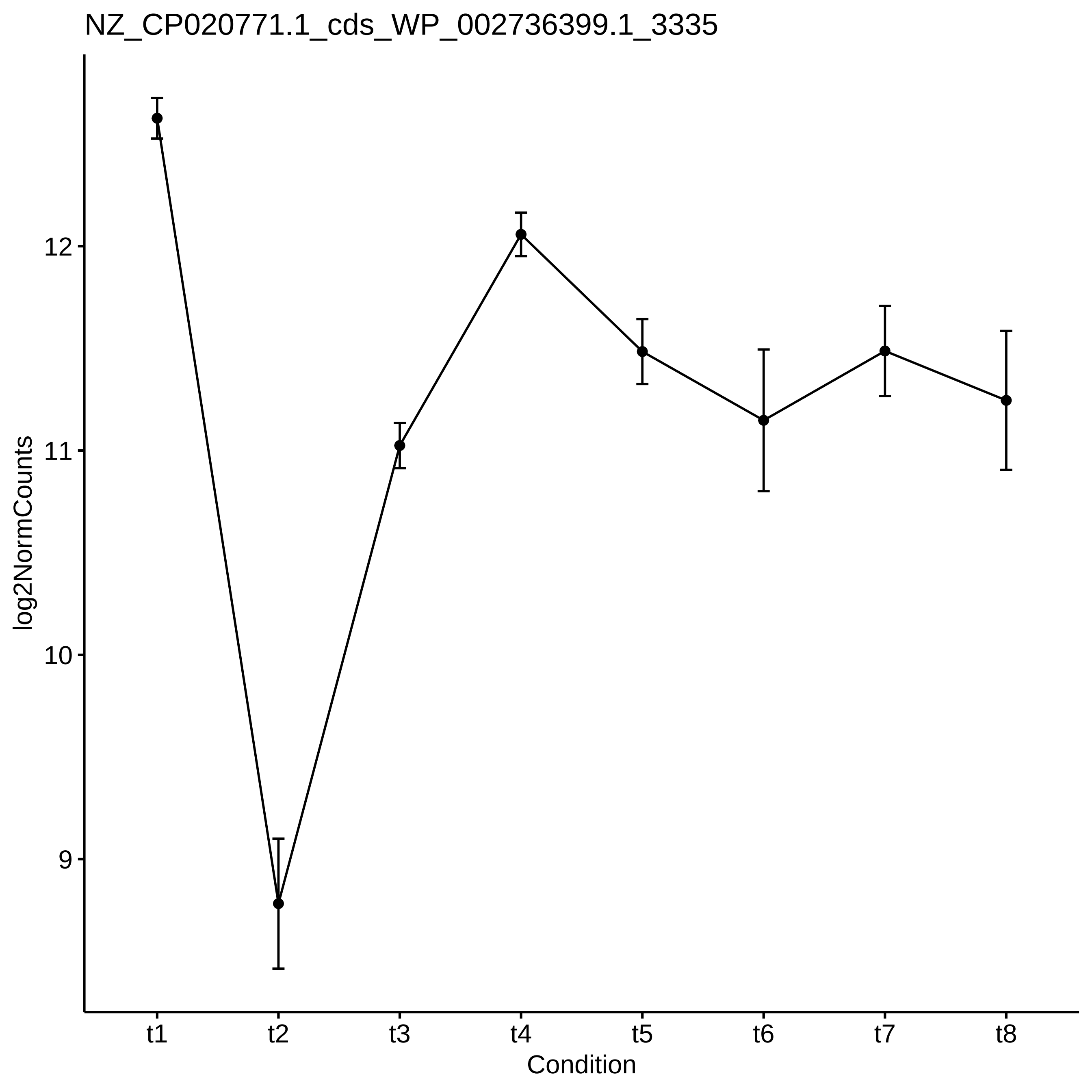
NZ_CP020771.1_cds_WP_002736399.1_3335
log2NormCounts
12
11
10
9
t1
t2
t3
t4
t5
t6
t7
t8
Condition

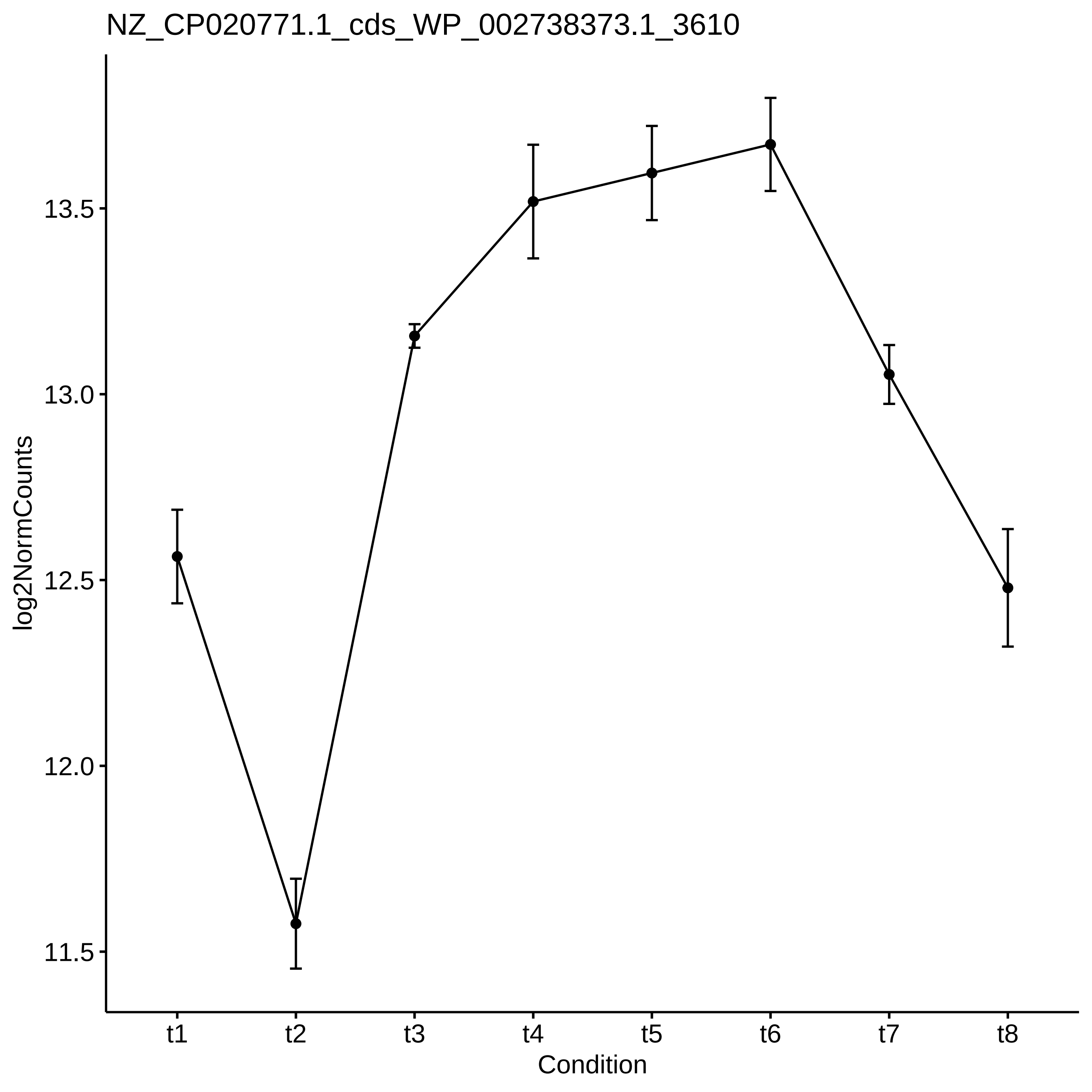
NZ_CP020771.1_cds_WP_002738373.1_3610
log2NormCounts
13.5
13.0
12.5
12.0
11.5
t1
t2
t3
t4
t5
t6
t7
t8
Condition
