## Supplementary_Data for "Diel changes in the expression of a marker gene and candidate genes for intracellular amorphous CaCO_3_ biomineralization in *Microcystis*": PCC7806_[t2-t8]_Top50_Profile.pdf

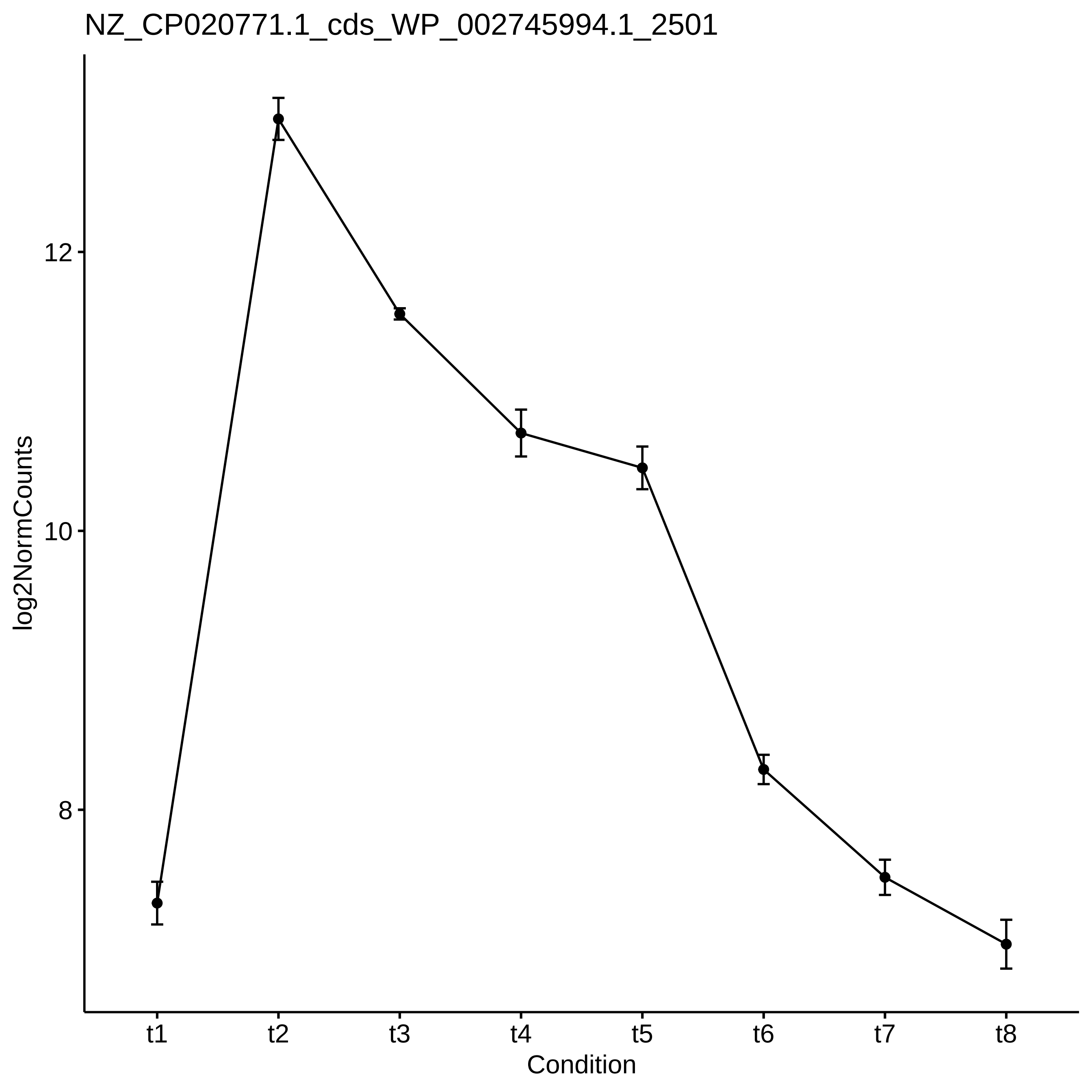
NZ_CP020771.1_cds_WP_002745994.1_2501
log2NormCounts
12
10
8
t1
t2
t3
t4
t5
t6
t7
t8
Condition

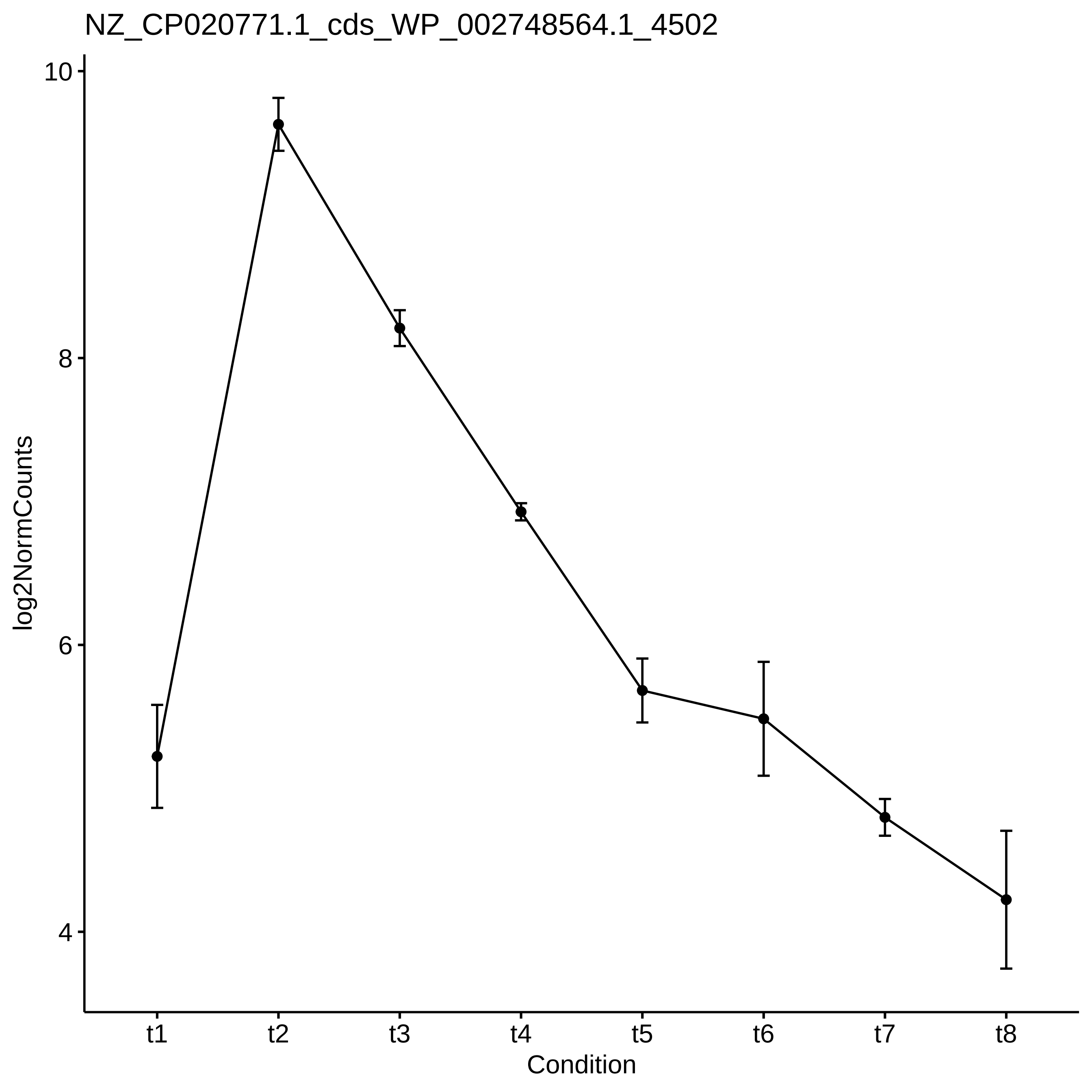

NZ_CP020771.1_cds_WP_002748564.1_4502
log2NormCounts
10
8
6
4
t1
t2
t3
t4
t5
t6
t7
t8
Condition

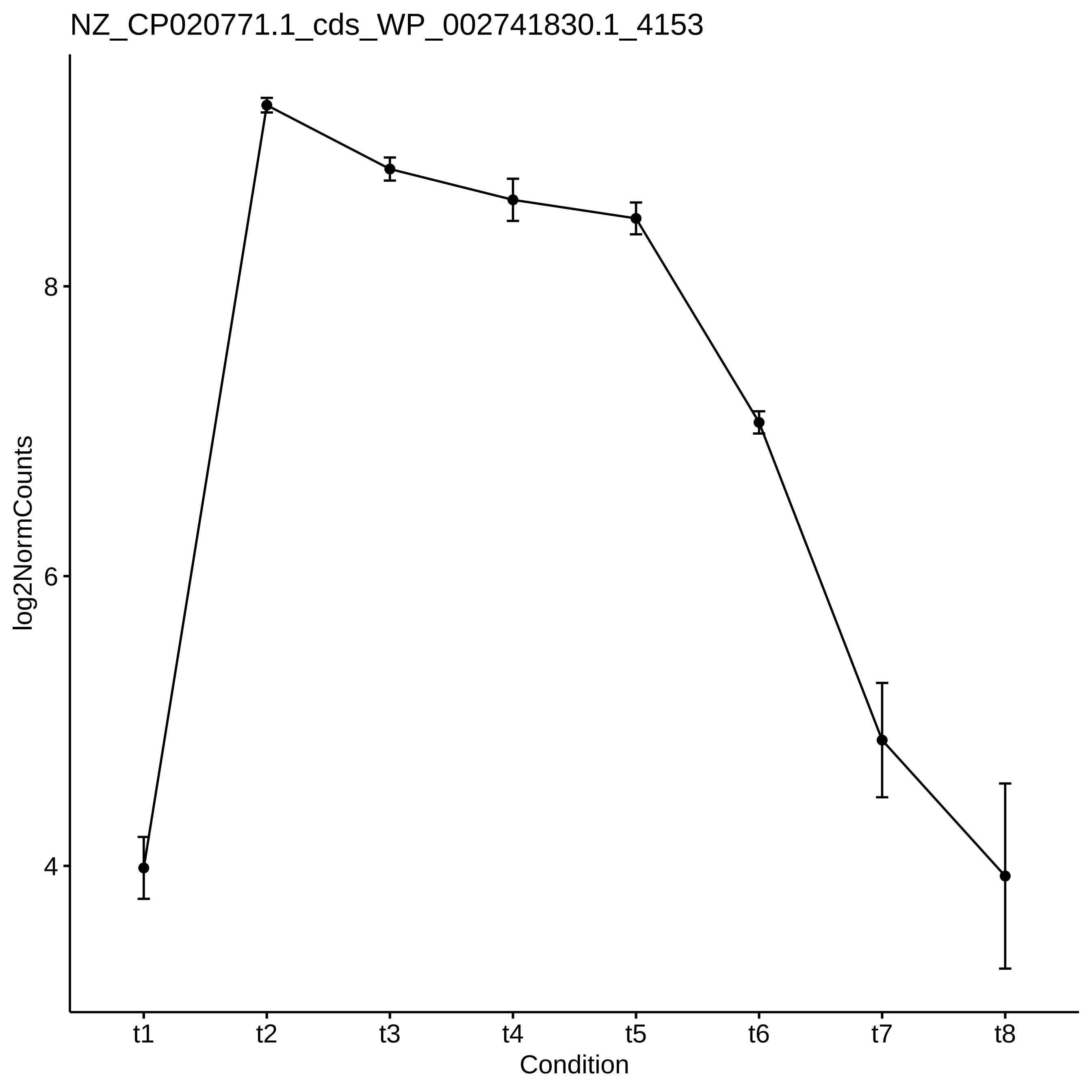
NZ_CP020771.1_cds_WP_002741830.1_4153
log2NormCounts
8
6
4
t1
t2
t3
t4
t5
t6
t7
t8
Condition

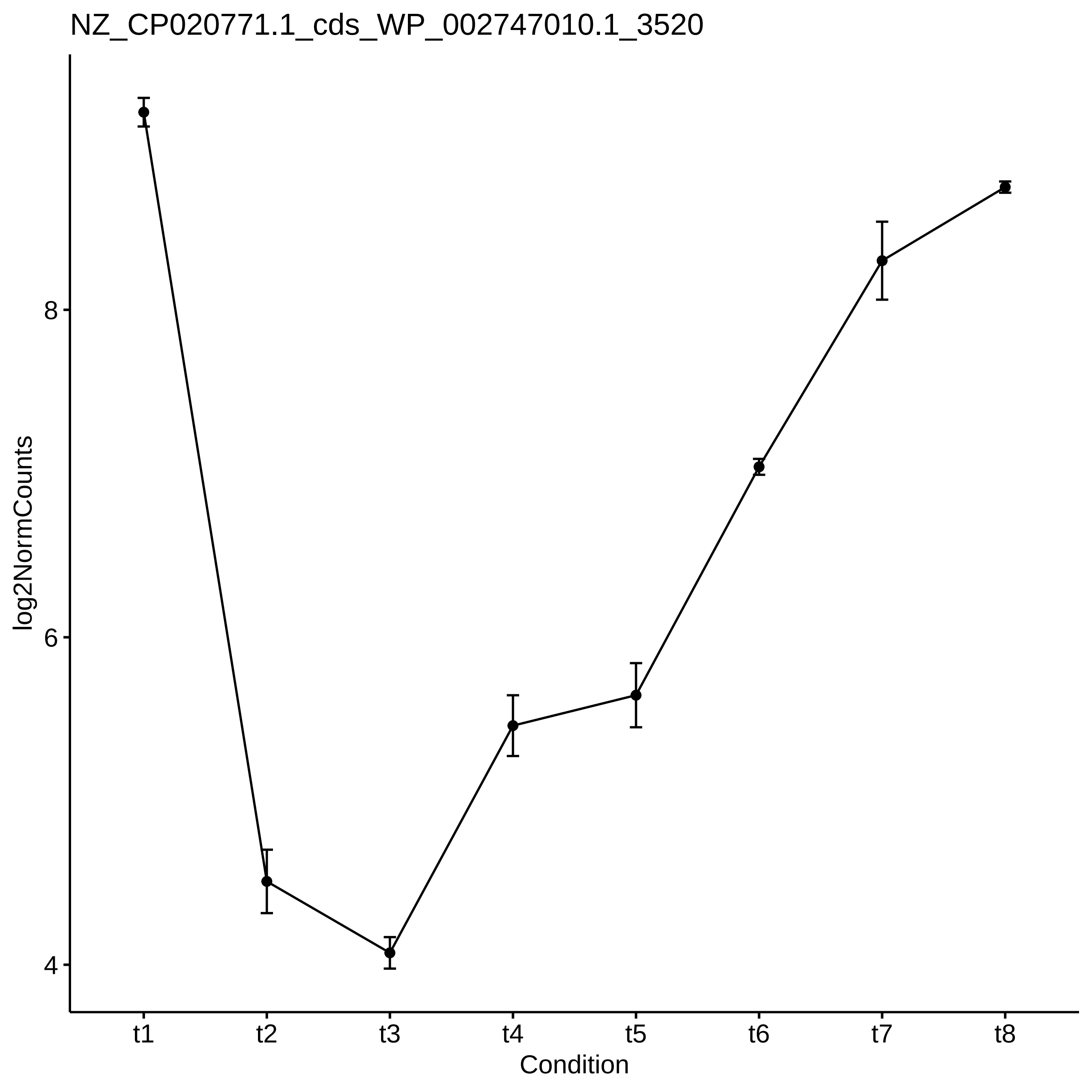
NZ_CP020771.1_cds_WP_002747010.1_3520
log2NormCounts
8
6
4
t1
t2
t3
t4
t5
t6
t7
t8
Condition

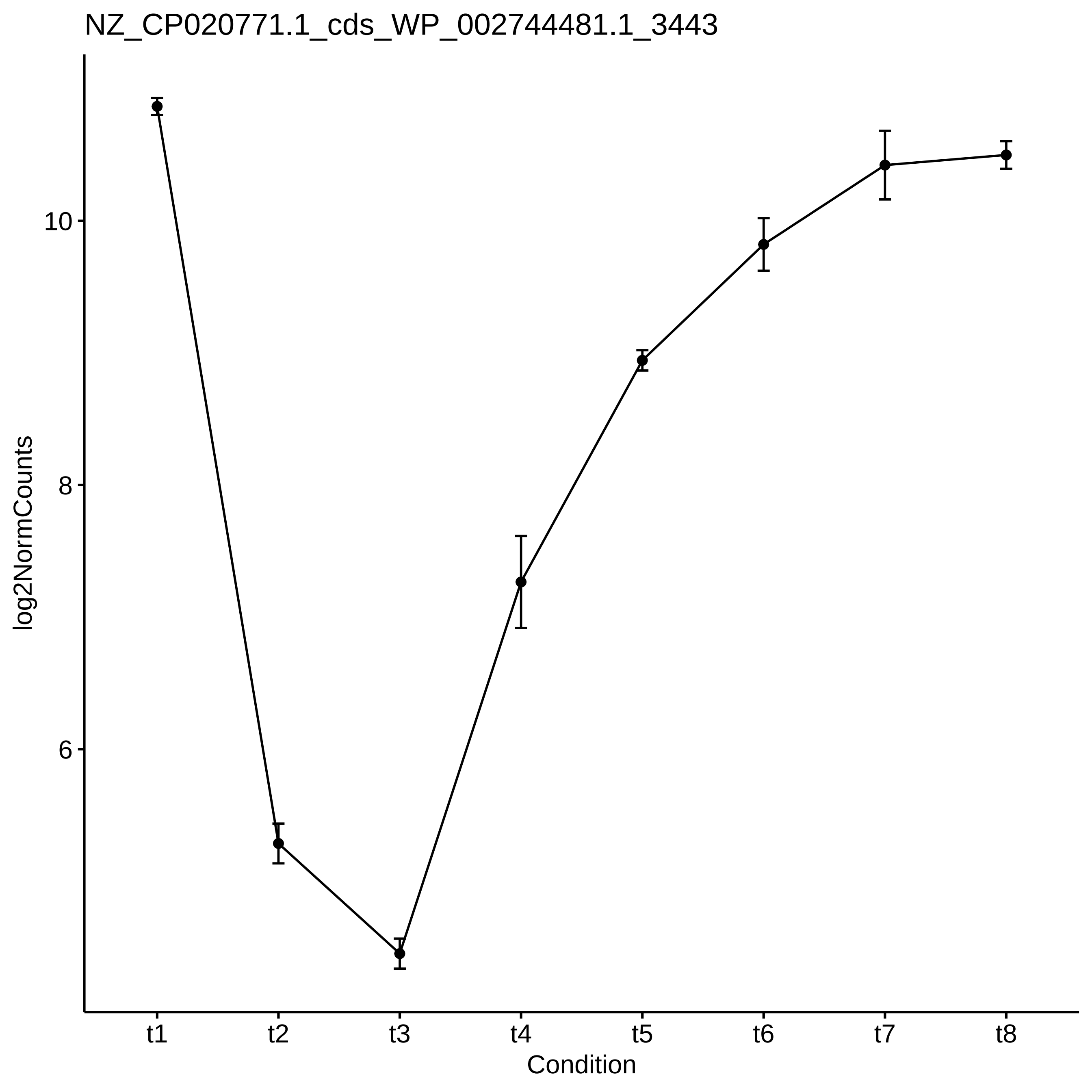
NZ_CP020771.1_cds_WP_002744481.1_3443
log2NormCounts
10
8
6
t1
t2
t3
t4
t5
t6
t7
t8
Condition

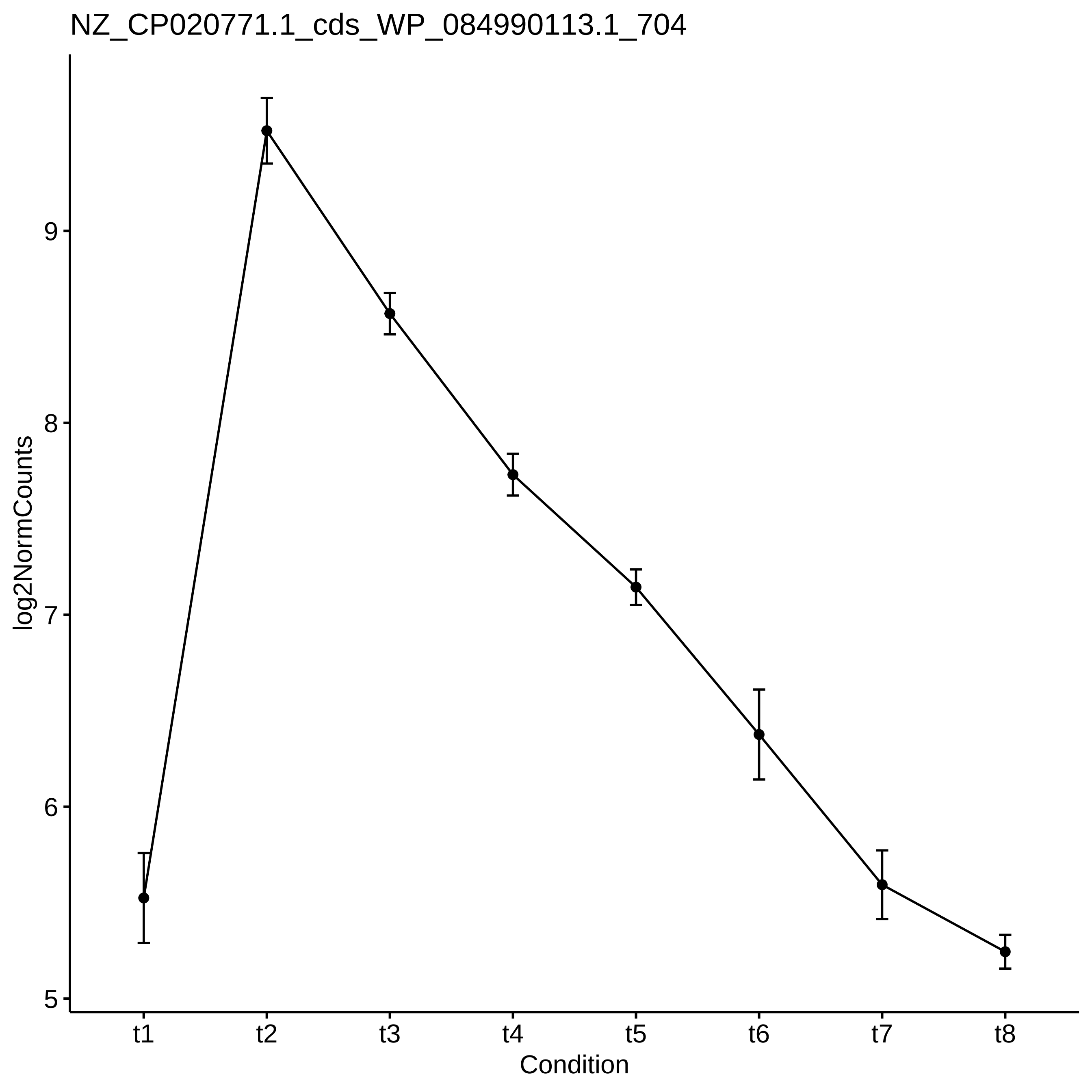

NZ_CP020771.1_cds_WP_084990113.1_704
log2NormCounts
9
8
7
6
5
t1
t2
t3
t4
t5
t6
t7
t8
Condition

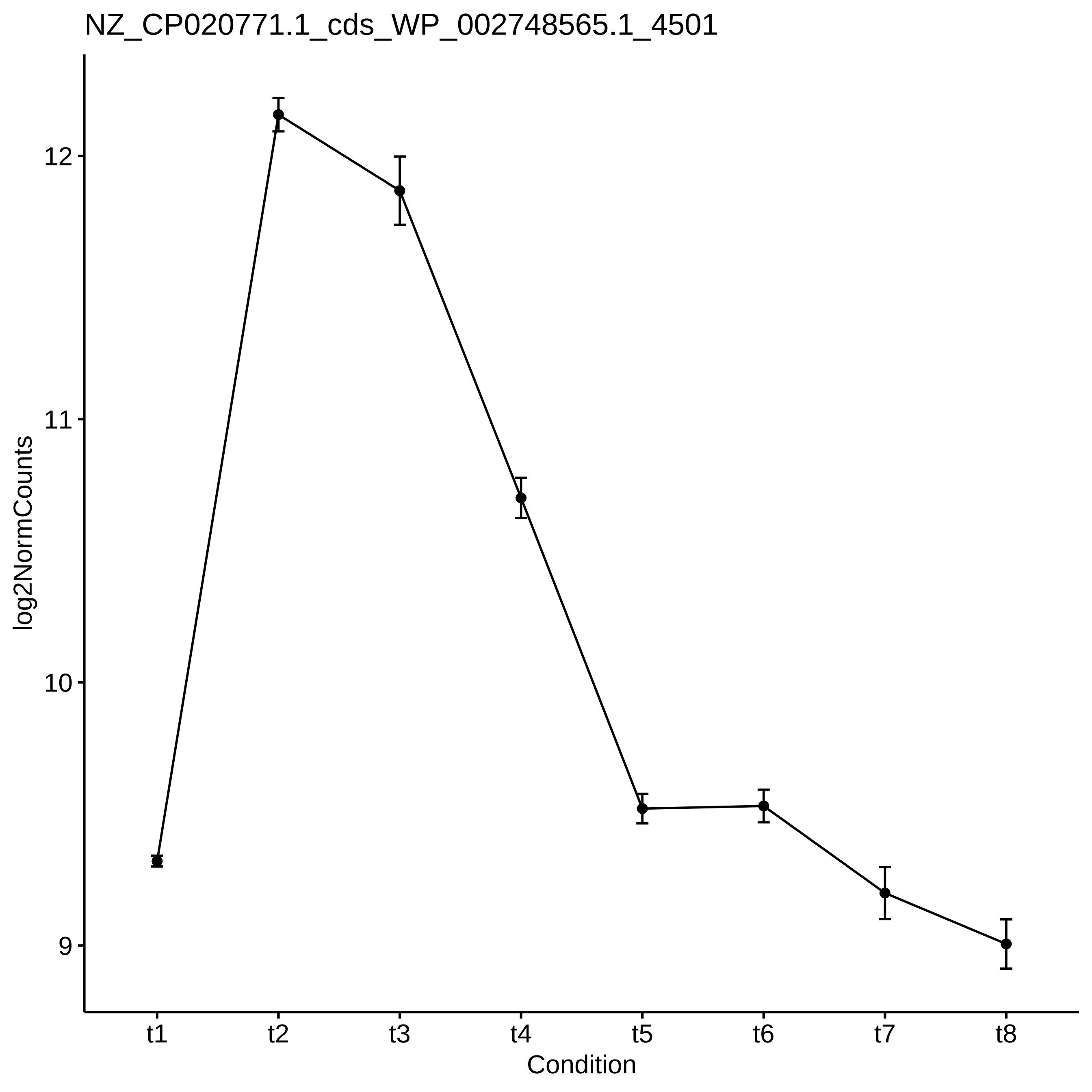
NZ_CP020771.1_cds_WP_002748565.1_4501
log2NormCounts
12
11
10
9
t1
t2
t3
t4
t5
t6
t7
t8
Condition

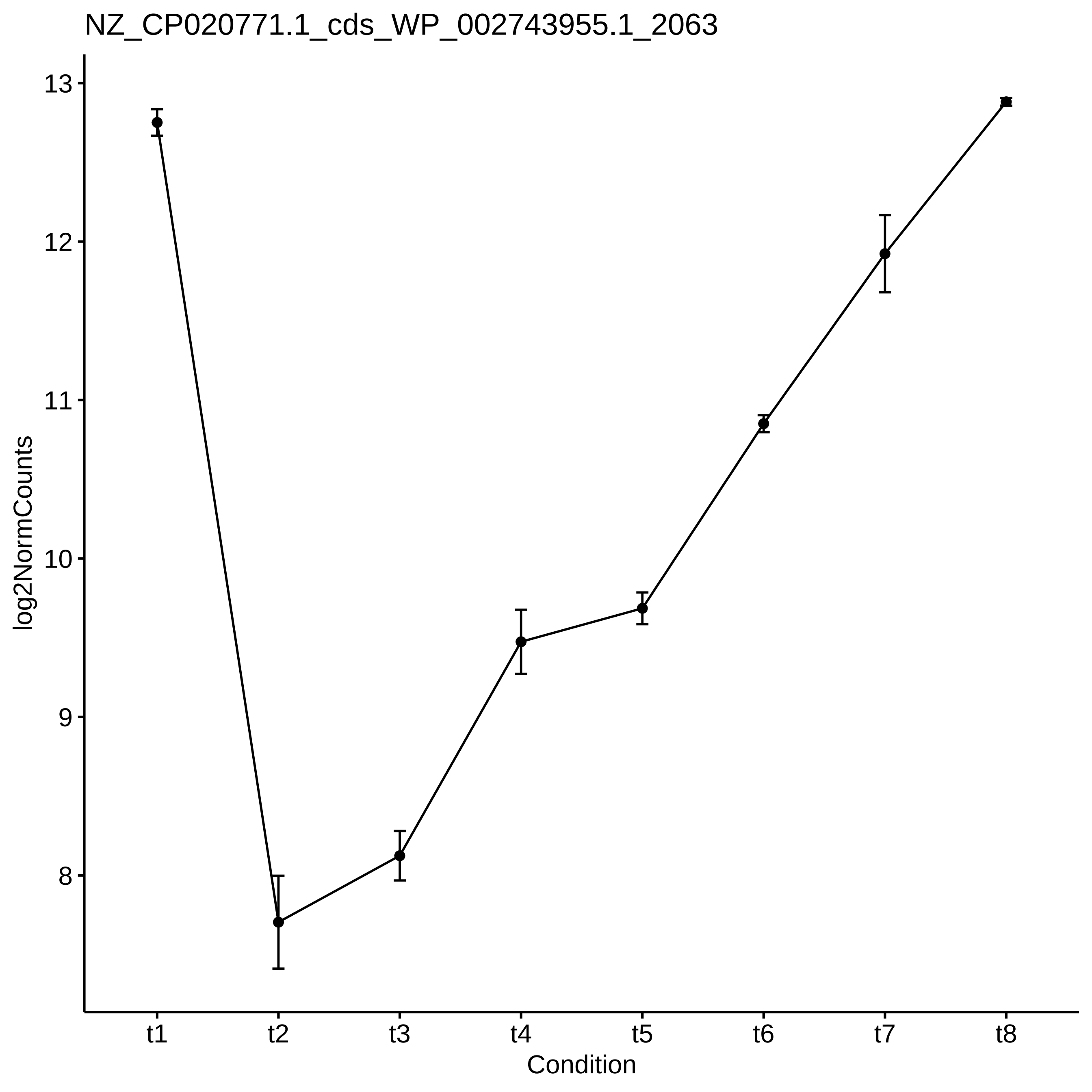
NZ_CP020771.1_cds_WP_002743955.1_2063
log2NormCounts
13
12
11
10
9
8
t1
t2
t3
t4
t5
t6
t7
t8
Condition

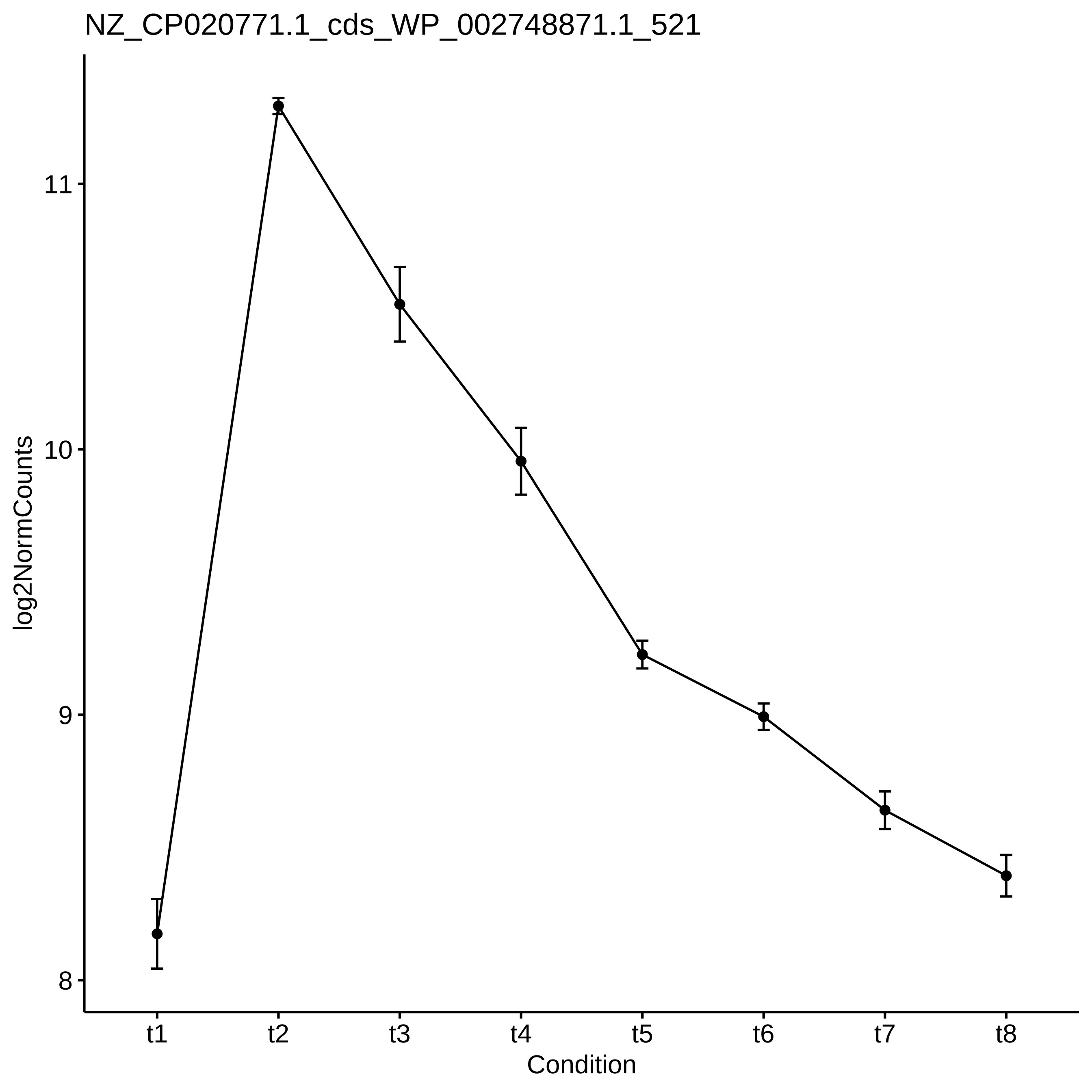
NZ_CP020771.1_cds_WP_002748871.1_521
log2NormCounts
11
10
9
8
t1
t2
t3
t4
t5
t6
t7
t8
Condition

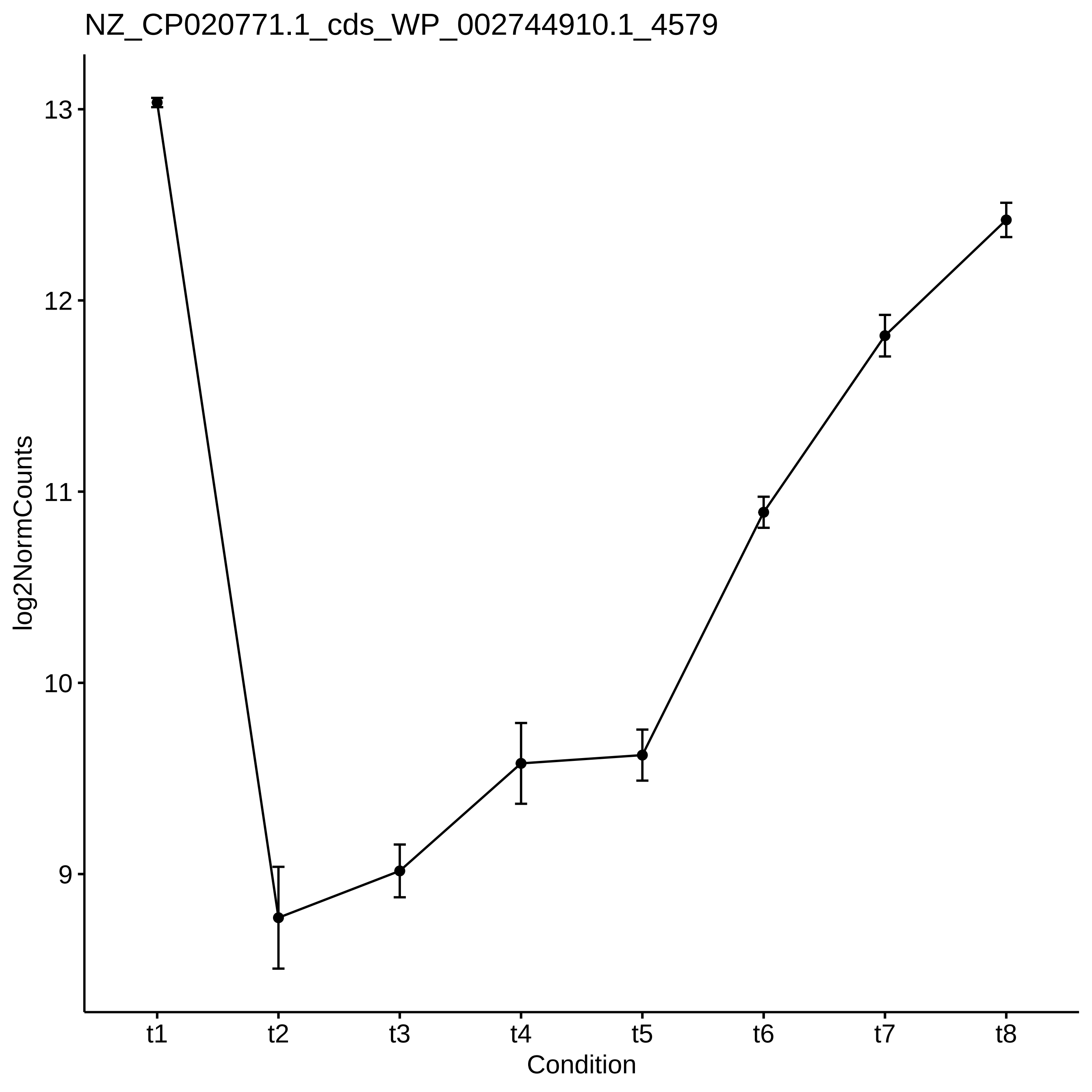
NZ_CP020771.1_cds_WP_002744910.1_4579
log2NormCounts
13
12
11
10
9
t1
t2
t3
t4
t5
t6
t7
t8
Condition

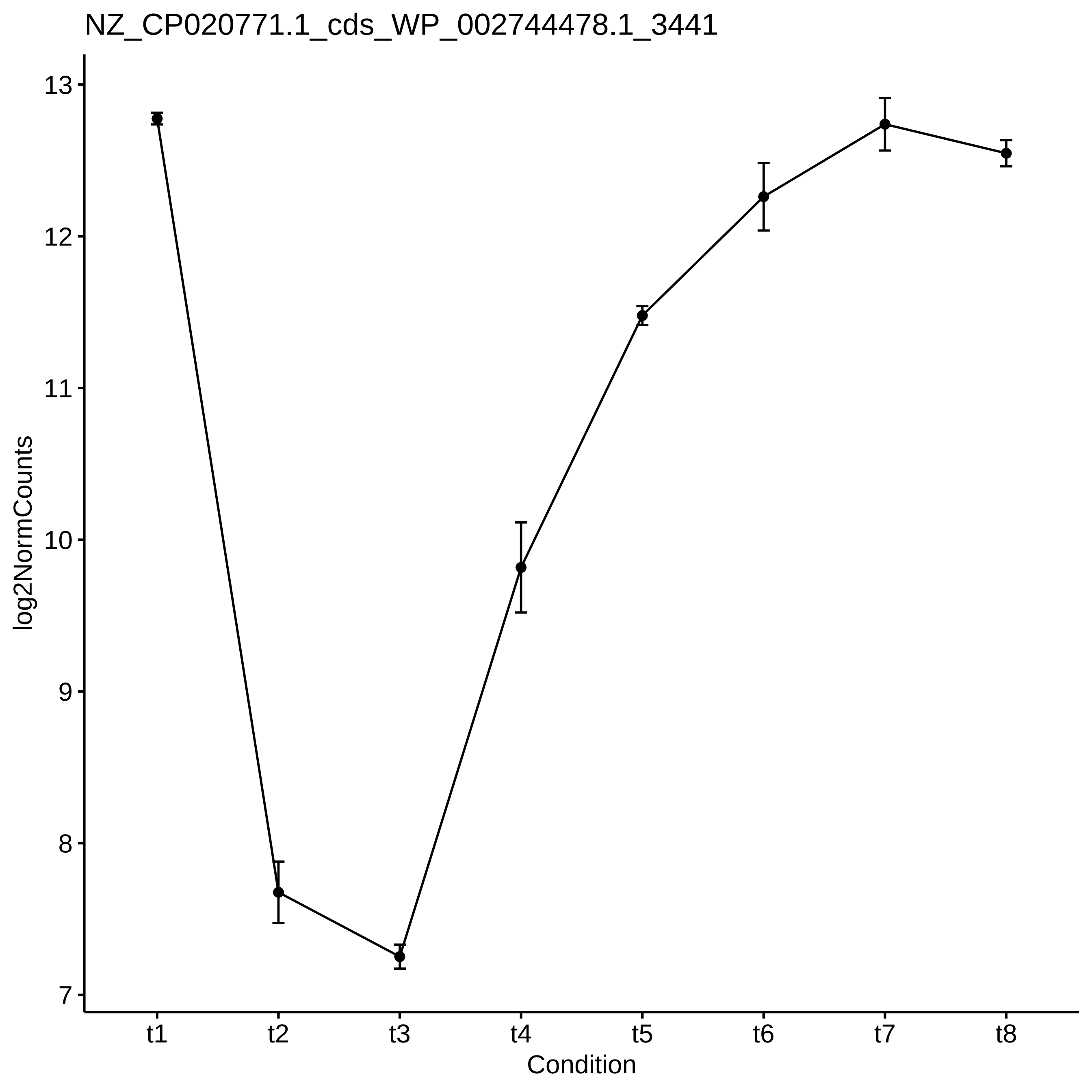
NZ_CP020771.1_cds_WP_002744478.1_3441
log2NormCounts
13
12
11
10
9
8
7
t1
t2
t3
t4
t5
t6
t7
t8
Condition

NZ_CP020771.1_cds_WP_002745275.1_222

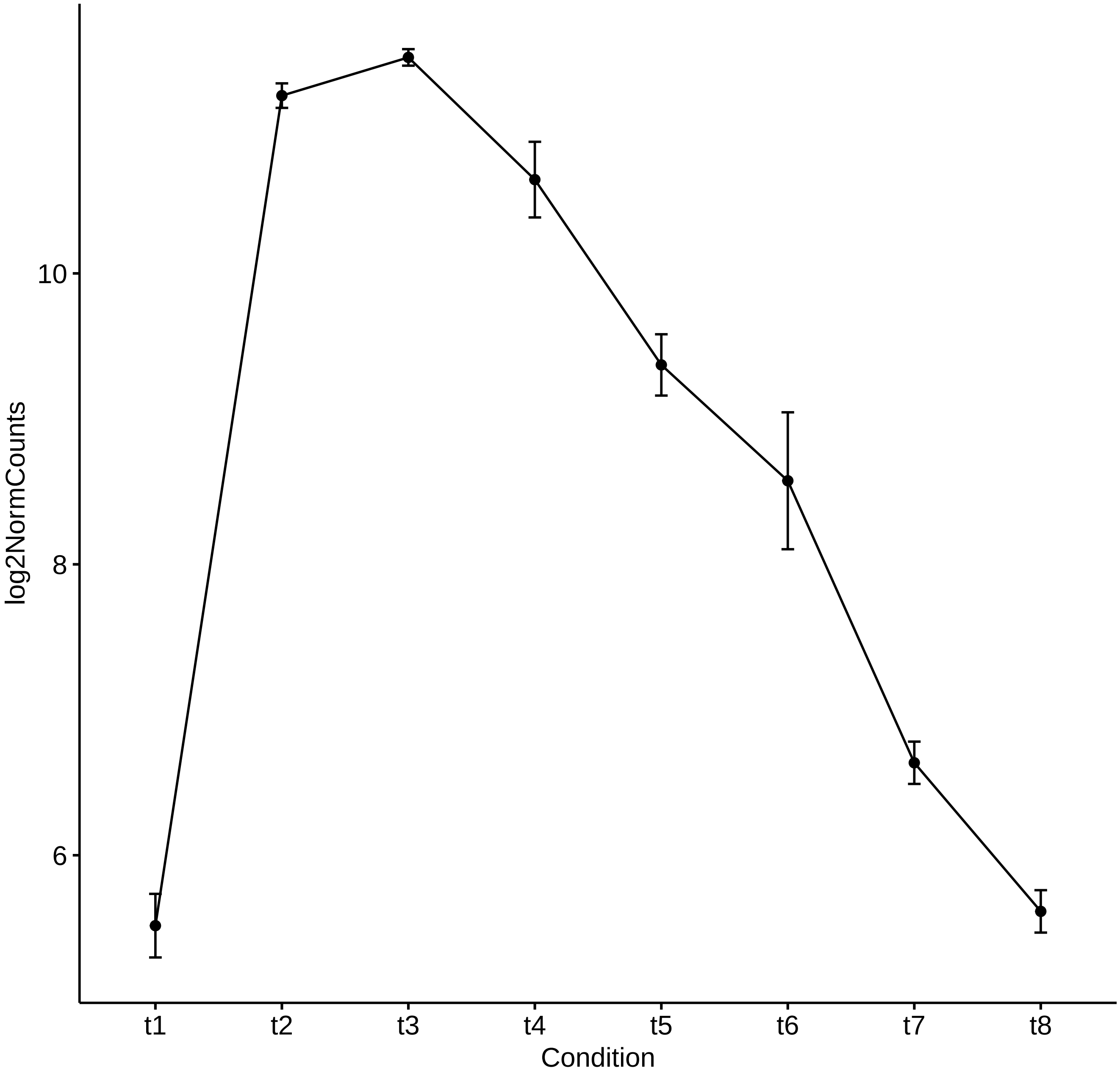

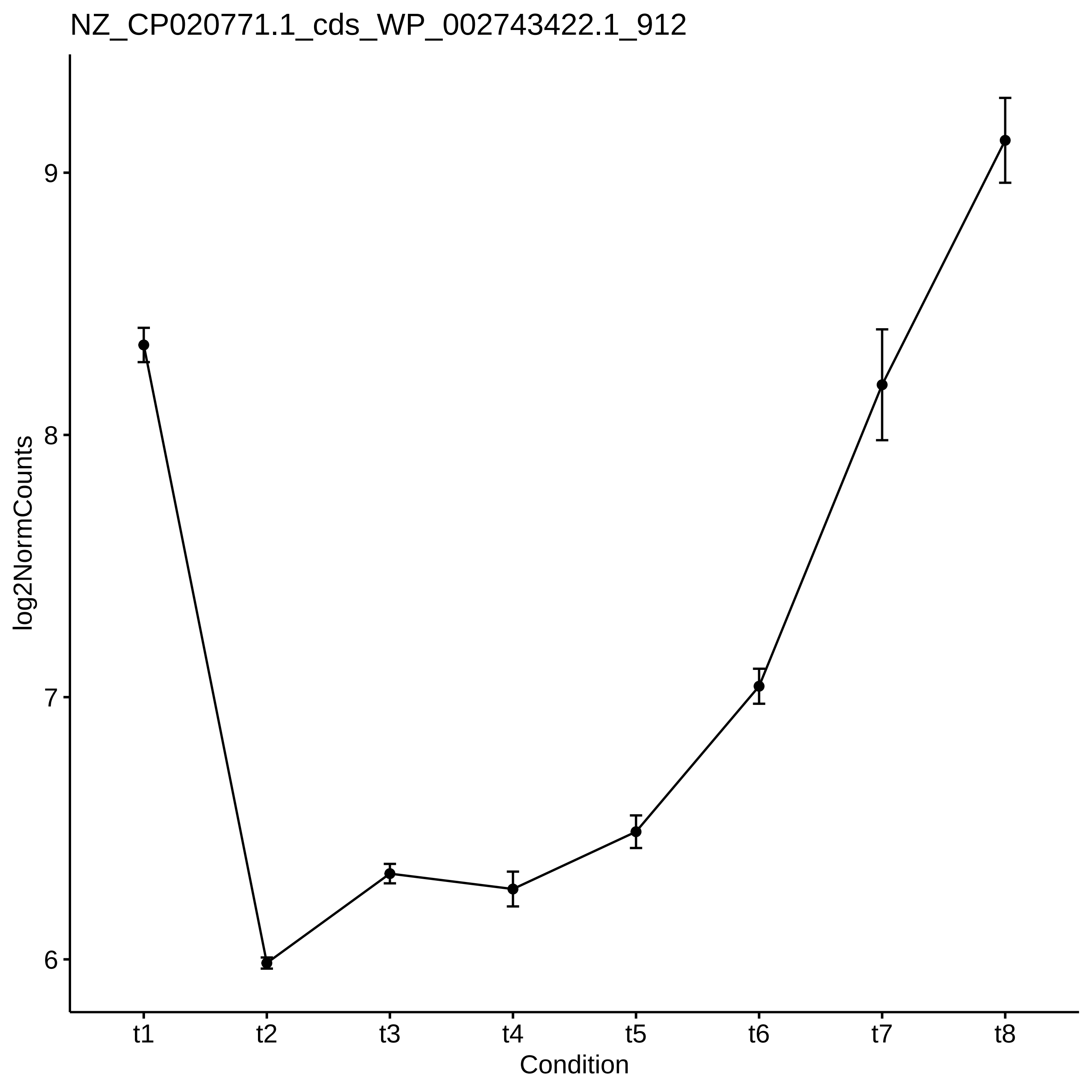

NZ_CP020771.1_cds_WP_002743422.1_912
log2NormCounts
9
8
7
6
t1
t2
t3
t4
t5
t6
t7
t8
Condition

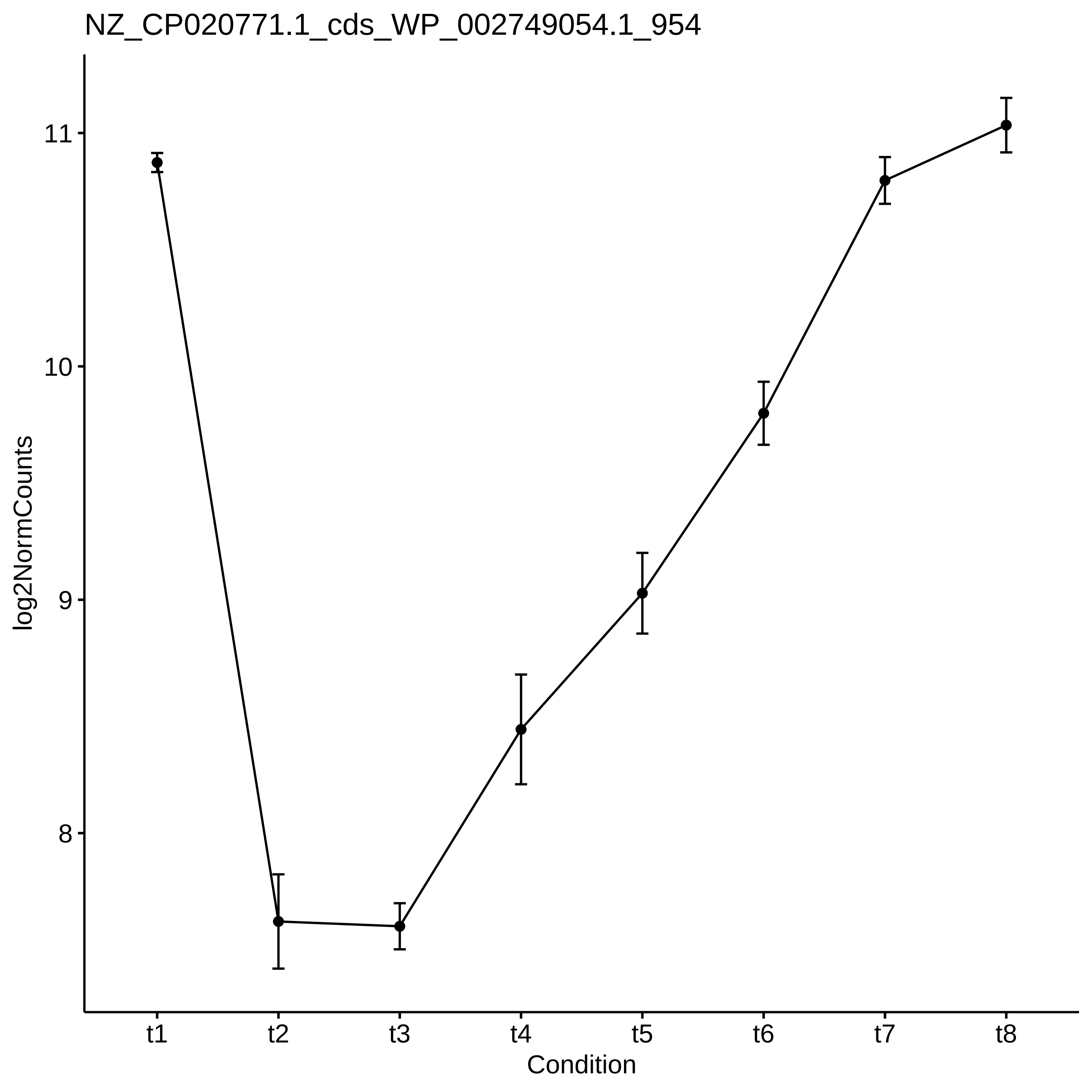
NZ_CP020771.1_cds_WP_002749054.1_954
log2NormCounts
11
10
9
8
t1
t2
t3
t4
t5
t6
t7
t8
Condition

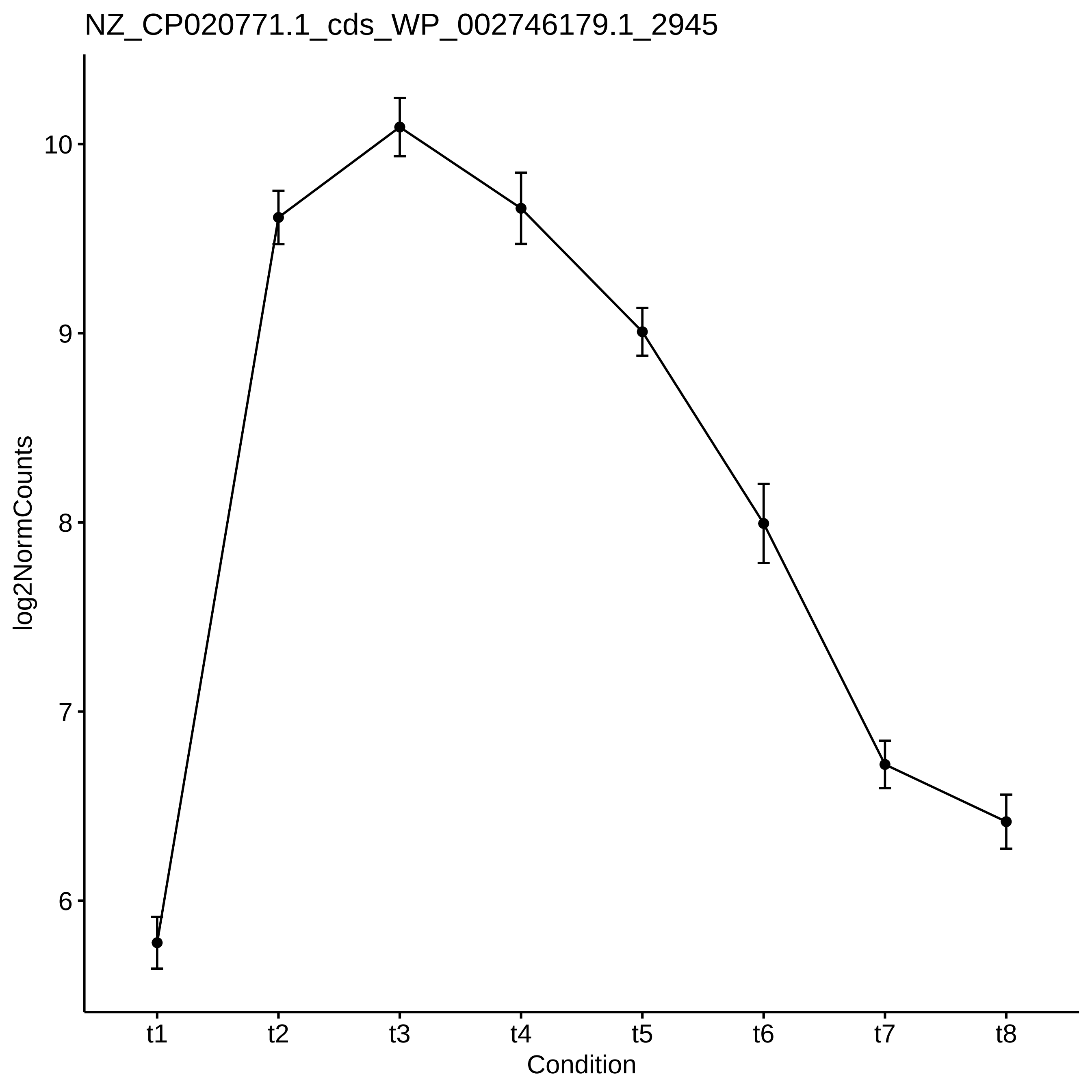
NZ_CP020771.1_cds_WP_002746179.1_2945
log2NormCounts
10
9
8
7
6
t1
t2
t3
t4
t5
t6
t7
t8
Condition

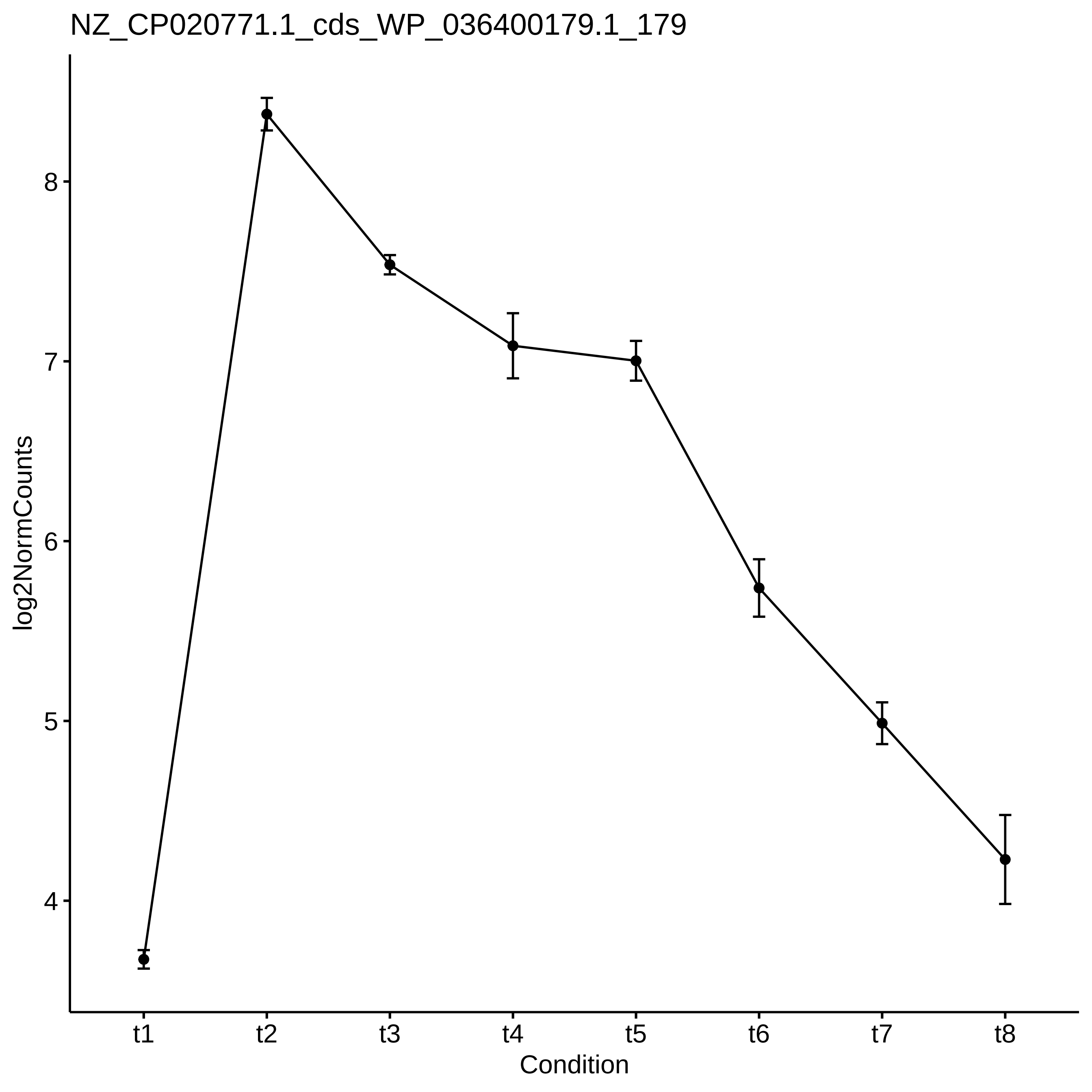
NZ_CP020771.1_cds_WP_036400179.1_179
log2NormCounts
8
7
6
5
4
t1
t2
t3
t4
t5
t6
t7
t8
Condition

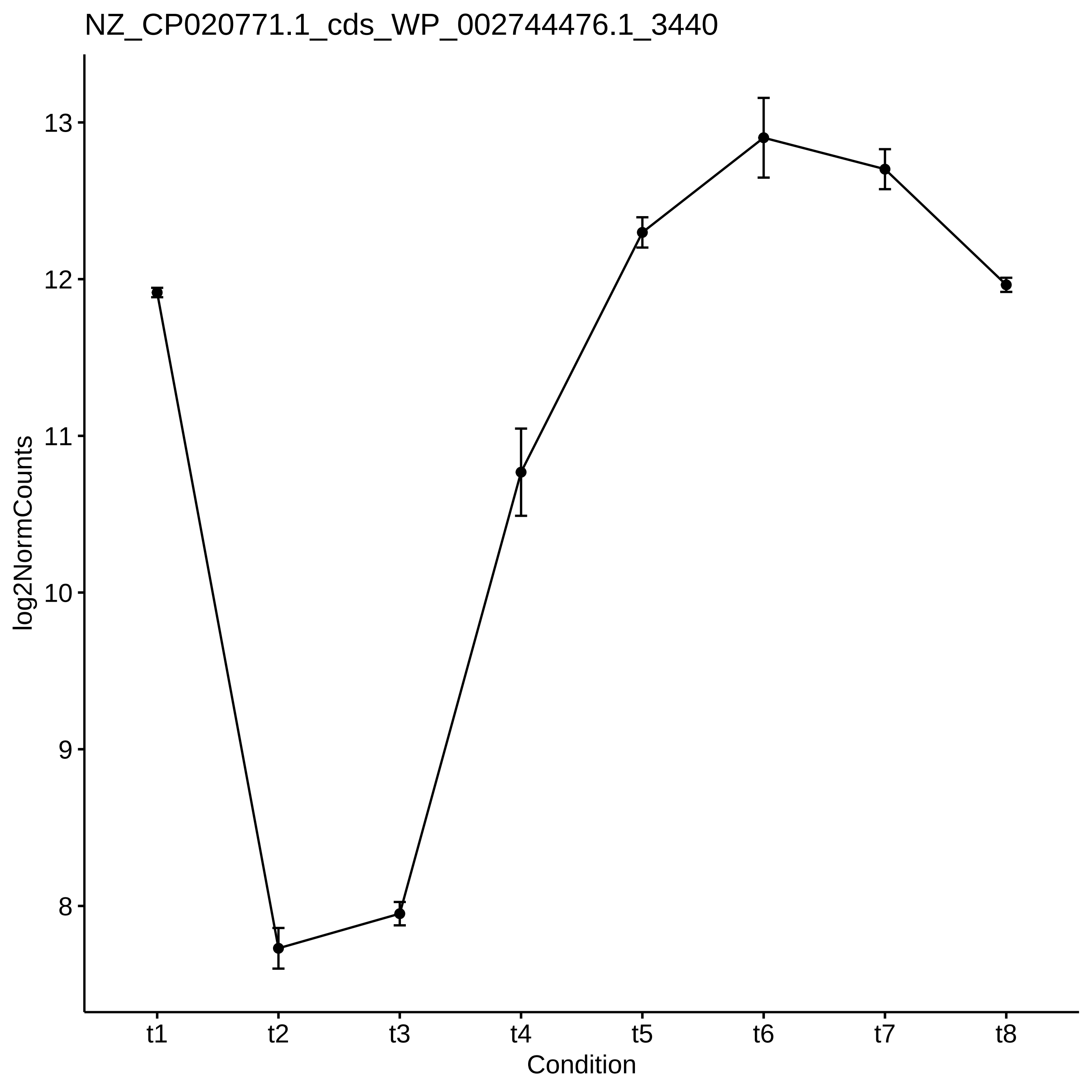

NZ_CP020771.1_cds_WP_002744476.1_3440
log2NormCounts
13
12
11
10
9
8
t1
t2
t3
t4
t5
t6
t7
t8
Condition

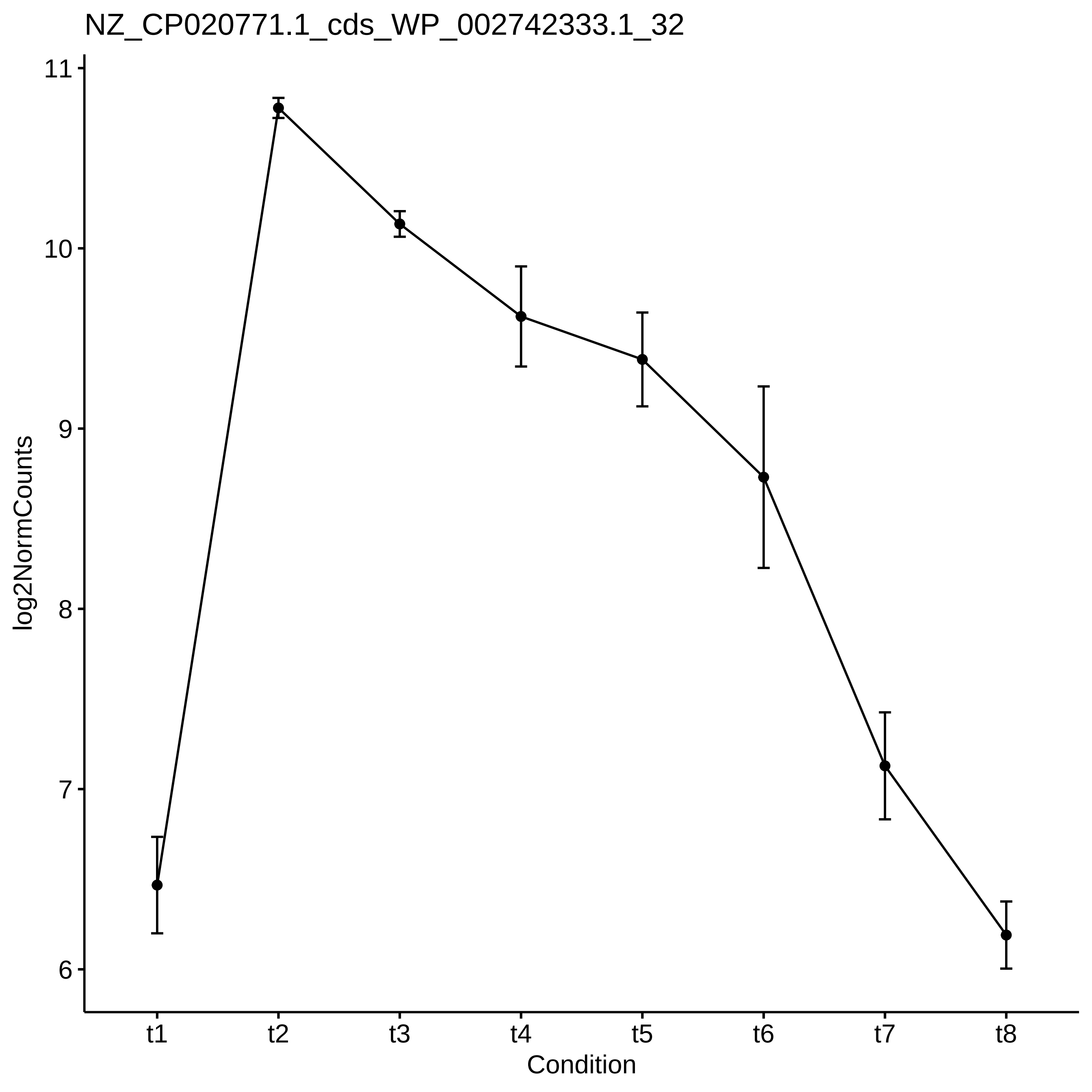

NZ_CP020771.1_cds_WP_002742333.1_32
log2NormCounts
11
10
9
8
7
6
t1
t2
t3
t4
t5
t6
t7
t8
Condition

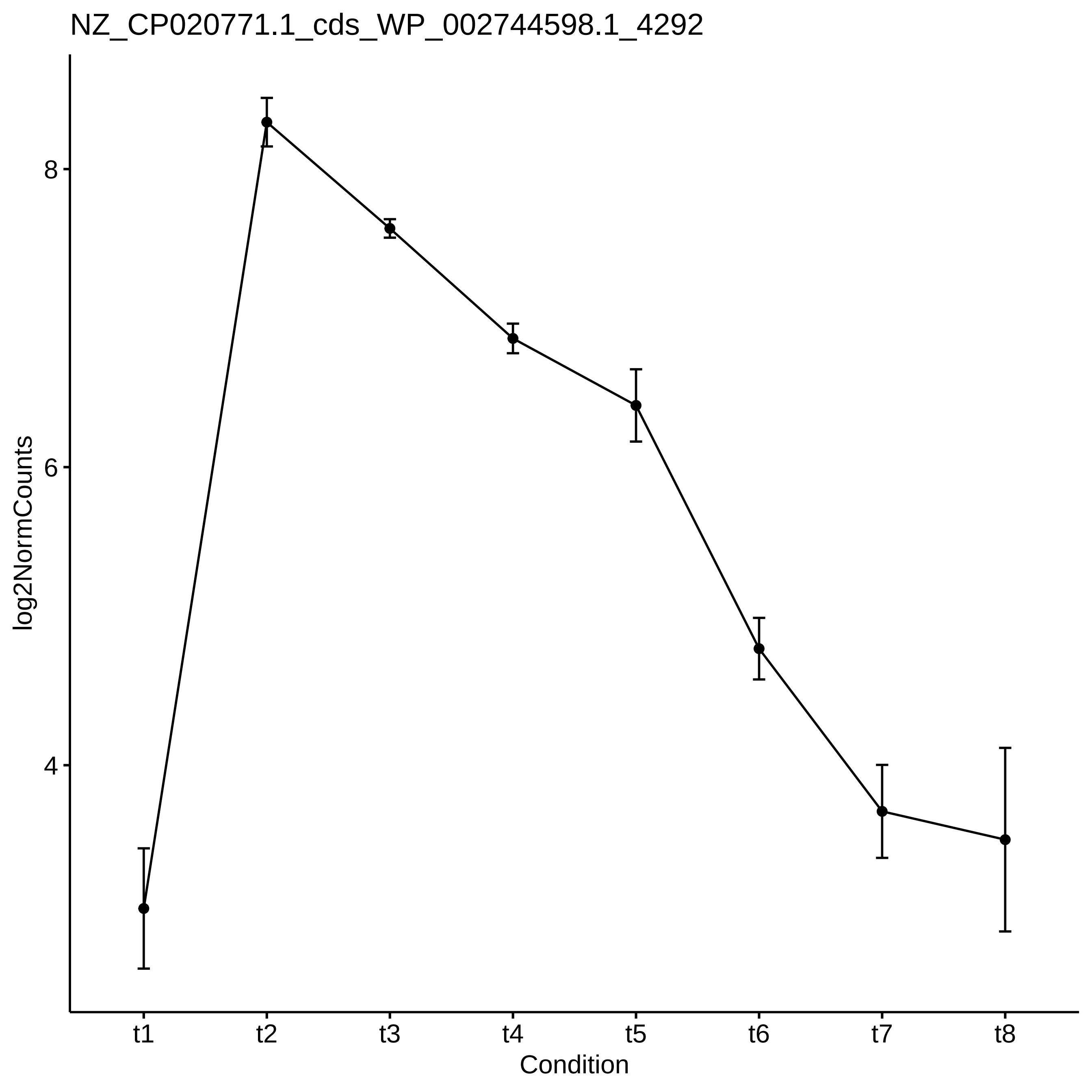
NZ_CP020771.1_cds_WP_002744598.1_4292
log2NormCounts
8
6
4
t1
t2
t3
t4
t5
t6
t7
t8
Condition

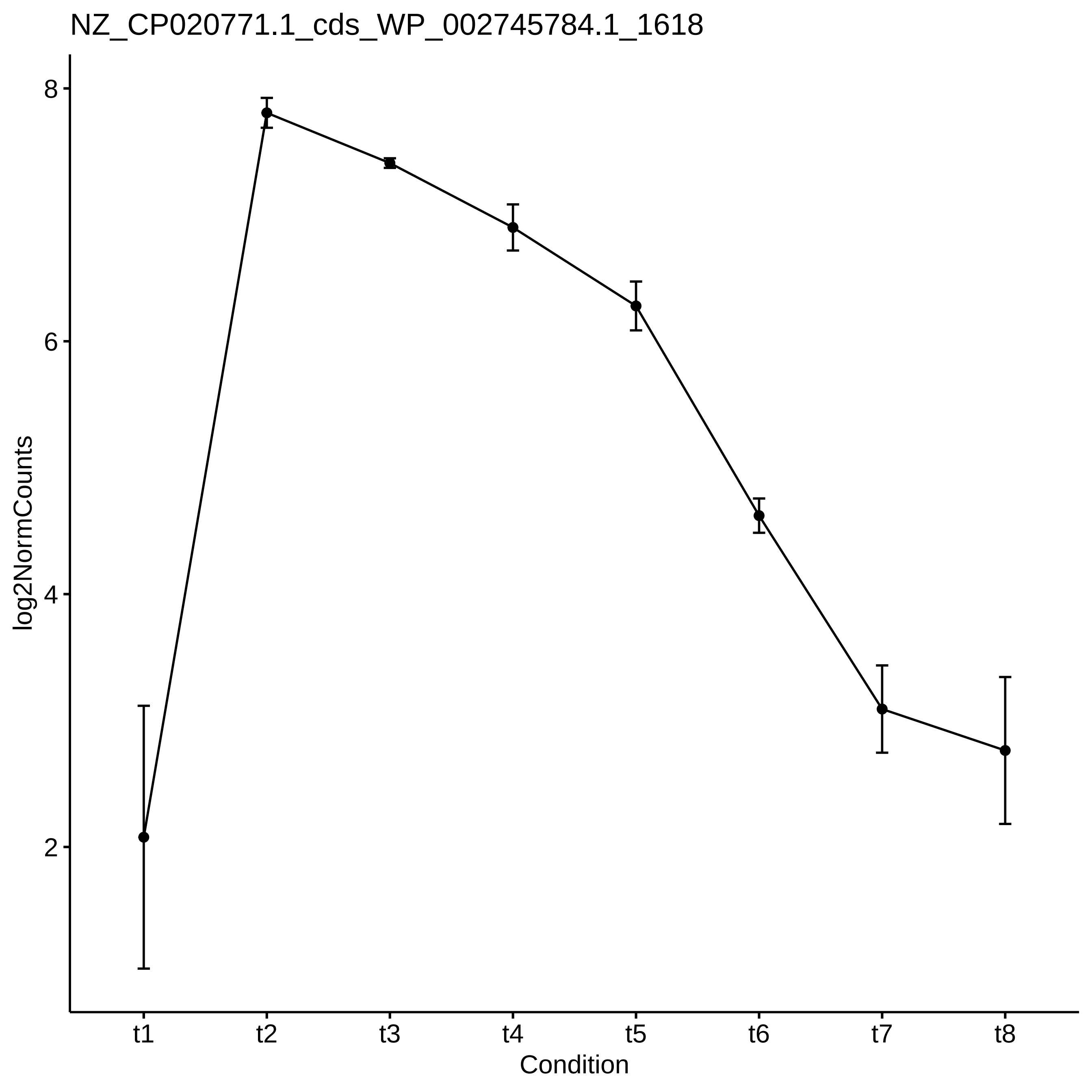
NZ_CP020771.1_cds_WP_002745784.1_1618
log2NormCounts
8
6
4
2
t1
t2
t3
t4
t5
t6
t7
t8
Condition

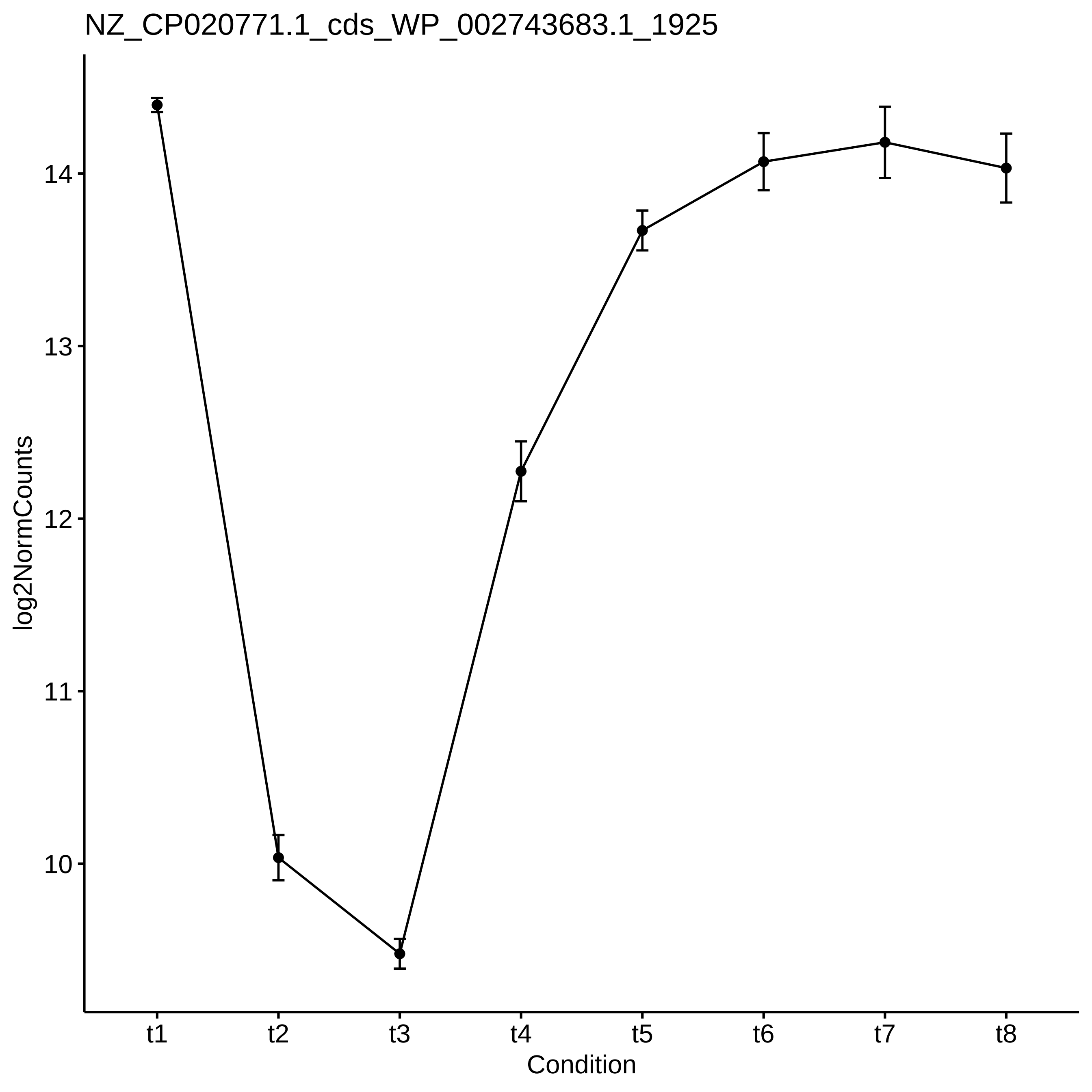
NZ_CP020771.1_cds_WP_002743683.1_1925
log2NormCounts
14
13
12
11
10
t1
t2
t3
t4
t5
t6
t7
t8
Condition

NZ_CP020771.1_cds_WP_002743745.1_1953

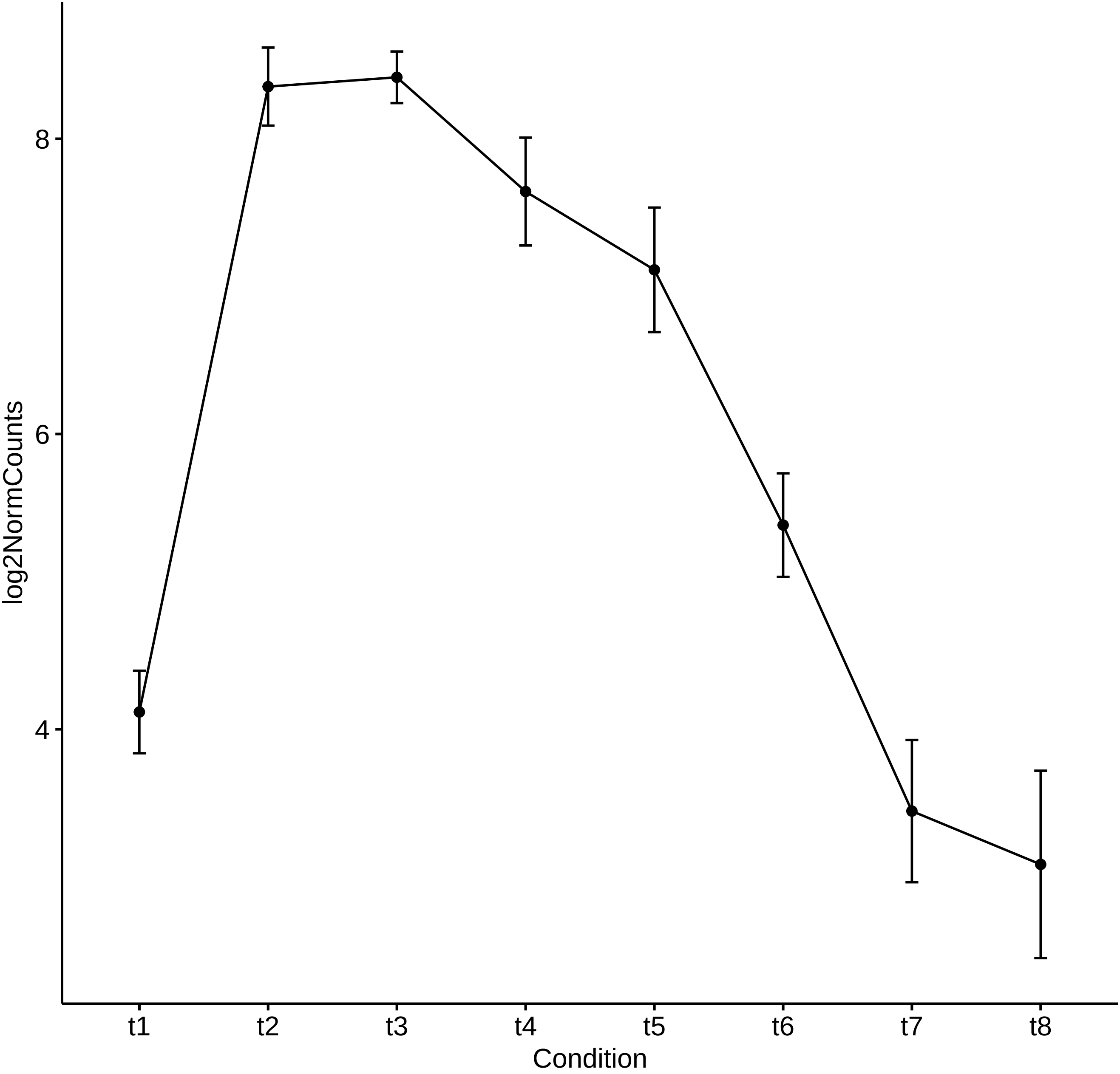

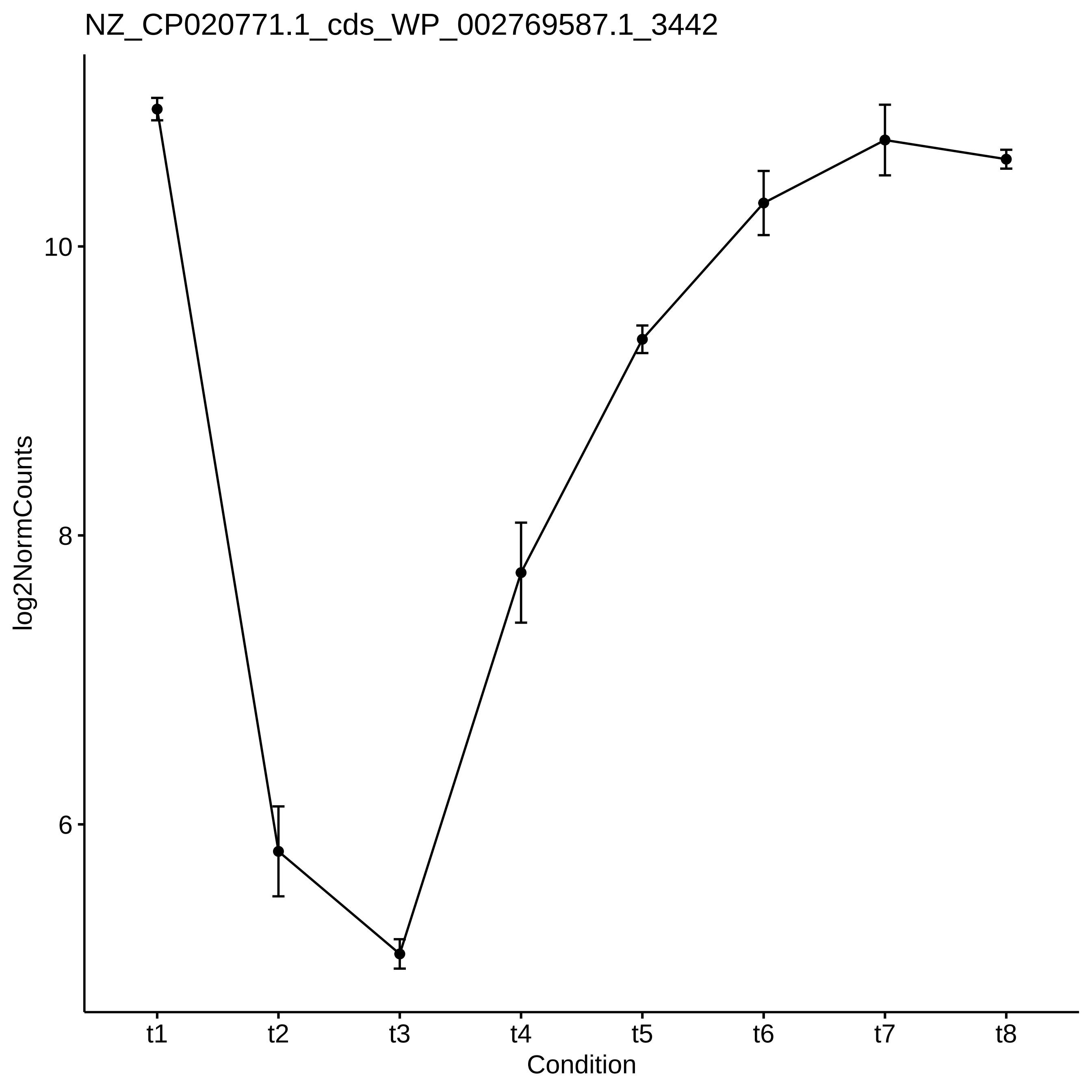
NZ_CP020771.1_cds_WP_002769587.1_3442
log2NormCounts
10
8
6
t1
t2
t3
t4
t5
t6
t7
t8
Condition

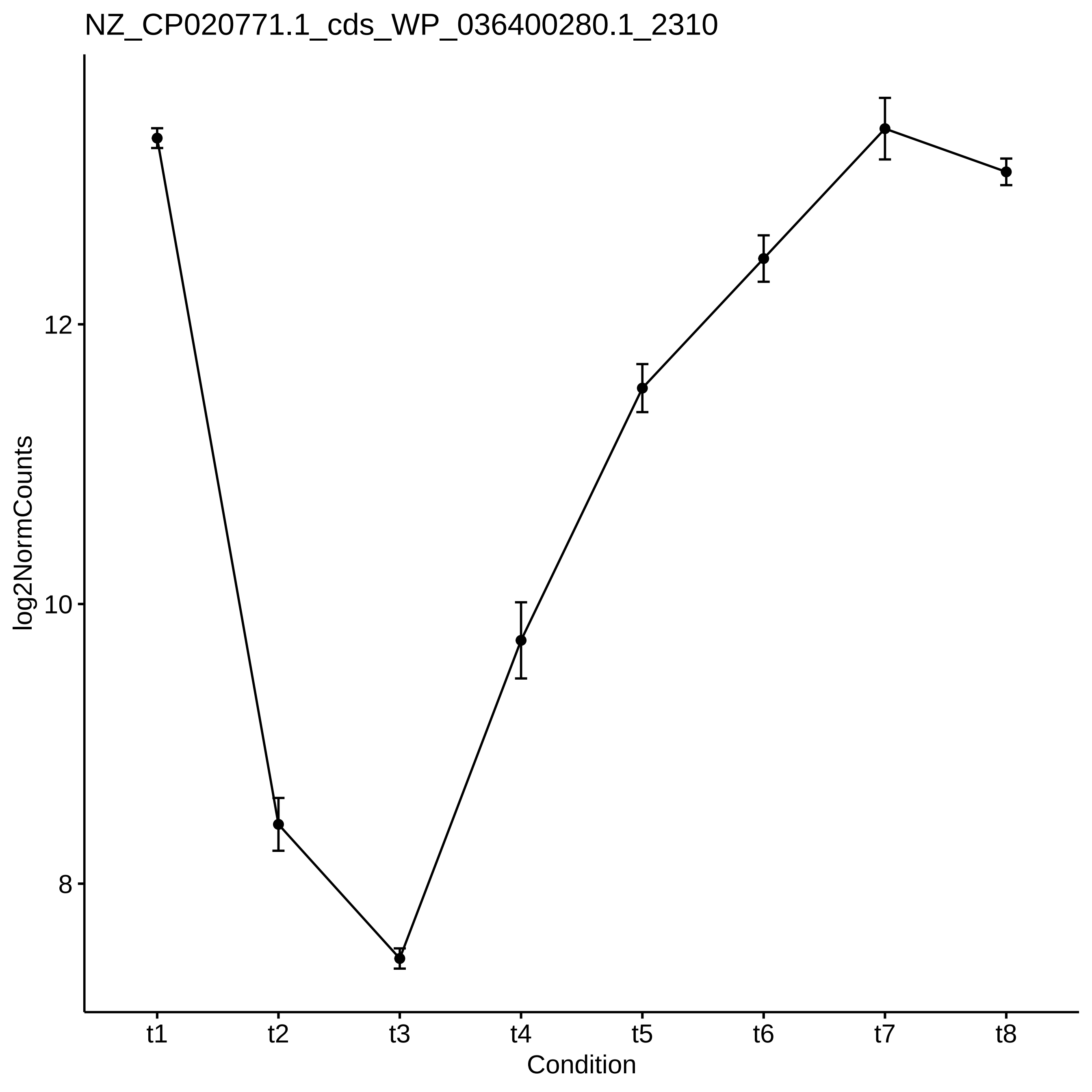
NZ_CP020771.1_cds_WP_036400280.1_2310
log2NormCounts
12
10
8
t1
t2
t3
t4
t5
t6
t7
t8
Condition

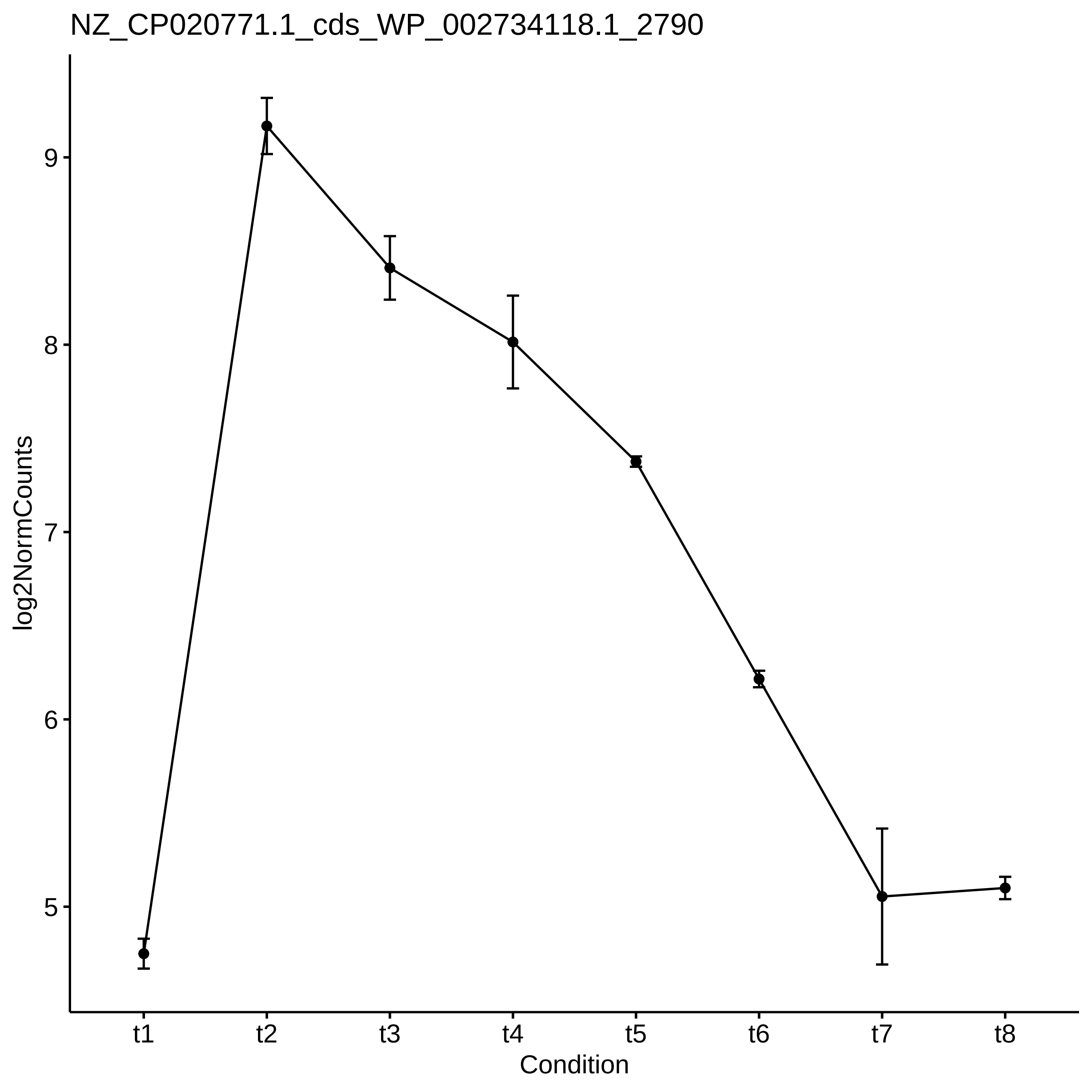
NZ_CP020771.1_cds_WP_002734118.1_2790
log2NormCounts
9
8
7
6
5
t1
t2
t3
t4
t5
t6
t7
t8
Condition

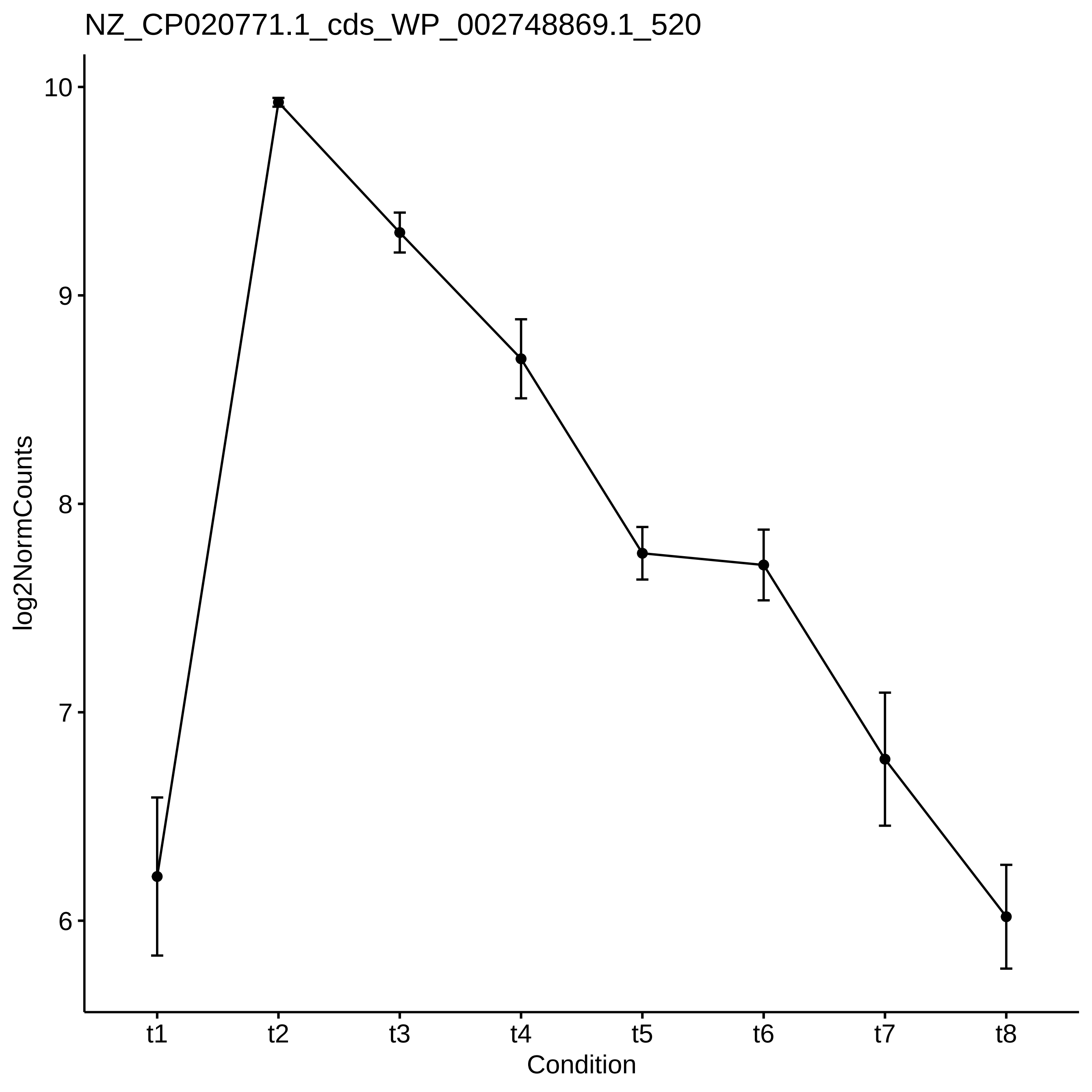

NZ_CP020771.1_cds_WP_002748869.1_520
10
9
8
7
6
log2NormCounts
t1
t2
t3
t4
t5
t6
t7
t8
Condition

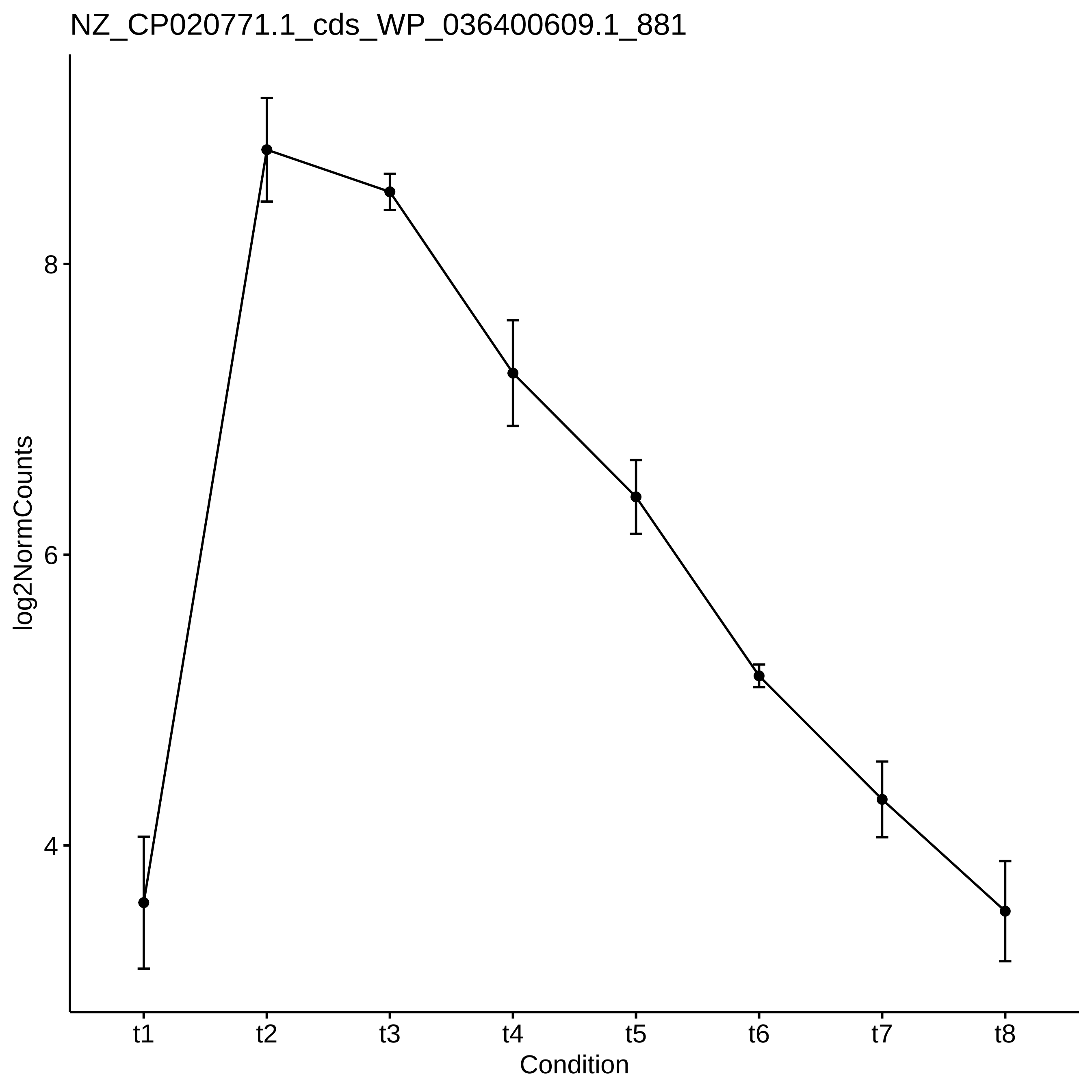

NZ_CP020771.1_cds_WP_036400609.1_881
log2NormCounts
8
6
4
t1
t2
t3
t4
t5
t6
t7
t8
Condition

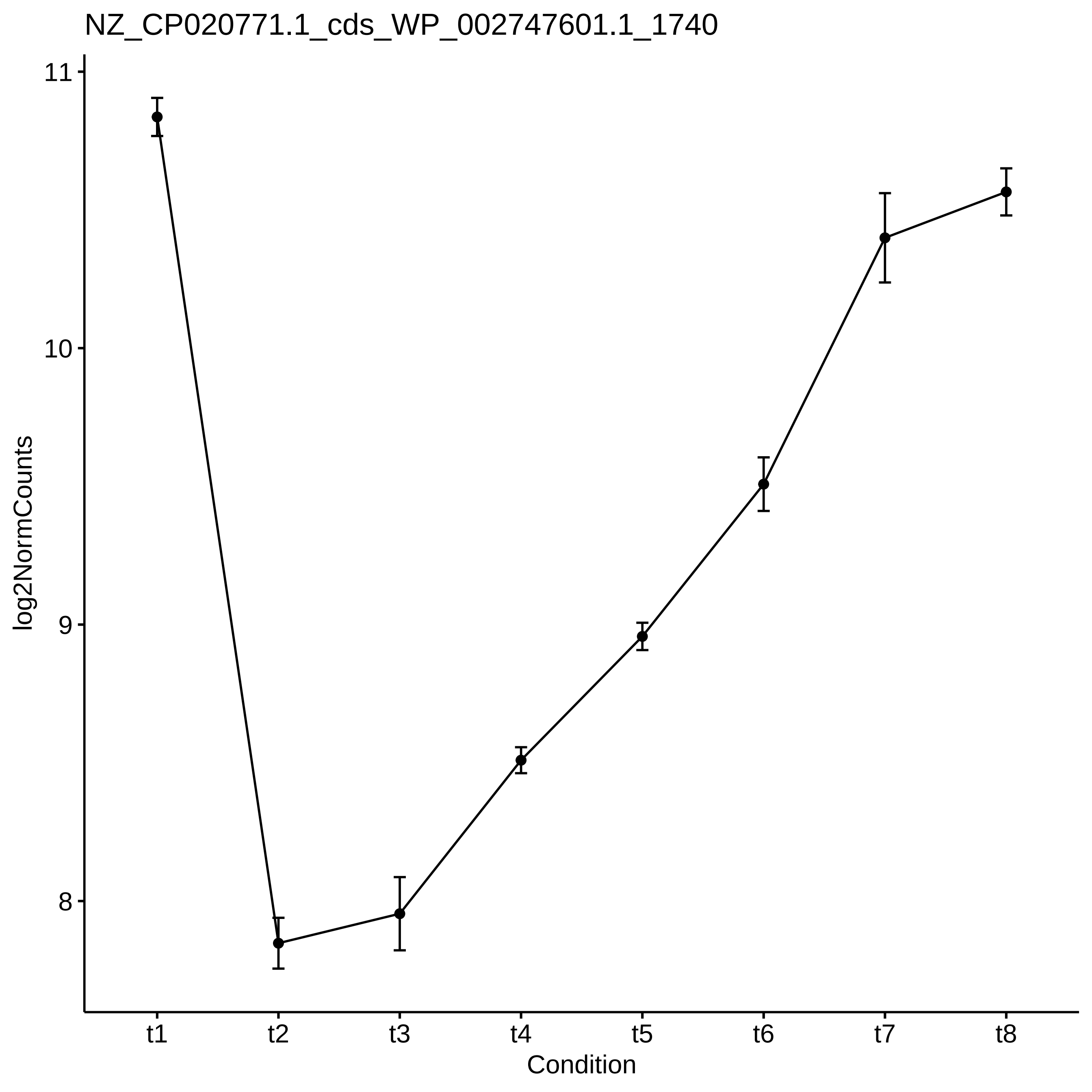

NZ_CP020771.1_cds_WP_002747601.1_1740
log2NormCounts
11
10
9
8
t1
t2
t3
t4
t5
t6
t7
t8
Condition

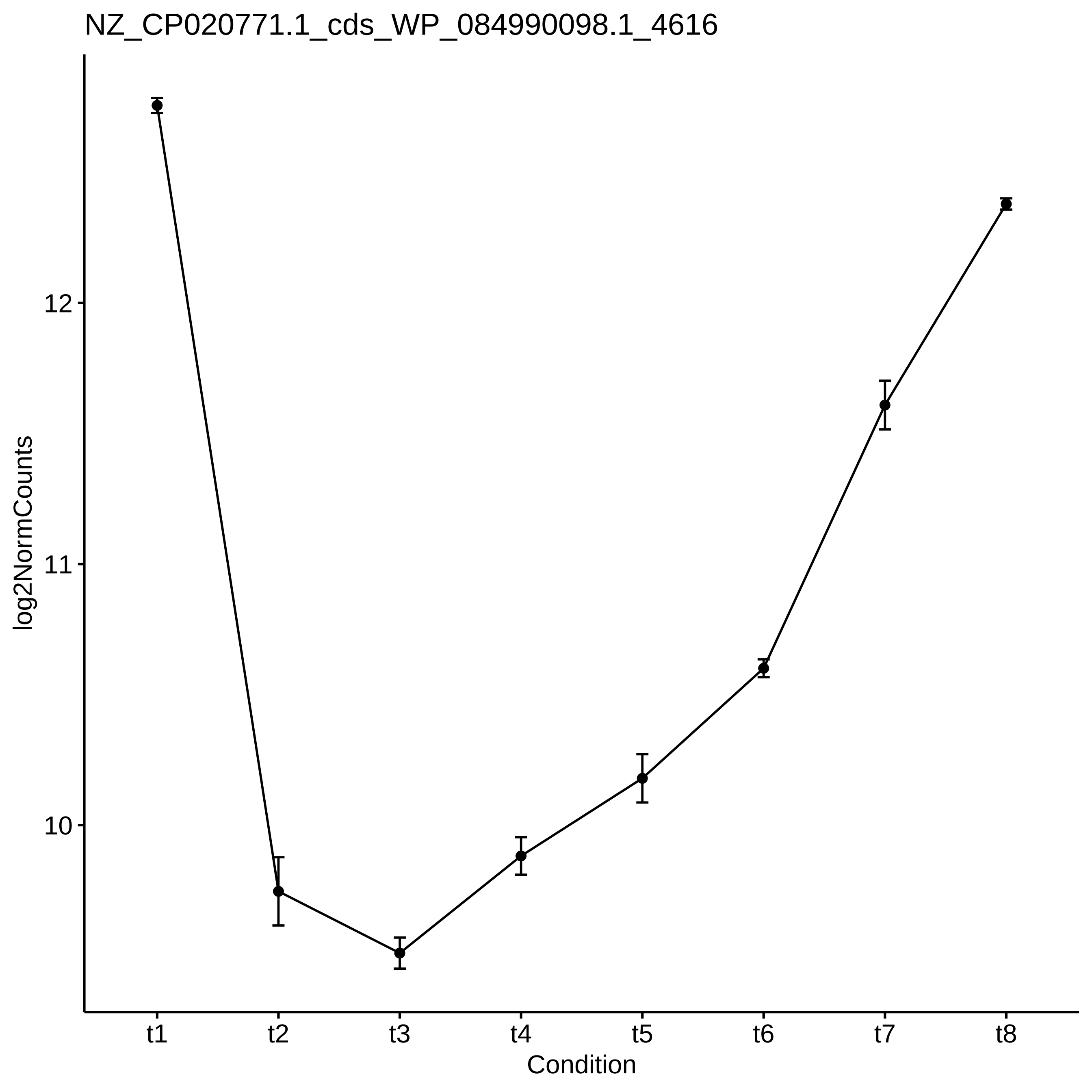

NZ_CP020771.1_cds_WP_084990098.1_4616
log2NormCounts
12
11
10
t1
t2
t3
t4
t5
t6
t7
t8
Condition

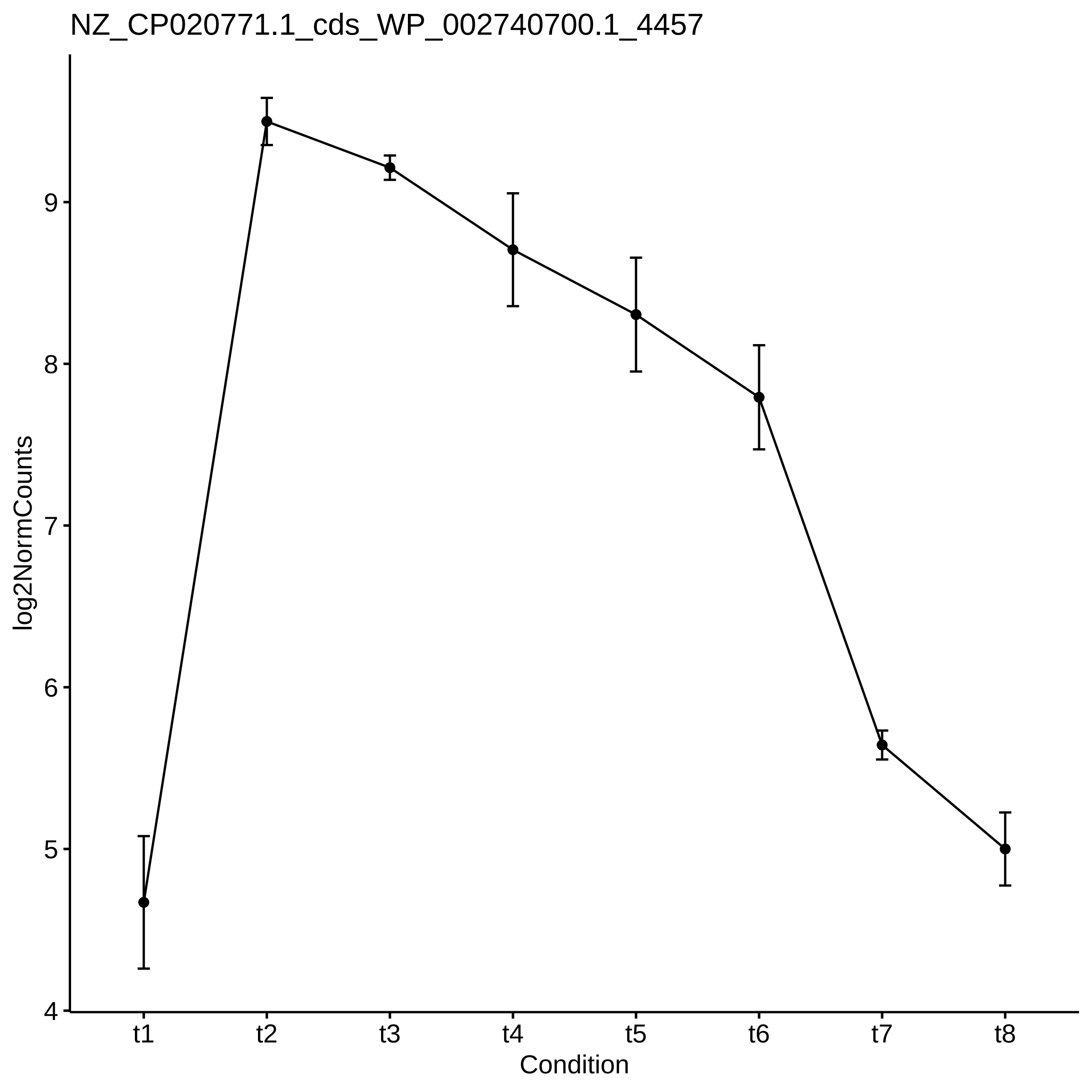

NZ_CP020771.1_cds_WP_002740700.1_4457
log2NormCounts
9
8
7
6
5
4
t1
t2
t3
t4
t5
t6
t7
t8
Condition

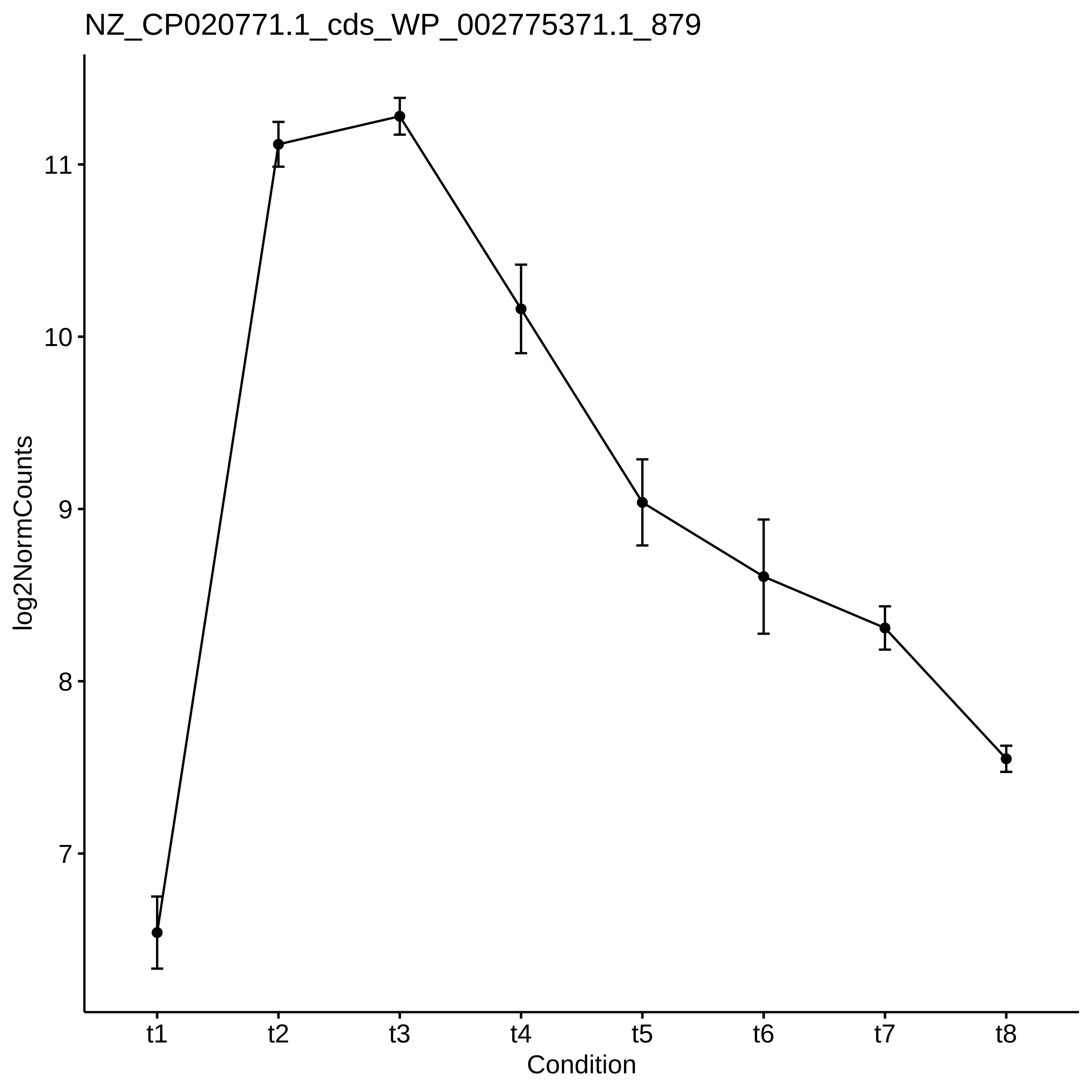

NZ_CP020771.1_cds_WP_002775371.1_879
log2NormCounts
11
10
9
8
7
t1
t2
t3
t4
t5
t6
t7
t8
Condition

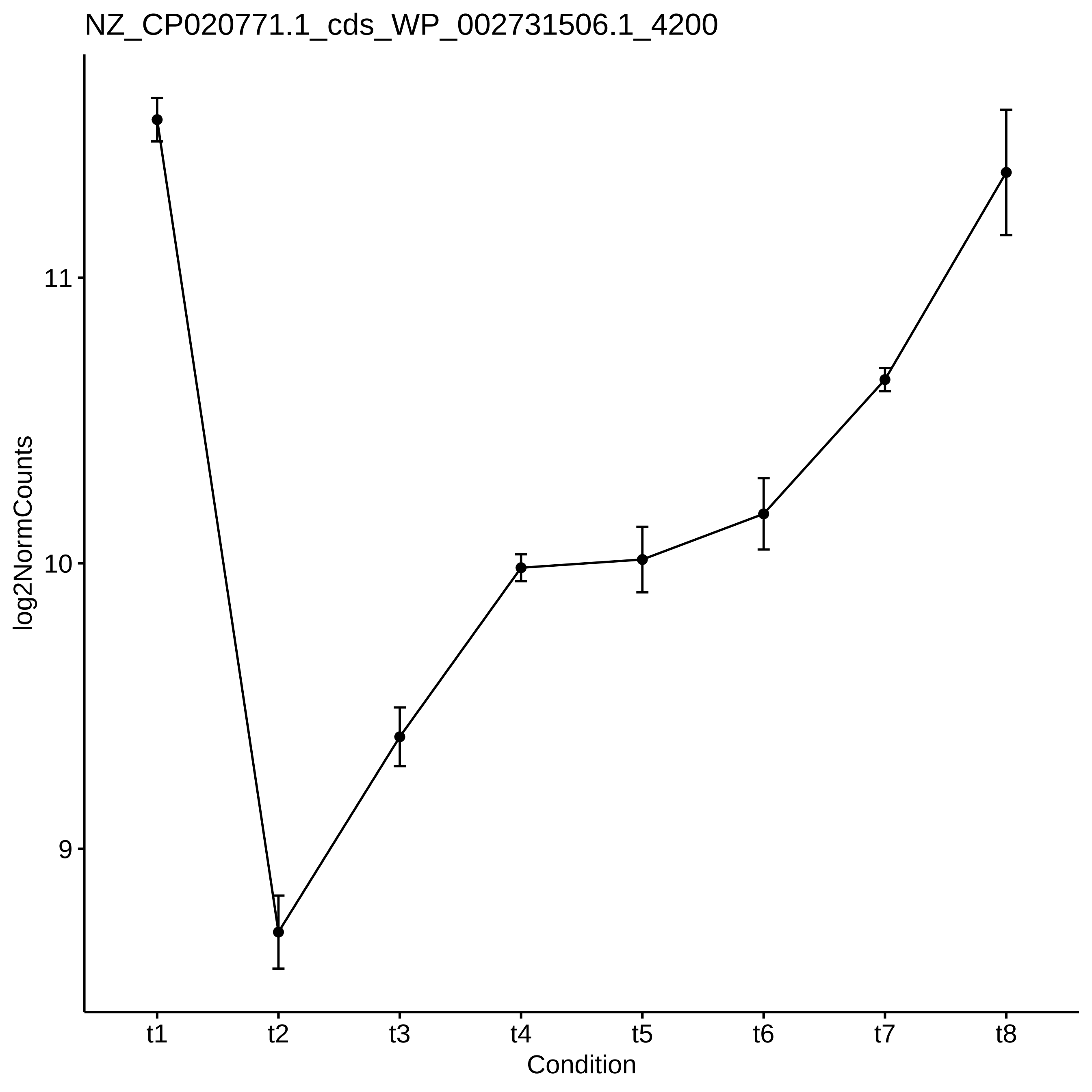
NZ_CP020771.1_cds_WP_002731506.1_4200
log2NormCounts
11
10
9
t1
t2
t3
t4
t5
t6
t7
t8
Condition

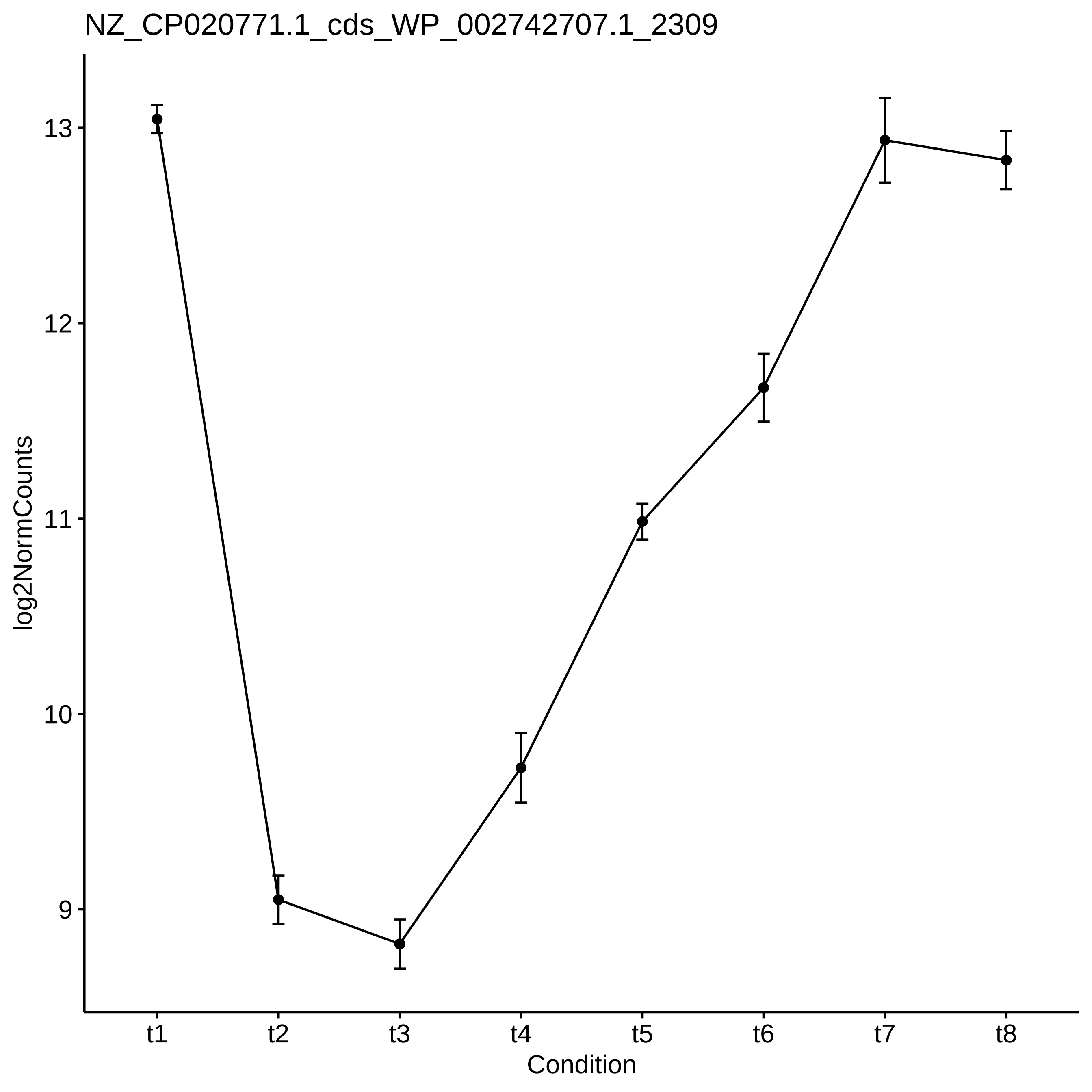

NZ_CP020771.1_cds_WP_002742707.1_2309
13
12
11
10
9
log2NormCounts
t1
t2
t3
t4
t5
t6
t7
t8
Condition

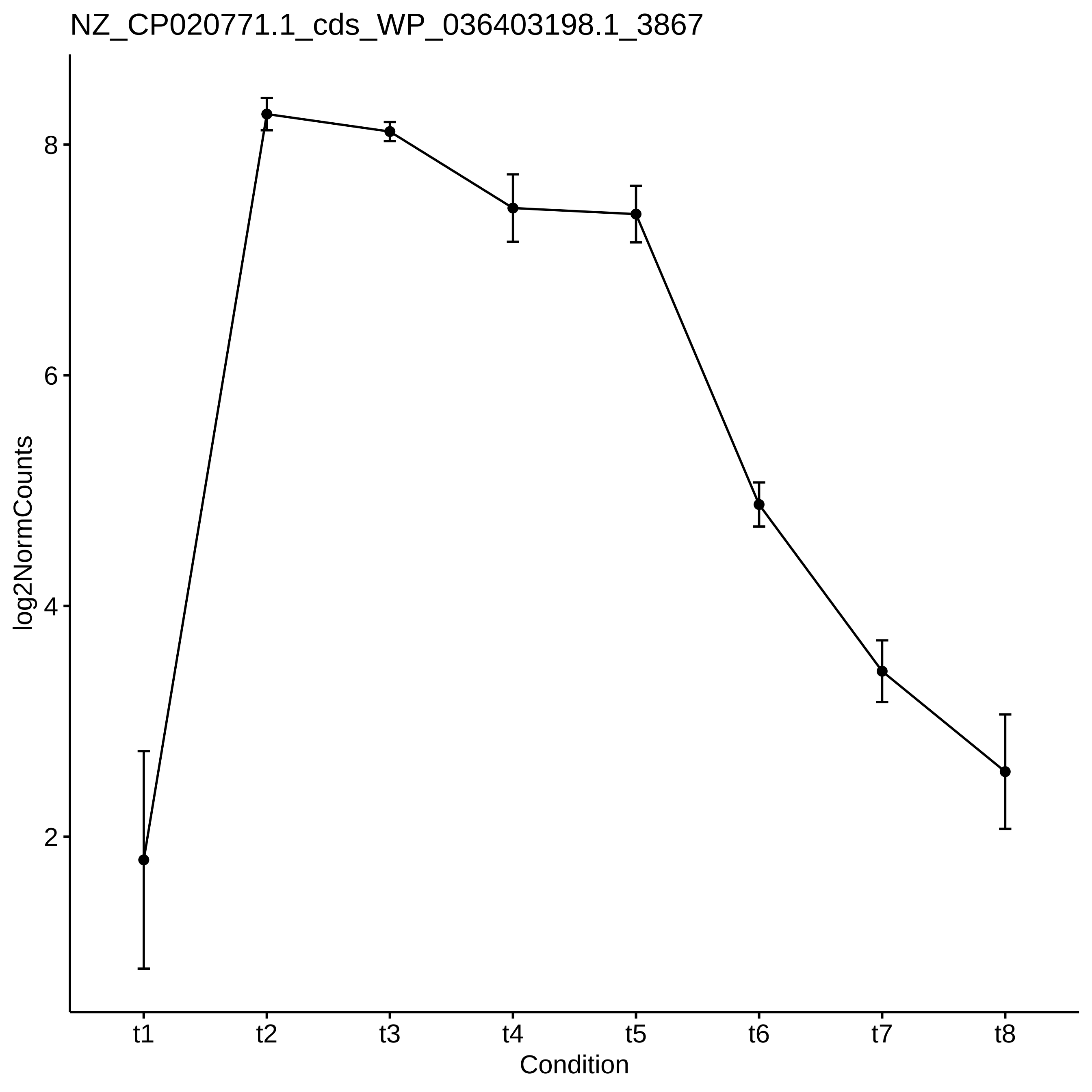

NZ_CP020771.1_cds_WP_036403198.1_3867
log2NormCounts
8
6
4
2
t1
t2
t3
t4
t5
t6
t7
t8
Condition

NZ_CP020771.1_cds_WP_002743493.1_935

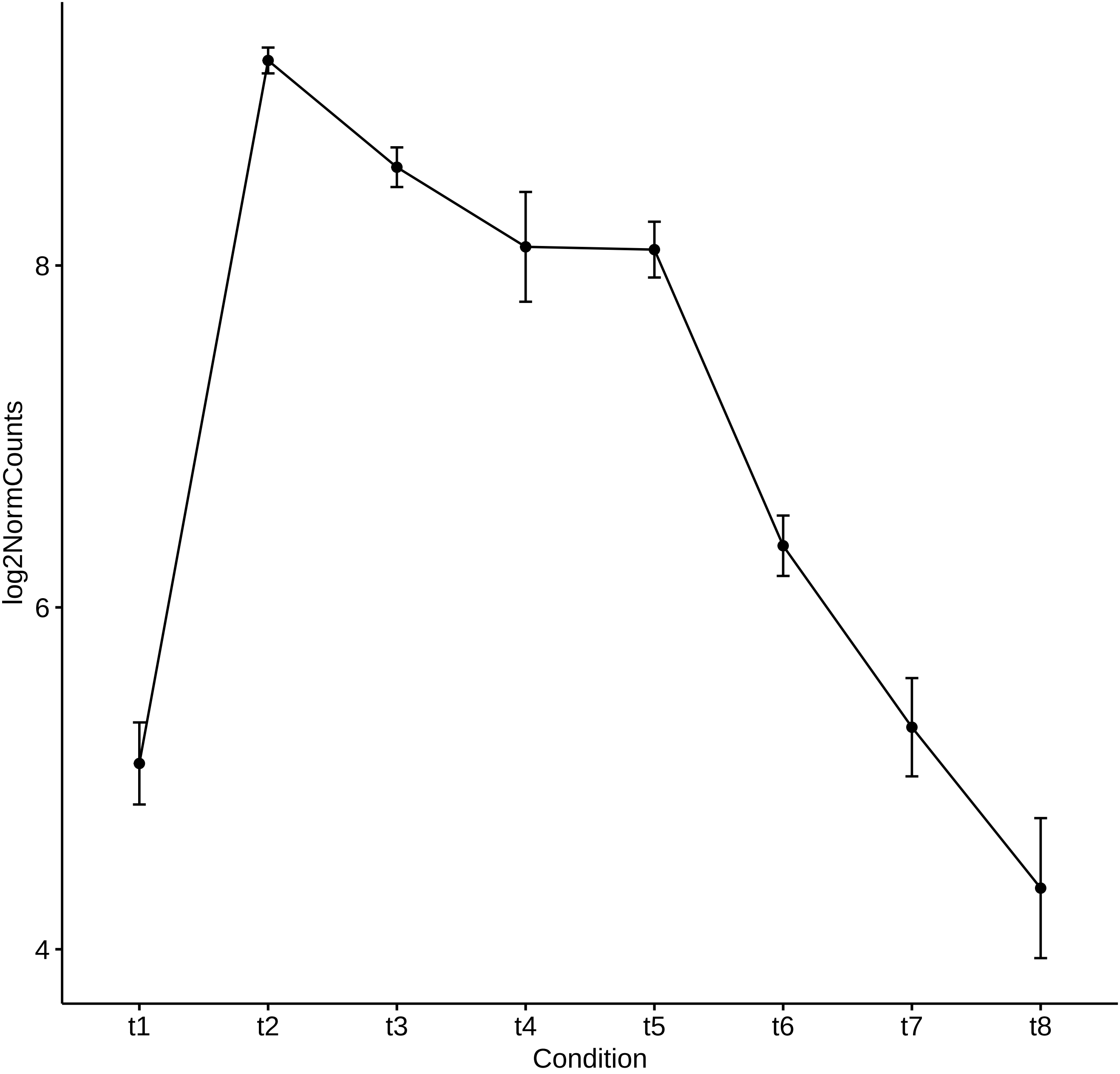

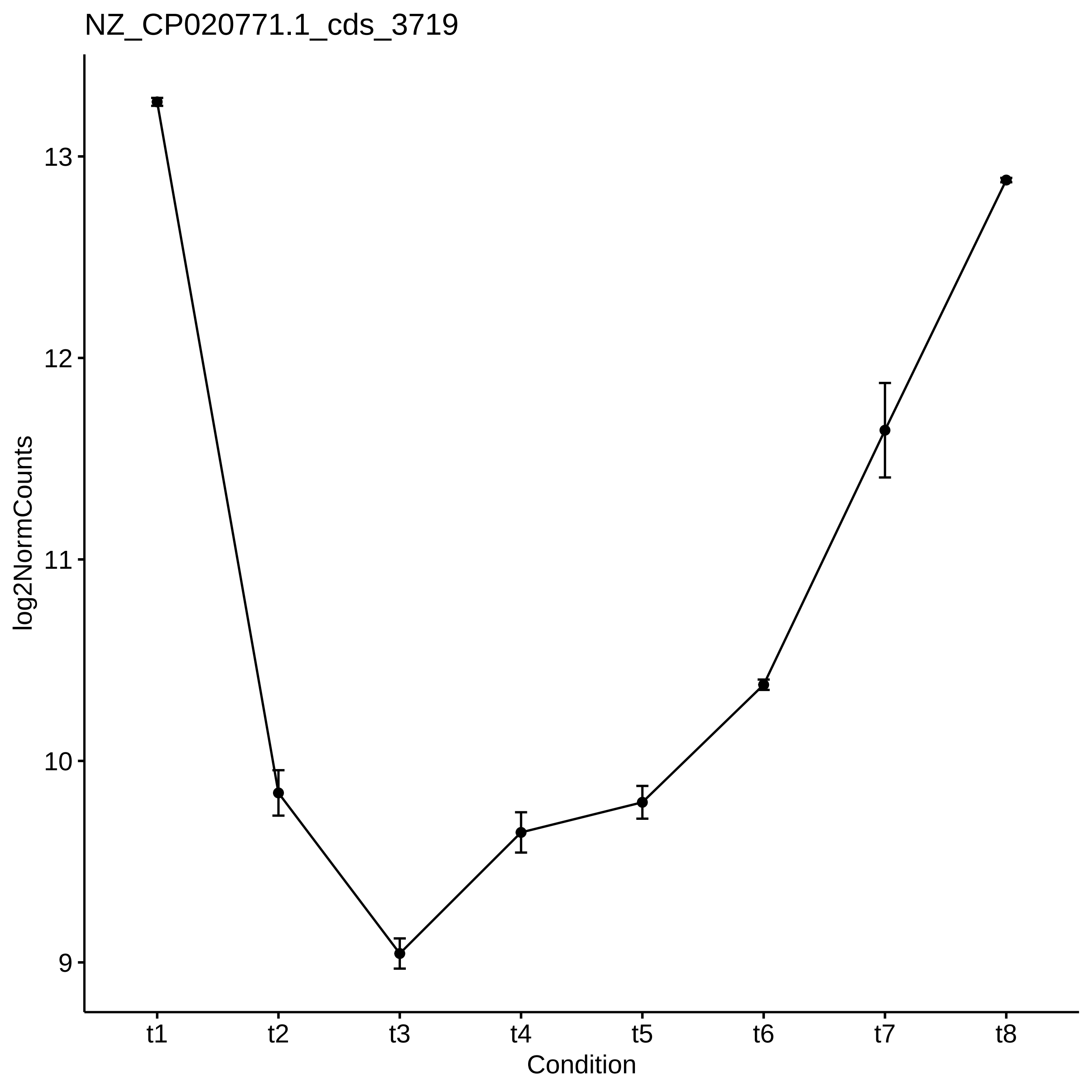
NZ_CP020771.1_cds_3719
log2NormCounts
13
12
11
10
9
t1
t2
t3
t4
t5
t6
t7
t8
Condition

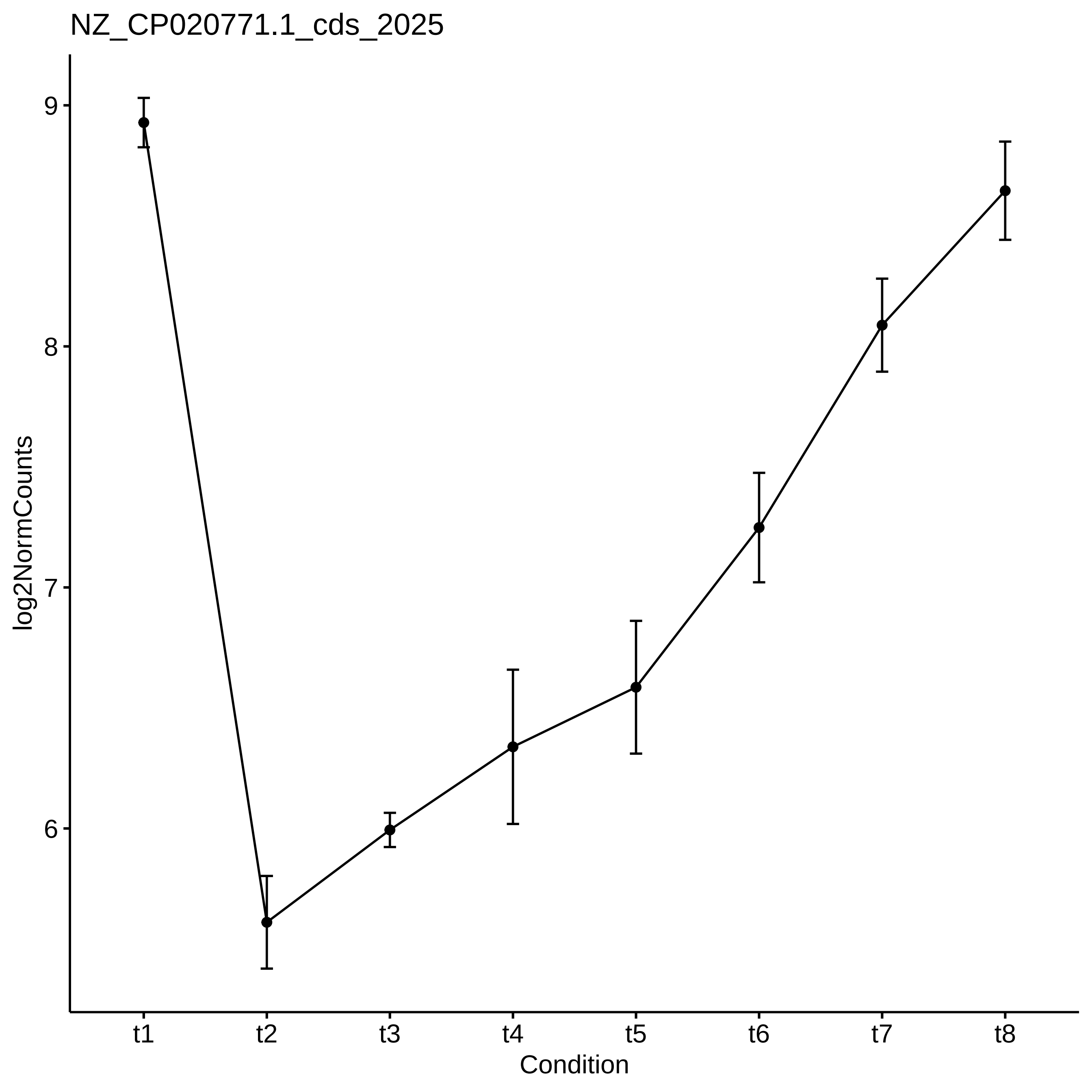
NZ_CP020771.1_cds_2025
log2NormCounts
9
8
7
6
t1
t2
t3
t4
t5
t6
t7
t8
Condition

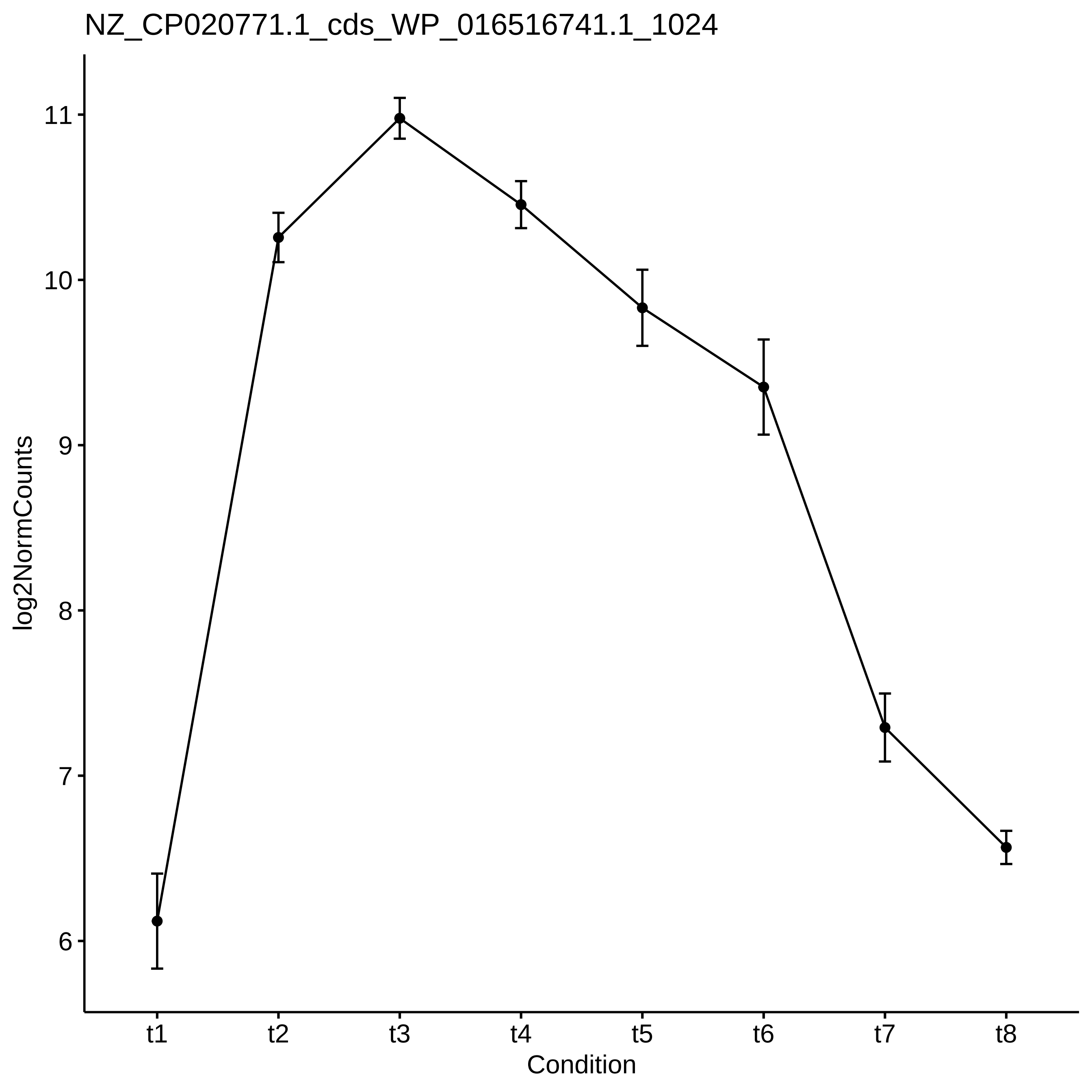

NZ_CP020771.1_cds_WP_016516741.1_1024
log2NormCounts
11
10
9
8
7
6
t1
t2
t3
t4
t5
t6
t7
t8
Condition

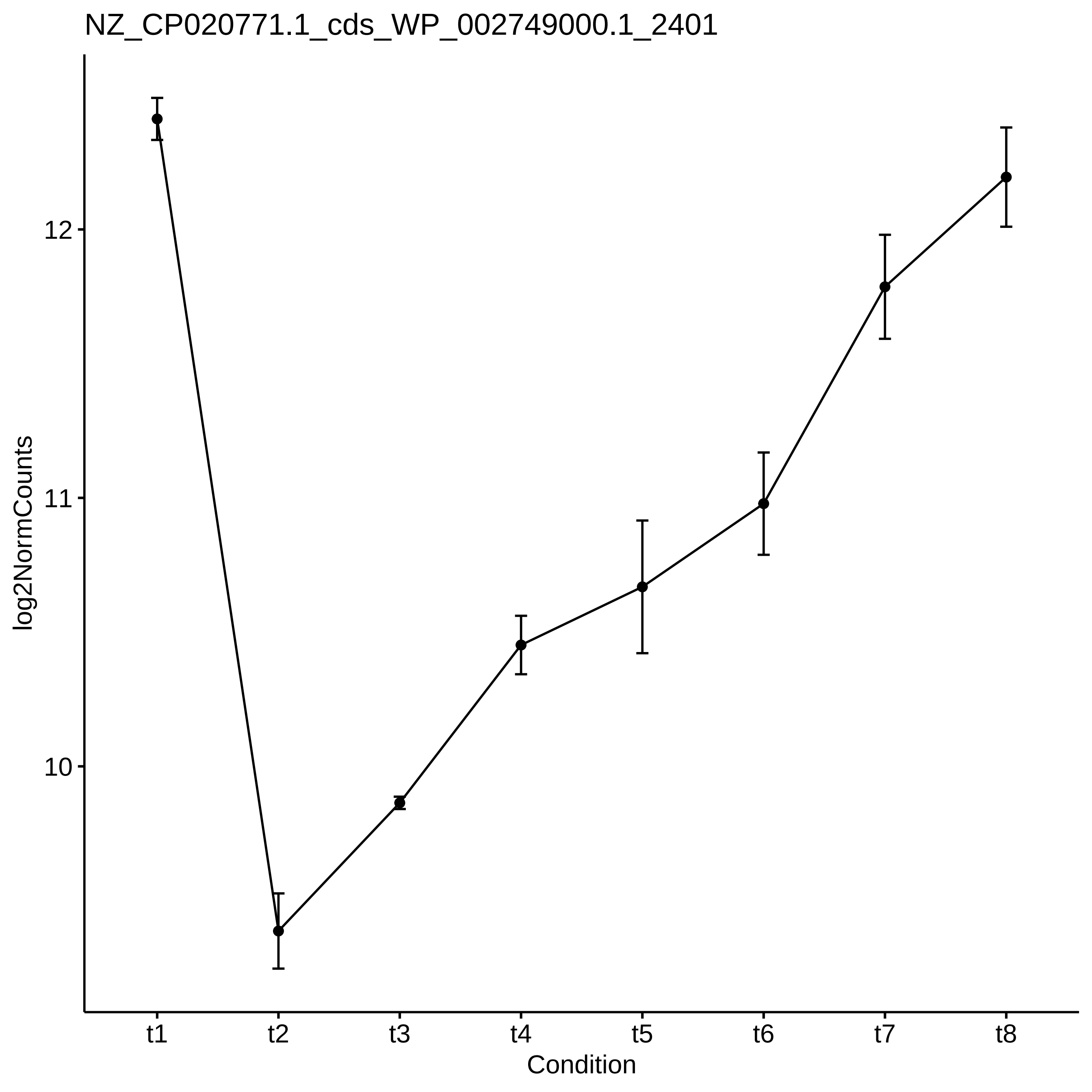

NZ_CP020771.1_cds_WP_002749000.1_2401
log2NormCounts
12
11
10
t1
t2
t3
t4
t5
t6
t7
t8
Condition

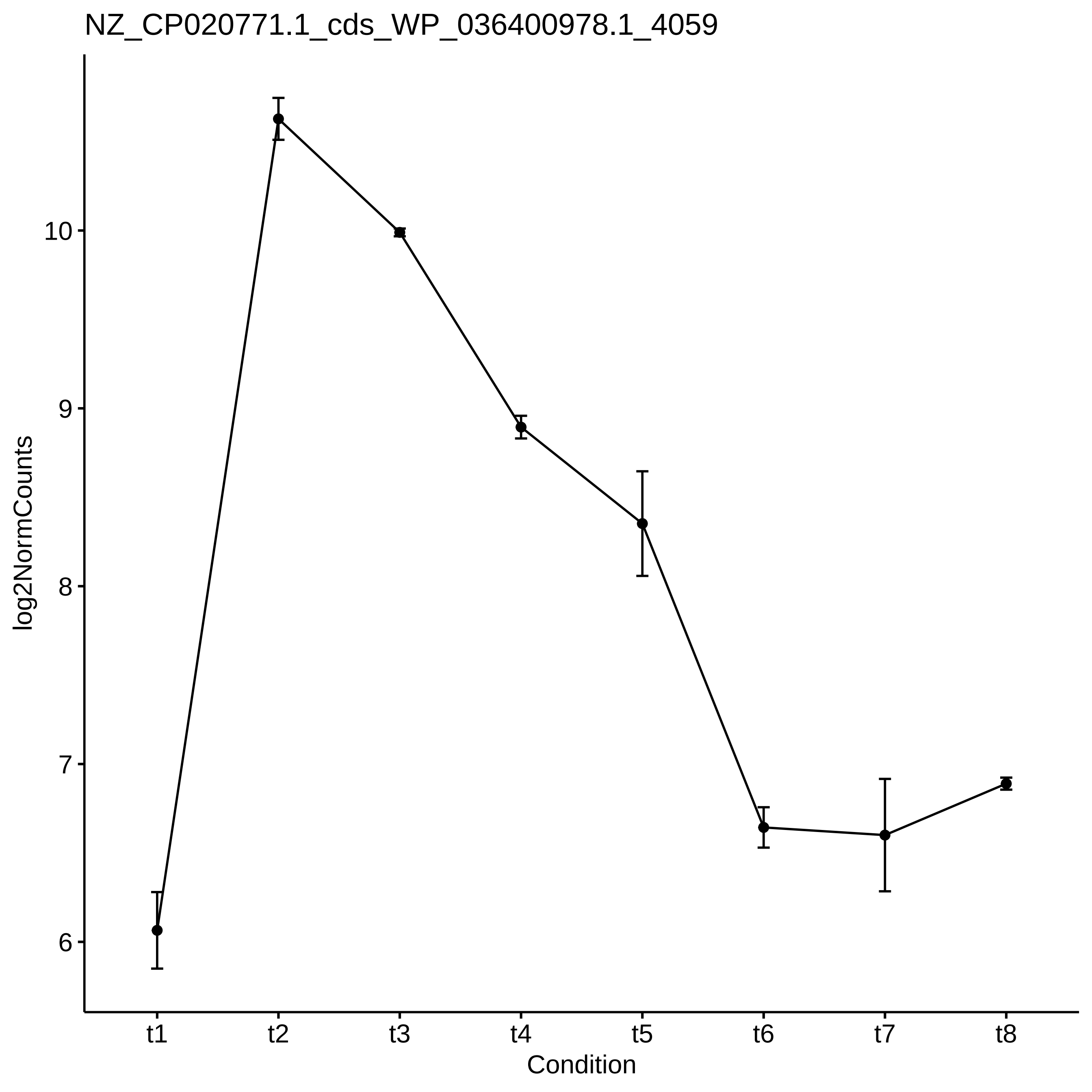
NZ_CP020771.1_cds_WP_036400978.1_4059
log2NormCounts
10
9
8
7
6
t1
t2
t3
t4
t5
t6
t7
t8
Condition

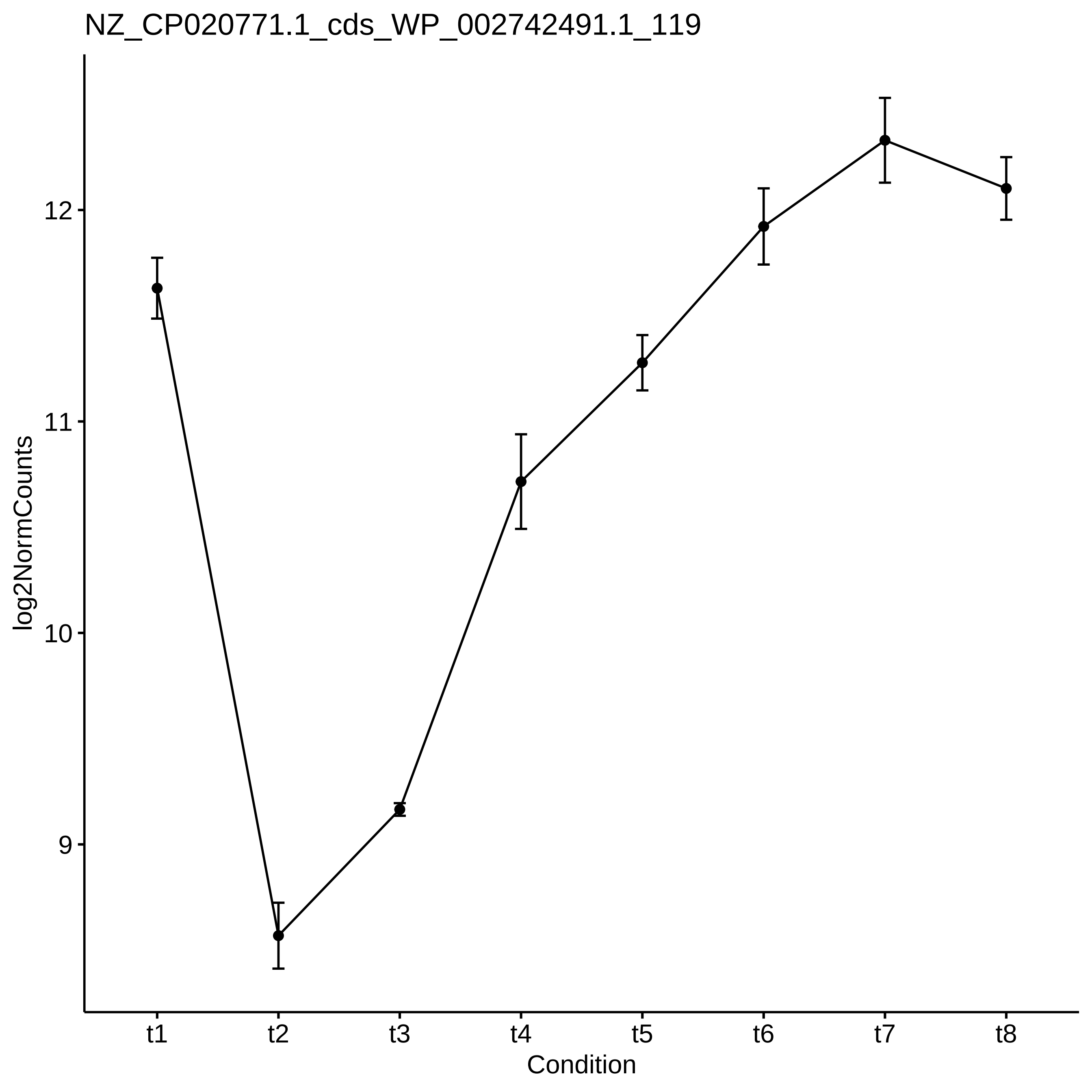

NZ_CP020771.1_cds_WP_002742491.1_119
log2NormCounts
12
11
10
9
t1
t2
t3
t4
t5
t6
t7
t8
Condition

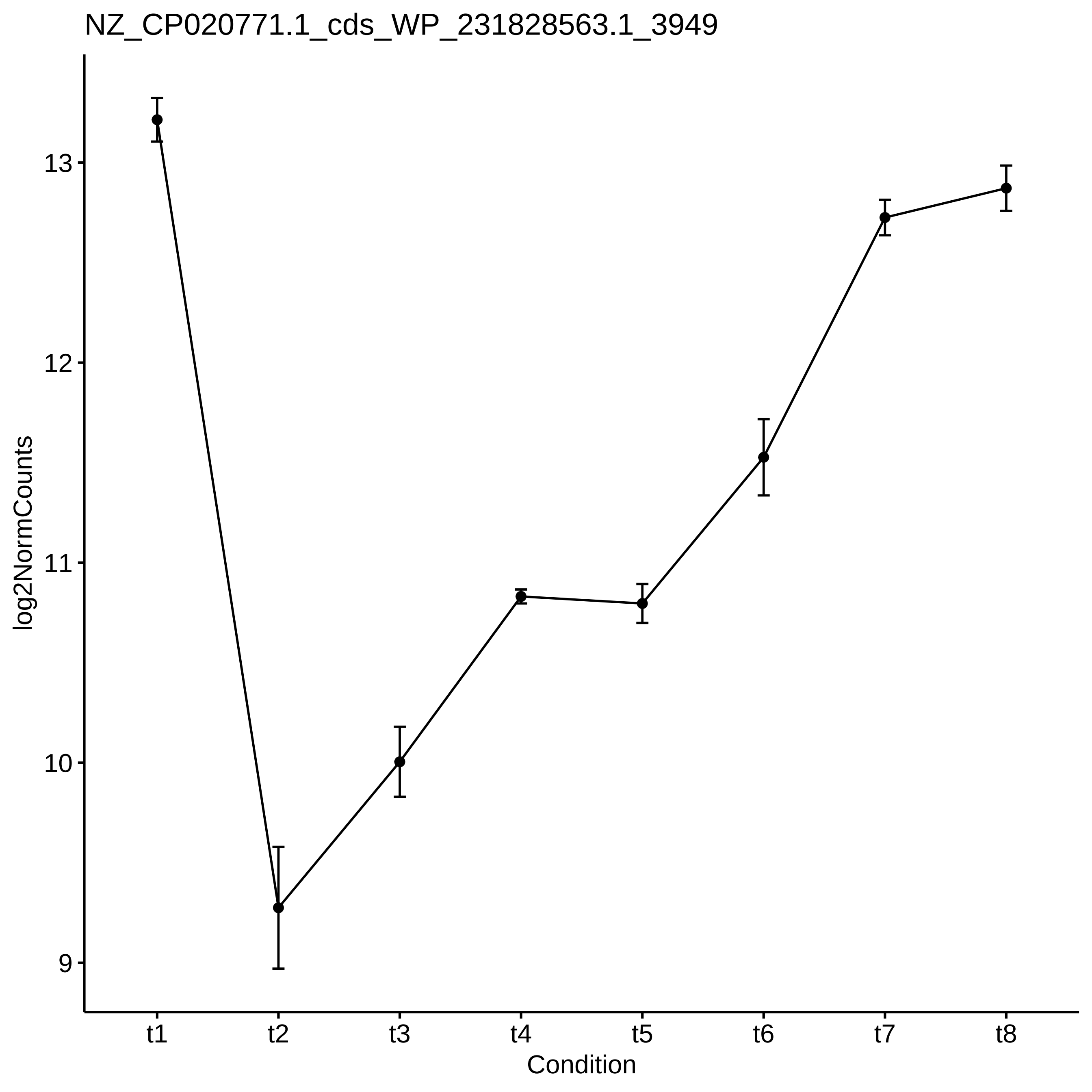
NZ_CP020771.1_cds_WP_231828563.1_3949
log2NormCounts
13
12
11
10
9
t1
t2
t3
t4
t5
t6
t7
t8
Condition

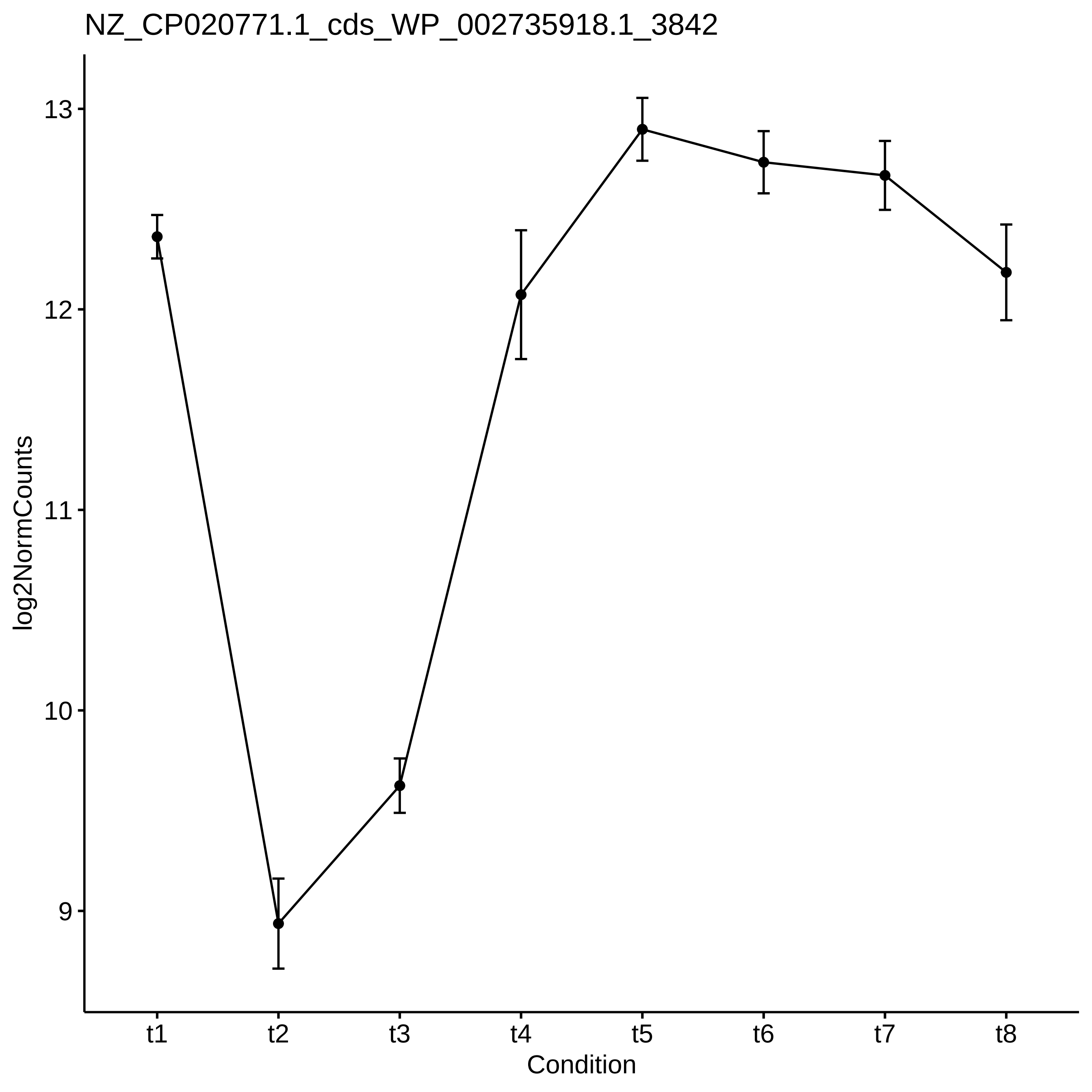

NZ_CP020771.1_cds_WP_002735918.1_3842
log2NormCounts
13
12
11
10
9
t1
t2
t3
t4
t5
t6
t7
t8
Condition

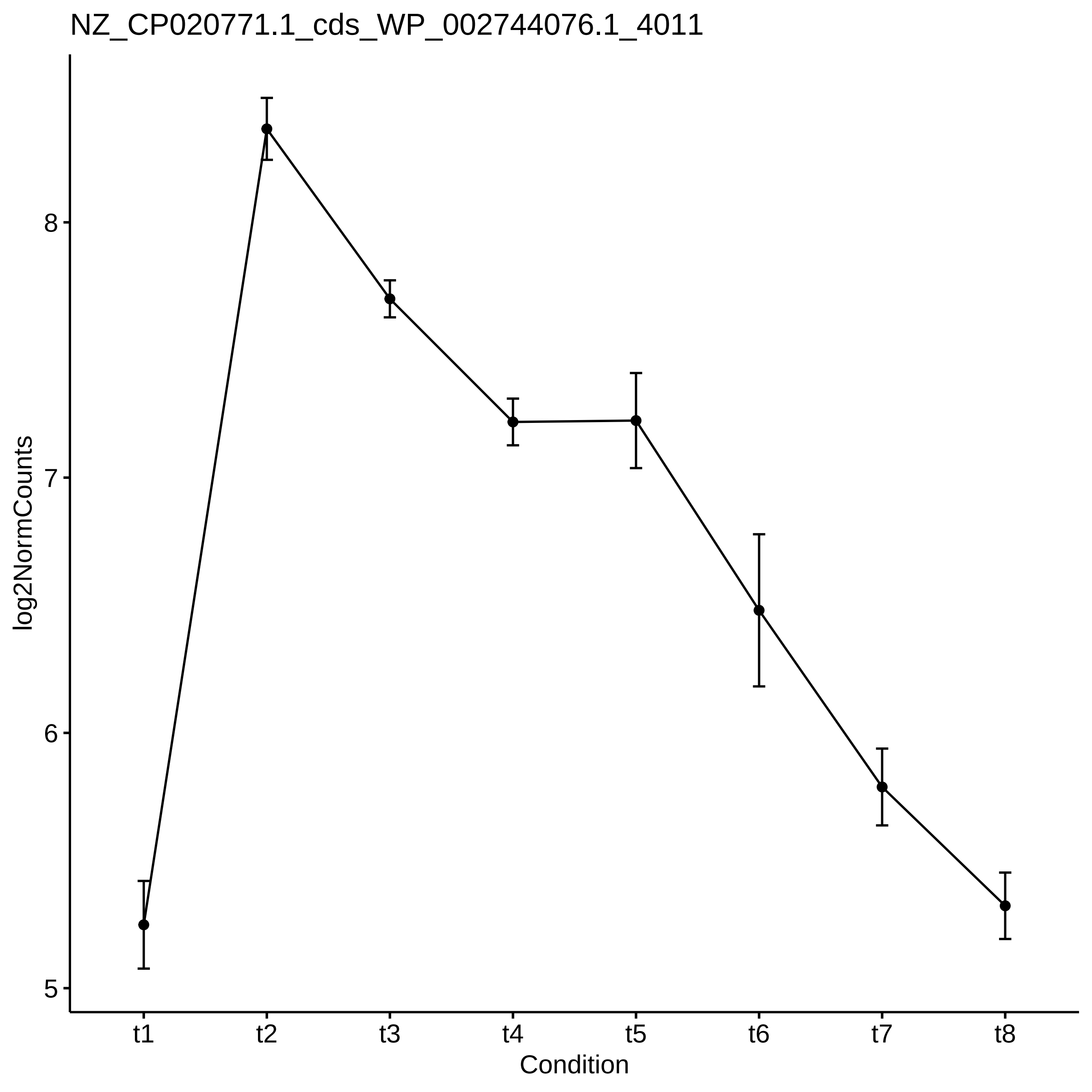
NZ_CP020771.1_cds_WP_002744076.1_4011
log2NormCounts
8
7
6
5
t1
t2
t3
t4
t5
t6
t7
t8
Condition

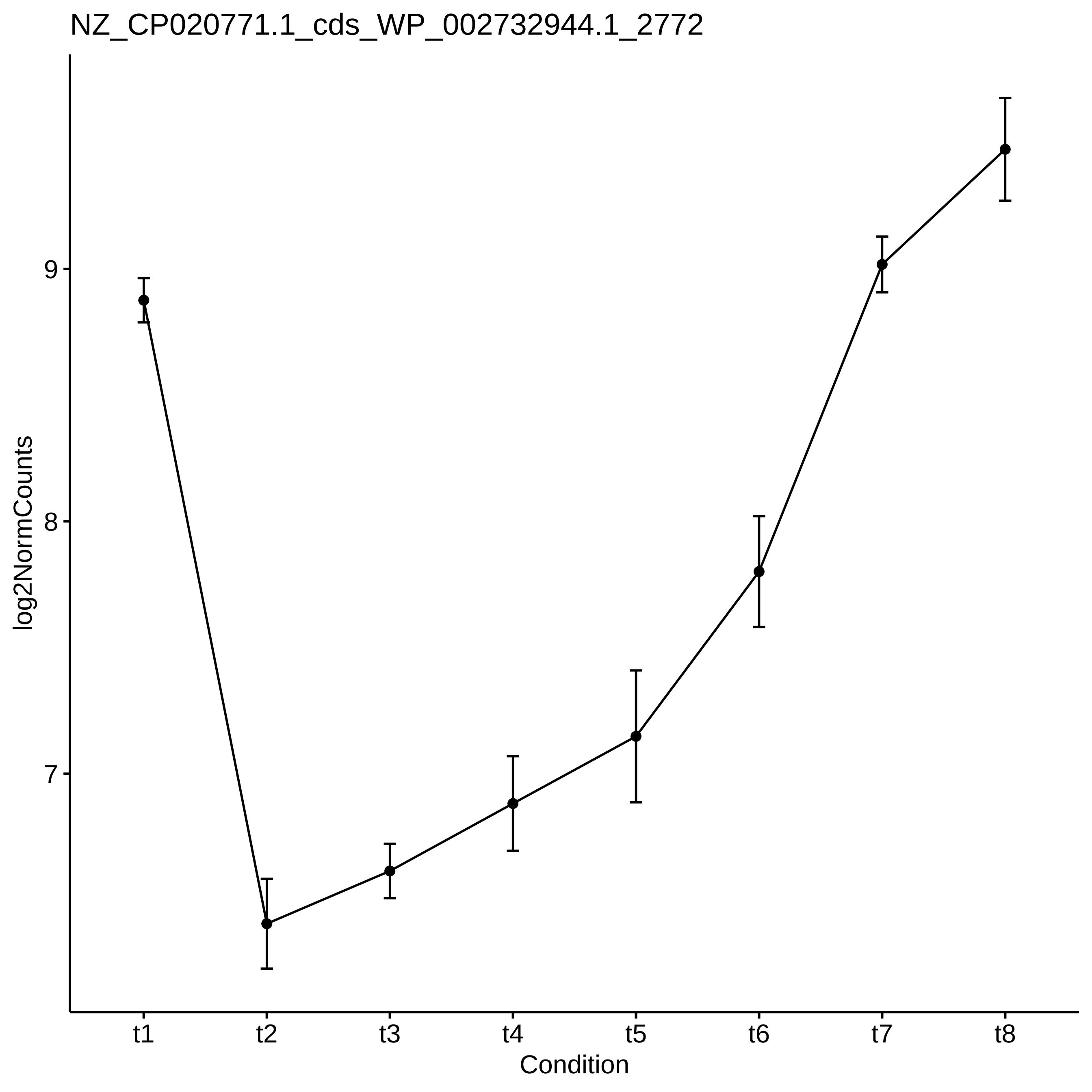
NZ_CP020771.1_cds_WP_002732944.1_2772
log2NormCounts
9
8
7
t1
t2
t3
t4
t5
t6
t7
t8
Condition

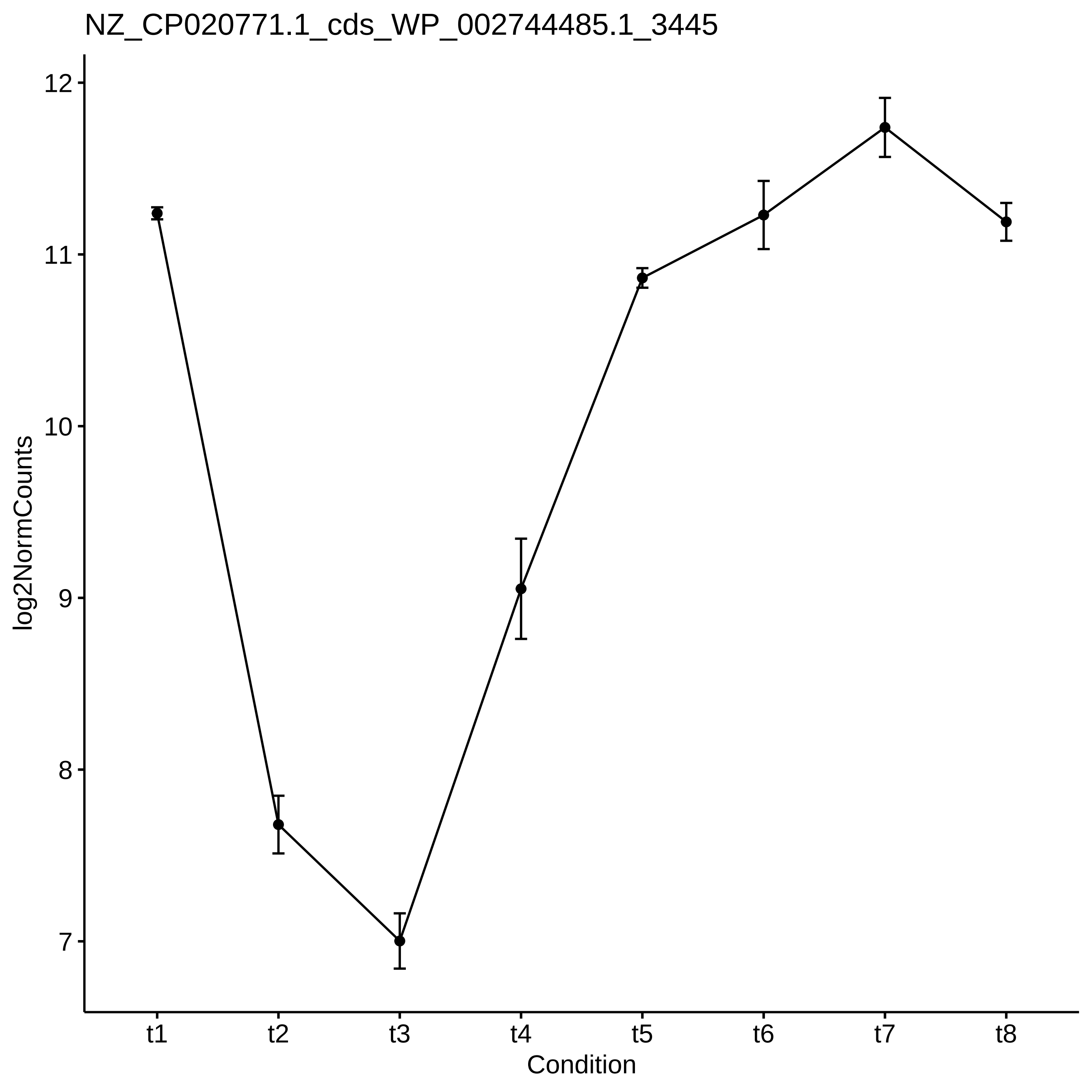
NZ_CP020771.1_cds_WP_002744485.1_3445
log2NormCounts
12
11
10
9
8
7
t1
t2
t3
t4
t5
t6
t7
t8
Condition

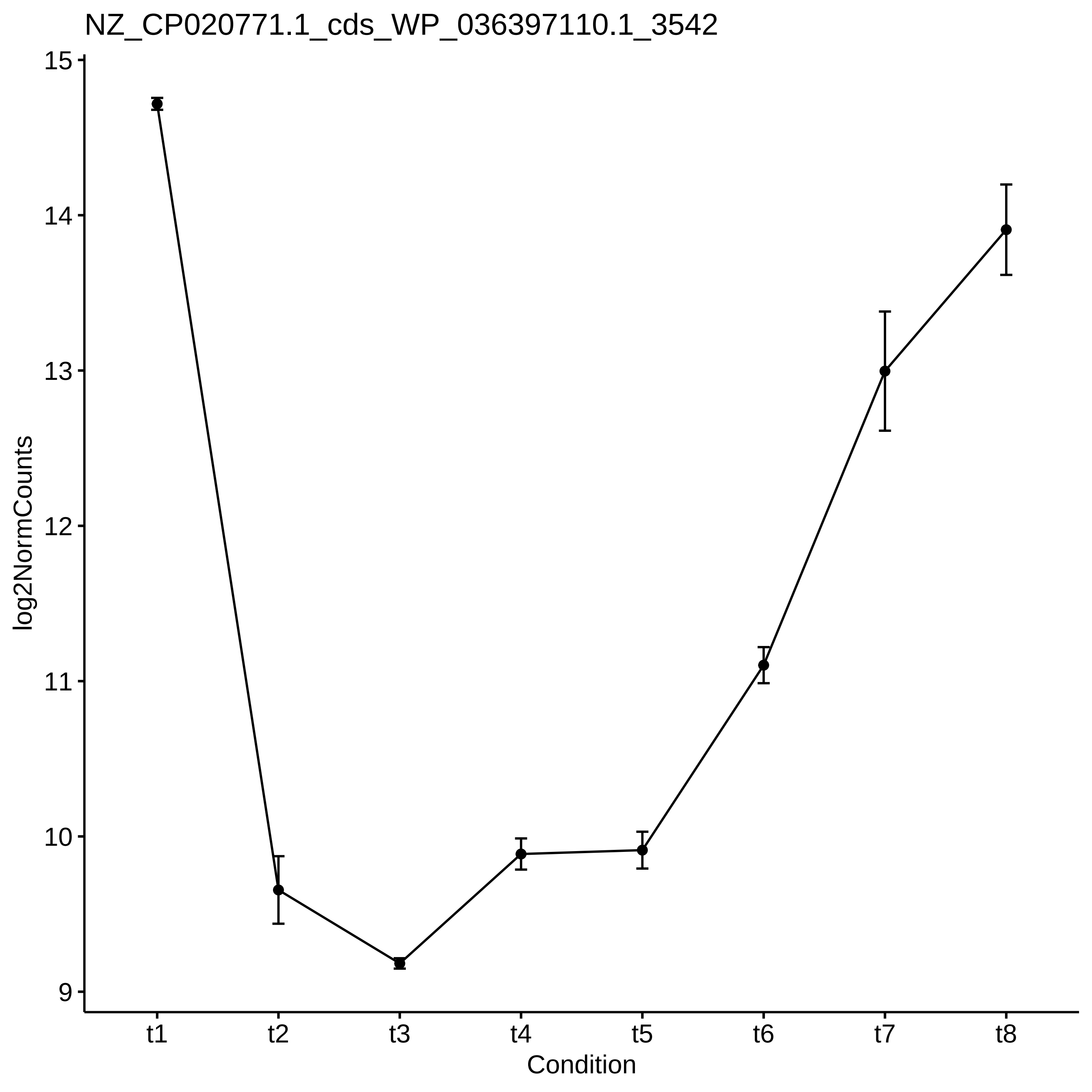

NZ_CP020771.1_cds_WP_036397110.1_3542
log2NormCounts
15
14
13
12
11
10
9
t1
t2
t3
t4
t5
t6
t7
t8
Condition

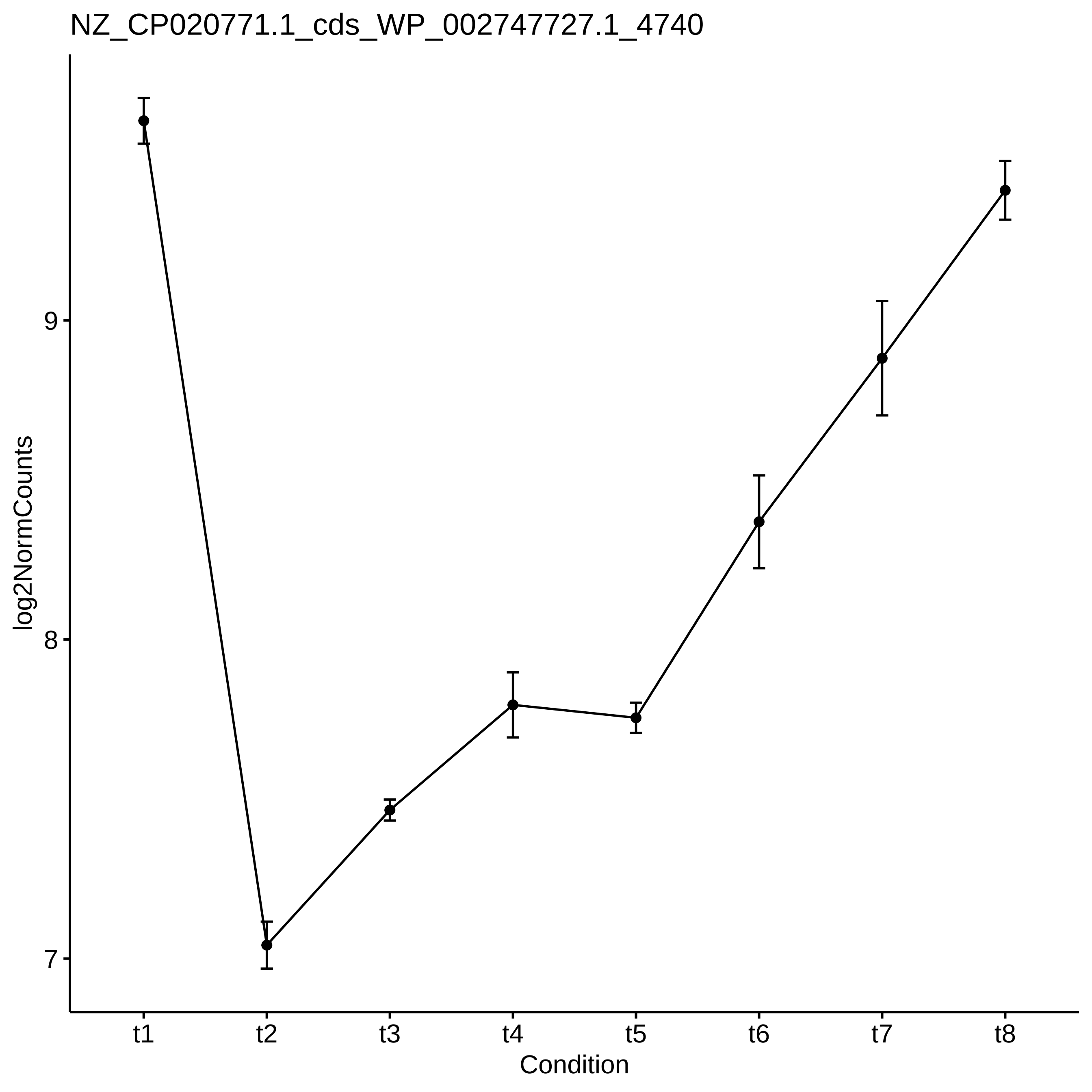
NZ_CP020771.1_cds_WP_002747727.1_4740
log2NormCounts
9
8
7
t1
t2
t3
t4
t5
t6
t7
t8
Condition

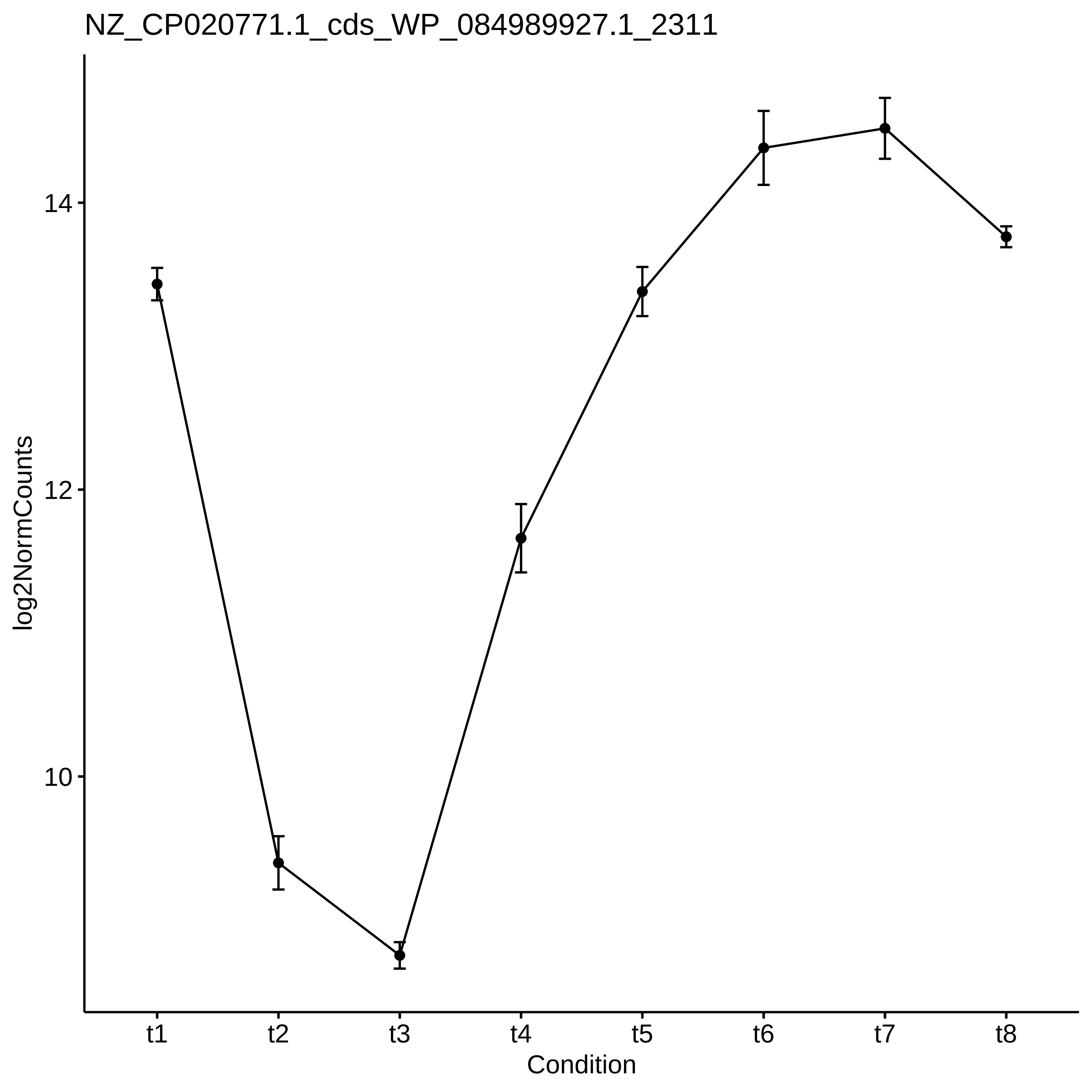

NZ_CP020771.1_cds_WP_084989927.1_2311
log2NormCounts
14
12
10
t1
t2
t3
t4
t5
t6
t7
t8
Condition

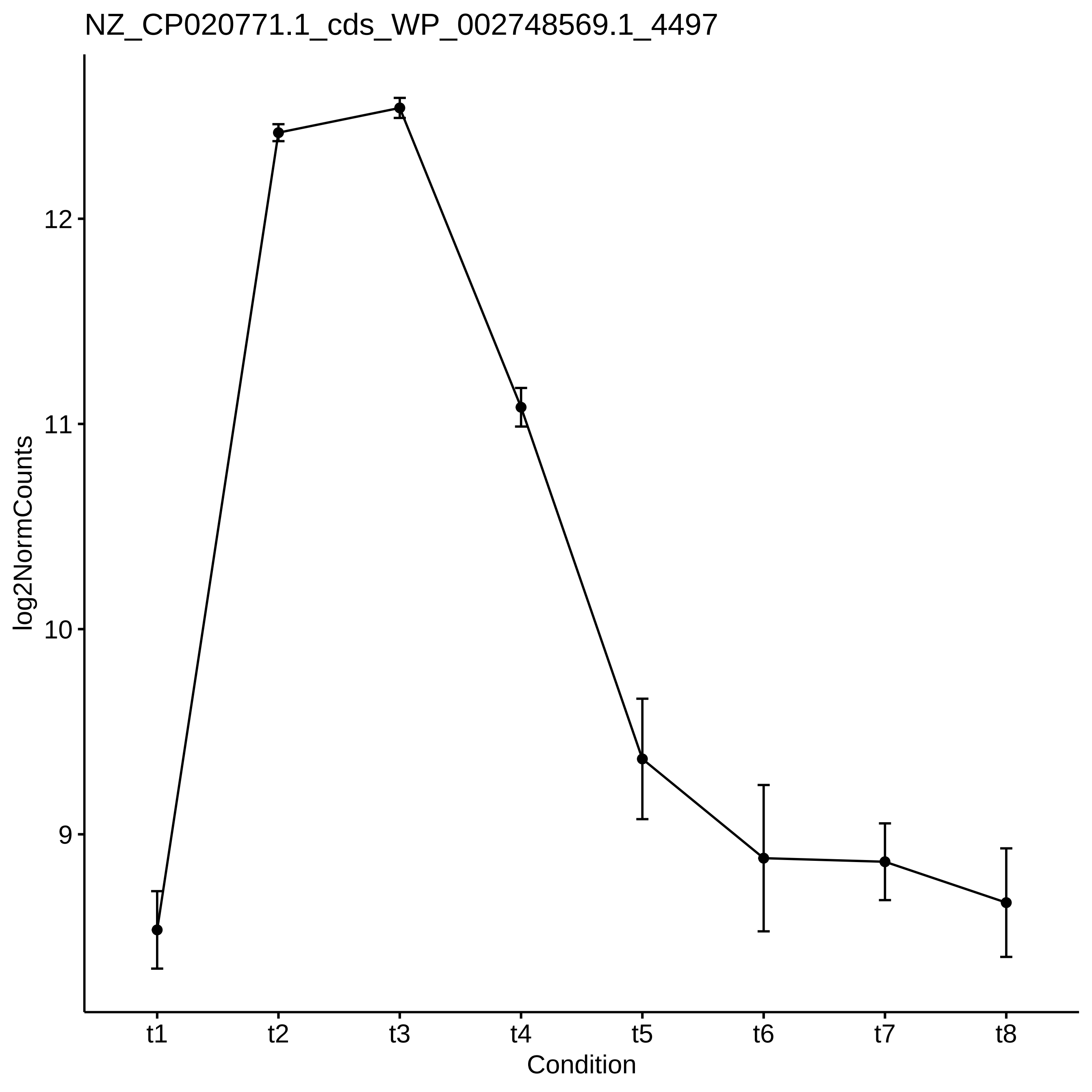
NZ_CP020771.1_cds_WP_002748569.1_4497
log2NormCounts
12
11
10
9
t1
t2
t3
t4
t5
t6
t7
t8
Condition
