## Supplementary_Data for "Diel changes in the expression of a marker gene and candidate genes for intracellular amorphous CaCO_3_ biomineralization in *Microcystis*": PCC7806_[t4-t6]_Top50_Profile.pdf

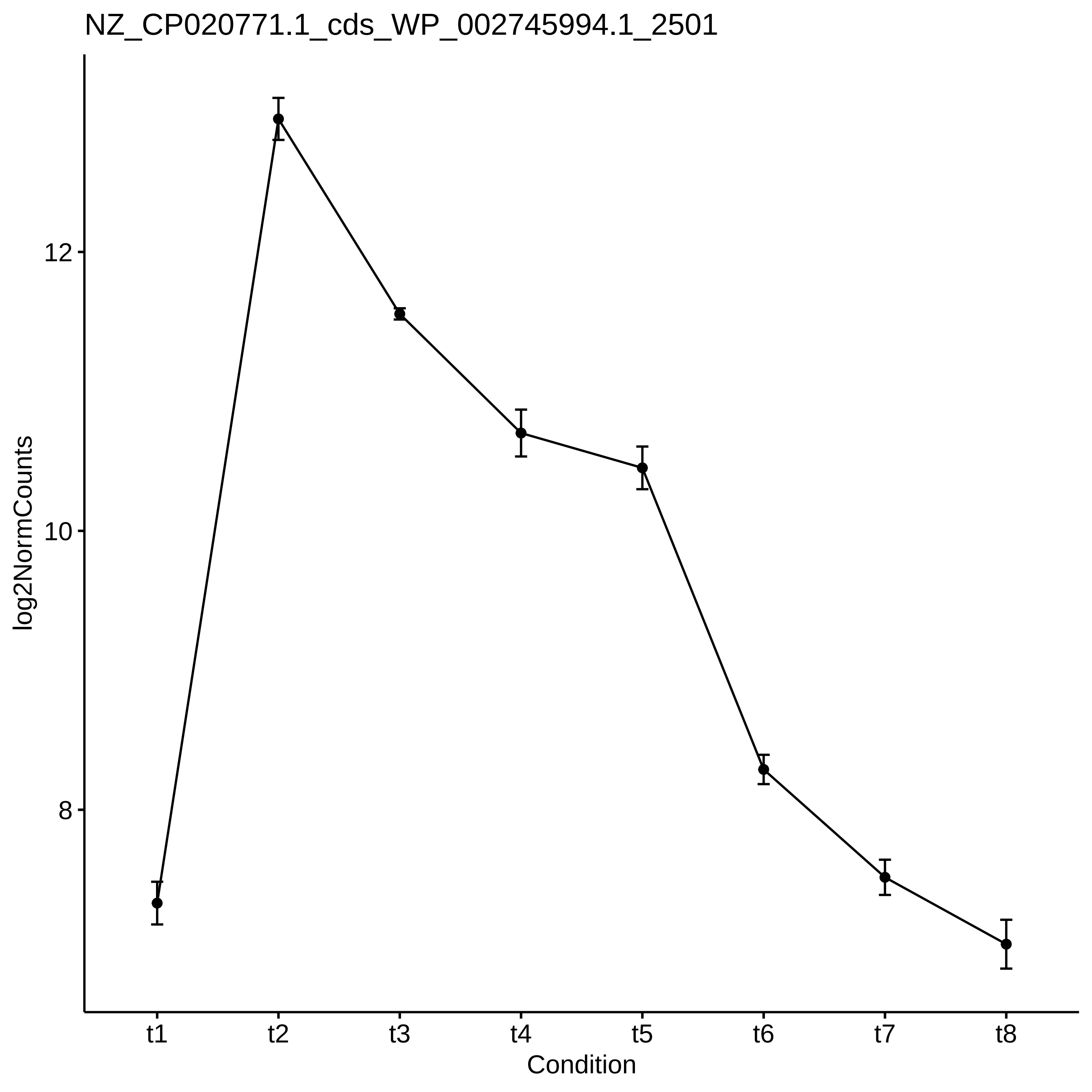

NZ_CP020771.1_cds_WP_002745994.1_2501
log2NormCounts
12
10
8
t1
t2
t3
t4
t5
t6
t7
t8
Condition

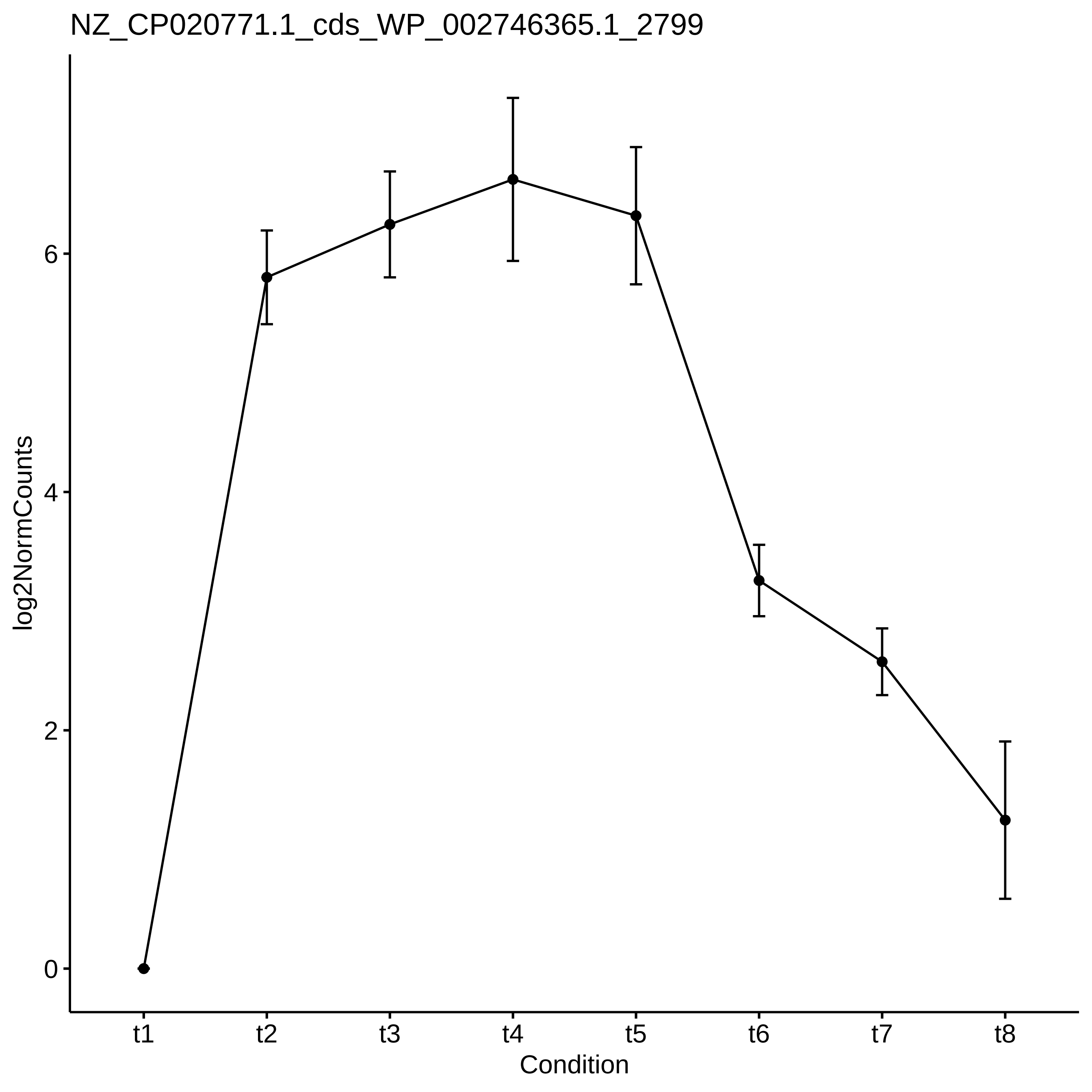
NZ_CP020771.1_cds_WP_002746365.1_2799
log2NormCounts
0
2
4
6
t1
t2
t3
t4
t5
t6
t7
t8
Condition

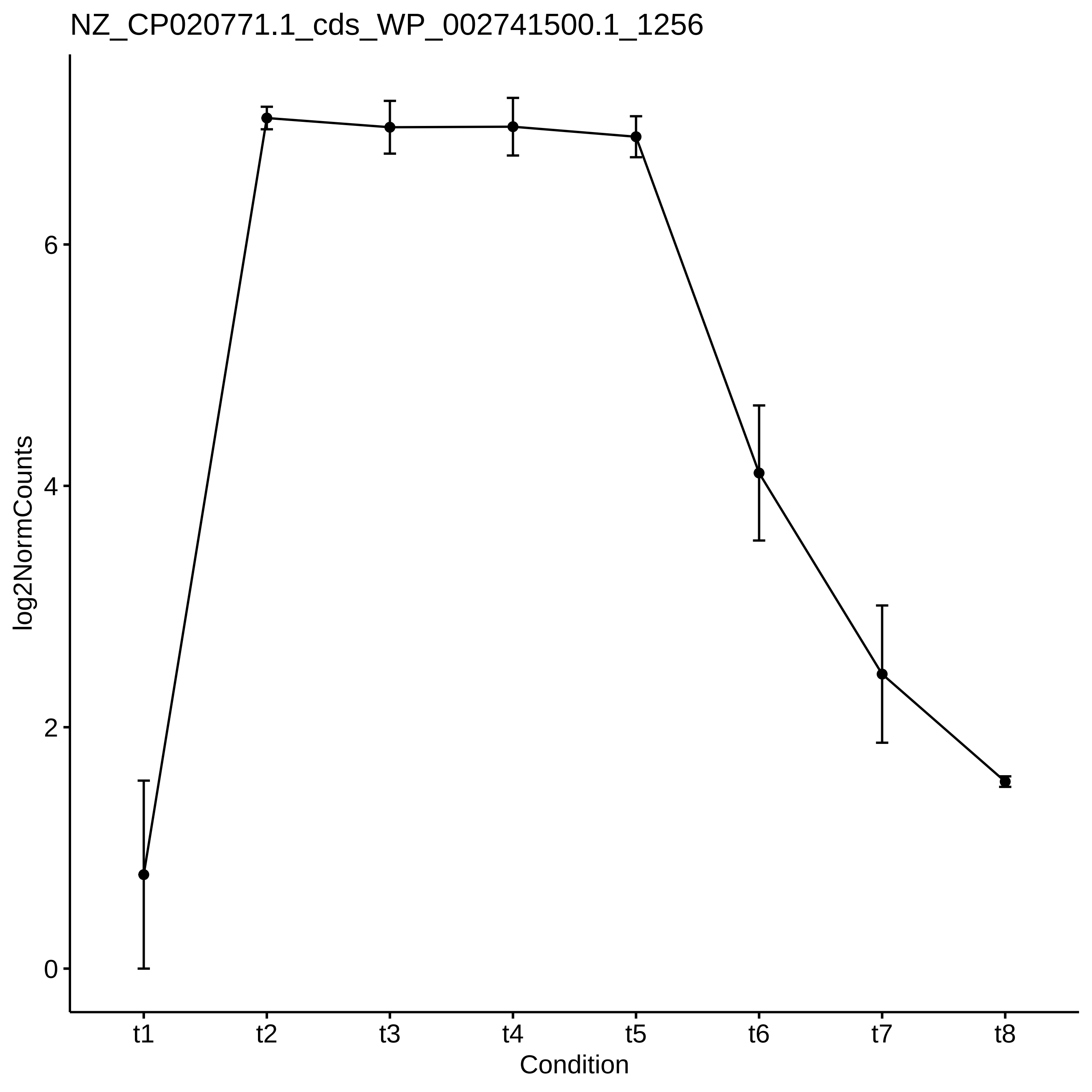

NZ_CP020771.1_cds_WP_002741500.1_1256
log2NormCounts
0
2
4
6
t1
t2
t3
t4
t5
t6
t7
t8
Condition

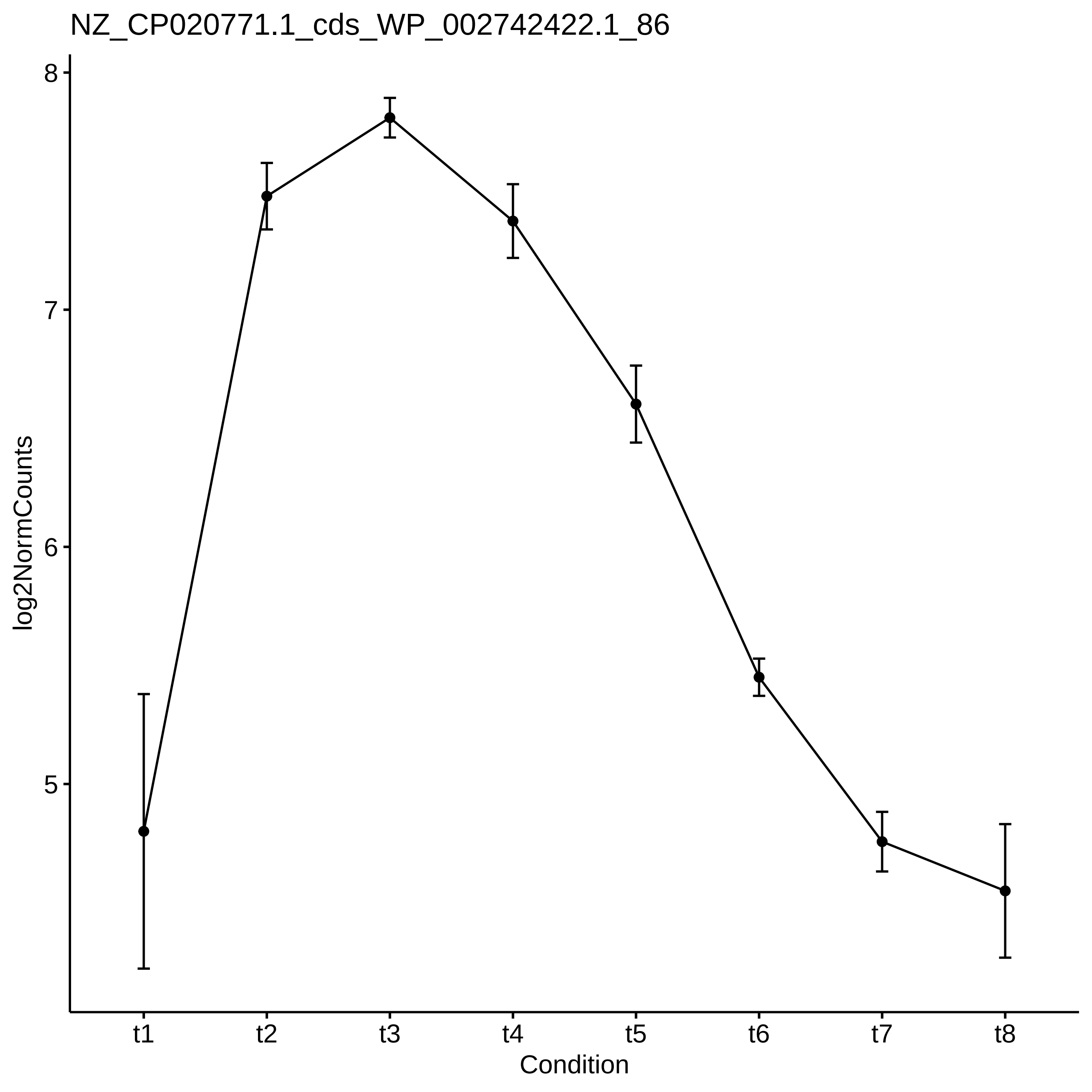

NZ_CP020771.1_cds_WP_002742422.1_86
log2NormCounts
8
7
6
5
t1
t2
t3
t4
t5
t6
t7
t8
Condition

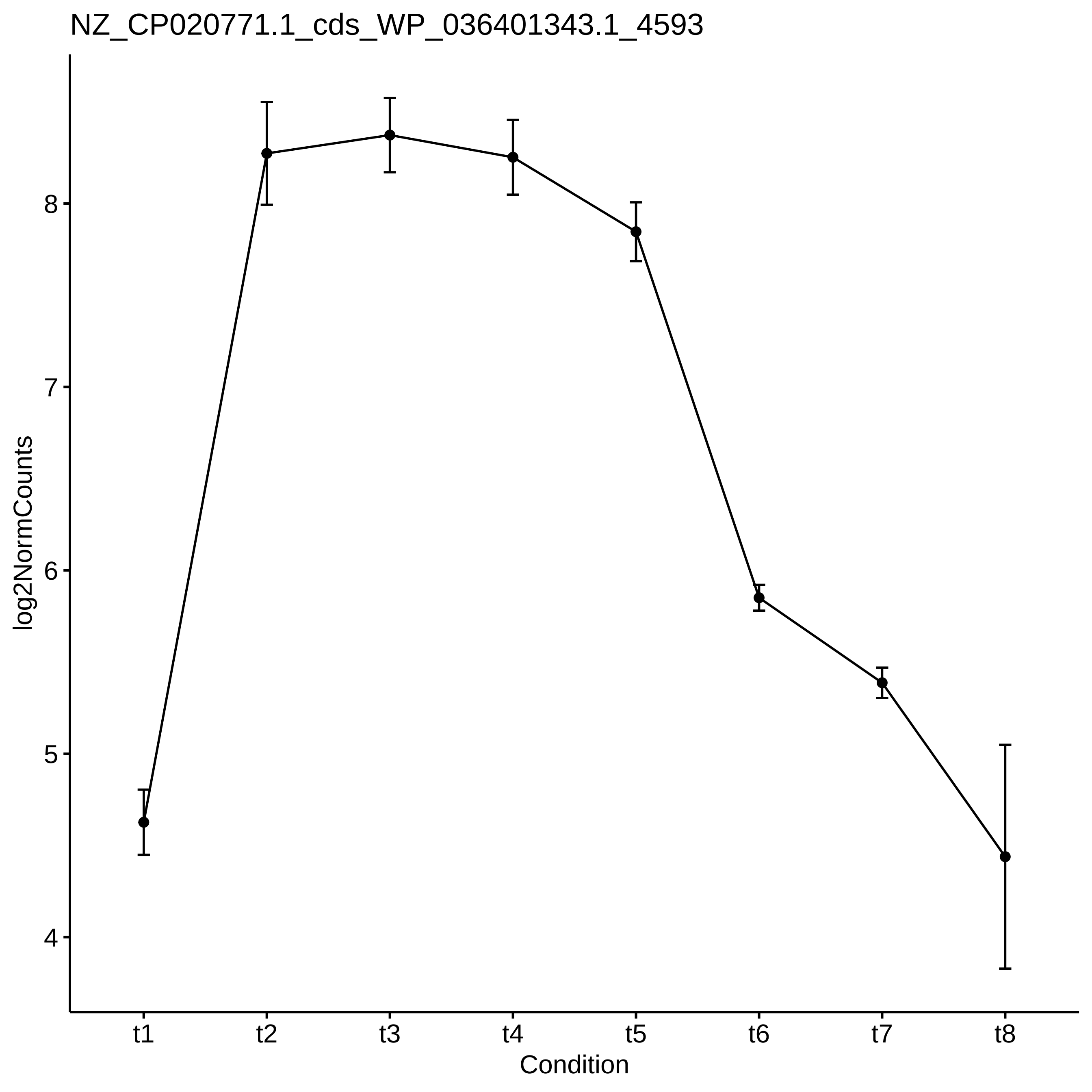
NZ_CP020771.1_cds_WP_036401343.1_4593
log2NormCounts
8
7
6
5
4
t1
t2
t3
t4
t5
t6
t7
t8
Condition

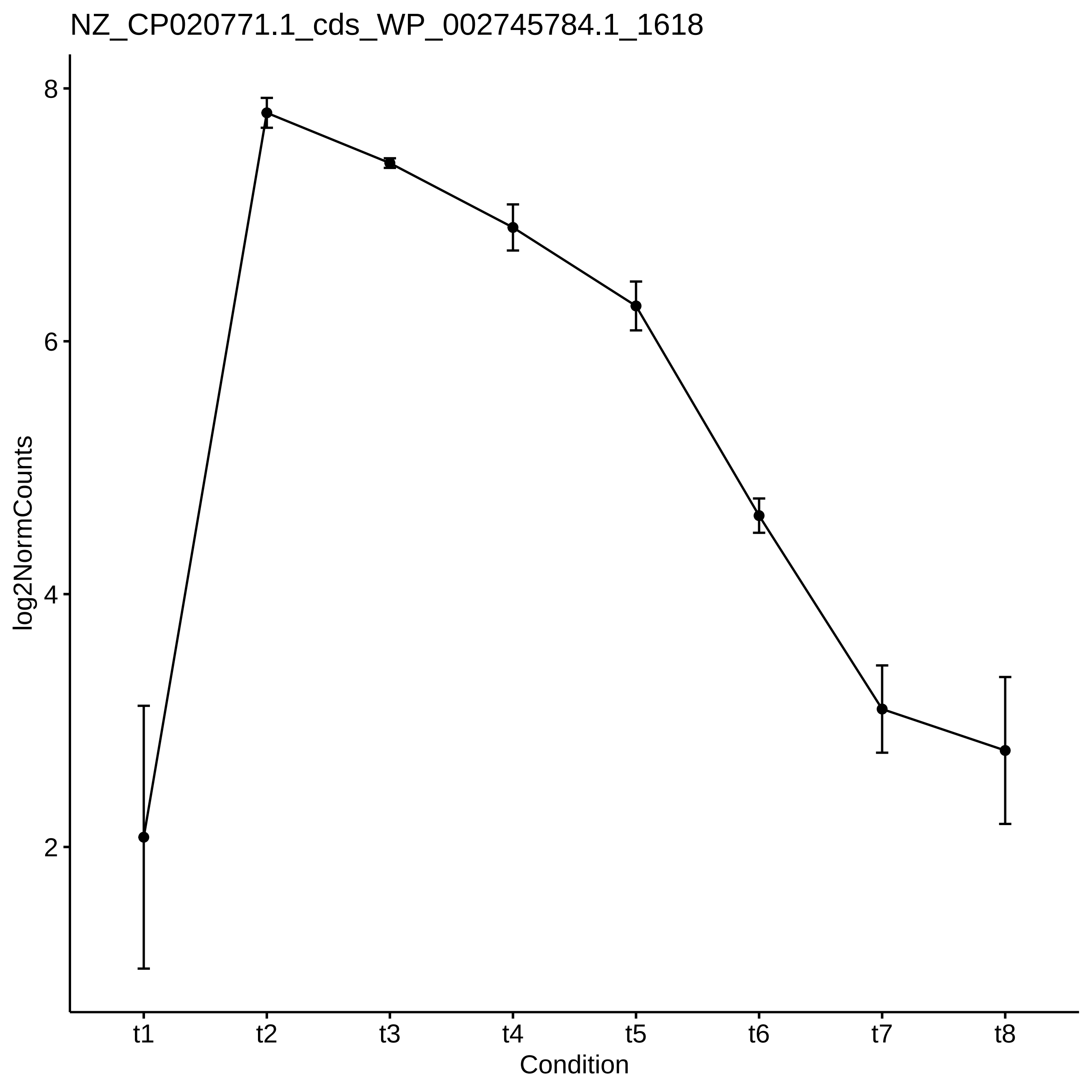

NZ_CP020771.1_cds_WP_002745784.1_1618
log2NormCounts
8
6
4
2
t1
t2
t3
t4
t5
t6
t7
t8
Condition

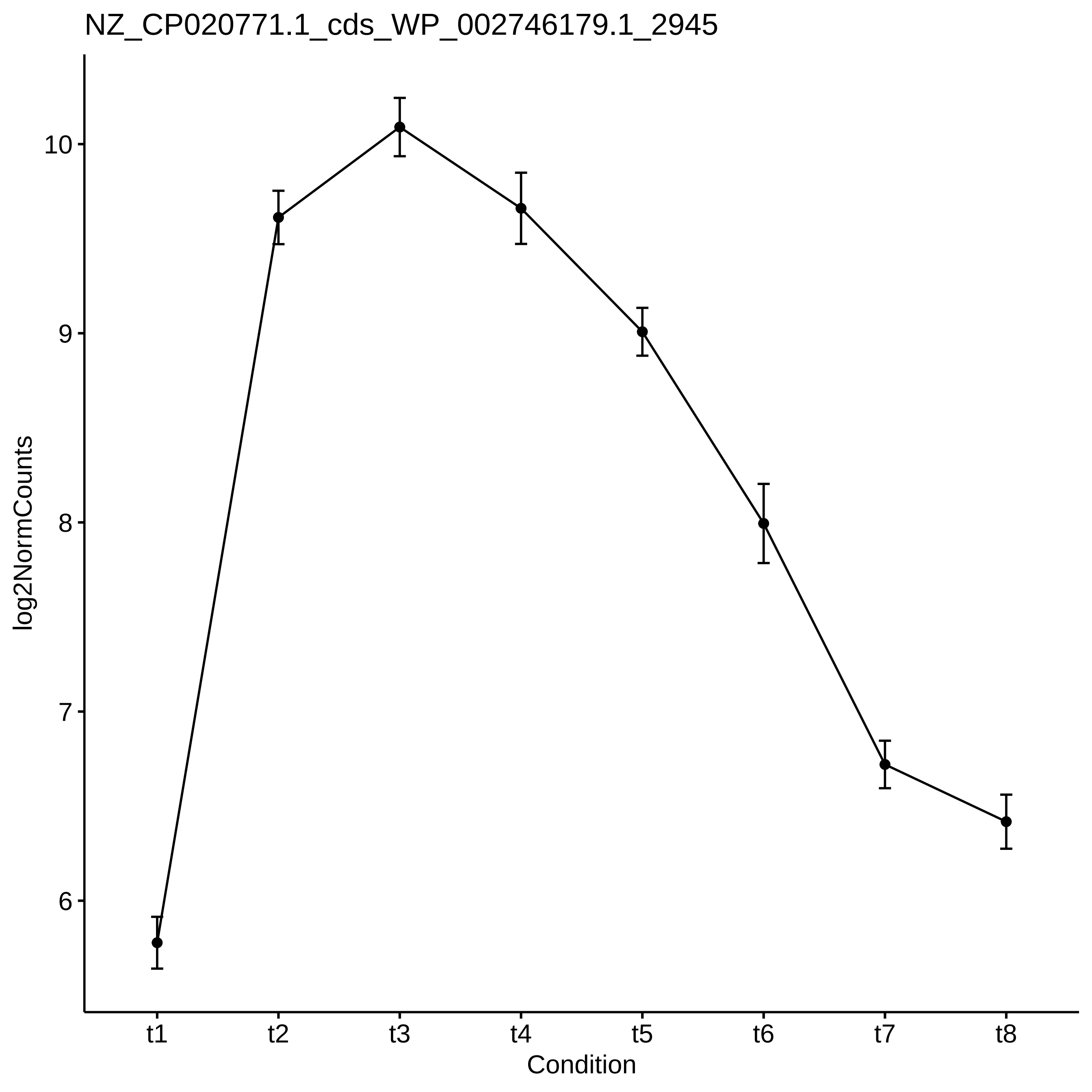

NZ_CP020771.1_cds_WP_002746179.1_2945
log2NormCounts
10
9
8
7
6
t1
t2
t3
t4
t5
t6
t7
t8
Condition

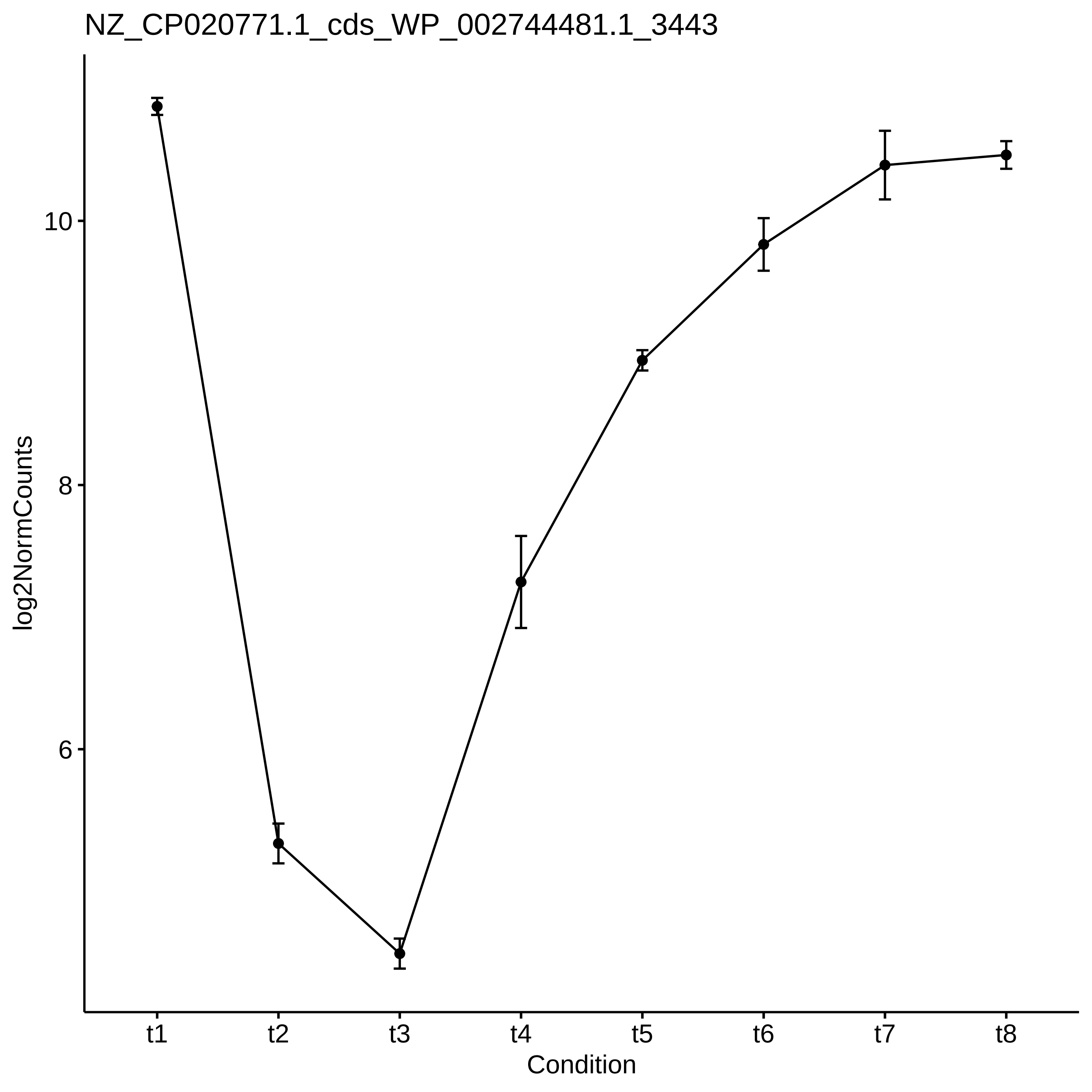
NZ_CP020771.1_cds_WP_002744481.1_3443
log2NormCounts
10
8
6
t1
t2
t3
t4
t5
t6
t7
t8
Condition

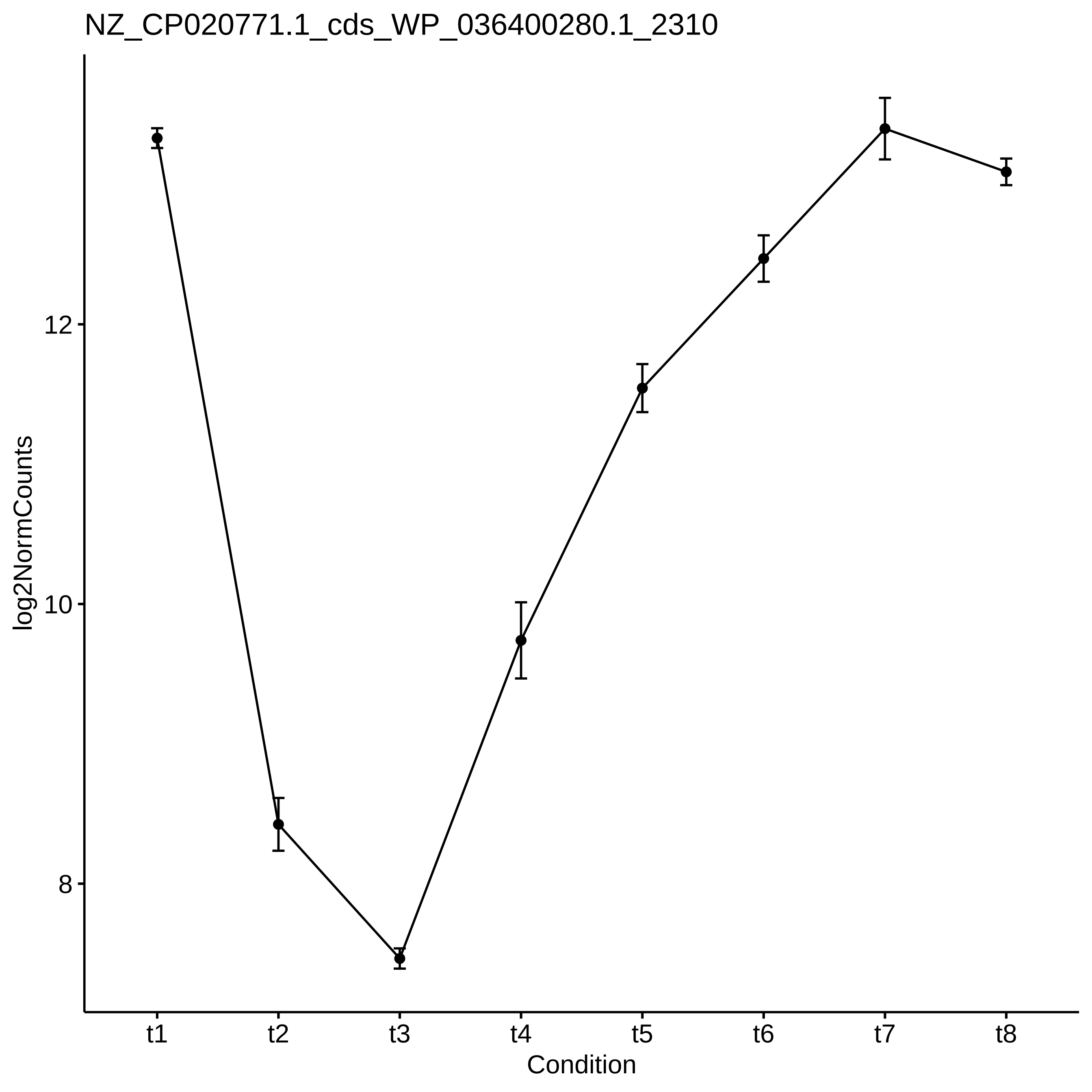
NZ_CP020771.1_cds_WP_036400280.1_2310
log2NormCounts
12
10
8
t1
t2
t3
t4
t5
t6
t7
t8
Condition

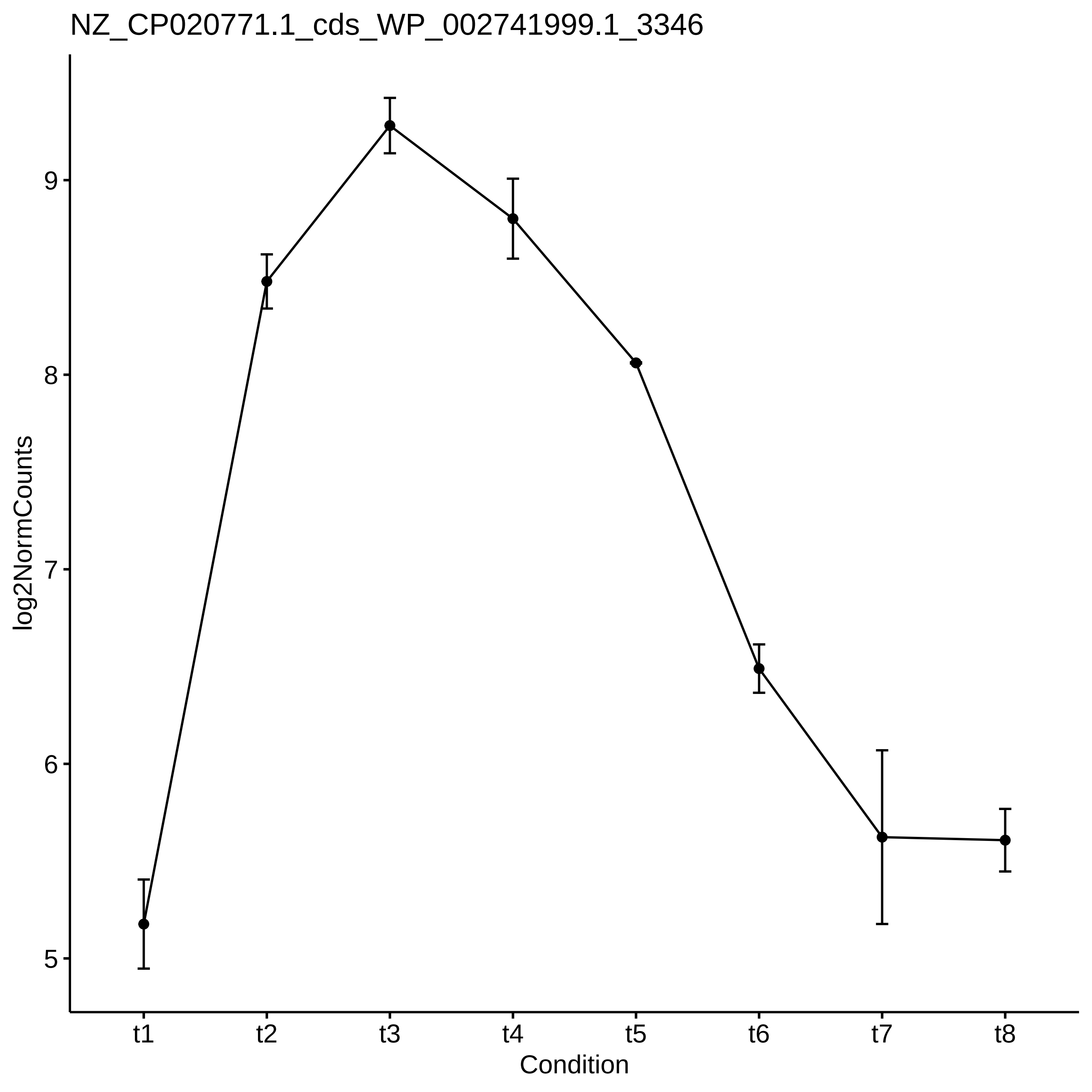
NZ_CP020771.1_cds_WP_002741999.1_3346
log2NormCounts
9
8
7
6
5
t1
t2
t3
t4
t5
t6
t7
t8
Condition

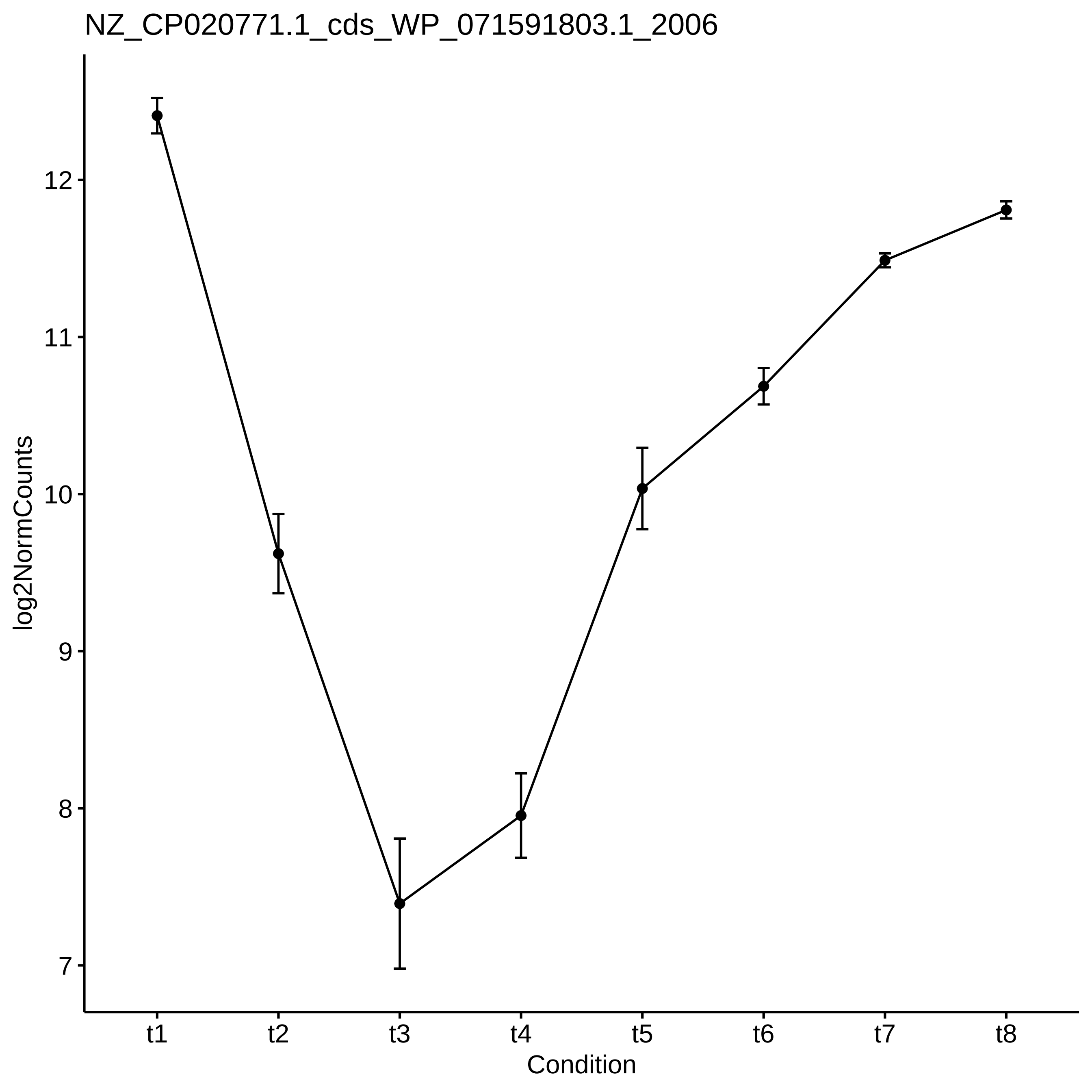

NZ_CP020771.1_cds_WP_071591803.1_2006
log2NormCounts
12
11
10
9
8
7
t1
t2
t3
t4
t5
t6
t7
t8
Condition

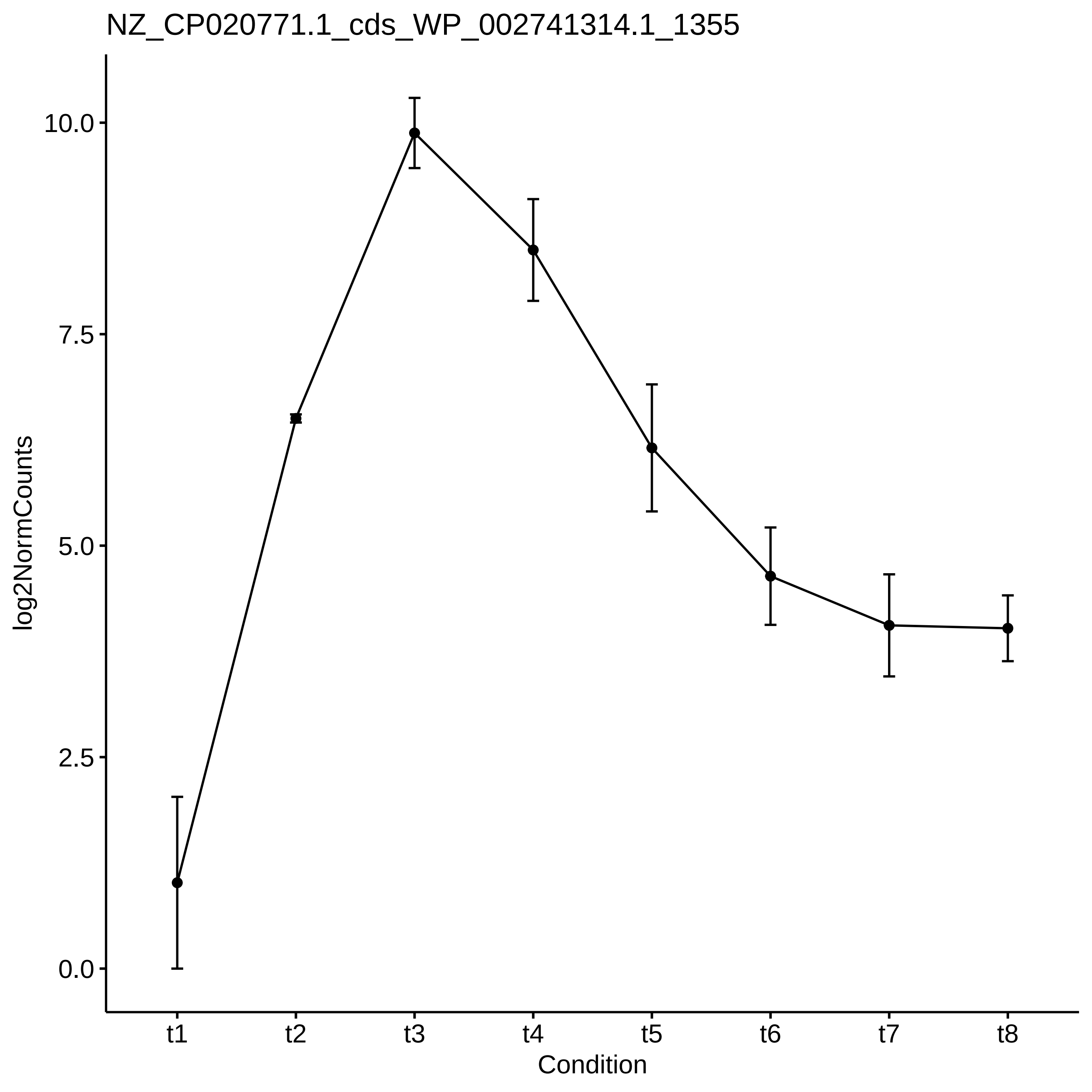
NZ_CP020771.1_cds_WP_002741314.1_1355
log2NormCounts
10.0
7.5
5.0
2.5
0.0
t1
t2
t3
t4
t5
t6
t7
t8
Condition

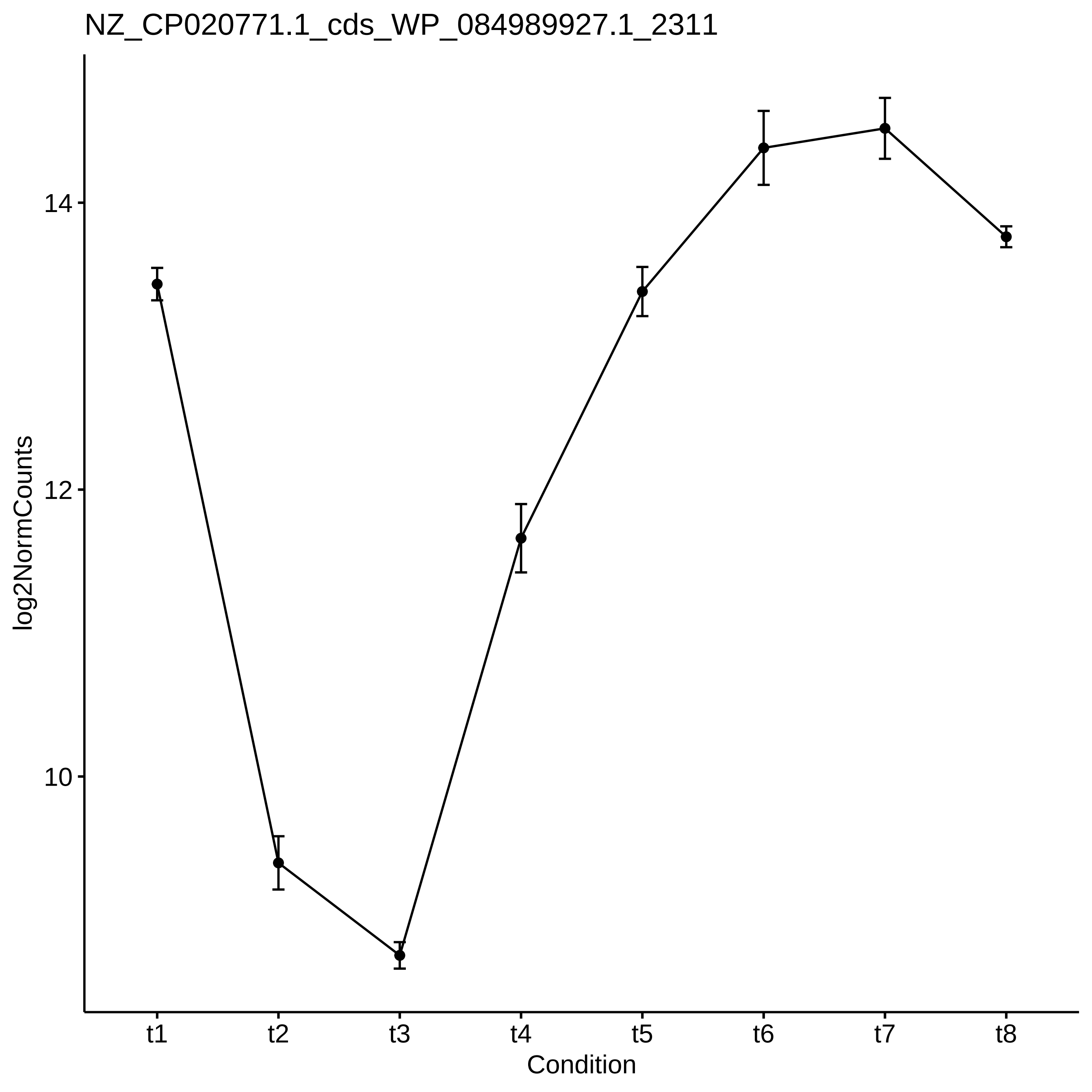
NZ_CP020771.1_cds_WP_084989927.1_2311
log2NormCounts
14
12
10
t1
t2
t3
t4
t5
t6
t7
t8
Condition

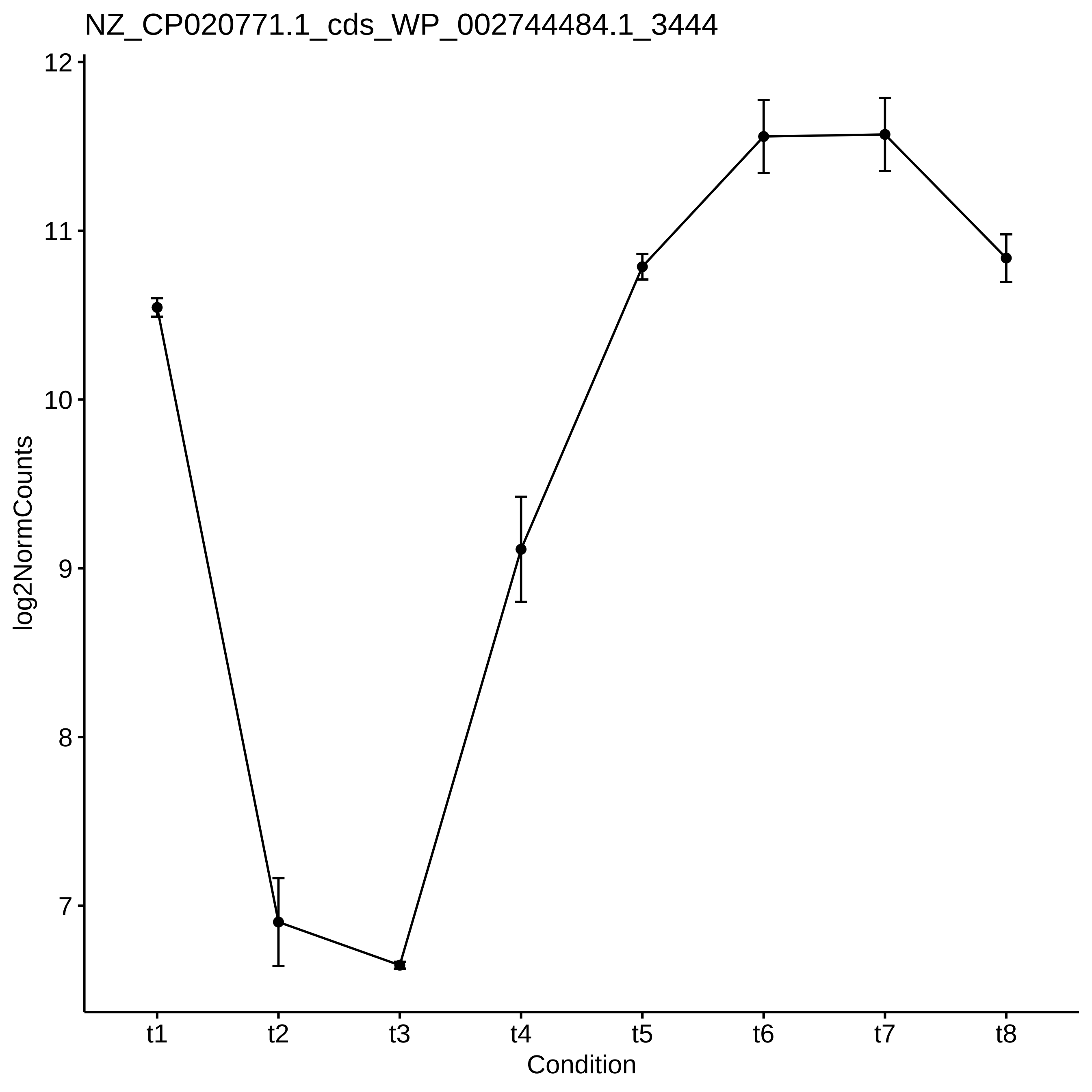
NZ_CP020771.1_cds_WP_002744484.1_3444
12
11
10
9
8
7
log2NormCounts
t1
t2
t3
t4
t5
t6
t7
t8
Condition

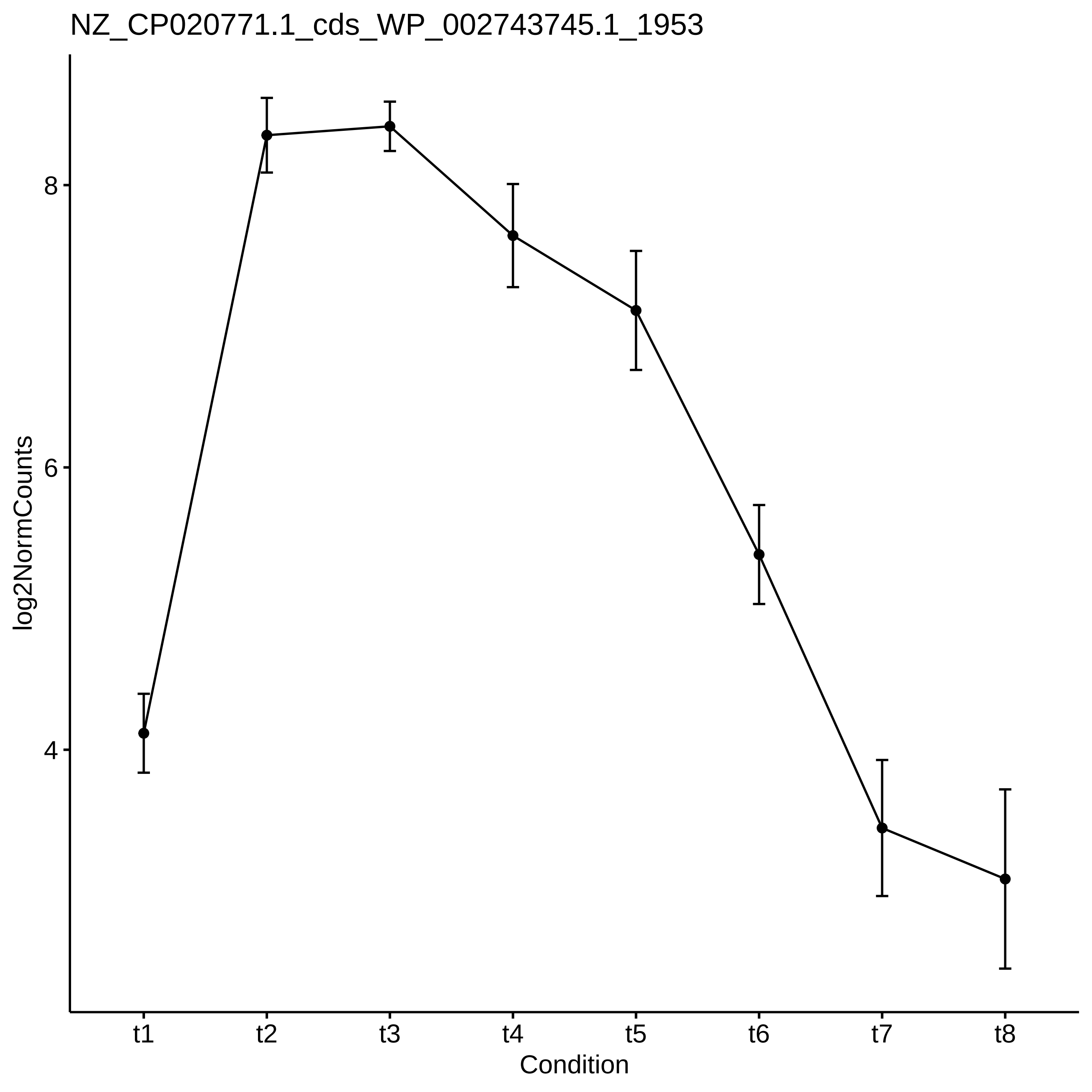
NZ_CP020771.1_cds_WP_002743745.1_1953
log2NormCounts
8
6
4
t1
t2
t3
t4
t5
t6
t7
t8
Condition

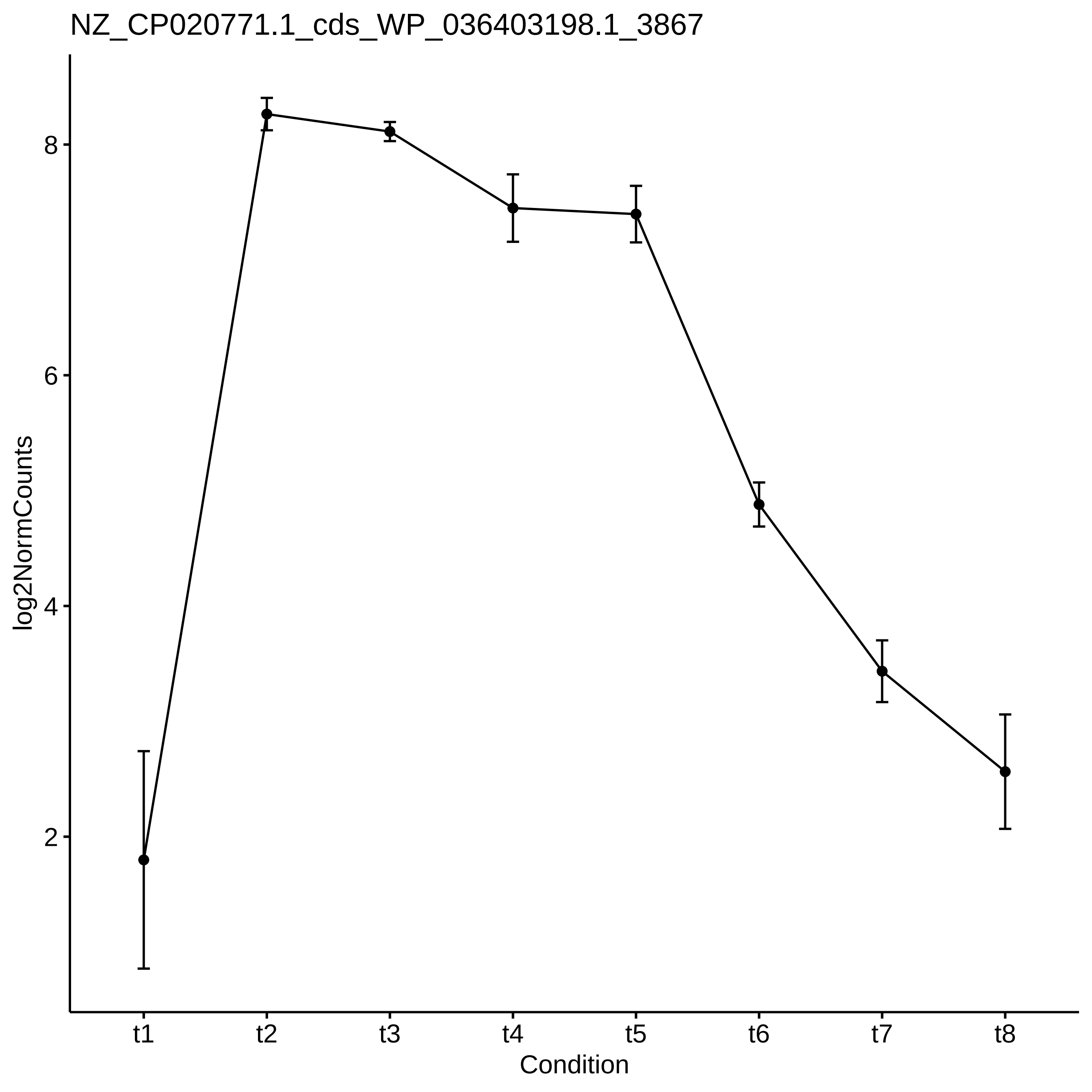

NZ_CP020771.1_cds_WP_036403198.1_3867
log2NormCounts
8
6
4
2
t1
t2
t3
t4
t5
t6
t7
t8
Condition

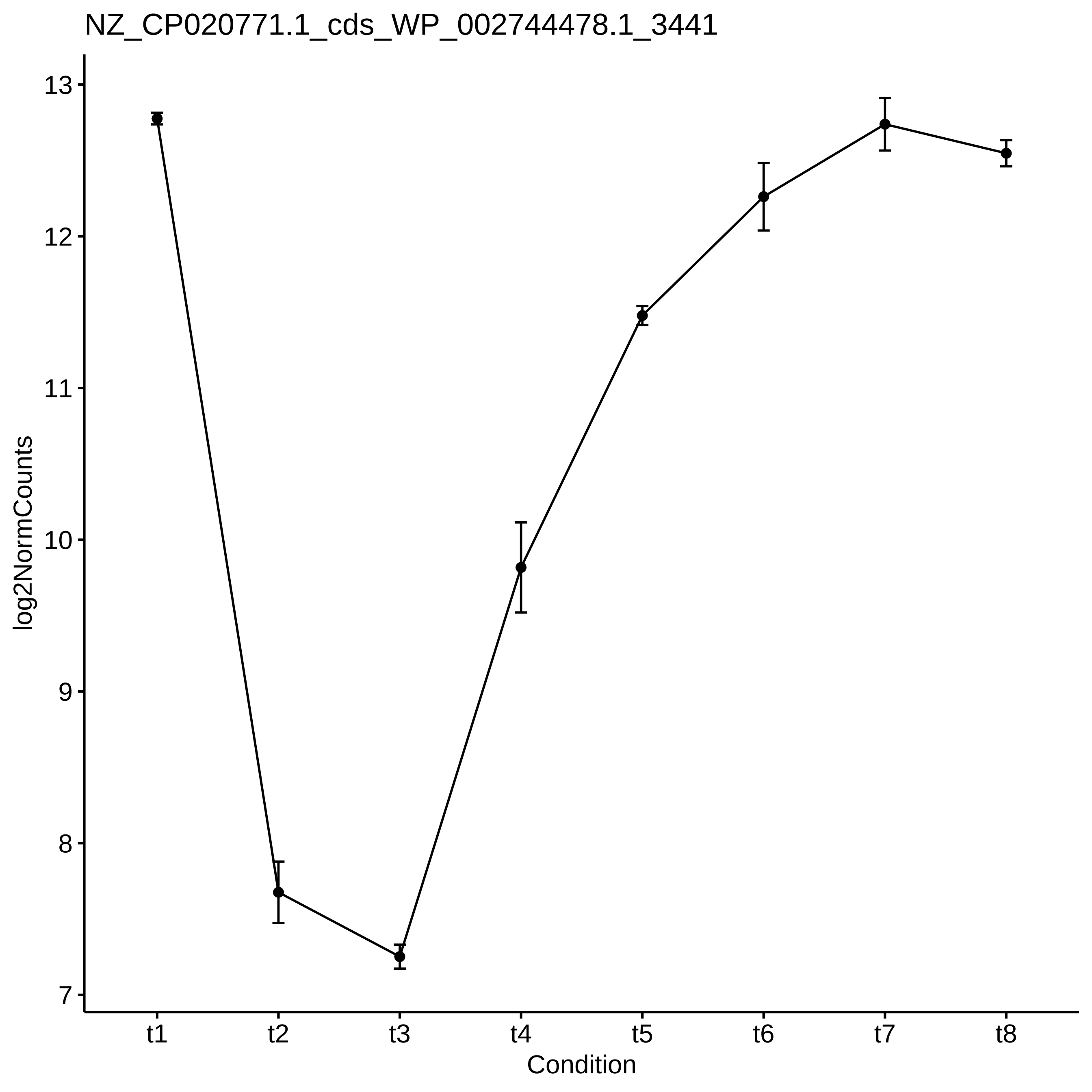

NZ_CP020771.1_cds_WP_002744478.1_3441
log2NormCounts
13
12
11
10
9
8
7
t1
t2
t3
t4
t5
t6
t7
t8
Condition

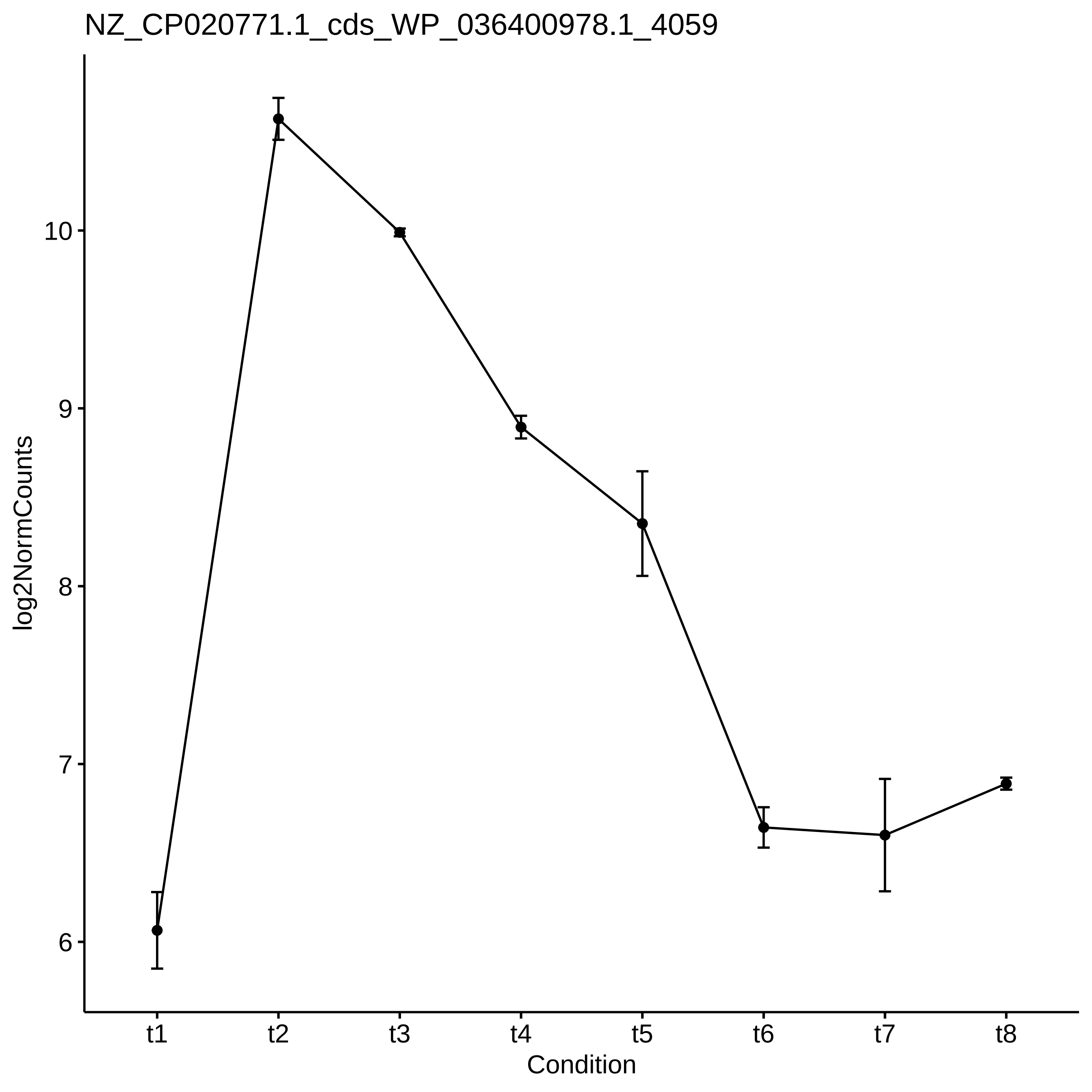

NZ_CP020771.1_cds_WP_036400978.1_4059
log2NormCounts
10
9
8
7
6
t1
t2
t3
t4
t5
t6
t7
t8
Condition

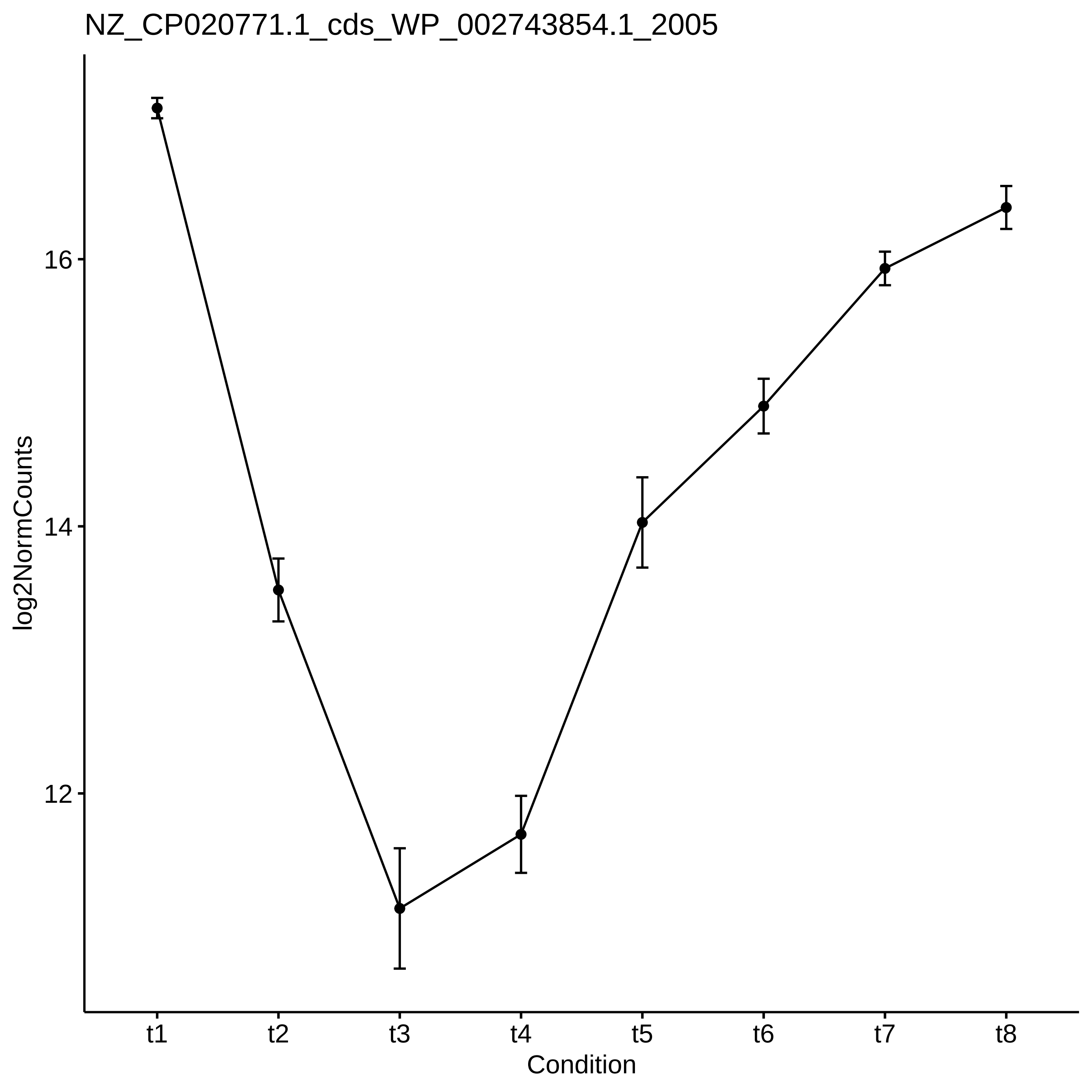

NZ_CP020771.1_cds_WP_002743854.1_2005
log2NormCounts
16
14
12
t1
t2
t3
t4
t5
t6
t7
t8
Condition

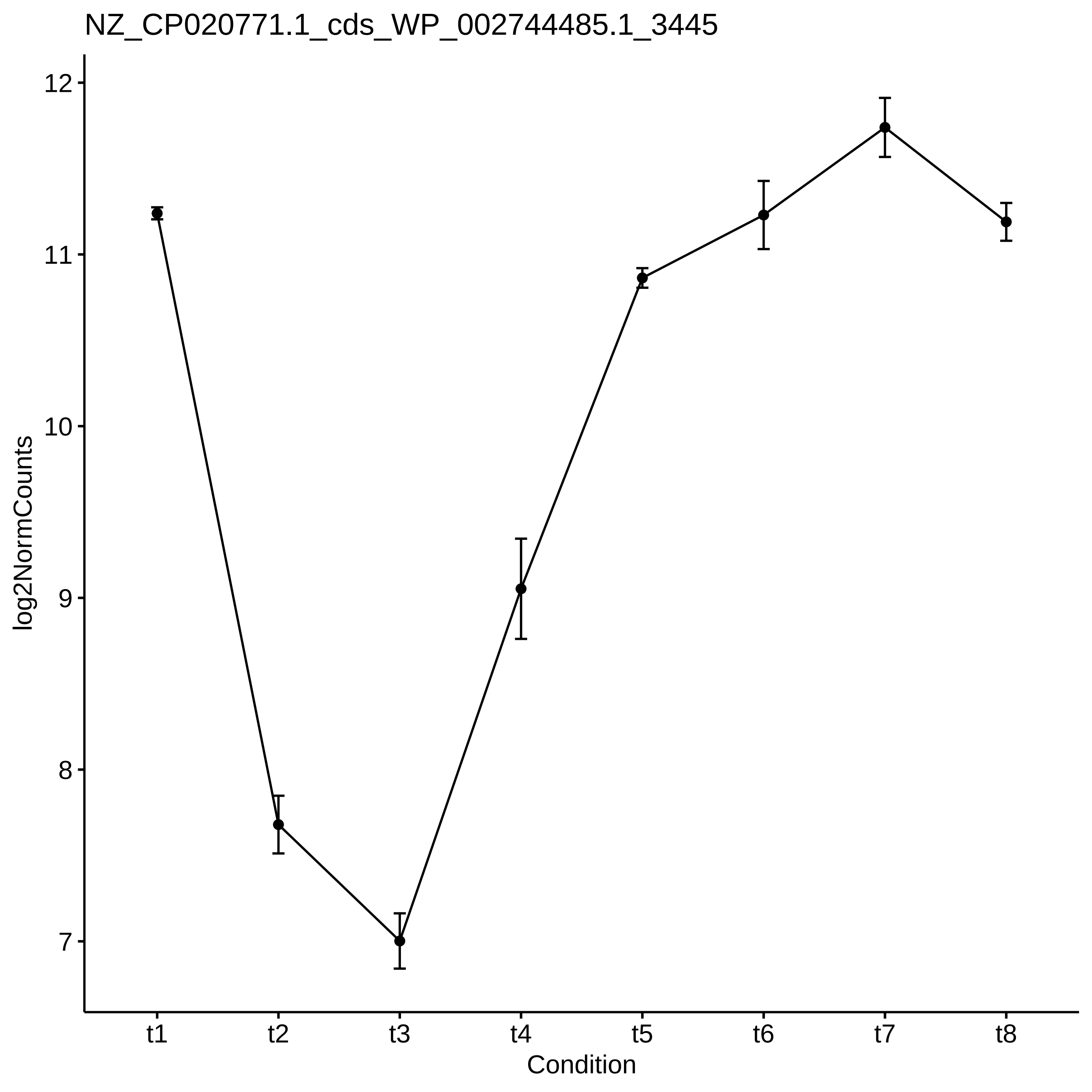

NZ_CP020771.1_cds_WP_002744485.1_3445
log2NormCounts
12
11
10
9
8
7
t1
t2
t3
t4
t5
t6
t7
t8
Condition

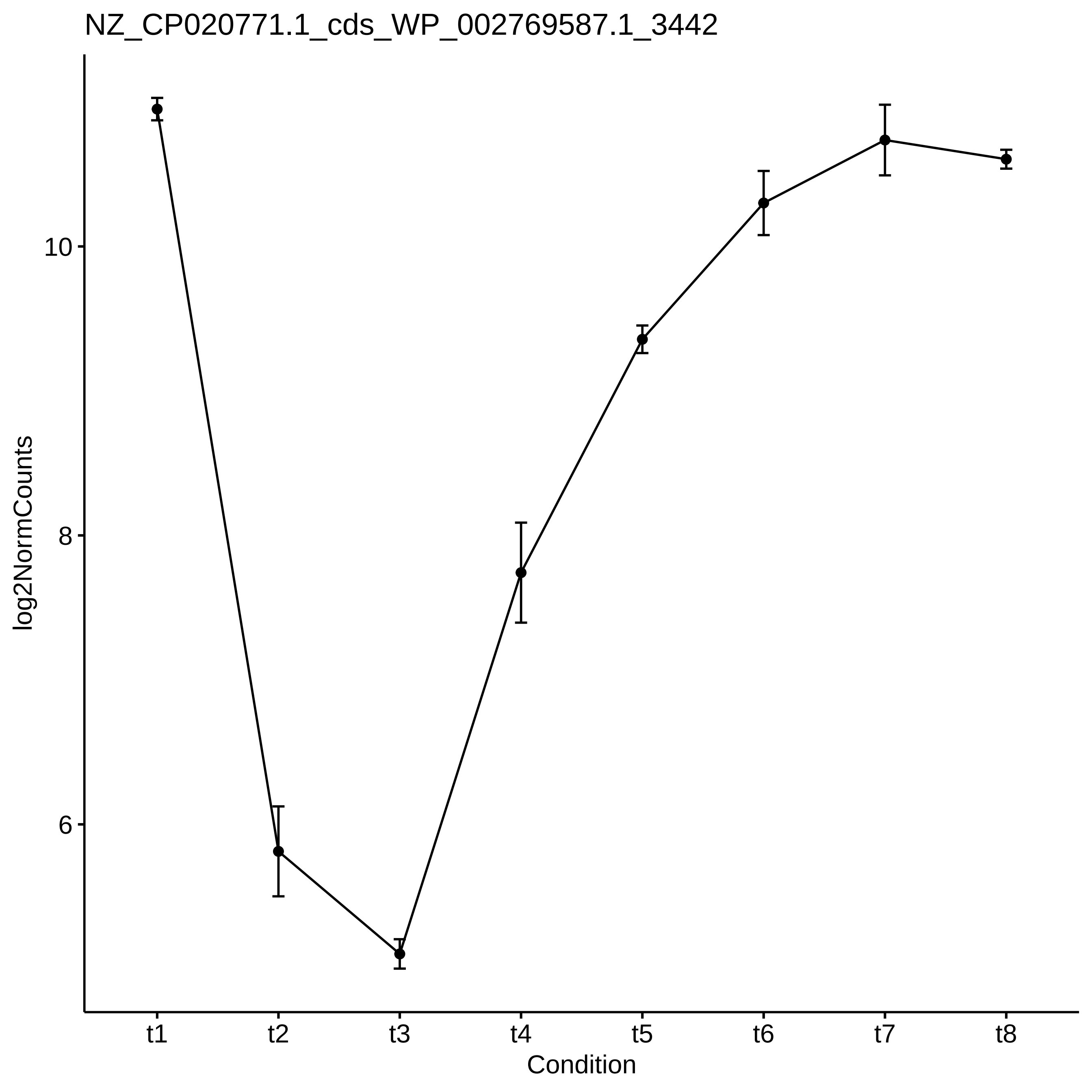

NZ_CP020771.1_cds_WP_002769587.1_3442
log2NormCounts
10
8
6
t1
t2
t3
t4
t5
t6
t7
t8
Condition

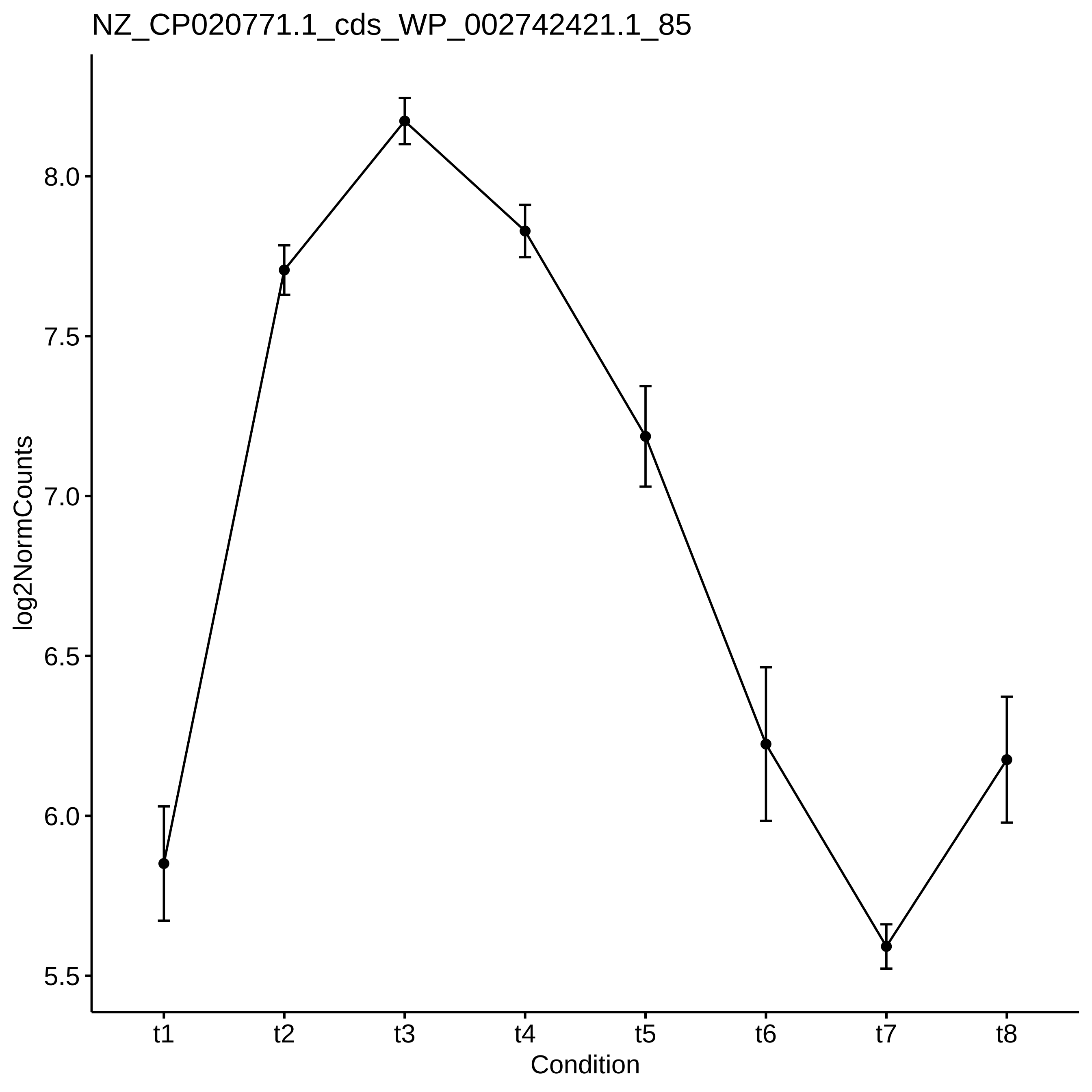

NZ_CP020771.1_cds_WP_002742421.1_85
log2NormCounts
8.0
7.5
7.0
6.5
6.0
5.5
t1
t2
t3
t4
t5
t6
t7
t8
Condition

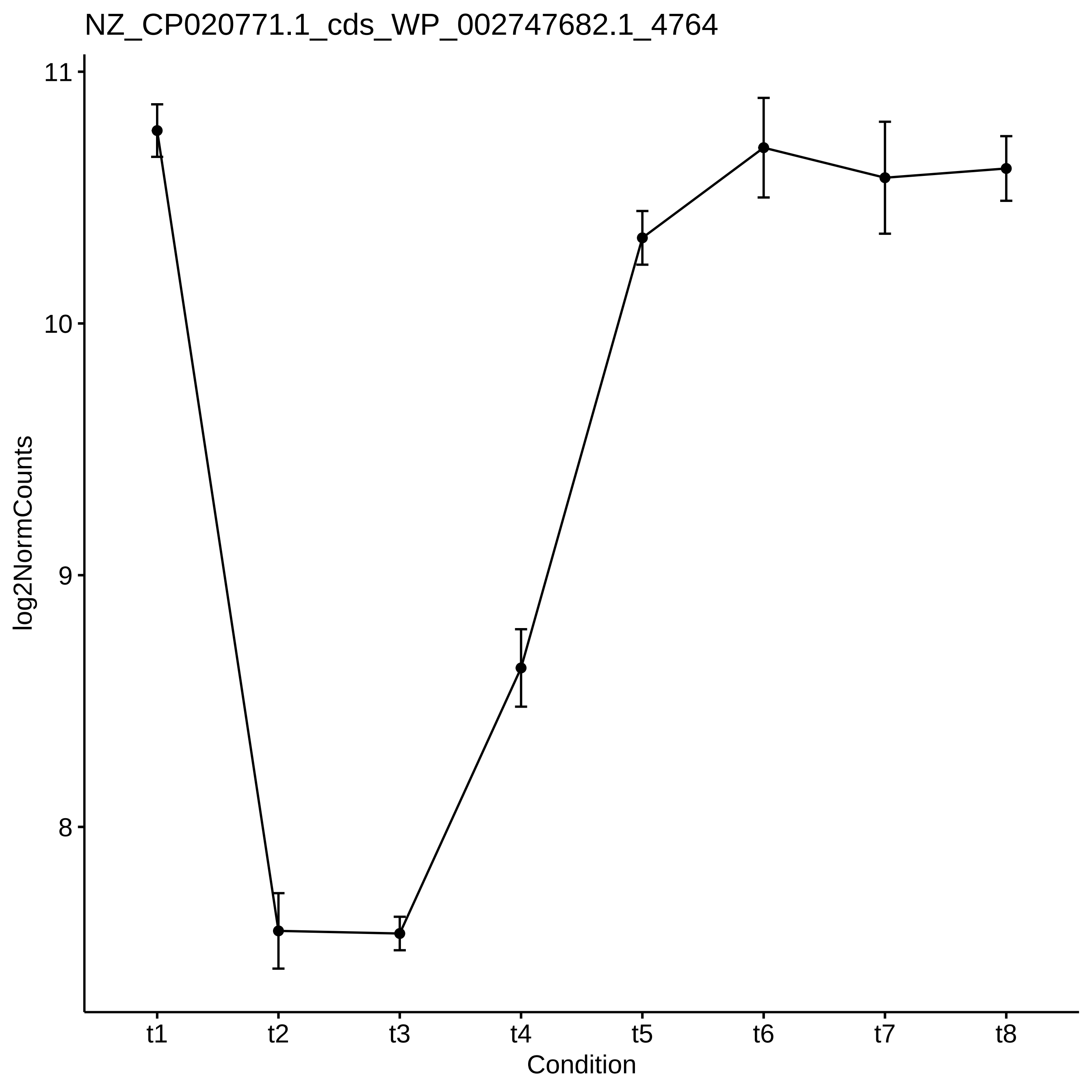

NZ_CP020771.1_cds_WP_002747682.1_4764
log2NormCounts
11
10
9
8
t1
t2
t3
t4
t5
t6
t7
t8
Condition

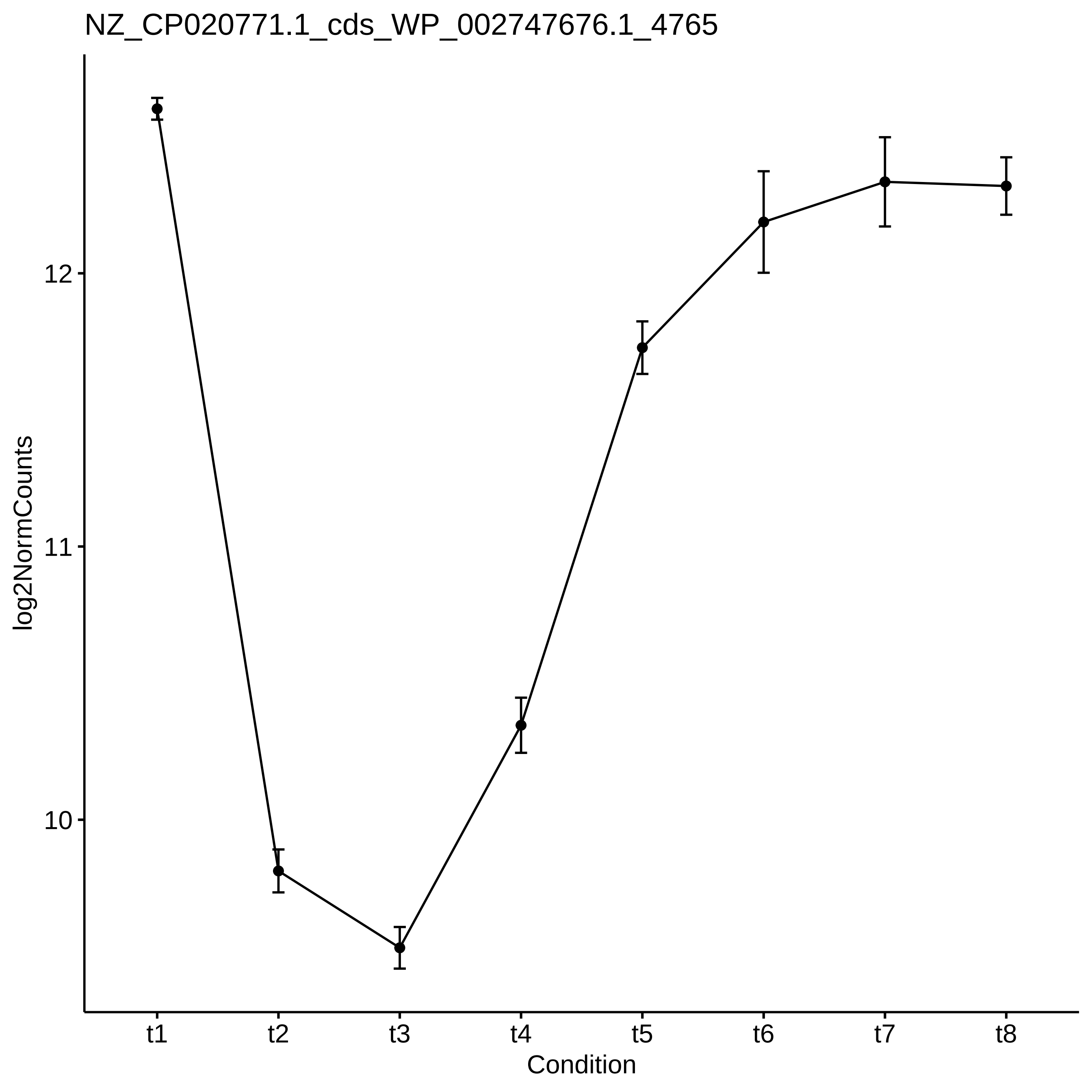

NZ_CP020771.1_cds_WP_002747676.1_4765
log2NormCounts
12
11
10
t1
t2
t3
t4
t5
t6
t7
t8
Condition

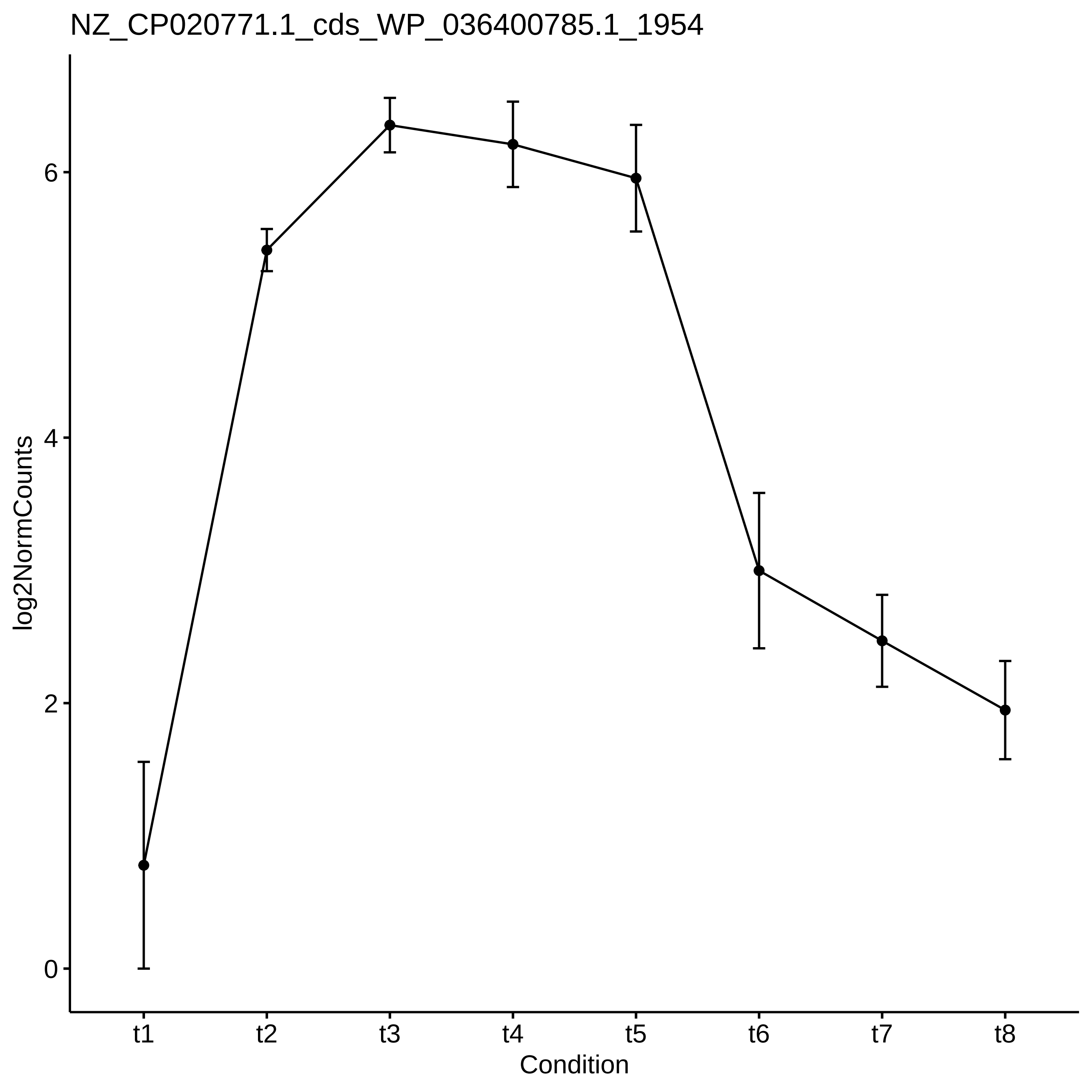

NZ_CP020771.1_cds_WP_036400785.1_1954
log2NormCounts
6
4
2
0
t1
t2
t3
t4
t5
t6
t7
t8
Condition

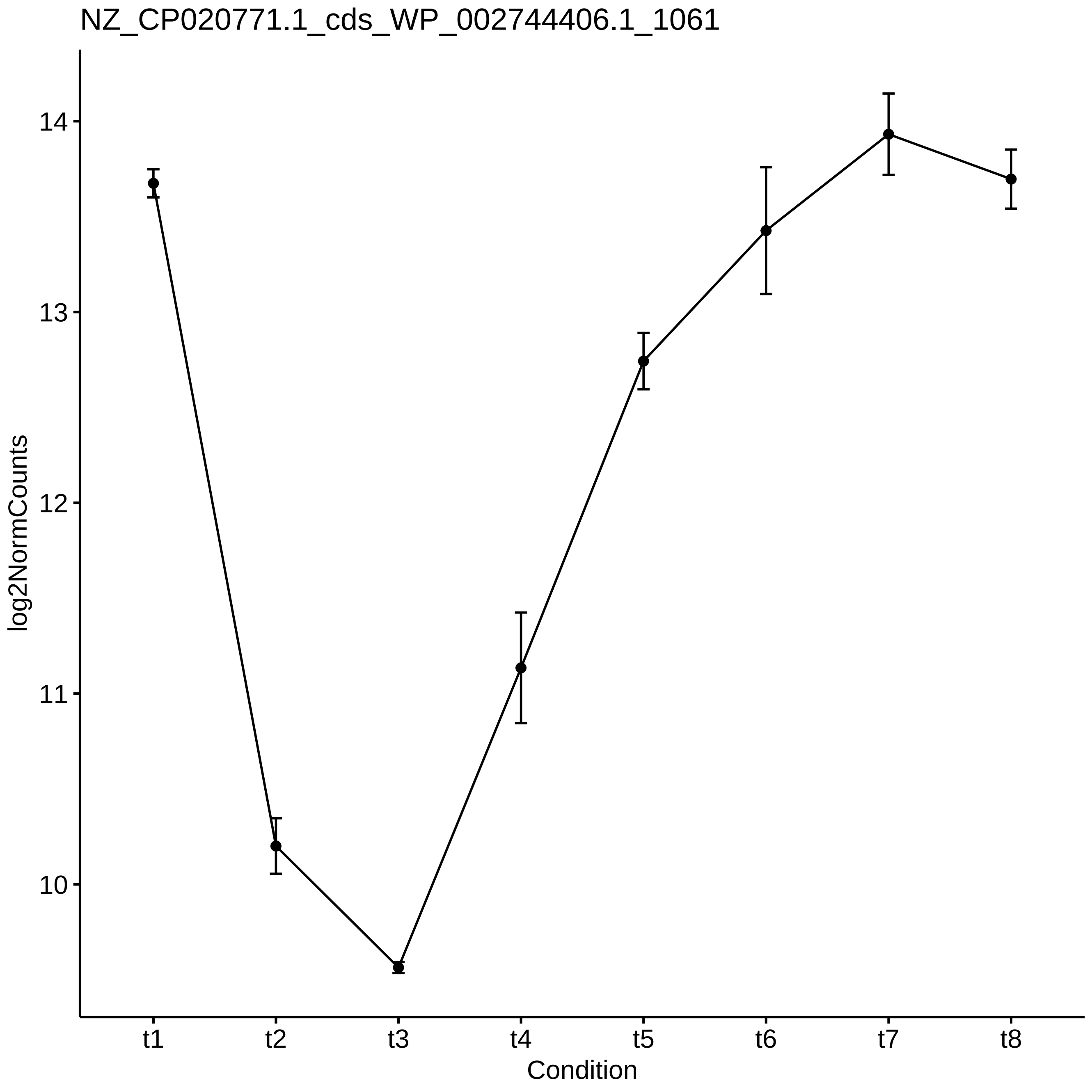
NZ_CP020771.1_cds_WP_002744406.1_1061
log2NormCounts
14
13
12
11
10
t1
t2
t3
t4
t5
t6
t7
t8
Condition

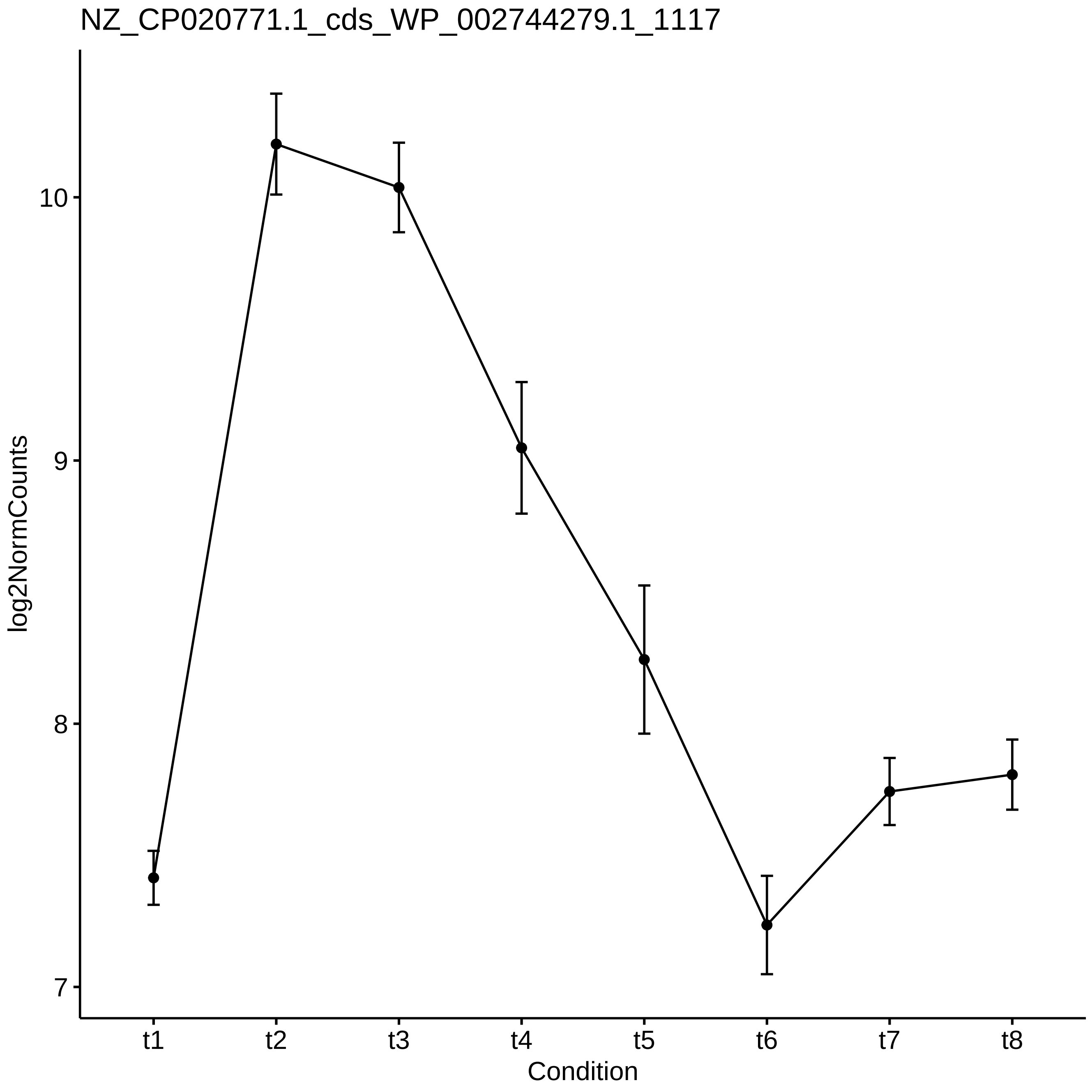

NZ_CP020771.1_cds_WP_002744279.1_1117
log2NormCounts
10
9
8
7
t1
t2
t3
t4
t5
t6
t7
t8
Condition

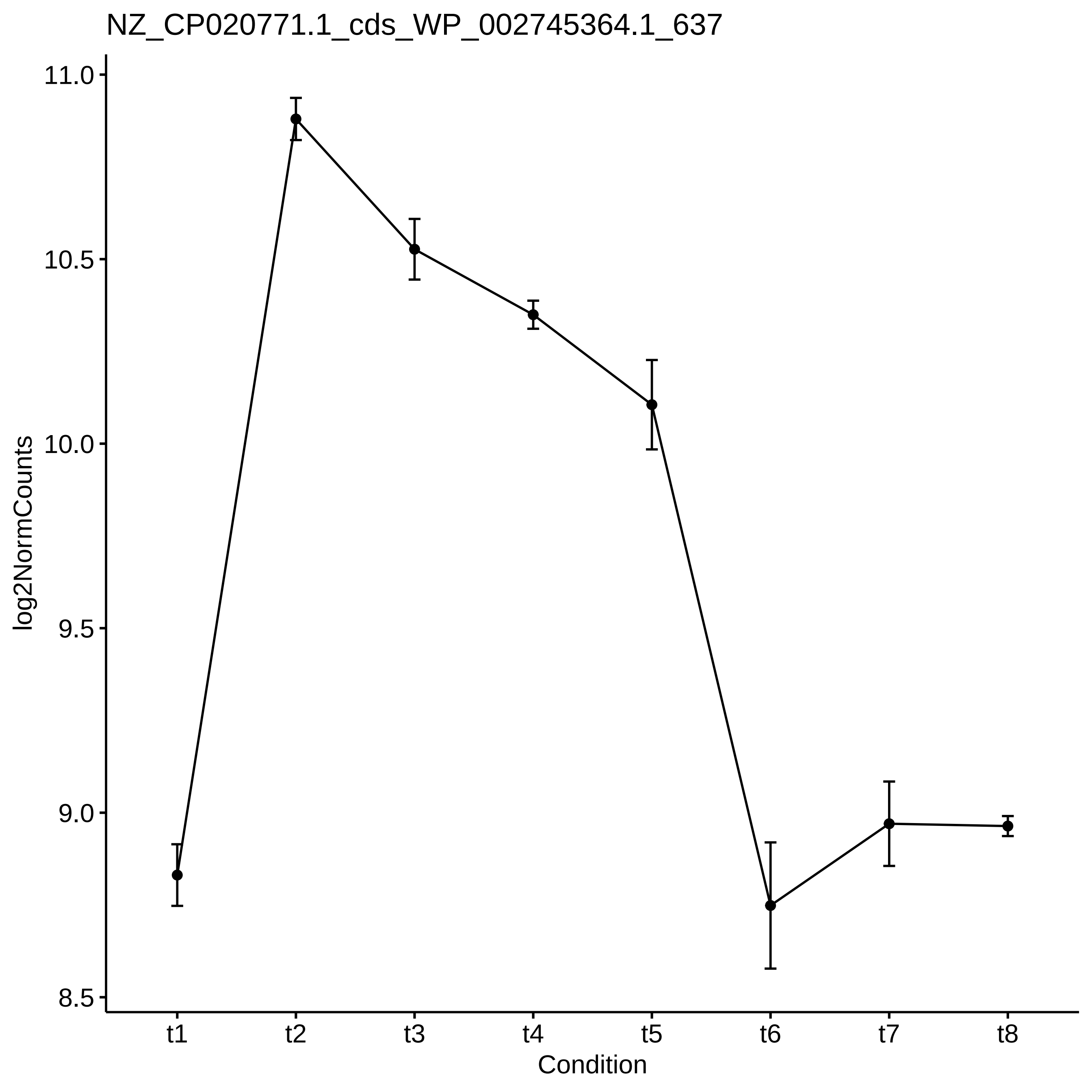

NZ_CP020771.1_cds_WP_002745364.1_637
11.0
10.5
10.0
9.5
9.0
8.5
log2NormCounts
t1
t2
t3
t4
t5
t6
t7
t8
Condition

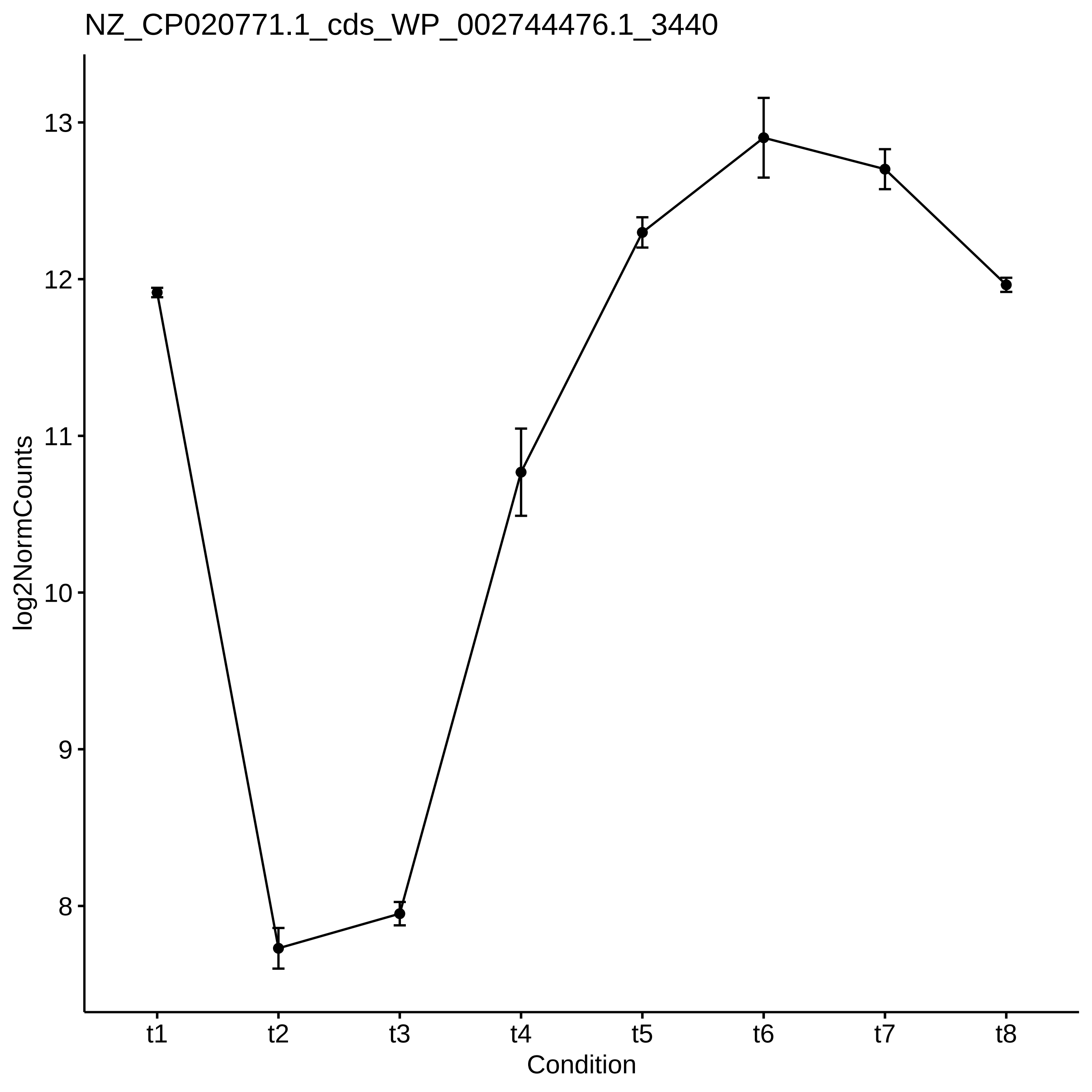
NZ_CP020771.1_cds_WP_002744476.1_3440
log2NormCounts
13
12
11
10
9
8
t1
t2
t3
t4
t5
t6
t7
t8
Condition

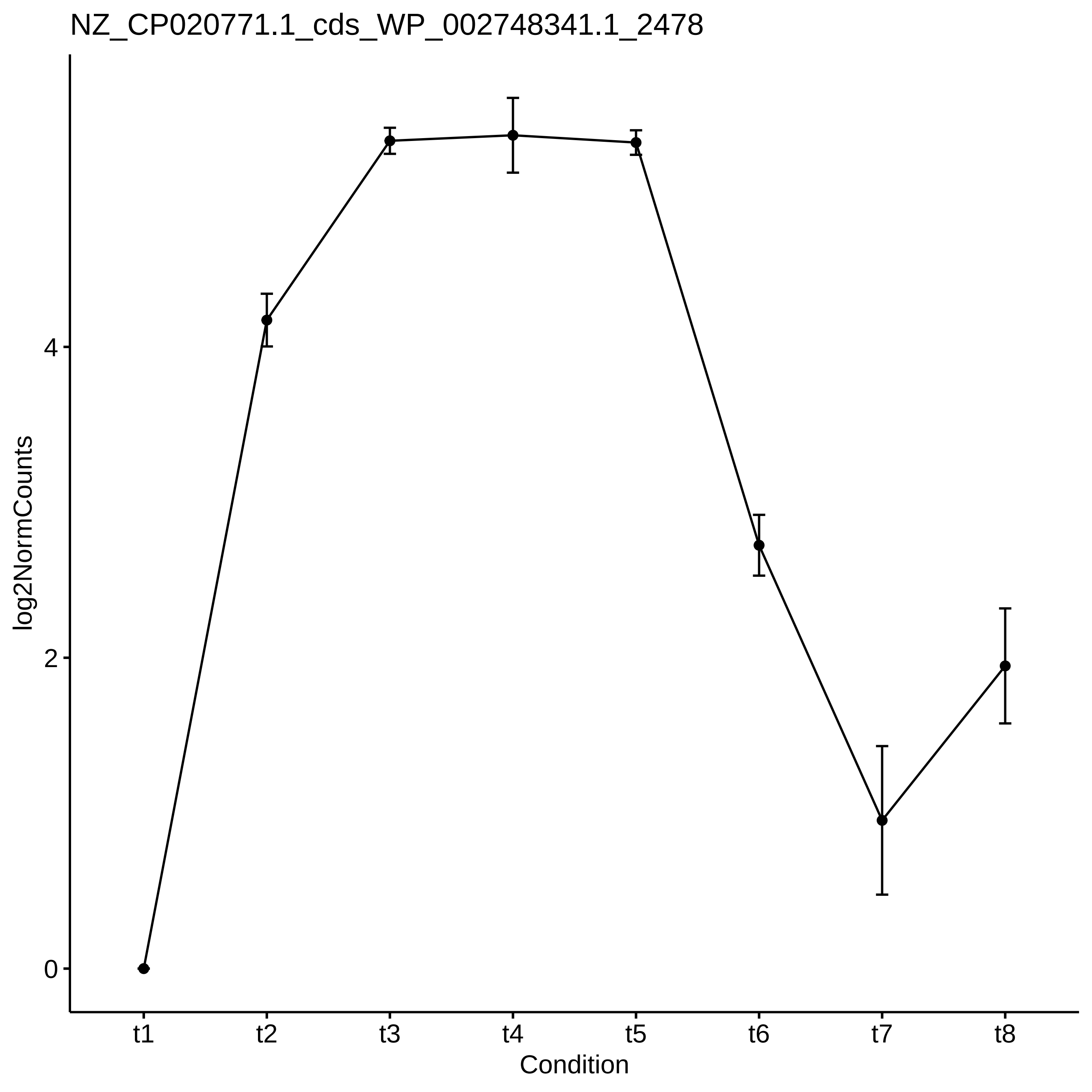

NZ_CP020771.1_cds_WP_002748341.1_2478
log2NormCounts
0
2
4
t1
t2
t3
t4
t5
t6
t7
t8
Condition

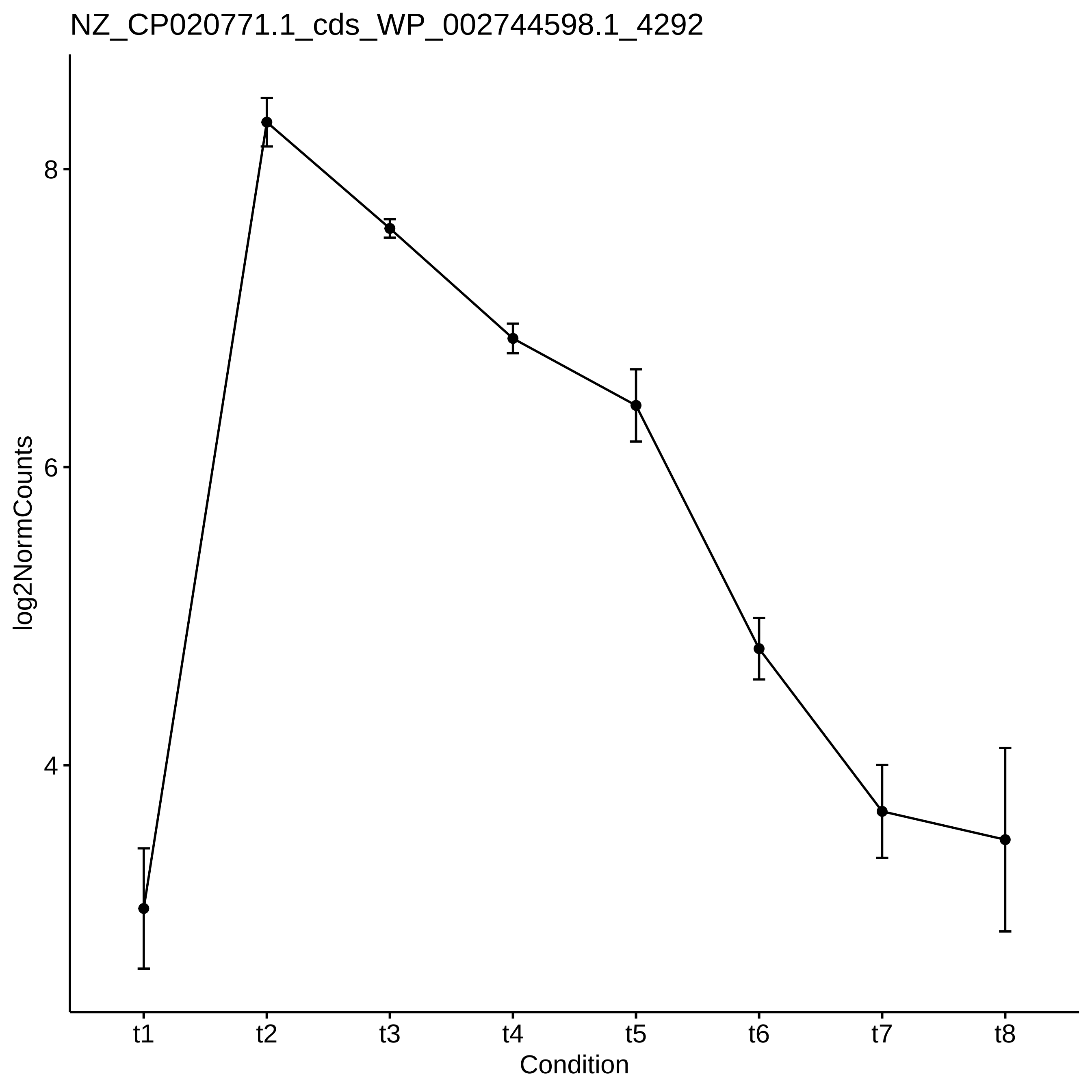

NZ_CP020771.1_cds_WP_002744598.1_4292
log2NormCounts
8
6
4
t1
t2
t3
t4
t5
t6
t7
t8
Condition

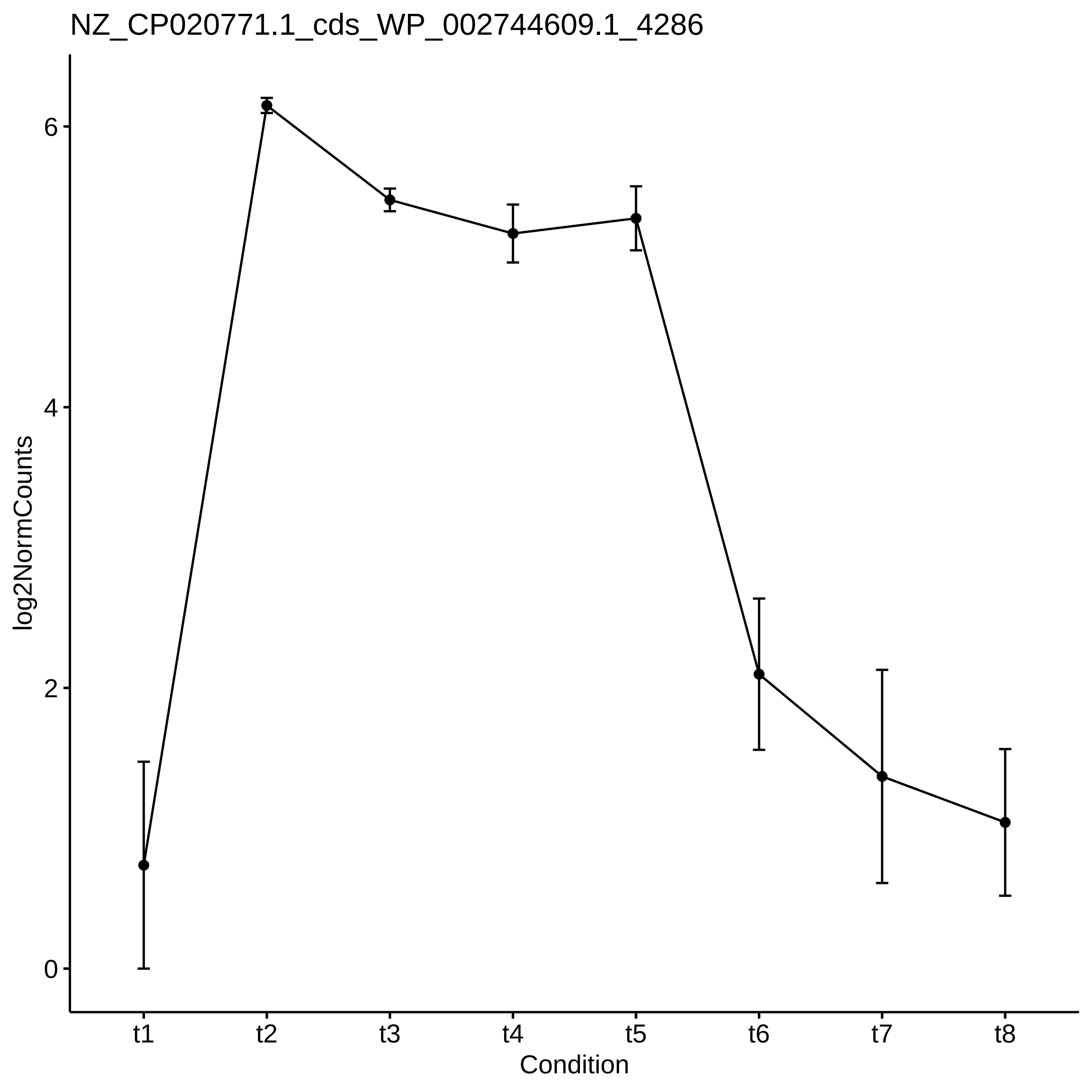

NZ_CP020771.1_cds_WP_002744609.1_4286
log2NormCounts
0
2
4
6
t1
t2
t3
t4
t5
t6
t7
t8
Condition

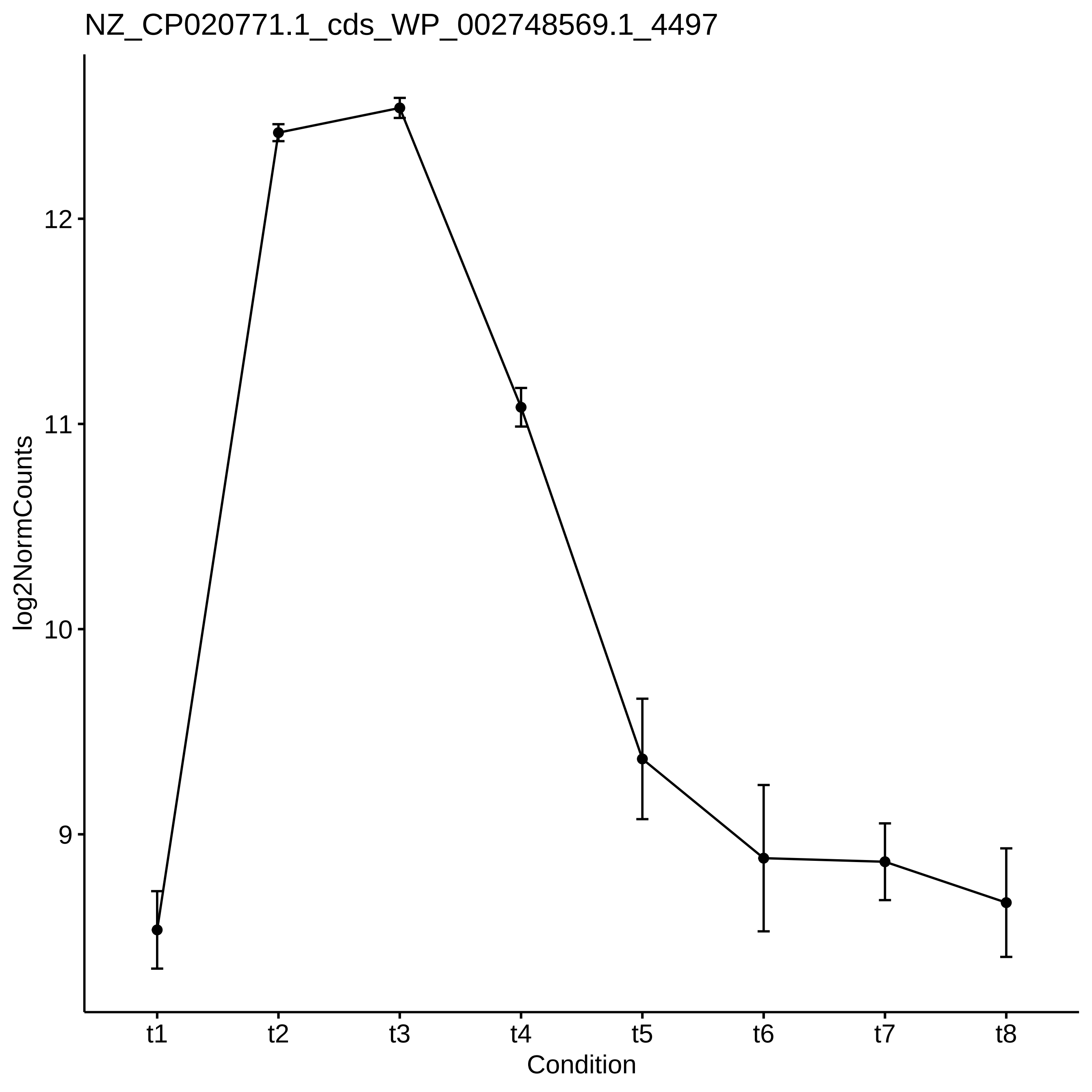

NZ_CP020771.1_cds_WP_002748569.1_4497
log2NormCounts
12
11
10
9
t1
t2
t3
t4
t5
t6
t7
t8
Condition

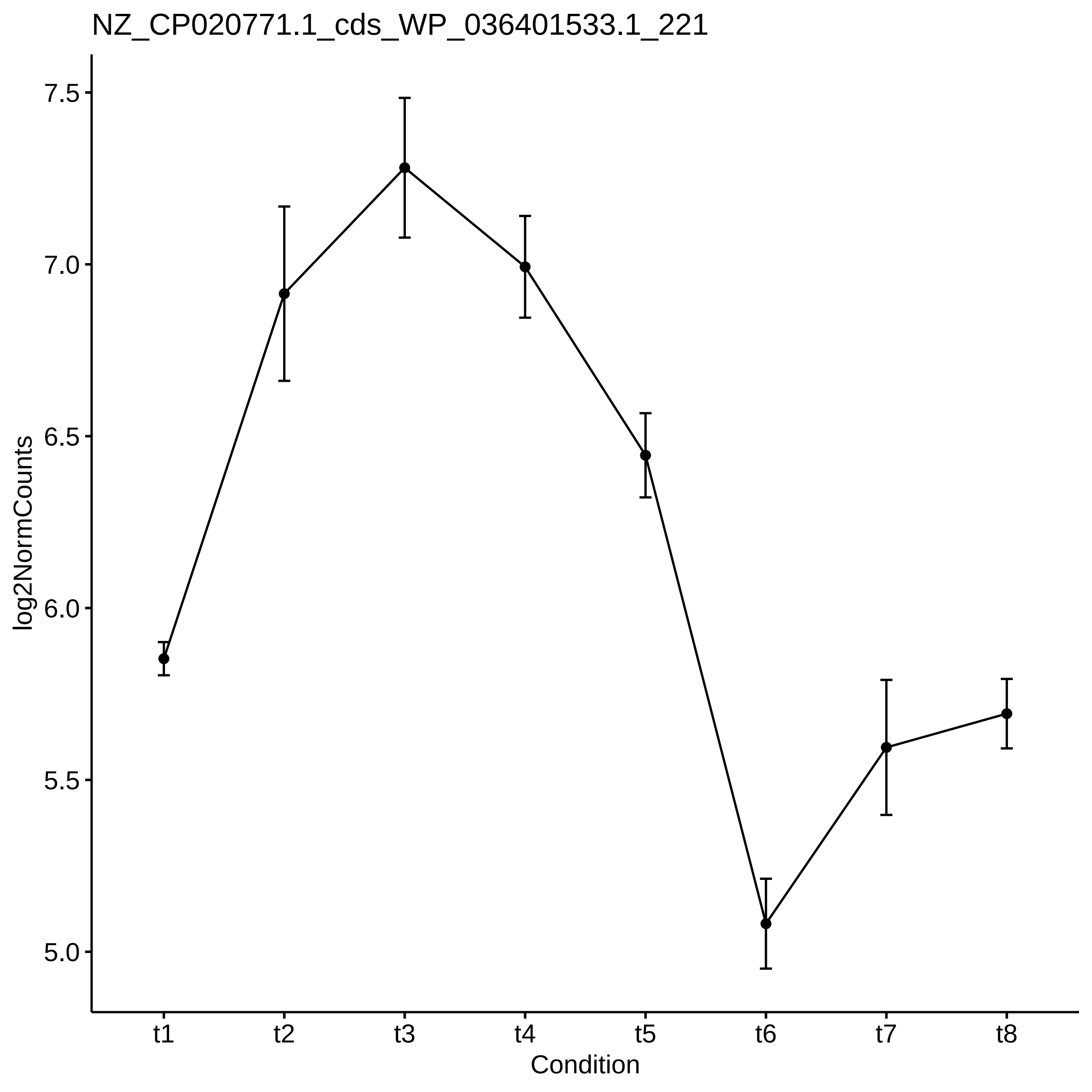
NZ_CP020771.1_cds_WP_036401533.1_221
log2NormCounts
7.5
7.0
6.5
6.0
5.5
5.0
t1
t2
t3
t4
t5
t6
t7
t8
Condition

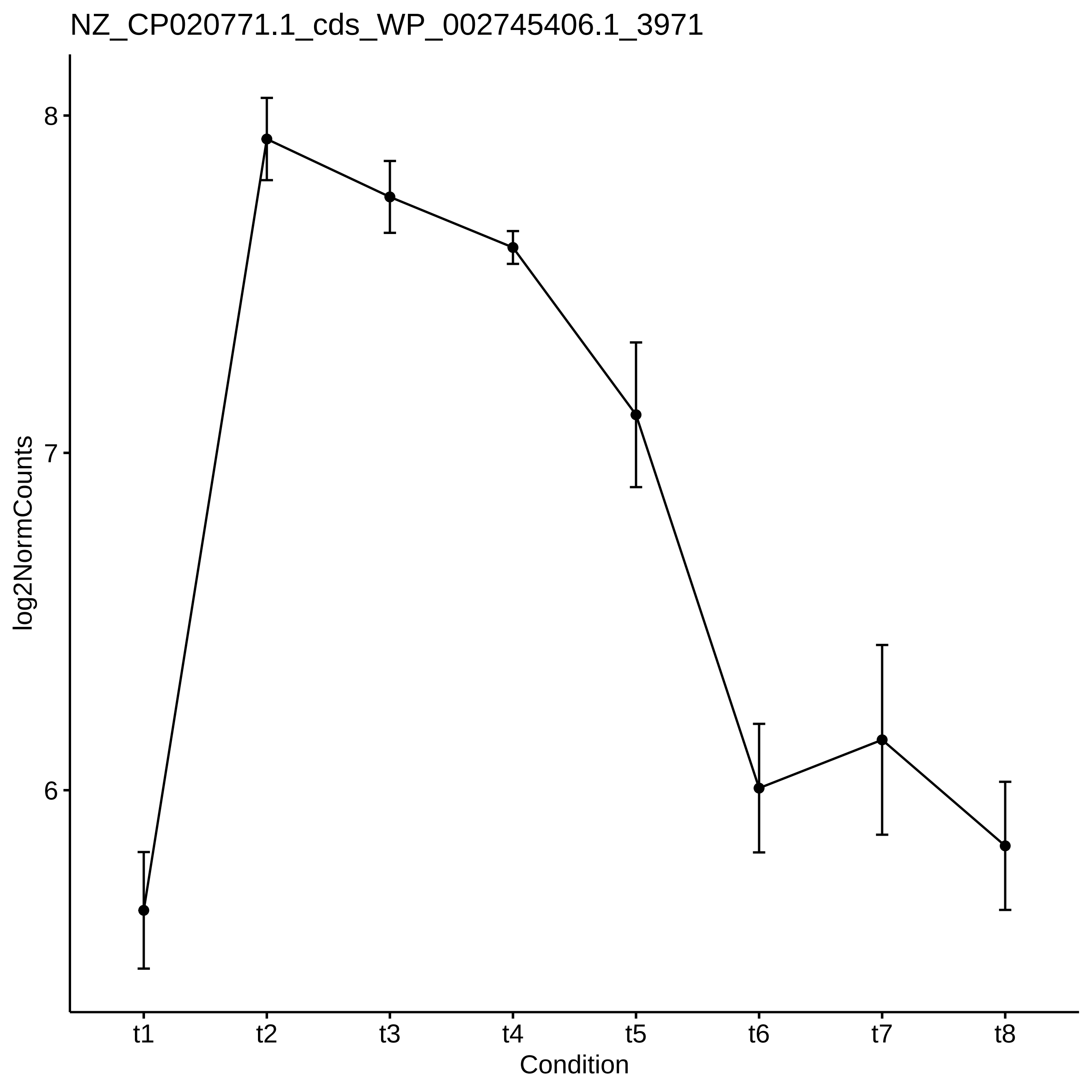
NZ_CP020771.1_cds_WP_002745406.1_3971
log2NormCounts
8
7
6
t1
t2
t3
t4
t5
t6
t7
t8
Condition

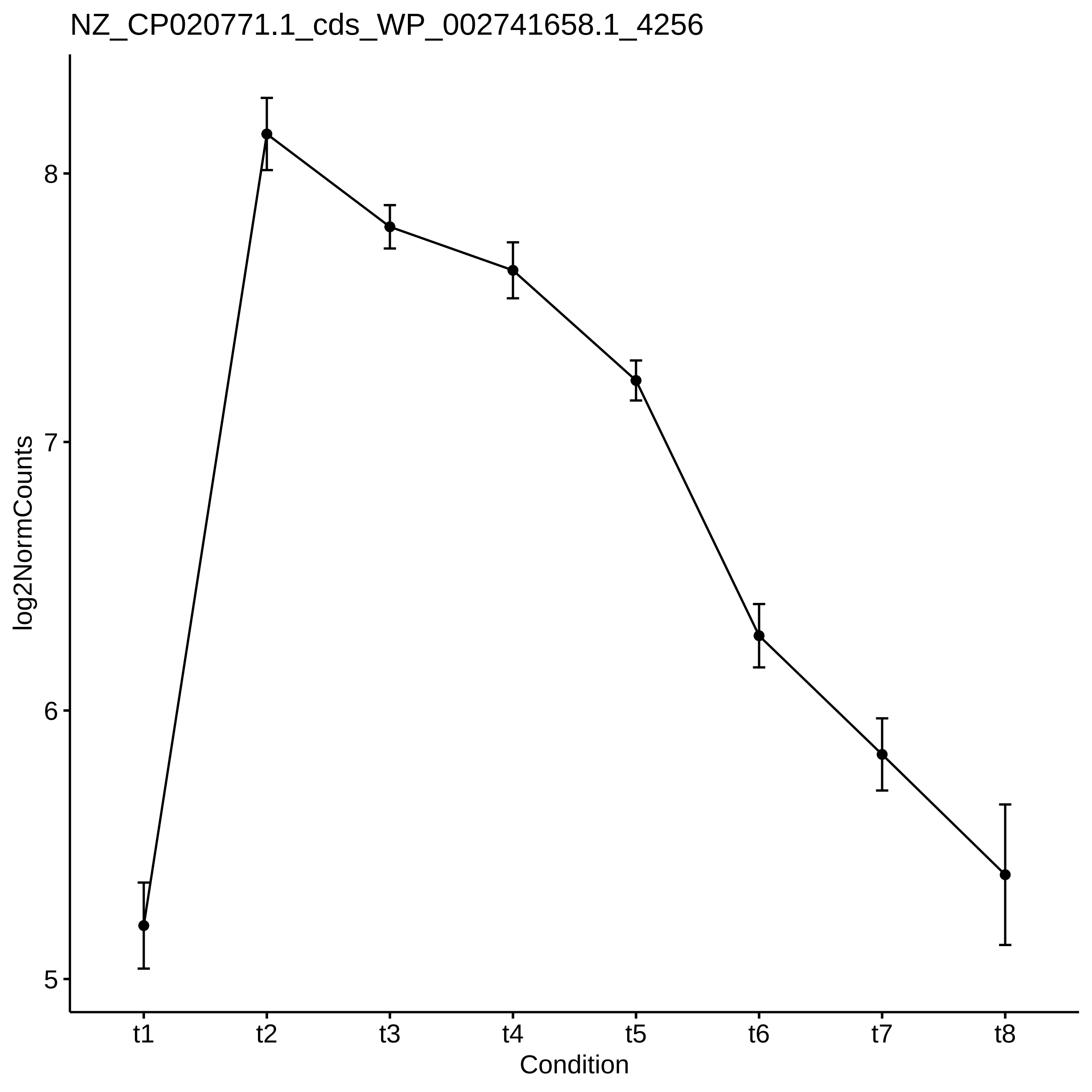

NZ_CP020771.1_cds_WP_002741658.1_4256
log2NormCounts
8
7
6
5
t1
t2
t3
t4
t5
t6
t7
t8
Condition

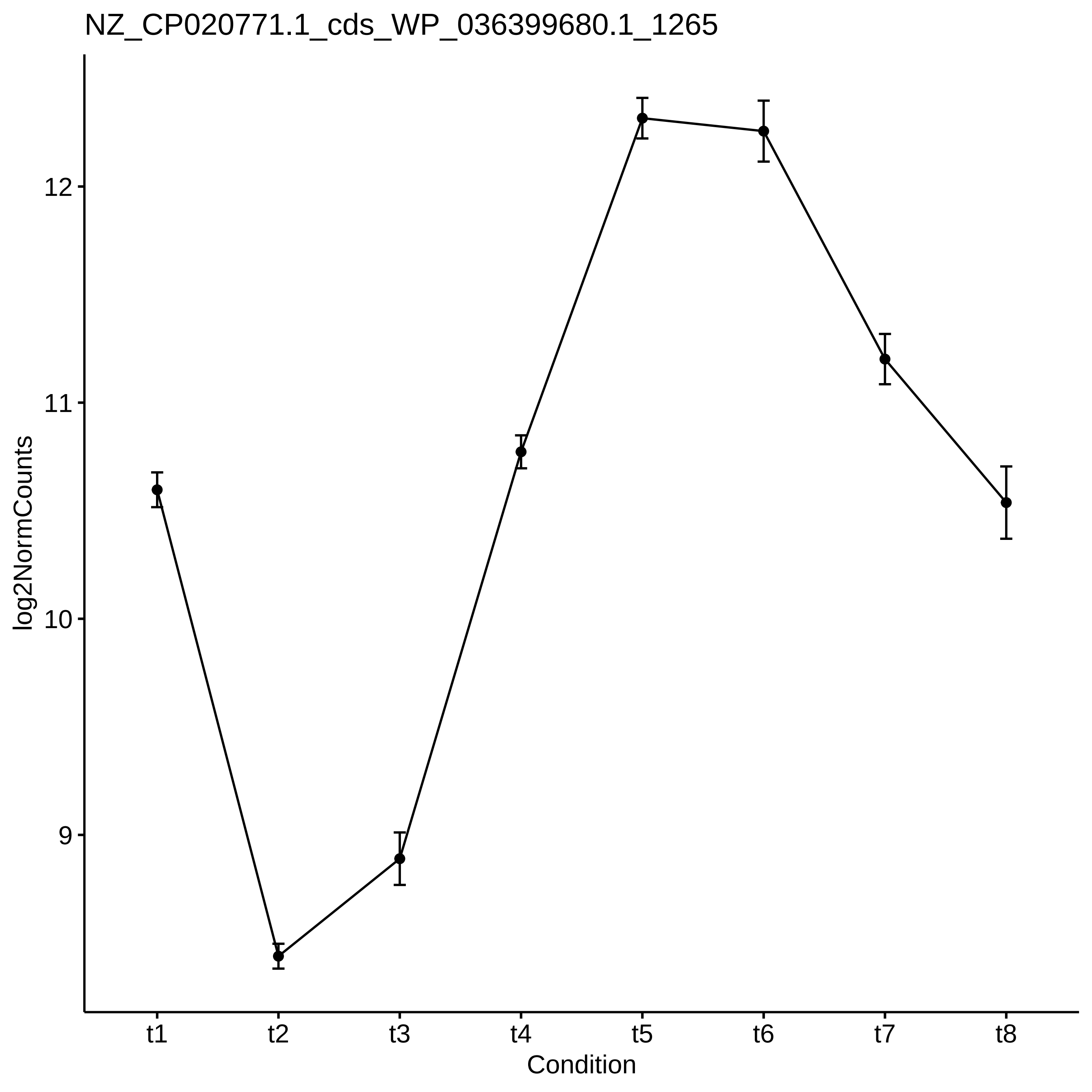
NZ_CP020771.1_cds_WP_036399680.1_1265
log2NormCounts
12
11
10
9
t1
t2
t3
t4
t5
t6
t7
t8
Condition

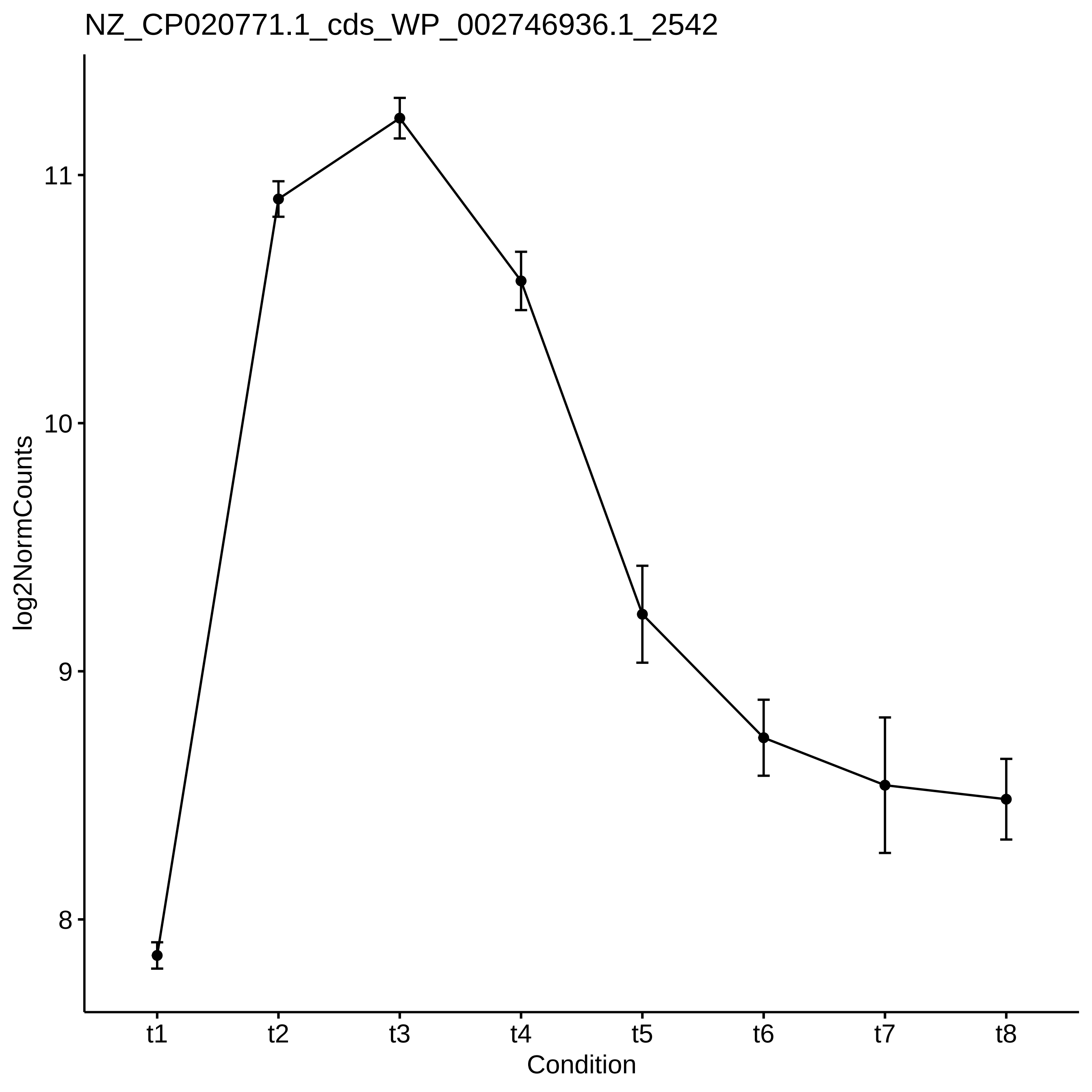

NZ_CP020771.1_cds_WP_002746936.1_2542
11
10
9
8
log2NormCounts
t1
t2
t3
t4
t5
t6
t7
t8
Condition

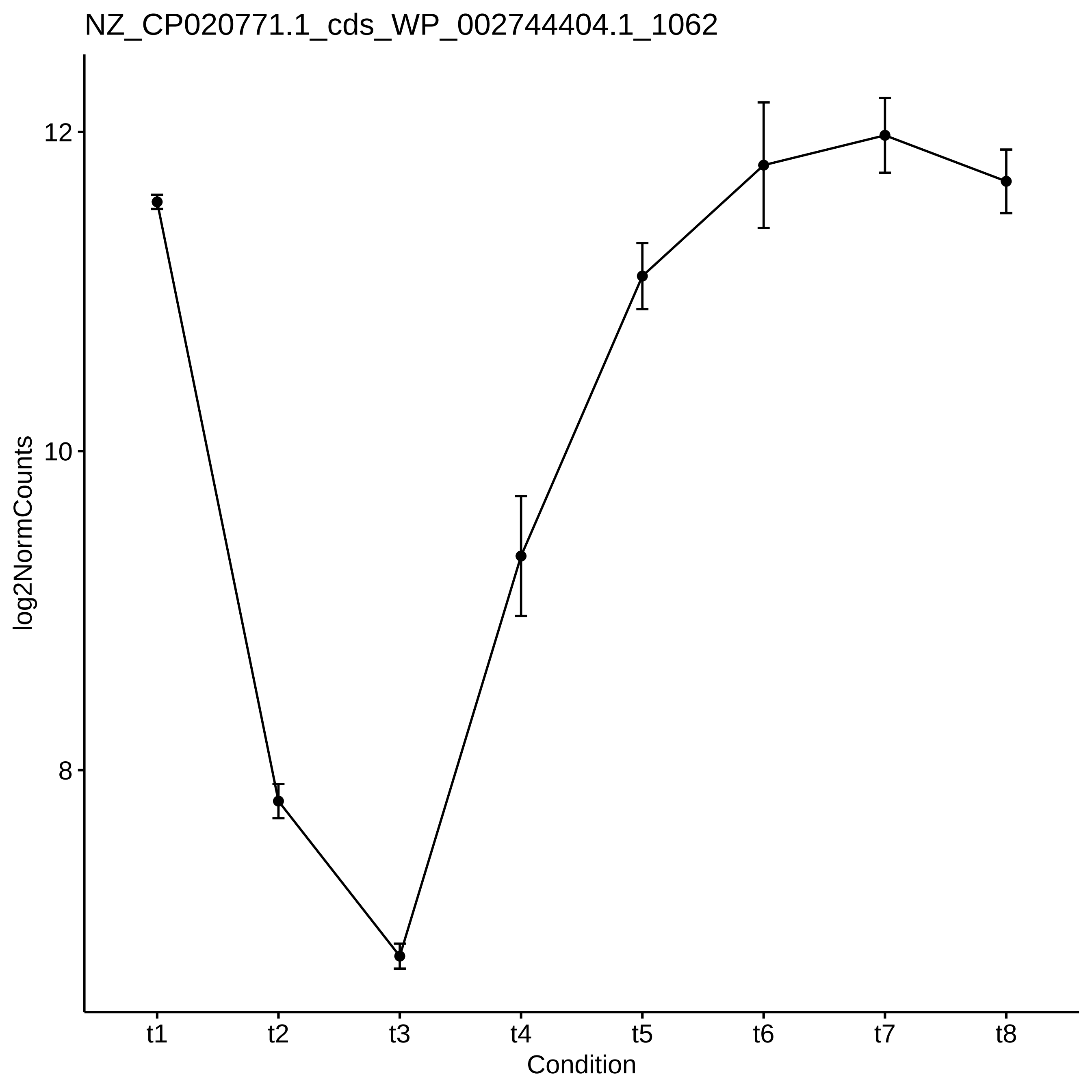

NZ_CP020771.1_cds_WP_002744404.1_1062
log2NormCounts
12
10
8
t1
t2
t3
t4
t5
t6
t7
t8
Condition

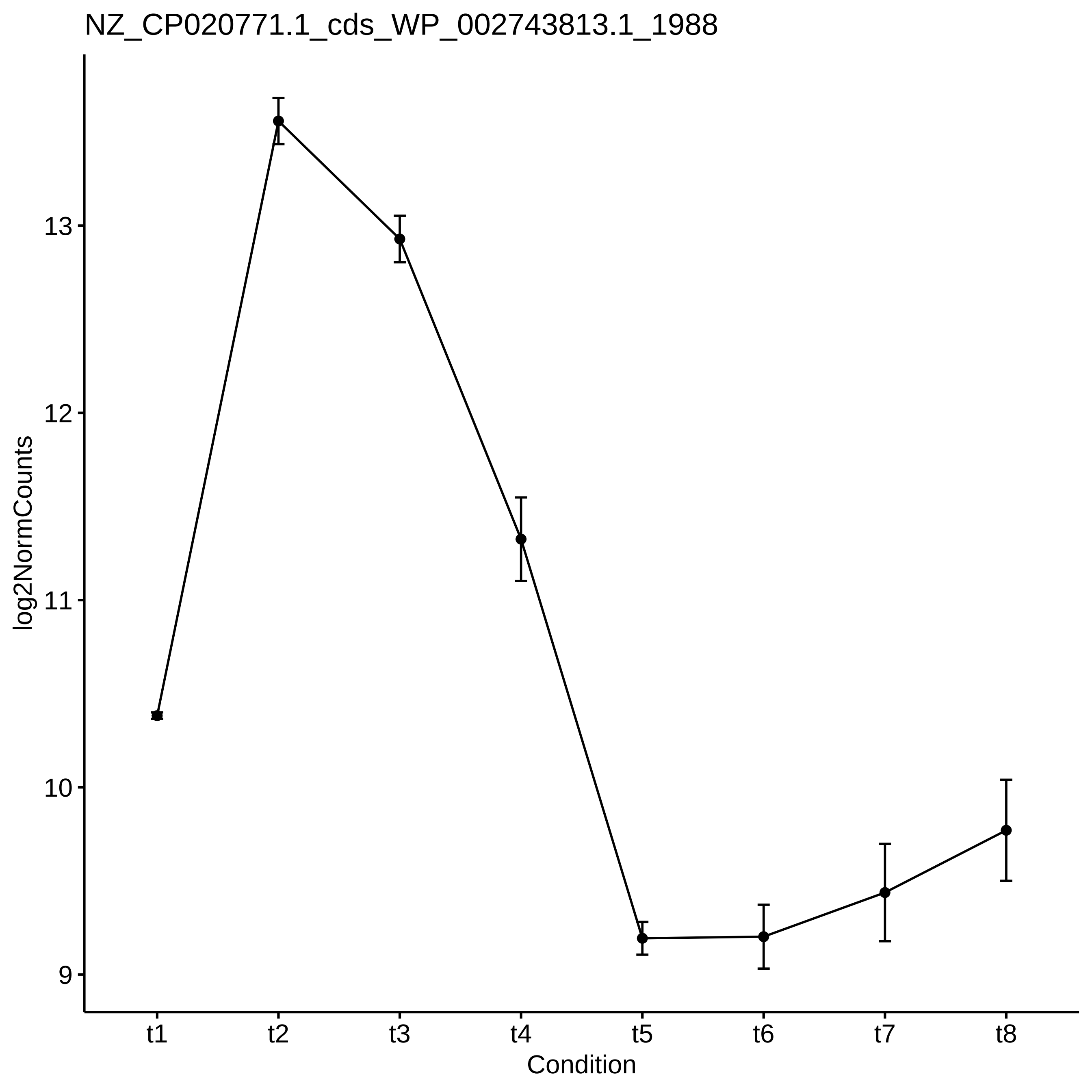
NZ_CP020771.1_cds_WP_002743813.1_1988
log2NormCounts
13
12
11
10
9
t1
t2
t3
t4
t5
t6
t7
t8
Condition

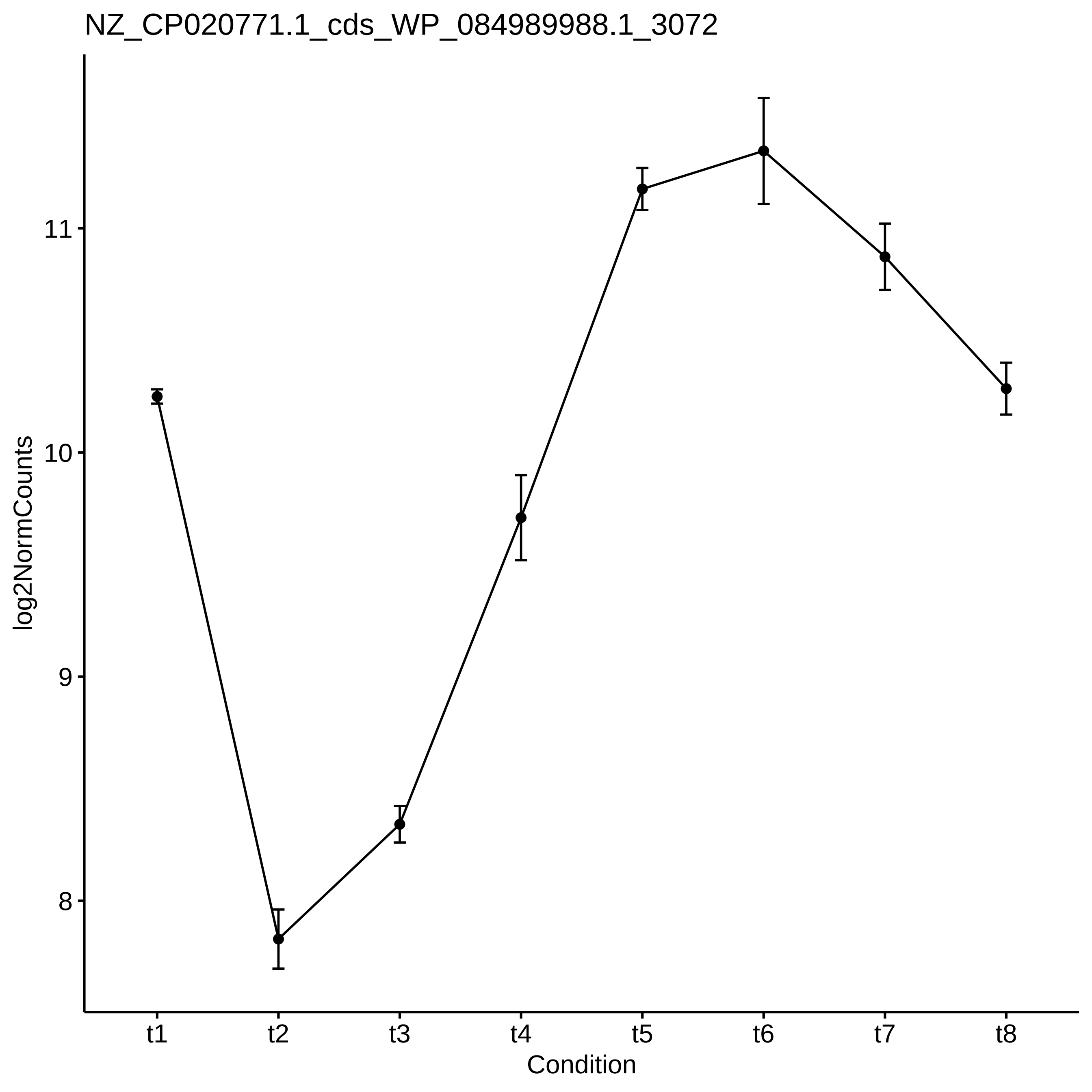
NZ_CP020771.1_cds_WP_084989988.1_3072
log2NormCounts
11
10
9
8
t1
t2
t3
t4
t5
t6
t7
t8
Condition

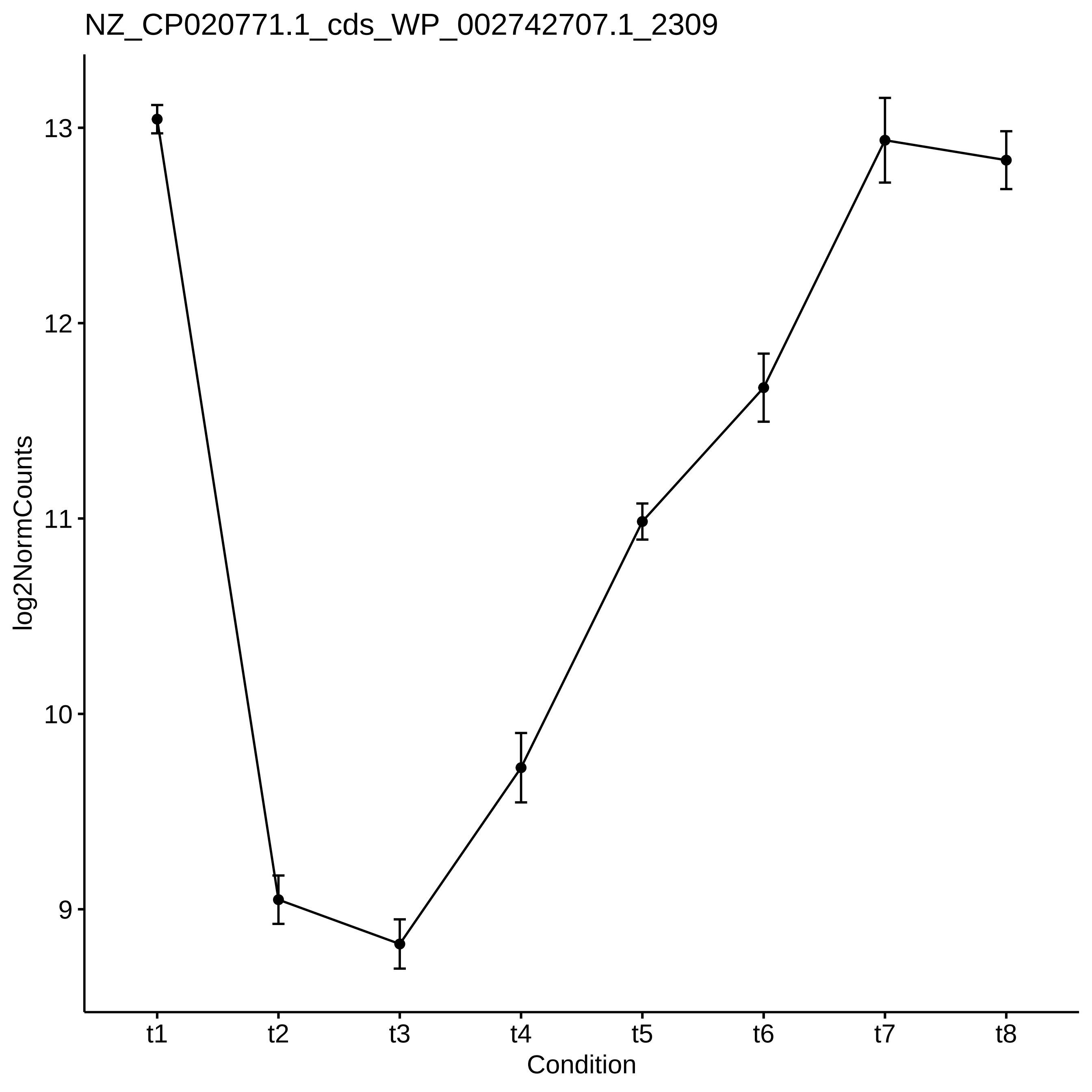

NZ_CP020771.1_cds_WP_002742707.1_2309
log2NormCounts
13
12
11
10
9
t1
t2
t3
t4
t5
t6
t7
t8
Condition

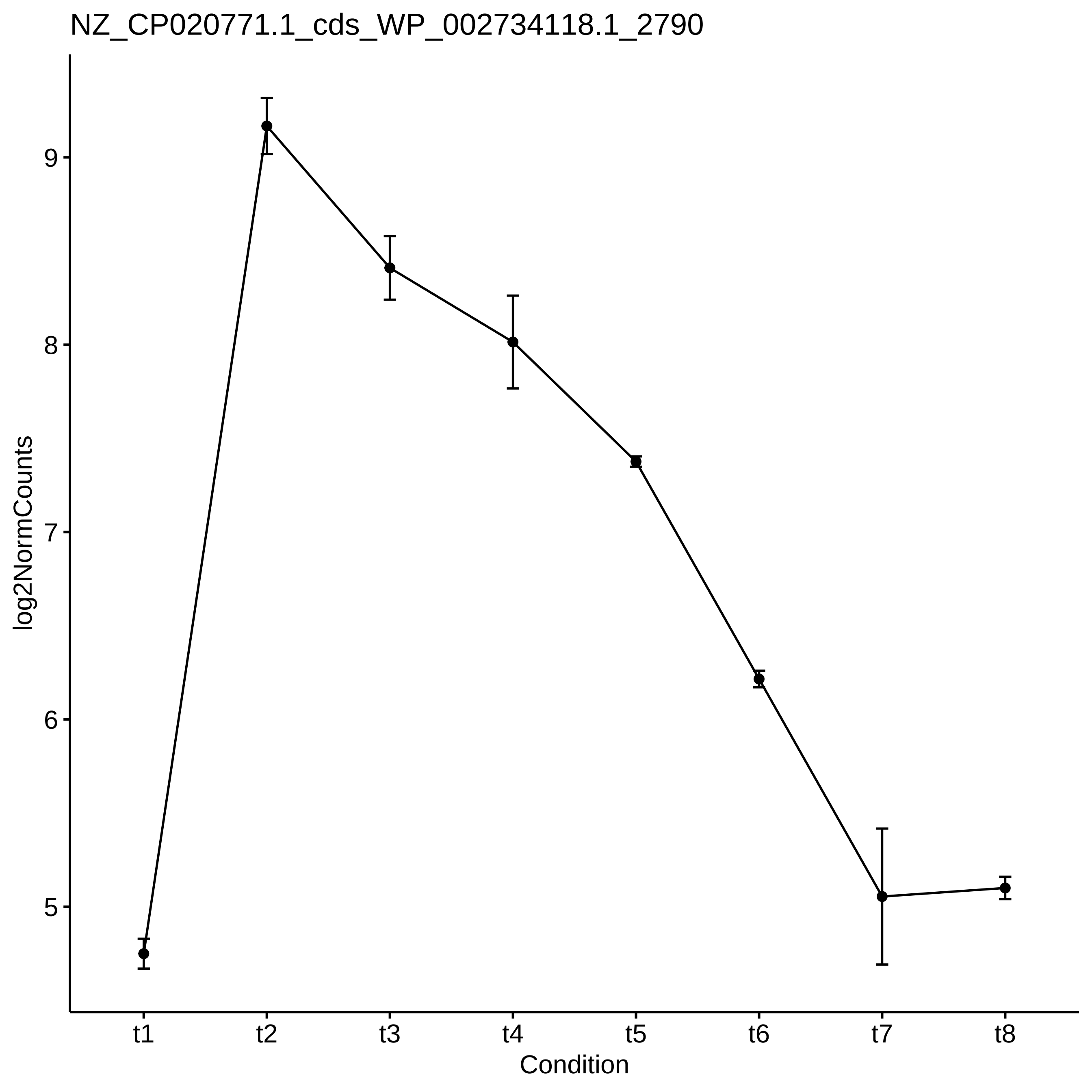

NZ_CP020771.1_cds_WP_002734118.1_2790
log2NormCounts
9
8
7
6
5
t1
t2
t3
t4
t5
t6
t7
t8
Condition

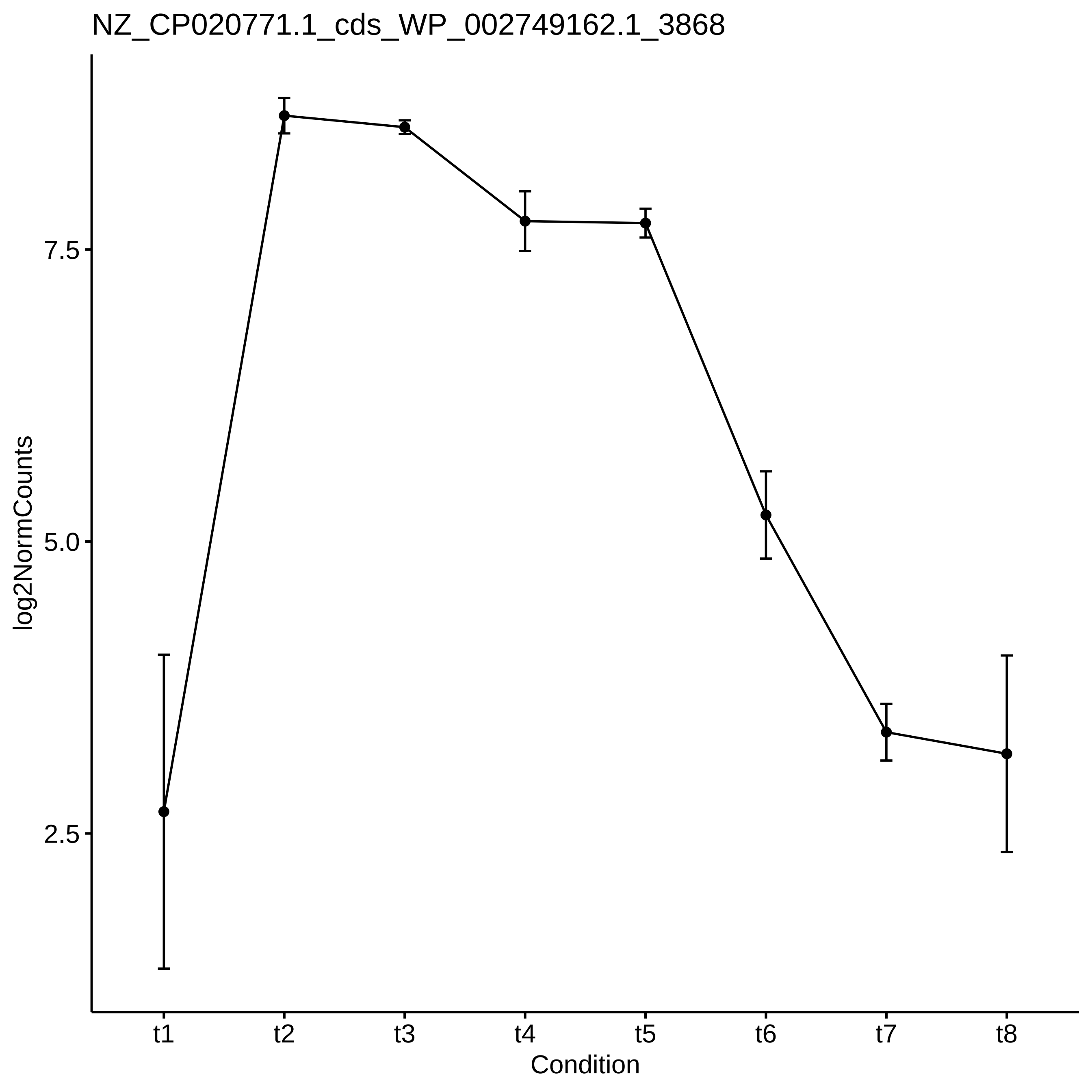
NZ_CP020771.1_cds_WP_002749162.1_3868
log2NormCounts
7.5
5.0
2.5
t1
t2
t3
t4
t5
t6
t7
t8
Condition

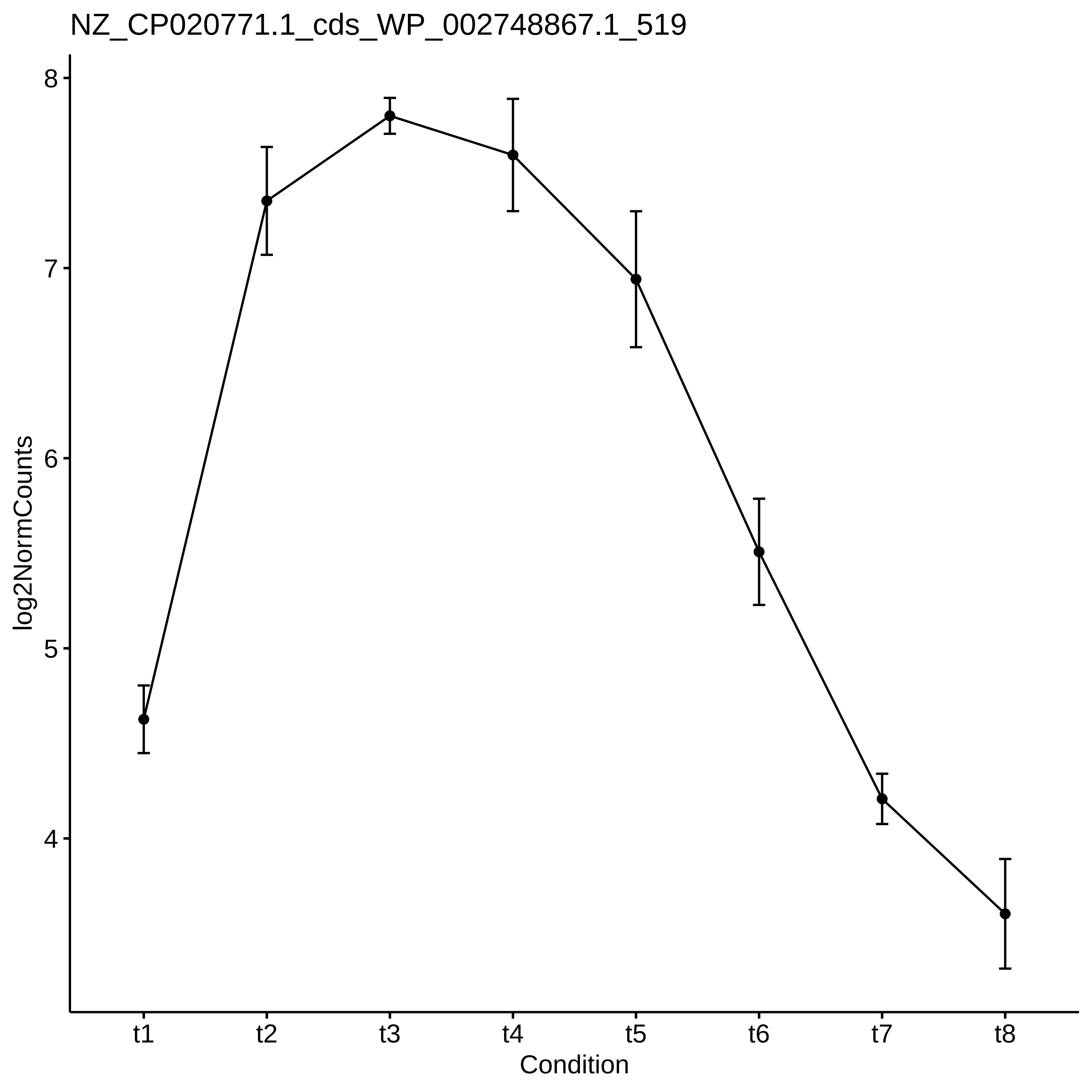

NZ_CP020771.1_cds_WP_002748867.1_519
log2NormCounts
8
7
6
5
4
t1
t2
t3
t4
t5
t6
t7
t8
Condition

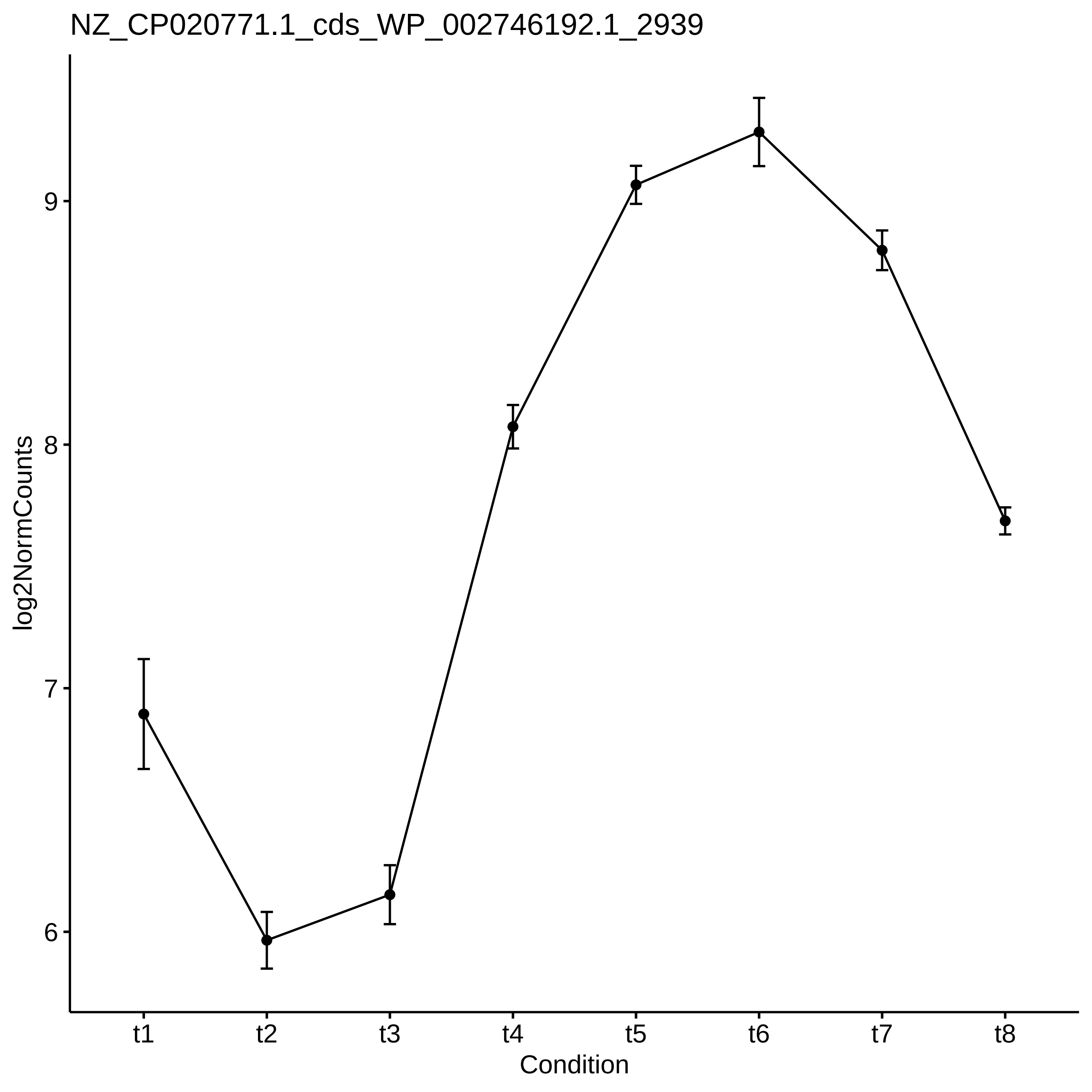

NZ_CP020771.1_cds_WP_002746192.1_2939
log2NormCounts
9
8
7
6
t1
t2
t3
t4
t5
t6
t7
t8
Condition

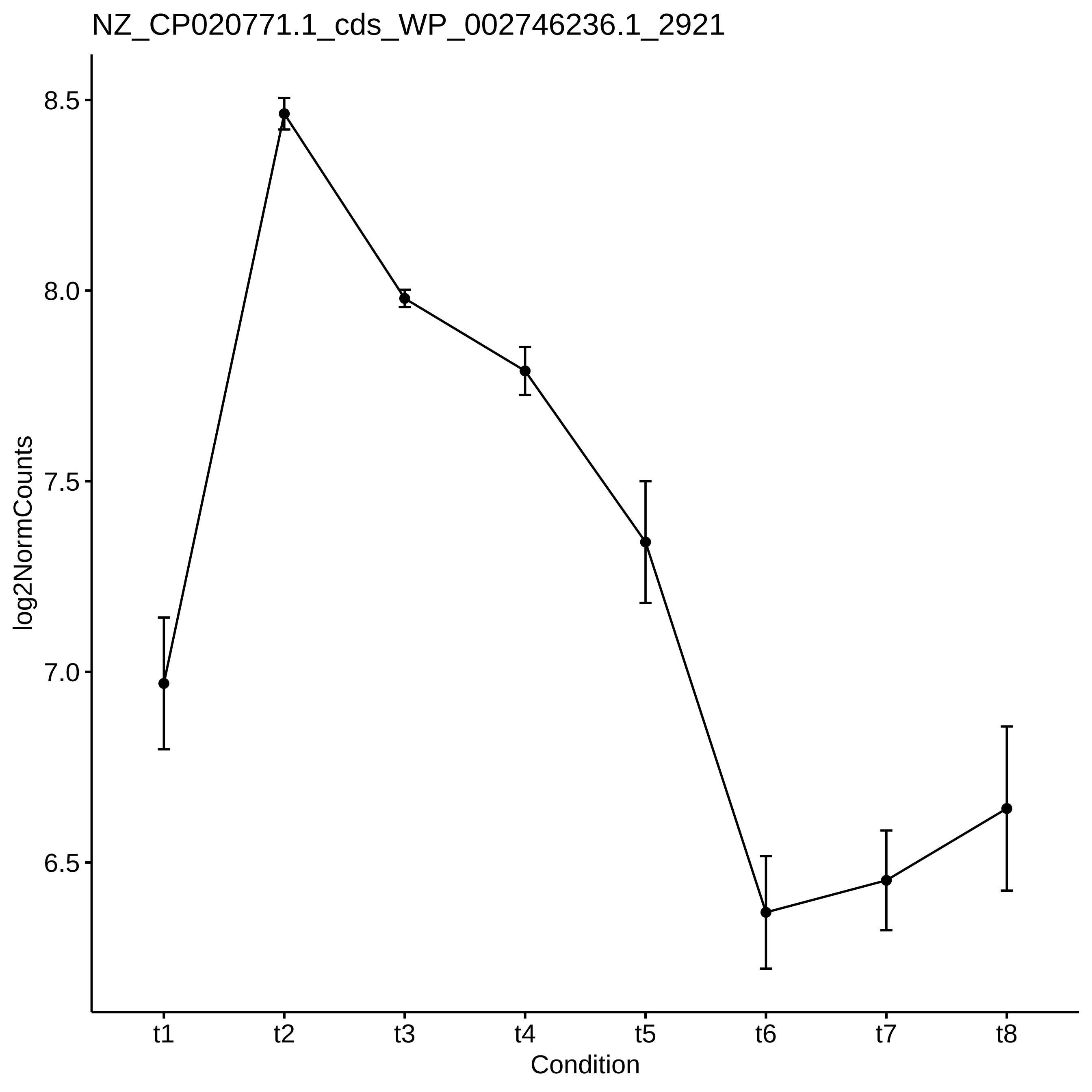
NZ_CP020771.1_cds_WP_002746236.1_2921
log2NormCounts
8.5
8.0
7.5
7.0
6.5
t1
t2
t3
t4
t5
t6
t7
t8
Condition

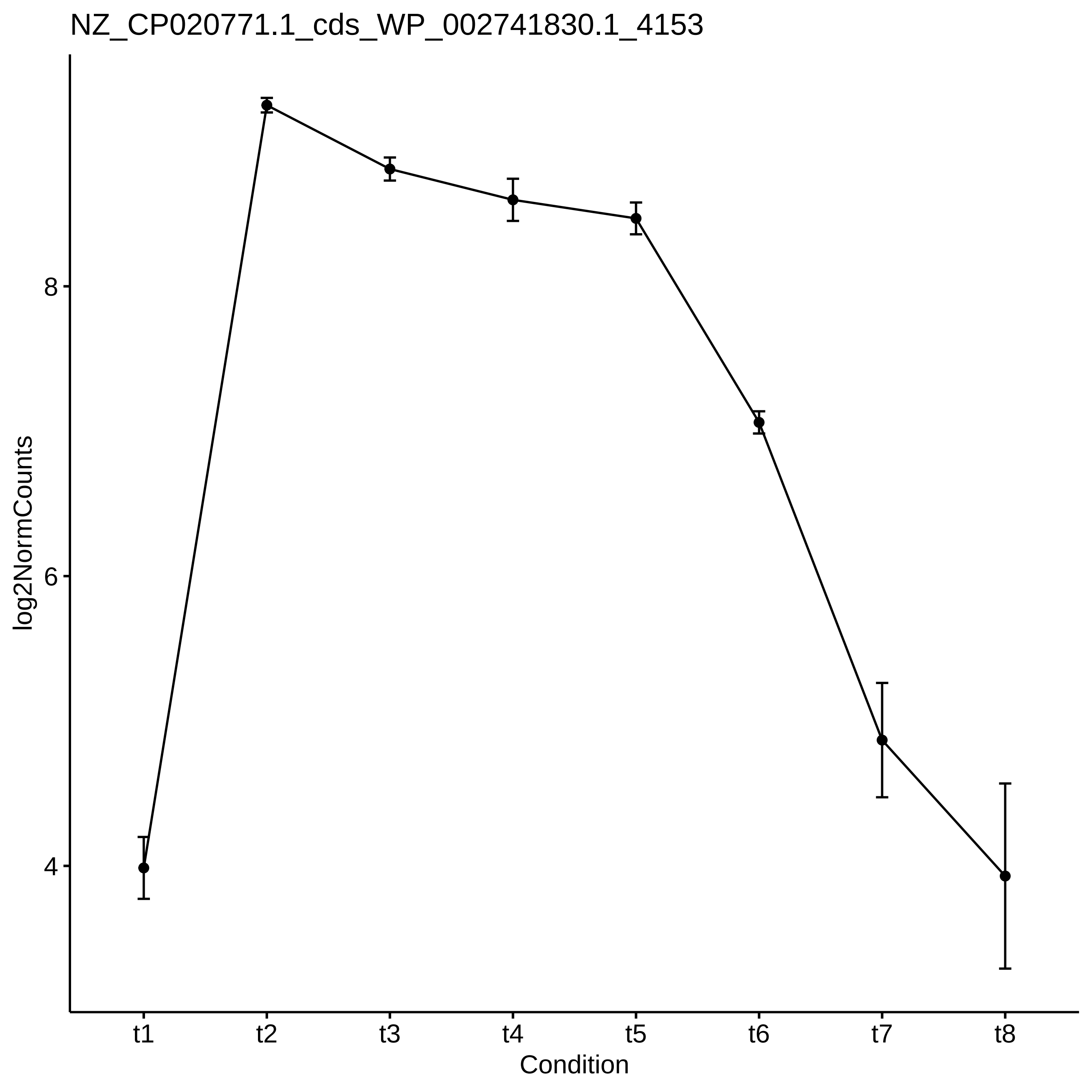

NZ_CP020771.1_cds_WP_002741830.1_4153
log2NormCounts
8
6
4
t1
t2
t3
t4
t5
t6
t7
t8
Condition

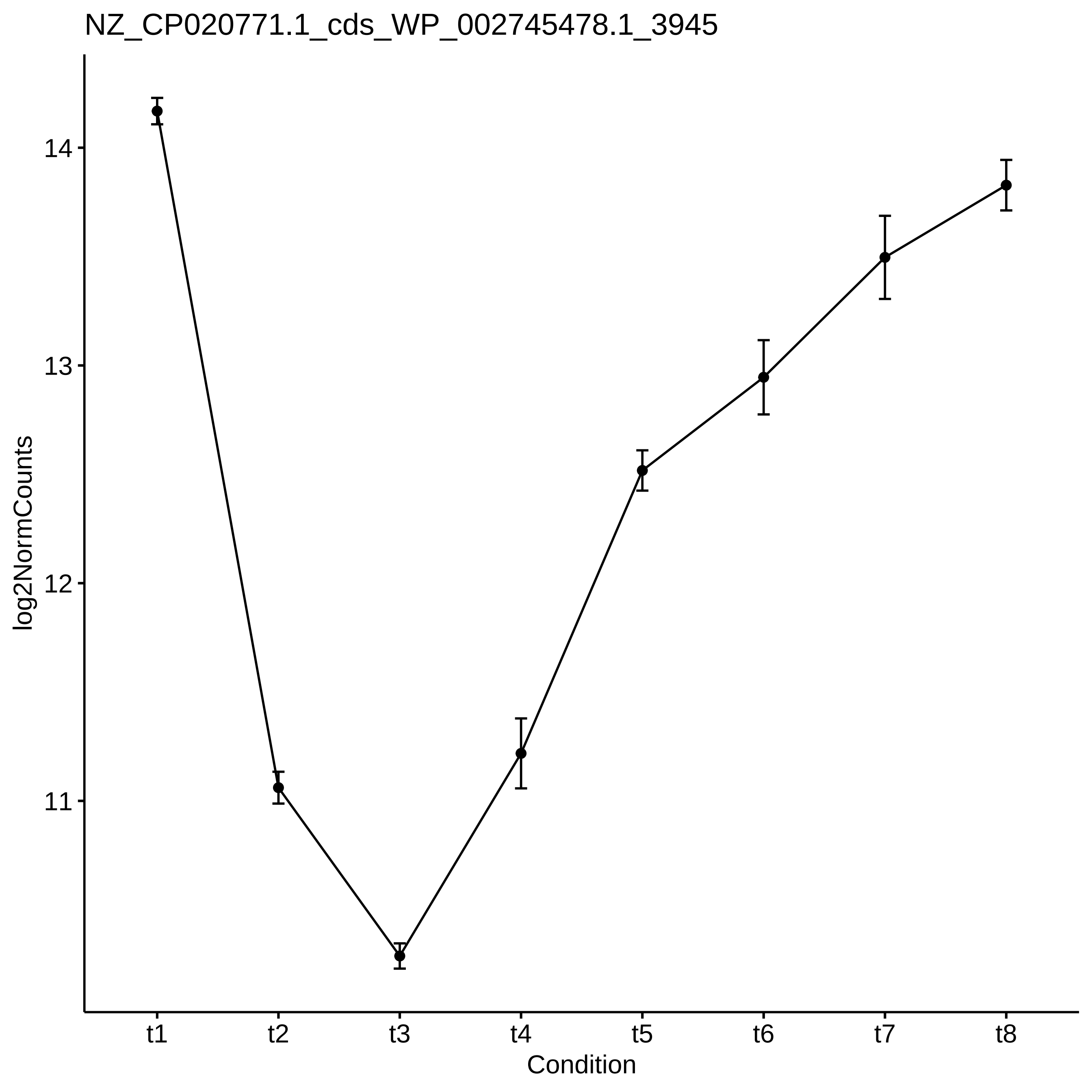

NZ_CP020771.1_cds_WP_002745478.1_3945
14
13
12
11
log2NormCounts
t1
t2
t3
t4
t5
t6
t7
t8
Condition

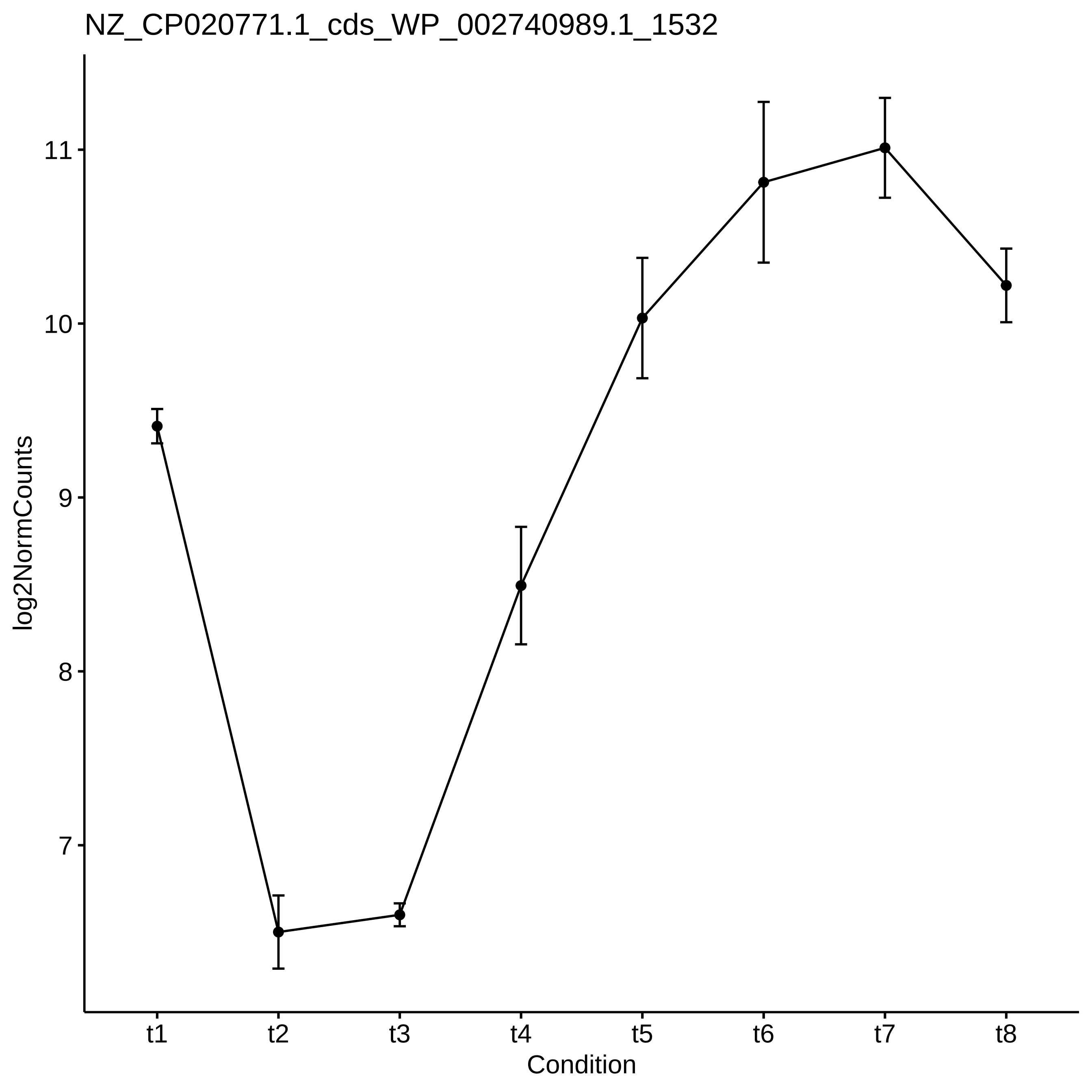
NZ_CP020771.1_cds_WP_002740989.1_1532
log2NormCounts
11
10
9
8
7
t1
t2
t3
t4
t5
t6
t7
t8
Condition
