## Supplementary figures and images for "Diel changes in the expression of a marker gene and candidate genes for intracellular amorphous CaCO_3_ biomineralization in *Microcystis*"

### Figure_S2.pdf

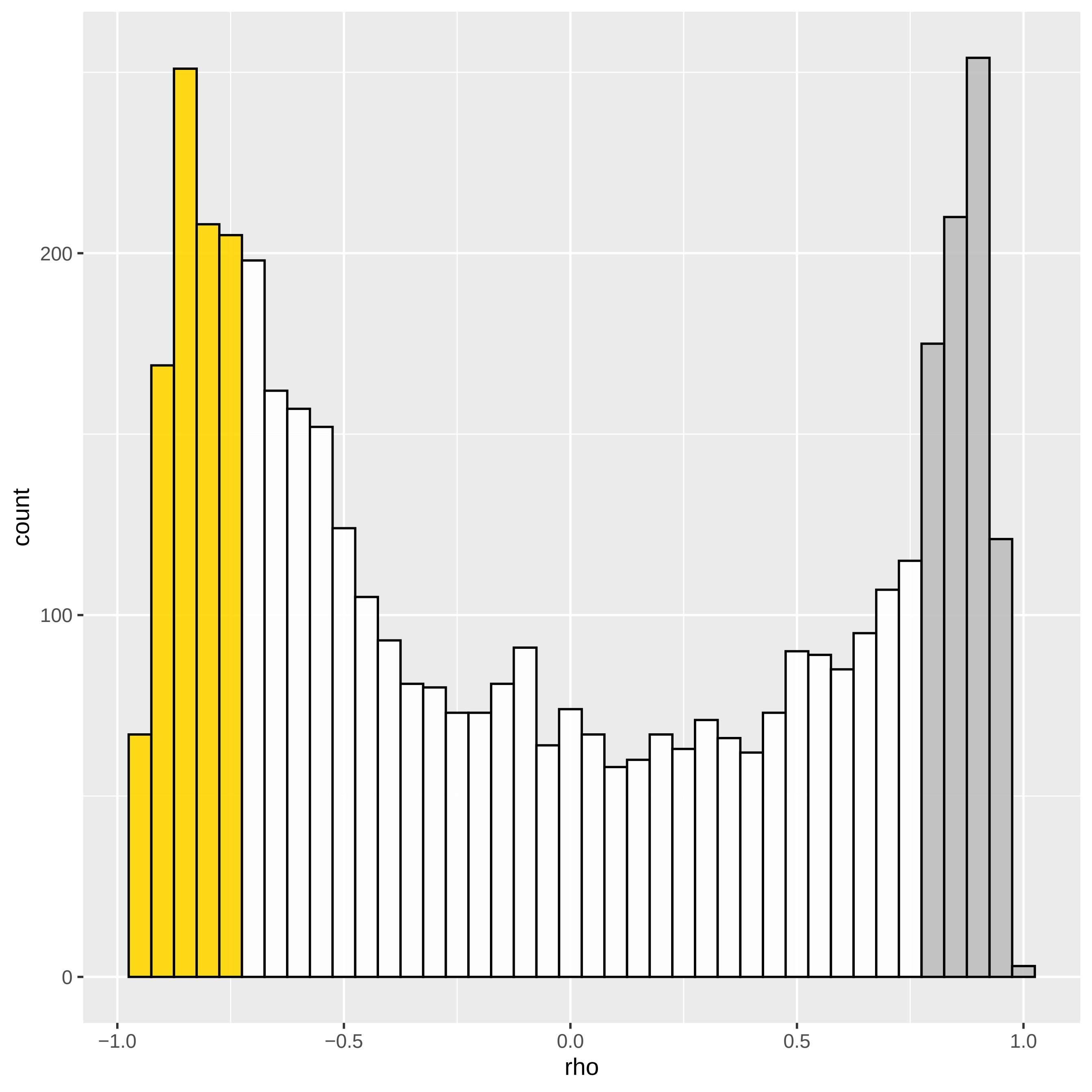

### PCC7806_Down_Up_DEG.pdf

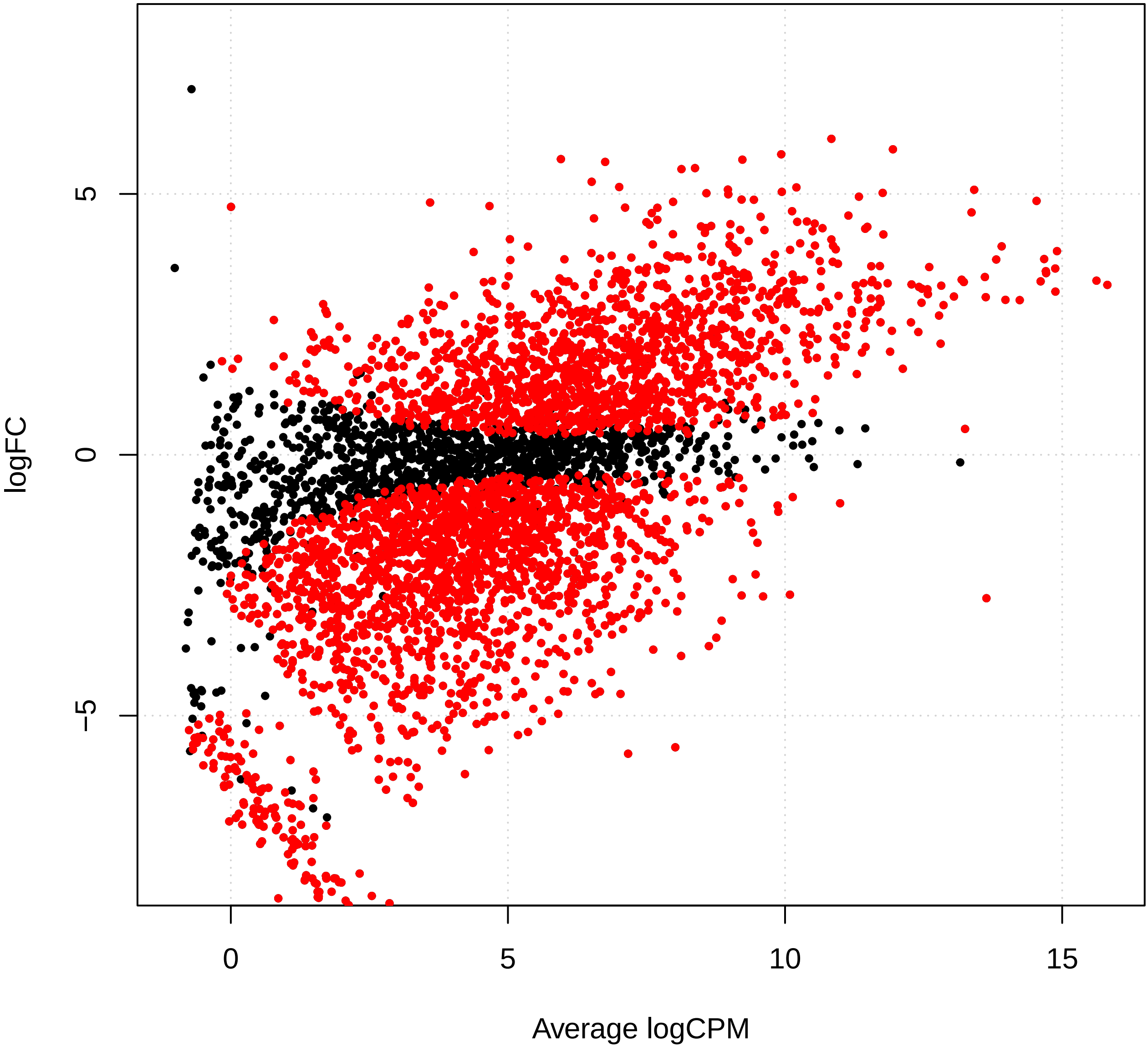

Supplement: Supplementary_Data [file 602159_file03.zip › Supplementary_DATA/DICOEXPRESS_output/DiffAnalysis/[t1-t2]/PCC7806_[t1-t2]_plotSmear.pdf]

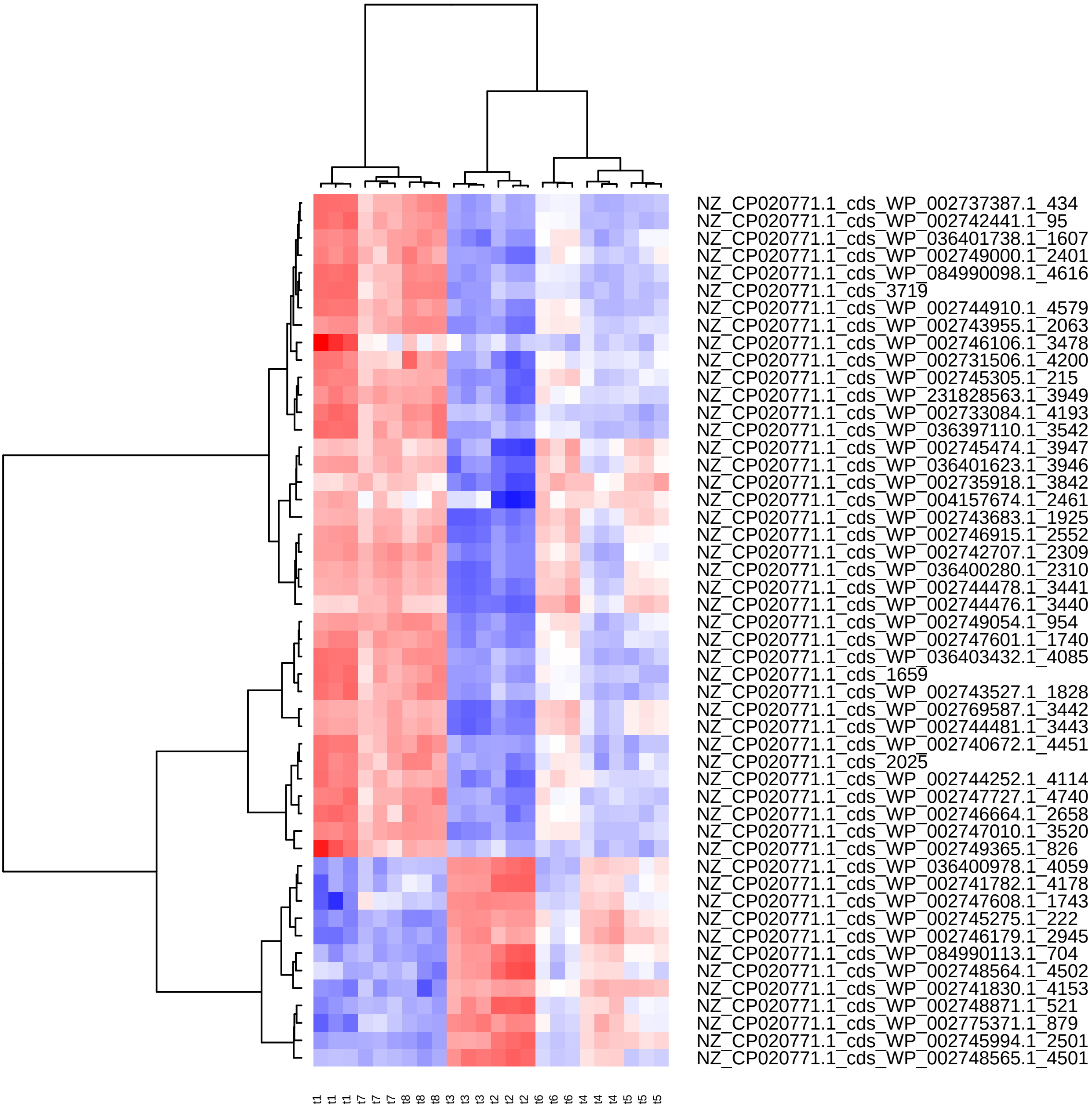

Supplement: Supplementary_Data [file 602159_file03.zip › Supplementary_DATA/DICOEXPRESS_output/DiffAnalysis/[t1-t2]/PCC7806_[t1-t2]_Top50_Clustering.pdf]

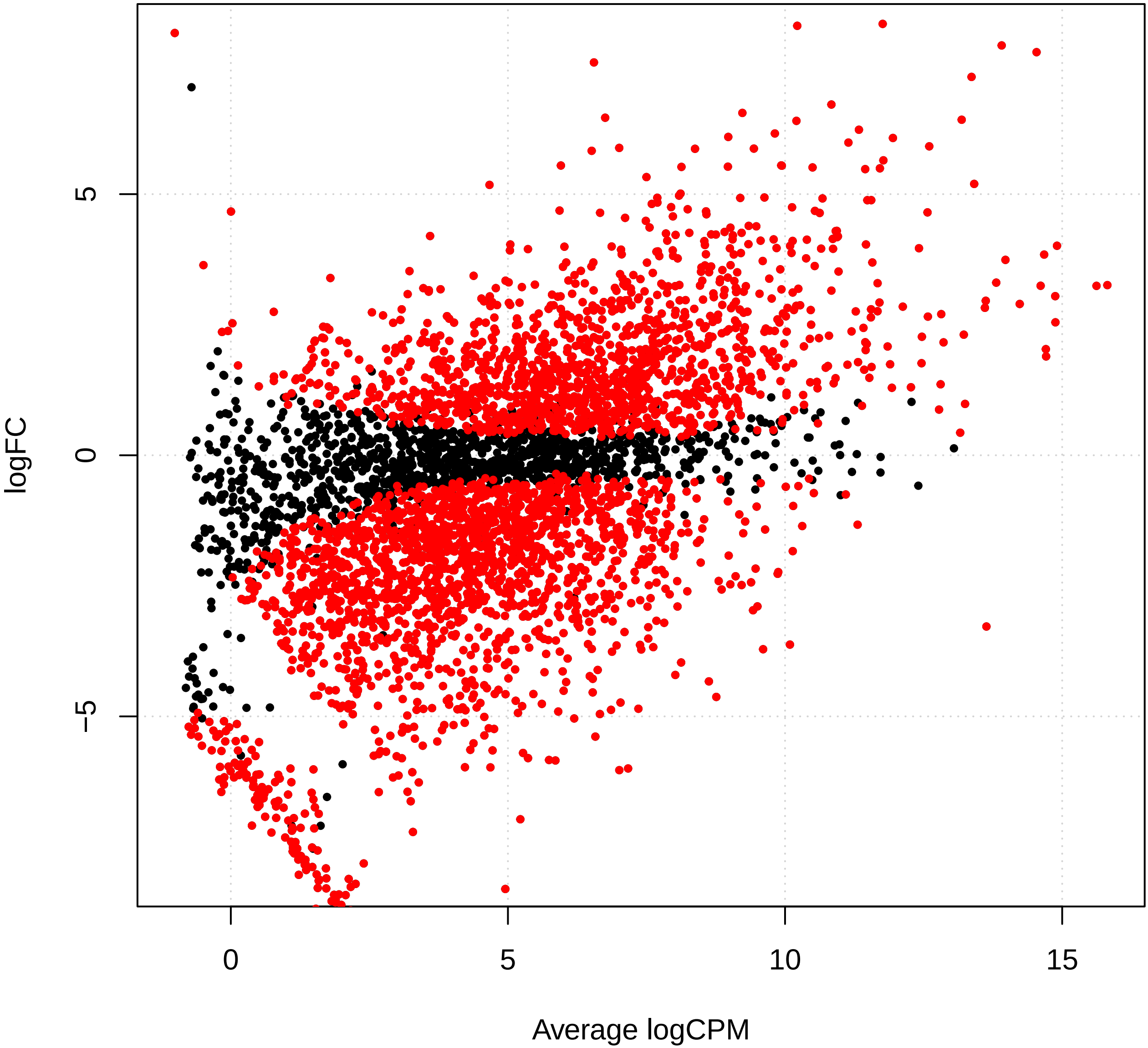

Supplement: Supplementary_Data [file 602159_file03.zip › Supplementary_DATA/DICOEXPRESS_output/DiffAnalysis/[t1-t3]/PCC7806_[t1-t3]_plotSmear.pdf]

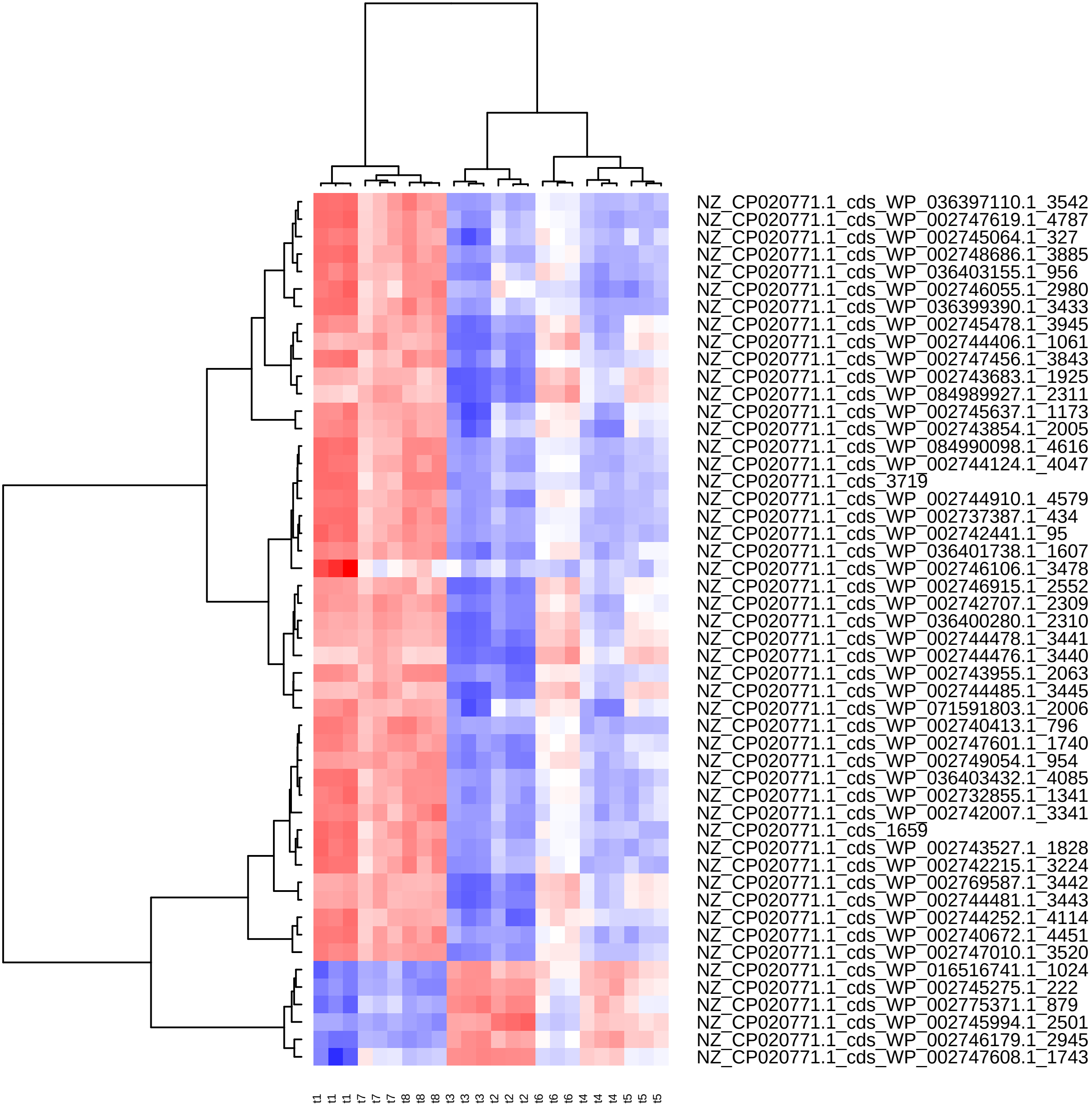

Supplement: Supplementary_Data [file 602159_file03.zip › Supplementary_DATA/DICOEXPRESS_output/DiffAnalysis/[t1-t3]/PCC7806_[t1-t3]_Top50_Clustering.pdf]

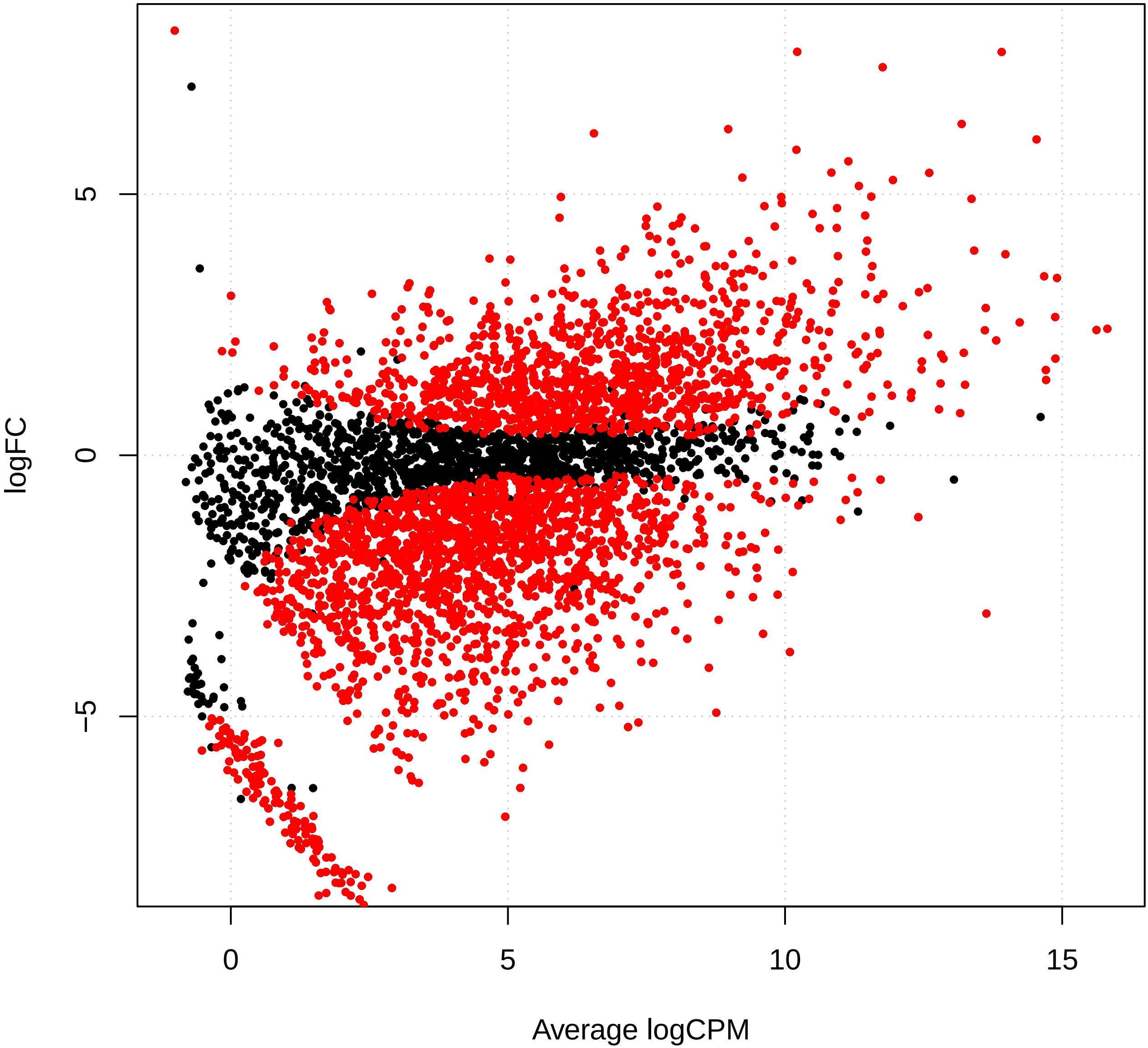

Supplement: Supplementary_Data [file 602159_file03.zip › Supplementary_DATA/DICOEXPRESS_output/DiffAnalysis/[t1-t4]/PCC7806_[t1-t4]_plotSmear.pdf]

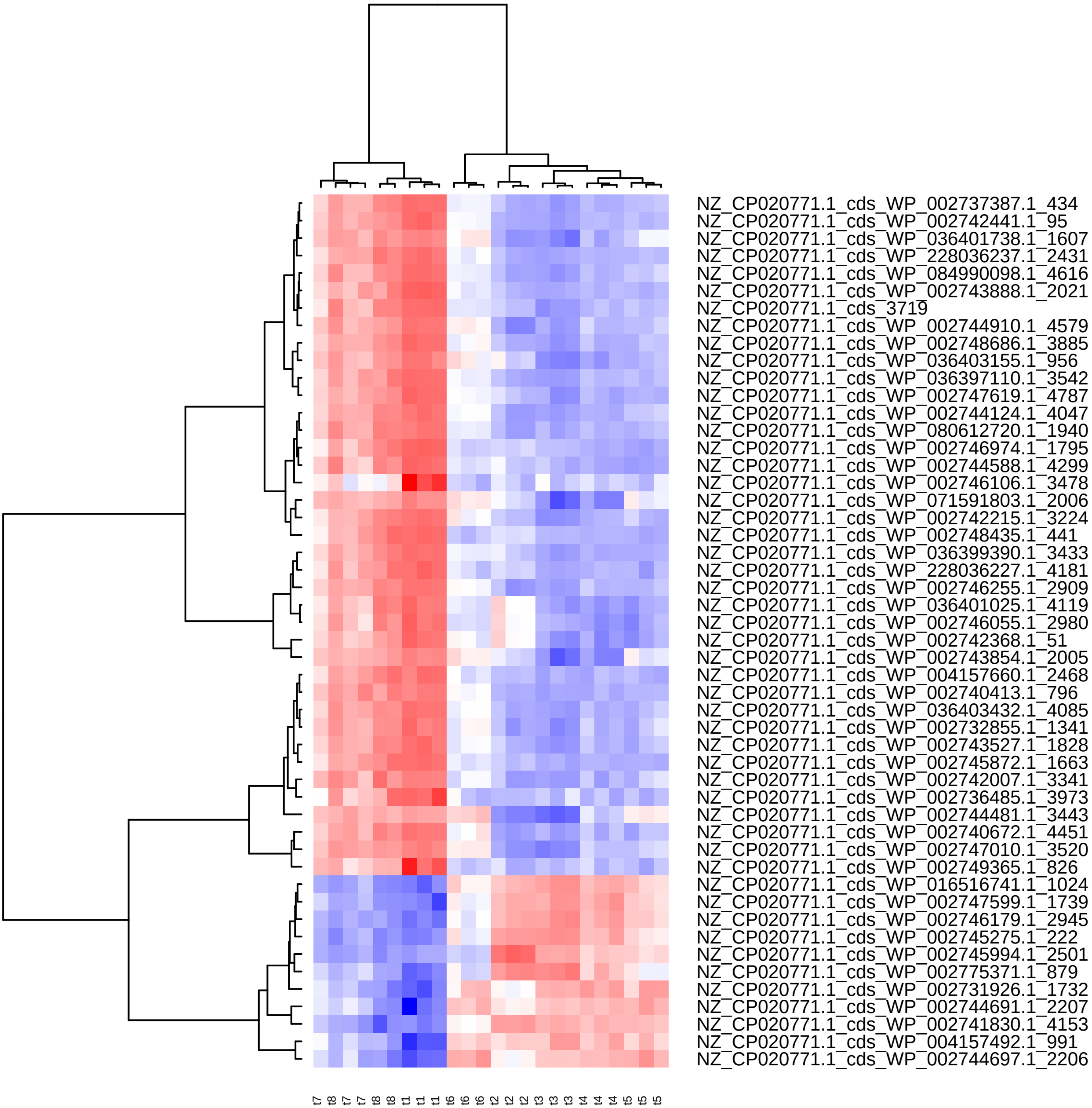

Supplement: Supplementary_Data [file 602159_file03.zip › Supplementary_DATA/DICOEXPRESS_output/DiffAnalysis/[t1-t4]/PCC7806_[t1-t4]_Top50_Clustering.pdf]

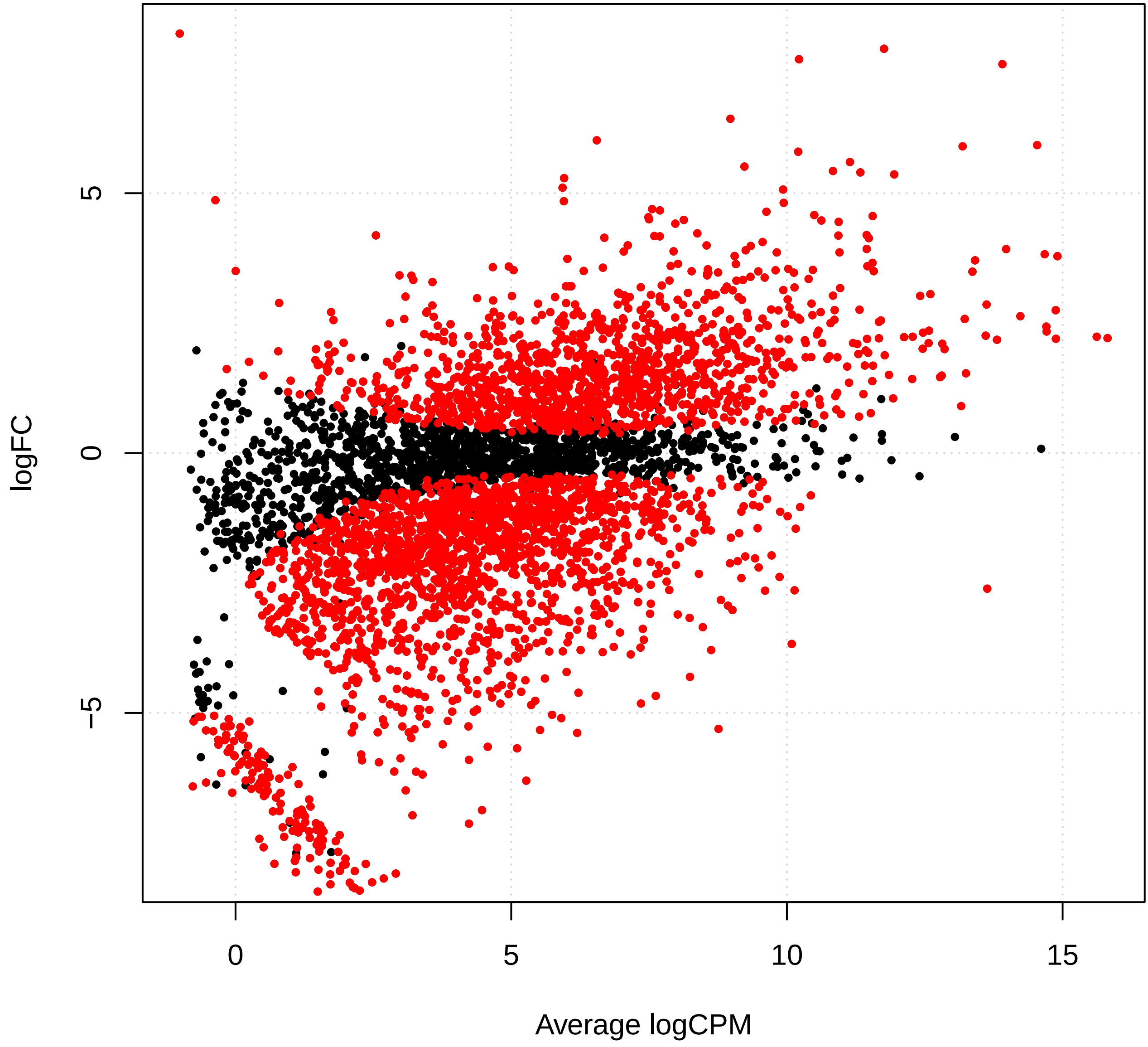

Supplement: Supplementary_Data [file 602159_file03.zip › Supplementary_DATA/DICOEXPRESS_output/DiffAnalysis/[t1-t5]/PCC7806_[t1-t5]_plotSmear.pdf]

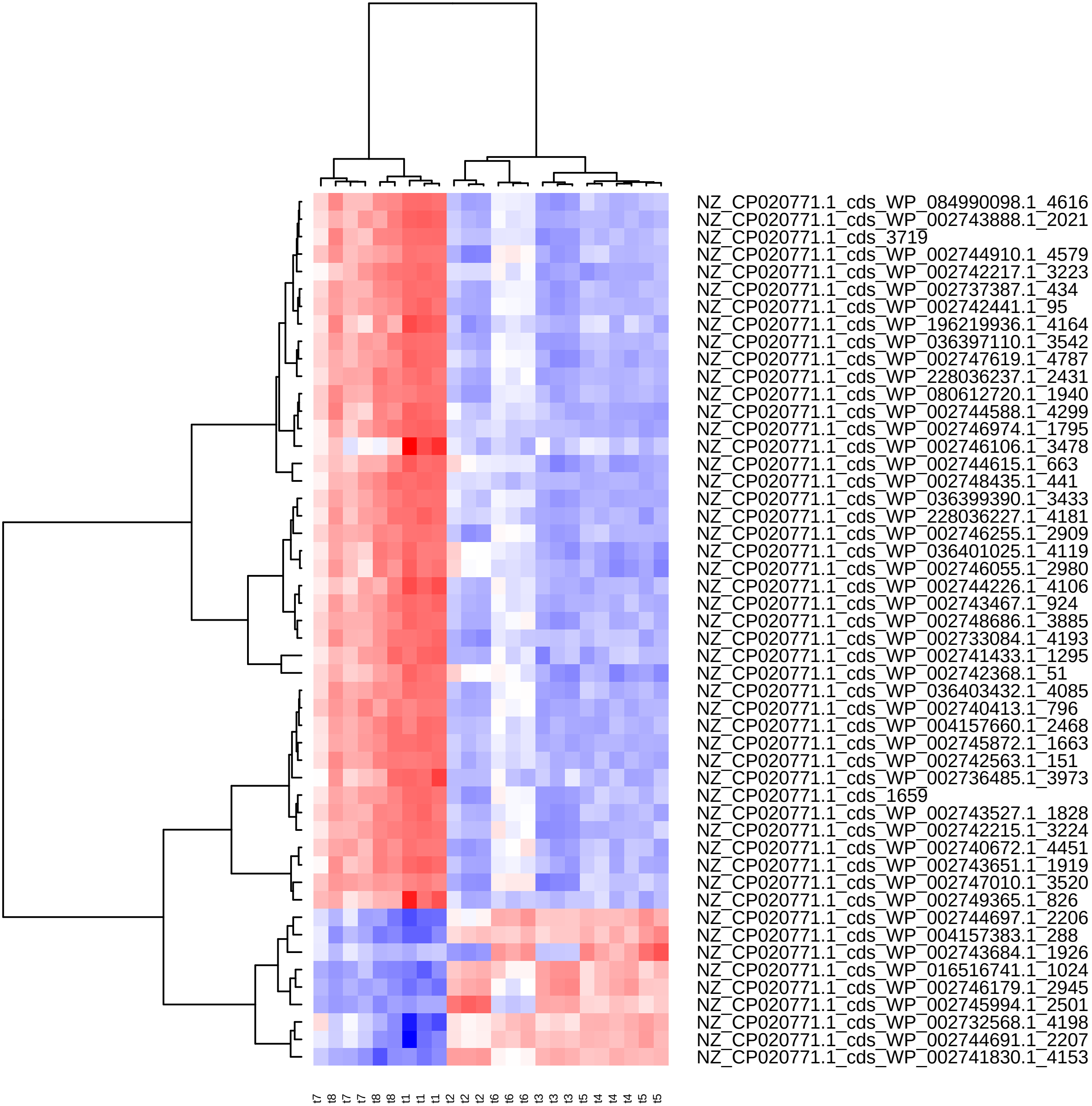

Supplement: Supplementary_Data [file 602159_file03.zip › Supplementary_DATA/DICOEXPRESS_output/DiffAnalysis/[t1-t5]/PCC7806_[t1-t5]_Top50_Clustering.pdf]

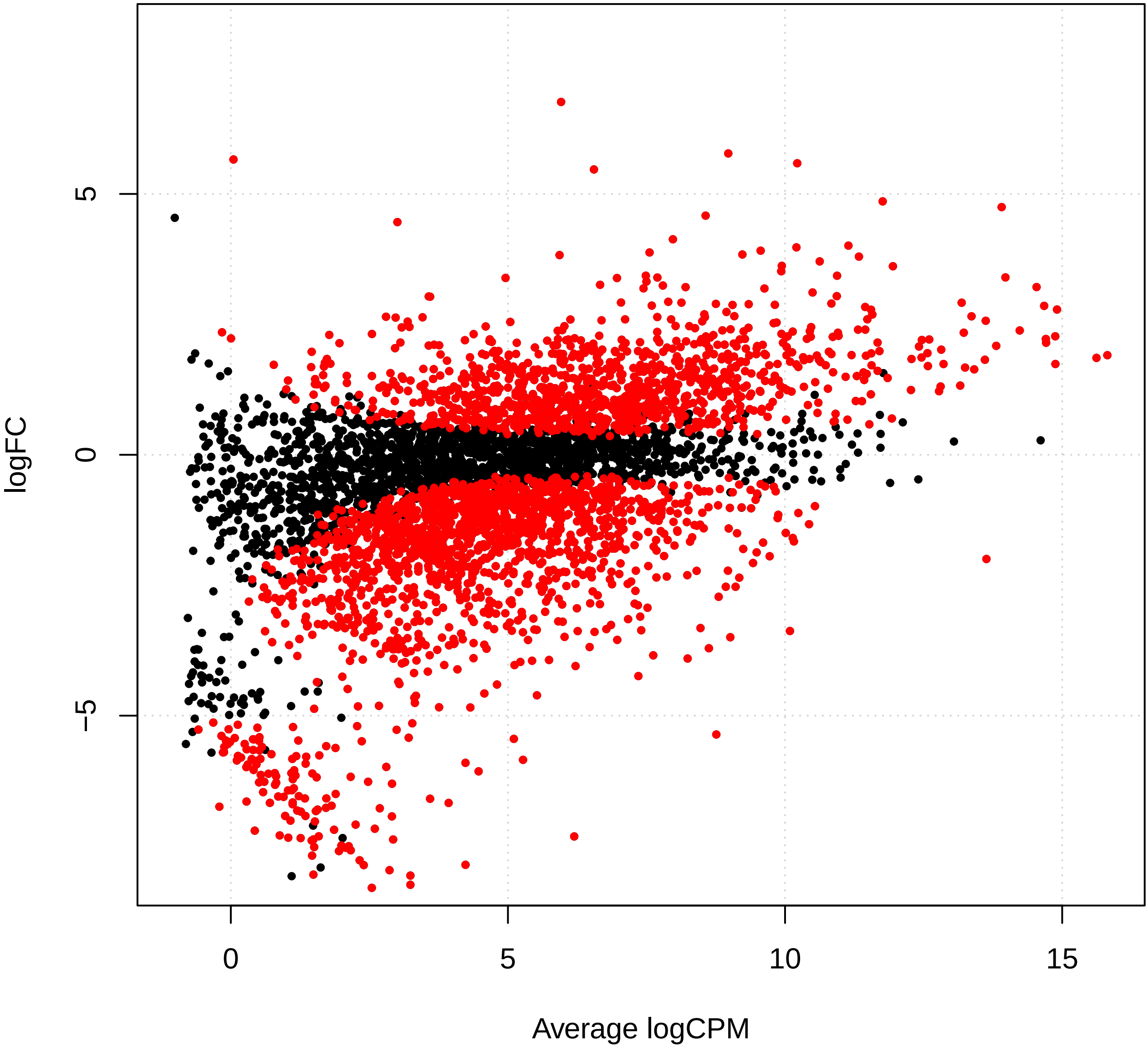

Supplement: Supplementary_Data [file 602159_file03.zip › Supplementary_DATA/DICOEXPRESS_output/DiffAnalysis/[t1-t6]/PCC7806_[t1-t6]_plotSmear.pdf]

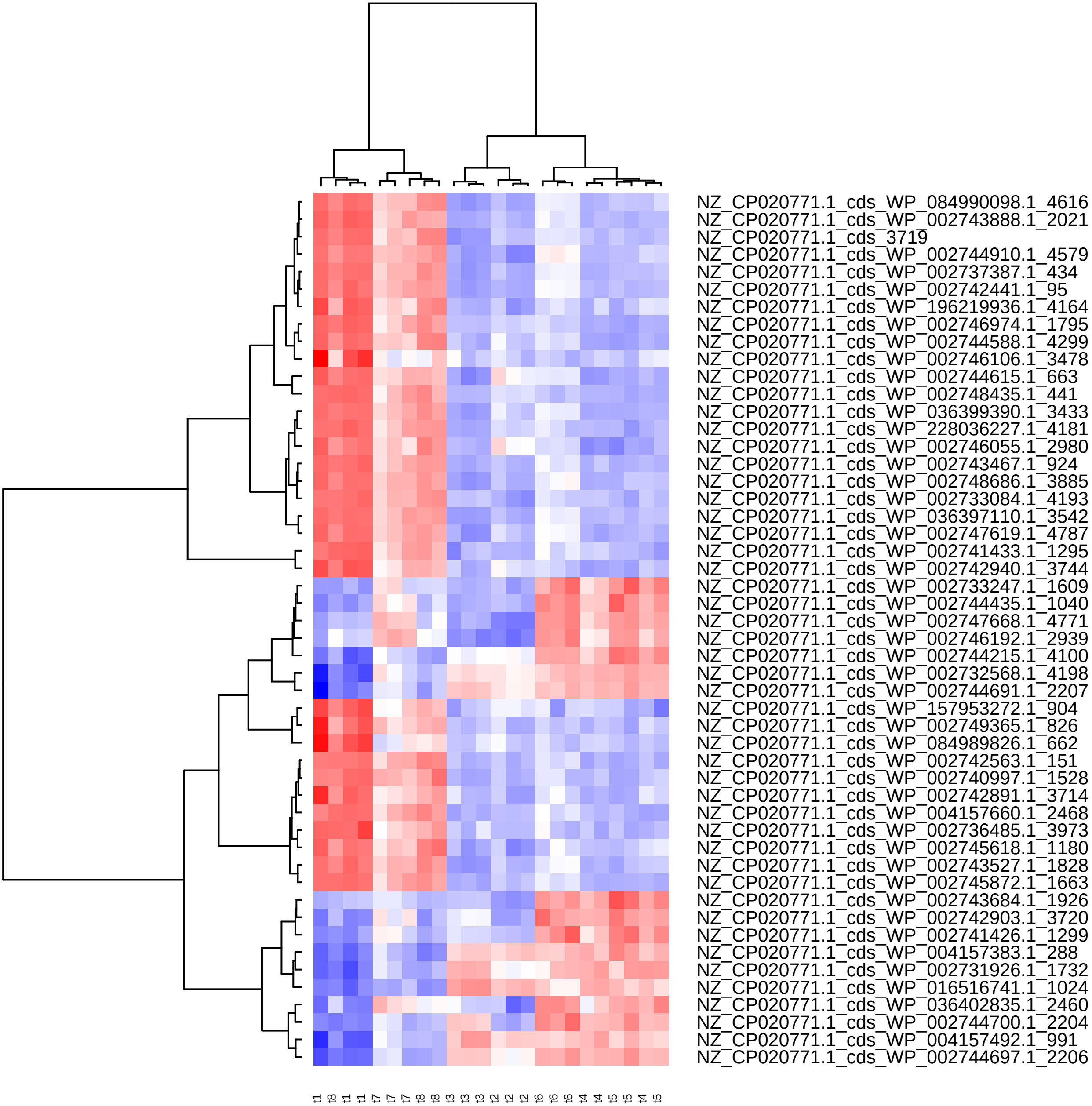

Supplement: Supplementary_Data [file 602159_file03.zip › Supplementary_DATA/DICOEXPRESS_output/DiffAnalysis/[t1-t6]/PCC7806_[t1-t6]_Top50_Clustering.pdf]

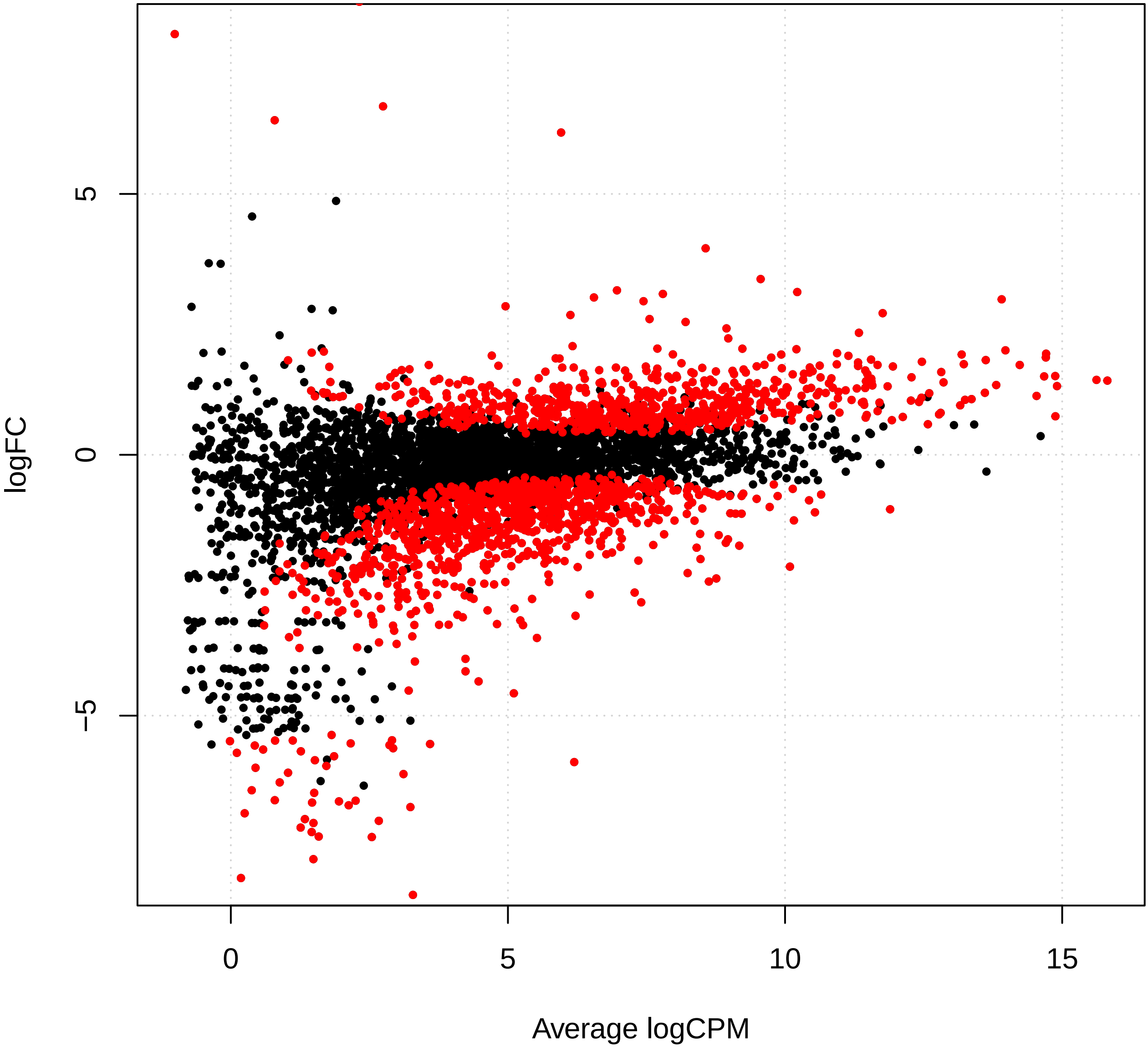

Supplement: Supplementary_Data [file 602159_file03.zip › Supplementary_DATA/DICOEXPRESS_output/DiffAnalysis/[t1-t7]/PCC7806_[t1-t7]_plotSmear.pdf]

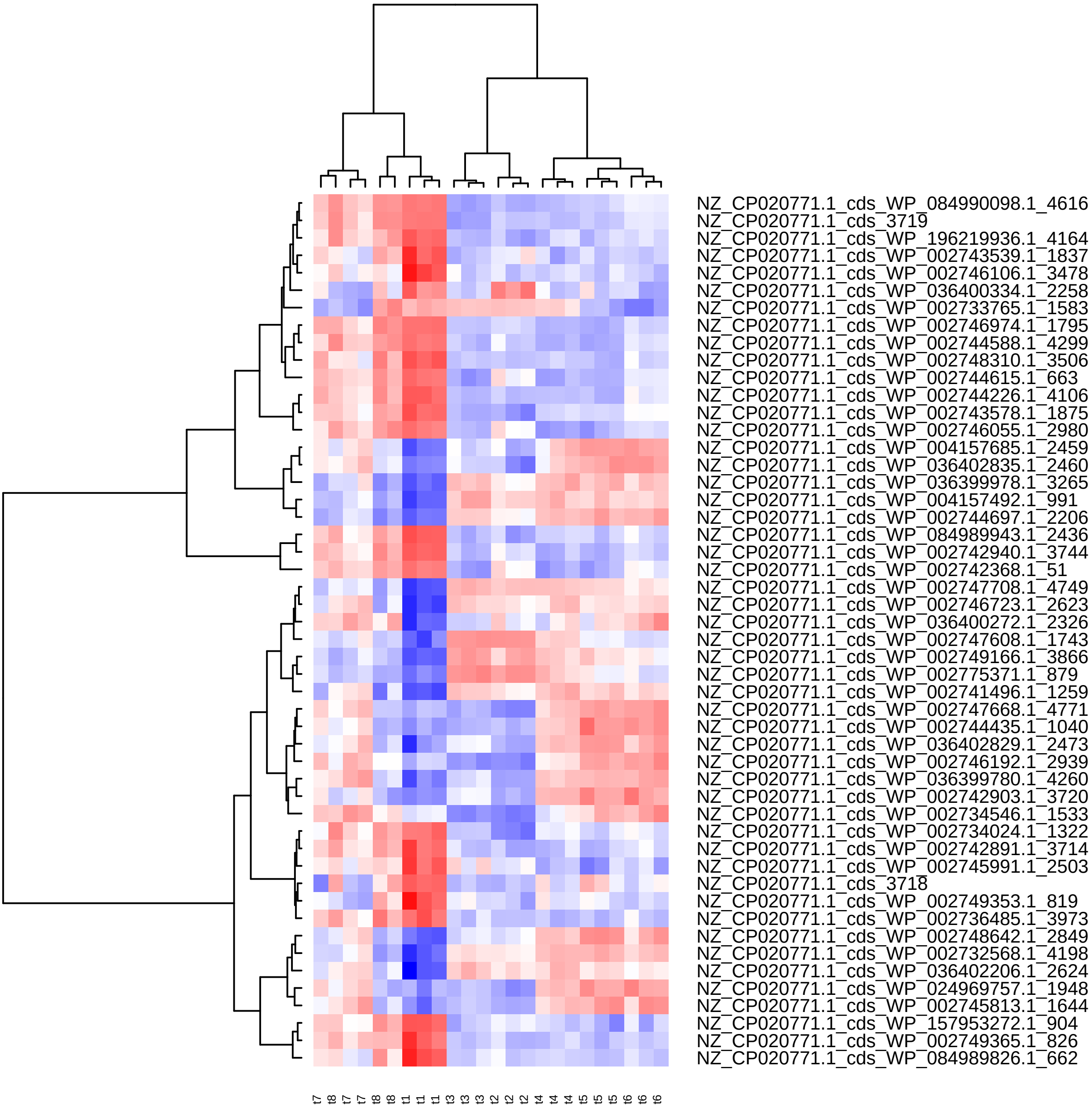

Supplement: Supplementary_Data [file 602159_file03.zip › Supplementary_DATA/DICOEXPRESS_output/DiffAnalysis/[t1-t7]/PCC7806_[t1-t7]_Top50_Clustering.pdf]

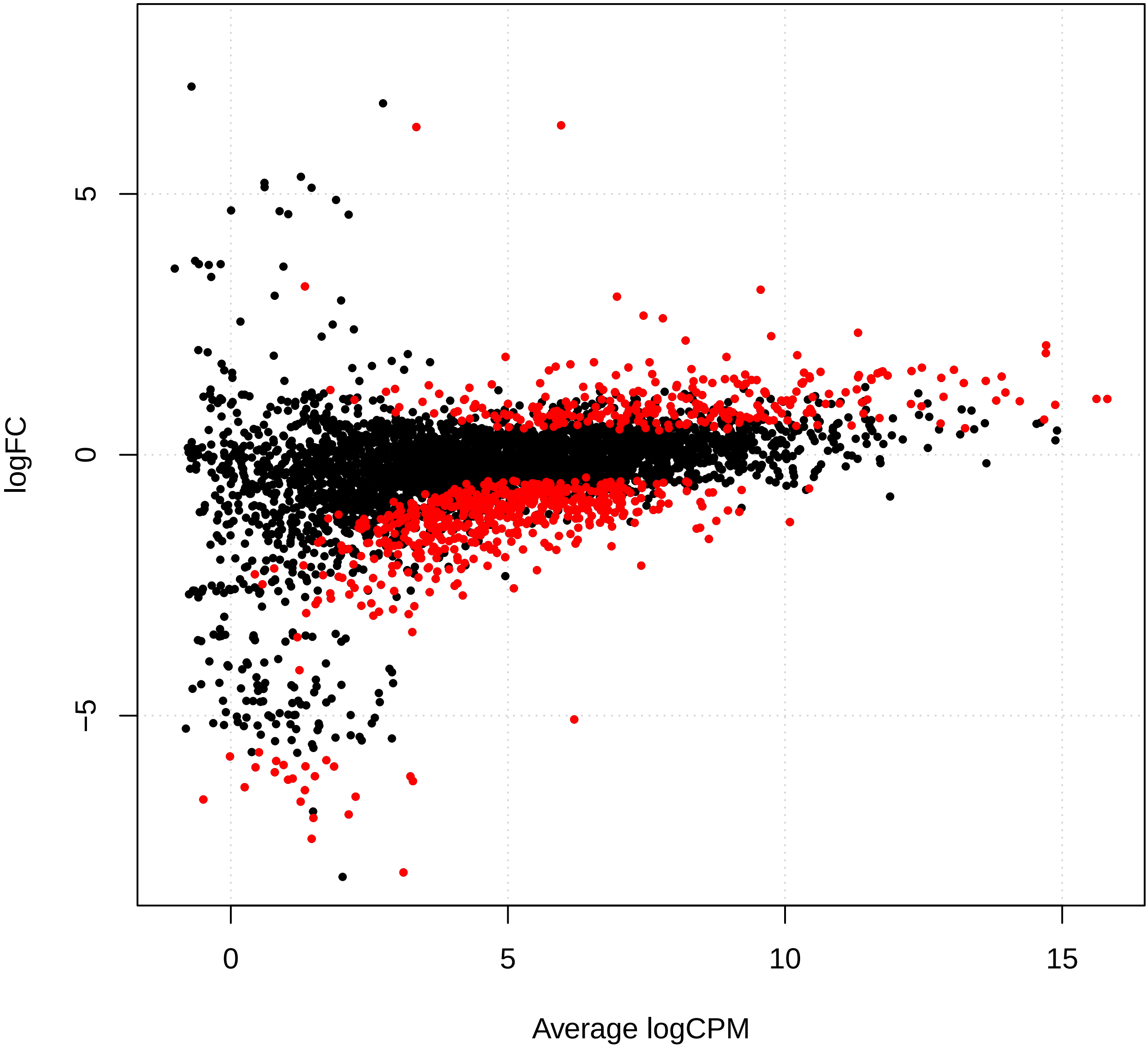

Supplement: Supplementary_Data [file 602159_file03.zip › Supplementary_DATA/DICOEXPRESS_output/DiffAnalysis/[t1-t8]/PCC7806_[t1-t8]_plotSmear.pdf]

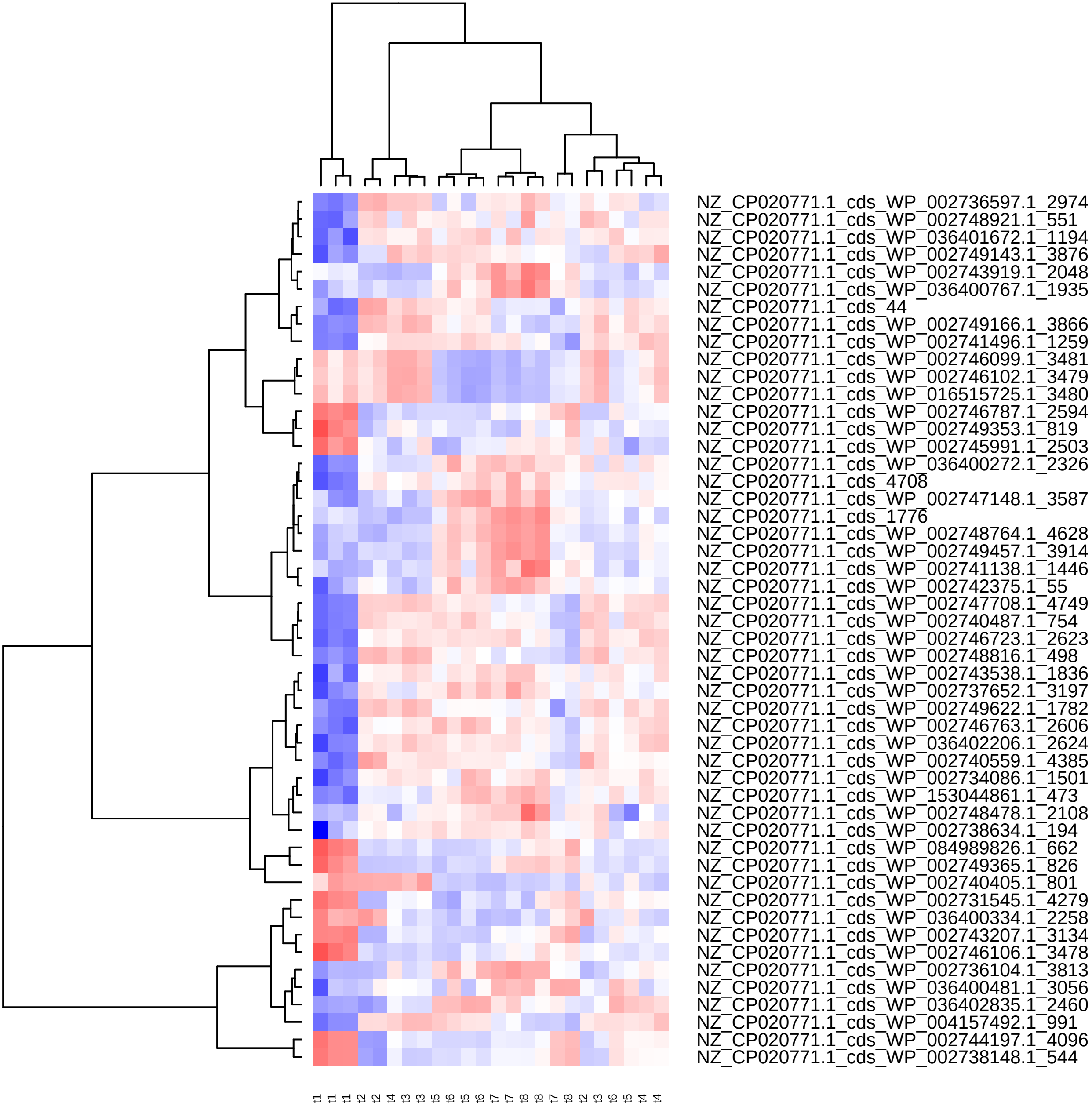

Supplement: Supplementary_Data [file 602159_file03.zip › Supplementary_DATA/DICOEXPRESS_output/DiffAnalysis/[t1-t8]/PCC7806_[t1-t8]_Top50_Clustering.pdf]

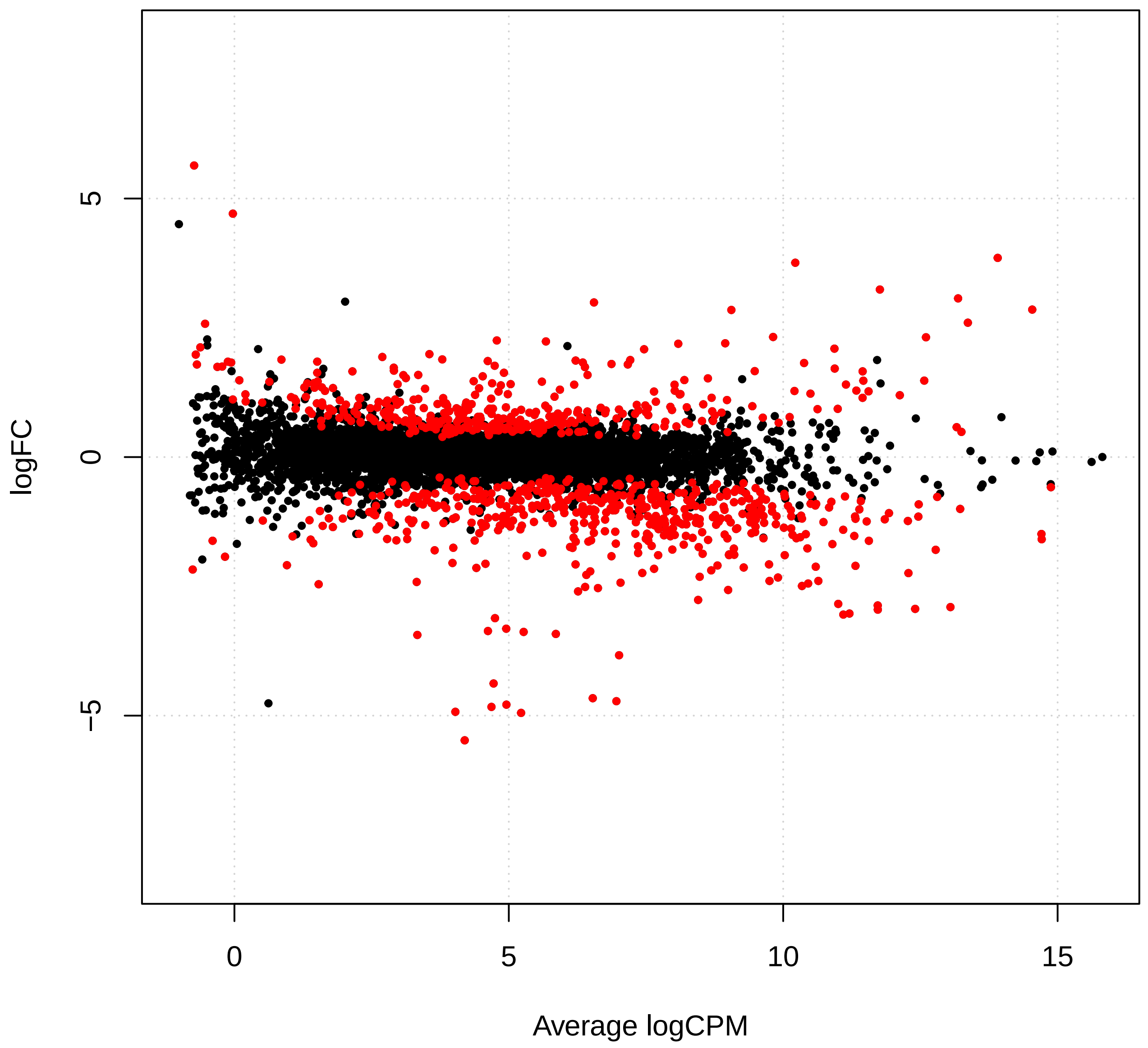

Supplement: Supplementary_Data [file 602159_file03.zip › Supplementary_DATA/DICOEXPRESS_output/DiffAnalysis/[t2-t3]/PCC7806_[t2-t3]_plotSmear.pdf]

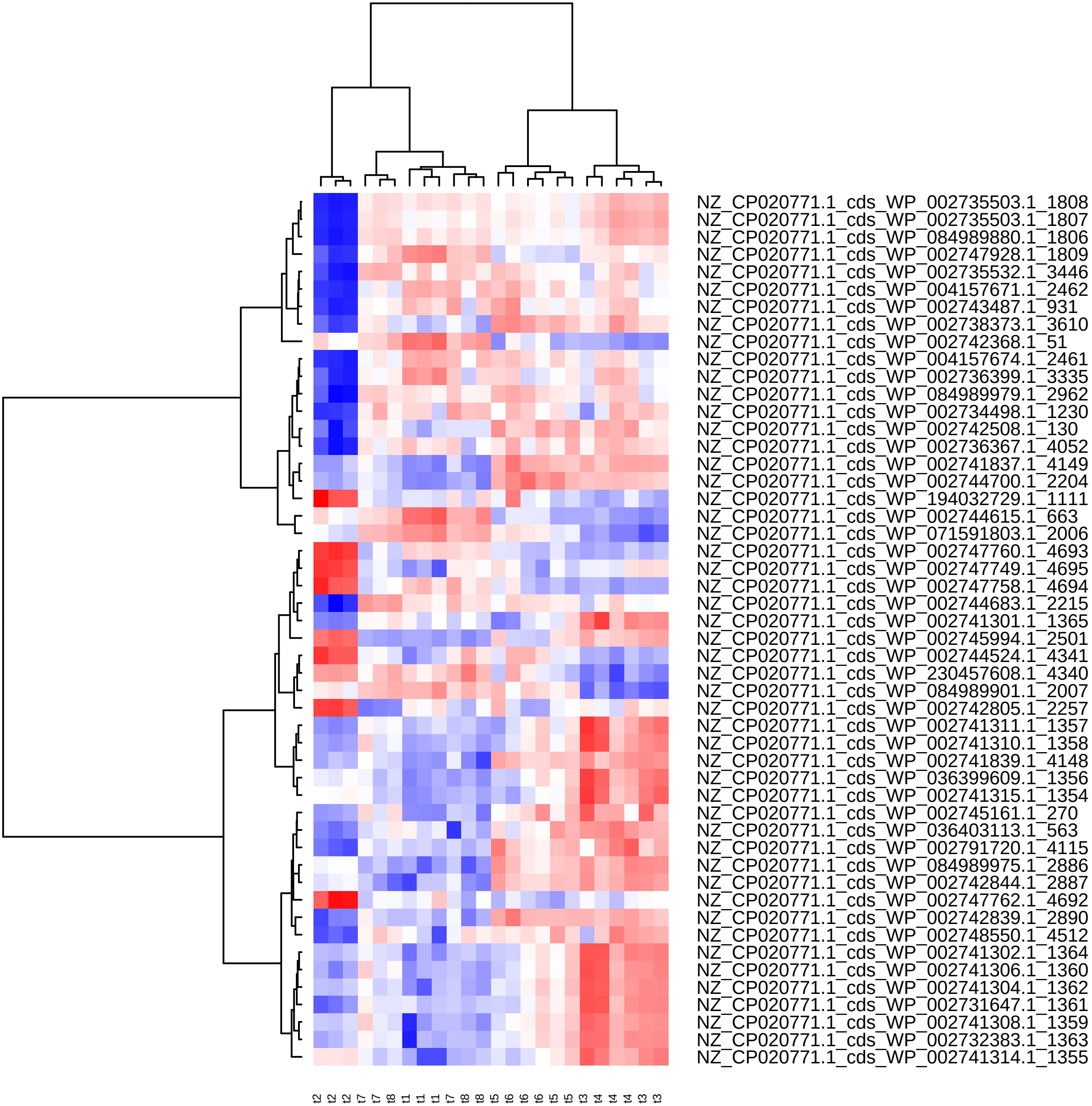

Supplement: Supplementary_Data [file 602159_file03.zip › Supplementary_DATA/DICOEXPRESS_output/DiffAnalysis/[t2-t3]/PCC7806_[t2-t3]_Top50_Clustering.pdf]

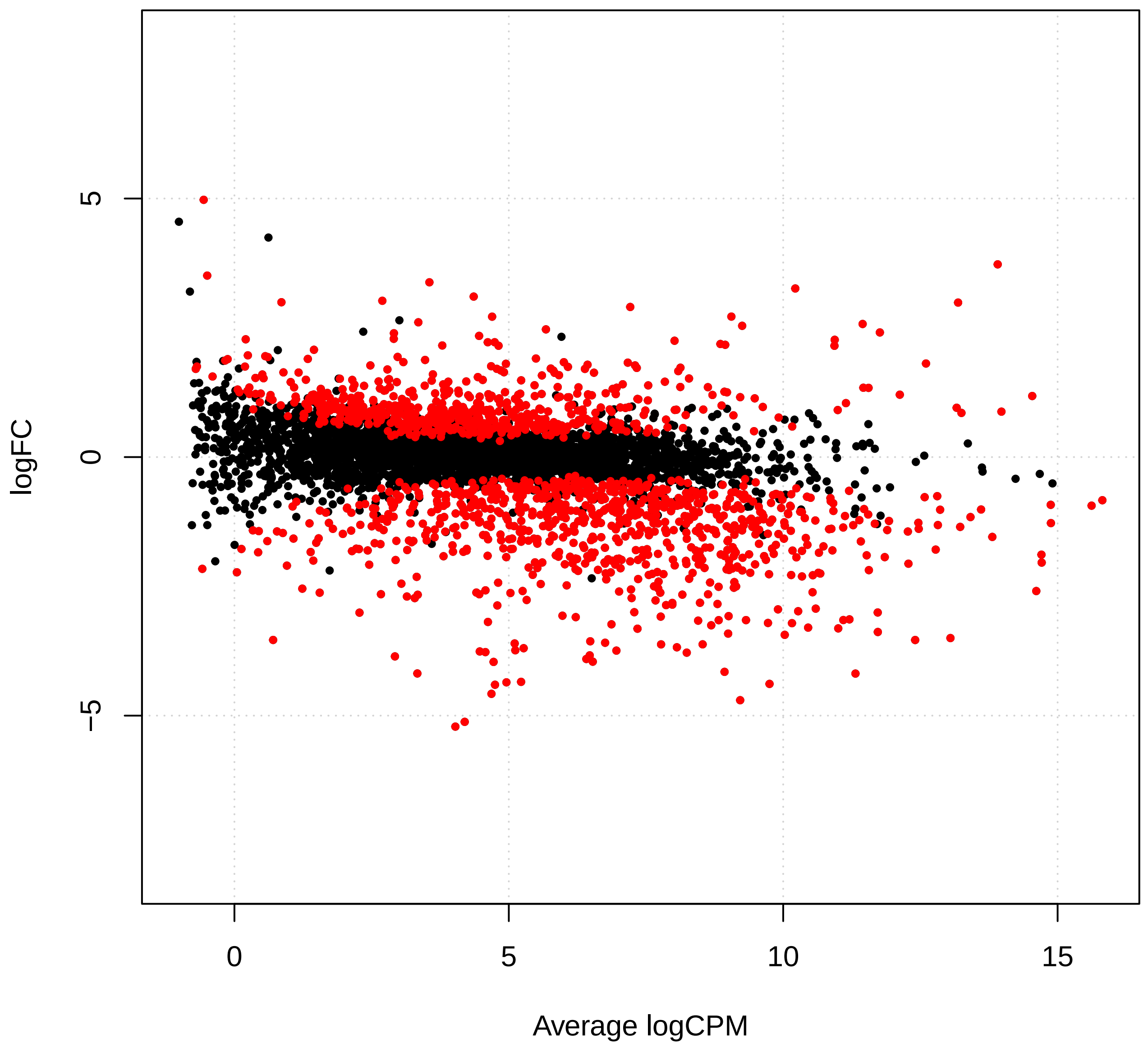

Supplement: Supplementary_Data [file 602159_file03.zip › Supplementary_DATA/DICOEXPRESS_output/DiffAnalysis/[t2-t4]/PCC7806_[t2-t4]_plotSmear.pdf]

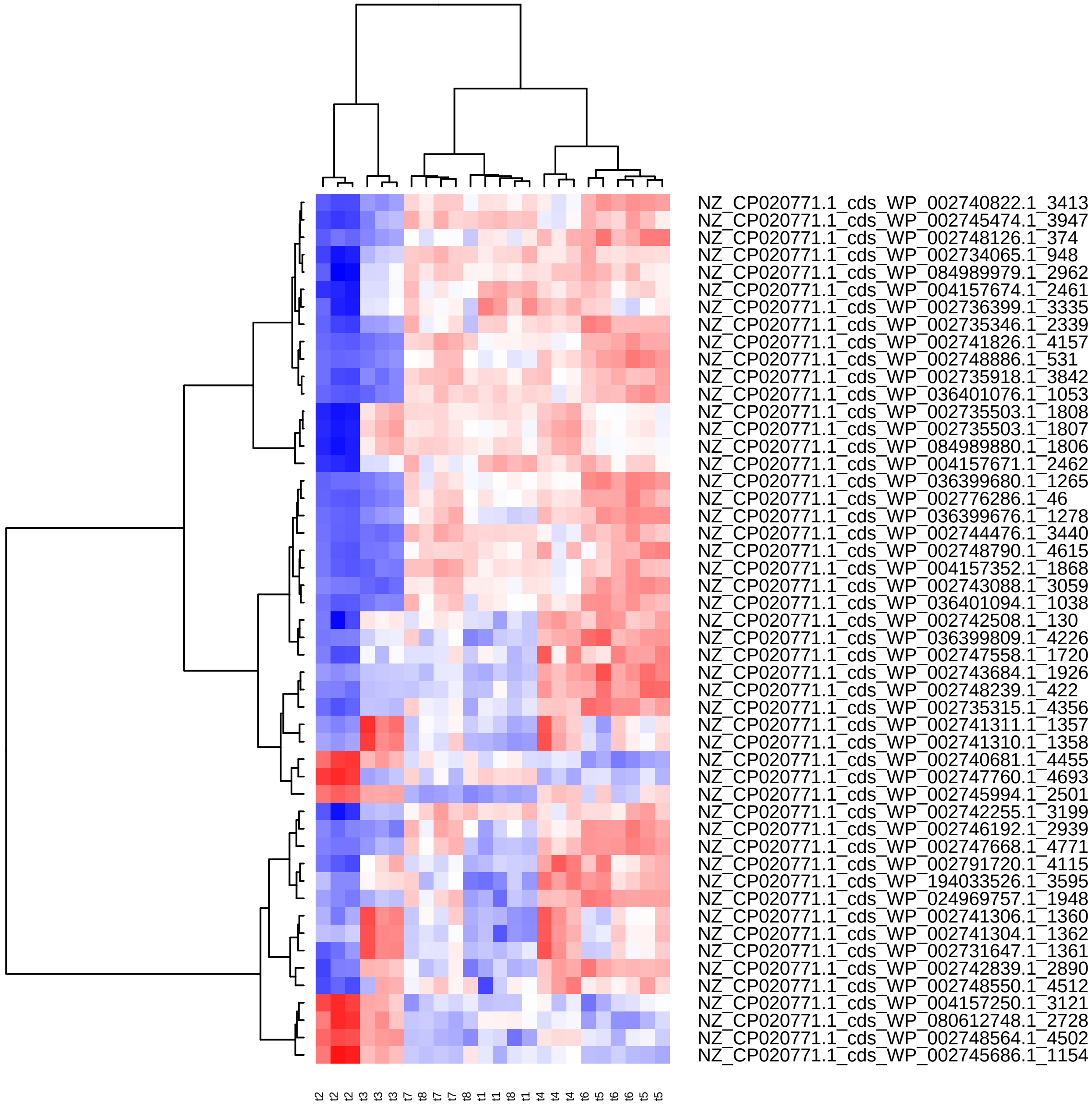

Supplement: Supplementary_Data [file 602159_file03.zip › Supplementary_DATA/DICOEXPRESS_output/DiffAnalysis/[t2-t4]/PCC7806_[t2-t4]_Top50_Clustering.pdf]

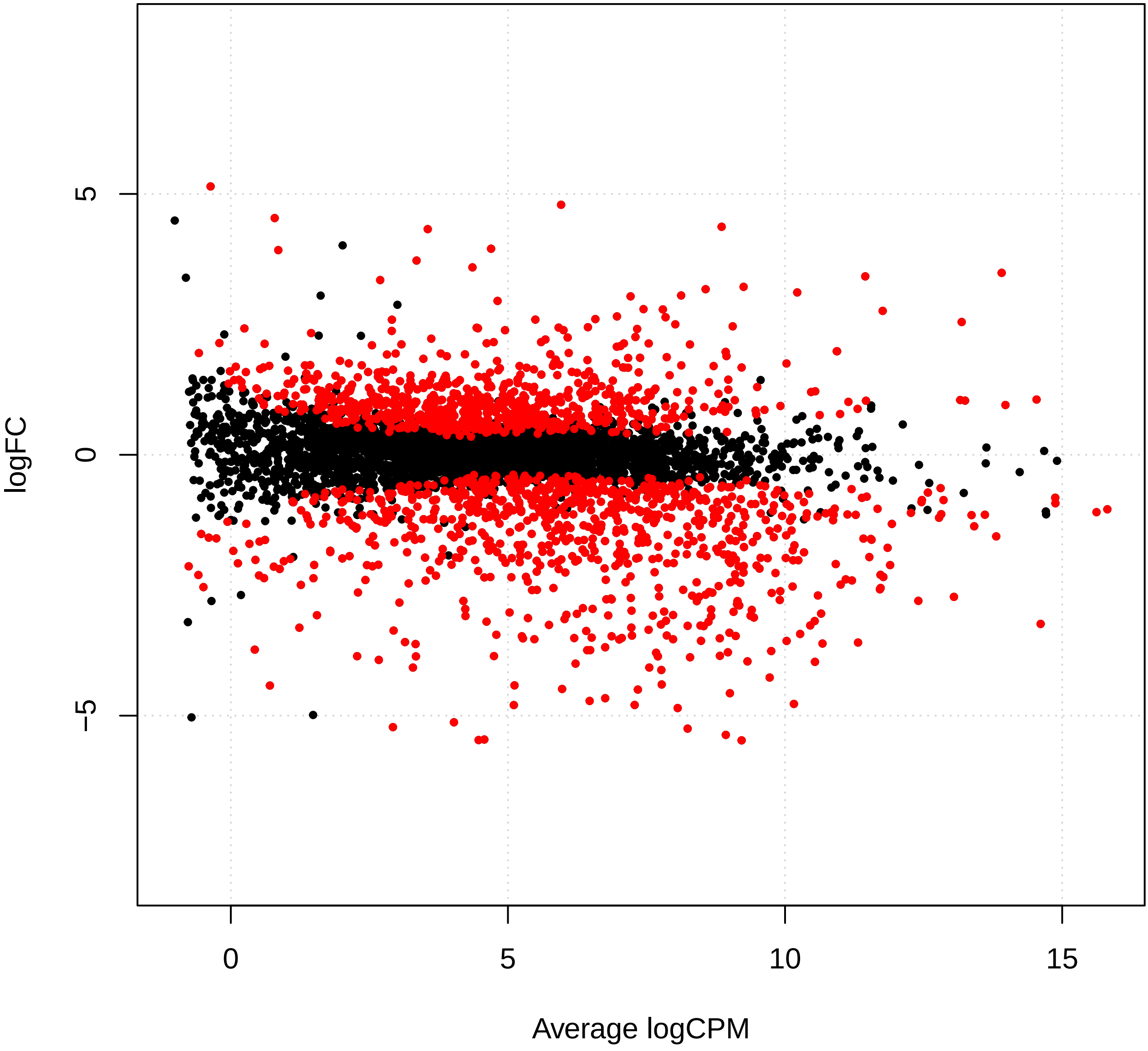

Supplement: Supplementary_Data [file 602159_file03.zip › Supplementary_DATA/DICOEXPRESS_output/DiffAnalysis/[t2-t5]/PCC7806_[t2-t5]_plotSmear.pdf]

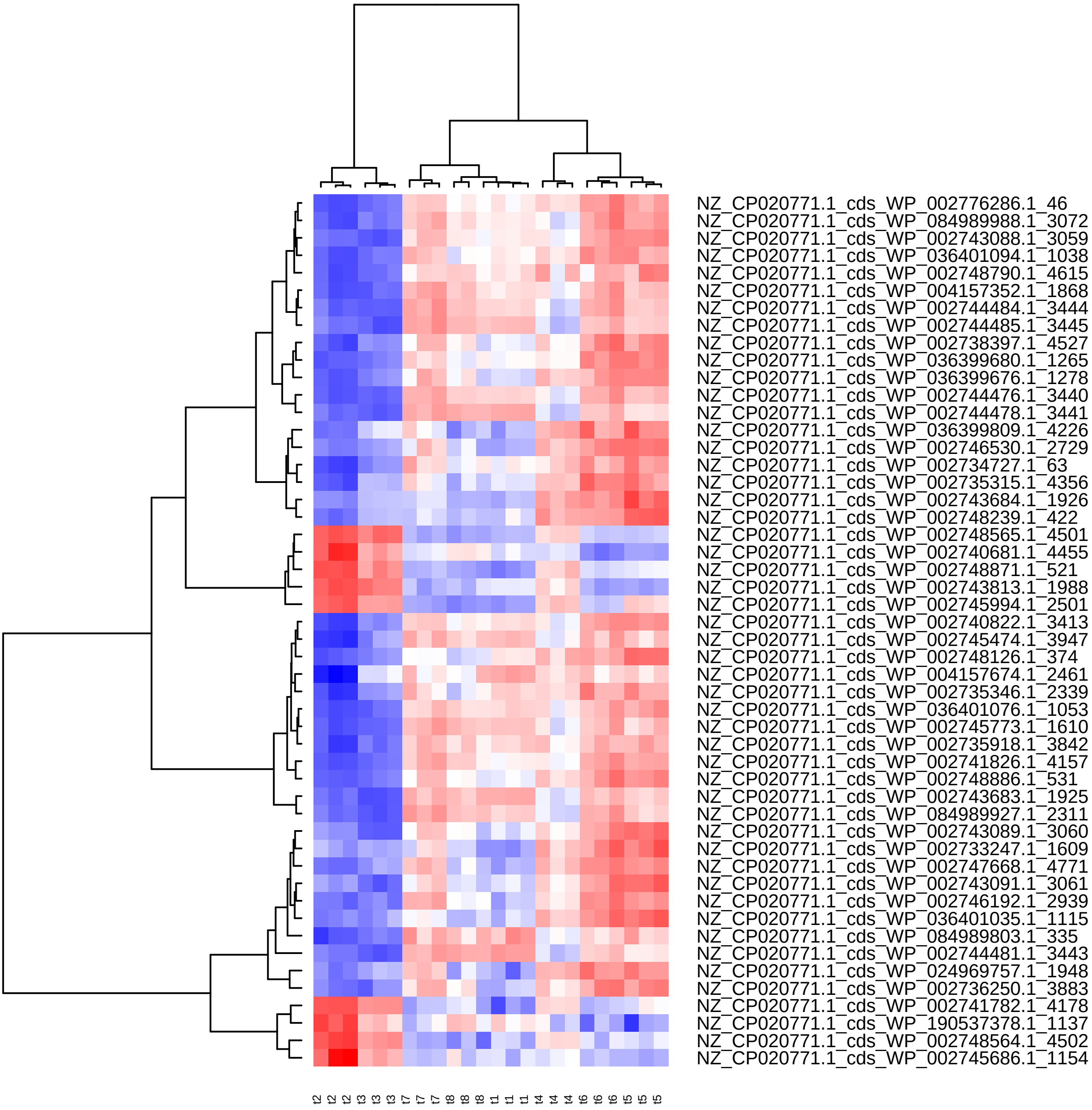

Supplement: Supplementary_Data [file 602159_file03.zip › Supplementary_DATA/DICOEXPRESS_output/DiffAnalysis/[t2-t5]/PCC7806_[t2-t5]_Top50_Clustering.pdf]

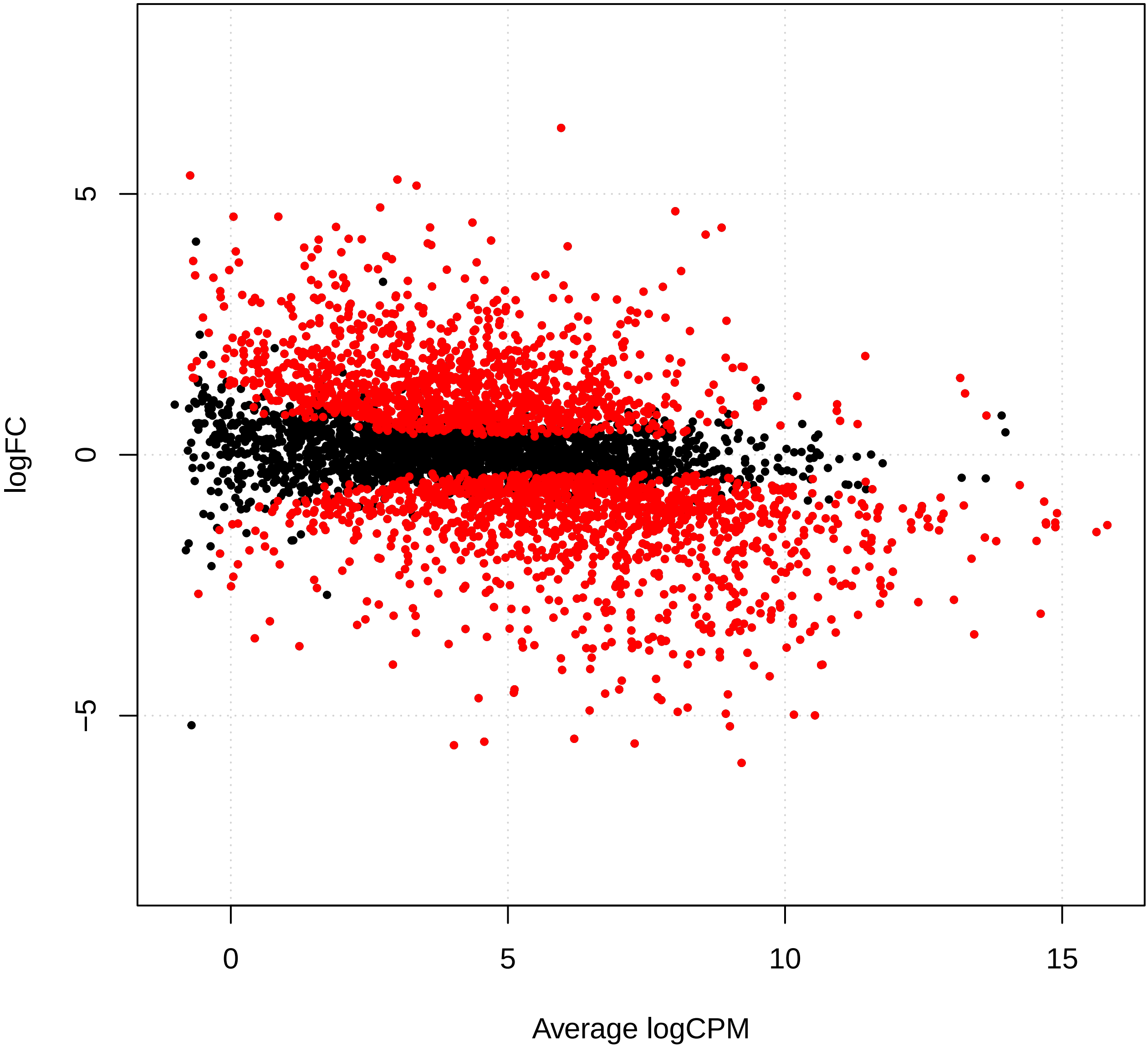

Supplement: Supplementary_Data [file 602159_file03.zip › Supplementary_DATA/DICOEXPRESS_output/DiffAnalysis/[t2-t6]/PCC7806_[t2-t6]_plotSmear.pdf]

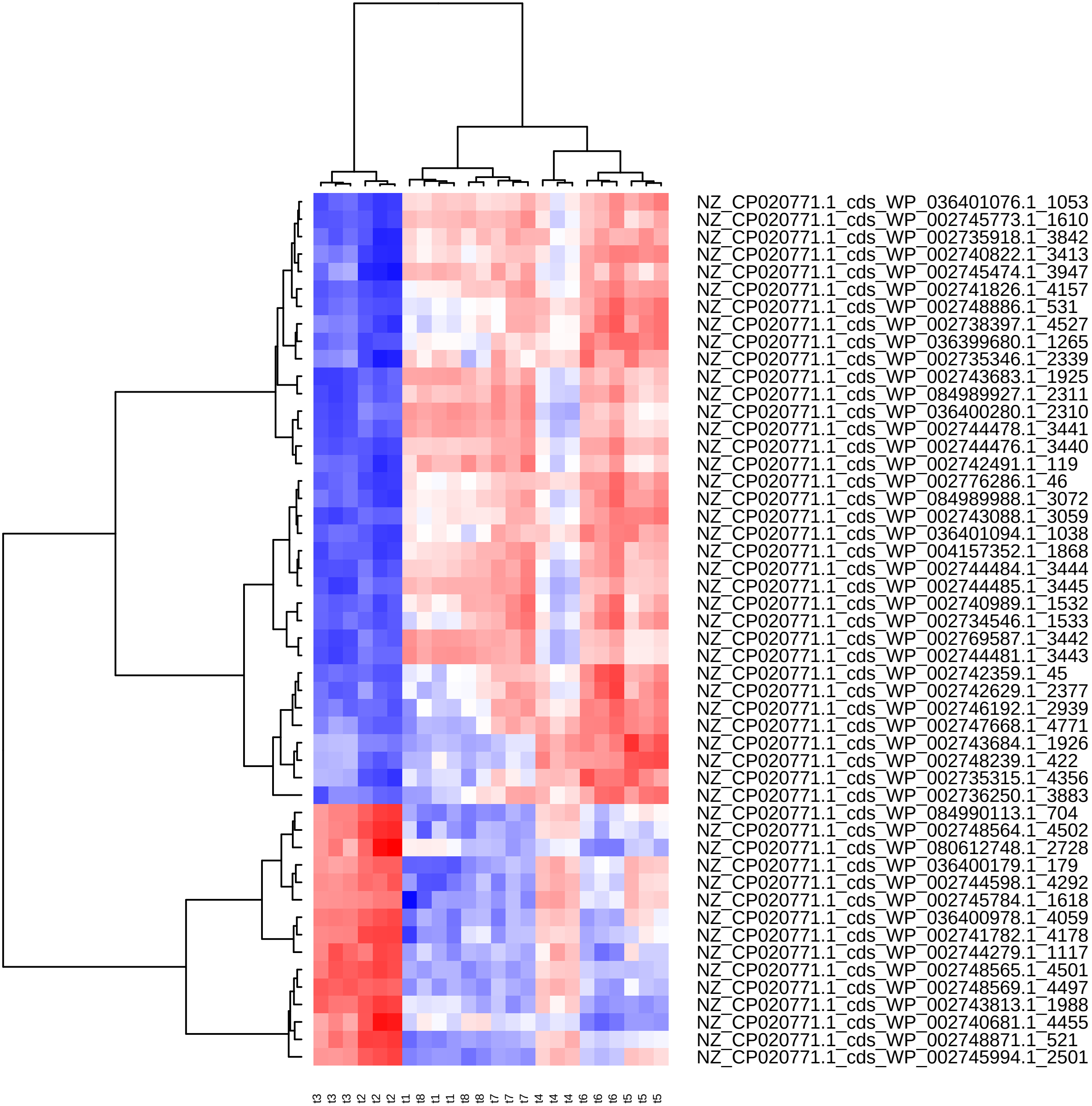

Supplement: Supplementary_Data [file 602159_file03.zip › Supplementary_DATA/DICOEXPRESS_output/DiffAnalysis/[t2-t6]/PCC7806_[t2-t6]_Top50_Clustering.pdf]

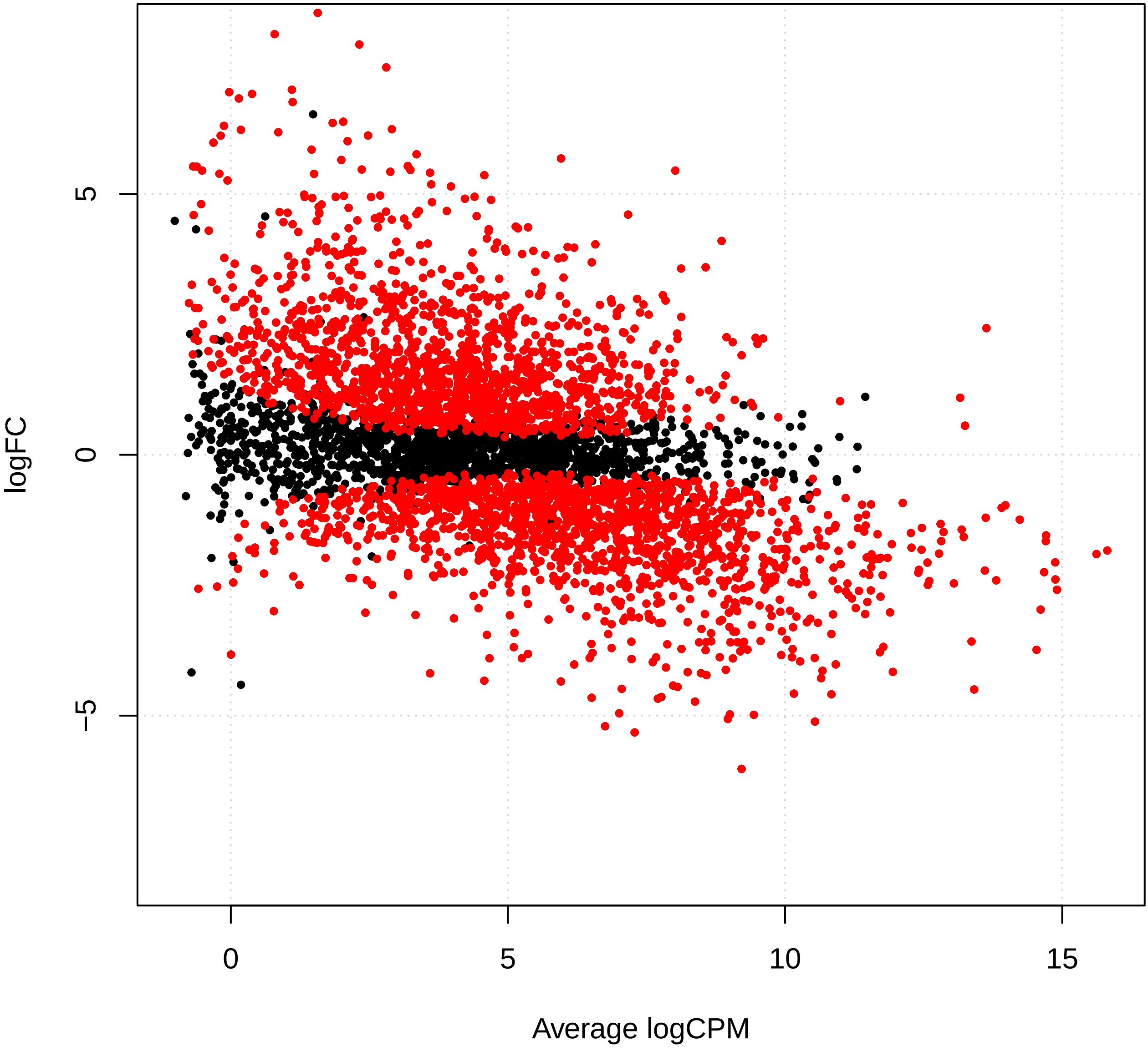

Supplement: Supplementary_Data [file 602159_file03.zip › Supplementary_DATA/DICOEXPRESS_output/DiffAnalysis/[t2-t7]/PCC7806_[t2-t7]_plotSmear.pdf]

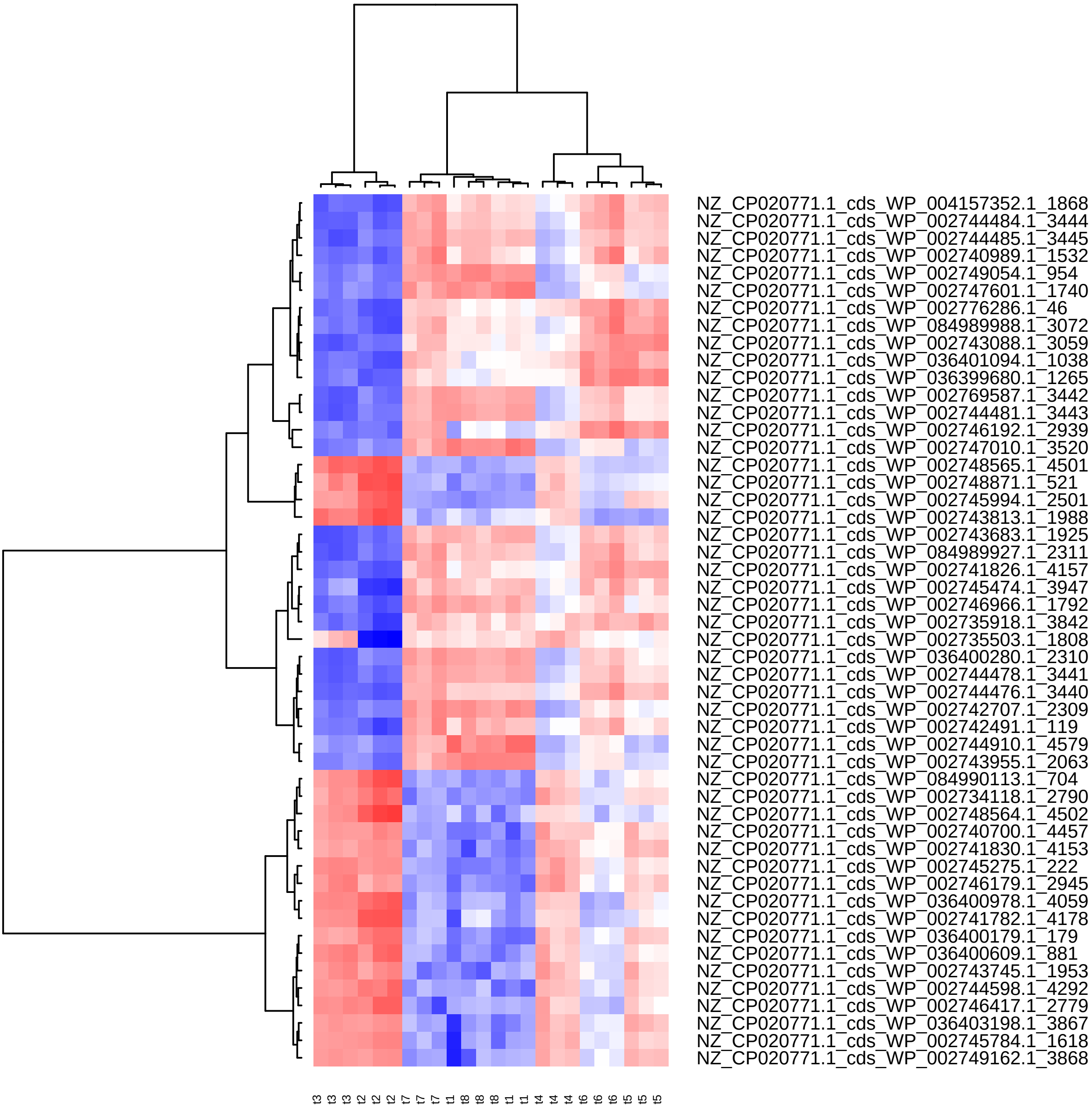

Supplement: Supplementary_Data [file 602159_file03.zip › Supplementary_DATA/DICOEXPRESS_output/DiffAnalysis/[t2-t7]/PCC7806_[t2-t7]_Top50_Clustering.pdf]

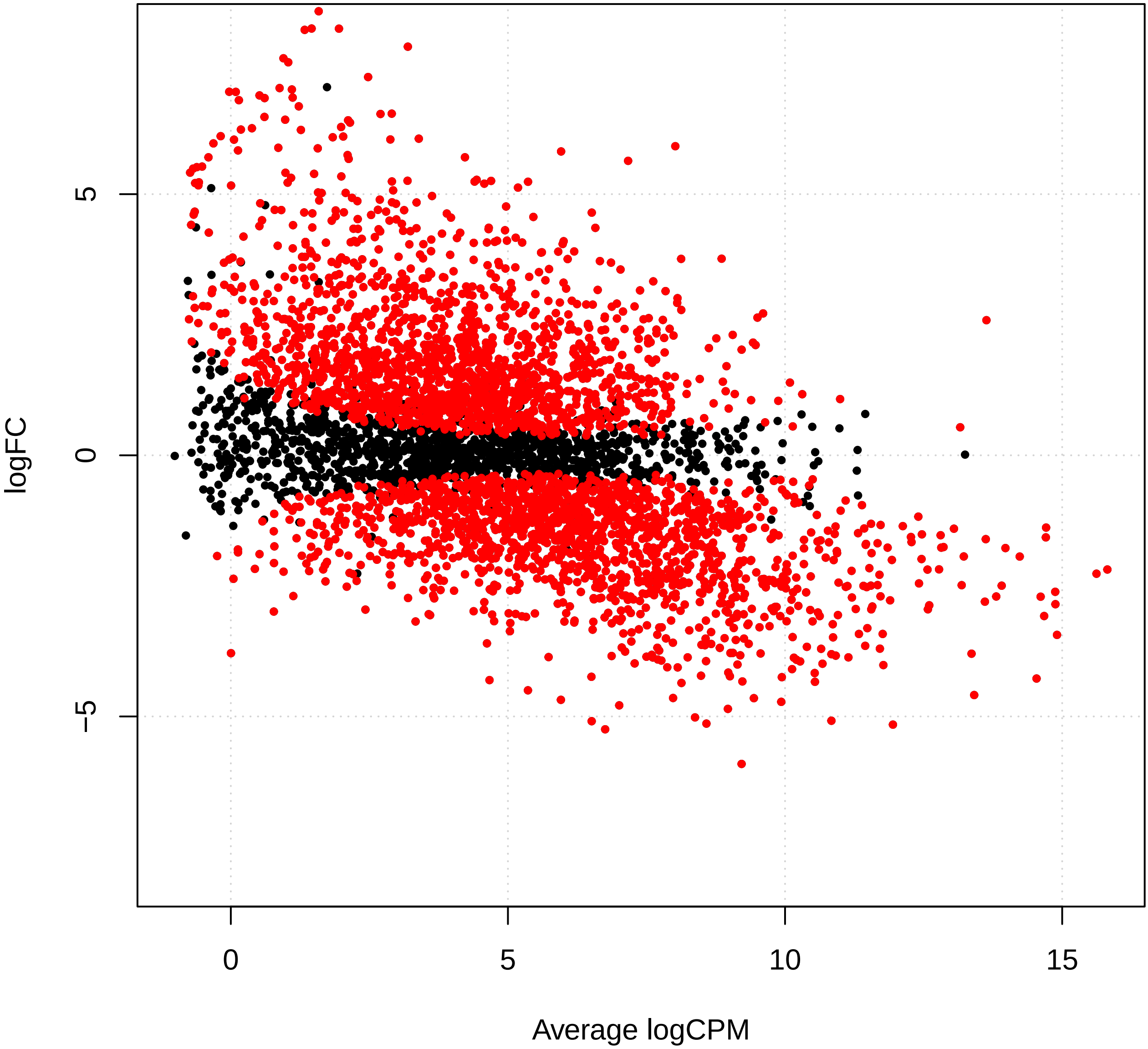

Supplement: Supplementary_Data [file 602159_file03.zip › Supplementary_DATA/DICOEXPRESS_output/DiffAnalysis/[t2-t8]/PCC7806_[t2-t8]_plotSmear.pdf]

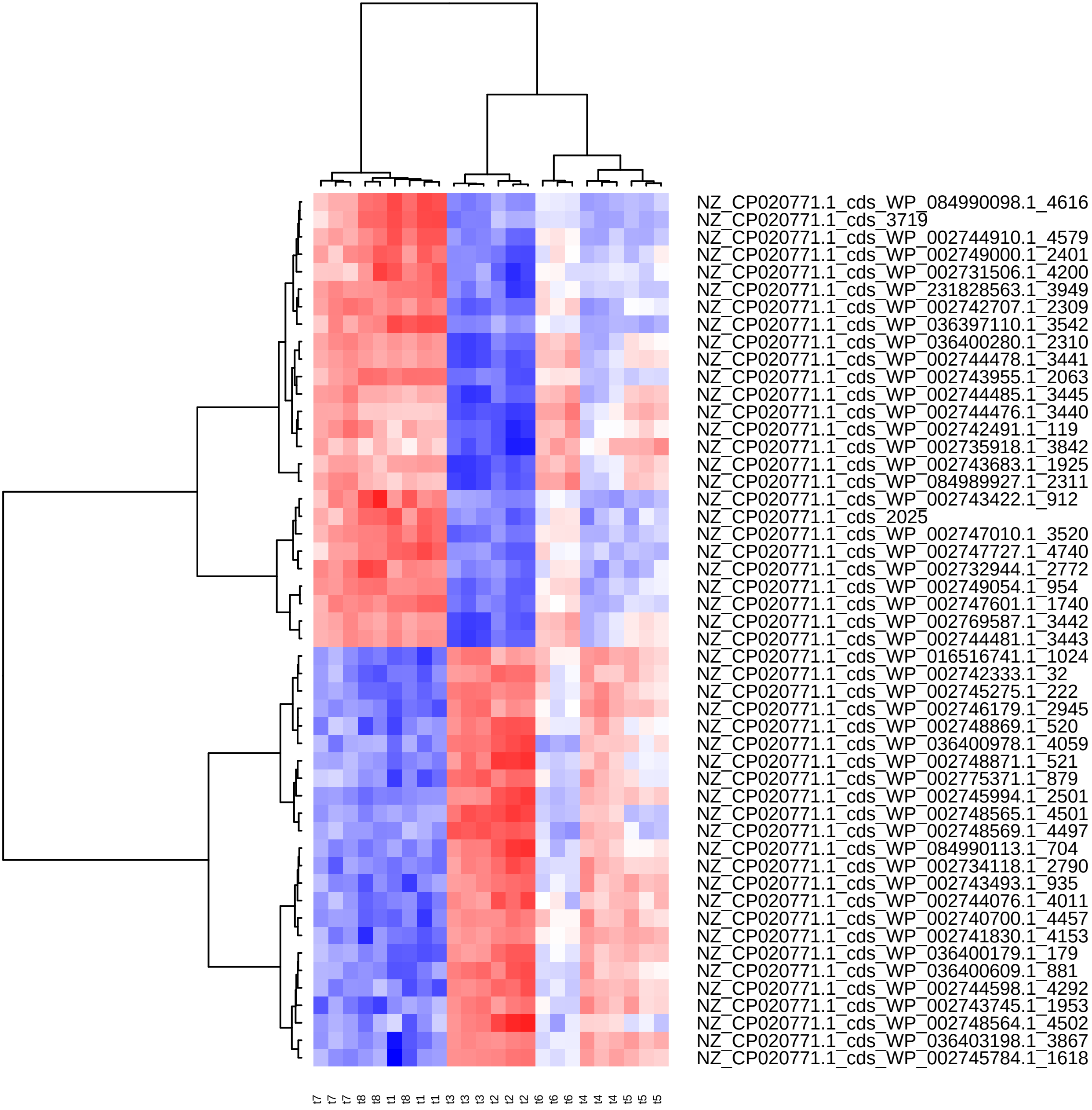

Supplement: Supplementary_Data [file 602159_file03.zip › Supplementary_DATA/DICOEXPRESS_output/DiffAnalysis/[t2-t8]/PCC7806_[t2-t8]_Top50_Clustering.pdf]

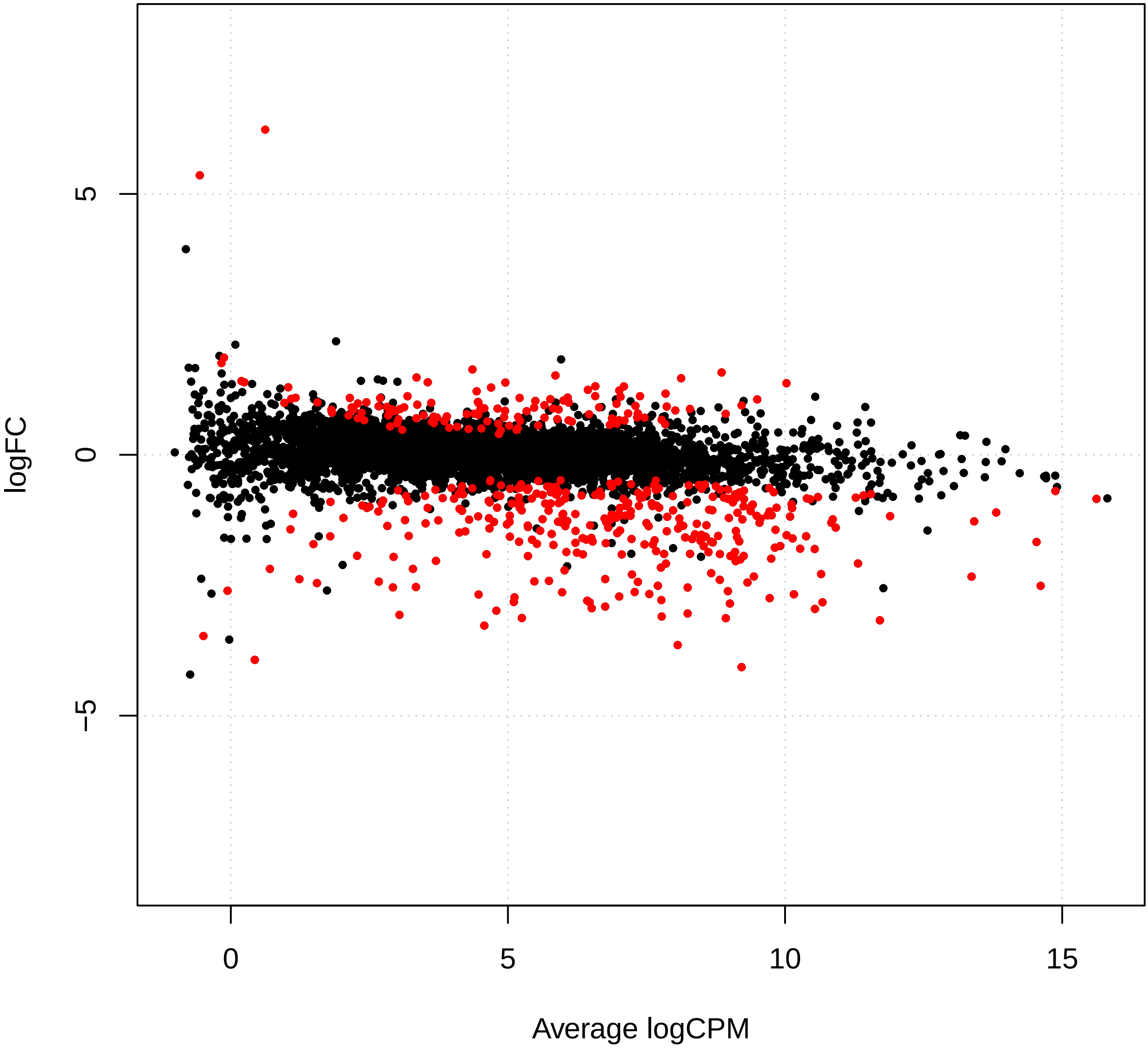

Supplement: Supplementary_Data [file 602159_file03.zip › Supplementary_DATA/DICOEXPRESS_output/DiffAnalysis/[t3-t4]/PCC7806_[t3-t4]_plotSmear.pdf]

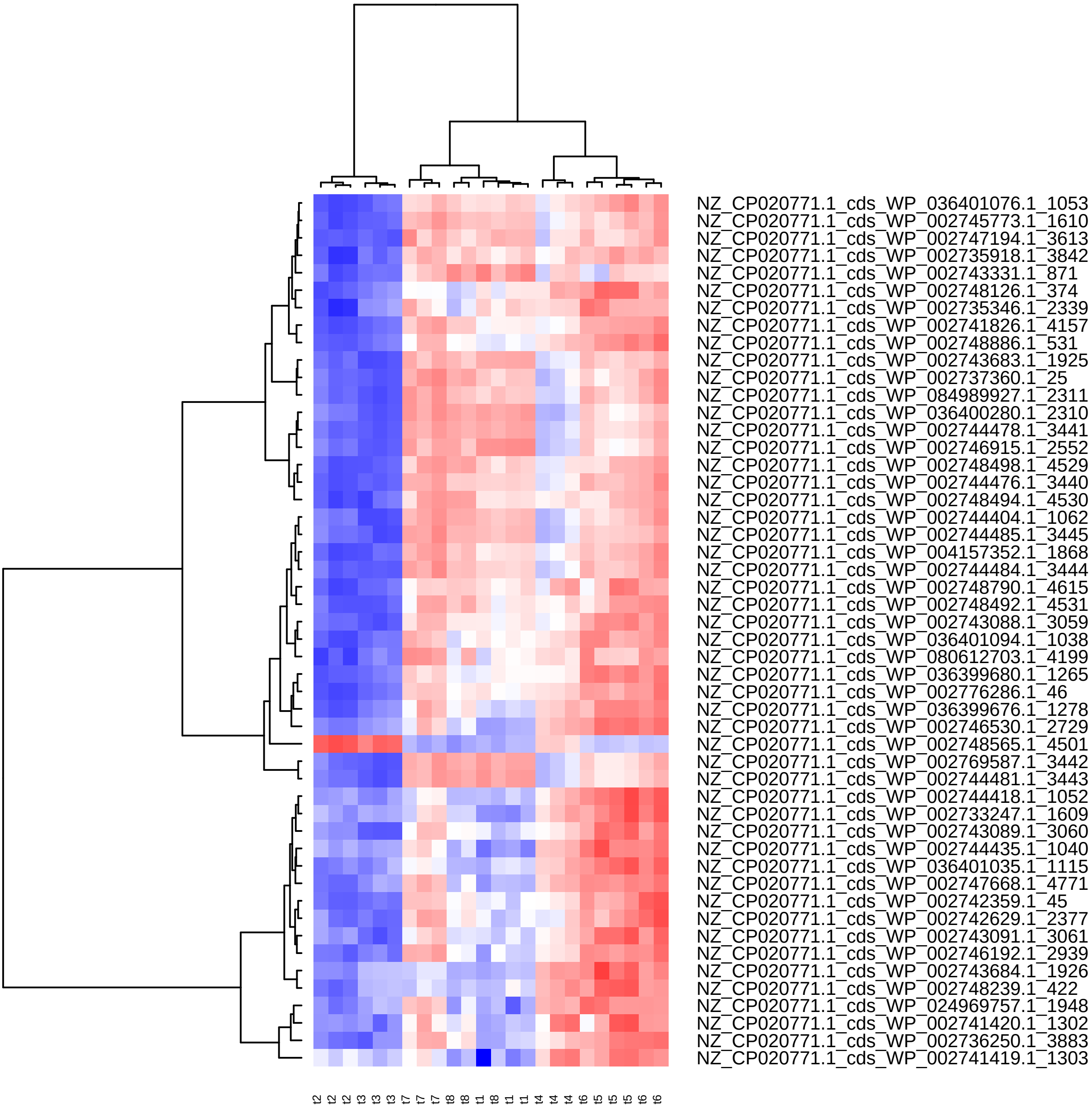

Supplement: Supplementary_Data [file 602159_file03.zip › Supplementary_DATA/DICOEXPRESS_output/DiffAnalysis/[t3-t4]/PCC7806_[t3-t4]_Top50_Clustering.pdf]

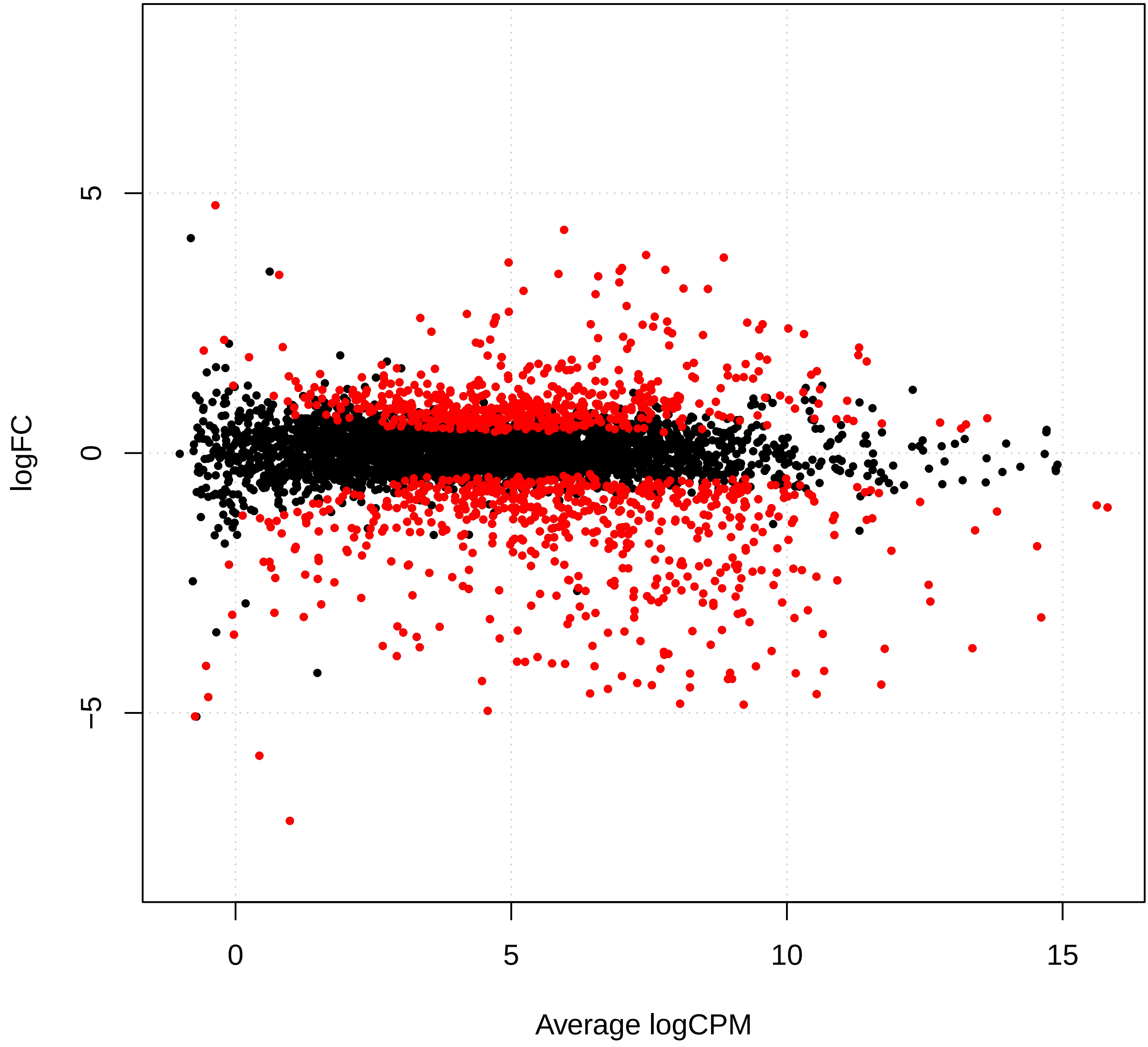

Supplement: Supplementary_Data [file 602159_file03.zip › Supplementary_DATA/DICOEXPRESS_output/DiffAnalysis/[t3-t5]/PCC7806_[t3-t5]_plotSmear.pdf]

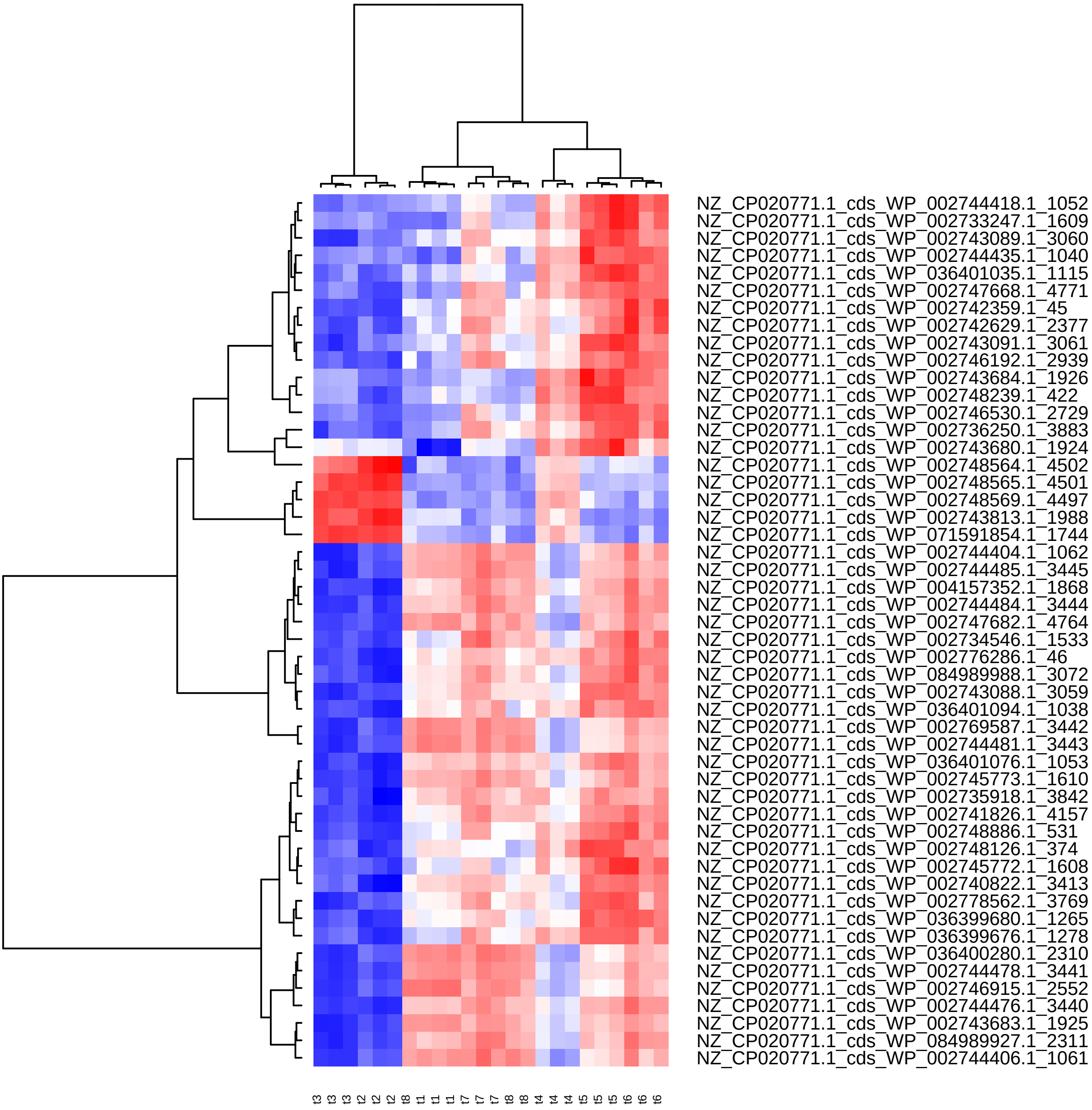

Supplement: Supplementary_Data [file 602159_file03.zip › Supplementary_DATA/DICOEXPRESS_output/DiffAnalysis/[t3-t5]/PCC7806_[t3-t5]_Top50_Clustering.pdf]

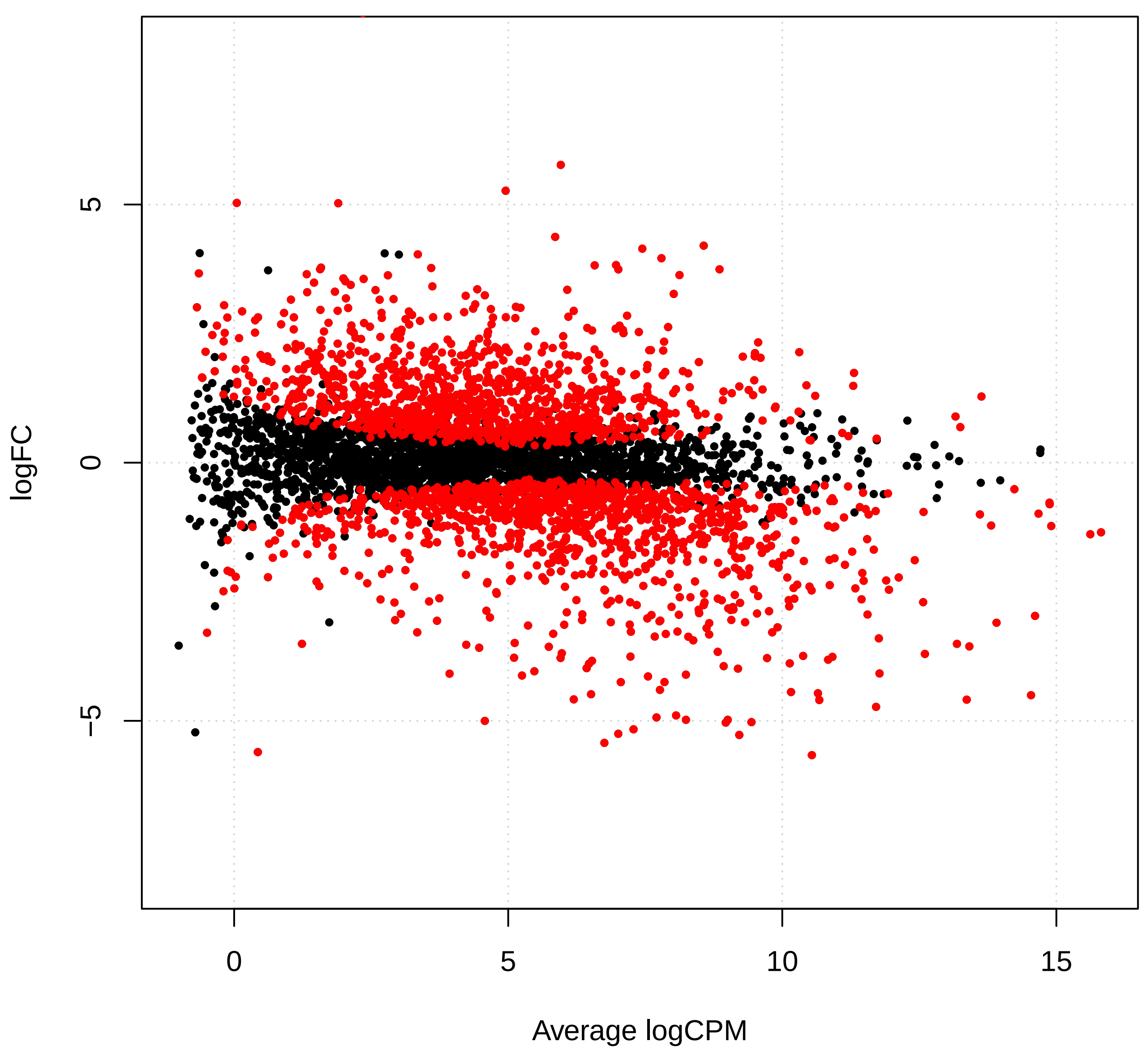

Supplement: Supplementary_Data [file 602159_file03.zip › Supplementary_DATA/DICOEXPRESS_output/DiffAnalysis/[t3-t6]/PCC7806_[t3-t6]_plotSmear.pdf]

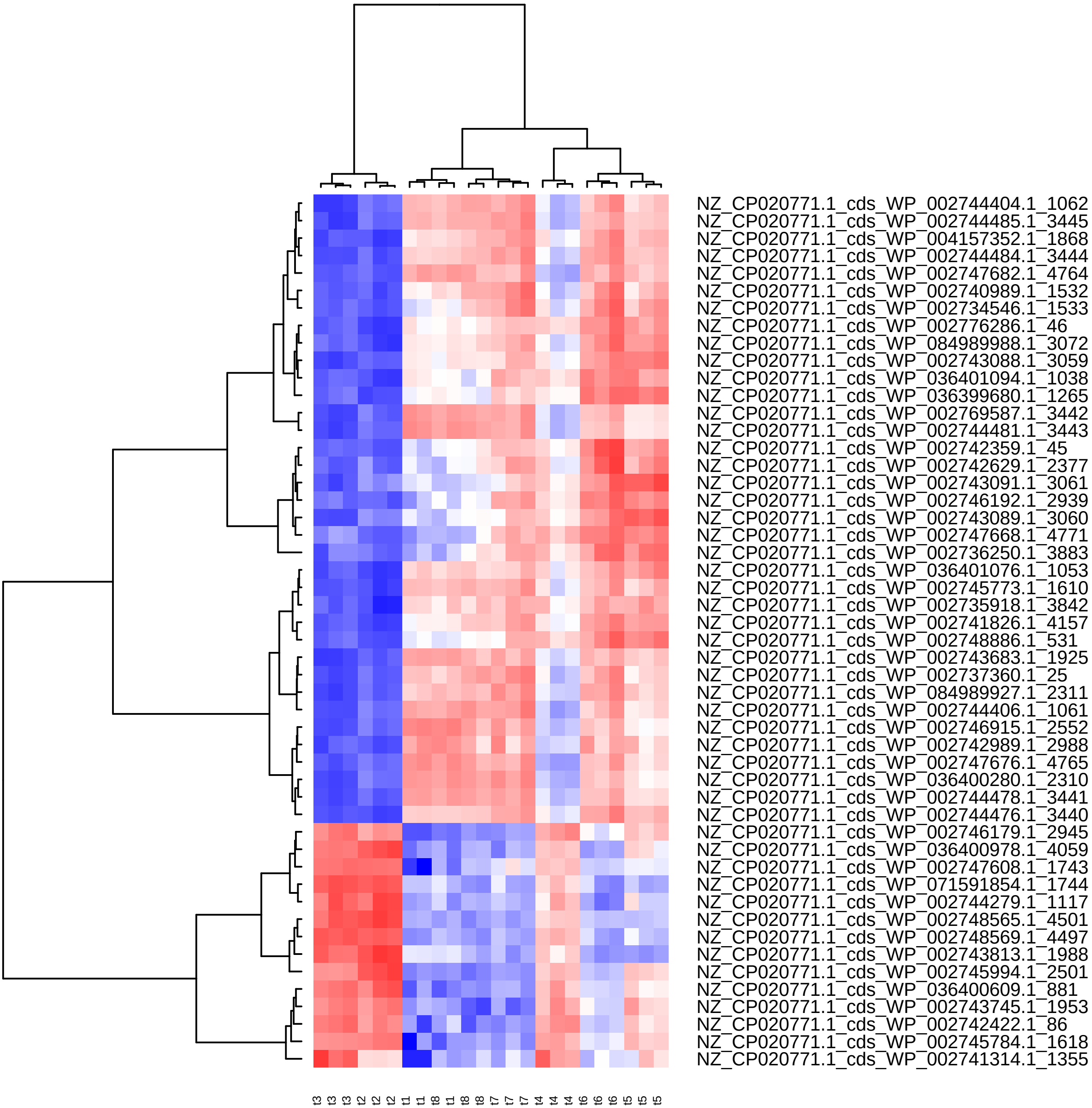

Supplement: Supplementary_Data [file 602159_file03.zip › Supplementary_DATA/DICOEXPRESS_output/DiffAnalysis/[t3-t6]/PCC7806_[t3-t6]_Top50_Clustering.pdf]

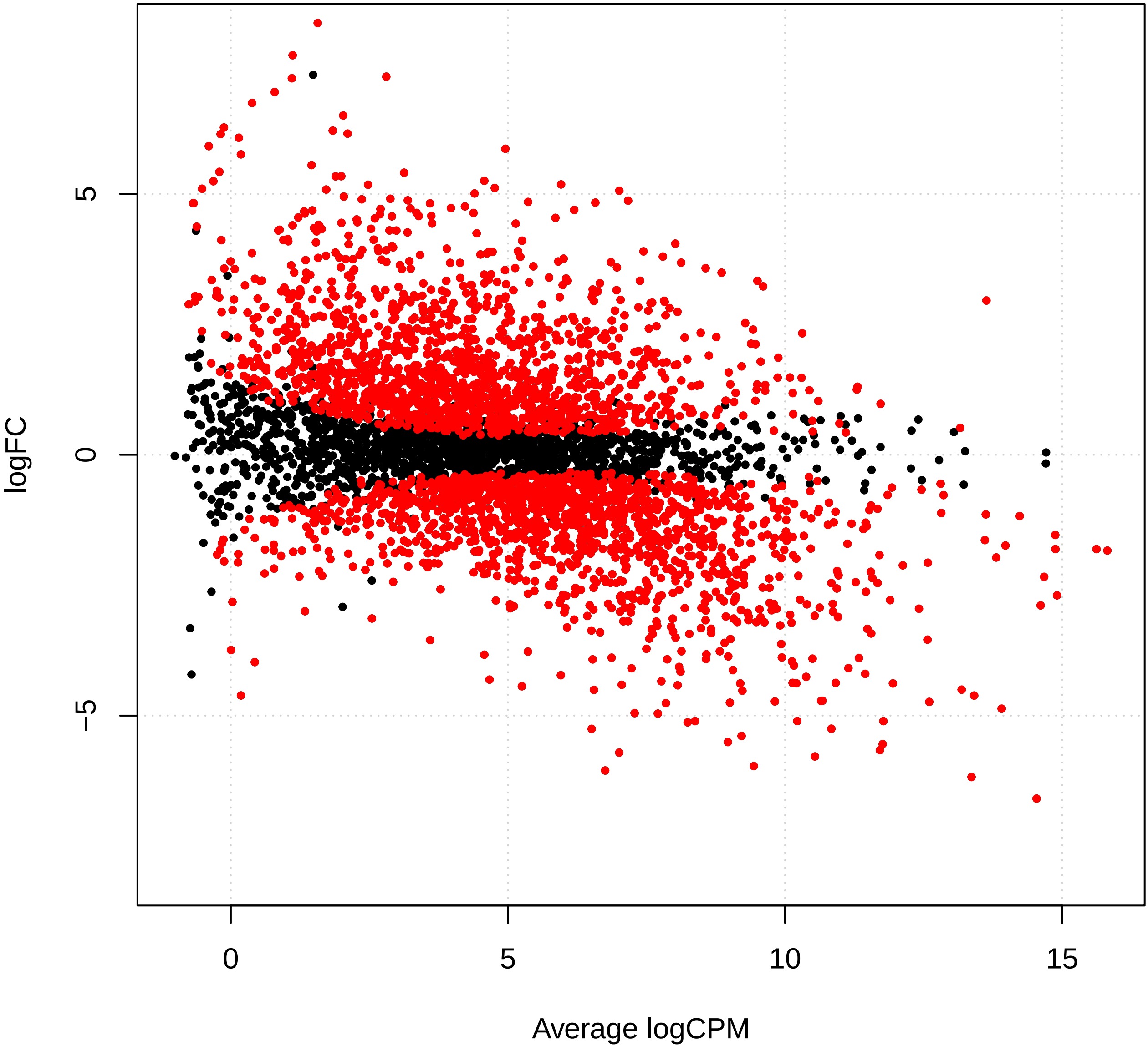

Supplement: Supplementary_Data [file 602159_file03.zip › Supplementary_DATA/DICOEXPRESS_output/DiffAnalysis/[t3-t7]/PCC7806_[t3-t7]_plotSmear.pdf]

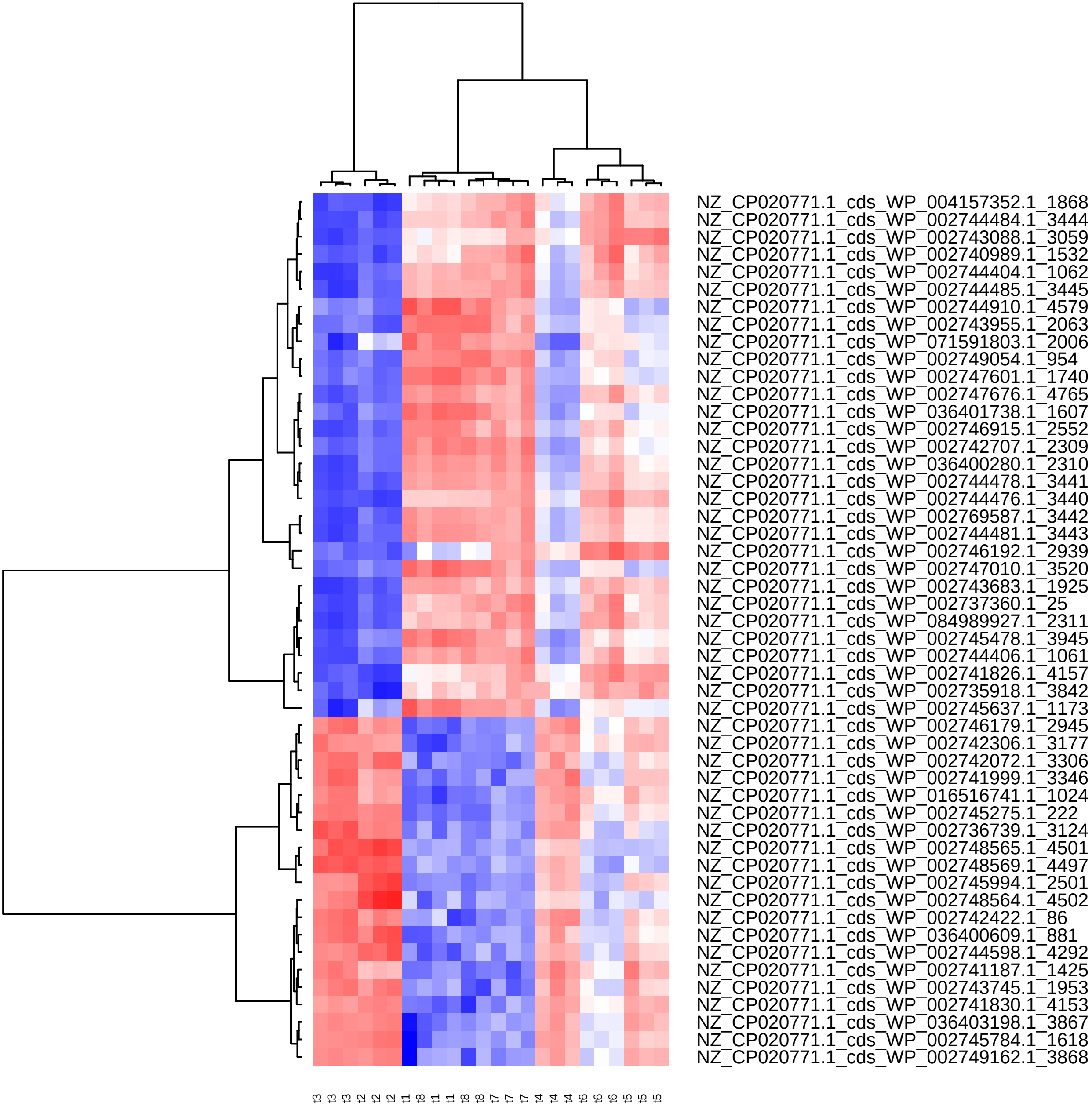

Supplement: Supplementary_Data [file 602159_file03.zip › Supplementary_DATA/DICOEXPRESS_output/DiffAnalysis/[t3-t7]/PCC7806_[t3-t7]_Top50_Clustering.pdf]

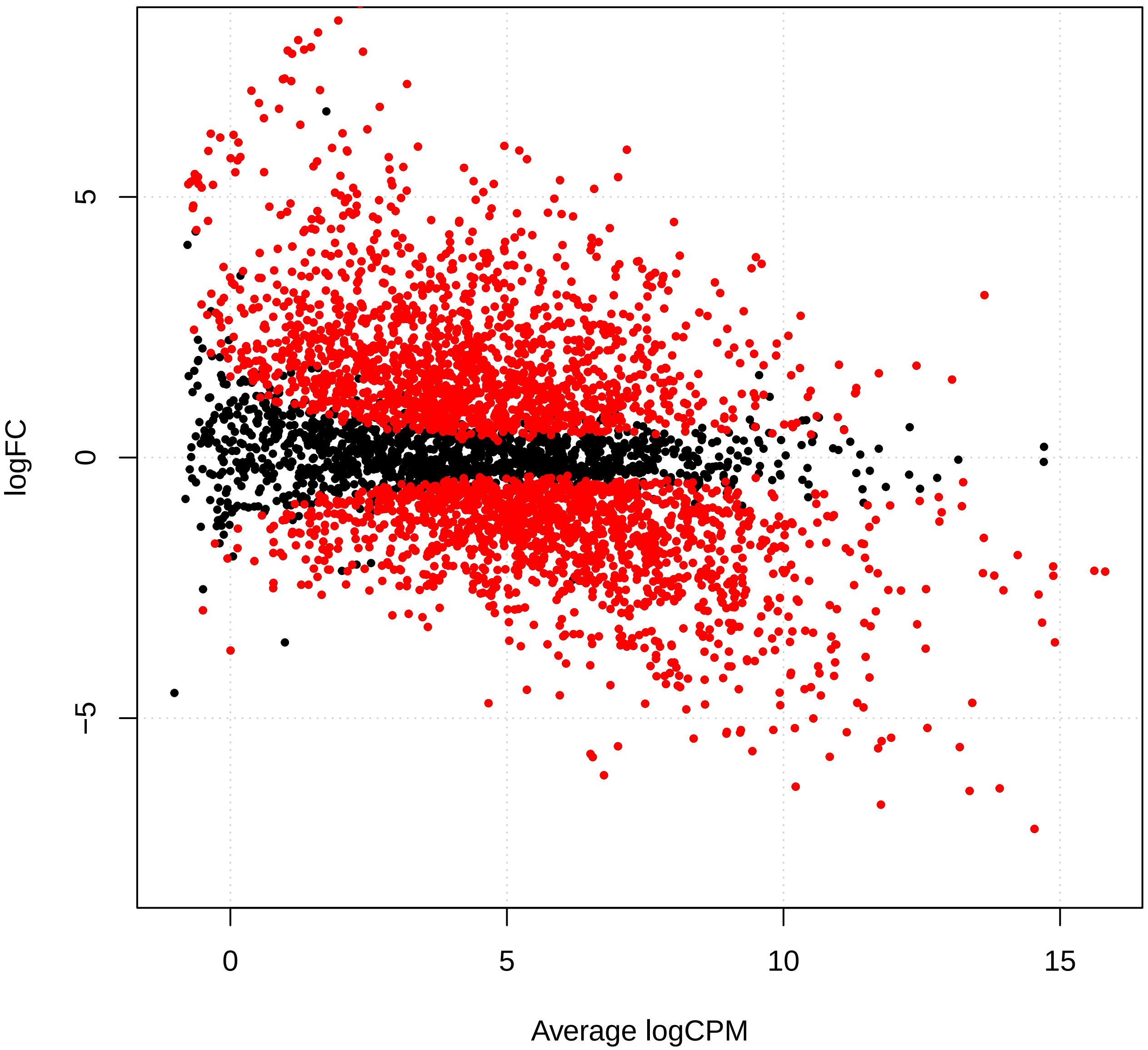

Supplement: Supplementary_Data [file 602159_file03.zip › Supplementary_DATA/DICOEXPRESS_output/DiffAnalysis/[t3-t8]/PCC7806_[t3-t8]_plotSmear.pdf]

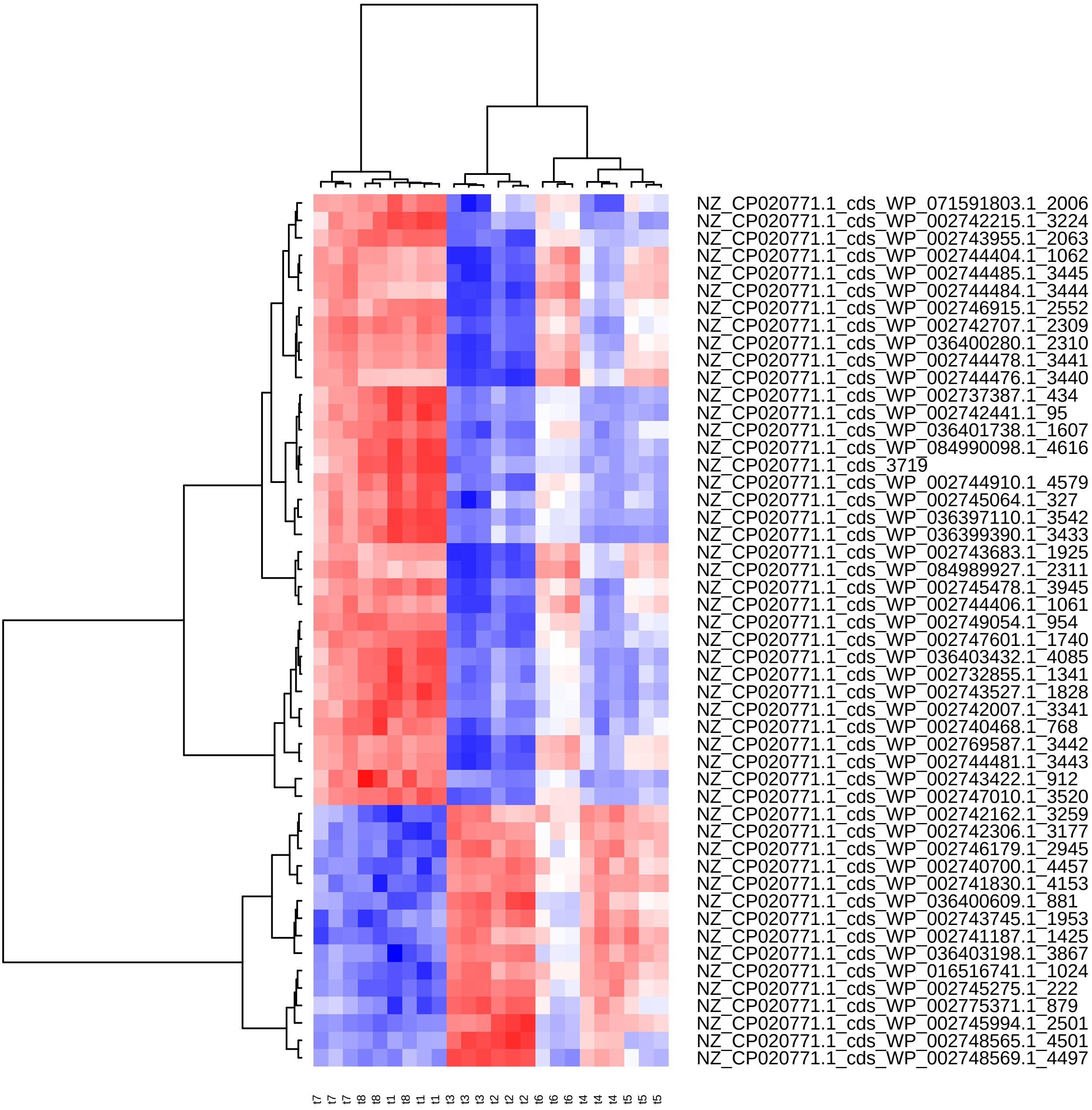

Supplement: Supplementary_Data [file 602159_file03.zip › Supplementary_DATA/DICOEXPRESS_output/DiffAnalysis/[t3-t8]/PCC7806_[t3-t8]_Top50_Clustering.pdf]

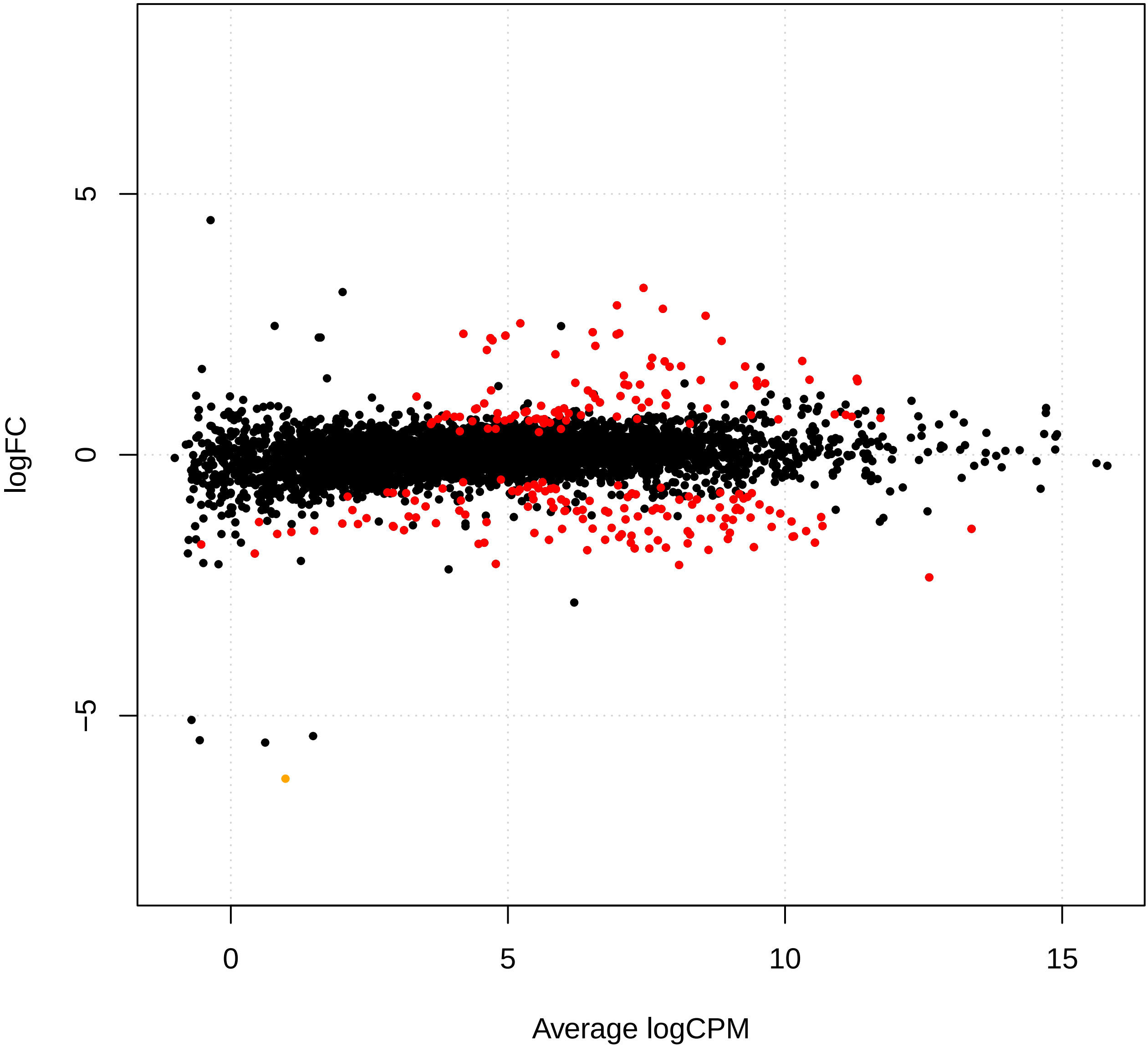

Supplement: Supplementary_Data [file 602159_file03.zip › Supplementary_DATA/DICOEXPRESS_output/DiffAnalysis/[t4-t5]/PCC7806_[t4-t5]_plotSmear.pdf]

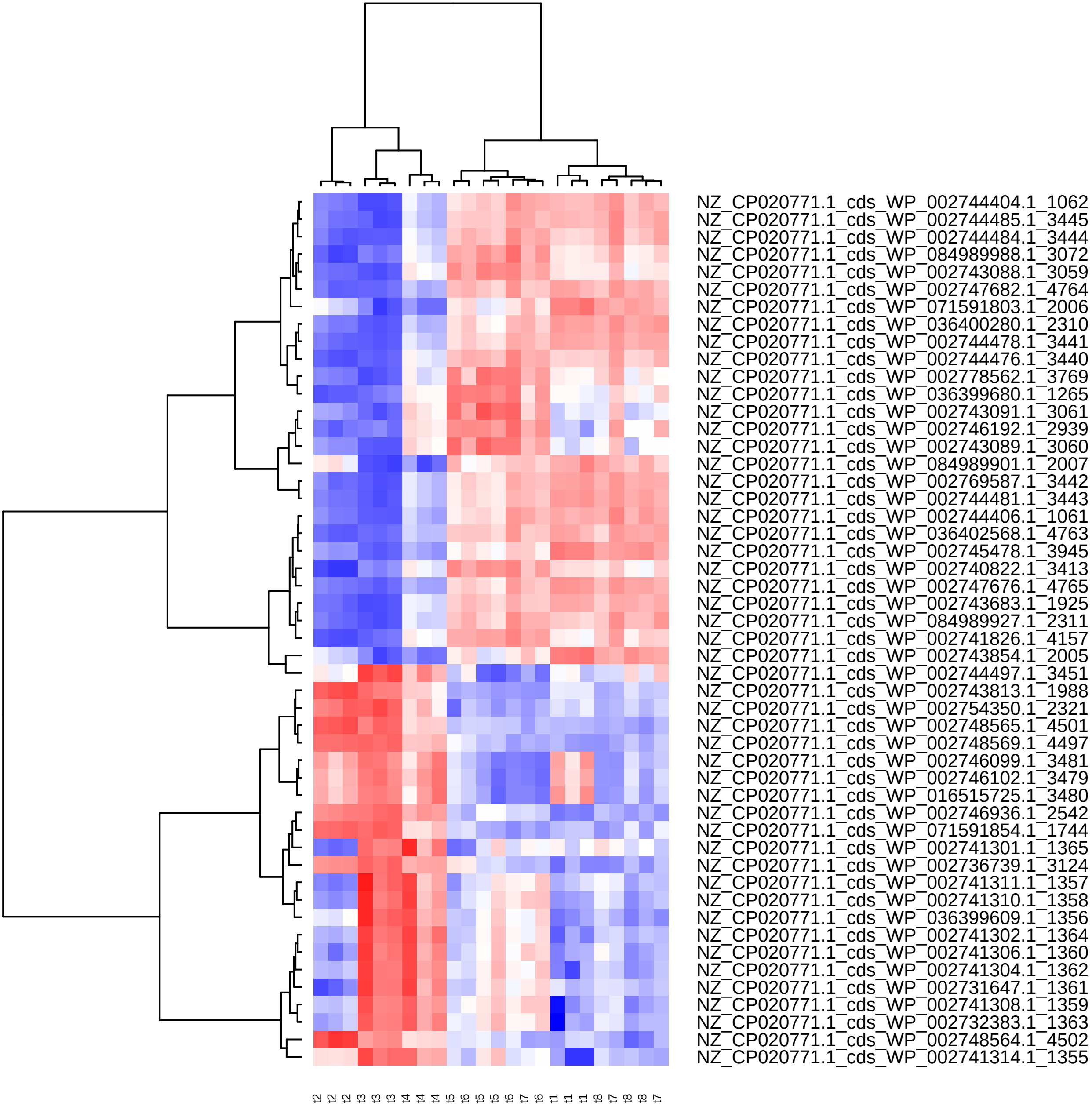

Supplement: Supplementary_Data [file 602159_file03.zip › Supplementary_DATA/DICOEXPRESS_output/DiffAnalysis/[t4-t5]/PCC7806_[t4-t5]_Top50_Clustering.pdf]

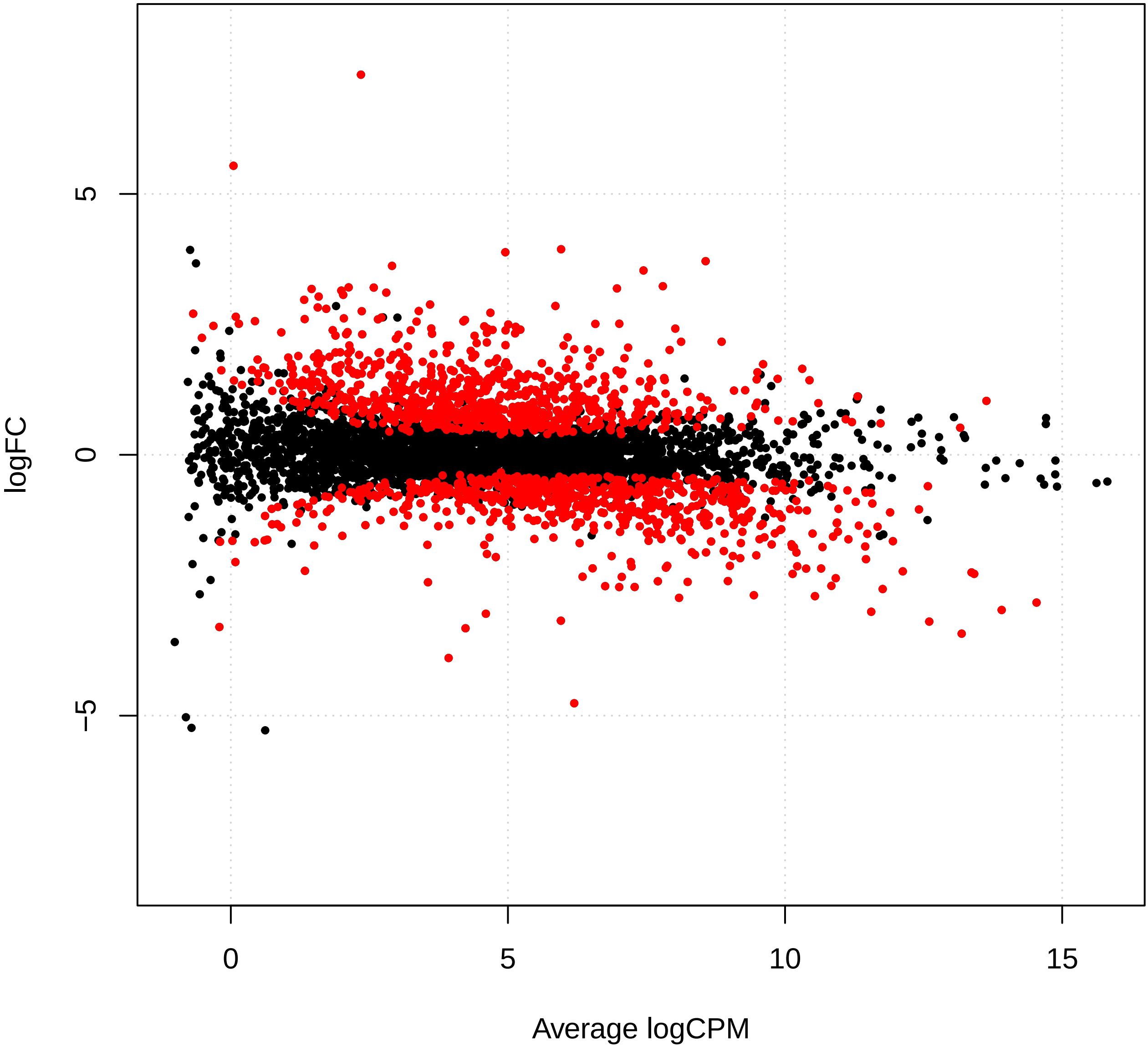

Supplement: Supplementary_Data [file 602159_file03.zip › Supplementary_DATA/DICOEXPRESS_output/DiffAnalysis/[t4-t6]/PCC7806_[t4-t6]_plotSmear.pdf]

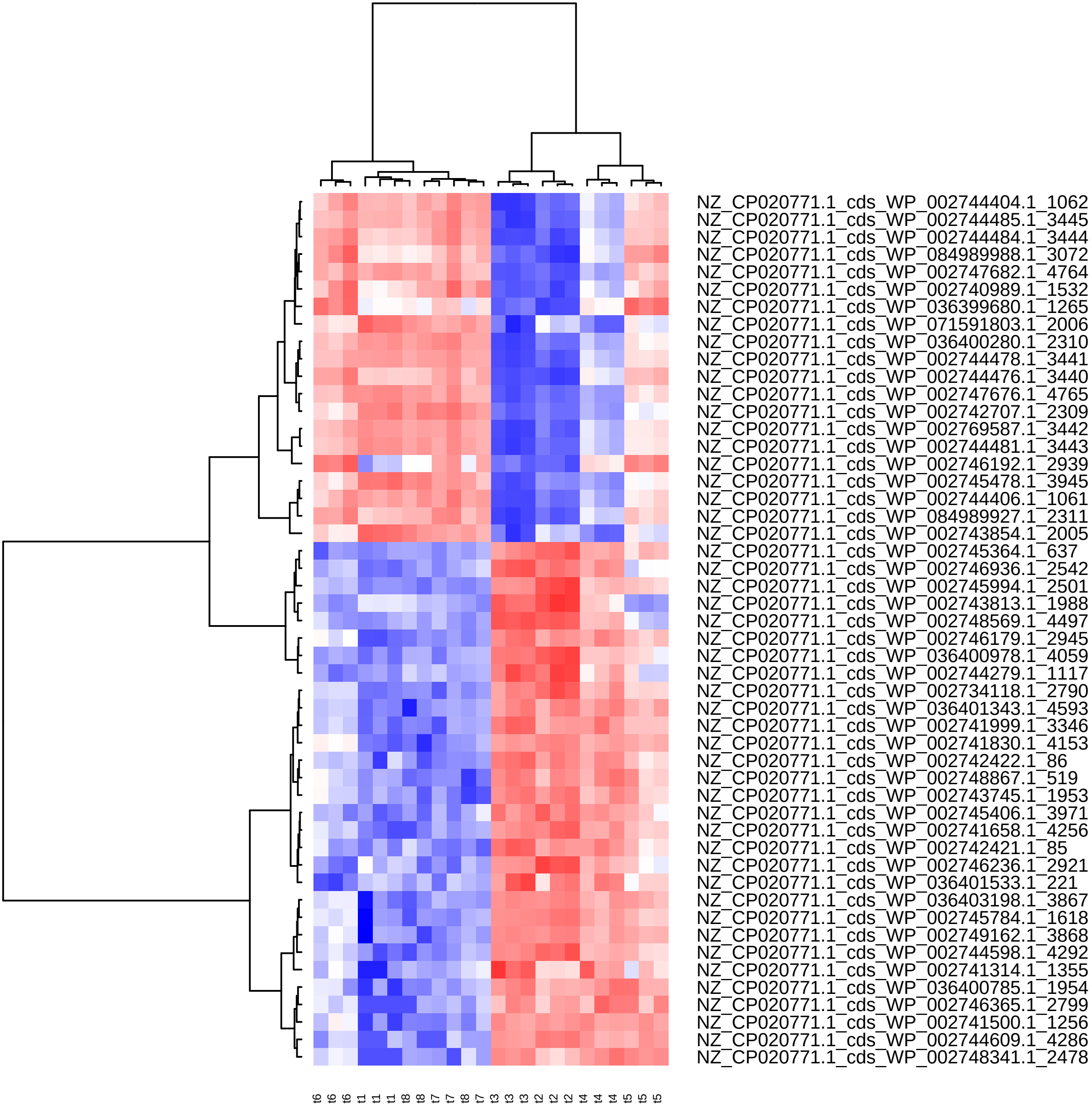

Supplement: Supplementary_Data [file 602159_file03.zip › Supplementary_DATA/DICOEXPRESS_output/DiffAnalysis/[t4-t6]/PCC7806_[t4-t6]_Top50_Clustering.pdf]

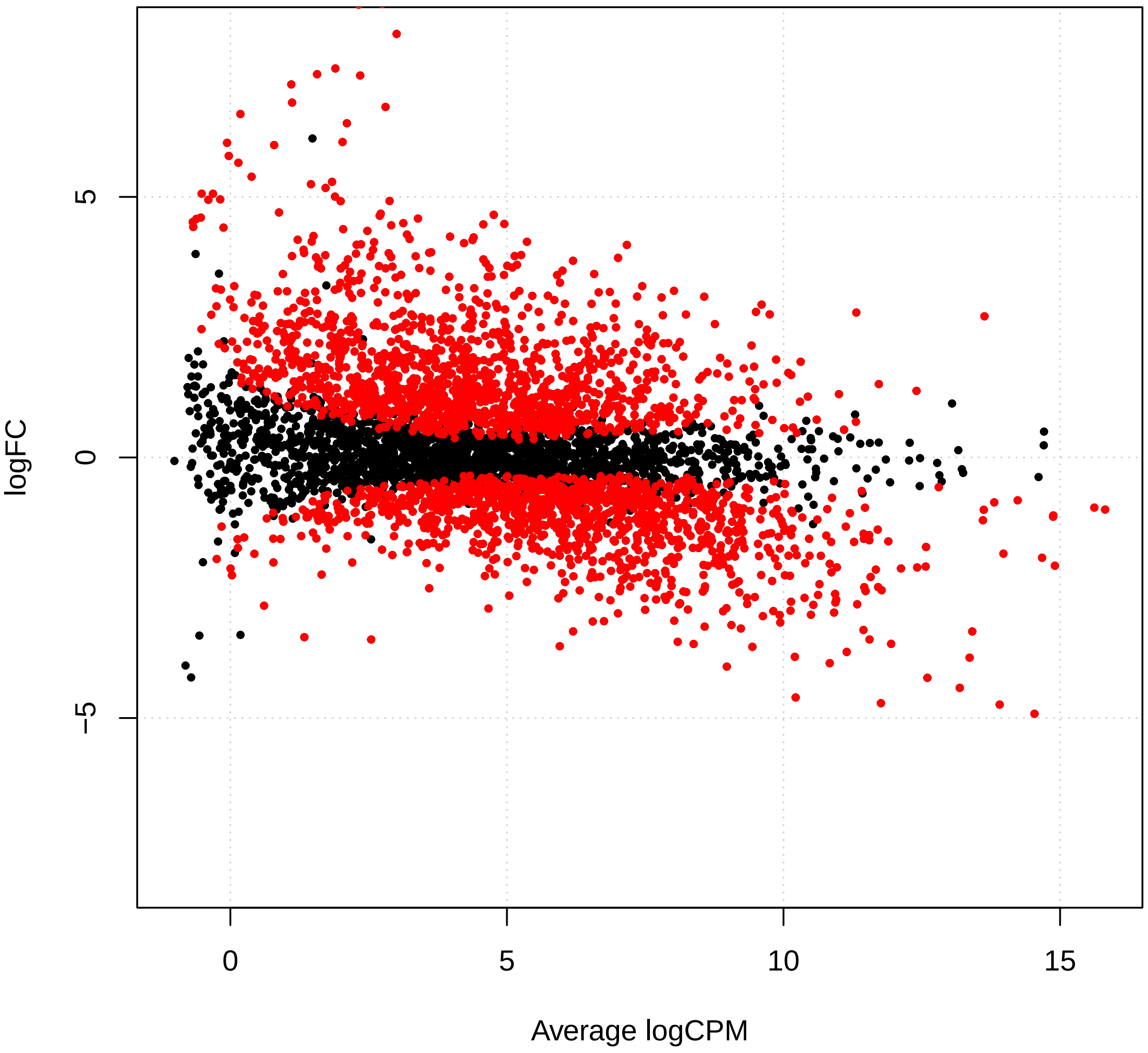

Supplement: Supplementary_Data [file 602159_file03.zip › Supplementary_DATA/DICOEXPRESS_output/DiffAnalysis/[t4-t7]/PCC7806_[t4-t7]_plotSmear.pdf]

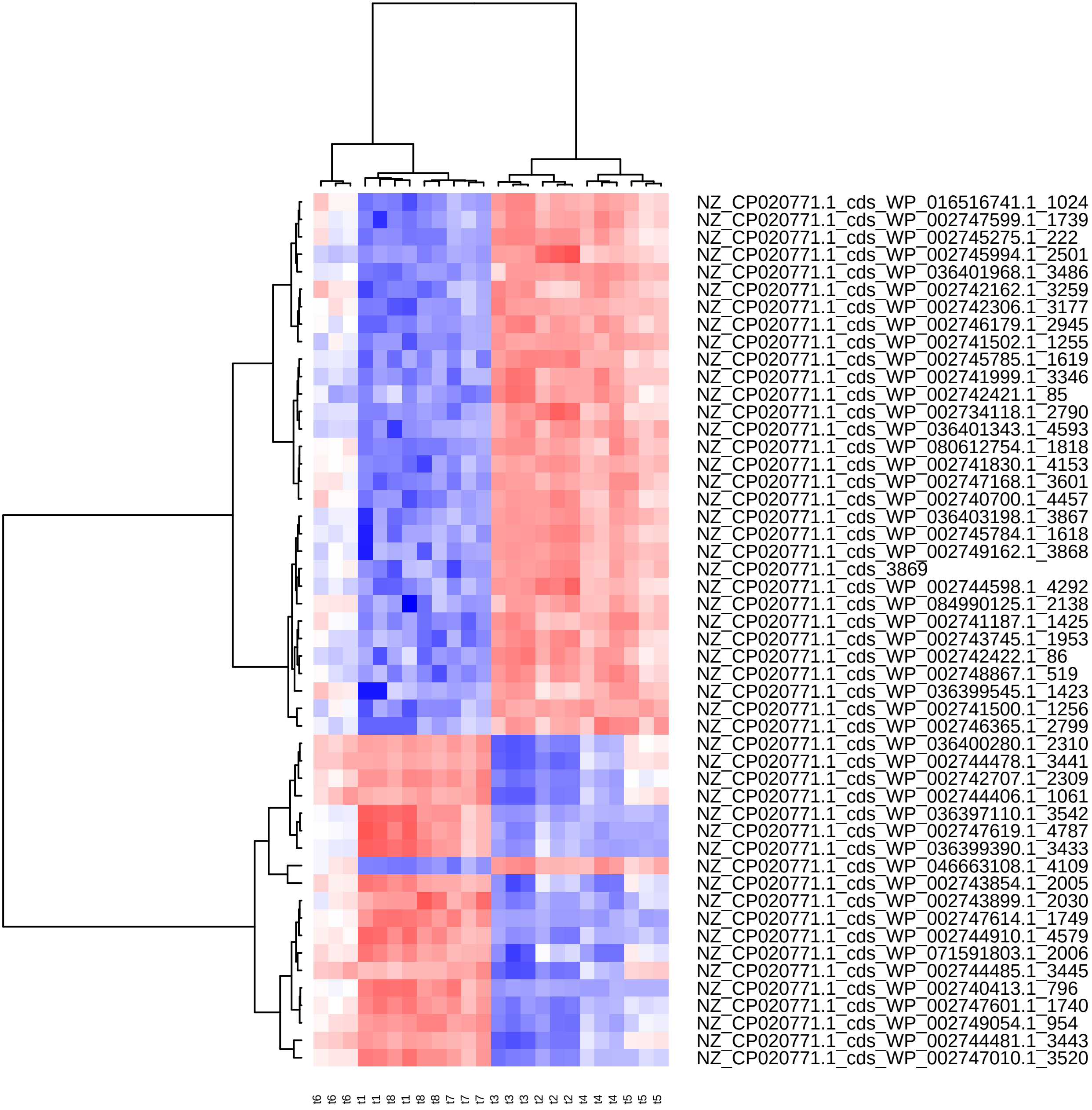

Supplement: Supplementary_Data [file 602159_file03.zip › Supplementary_DATA/DICOEXPRESS_output/DiffAnalysis/[t4-t7]/PCC7806_[t4-t7]_Top50_Clustering.pdf]

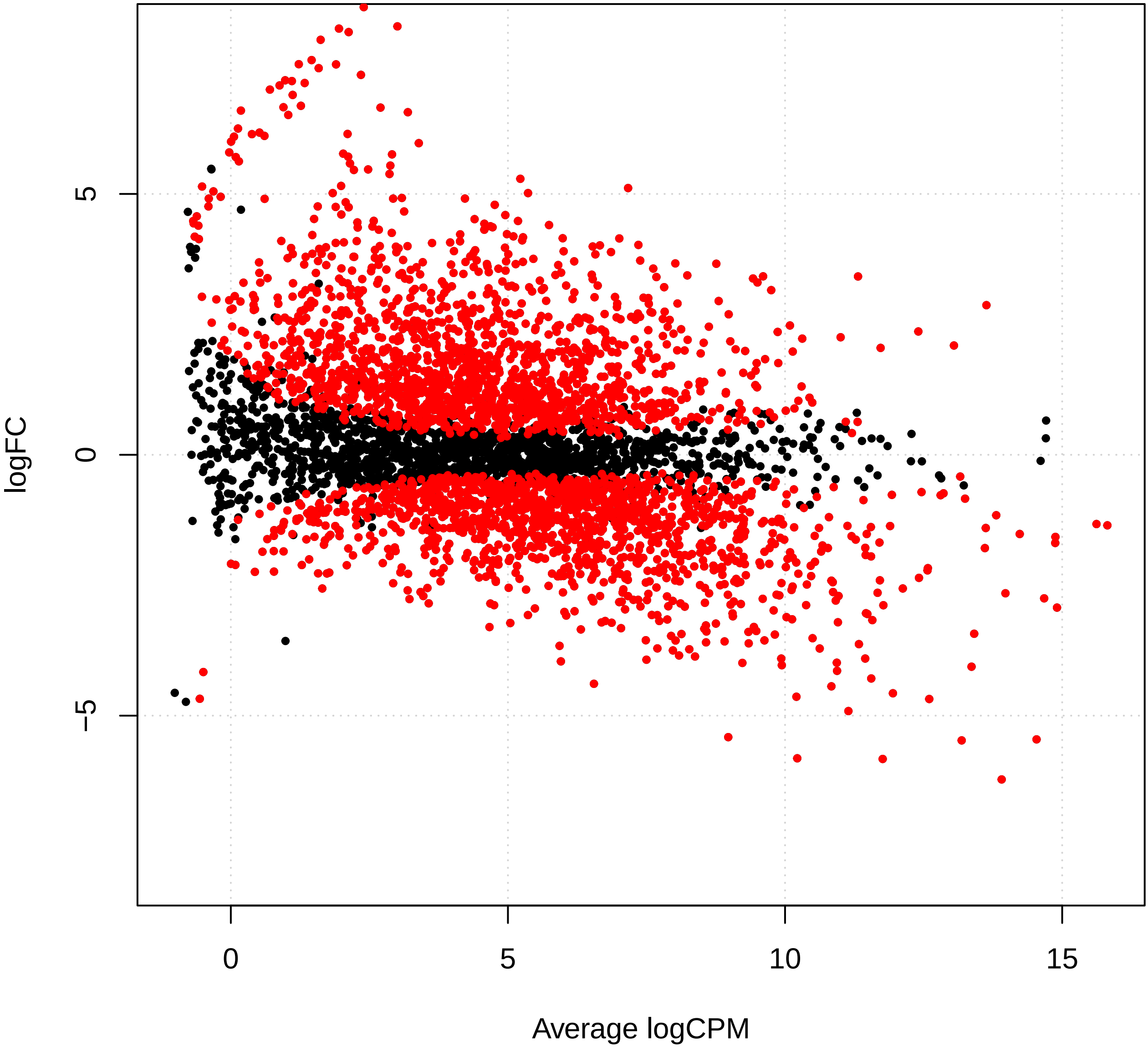

Supplement: Supplementary_Data [file 602159_file03.zip › Supplementary_DATA/DICOEXPRESS_output/DiffAnalysis/[t4-t8]/PCC7806_[t4-t8]_plotSmear.pdf]

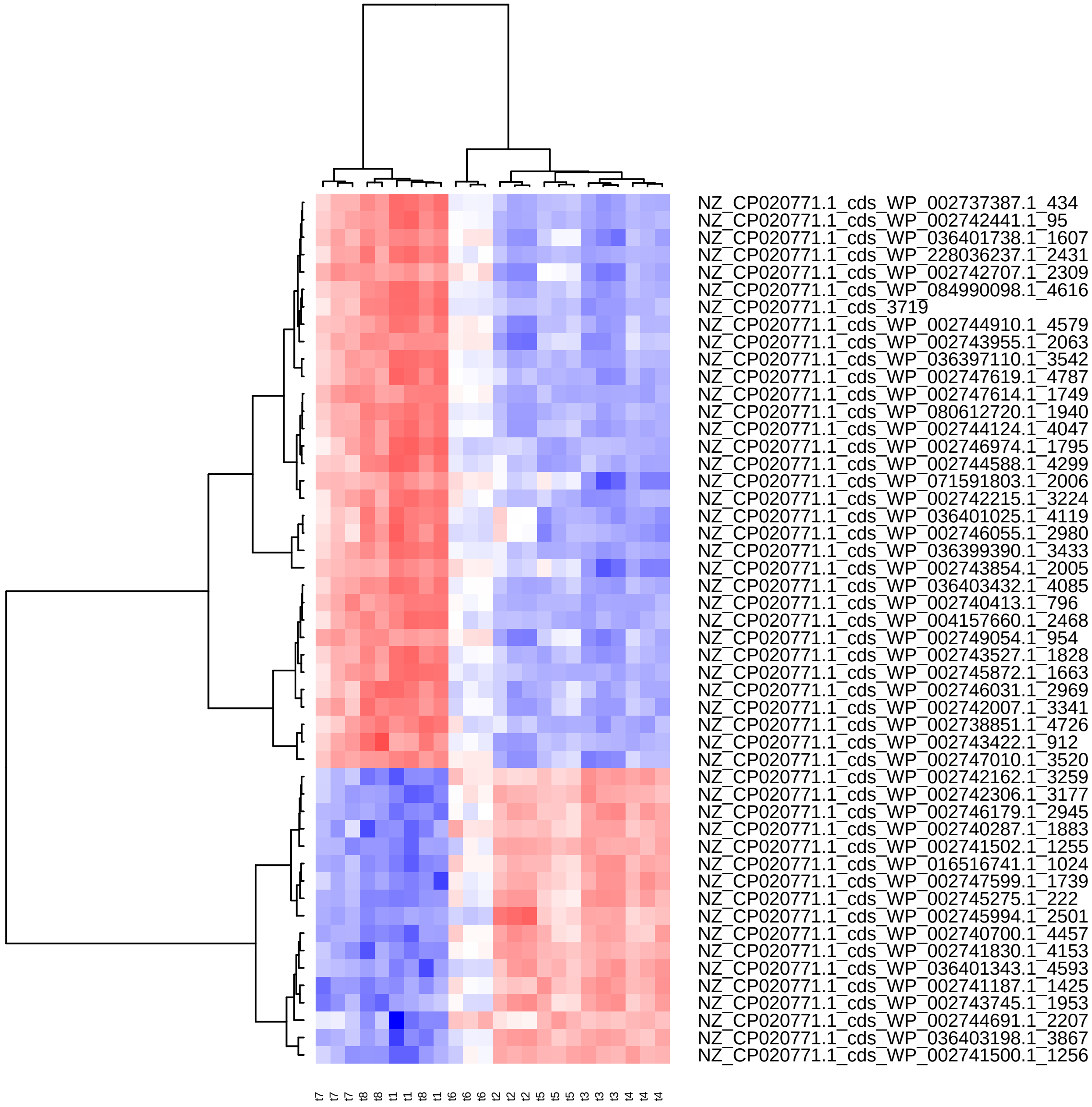

Supplement: Supplementary_Data [file 602159_file03.zip › Supplementary_DATA/DICOEXPRESS_output/DiffAnalysis/[t4-t8]/PCC7806_[t4-t8]_Top50_Clustering.pdf]

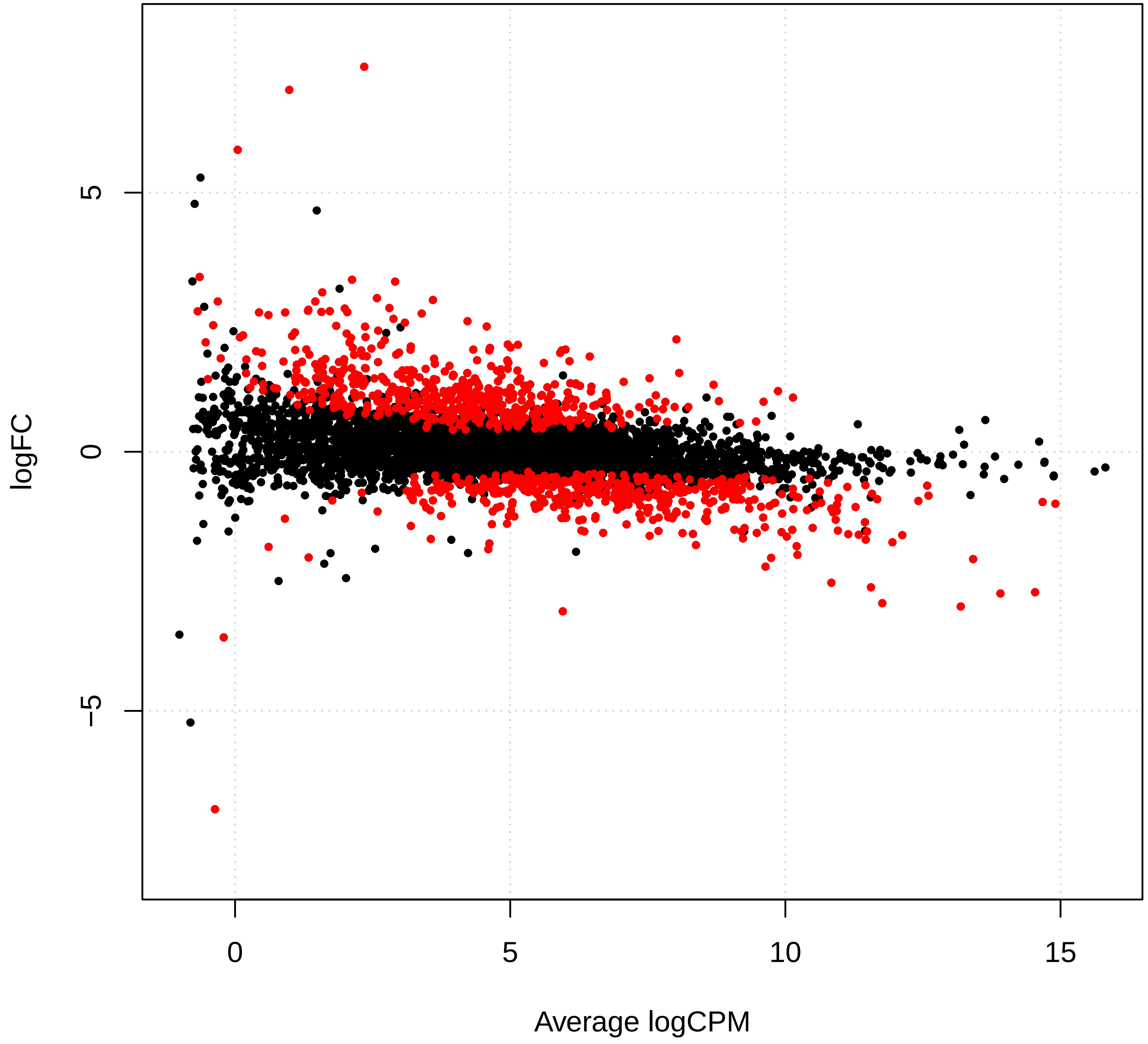

Supplement: Supplementary_Data [file 602159_file03.zip › Supplementary_DATA/DICOEXPRESS_output/DiffAnalysis/[t5-t6]/PCC7806_[t5-t6]_plotSmear.pdf]

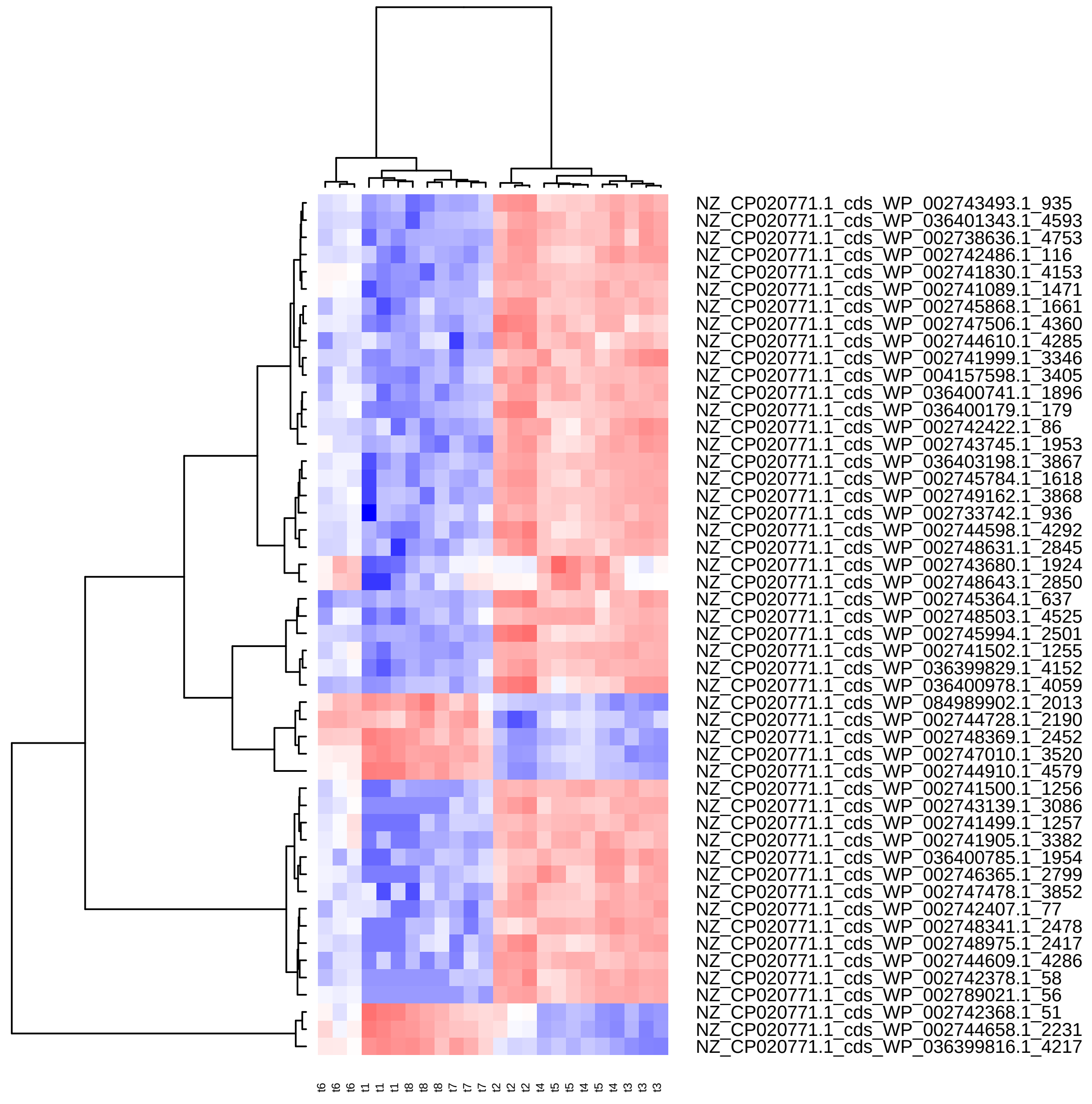

Supplement: Supplementary_Data [file 602159_file03.zip › Supplementary_DATA/DICOEXPRESS_output/DiffAnalysis/[t5-t6]/PCC7806_[t5-t6]_Top50_Clustering.pdf]

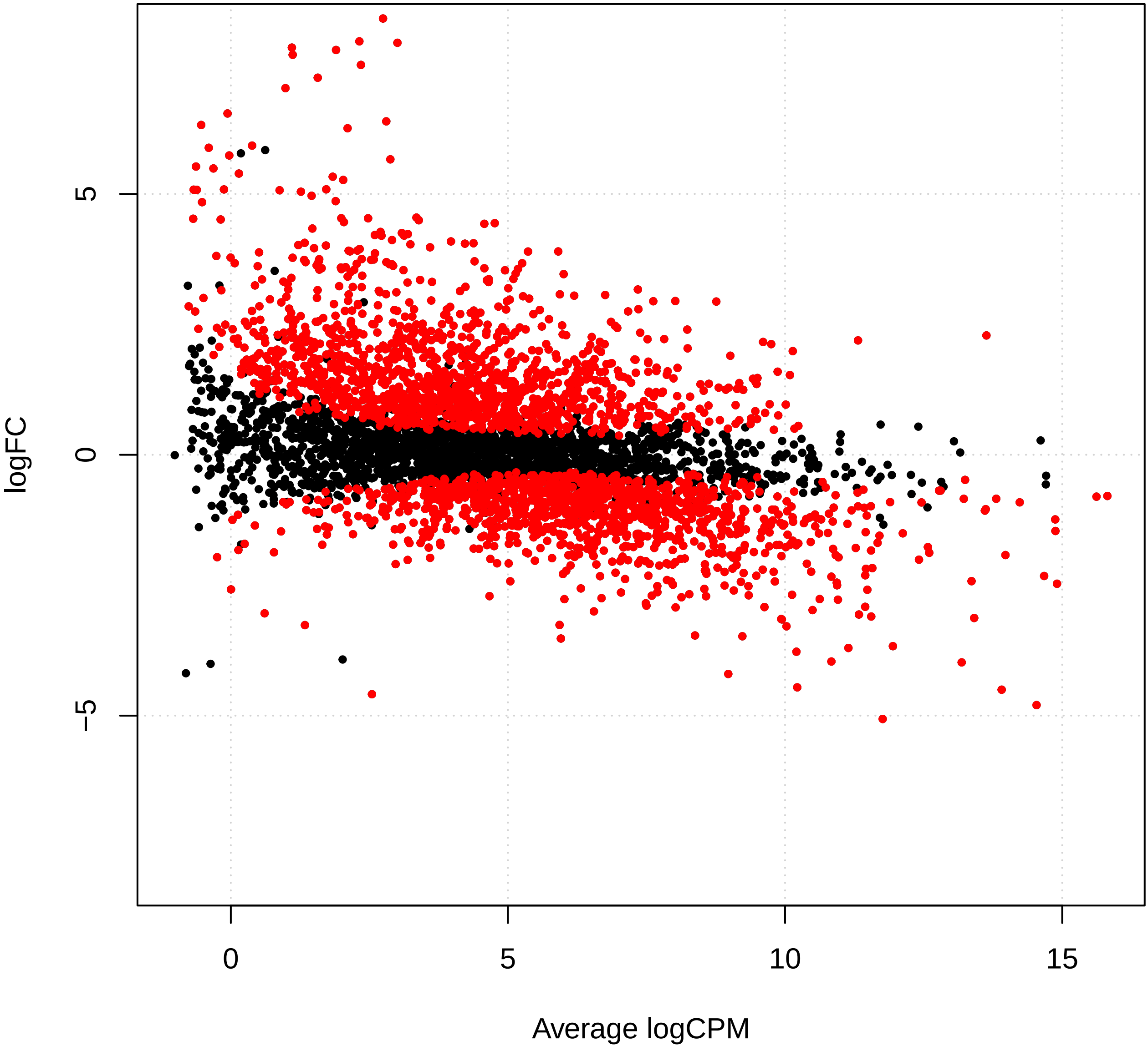

Supplement: Supplementary_Data [file 602159_file03.zip › Supplementary_DATA/DICOEXPRESS_output/DiffAnalysis/[t5-t7]/PCC7806_[t5-t7]_plotSmear.pdf]

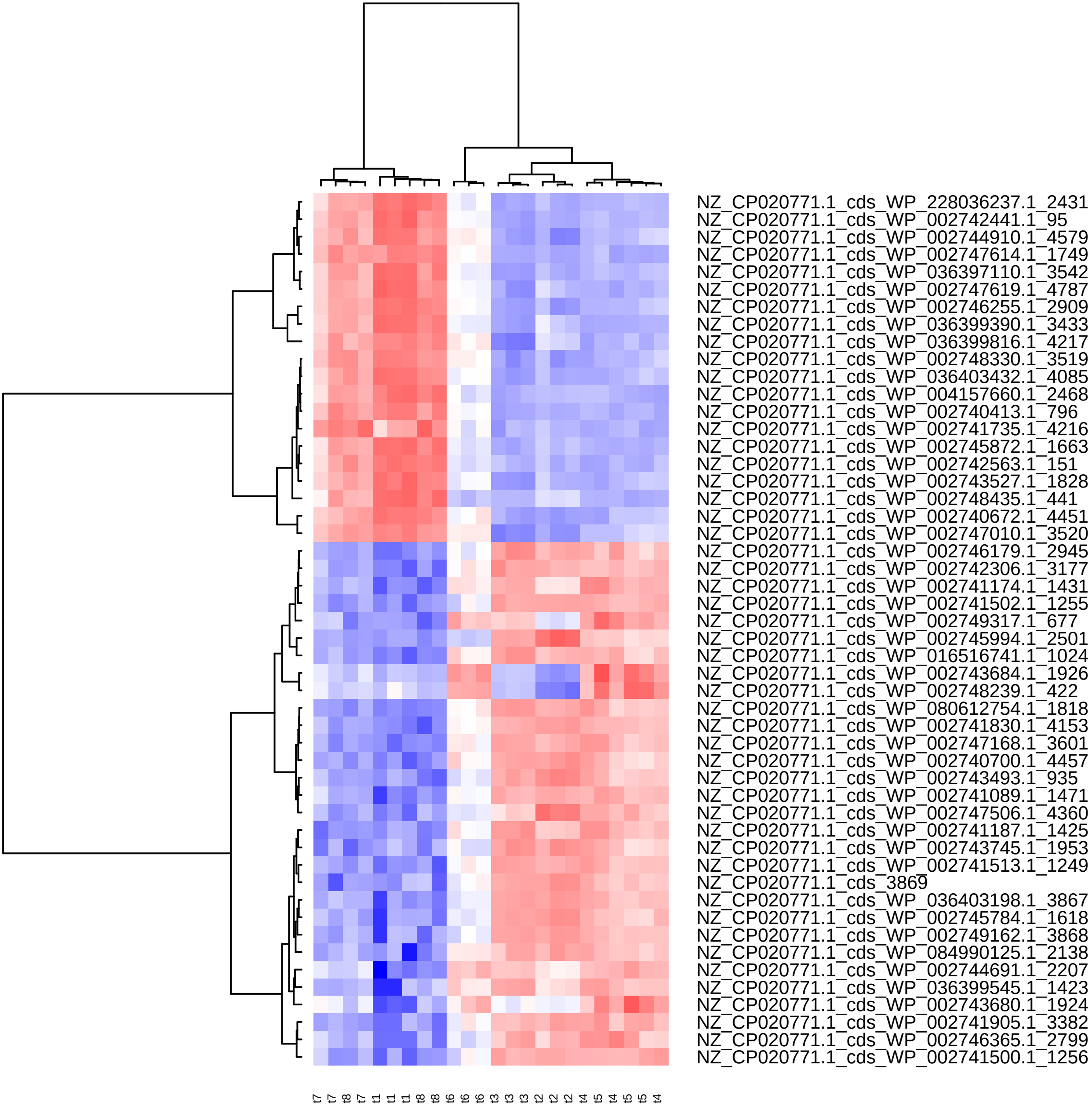

Supplement: Supplementary_Data [file 602159_file03.zip › Supplementary_DATA/DICOEXPRESS_output/DiffAnalysis/[t5-t7]/PCC7806_[t5-t7]_Top50_Clustering.pdf]

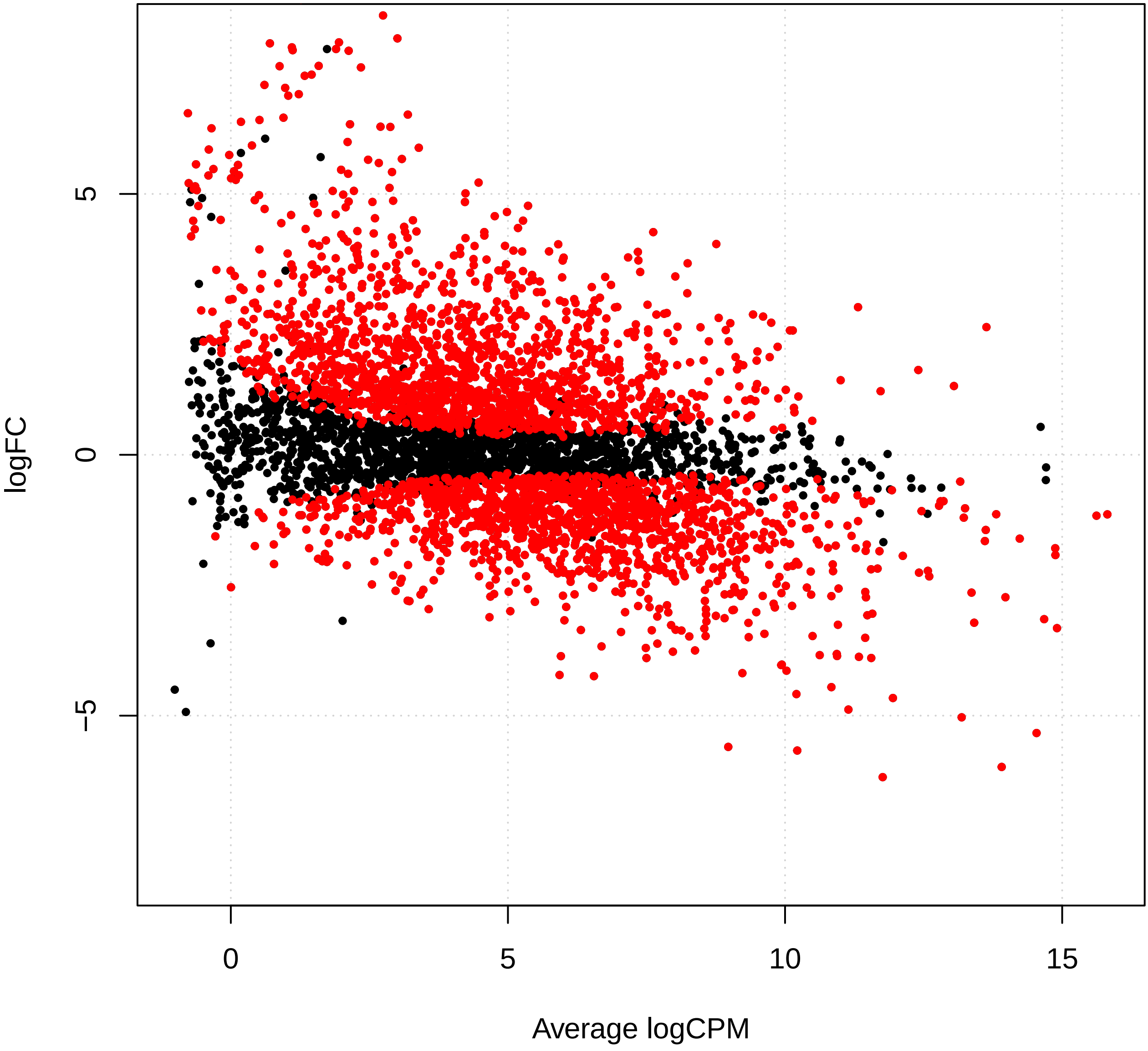

Supplement: Supplementary_Data [file 602159_file03.zip › Supplementary_DATA/DICOEXPRESS_output/DiffAnalysis/[t5-t8]/PCC7806_[t5-t8]_plotSmear.pdf]

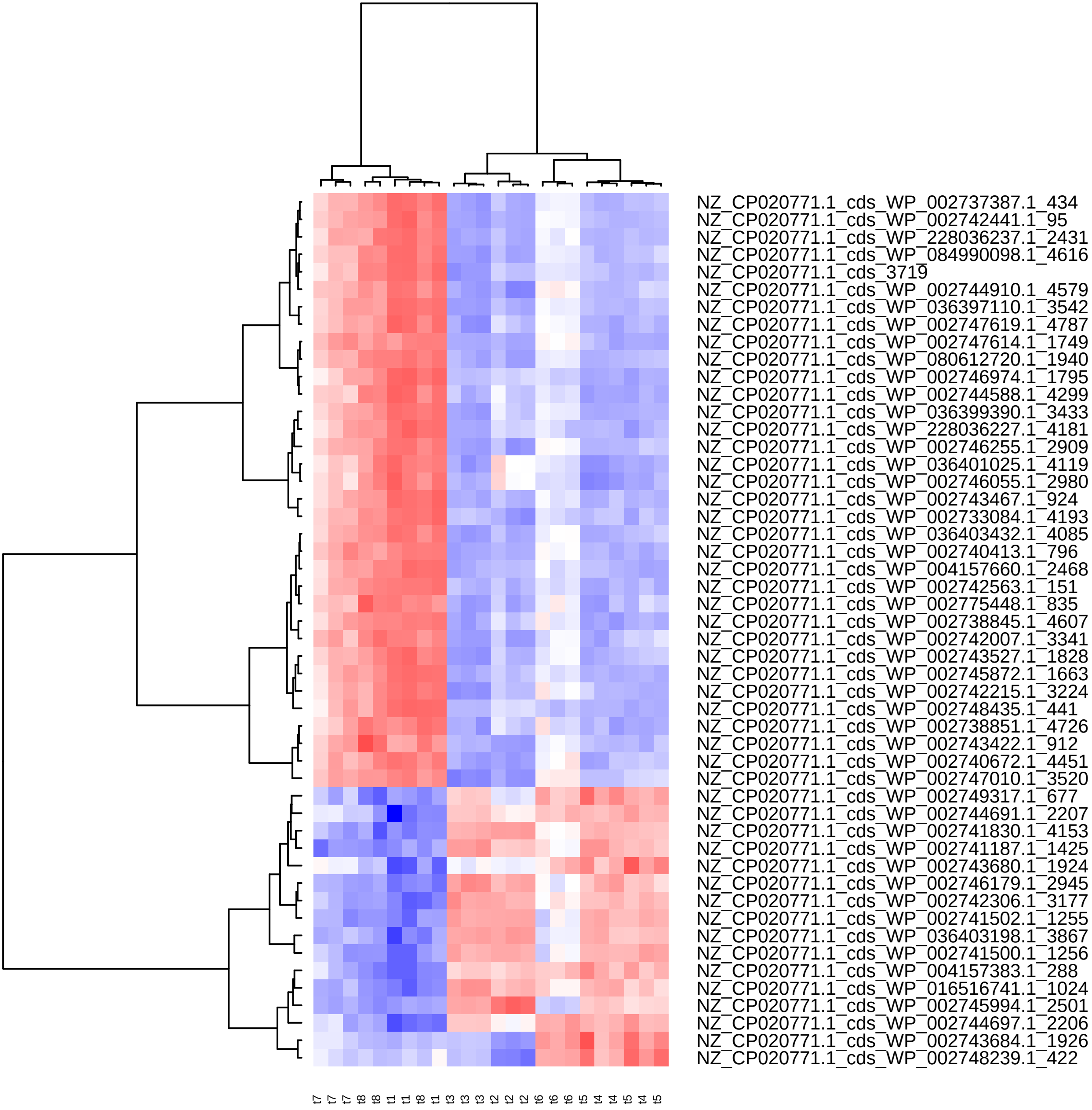

Supplement: Supplementary_Data [file 602159_file03.zip › Supplementary_DATA/DICOEXPRESS_output/DiffAnalysis/[t5-t8]/PCC7806_[t5-t8]_Top50_Clustering.pdf]

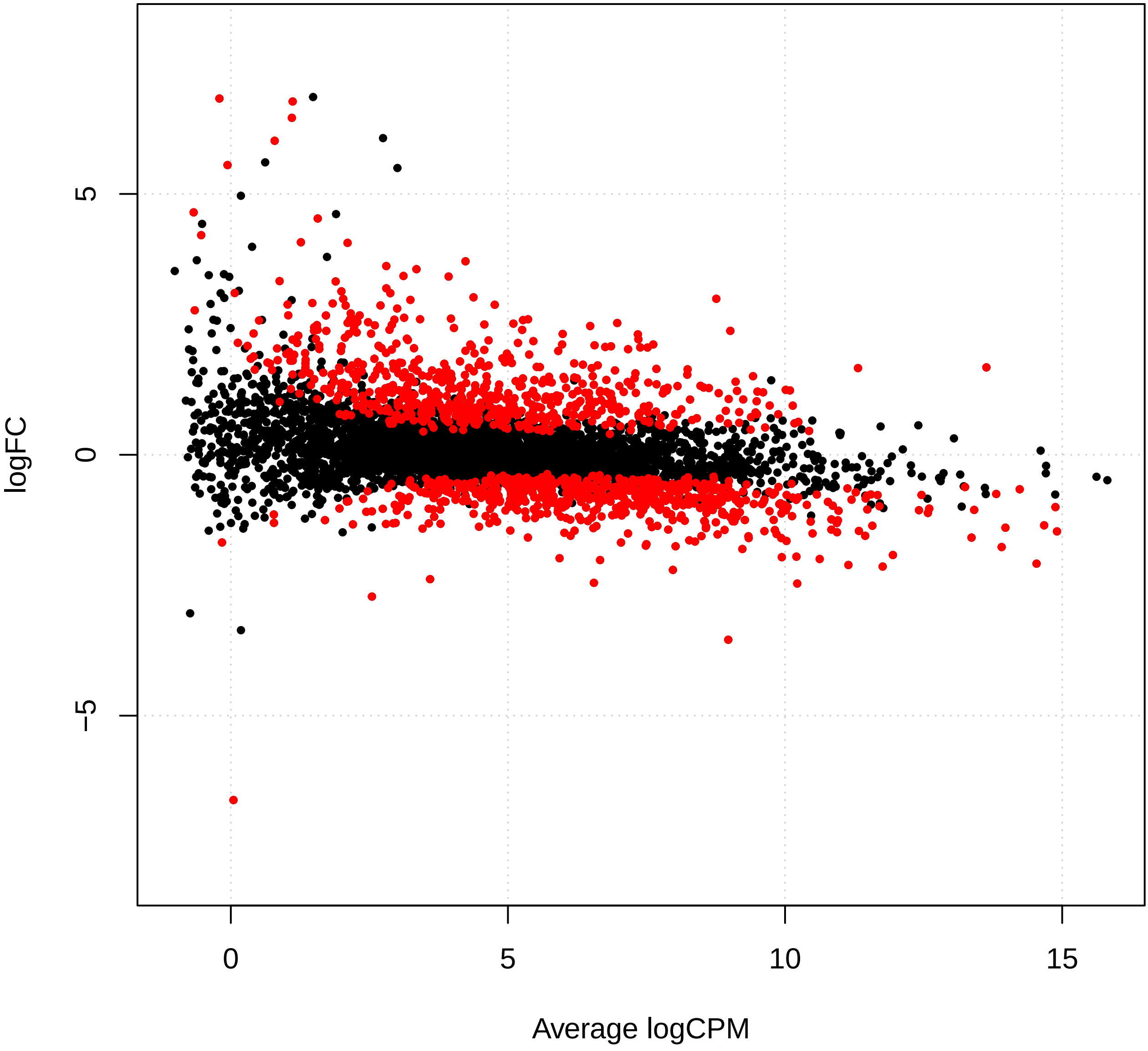

Supplement: Supplementary_Data [file 602159_file03.zip › Supplementary_DATA/DICOEXPRESS_output/DiffAnalysis/[t6-t7]/PCC7806_[t6-t7]_plotSmear.pdf]

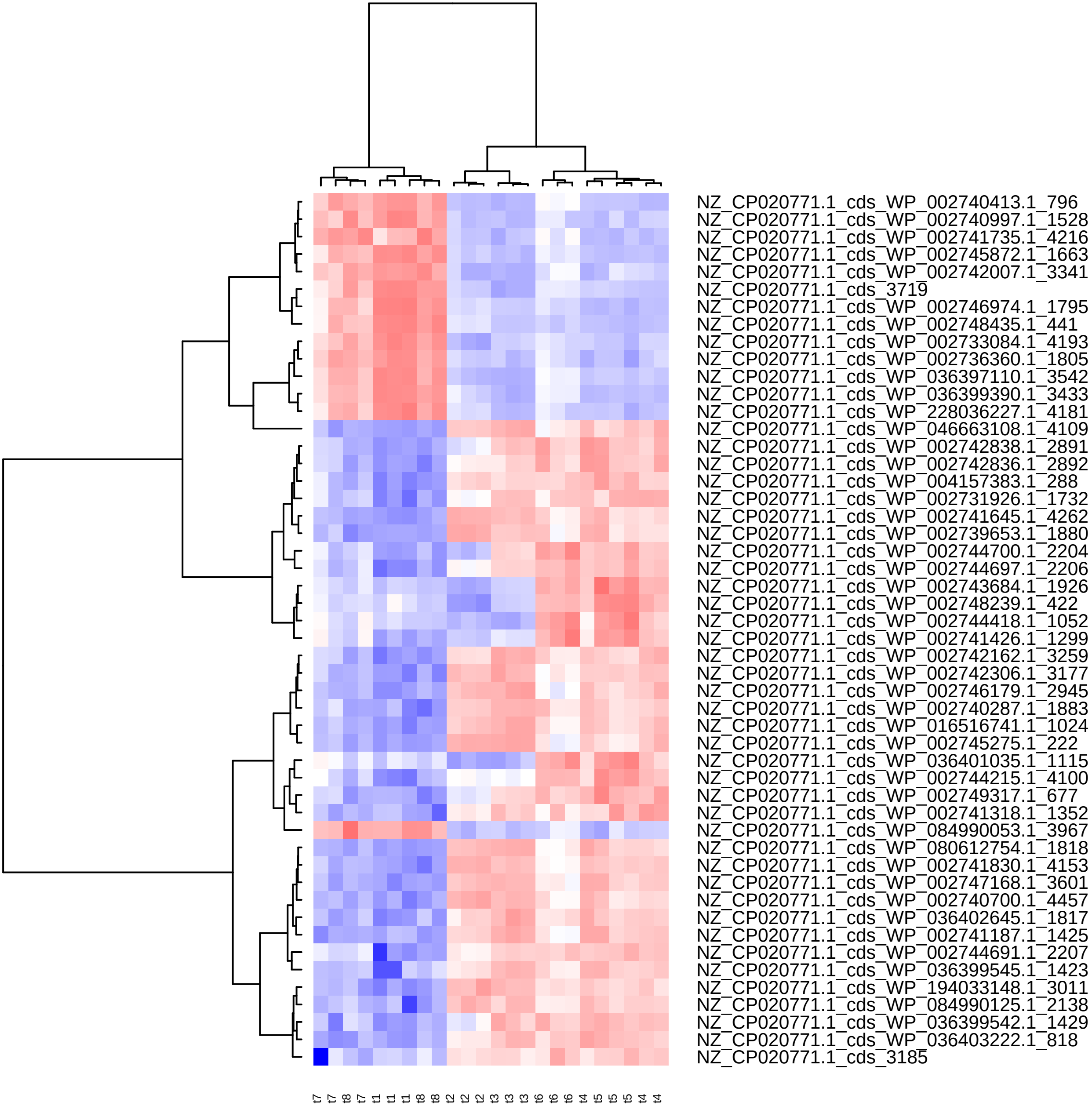

Supplement: Supplementary_Data [file 602159_file03.zip › Supplementary_DATA/DICOEXPRESS_output/DiffAnalysis/[t6-t7]/PCC7806_[t6-t7]_Top50_Clustering.pdf]

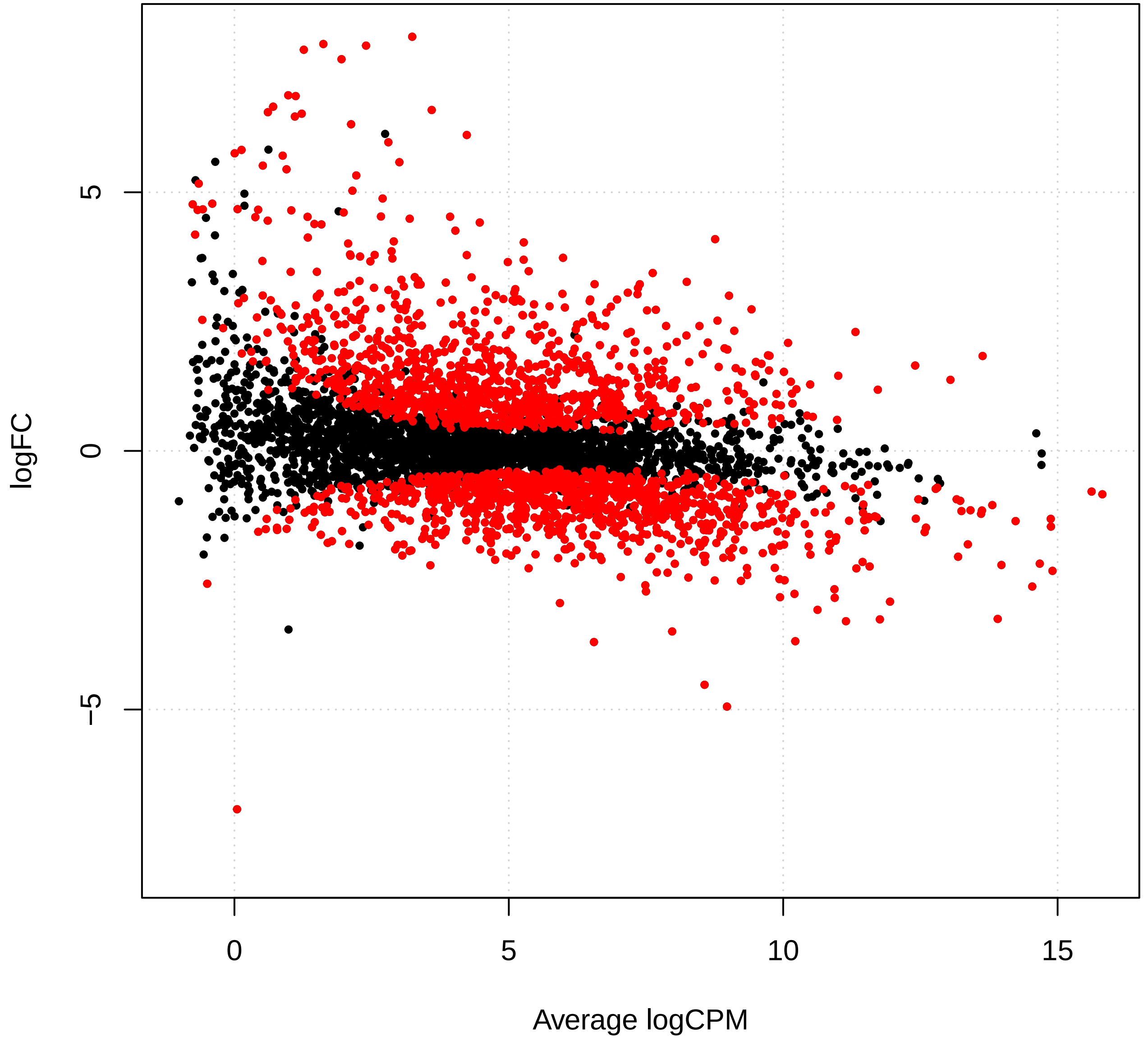

Supplement: Supplementary_Data [file 602159_file03.zip › Supplementary_DATA/DICOEXPRESS_output/DiffAnalysis/[t6-t8]/PCC7806_[t6-t8]_plotSmear.pdf]

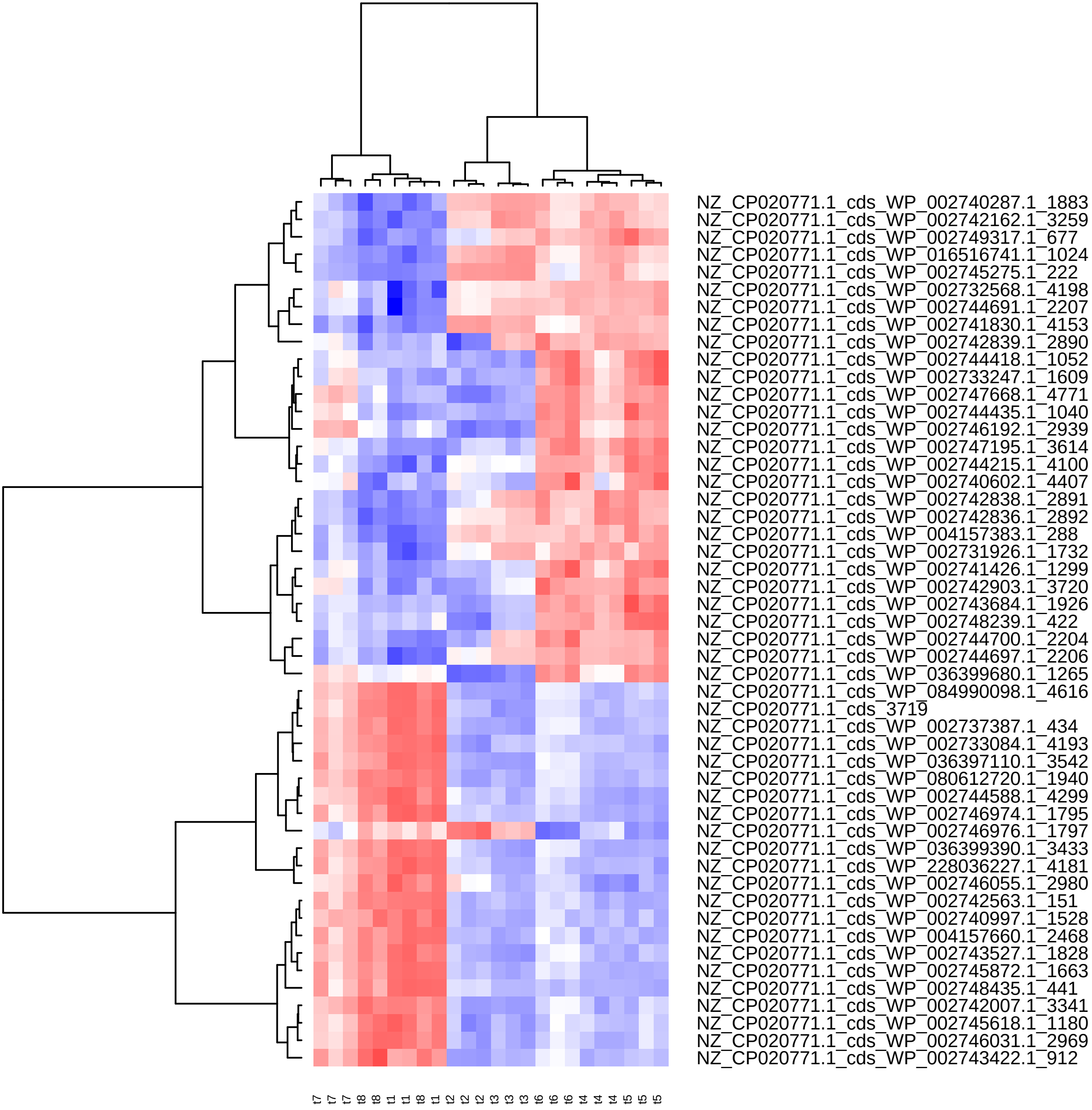

Supplement: Supplementary_Data [file 602159_file03.zip › Supplementary_DATA/DICOEXPRESS_output/DiffAnalysis/[t6-t8]/PCC7806_[t6-t8]_Top50_Clustering.pdf]

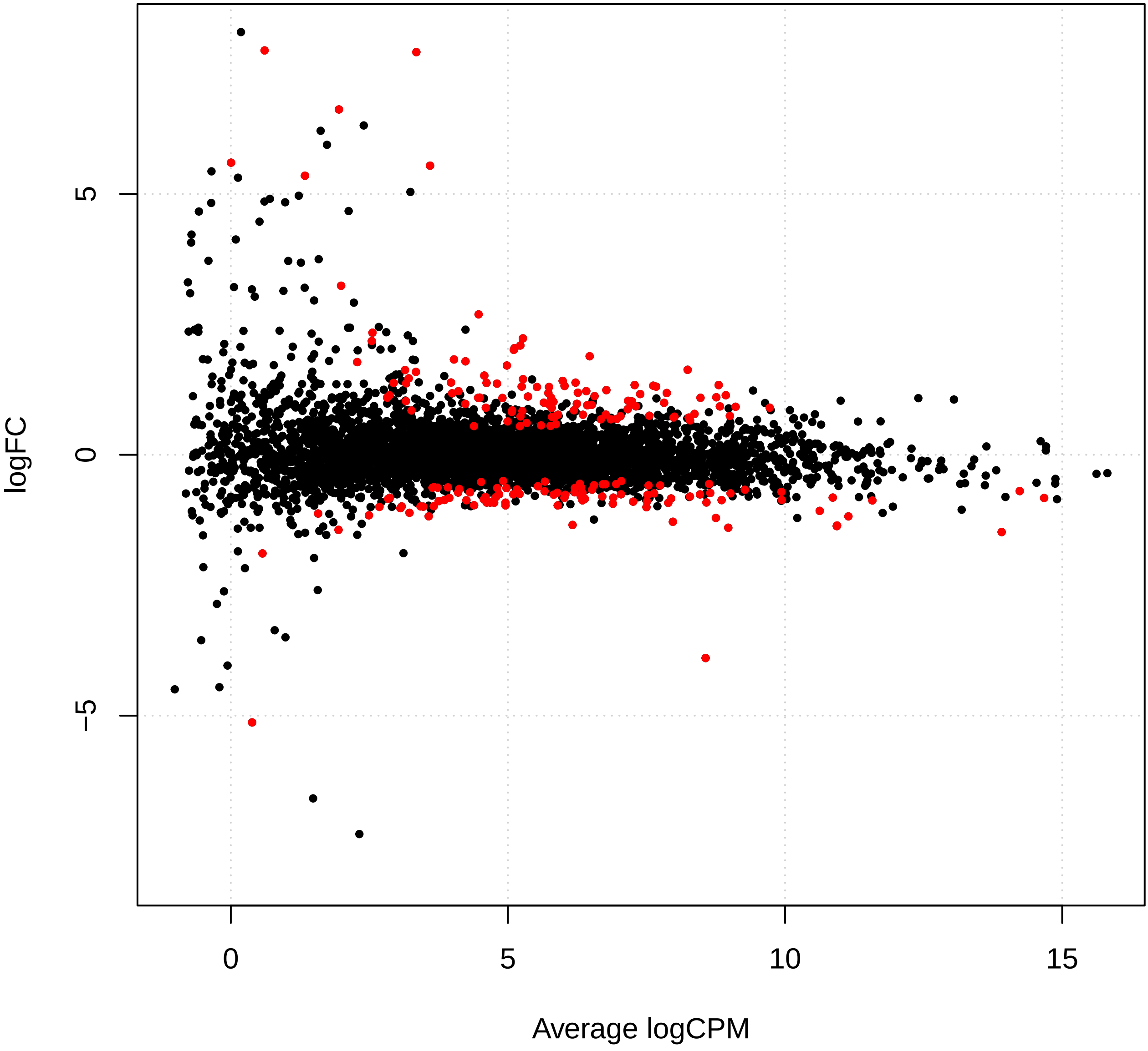

Supplement: Supplementary_Data [file 602159_file03.zip › Supplementary_DATA/DICOEXPRESS_output/DiffAnalysis/[t7-t8]/PCC7806_[t7-t8]_plotSmear.pdf]

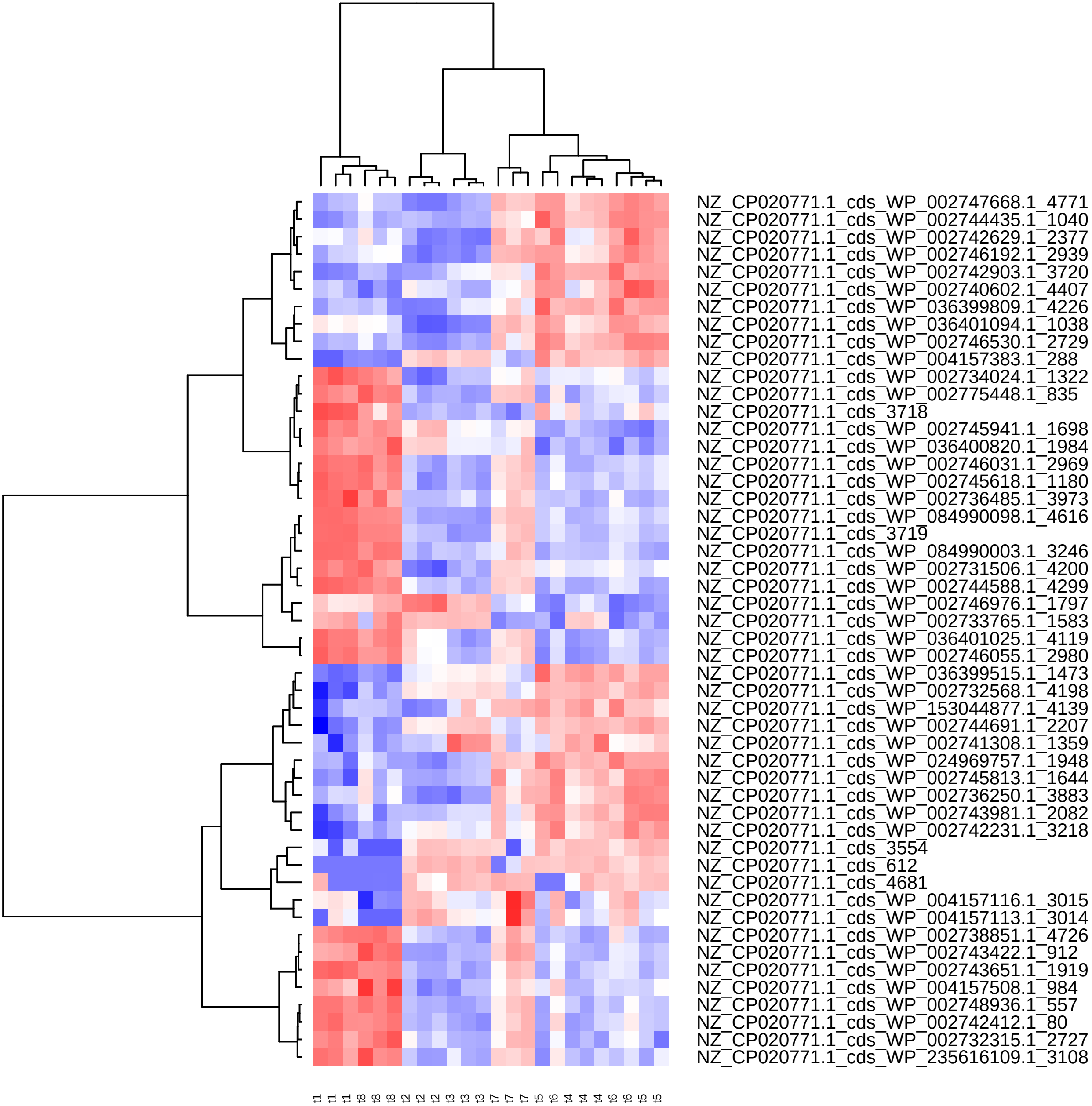

Supplement: Supplementary_Data [file 602159_file03.zip › Supplementary_DATA/DICOEXPRESS_output/DiffAnalysis/[t7-t8]/PCC7806_[t7-t8]_Top50_Clustering.pdf]

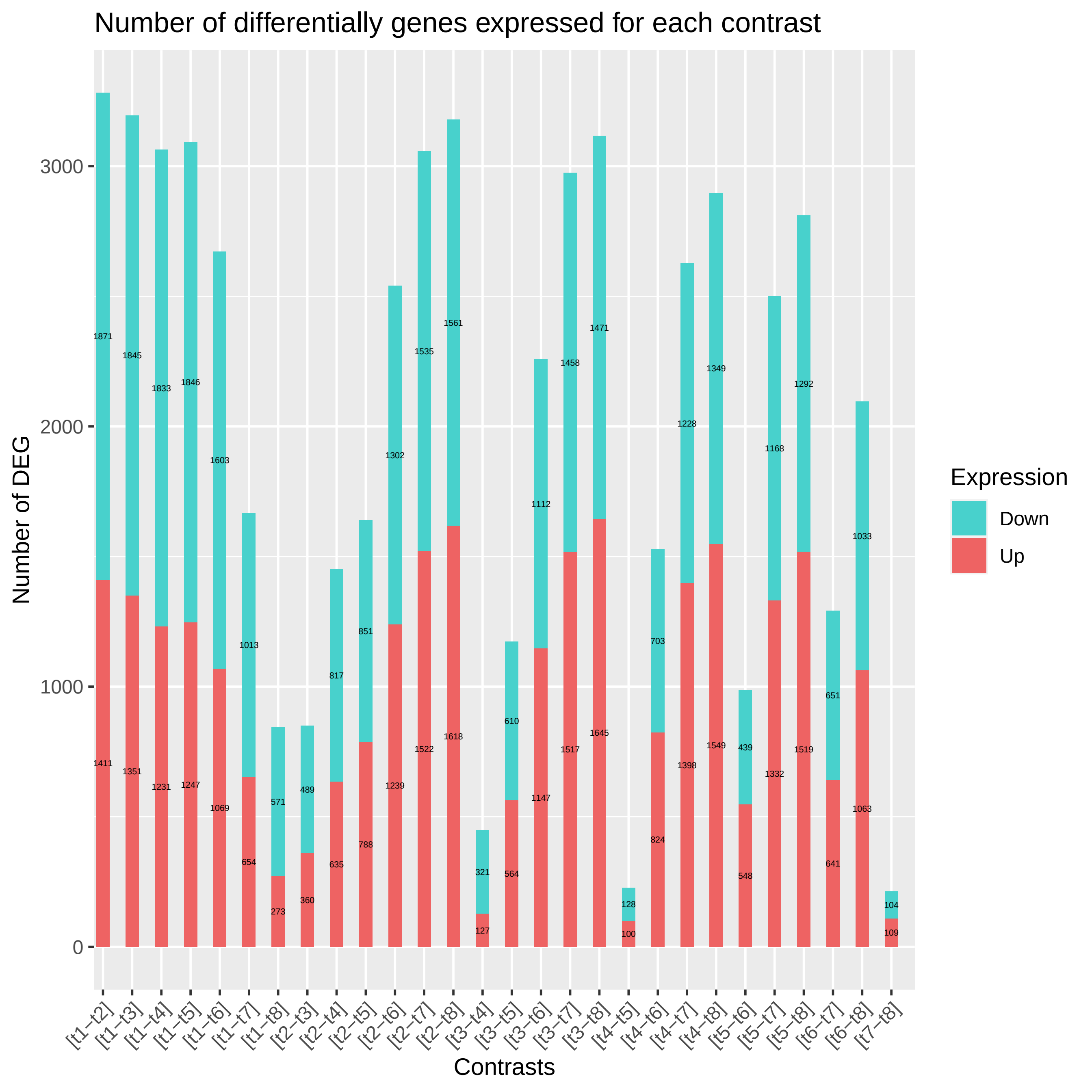
