## Supplementary_Figures for "Diel changes in the expression of a marker gene and candidate genes for intracellular amorphous CaCO_3_ biomineralization in *Microcystis*": Figure_S1.pdf

# WP_002731545.1

### DUF1269 domain-containing protein

#### GlyZip

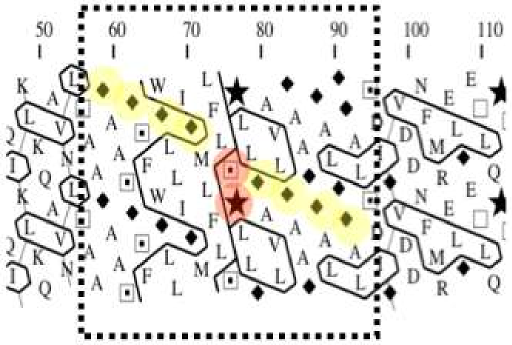


```
WP_002731545.1   KQAVNLTAAGAASGSFWGLLIGMLFLSPLLGAVLGAAGGALGGALSDIGVDD
WP_084990073.1   GGAFLGMSAGQVAGGMIGGAIGSAL.GP.GGAVLGAQVGAFAGSTTGAKLGY
```
