## Supplementary_Figures for "Diel changes in the expression of a marker gene and candidate genes for intracellular amorphous CaCO_3_ biomineralization in *Microcystis*": Figure_S3_SI_voisinsccyA.pdf

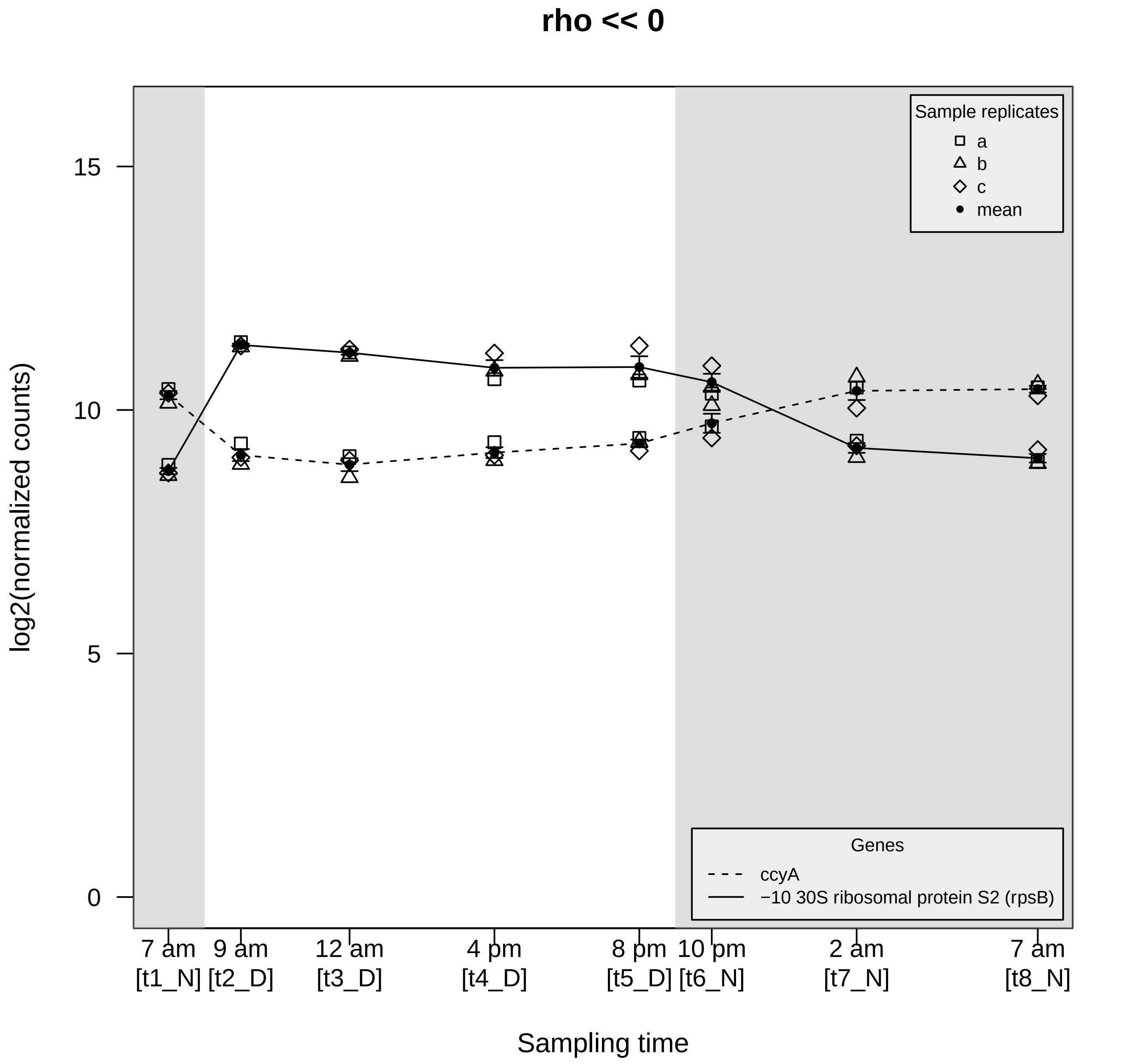

rho << 0
Sample replicates
a
b
c
mean
Genes
ccyA
−10 30S ribosomal protein S2 (rpsB)
log2(normalized counts)
0
5
10
15
7 am [t1_N]
9 am [t2_D]
12 am [t3_D]
4 pm [t4_D]
8 pm [t5_D]
10 pm [t6_N]
2 am [t7_N]
7 am [t8_N]
Sampling time

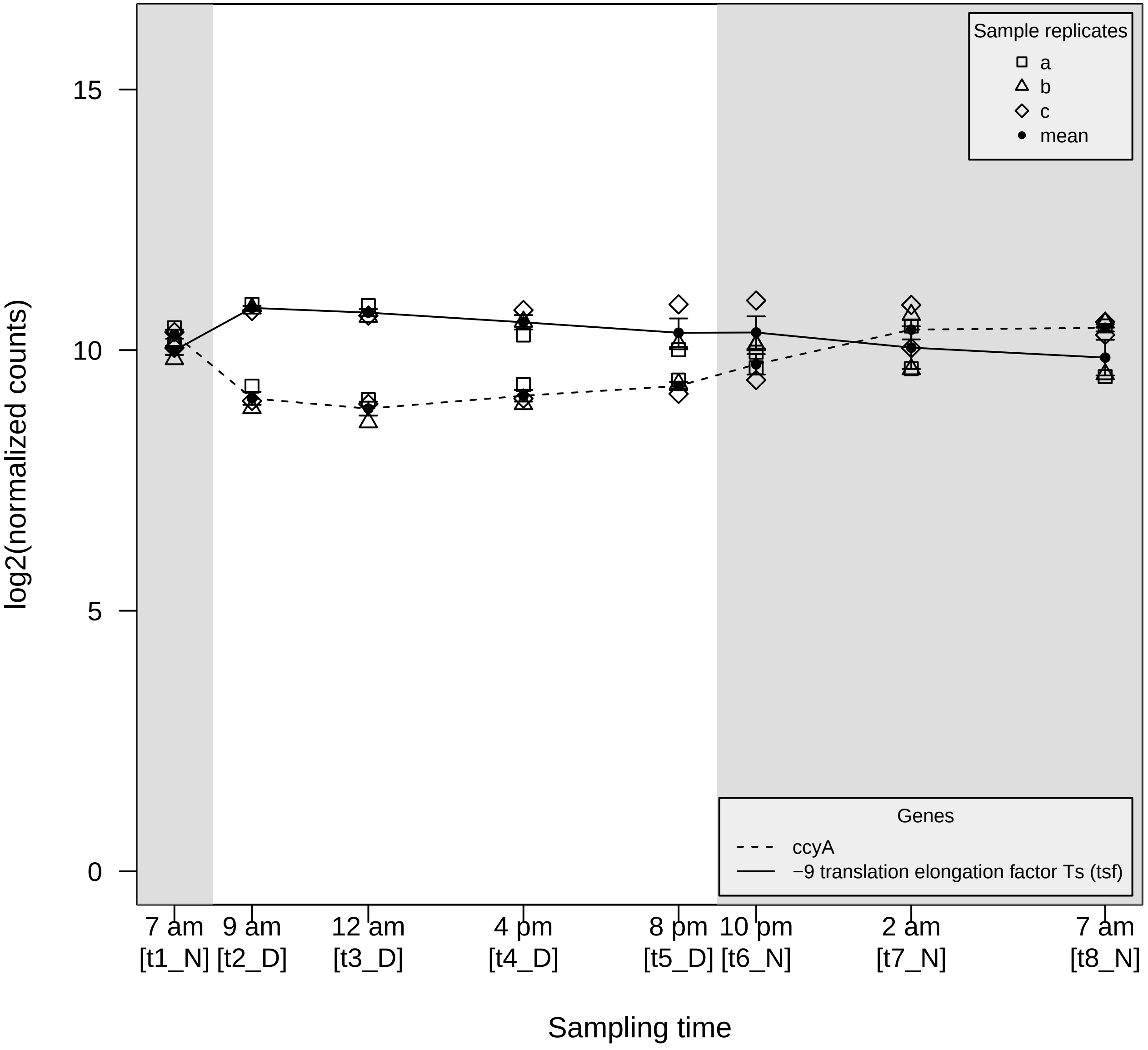

Sample replicates
a
b
c
mean
Genes
ccyA
−9 translation elongation factor Ts (tsf)
log2(normalized counts)
15
10
5
0
7 am [t1_N]
9 am [t2_D]
12 am [t3_D]
4 pm [t4_D]
8 pm [t5_D]
10 pm [t6_N]
2 am [t7_N]
7 am [t8_N]
Sampling time

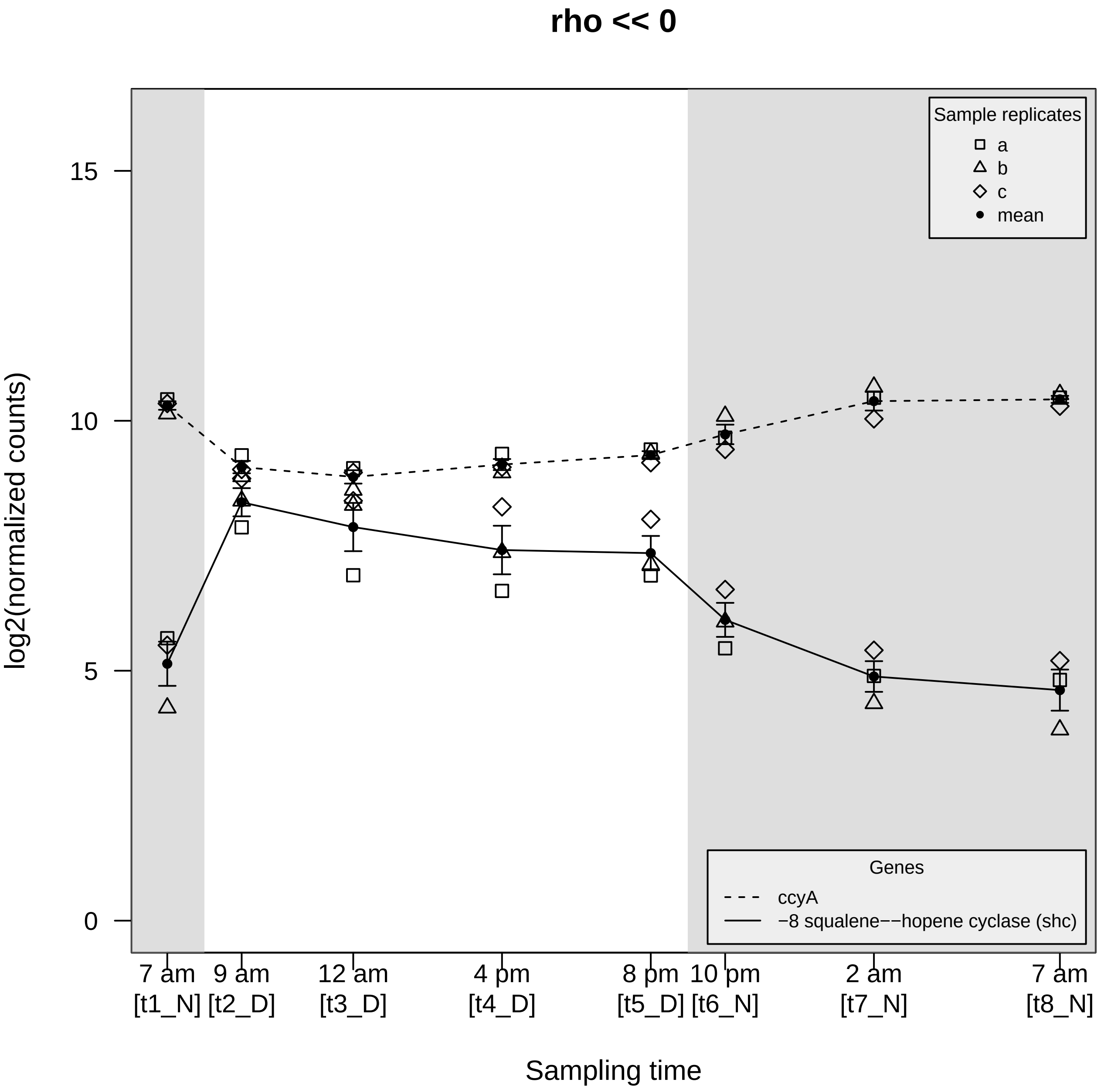
rho << 0
log2(normalized counts)
15
10
5
0
Sample replicates
a
b
c
mean
Genes
ccyA
−8 squalene−−hopene cyclase (shc)
7 am [t1_N]
9 am [t2_D]
12 am [t3_D]
4 pm [t4_D]
8 pm [t5_D]
10 pm [t6_N]
2 am [t7_N]
7 am [t8_N]
Sampling time

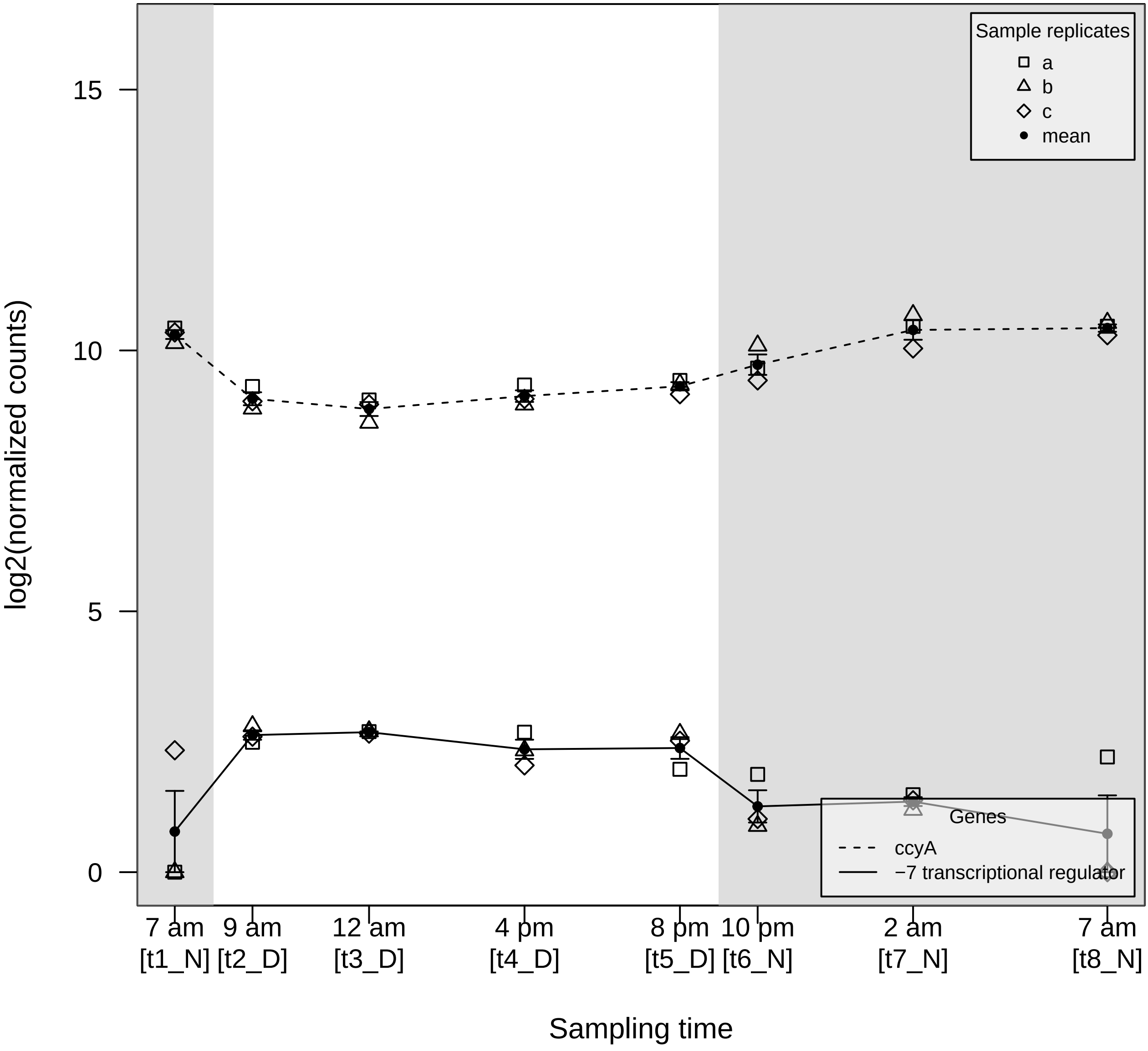

Sample replicates
a
b
c
mean
Genes
ccyA
−7 transcriptional regulator
15
10
5
0
log2(normalized counts)
7 am [t1_N]
9 am [t2_D]
12 am [t3_D]
4 pm [t4_D]
8 pm [t5_D]
10 pm [t6_N]
2 am [t7_N]
7 am [t8_N]
Sampling time

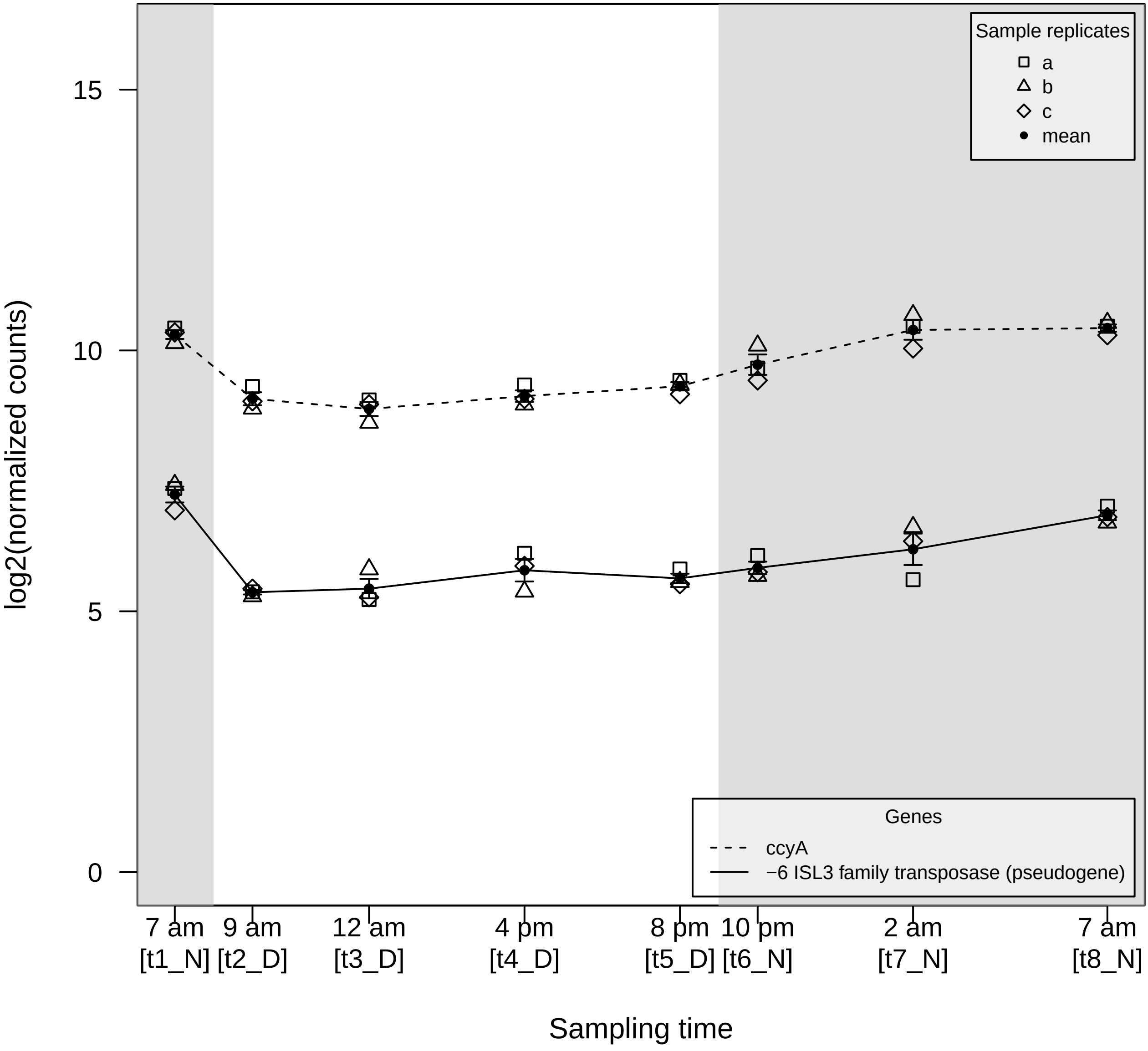

log2(normalized counts)
15
10
5
0
Sample replicates
a
b
c
mean
Genes
ccyA
−6 ISL3 family transposase (pseudogene)
7 am [t1_N]
9 am [t2_D]
12 am [t3_D]
4 pm [t4_D]
8 pm [t5_D]
10 pm [t6_N]
2 am [t7_N]
7 am [t8_N]
Sampling time

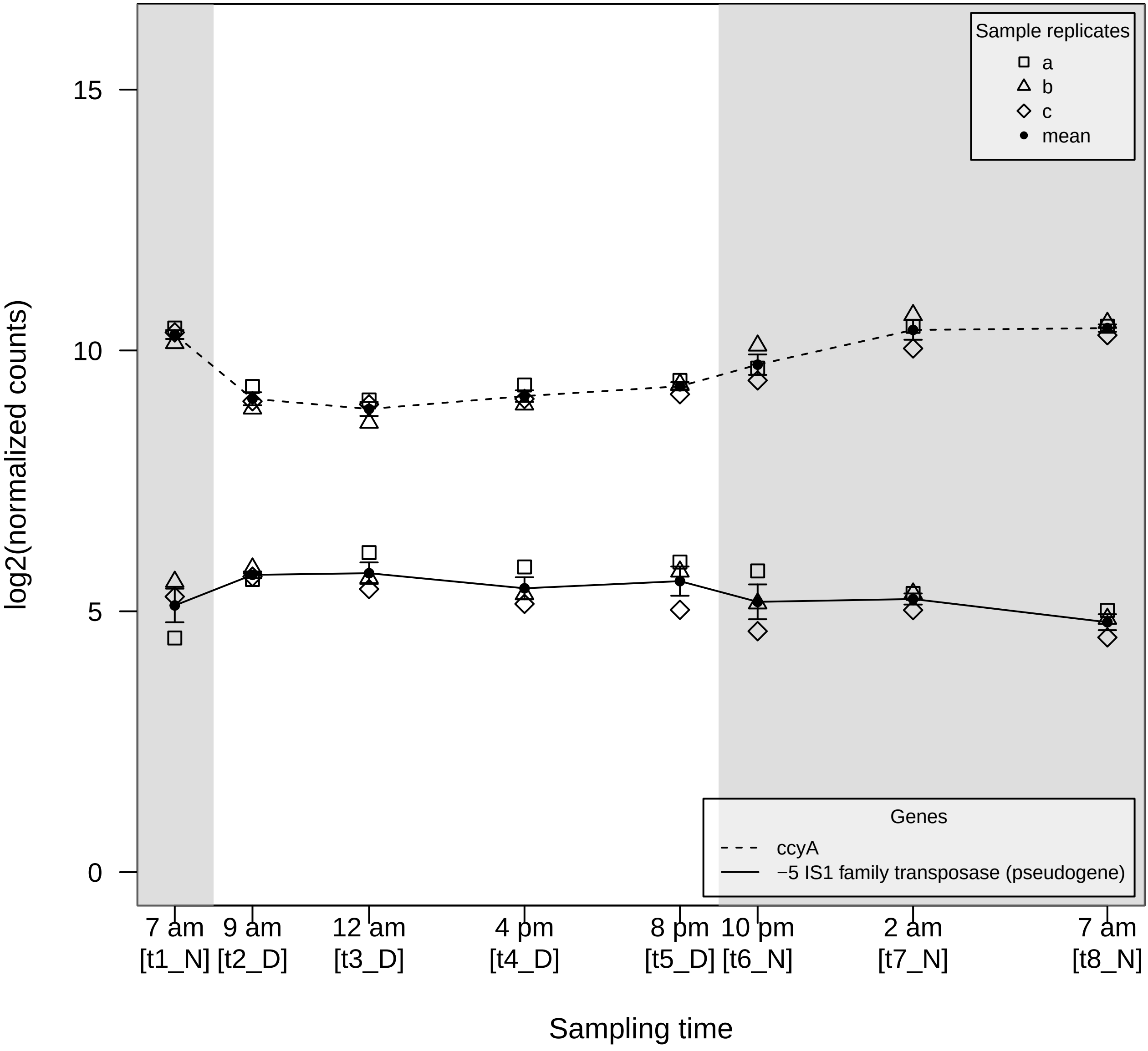

log2(normalized counts)
15
10
5
0
Sample replicates
a
b
c
mean
Genes
ccyA
−5 IS1 family transposase (pseudogene)
7 am [t1_N]
9 am [t2_D]
12 am [t3_D]
4 pm [t4_D]
8 pm [t5_D]
10 pm [t6_N]
2 am [t7_N]
7 am [t8_N]
Sampling time

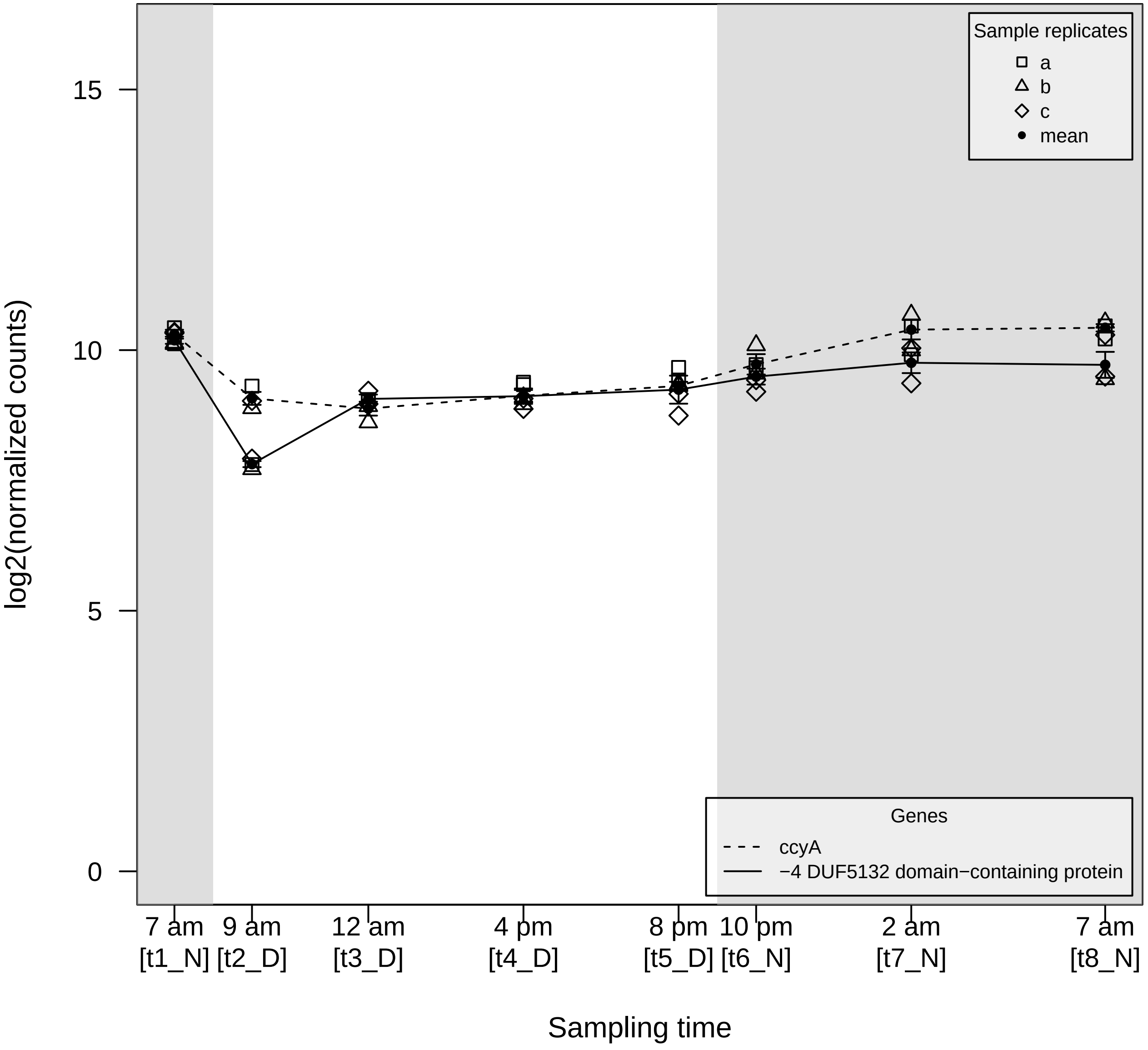

log2(normalized counts)
15
10
5
0
Sample replicates
a
b
c
mean
Genes
ccyA
−4 DUF5132 domain−containing protein
7 am [t1_N]
9 am [t2_D]
12 am [t3_D]
4 pm [t4_D]
8 pm [t5_D]
10 pm [t6_N]
2 am [t7_N]
7 am [t8_N]
Sampling time

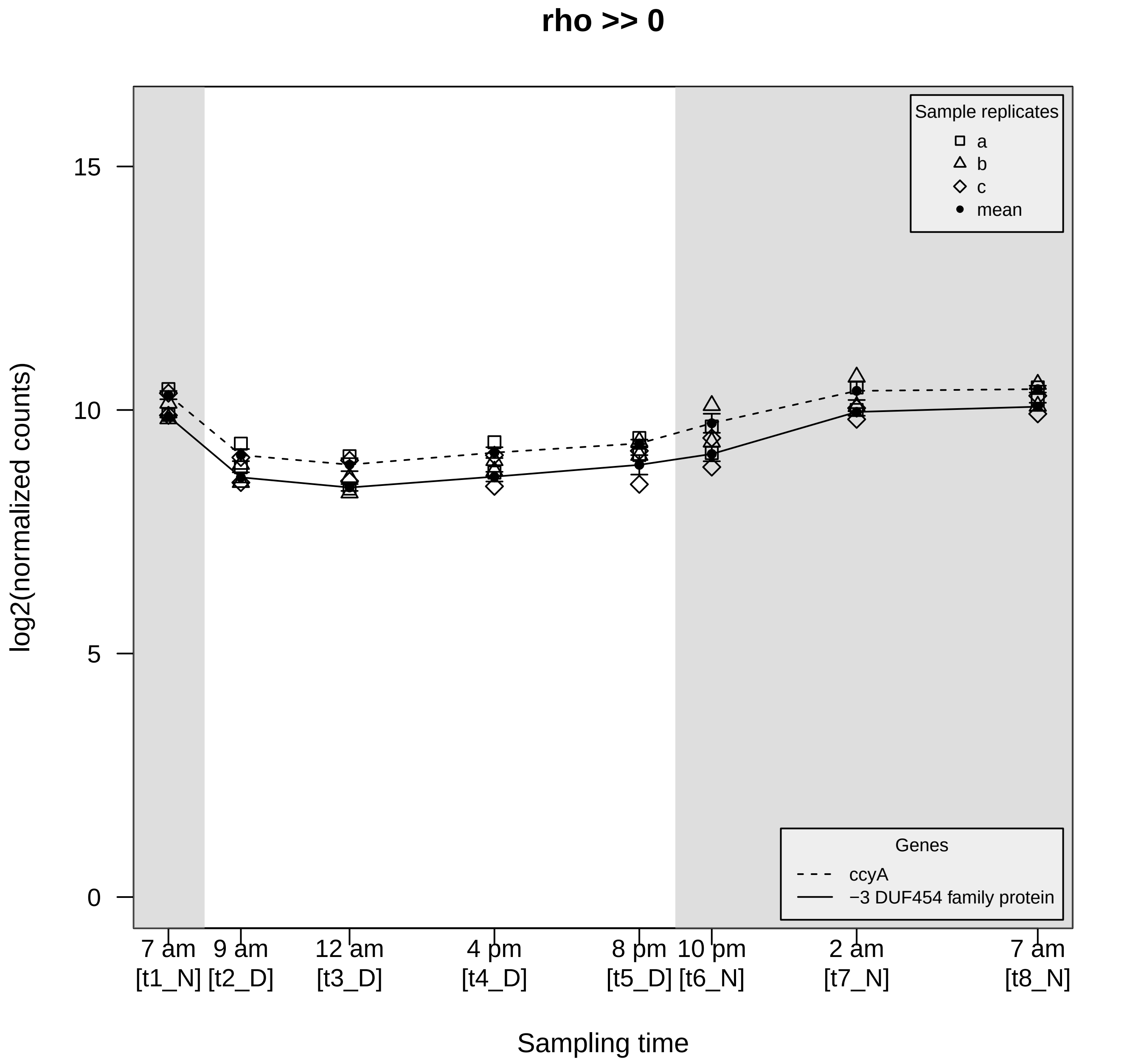
rho >> 0
Sample replicates
a
b
c
mean
Genes
ccyA
−3 DUF454 family protein
log2(normalized counts)
15
10
5
0
7 am
[t1_N]
9 am
[t2_D]
12 am
[t3_D]
4 pm
[t4_D]
8 pm
[t5_D]
10 pm
[t6_N]
2 am
[t7_N]
7 am
[t8_N]
Sampling time

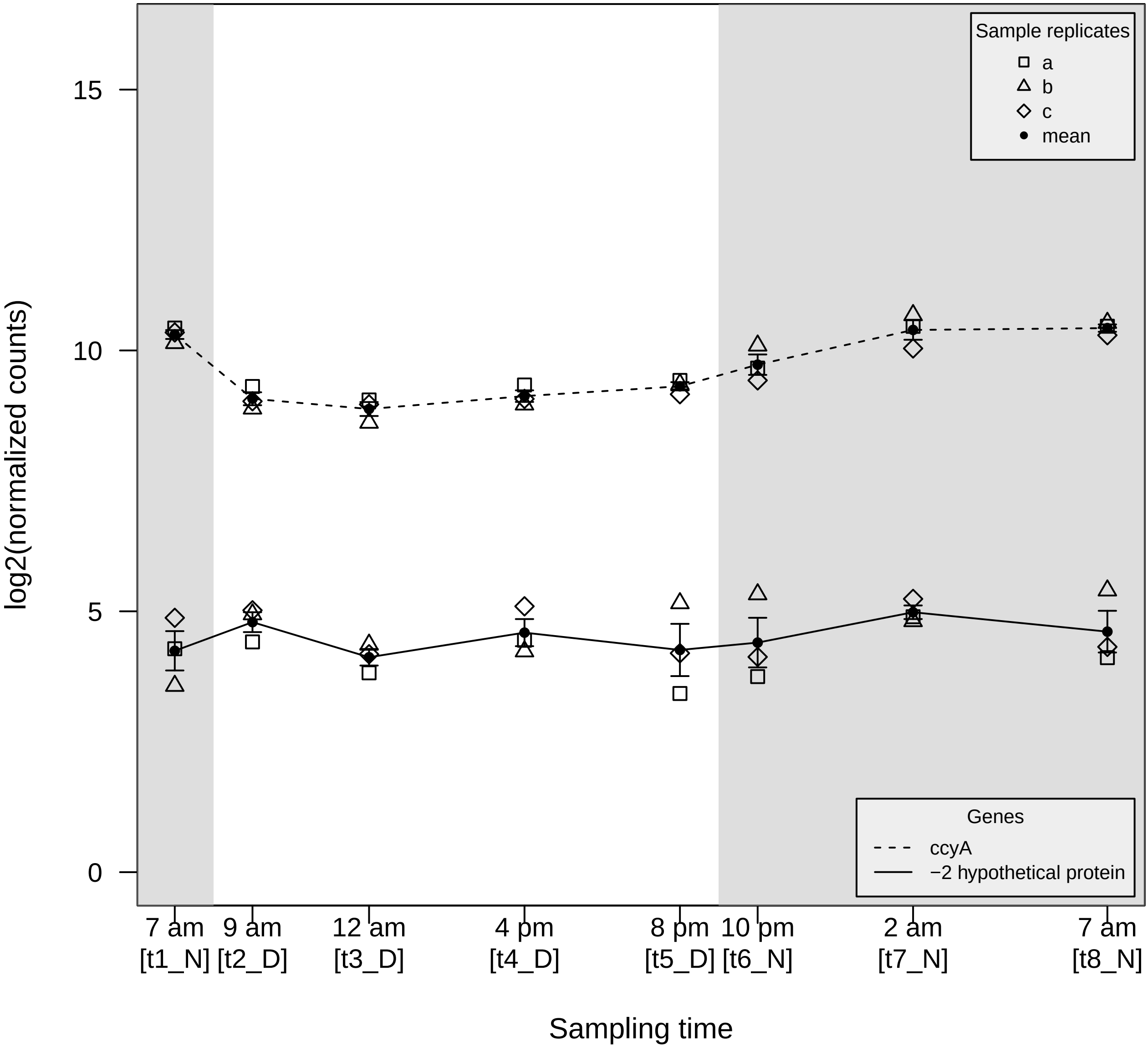

Sample replicates
a
b
c
mean
Genes
ccyA
−2 hypothetical protein
log2(normalized counts)
15
10
5
0
7 am [t1_N]
9 am [t2_D]
12 am [t3_D]
4 pm [t4_D]
8 pm [t5_D]
10 pm [t6_N]
2 am [t7_N]
7 am [t8_N]
Sampling time

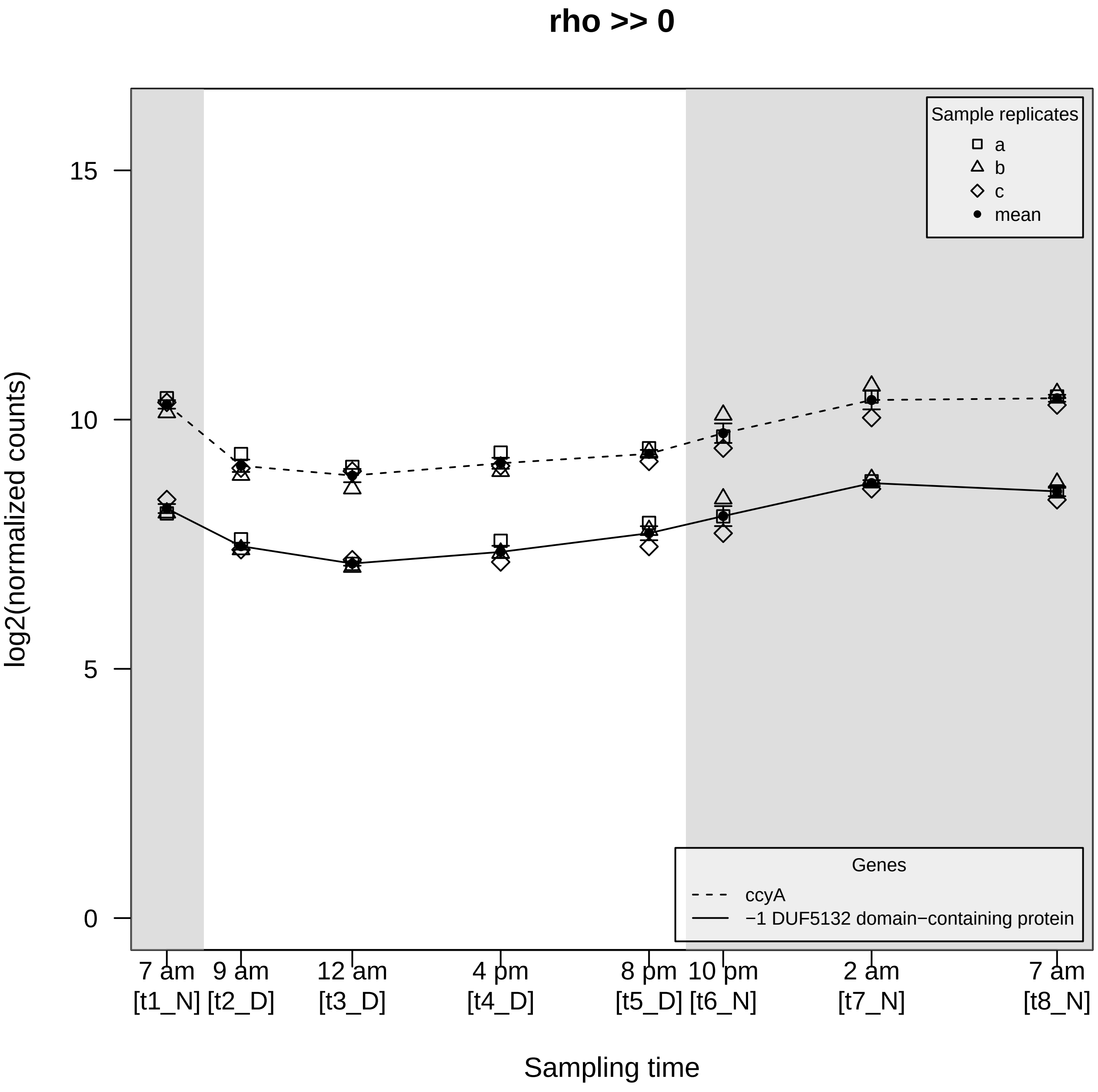

rho >> 0
Sample replicates
a
b
c
mean
Genes
ccyA
−1 DUF5132 domain−containing protein
log2(normalized counts)
15
10
5
0
7 am [t1_N]
9 am [t2_D]
12 am [t3_D]
4 pm [t4_D]
8 pm [t5_D]
10 pm [t6_N]
2 am [t7_N]
7 am [t8_N]
Sampling time

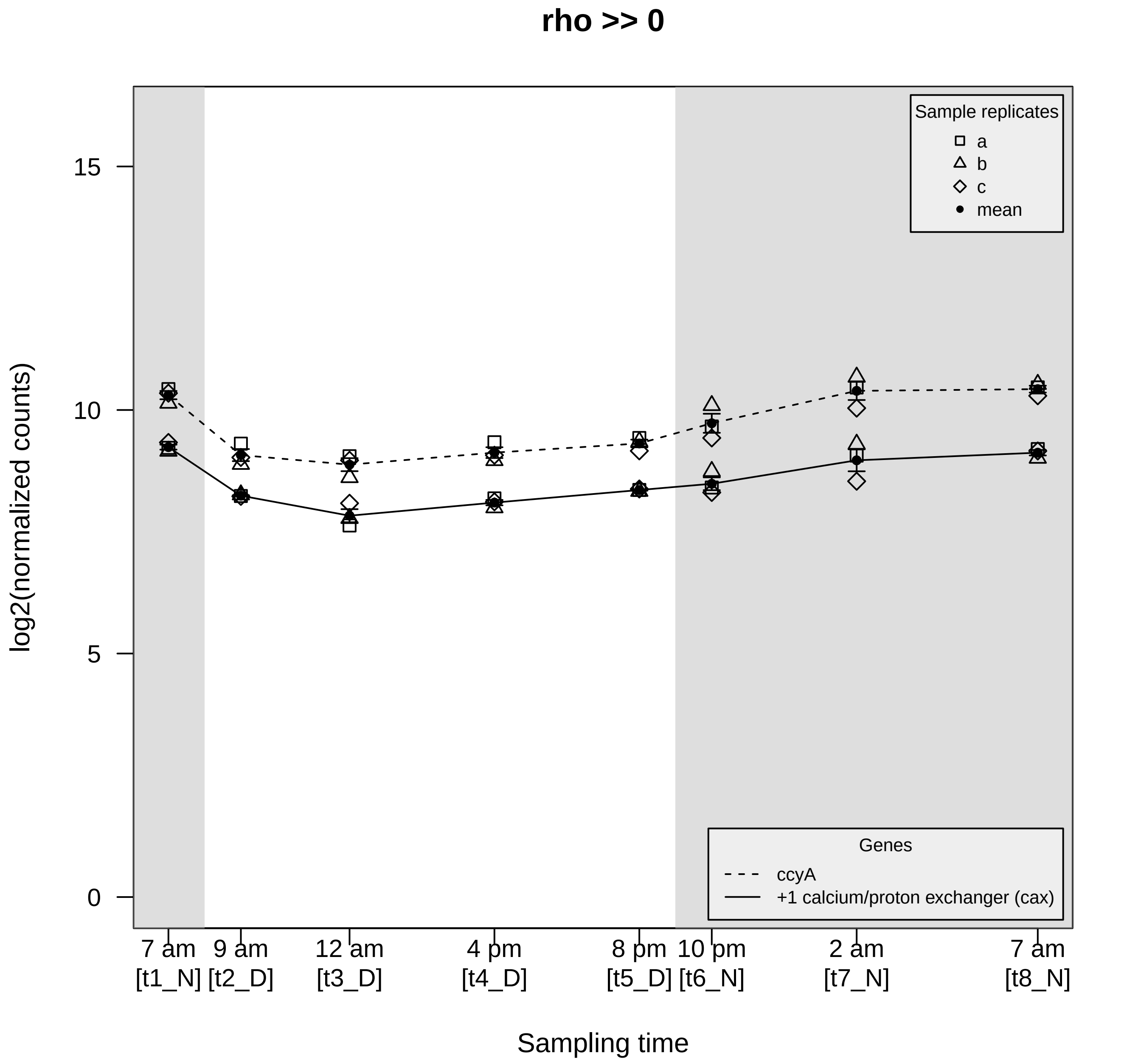
rho >> 0
Sample replicates
a
b
c
mean
Genes
ccyA
+1 calcium/proton exchanger (cax)
log2(normalized counts)
15
10
5
0
7 am [t1_N]
9 am [t2_D]
12 am [t3_D]
4 pm [t4_D]
8 pm [t5_D]
10 pm [t6_N]
2 am [t7_N]
7 am [t8_N]
Sampling time

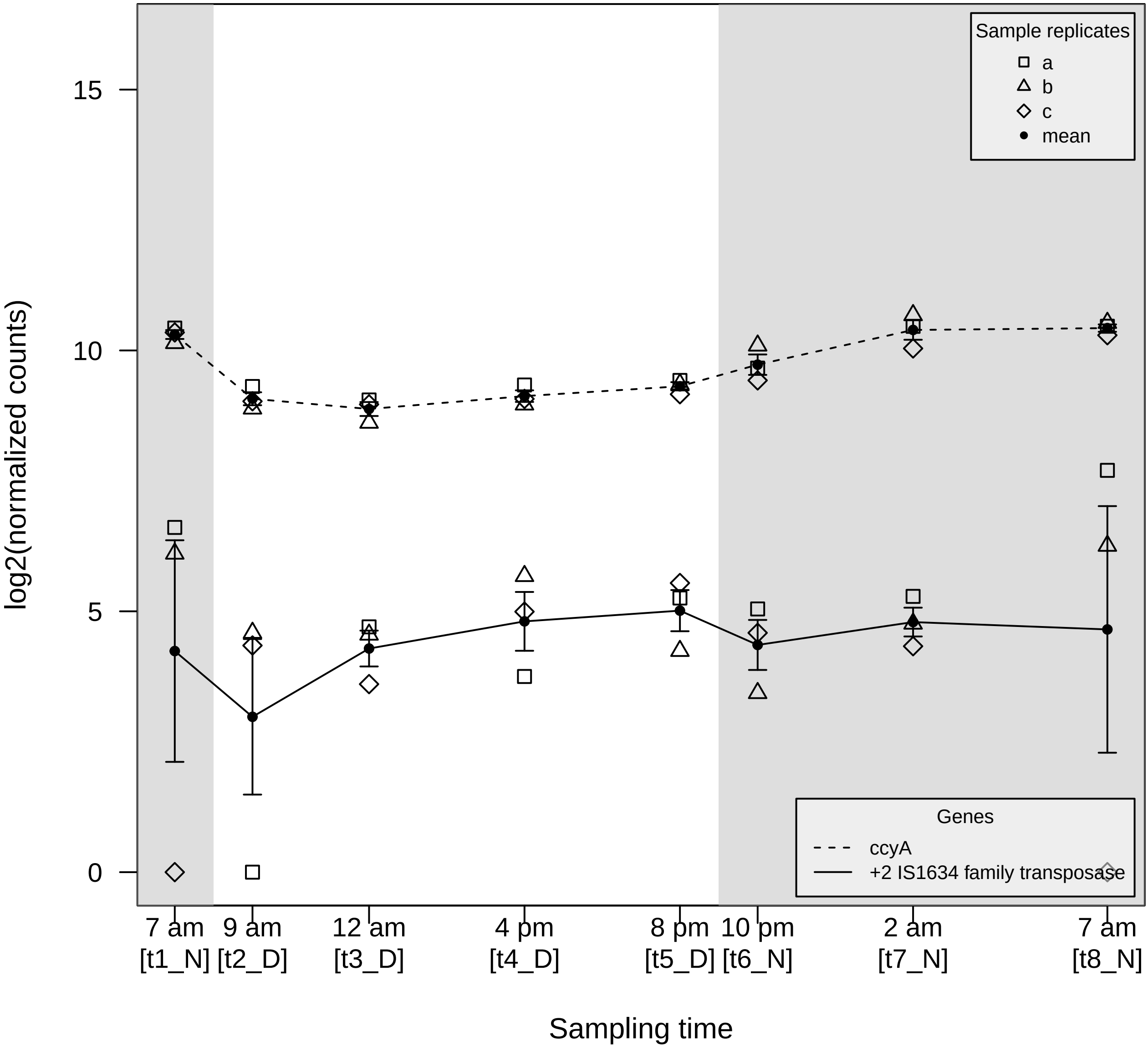

Sample replicates
a
b
c
mean
Genes
ccyA
+2 IS1634 family transposase
log2(normalized counts)
0
5
10
15
7 am [t1_N]
9 am [t2_D]
12 am [t3_D]
4 pm [t4_D]
8 pm [t5_D]
10 pm [t6_N]
2 am [t7_N]
7 am [t8_N]
Sampling time

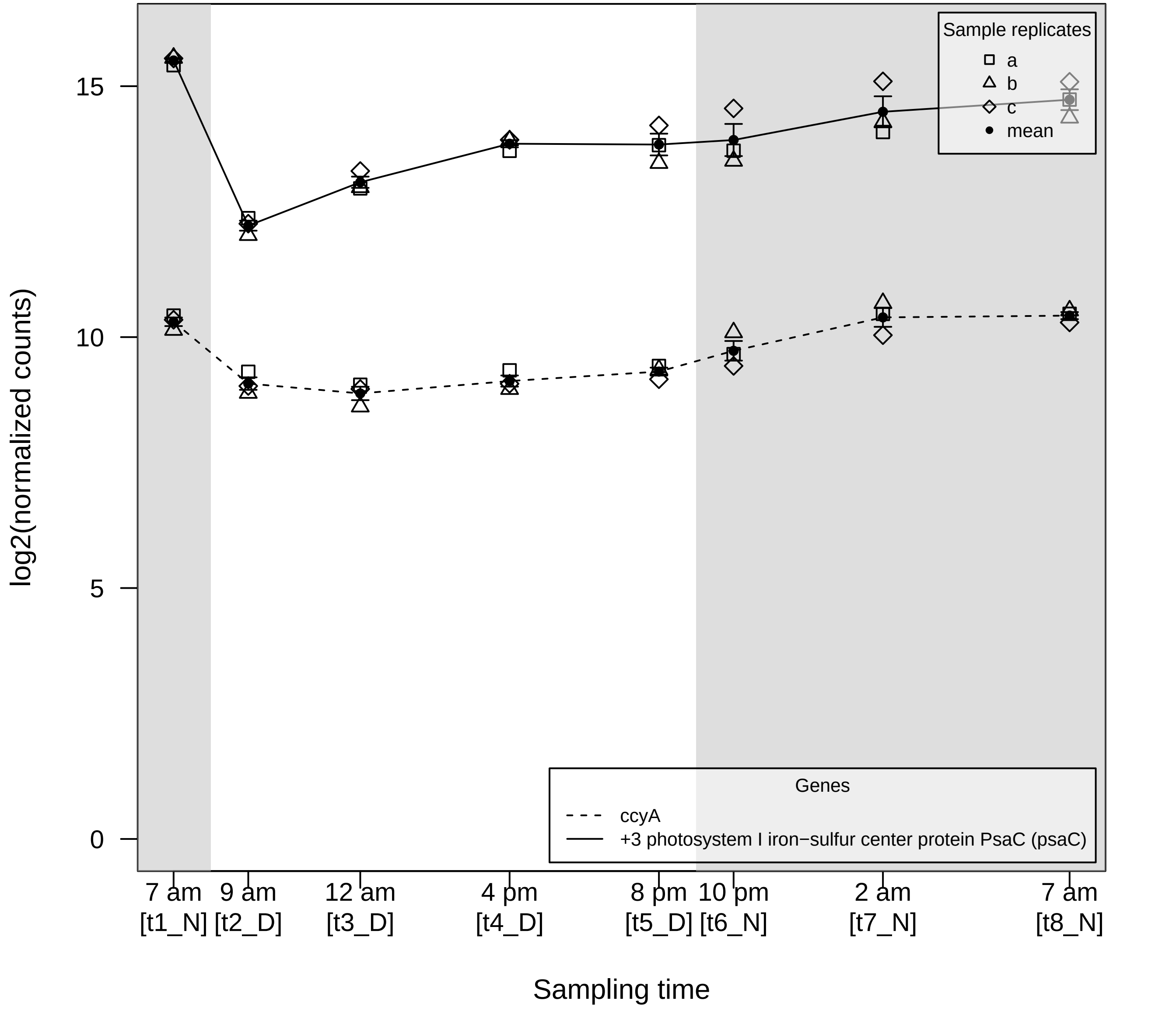

log2(normalized counts)
0
5
10
15
Sample replicates
a
b
c
mean
Genes
ccyA
+3 photosystem I iron−sulfur center protein PsaC (psaC)
7 am [t1_N]
9 am [t2_D]
12 am [t3_D]
4 pm [t4_D]
8 pm [t5_D]
10 pm [t6_N]
2 am [t7_N]
7 am [t8_N]
Sampling time

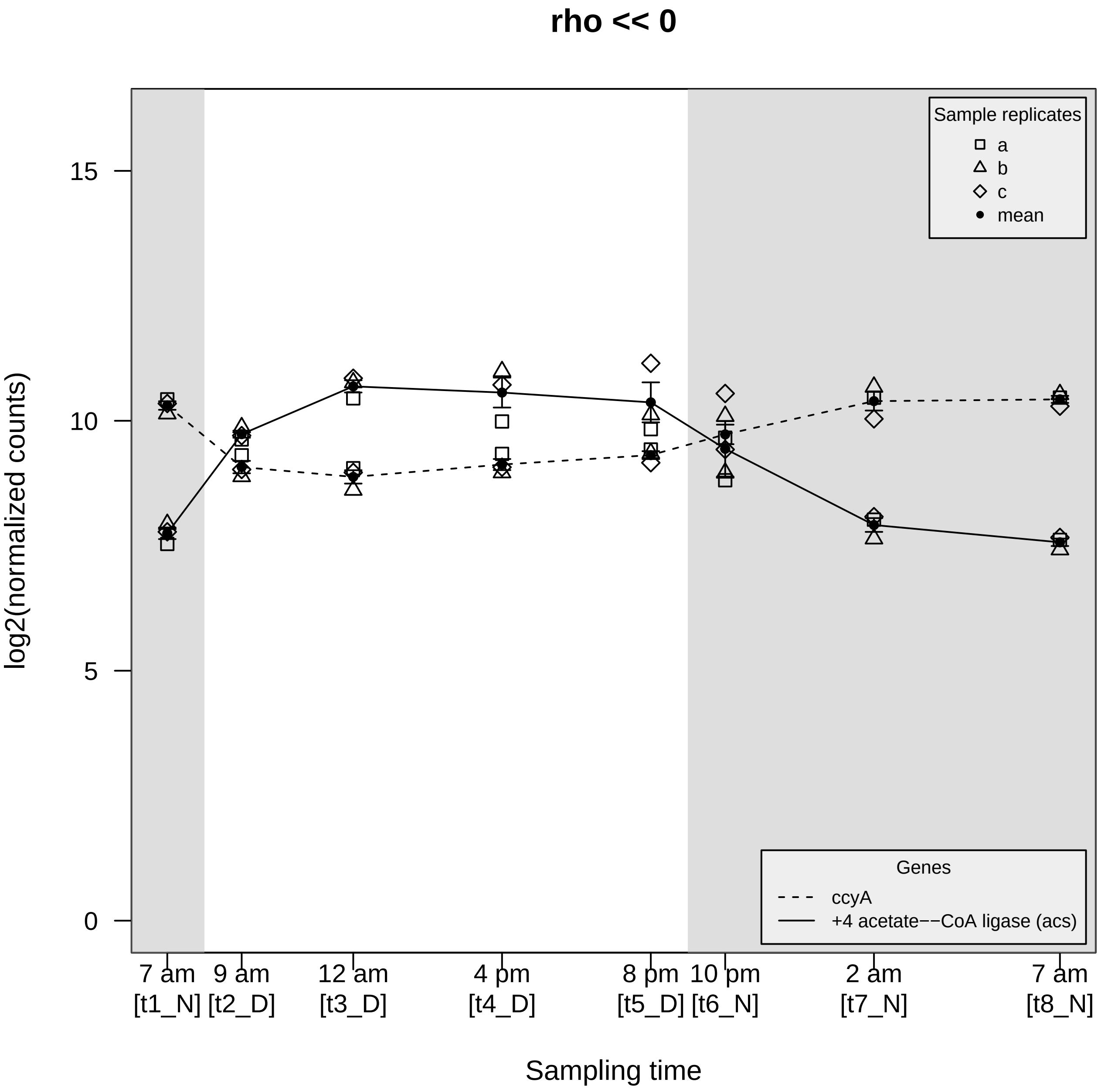

rho << 0
log2(normalized counts)
15
10
5
0
Sample replicates
a
b
c
mean
Genes
ccyA
+4 acetate−−CoA ligase (acs)
7 am [t1_N]
9 am [t2_D]
12 am [t3_D]
4 pm [t4_D]
8 pm [t5_D]
10 pm [t6_N]
2 am [t7_N]
7 am [t8_N]
Sampling time

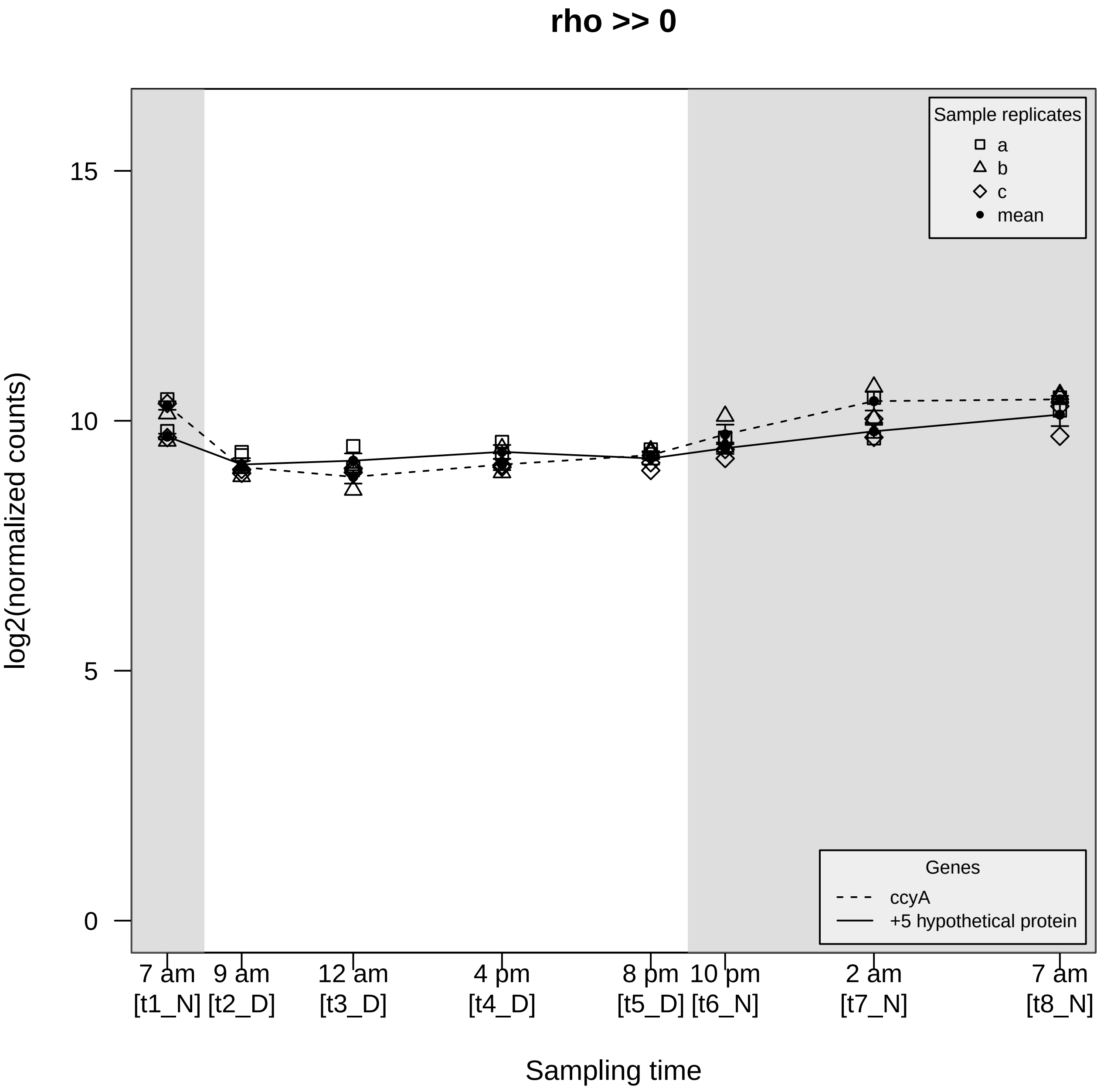

rho >> 0
Sample replicates
a
b
c
mean
Genes
ccyA
+5 hypothetical protein
15
10
5
0
log2(normalized counts)
7 am [t1_N]
9 am [t2_D]
12 am [t3_D]
4 pm [t4_D]
8 pm [t5_D]
10 pm [t6_N]
2 am [t7_N]
7 am [t8_N]
Sampling time

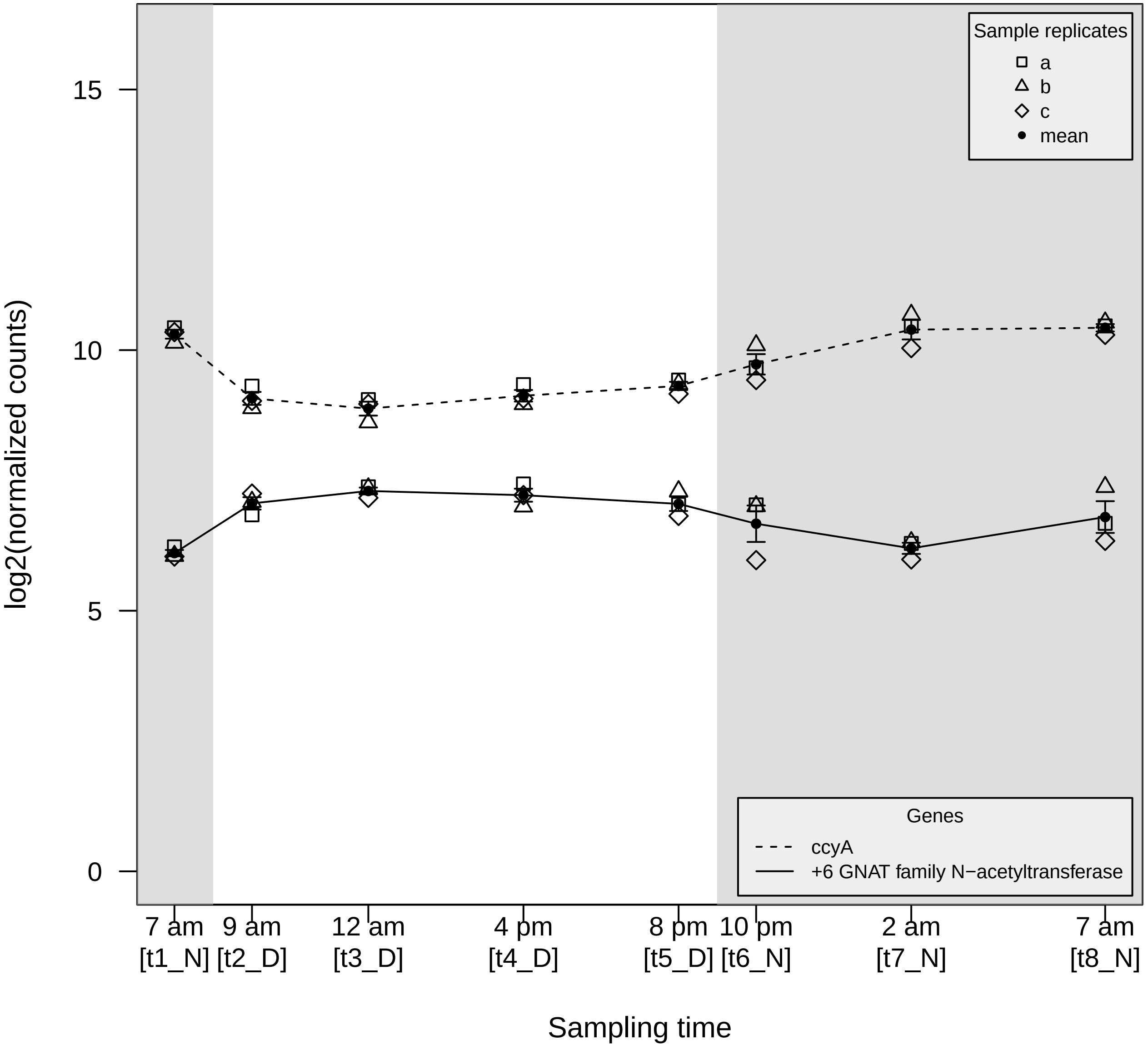

log2(normalized counts)
15
10
5
0
Sample replicates
a
b
c
mean
Genes
ccyA
+6 GNAT family N−acetyltransferase
7 am [t1_N]
9 am [t2_D]
12 am [t3_D]
4 pm [t4_D]
8 pm [t5_D]
10 pm [t6_N]
2 am [t7_N]
7 am [t8_N]
Sampling time

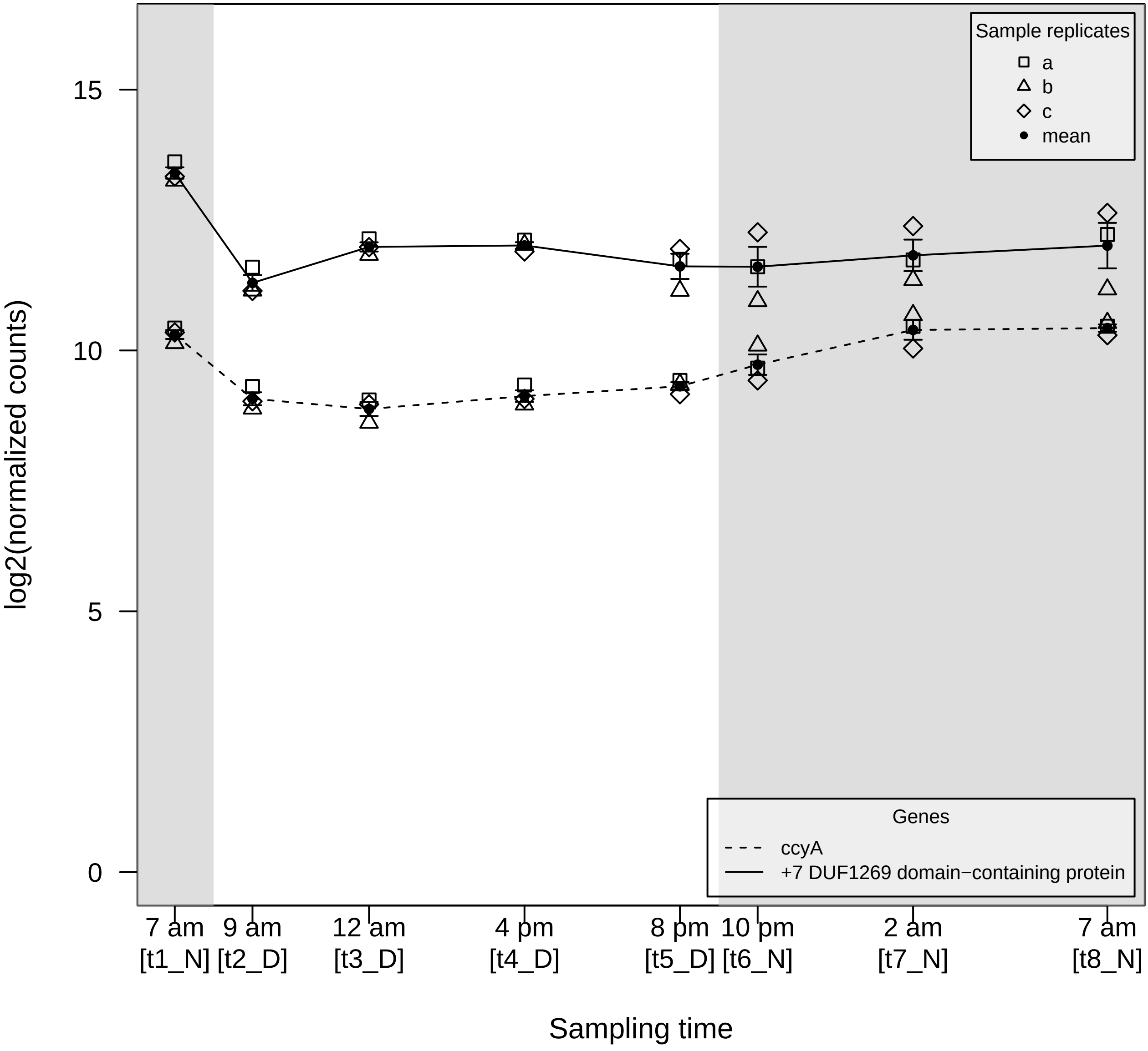

Sample replicates
a
b
c
mean
Genes
ccyA
+7 DUF1269 domain−containing protein
15
10
5
0
log2(normalized counts)
7 am [t1_N]
9 am [t2_D]
12 am [t3_D]
4 pm [t4_D]
8 pm [t5_D]
10 pm [t6_N]
2 am [t7_N]
7 am [t8_N]
Sampling time

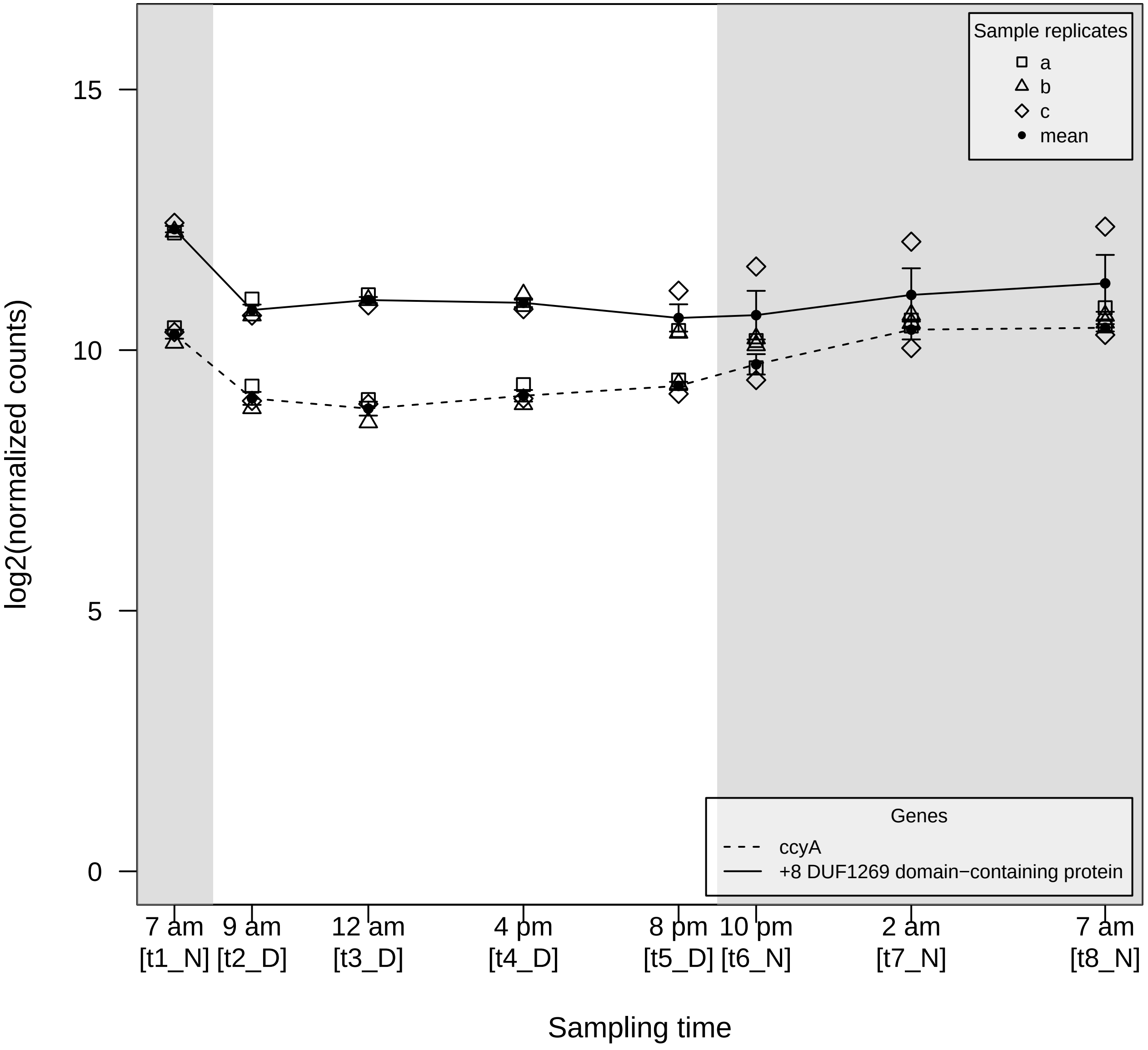

log2(normalized counts)
15
10
5
0
Sample replicates
a
b
c
mean
Genes
ccyA
+8 DUF1269 domain−containing protein
7 am [t1_N]
9 am [t2_D]
12 am [t3_D]
4 pm [t4_D]
8 pm [t5_D]
10 pm [t6_N]
2 am [t7_N]
7 am [t8_N]
Sampling time

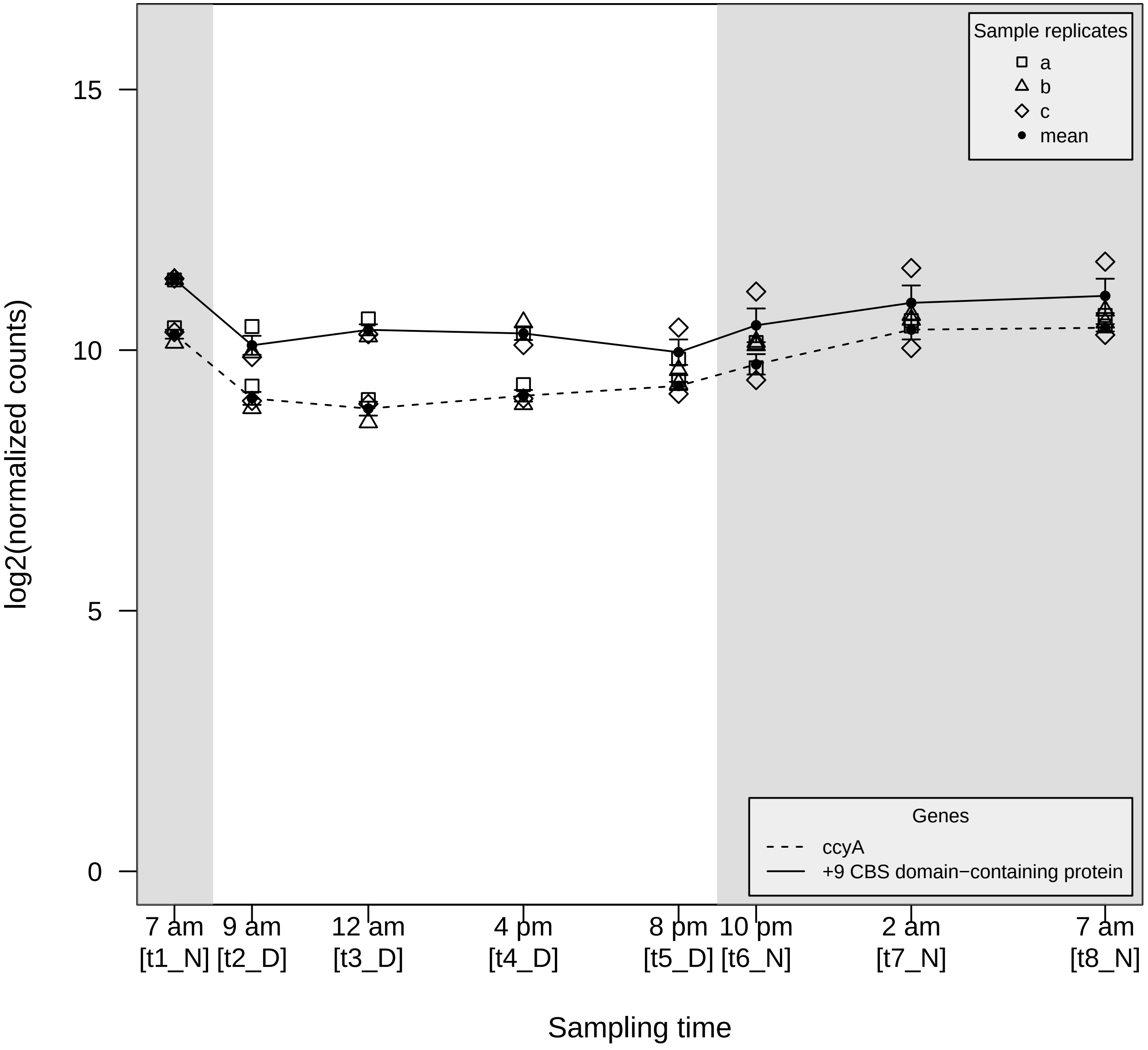

Sample replicates
a
b
c
mean
Genes
ccyA
+9 CBS domain−containing protein
log2(normalized counts)
15
10
5
0
7 am [t1_N]
9 am [t2_D]
12 am [t3_D]
4 pm [t4_D]
8 pm [t5_D]
10 pm [t6_N]
2 am [t7_N]
7 am [t8_N]
Sampling time

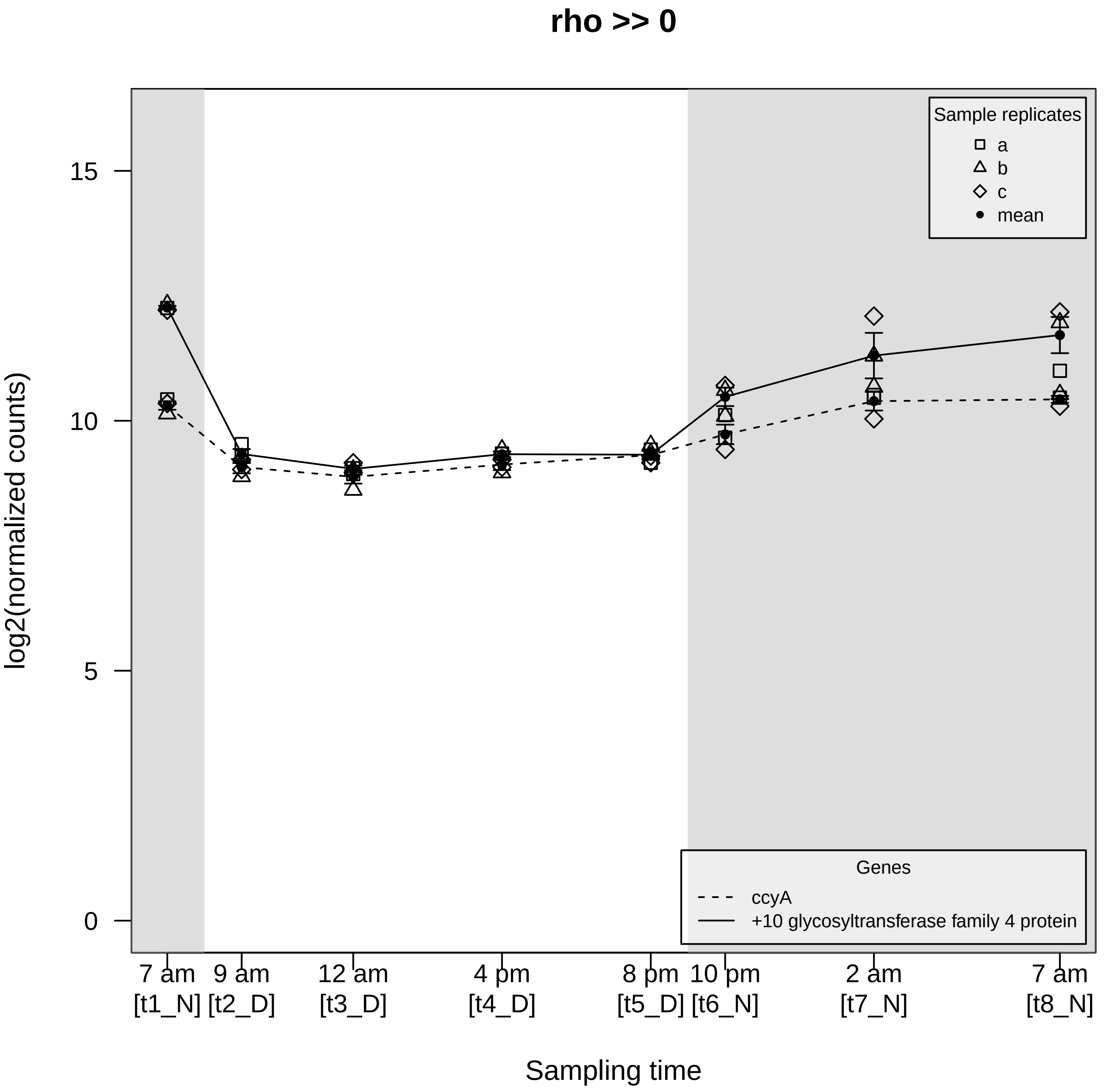
rho >> 0
Sample replicates
a
b
c
mean
15
10
5
0
log2(normalized counts)
Genes
ccyA
+10 glycosyltransferase family 4 protein
7 am [t1_N]
9 am [t2_D]
12 am [t3_D]
4 pm [t4_D]
8 pm [t5_D]
10 pm [t6_N]
2 am [t7_N]
7 am [t8_N]
Sampling time
