## Supplementary_Figures for "Diel changes in the expression of a marker gene and candidate genes for intracellular amorphous CaCO_3_ biomineralization in *Microcystis*": Figure_S4_SI_ccm.pdf

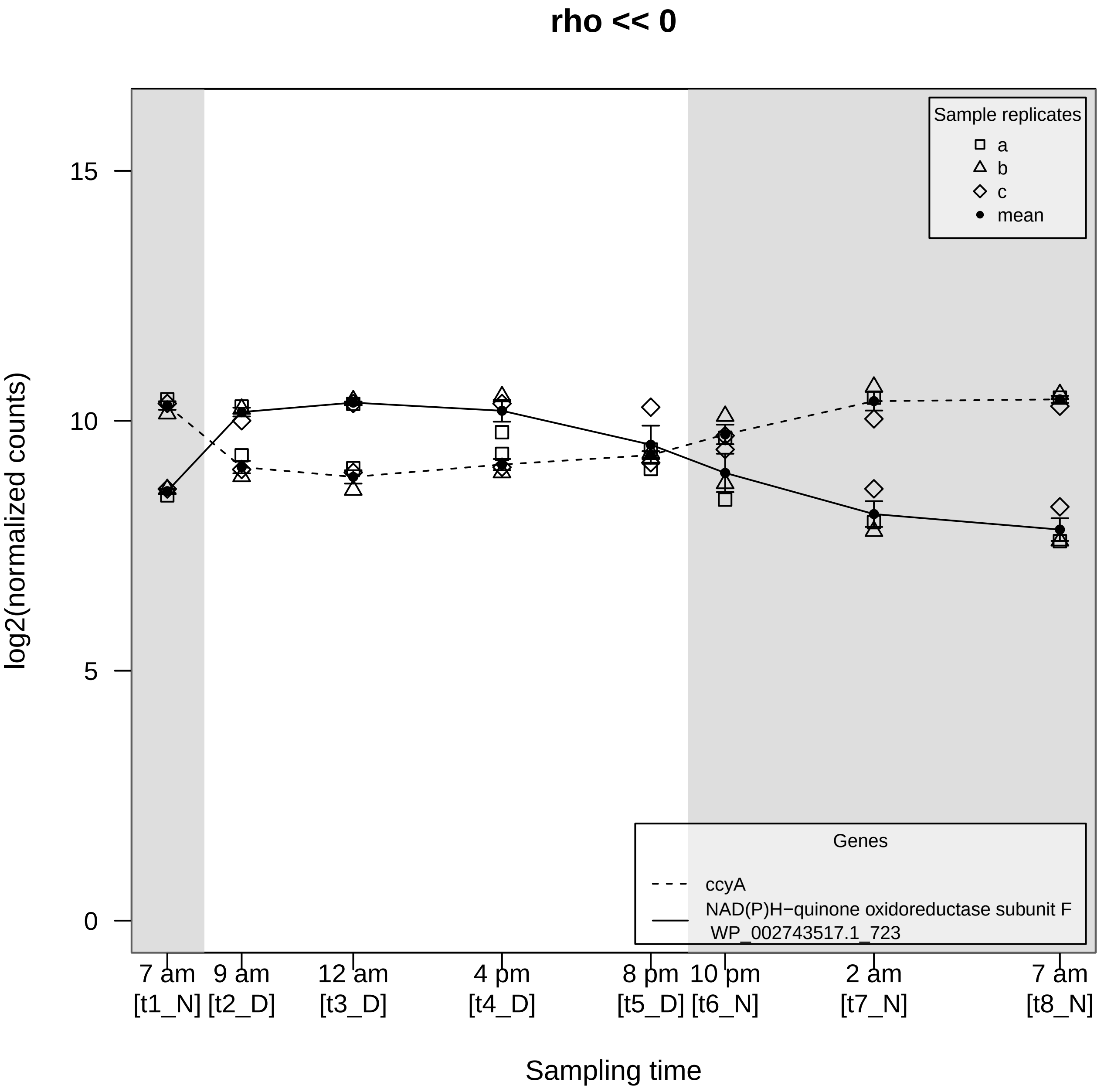

rho << 0
log2(normalized counts)
15
10
5
0
Sample replicates
a
b
c
mean
Genes
ccyA
NAD(P)H−quinone oxidoreductase subunit F
WP_002743517.1_723
7 am [t1_N]
9 am [t2_D]
12 am [t3_D]
4 pm [t4_D]
8 pm [t5_D]
10 pm [t6_N]
2 am [t7_N]
7 am [t8_N]
Sampling time

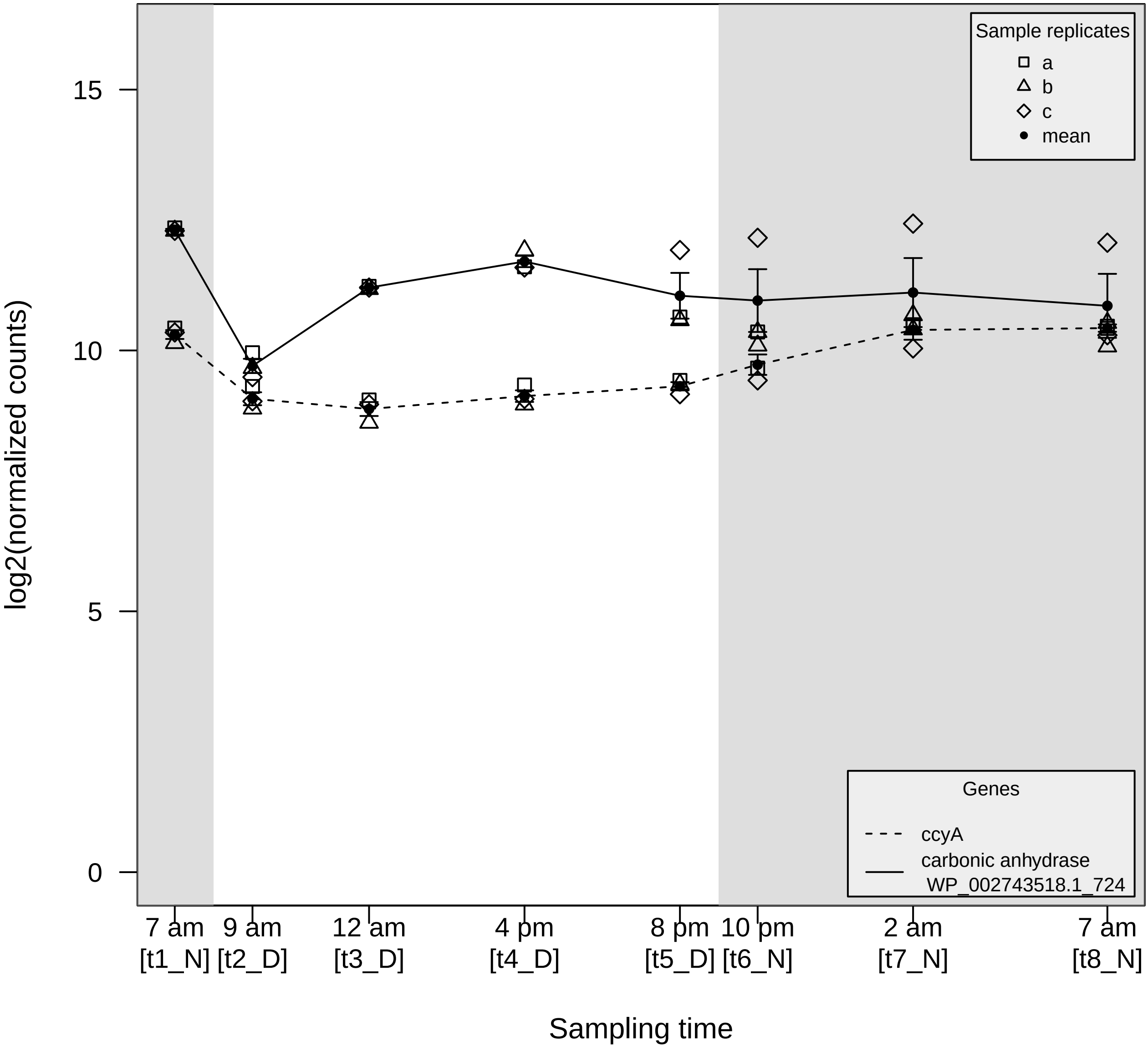

Sample replicates
a
b
c
mean
Genes
ccyA
carbonic anhydrase
WP_002743518.1_724
log2(normalized counts)
15
10
5
0
7 am [t1_N]
9 am [t2_D]
12 am [t3_D]
4 pm [t4_D]
8 pm [t5_D]
10 pm [t6_N]
2 am [t7_N]
7 am [t8_N]
Sampling time

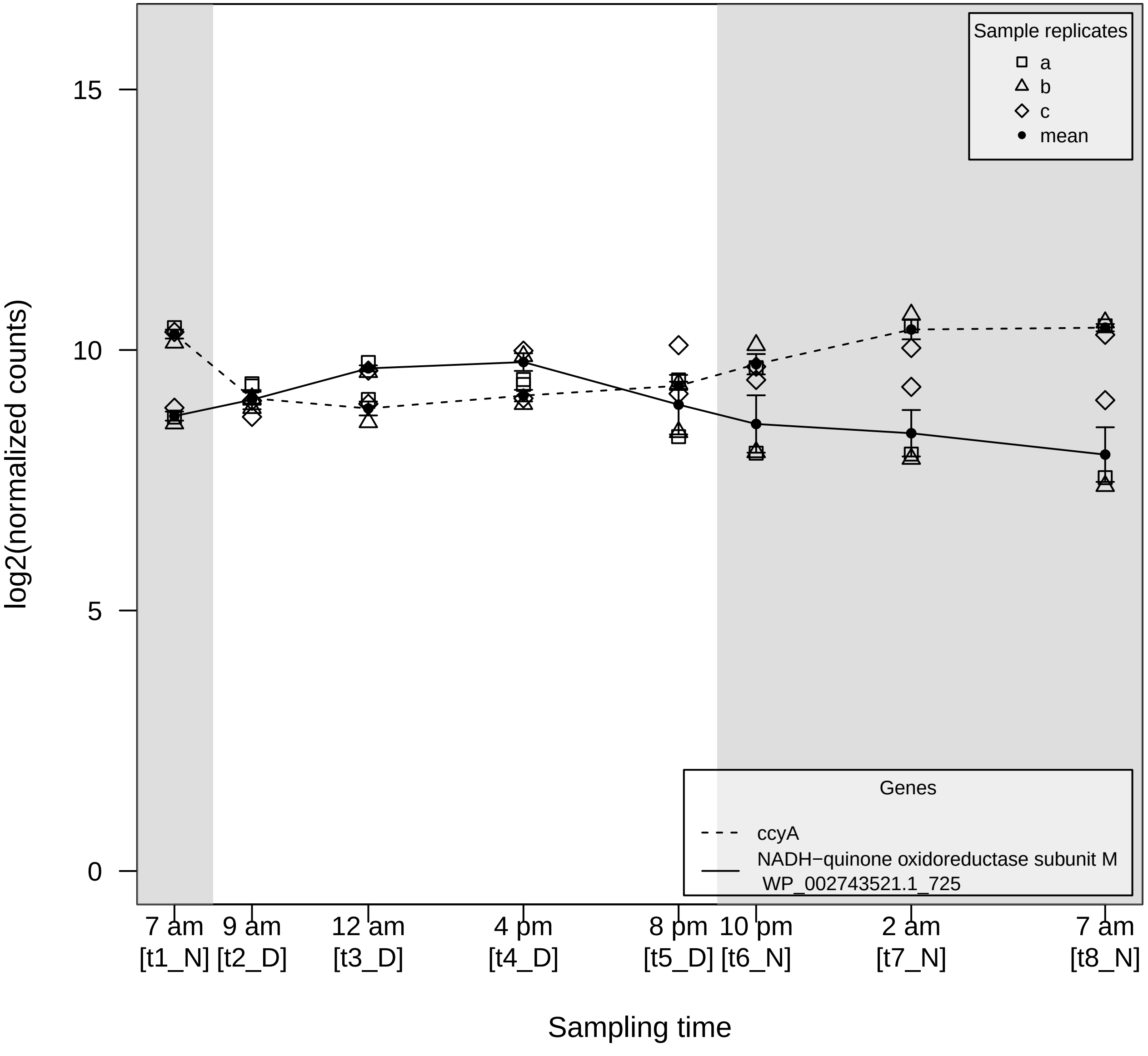

Sample replicates
a
b
c
mean
Genes
ccyA
NADH−quinone oxidoreductase subunit M
WP_002743521.1_725
log2(normalized counts)
15
10
5
0
7 am [t1_N]
9 am [t2_D]
12 am [t3_D]
4 pm [t4_D]
8 pm [t5_D]
10 pm [t6_N]
2 am [t7_N]
7 am [t8_N]
Sampling time

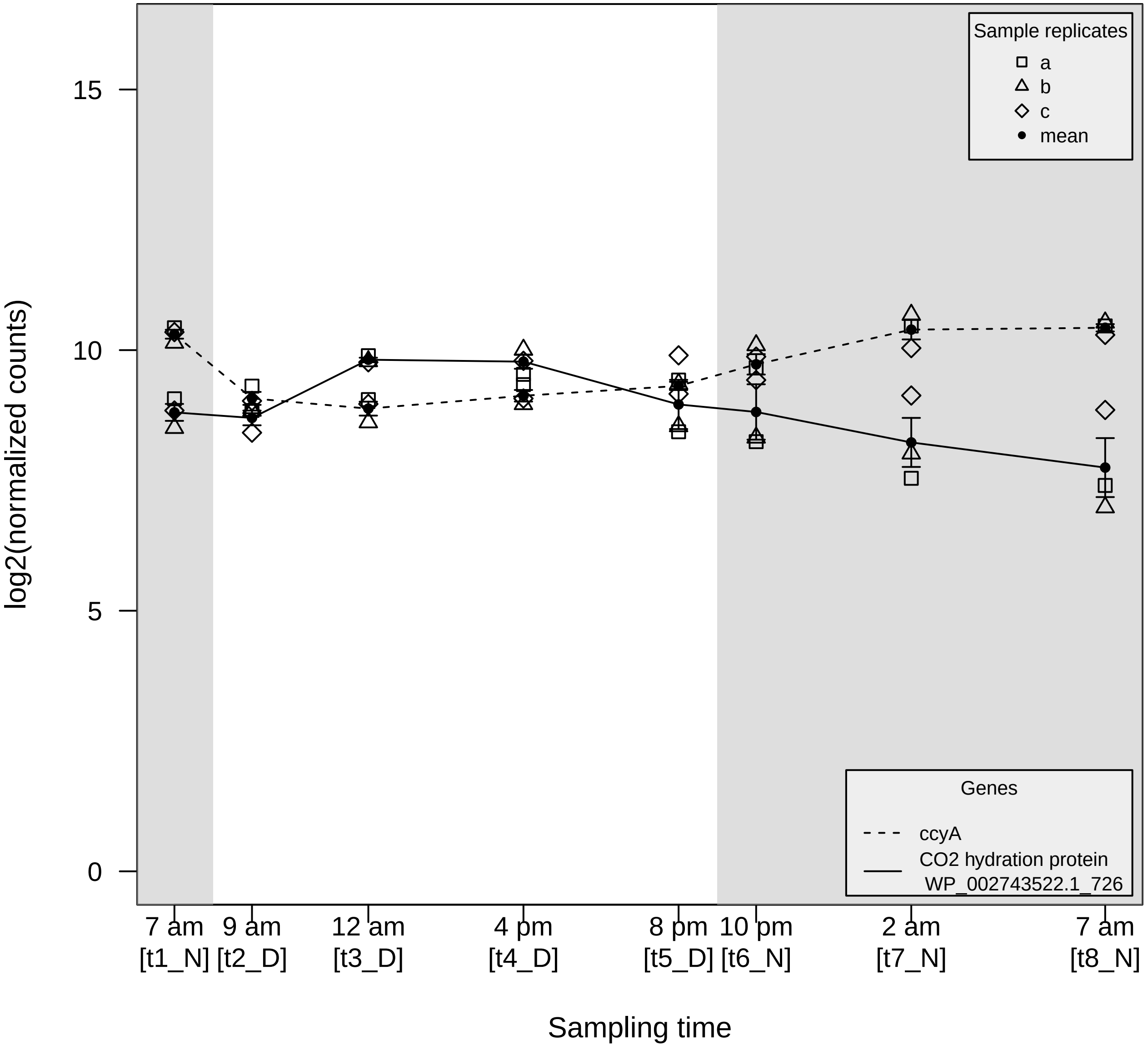

Sample replicates
a
b
c
mean
Genes
ccyA
CO2 hydration protein
WP_002743522.1_726
15
10
5
0
log2(normalized counts)
7 am [t1_N]
9 am [t2_D]
12 am [t3_D]
4 pm [t4_D]
8 pm [t5_D]
10 pm [t6_N]
2 am [t7_N]
7 am [t8_N]
Sampling time

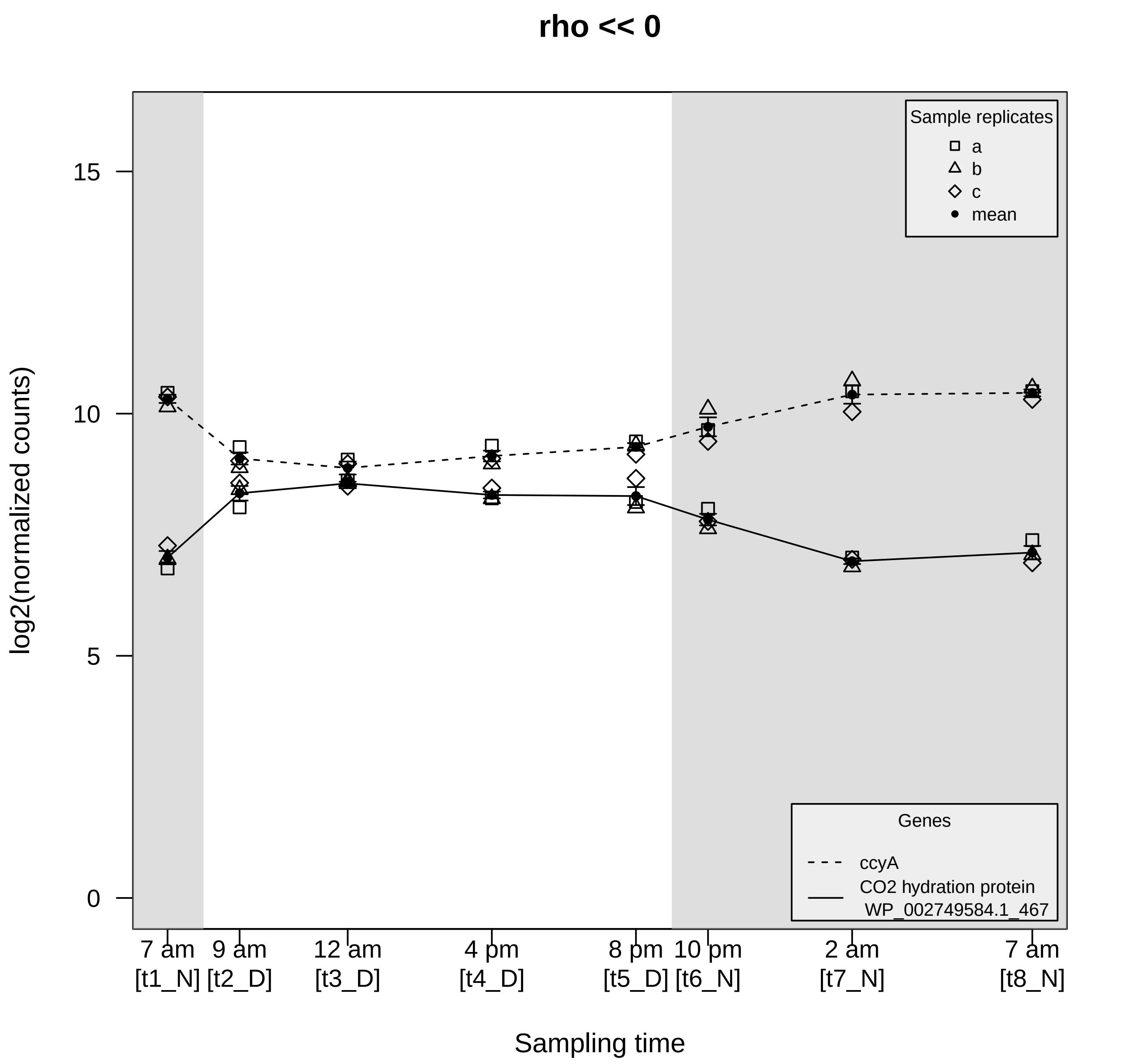
rho << 0
Sample replicates
a
b
c
mean
Genes
ccyA
CO2 hydration protein
WP_002749584.1_467
log2(normalized counts)
15
10
5
0
7 am [t1_N]
9 am [t2_D]
12 am [t3_D]
4 pm [t4_D]
8 pm [t5_D]
10 pm [t6_N]
2 am [t7_N]
7 am [t8_N]
Sampling time

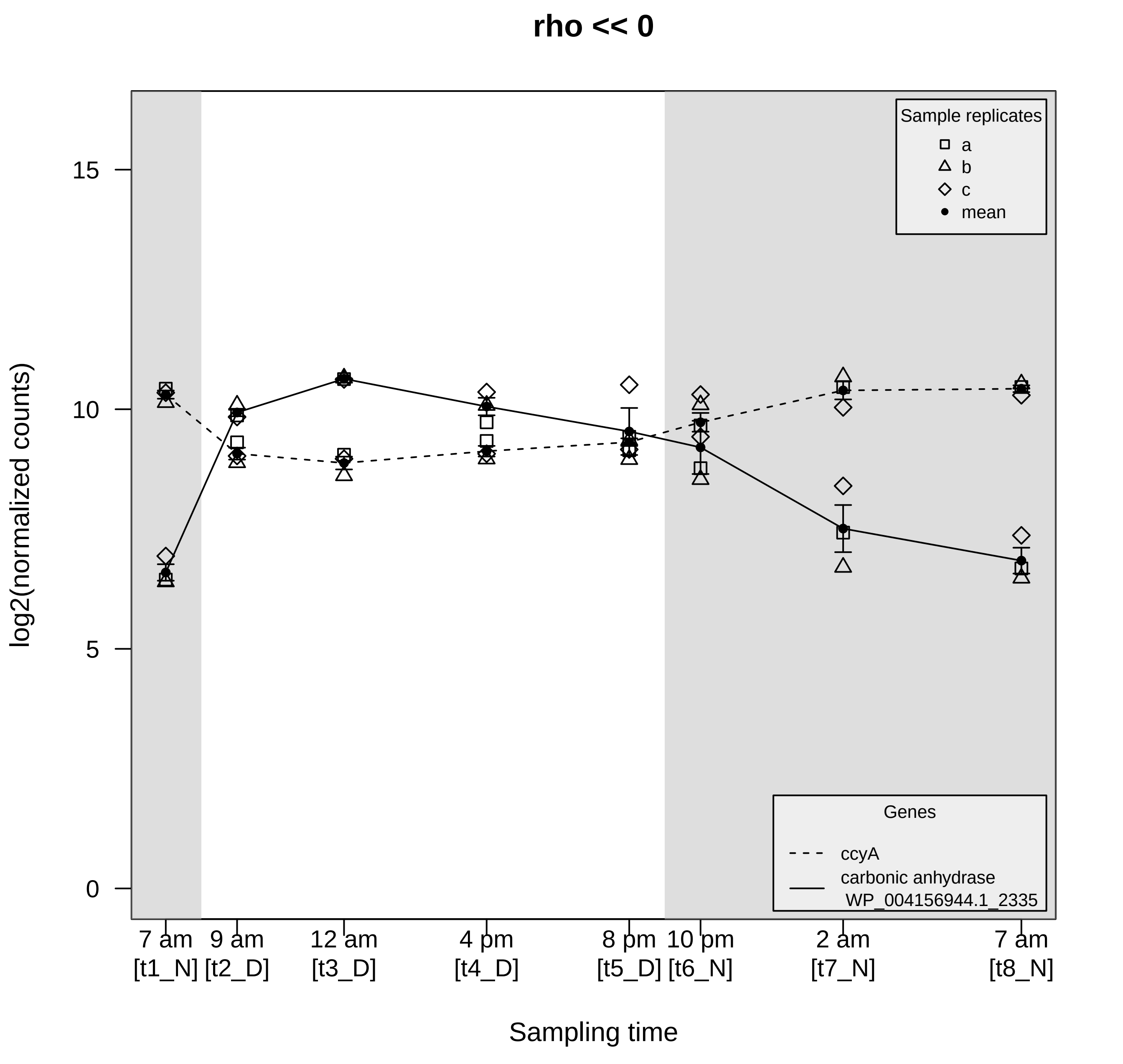

rho << 0
Sample replicates
a
b
c
mean
Genes
ccyA
carbonic anhydrase
WP_004156944.1_2335
log2(normalized counts)
15
10
5
0
7 am
[t1_N]
9 am
[t2_D]
12 am
[t3_D]
4 pm
[t4_D]
8 pm
[t5_D]
10 pm
[t6_N]
2 am
[t7_N]
7 am
[t8_N]
Sampling time

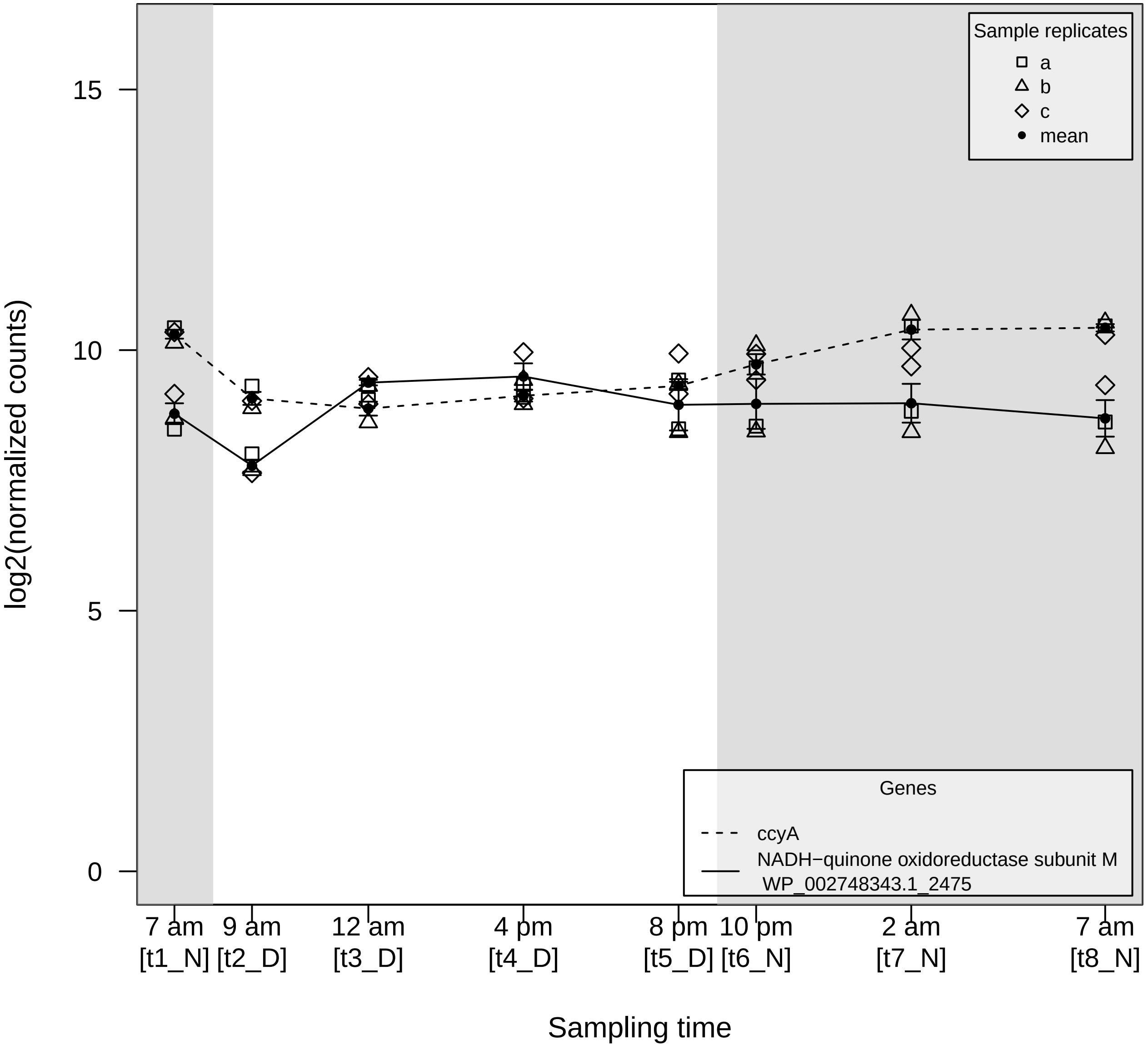
log2(normalized counts)
15
10
5
0
Sample replicates
a
b
c
mean
Genes
ccyA
NADH−quinone oxidoreductase subunit M
WP_002748343.1_2475
7 am
[t1_N]
9 am
[t2_D]
12 am
[t3_D]
4 pm
[t4_D]
8 pm
[t5_D]
10 pm
[t6_N]
2 am
[t7_N]
7 am
[t8_N]
Sampling time

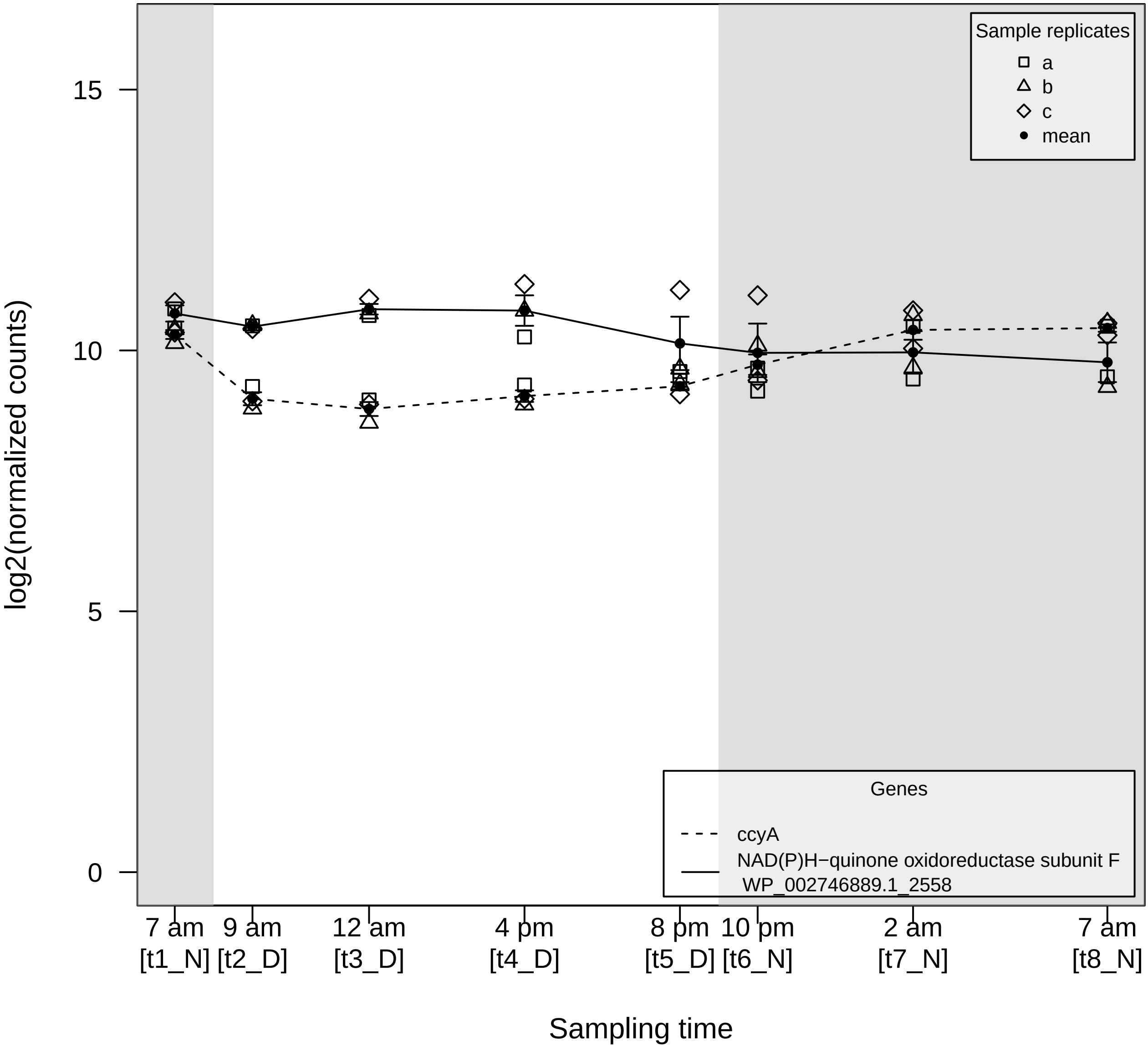
Sample replicates
a
b
c
mean
15
10
5
0
log2(normalized counts)
Genes
ccyA
NAD(P)H−quinone oxidoreductase subunit F
WP_002746889.1_2558
7 am [t1_N]
9 am [t2_D]
12 am [t3_D]
4 pm [t4_D]
8 pm [t5_D]
10 pm [t6_N]
2 am [t7_N]
7 am [t8_N]
Sampling time

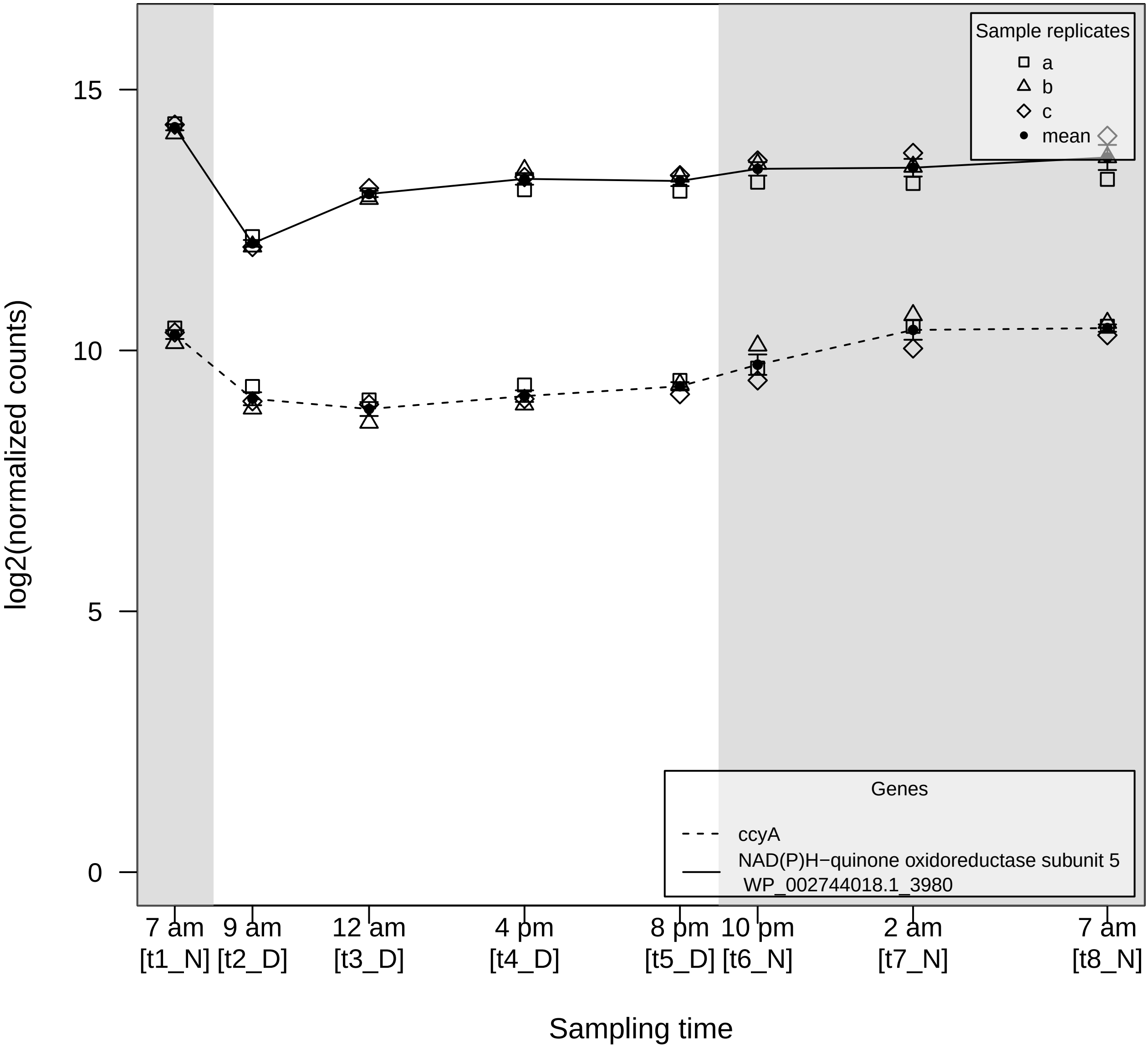

Sample replicates
a
b
c
mean
Genes
ccyA
NAD(P)H−quinone oxidoreductase subunit 5
WP_002744018.1_3980
log2(normalized counts)
15
10
5
0
7 am [t1_N]
9 am [t2_D]
12 am [t3_D]
4 pm [t4_D]
8 pm [t5_D]
10 pm [t6_N]
2 am [t7_N]
7 am [t8_N]
Sampling time

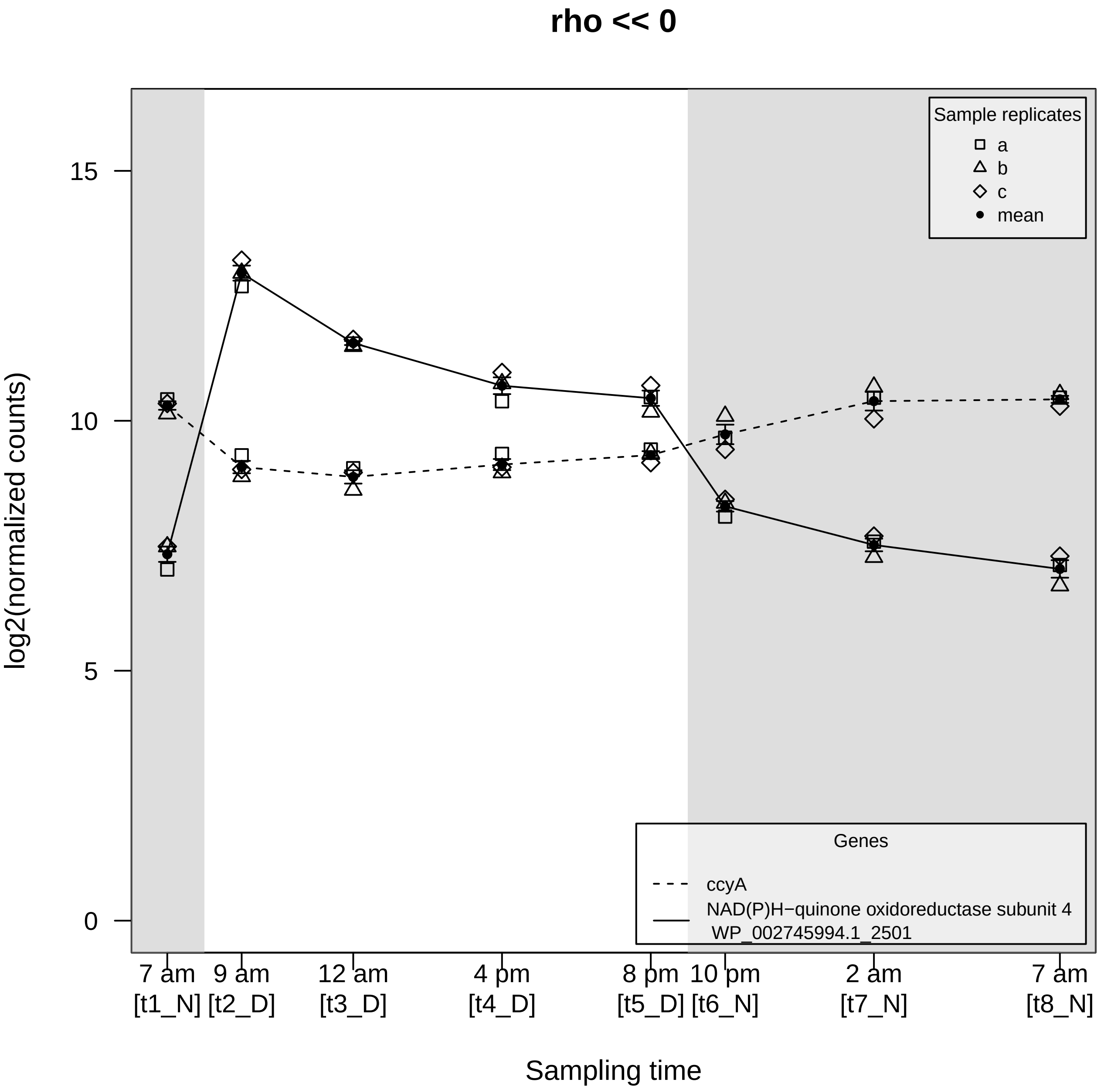

rho << 0
Sample replicates
a
b
c
mean
Genes
ccyA
NAD(P)H−quinone oxidoreductase subunit 4
WP_002745994.1_2501
log2(normalized counts)
15
10
5
0
7 am [t1_N]
9 am [t2_D]
12 am [t3_D]
4 pm [t4_D]
8 pm [t5_D]
10 pm [t6_N]
2 am [t7_N]
7 am [t8_N]
Sampling time

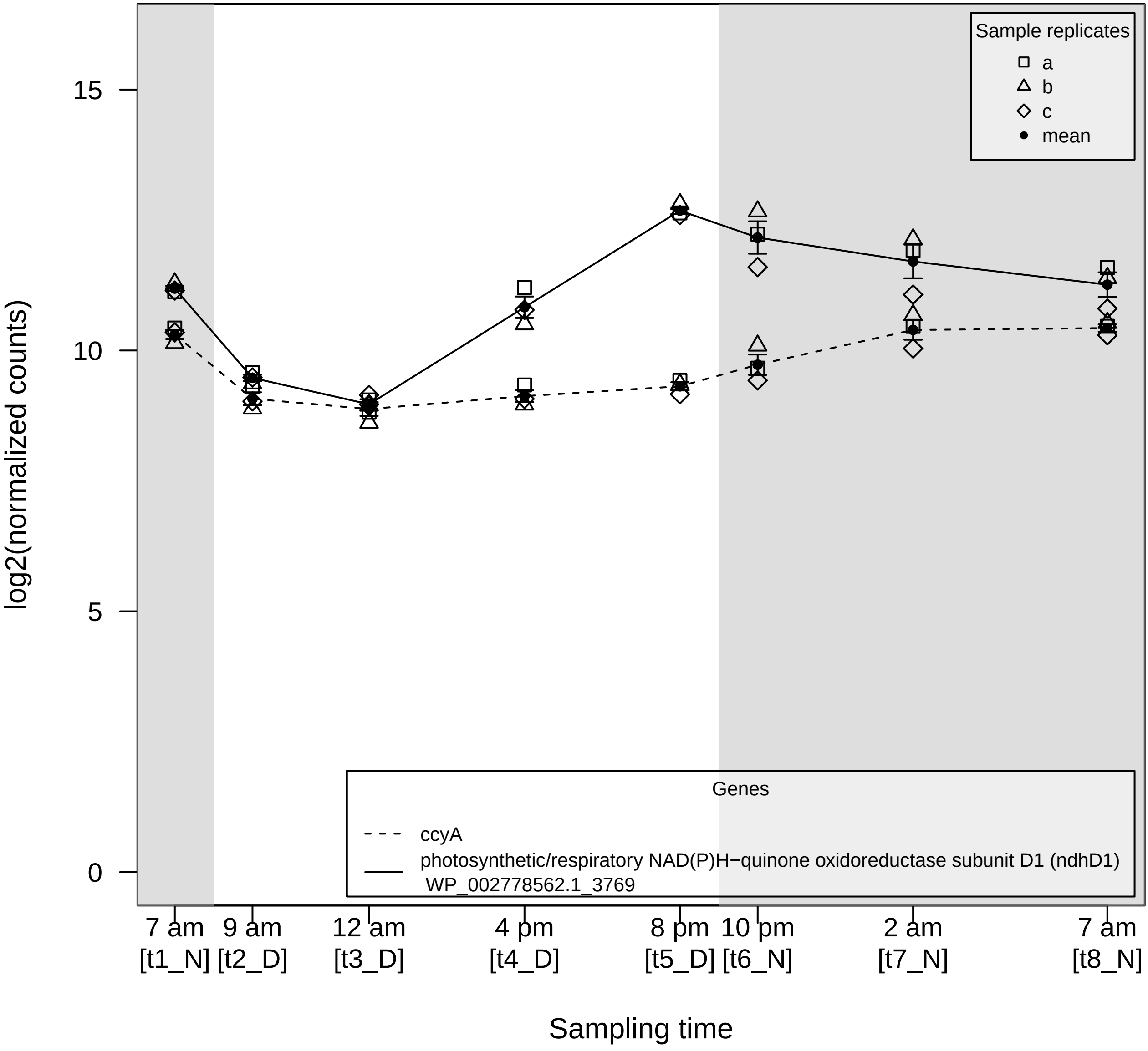
Sample replicates
a
b
c
mean
Genes
ccyA
photosynthetic/respiratory NAD(P)H−quinone oxidoreductase subunit D1 (ndhD1)
WP_002778562.1_3769
log2(normalized counts)
15
10
5
0
7 am [t1_N]
9 am [t2_D]
12 am [t3_D]
4 pm [t4_D]
8 pm [t5_D]
10 pm [t6_N]
2 am [t7_N]
7 am [t8_N]
Sampling time

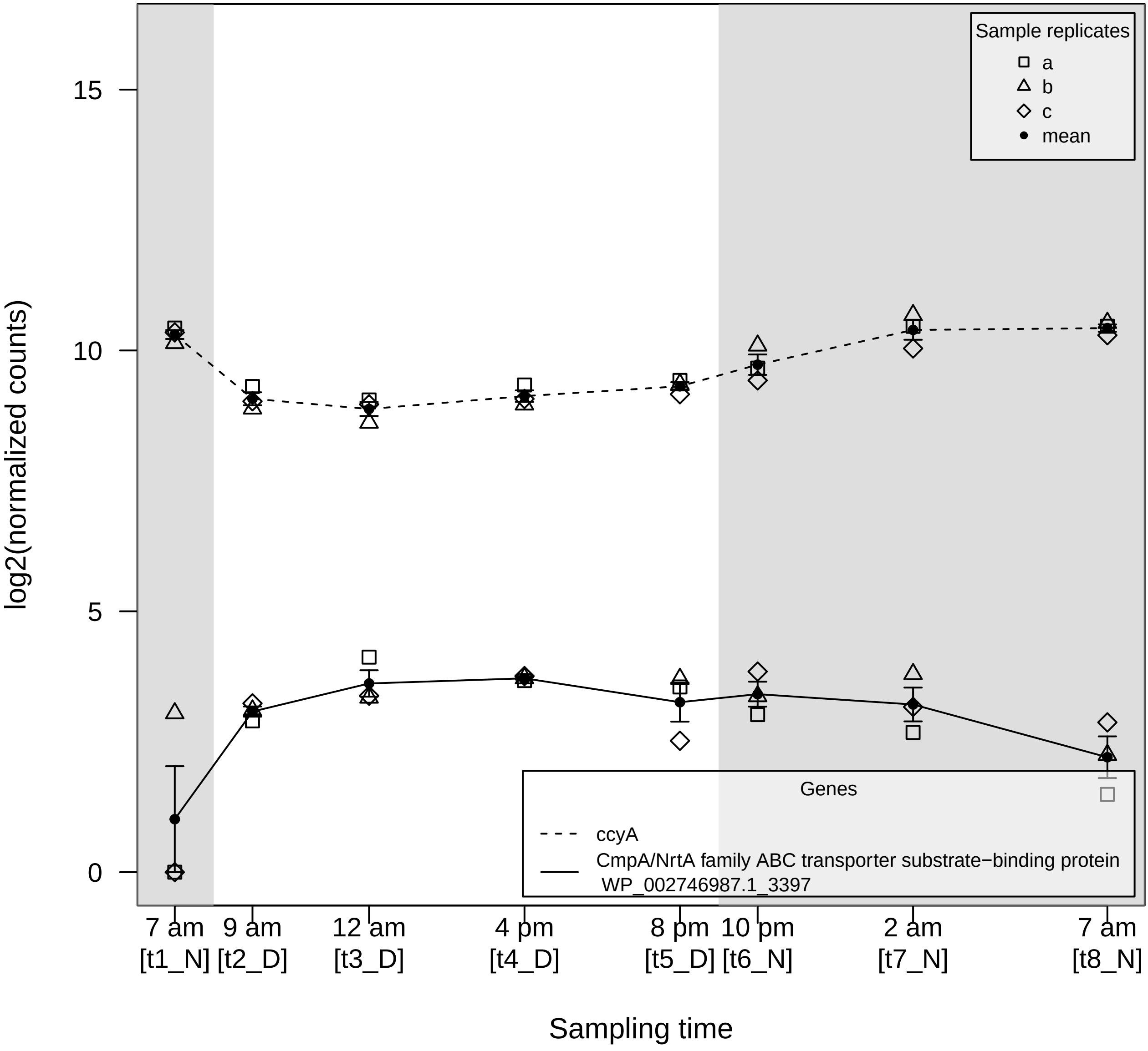

Sample replicates
a
b
c
mean
Genes
ccyA
CmpA/NrtA family ABC transporter substrate−binding protein
WP_002746987.1_3397
log2(normalized counts)
15
10
5
0
7 am
[t1_N]
9 am
[t2_D]
12 am
[t3_D]
4 pm
[t4_D]
8 pm
[t5_D]
10 pm
[t6_N]
2 am
[t7_N]
7 am
[t8_N]
Sampling time

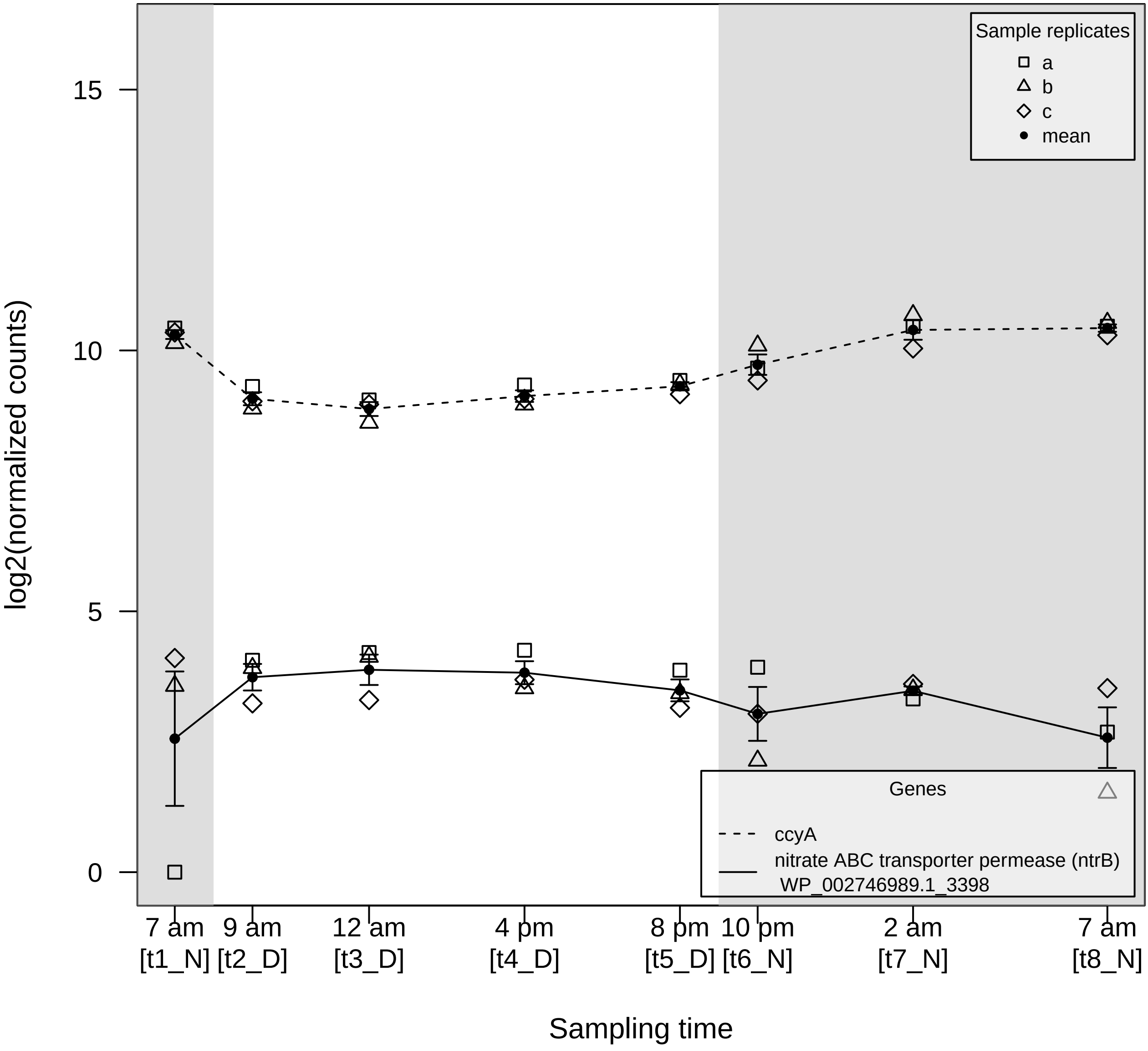
Sample replicates
a
b
c
mean
15
10
5
0
log2(normalized counts)
Genes
ccyA
nitrate ABC transporter permease (ntrB)
WP_002746989.1_3398
7 am [t1_N]
9 am [t2_D]
12 am [t3_D]
4 pm [t4_D]
8 pm [t5_D]
10 pm [t6_N]
2 am [t7_N]
7 am [t8_N]
Sampling time

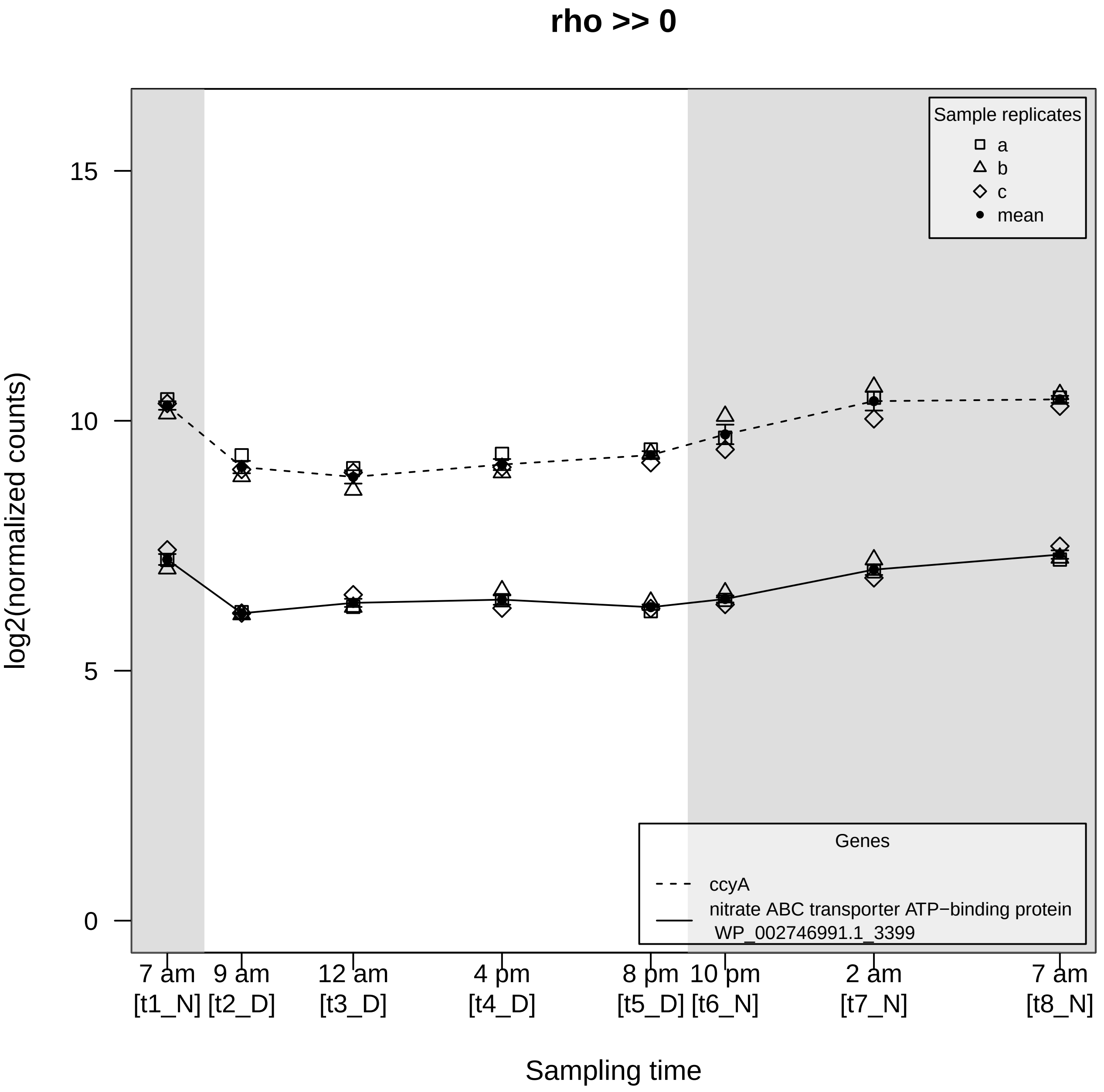
rho >> 0
Sample replicates
a
b
c
mean
Genes
ccyA
nitrate ABC transporter ATP−binding protein
WP_002746991.1_3399
log2(normalized counts)
15
10
5
0
7 am [t1_N]
9 am [t2_D]
12 am [t3_D]
4 pm [t4_D]
8 pm [t5_D]
10 pm [t6_N]
2 am [t7_N]
7 am [t8_N]
Sampling time

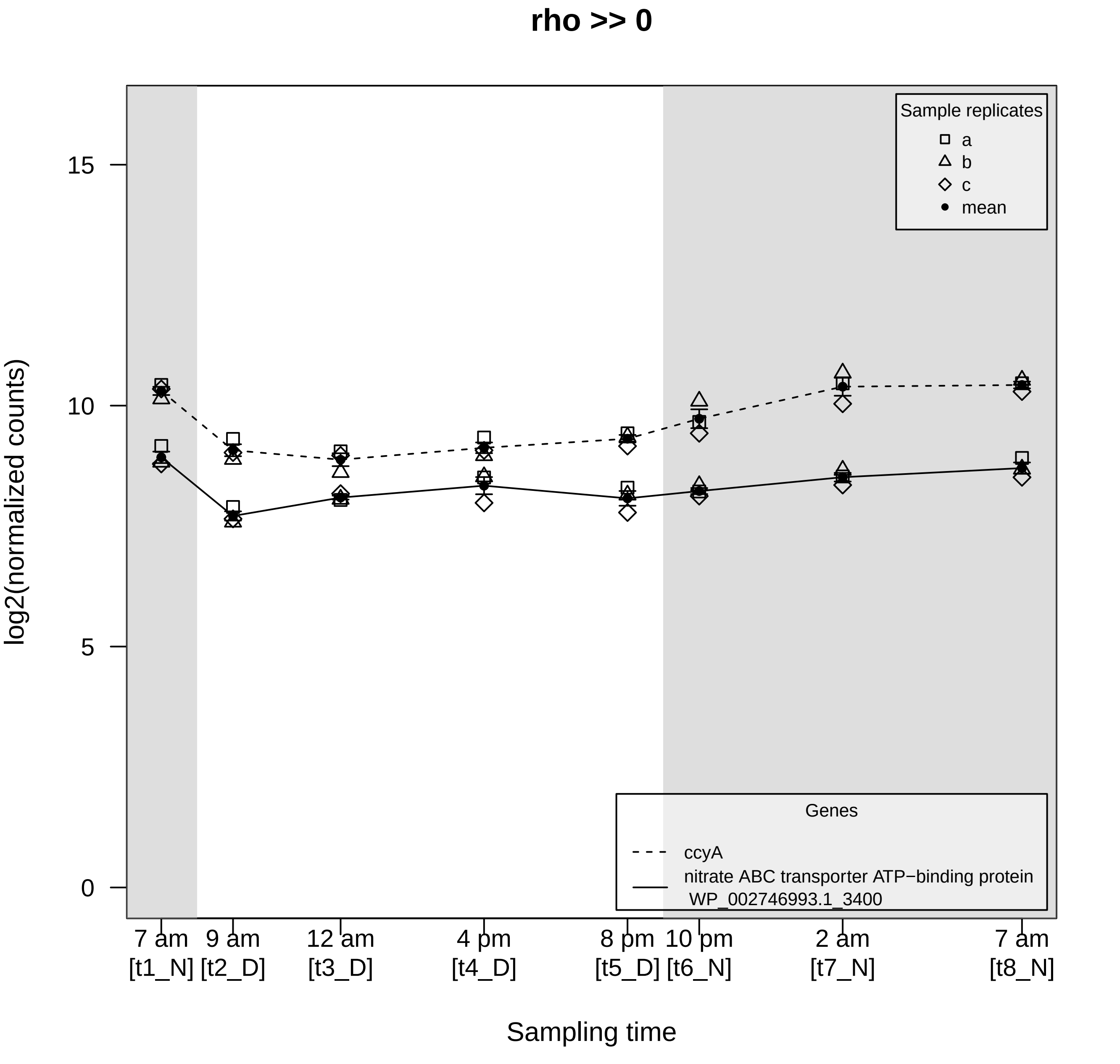

rho >> 0
log2(normalized counts)
15
10
5
0
Sample replicates
a
b
c
mean
Genes
ccyA
nitrate ABC transporter ATP−binding protein
WP_002746993.1_3400
7 am [t1_N]
9 am [t2_D]
12 am [t3_D]
4 pm [t4_D]
8 pm [t5_D]
10 pm [t6_N]
2 am [t7_N]
7 am [t8_N]
Sampling time

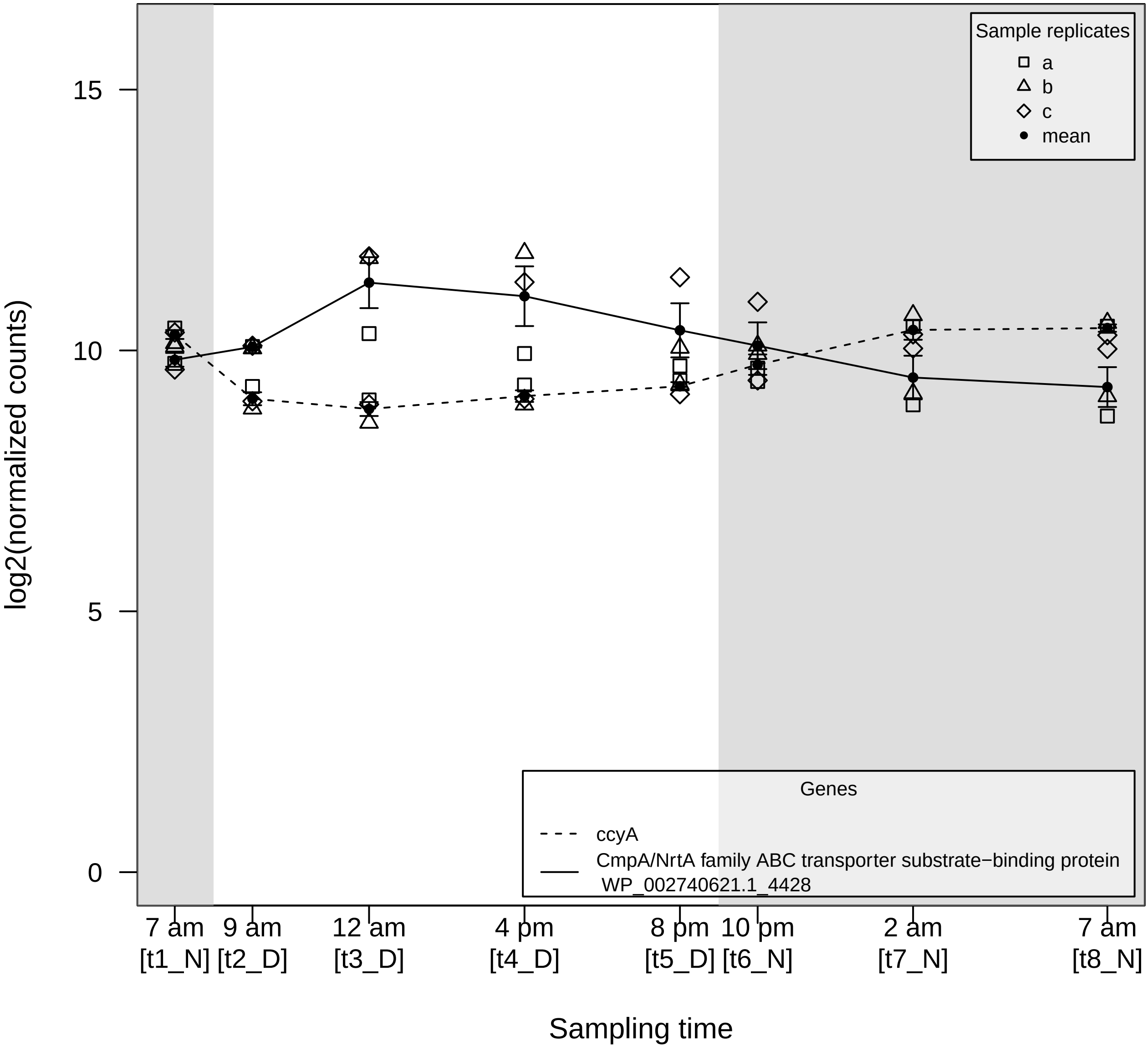
log2(normalized counts)
15
10
5
0
Sample replicates
a
b
c
mean
Genes
ccyA
CmpA/NrtA family ABC transporter substrate−binding protein
WP_002740621.1_4428
7 am [t1_N]
9 am [t2_D]
12 am [t3_D]
4 pm [t4_D]
8 pm [t5_D]
10 pm [t6_N]
2 am [t7_N]
7 am [t8_N]
Sampling time

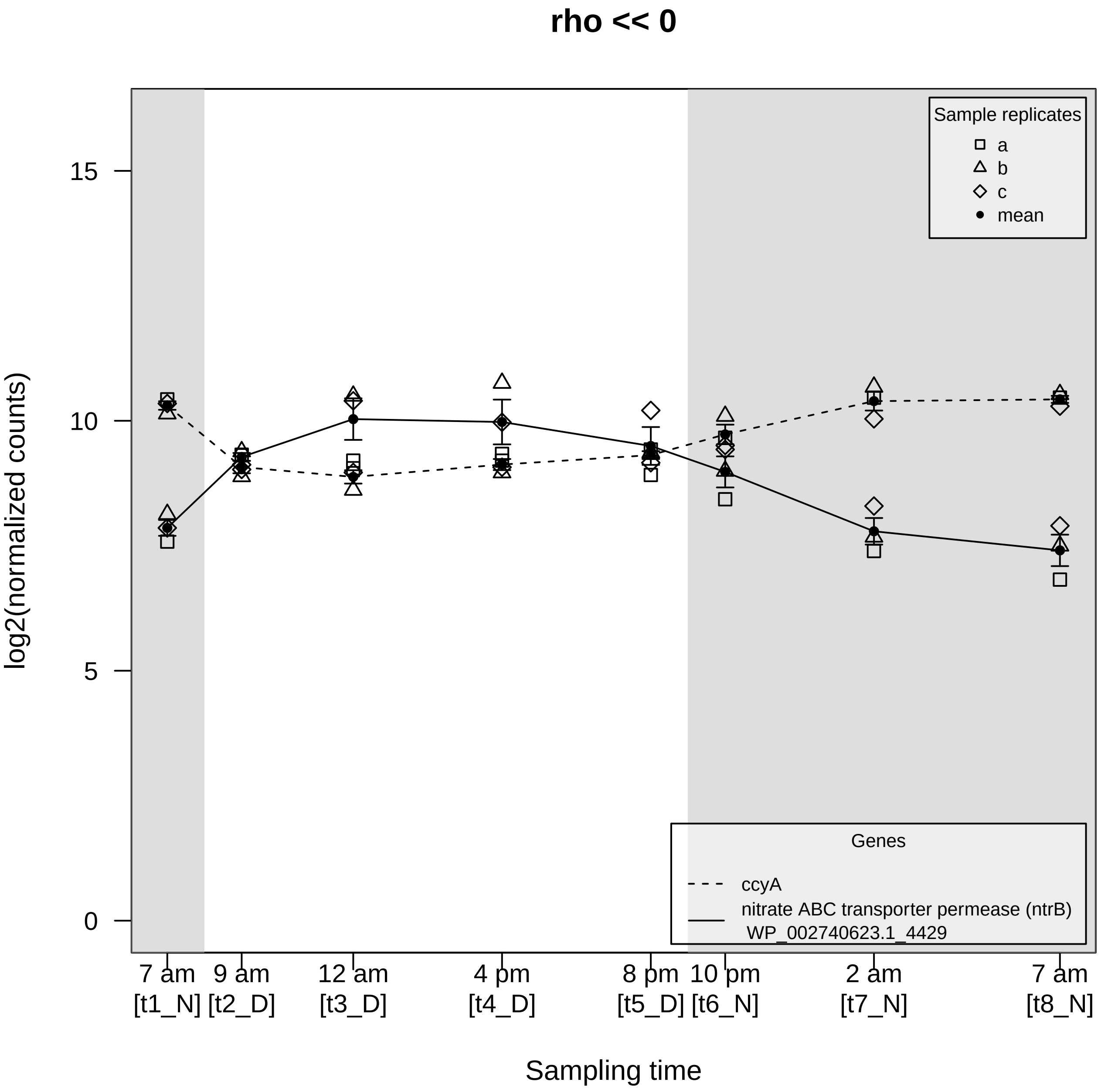
rho << 0
log2(normalized counts)
15
10
5
0
Sample replicates
a
b
c
mean
Genes
ccyA
nitrate ABC transporter permease (ntrB)
WP_002740623.1_4429
7 am
[t1_N]
9 am
[t2_D]
12 am
[t3_D]
4 pm
[t4_D]
8 pm
[t5_D]
10 pm
[t6_N]
2 am
[t7_N]
7 am
[t8_N]
Sampling time

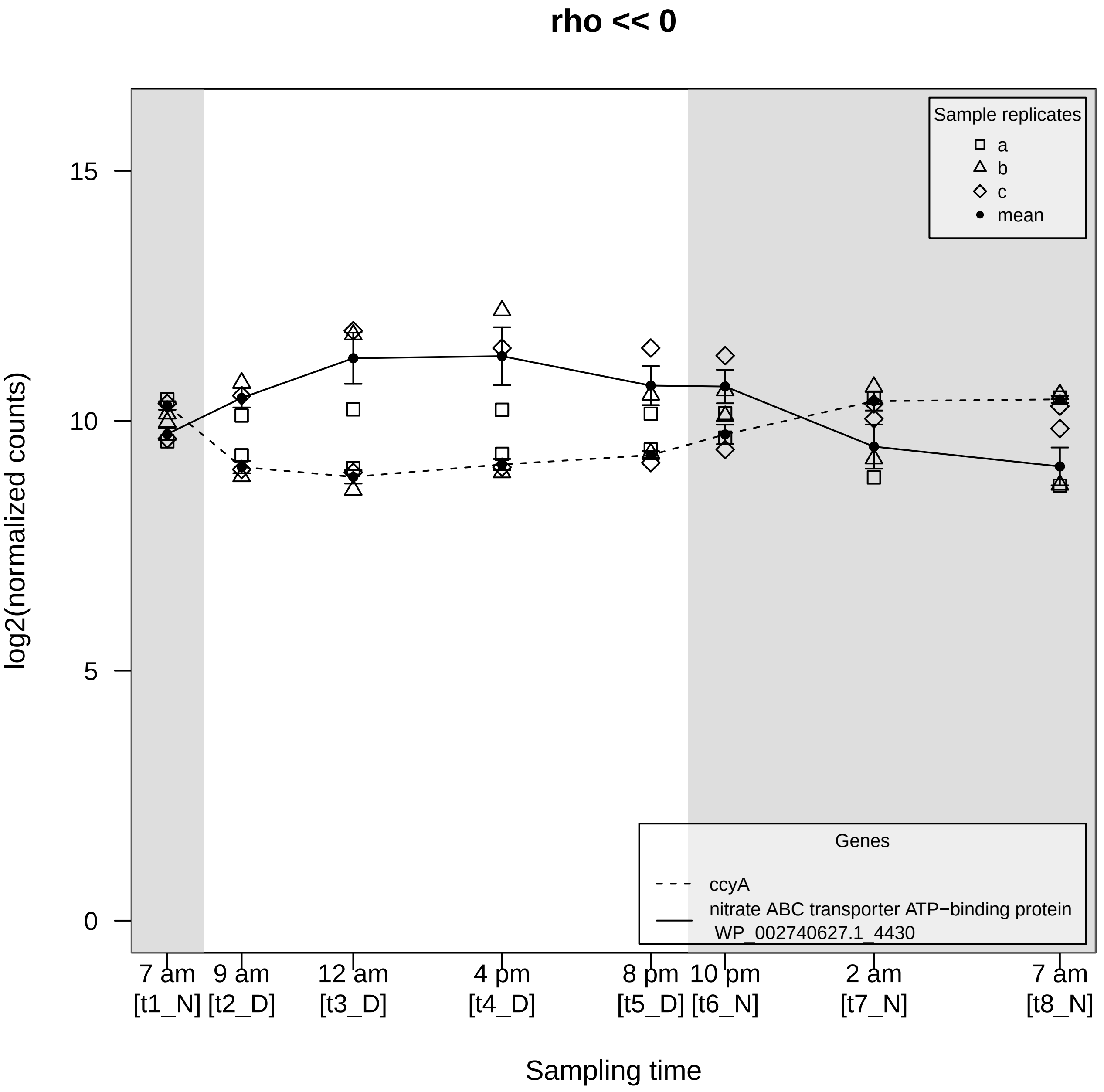

rho << 0
Sample replicates
a
b
c
mean
Genes
ccyA
nitrate ABC transporter ATP−binding protein
WP_002740627.1_4430
log2(normalized counts)
0
5
10
15
7 am
[t1_N]
9 am
[t2_D]
12 am
[t3_D]
4 pm
[t4_D]
8 pm
[t5_D]
10 pm
[t6_N]
2 am
[t7_N]
7 am
[t8_N]
Sampling time

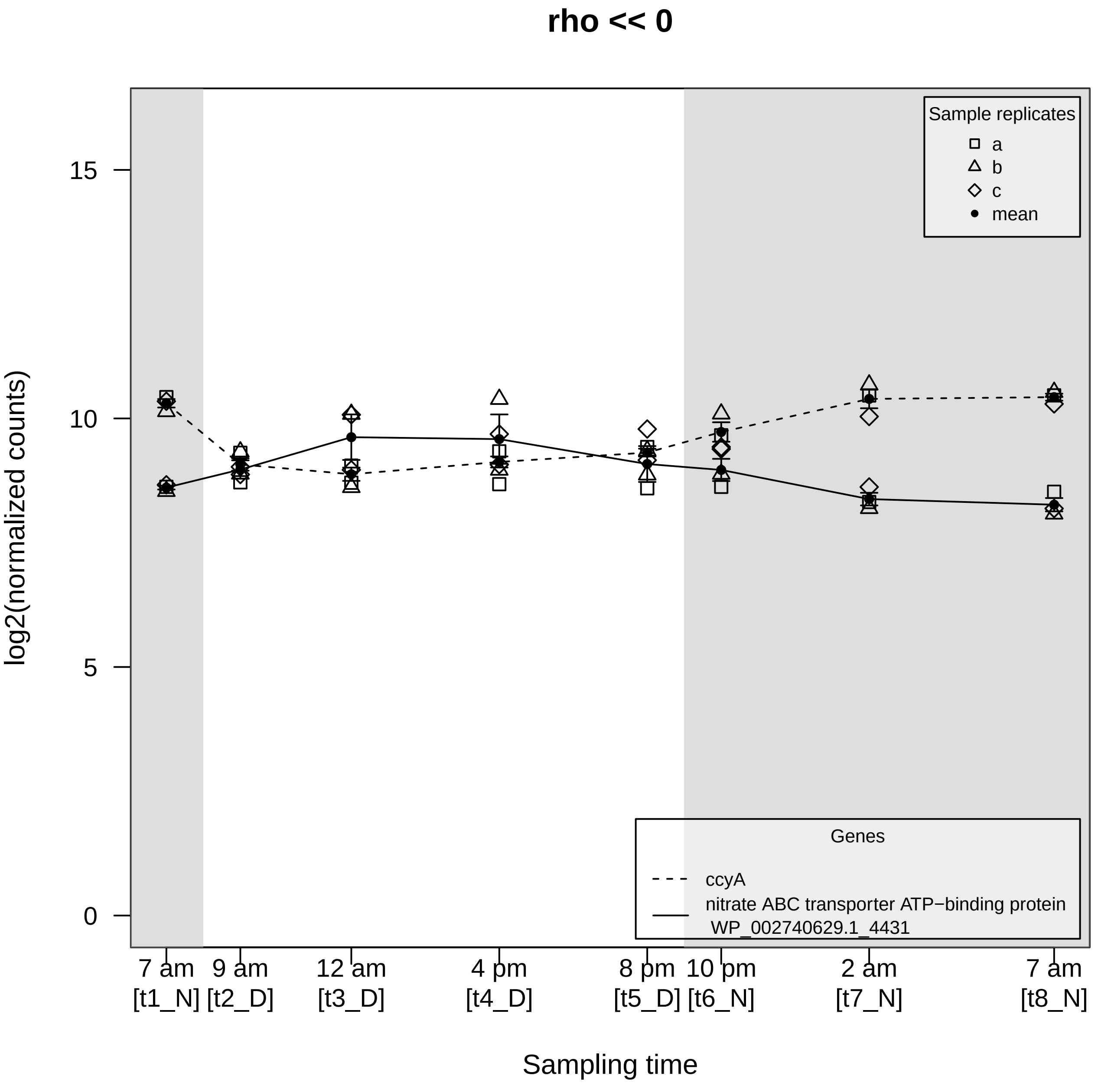

rho << 0
Sample replicates
a
b
c
mean
15
10
5
0
log2(normalized counts)
Genes
ccyA
nitrate ABC transporter ATP−binding protein
WP_002740629.1_4431
7 am [t1_N]
9 am [t2_D]
12 am [t3_D]
4 pm [t4_D]
8 pm [t5_D]
10 pm [t6_N]
2 am [t7_N]
7 am [t8_N]
Sampling time

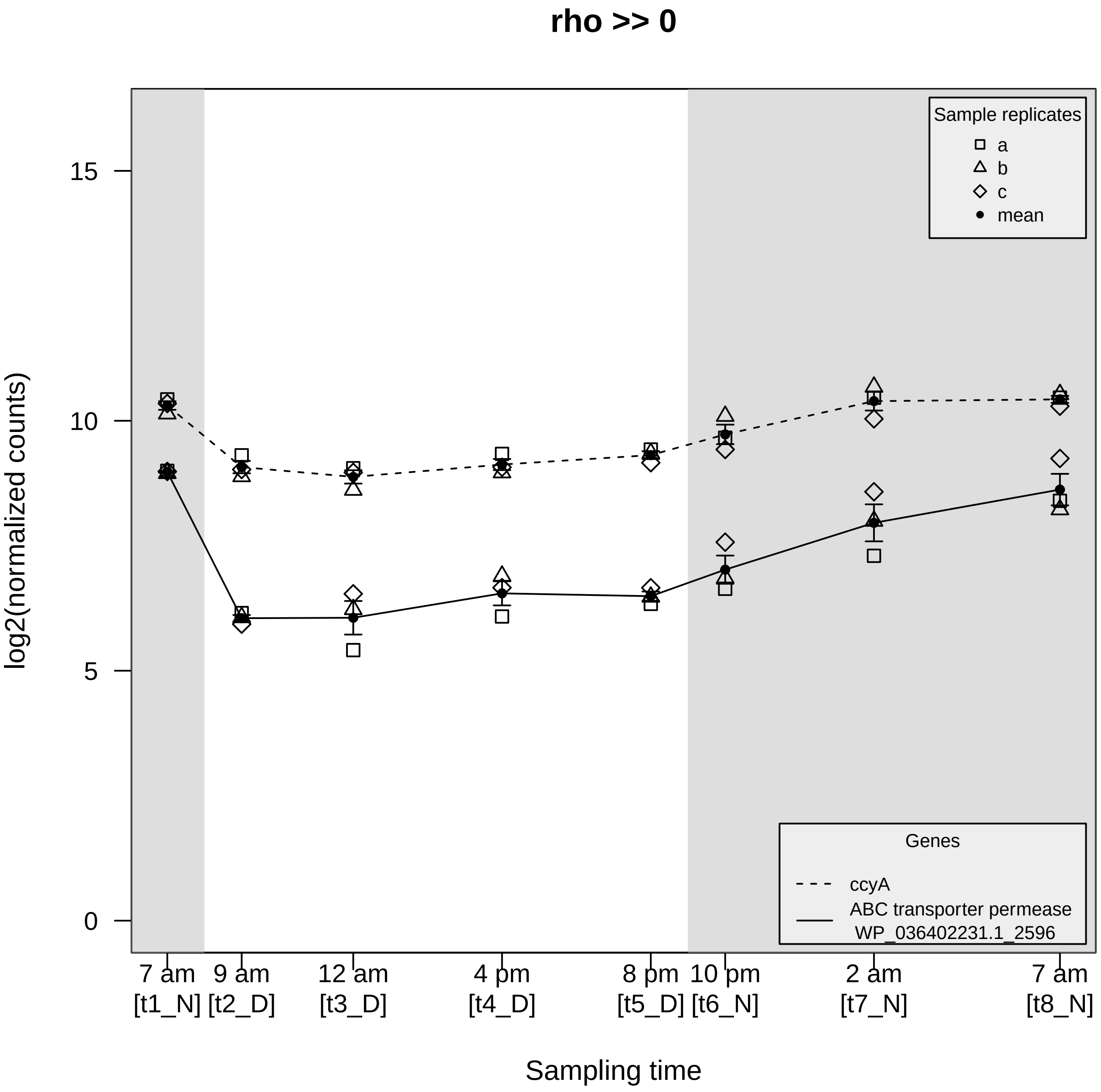

rho >> 0
log2(normalized counts)
15
10
5
0
Sample replicates
a
b
c
mean
Genes
ccyA
ABC transporter permease
WP_036402231.1_2596
7 am
[t1_N]
9 am
[t2_D]
12 am
[t3_D]
4 pm
[t4_D]
8 pm
[t5_D]
10 pm
[t6_N]
2 am
[t7_N]
7 am
[t8_N]
Sampling time

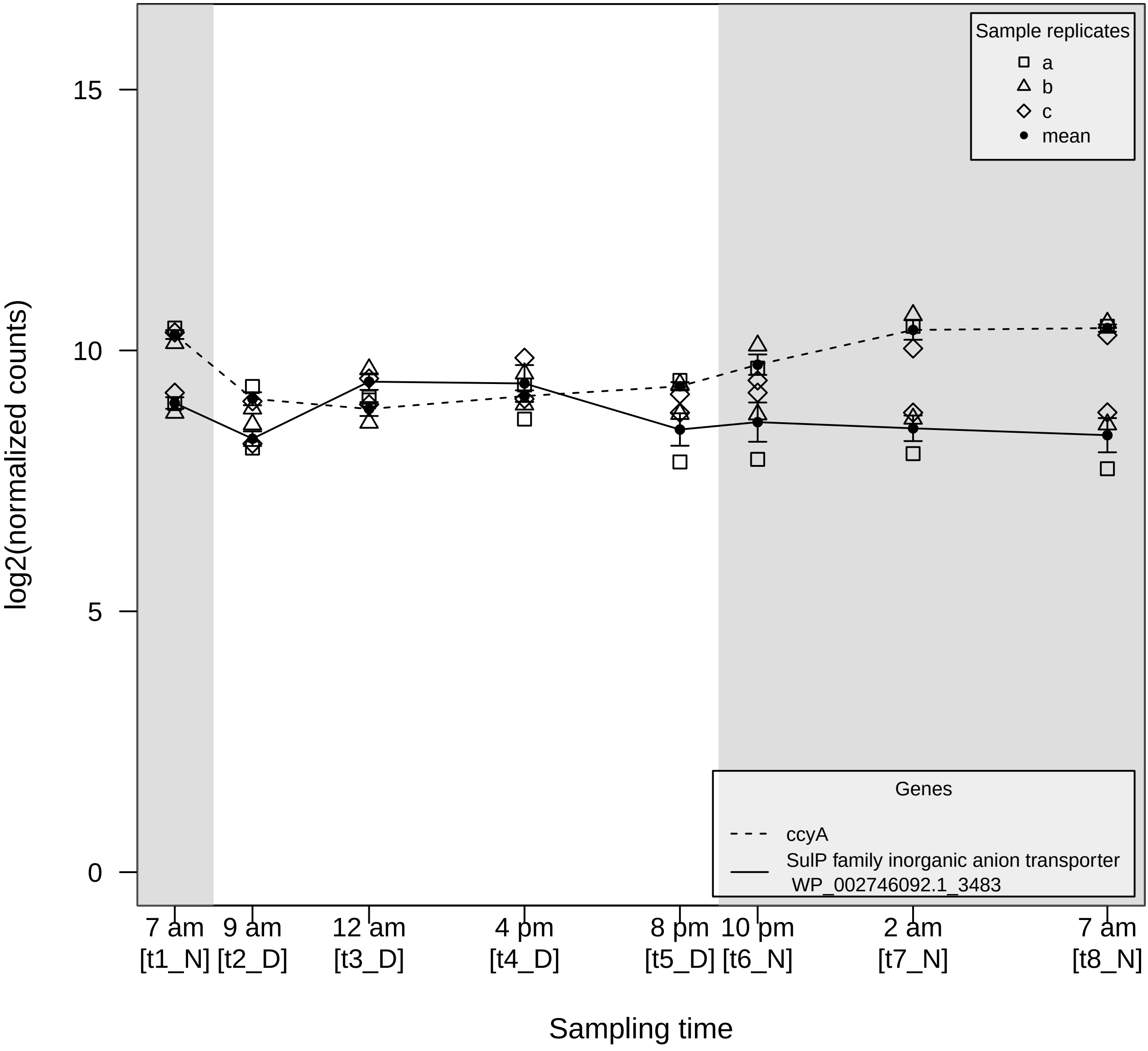
Sample replicates
a
b
c
mean
Genes
ccyA
SulP family inorganic anion transporter
WP_002746092.1_3483
log2(normalized counts)
15
10
5
0
7 am [t1_N]
9 am [t2_D]
12 am [t3_D]
4 pm [t4_D]
8 pm [t5_D]
10 pm [t6_N]
2 am [t7_N]
7 am [t8_N]
Sampling time

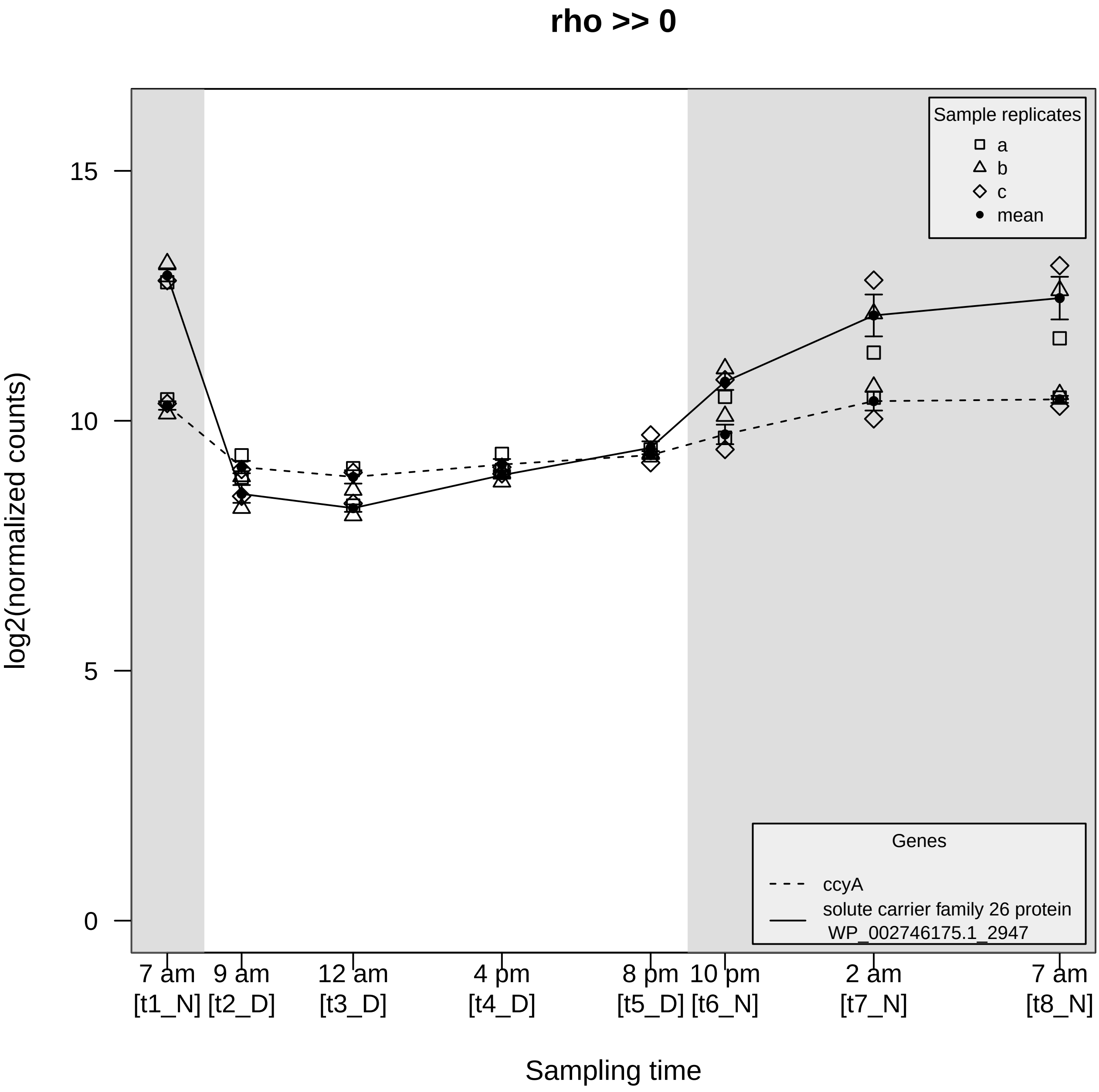

rho >> 0
log2(normalized counts)
15
10
5
0
Sample replicates
a
b
c
mean
Genes
ccyA
solute carrier family 26 protein
WP_002746175.1_2947
7 am [t1_N]
9 am [t2_D]
12 am [t3_D]
4 pm [t4_D]
8 pm [t5_D]
10 pm [t6_N]
2 am [t7_N]
7 am [t8_N]
Sampling time

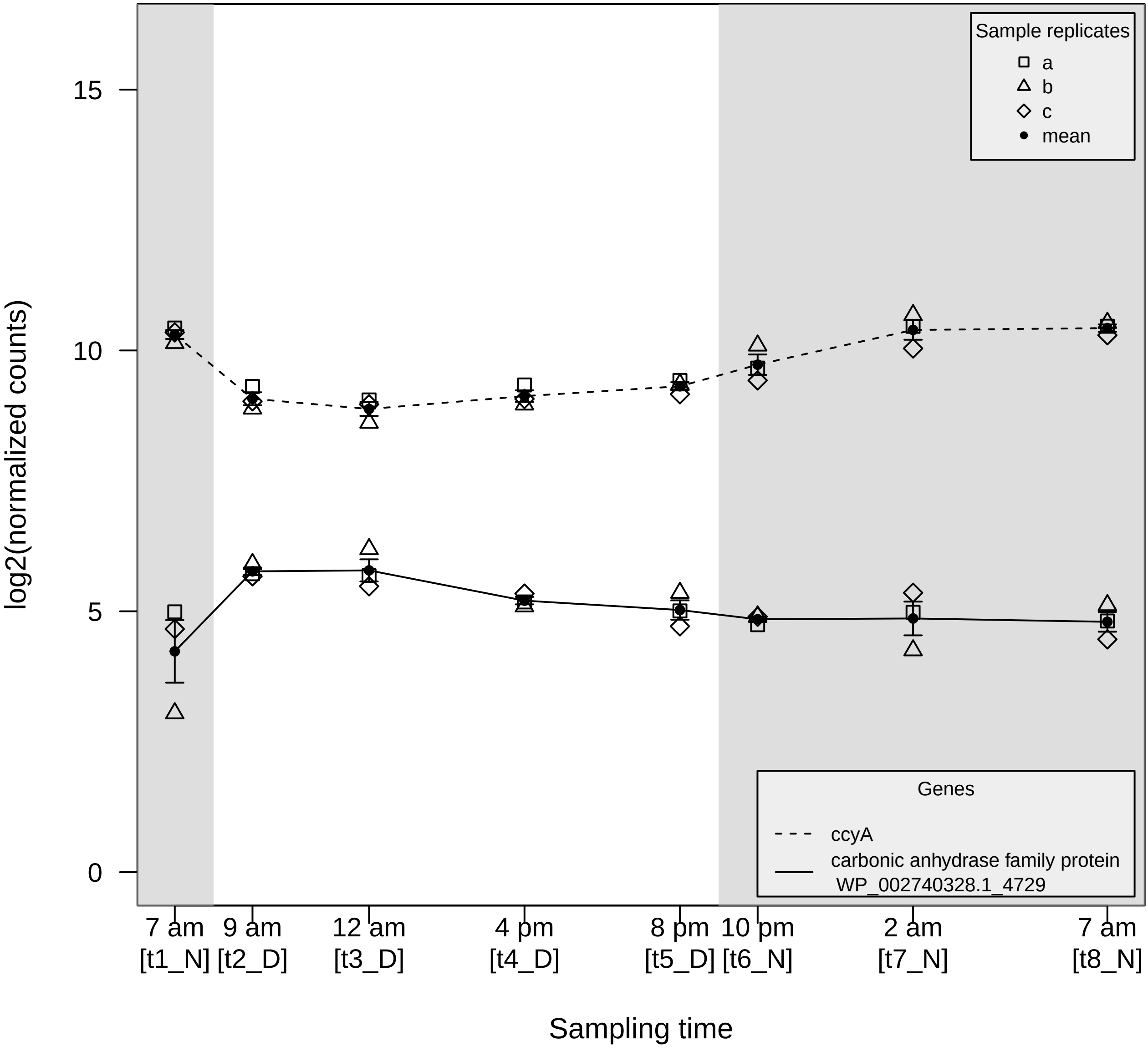

Sample replicates
a
b
c
mean
Genes
ccyA
carbonic anhydrase family protein
WP_002740328.1_4729
log2(normalized counts)
15
10
5
0
7 am [t1_N]
9 am [t2_D]
12 am [t3_D]
4 pm [t4_D]
8 pm [t5_D]
10 pm [t6_N]
2 am [t7_N]
7 am [t8_N]
Sampling time
