## Supplementary_Figures for "Diel changes in the expression of a marker gene and candidate genes for intracellular amorphous CaCO_3_ biomineralization in *Microcystis*": Figure_S5_SI_ca.pdf

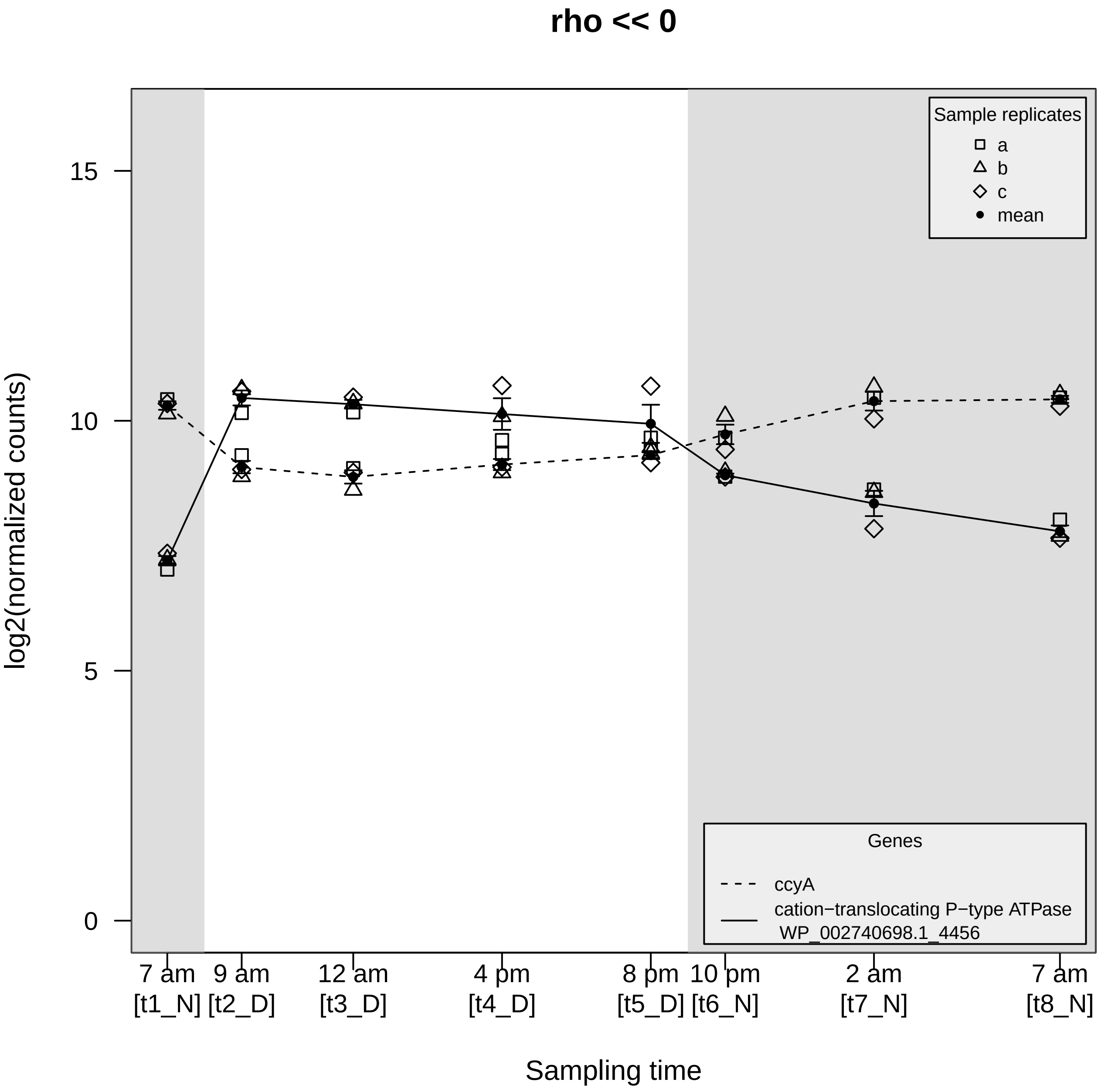

rho << 0
Sample replicates
a
b
c
mean
Genes
ccyA
cation−translocating P−type ATPase
WP_002740698.1_4456
log2(normalized counts)
15
10
5
0
7 am
[t1_N]
9 am
[t2_D]
12 am
[t3_D]
4 pm
[t4_D]
8 pm
[t5_D]
10 pm
[t6_N]
2 am
[t7_N]
7 am
[t8_N]
Sampling time

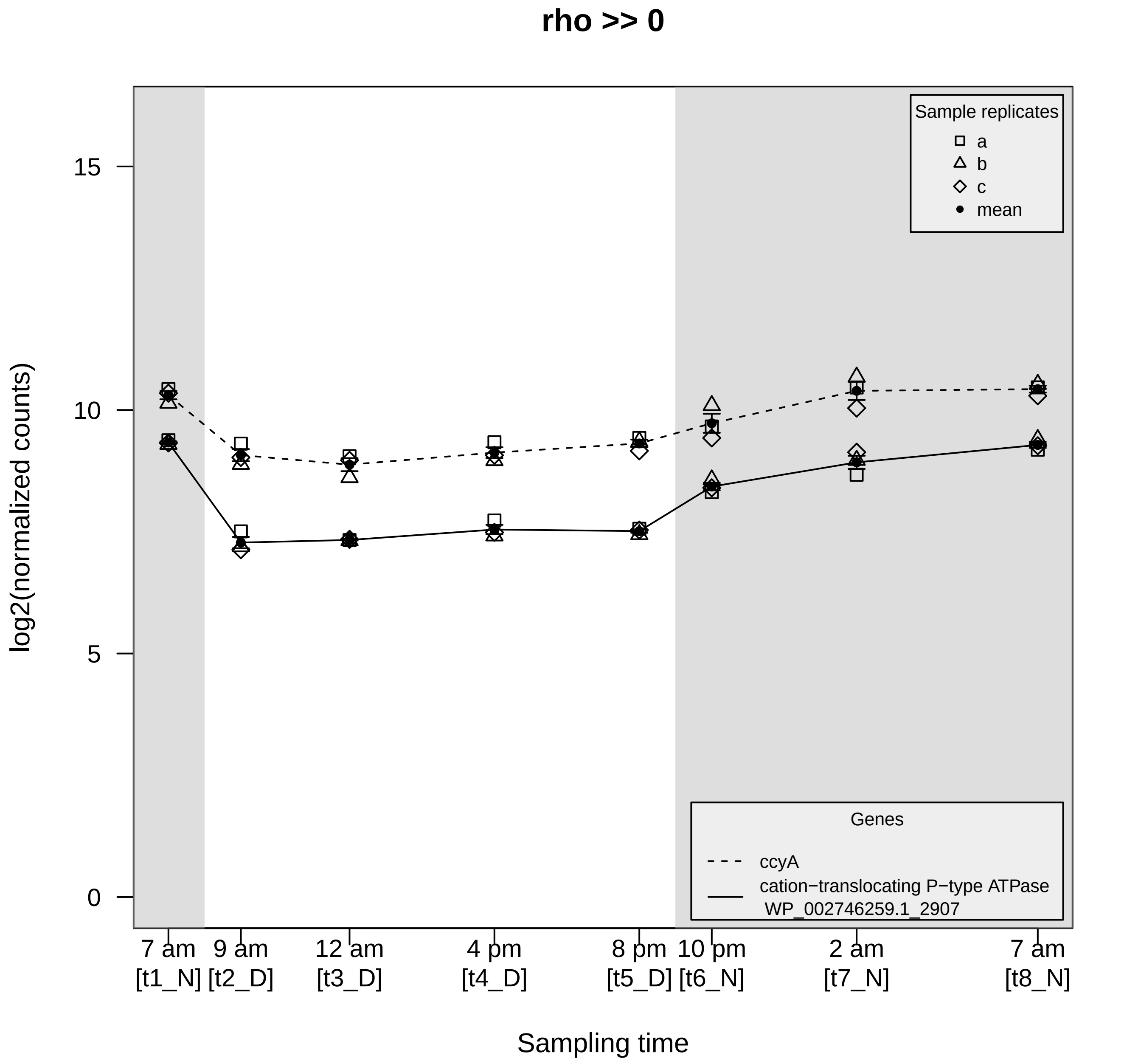

rho >> 0
Sample replicates
a
b
c
mean
Genes
ccyA
cation−translocating P−type ATPase WP_002746259.1_2907
log2(normalized counts)
0
5
10
15
7 am [t1_N]
9 am [t2_D]
12 am [t3_D]
4 pm [t4_D]
8 pm [t5_D]
10 pm [t6_N]
2 am [t7_N]
7 am [t8_N]
Sampling time

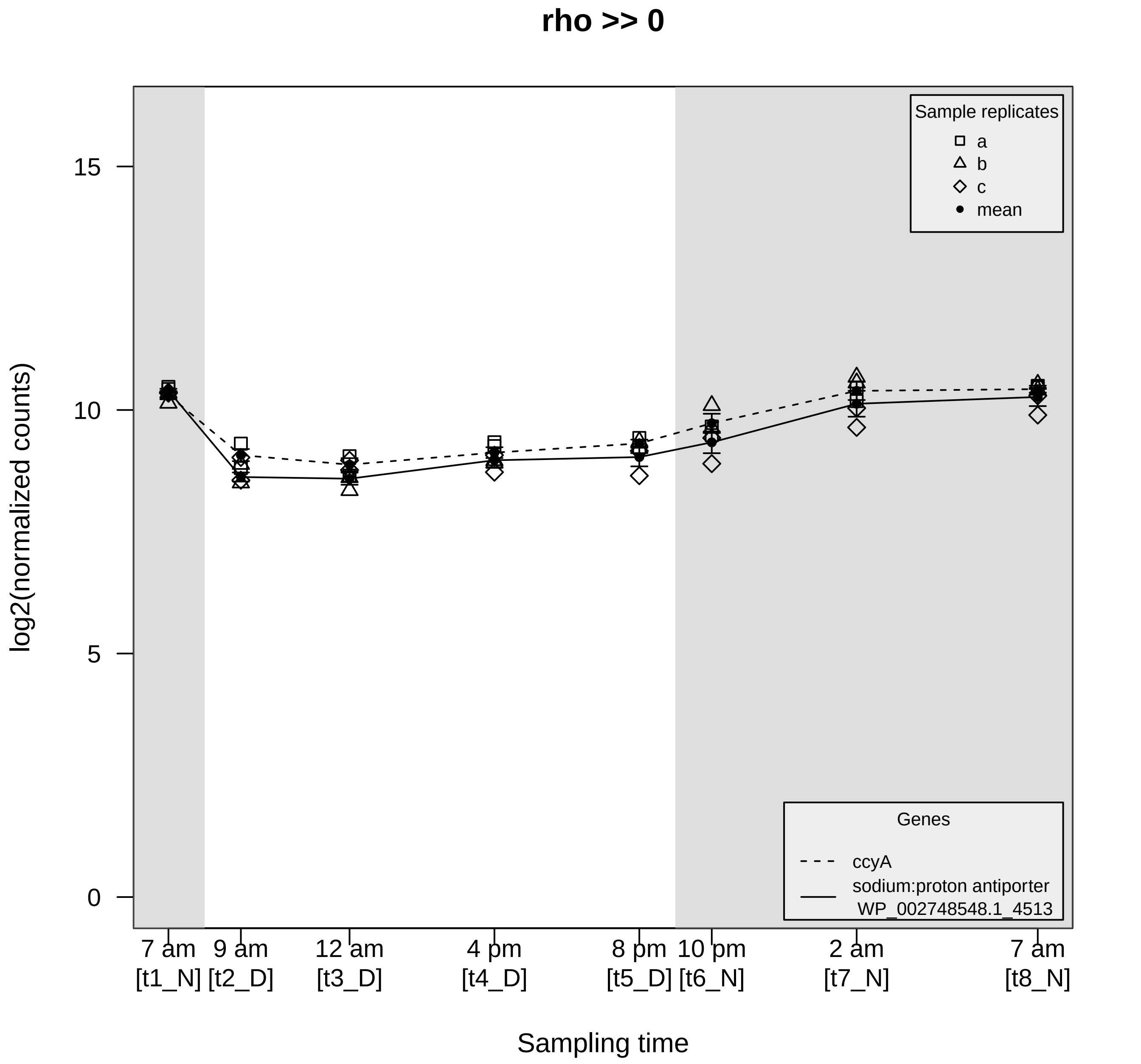

rho >> 0
log2(normalized counts)
15
10
5
0
Sample replicates
a
b
c
mean
Genes
ccyA
sodium:proton antiporter
WP_002748548.1_4513
7 am [t1_N]
9 am [t2_D]
12 am [t3_D]
4 pm [t4_D]
8 pm [t5_D]
10 pm [t6_N]
2 am [t7_N]
7 am [t8_N]
Sampling time

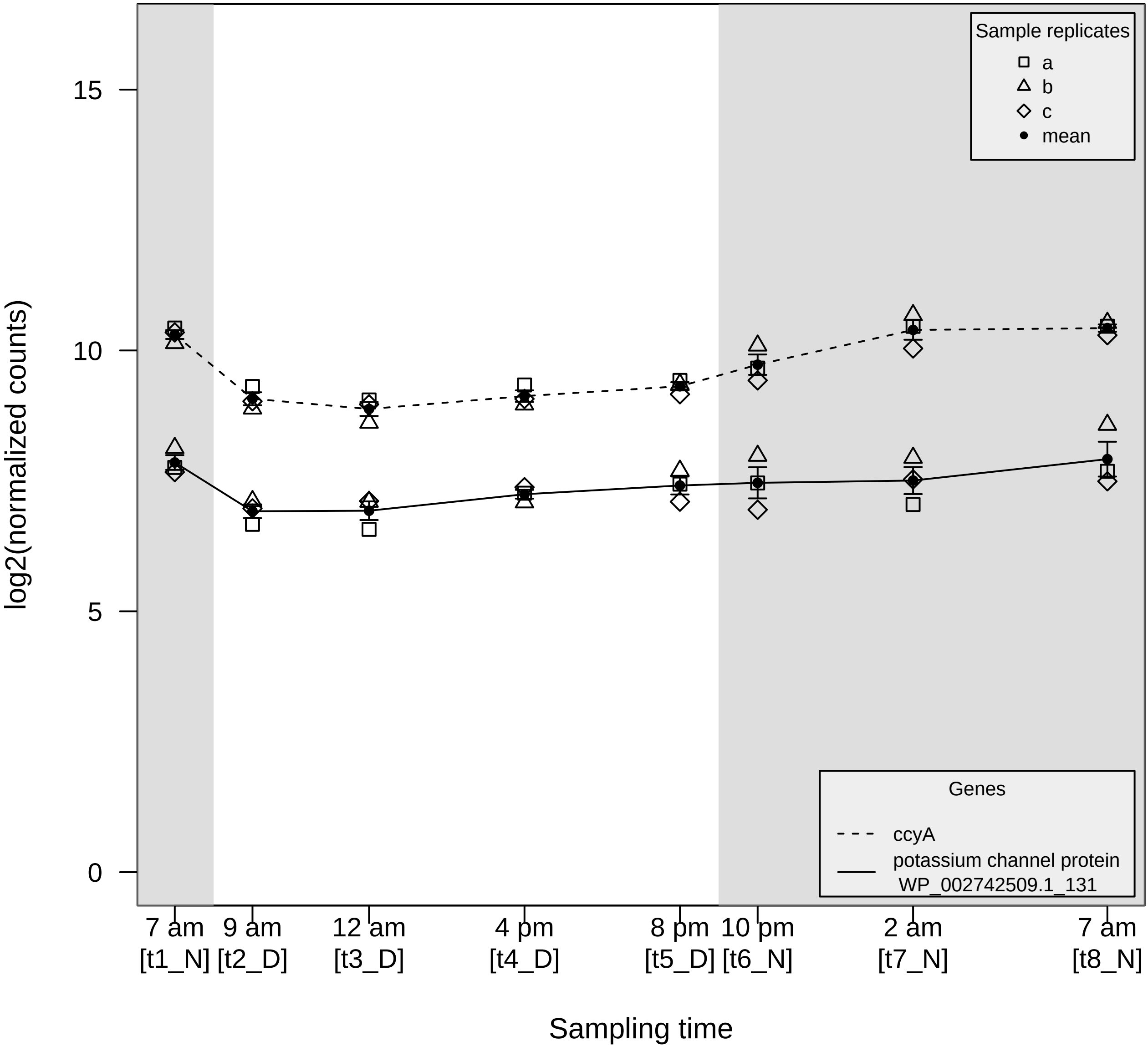

Sample replicates
a
b
c
mean
Genes
ccyA
potassium channel protein
WP_002742509.1_131
log2(normalized counts)
0
5
10
15
7 am
[t1_N]
9 am
[t2_D]
12 am
[t3_D]
4 pm
[t4_D]
8 pm
[t5_D]
10 pm
[t6_N]
2 am
[t7_N]
7 am
[t8_N]
Sampling time

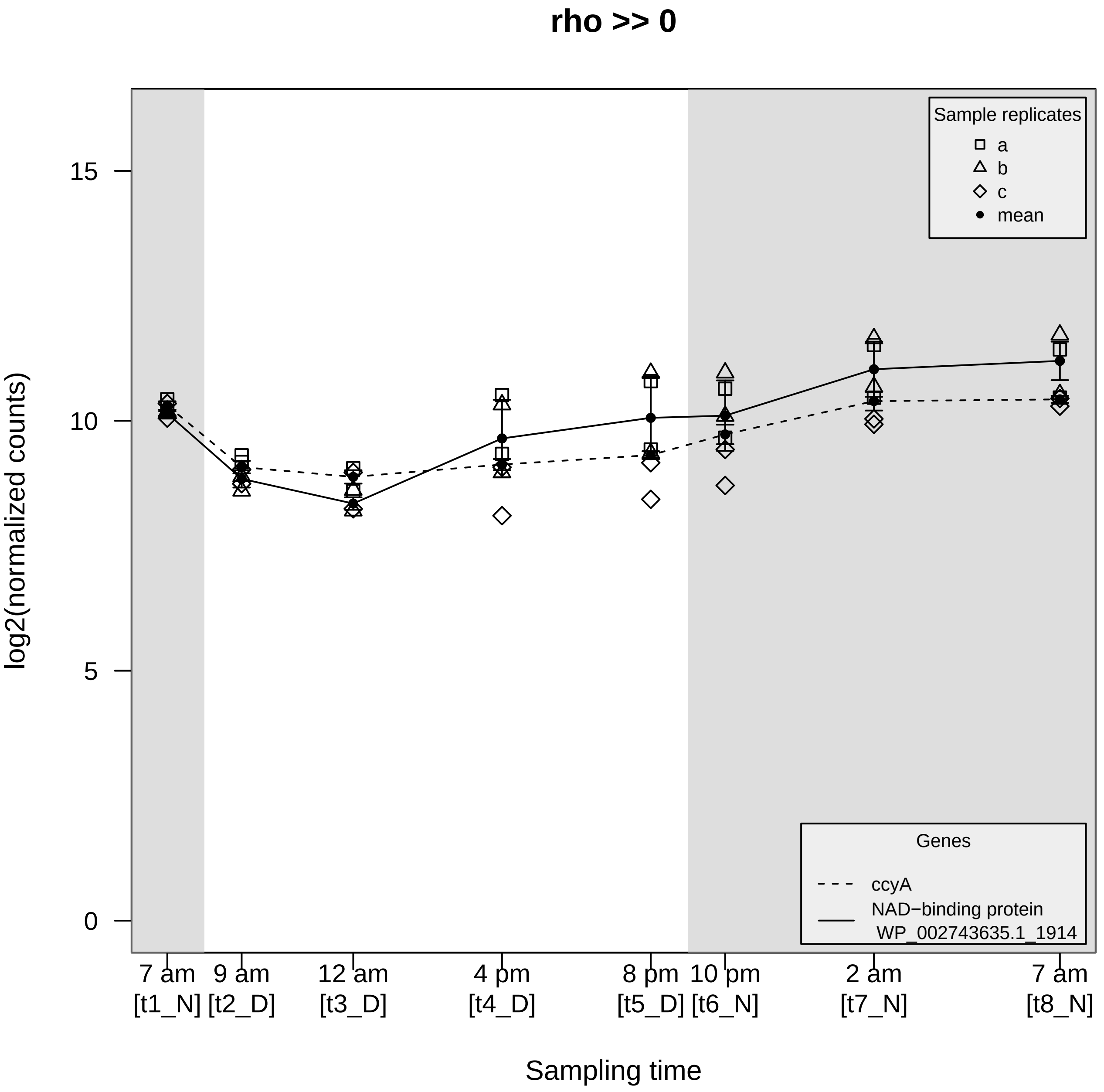
rho >> 0
log2(normalized counts)
15
10
5
0
Sample replicates
a
b
c
mean
Genes
ccyA
NAD−binding protein
WP_002743635.1_1914
7 am [t1_N]
9 am [t2_D]
12 am [t3_D]
4 pm [t4_D]
8 pm [t5_D]
10 pm [t6_N]
2 am [t7_N]
7 am [t8_N]
Sampling time

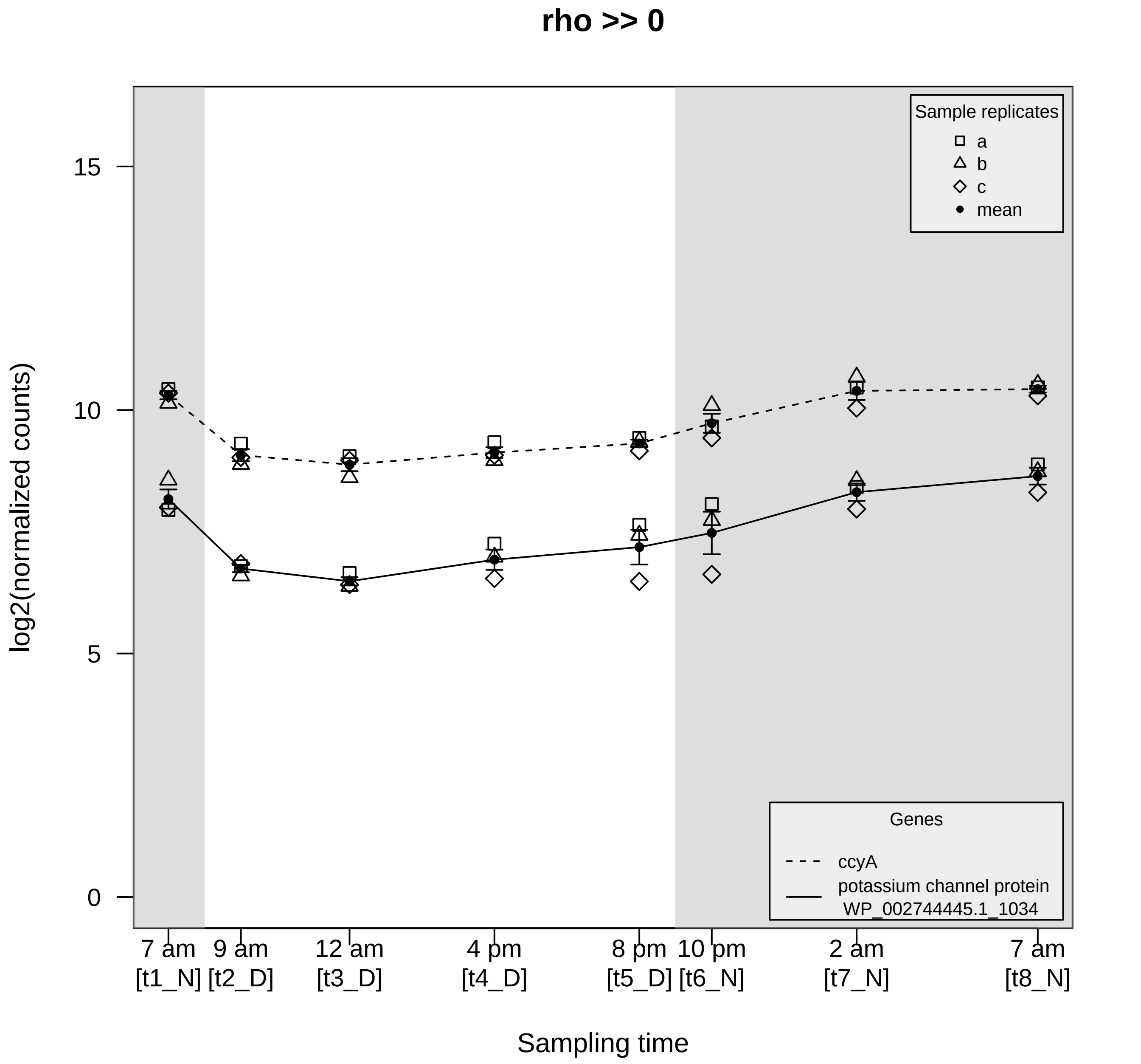

rho >> 0
Sample replicates
a
b
c
mean
Genes
ccyA
potassium channel protein
WP_002744445.1_1034
log2(normalized counts)
15
10
5
0
7 am
[t1_N]
9 am
[t2_D]
12 am
[t3_D]
4 pm
[t4_D]
8 pm
[t5_D]
10 pm
[t6_N]
2 am
[t7_N]
7 am
[t8_N]
Sampling time

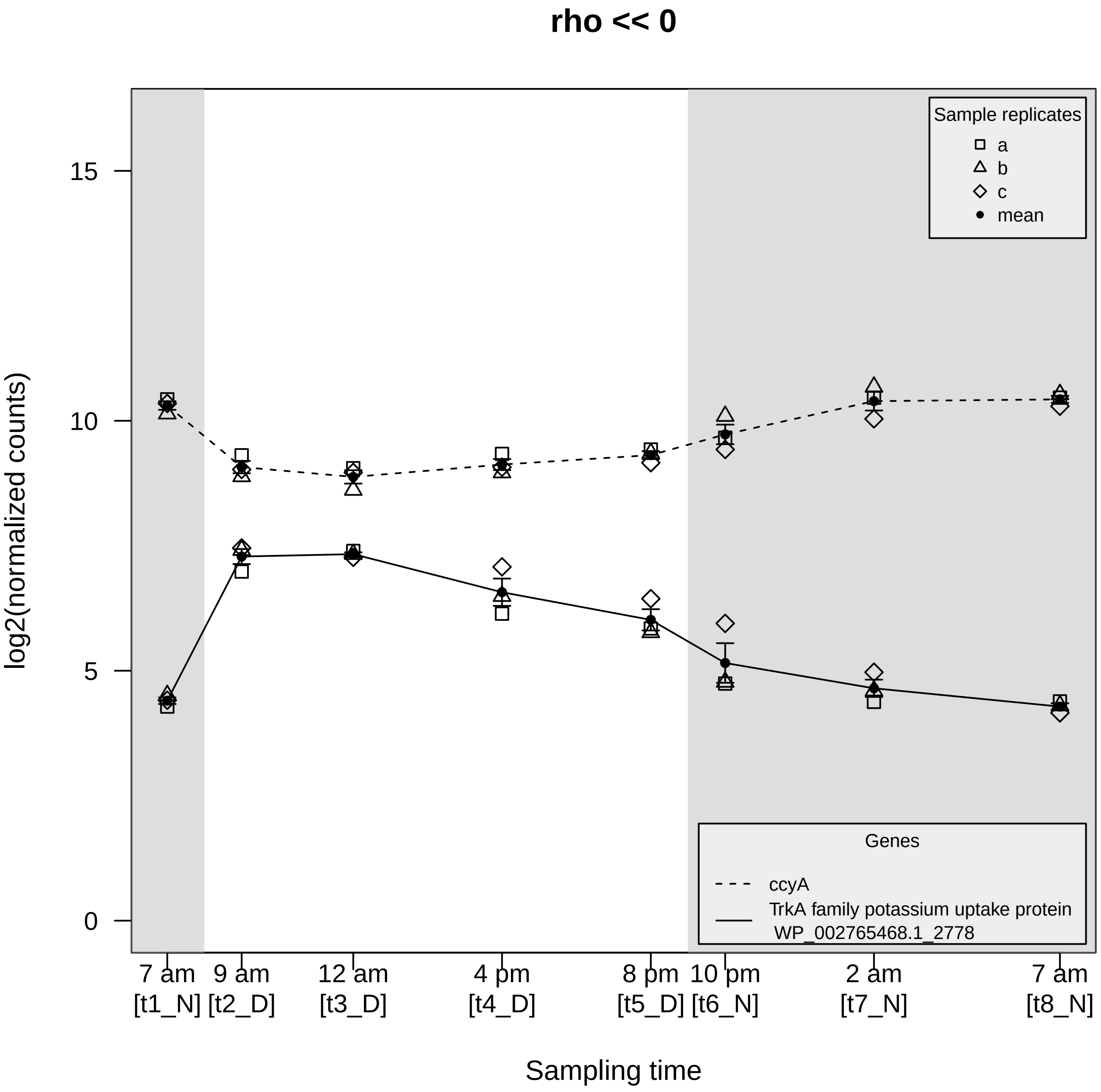

rho << 0
Sample replicates
a
b
c
mean
Genes
ccyA
TrkA family potassium uptake protein
WP_002765468.1_2778
log2(normalized counts)
15
10
5
0
7 am
[t1_N]
9 am
[t2_D]
12 am
[t3_D]
4 pm
[t4_D]
8 pm
[t5_D]
10 pm
[t6_N]
2 am
[t7_N]
7 am
[t8_N]
Sampling time

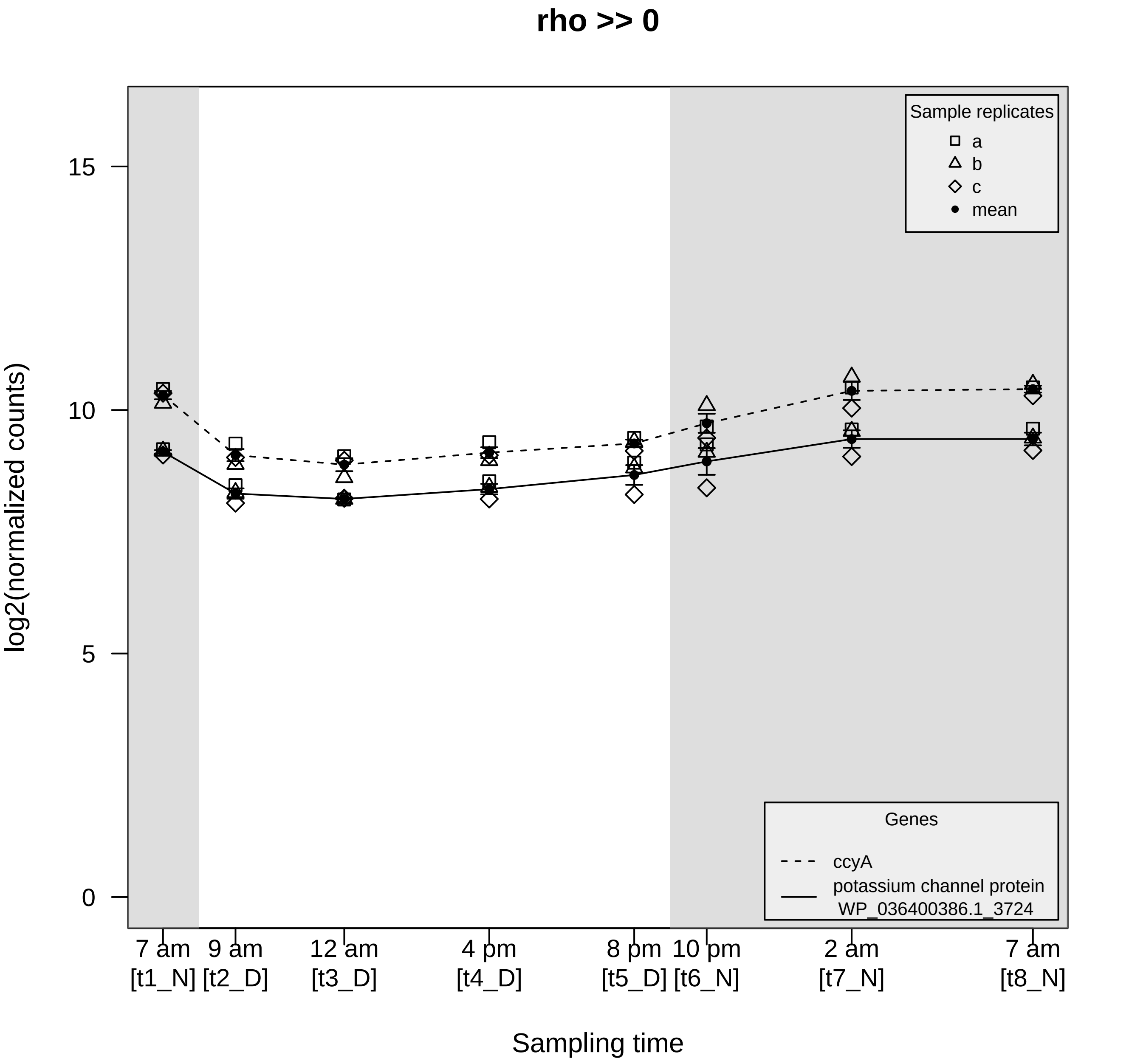
rho >> 0
Sample replicates
a
b
c
mean
Genes
ccyA
potassium channel protein WP_036400386.1_3724
log2(normalized counts)
15
10
5
0
7 am [t1_N]
9 am [t2_D]
12 am [t3_D]
4 pm [t4_D]
8 pm [t5_D]
10 pm [t6_N]
2 am [t7_N]
7 am [t8_N]
Sampling time

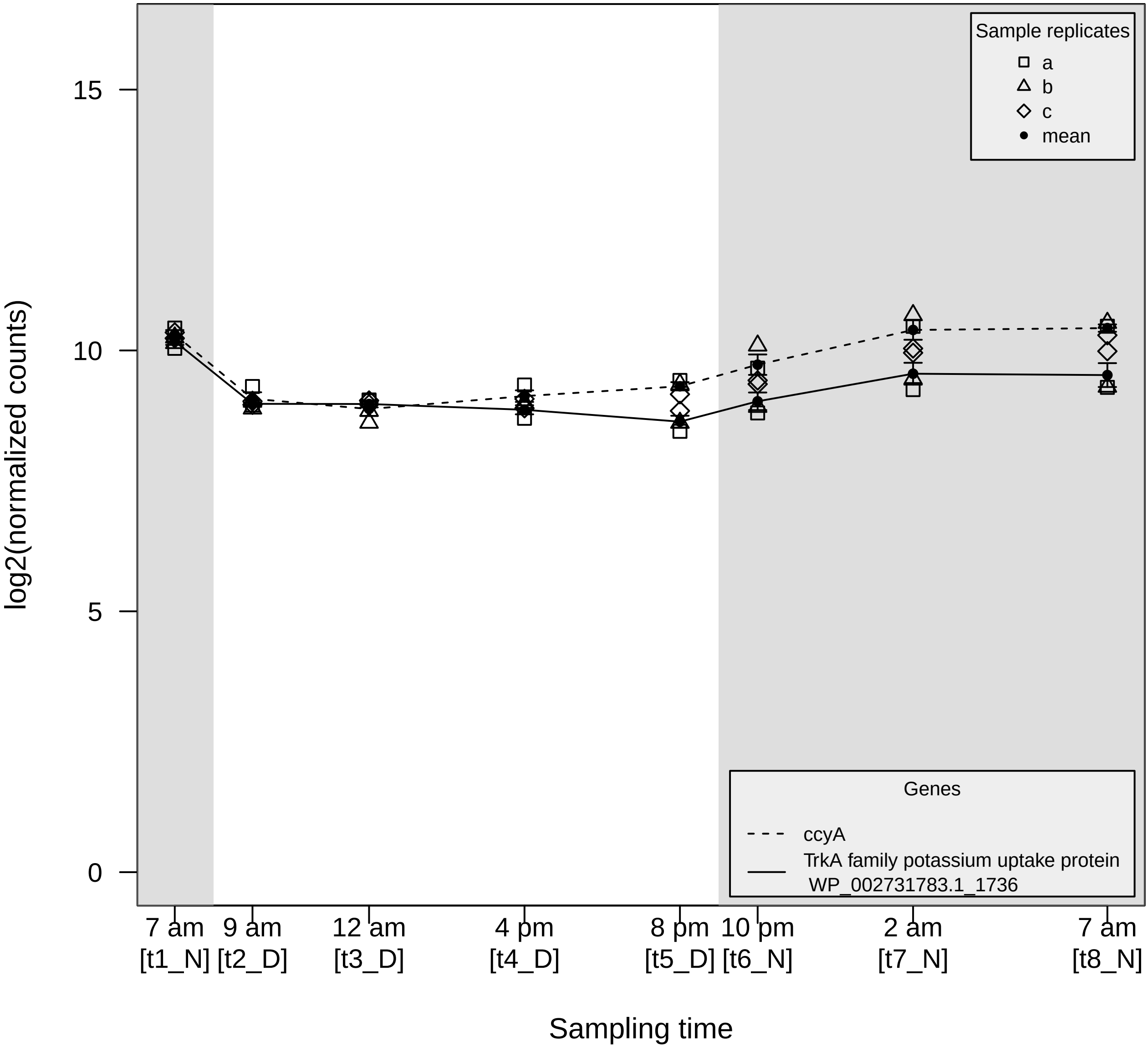

Sample replicates
a
b
c
mean
Genes
ccyA
TrkA family potassium uptake protein
WP_002731783.1_1736
log2(normalized counts)
15
10
5
0
7 am [t1_N]
9 am [t2_D]
12 am [t3_D]
4 pm [t4_D]
8 pm [t5_D]
10 pm [t6_N]
2 am [t7_N]
7 am [t8_N]
Sampling time

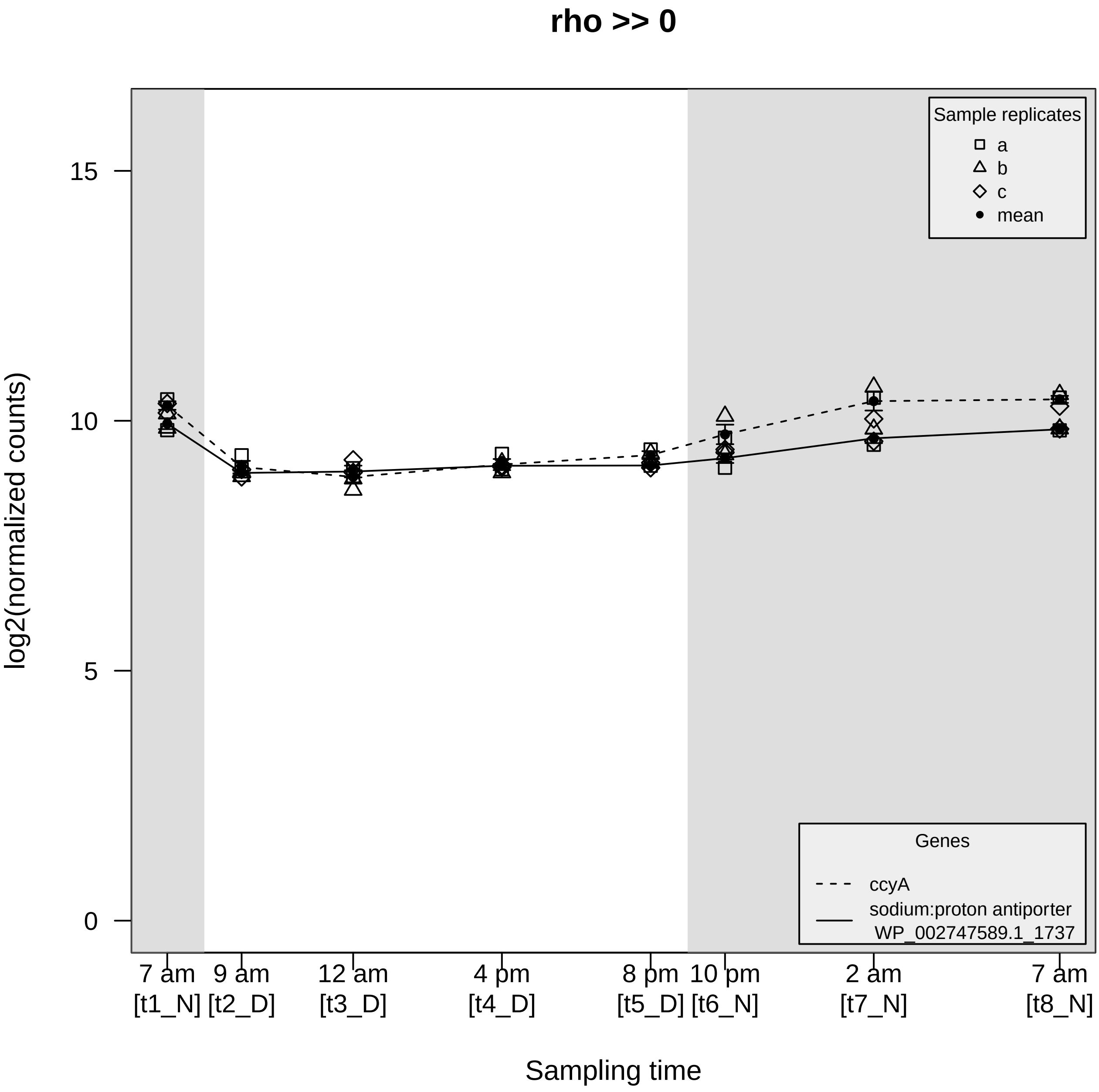

rho >> 0
Sample replicates
a
b
c
mean
Genes
ccyA
sodium:proton antiporter
WP_002747589.1_1737
log2(normalized counts)
15
10
5
0
7 am [t1_N]
9 am [t2_D]
12 am [t3_D]
4 pm [t4_D]
8 pm [t5_D]
10 pm [t6_N]
2 am [t7_N]
7 am [t8_N]
Sampling time

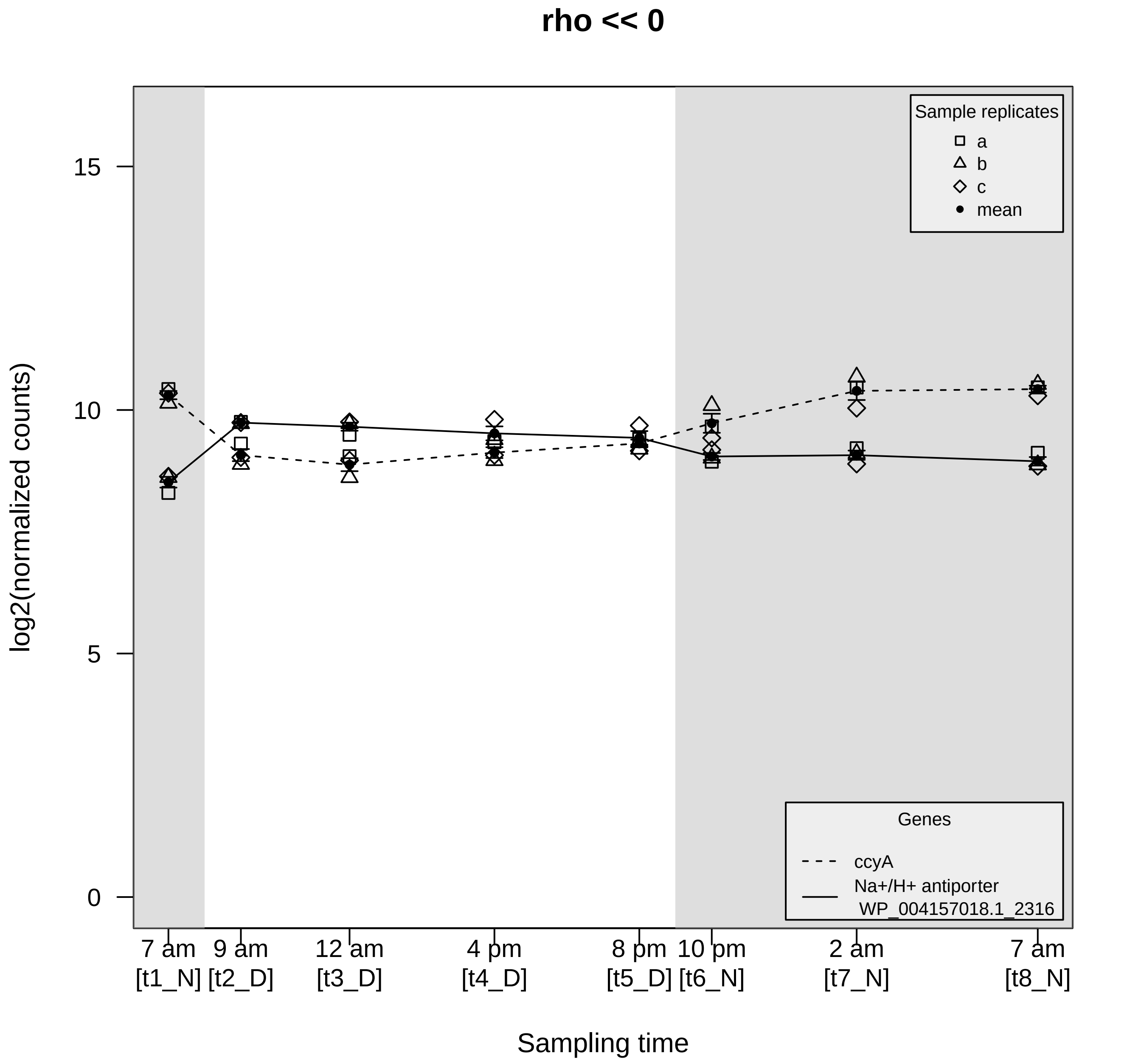
rho << 0
Sample replicates
a
b
c
mean
Genes
ccyA
Na+/H+ antiporter
WP_004157018.1_2316
log2(normalized counts)
15
10
5
0
7 am [t1_N]
9 am [t2_D]
12 am [t3_D]
4 pm [t4_D]
8 pm [t5_D]
10 pm [t6_N]
2 am [t7_N]
7 am [t8_N]
Sampling time

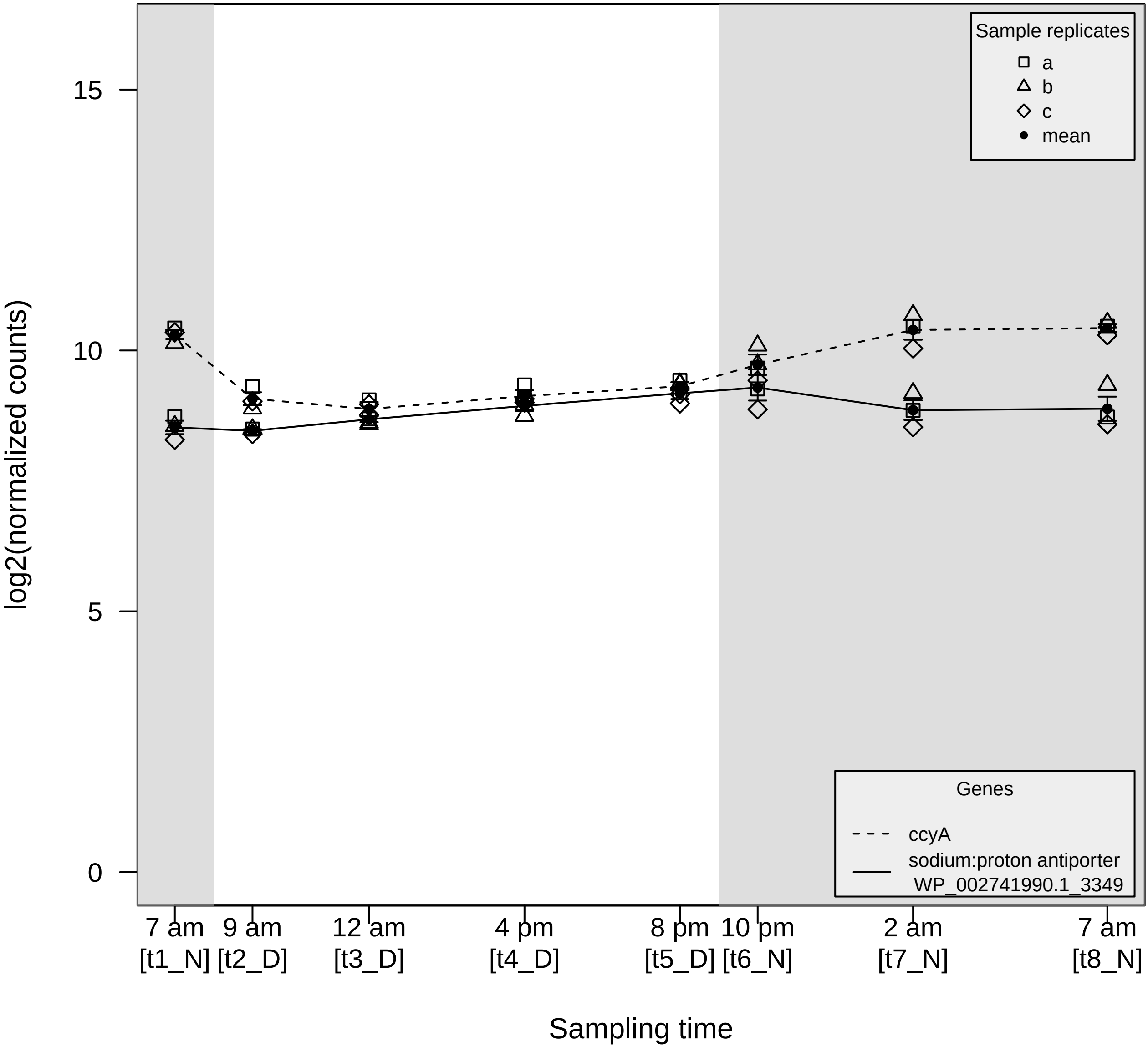

Sample replicates
a
b
c
mean
15
10
5
0
log2(normalized counts)
Genes
ccyA
sodium:proton antiporter
WP_002741990.1_3349
7 am
[t1_N]
9 am
[t2_D]
12 am
[t3_D]
4 pm
[t4_D]
8 pm
[t5_D]
10 pm
[t6_N]
2 am
[t7_N]
7 am
[t8_N]
Sampling time

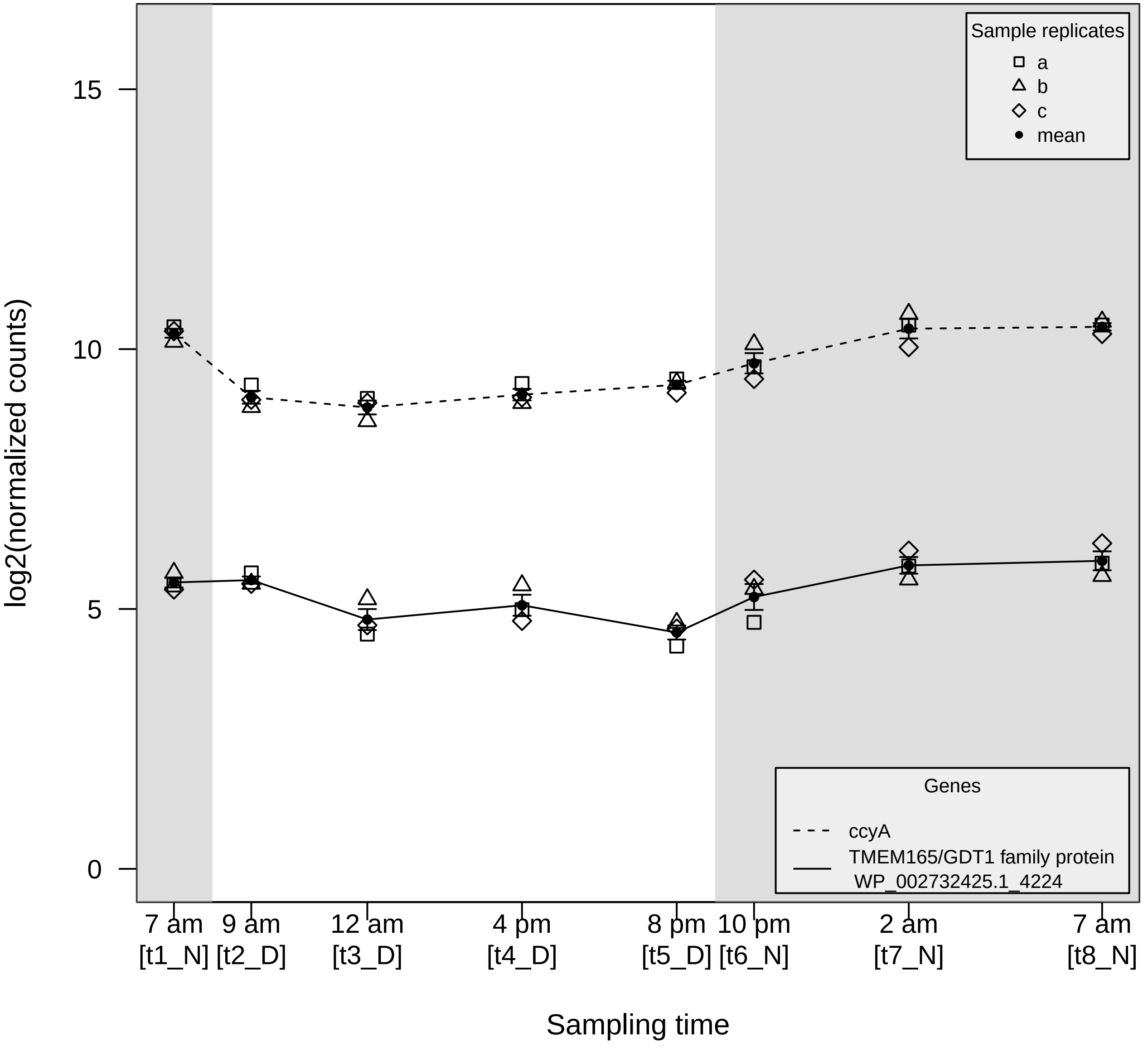

Sample replicates
a
b
c
mean
Genes
ccyA
TMEM165/GDT1 family protein
WP_002732425.1_4224
log2(normalized counts)
15
10
5
0
7 am [t1_N]
9 am [t2_D]
12 am [t3_D]
4 pm [t4_D]
8 pm [t5_D]
10 pm [t6_N]
2 am [t7_N]
7 am [t8_N]
Sampling time

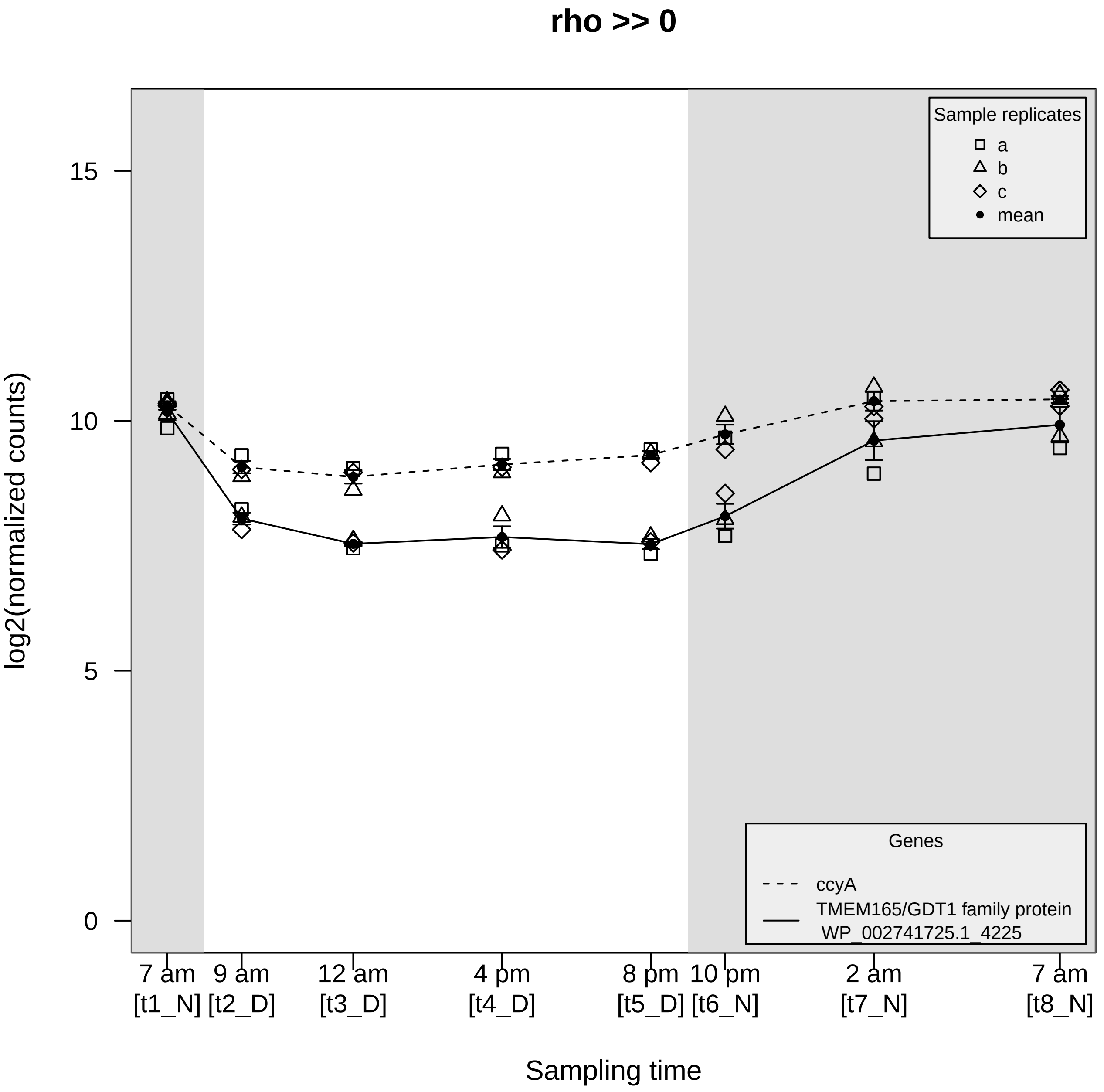
rho >> 0
Sample replicates
a
b
c
mean
Genes
ccyA
TMEM165/GDT1 family protein
WP_002741725.1_4225
log2(normalized counts)
15
10
5
0
7 am [t1_N]
9 am [t2_D]
12 am [t3_D]
4 pm [t4_D]
8 pm [t5_D]
10 pm [t6_N]
2 am [t7_N]
7 am [t8_N]
Sampling time

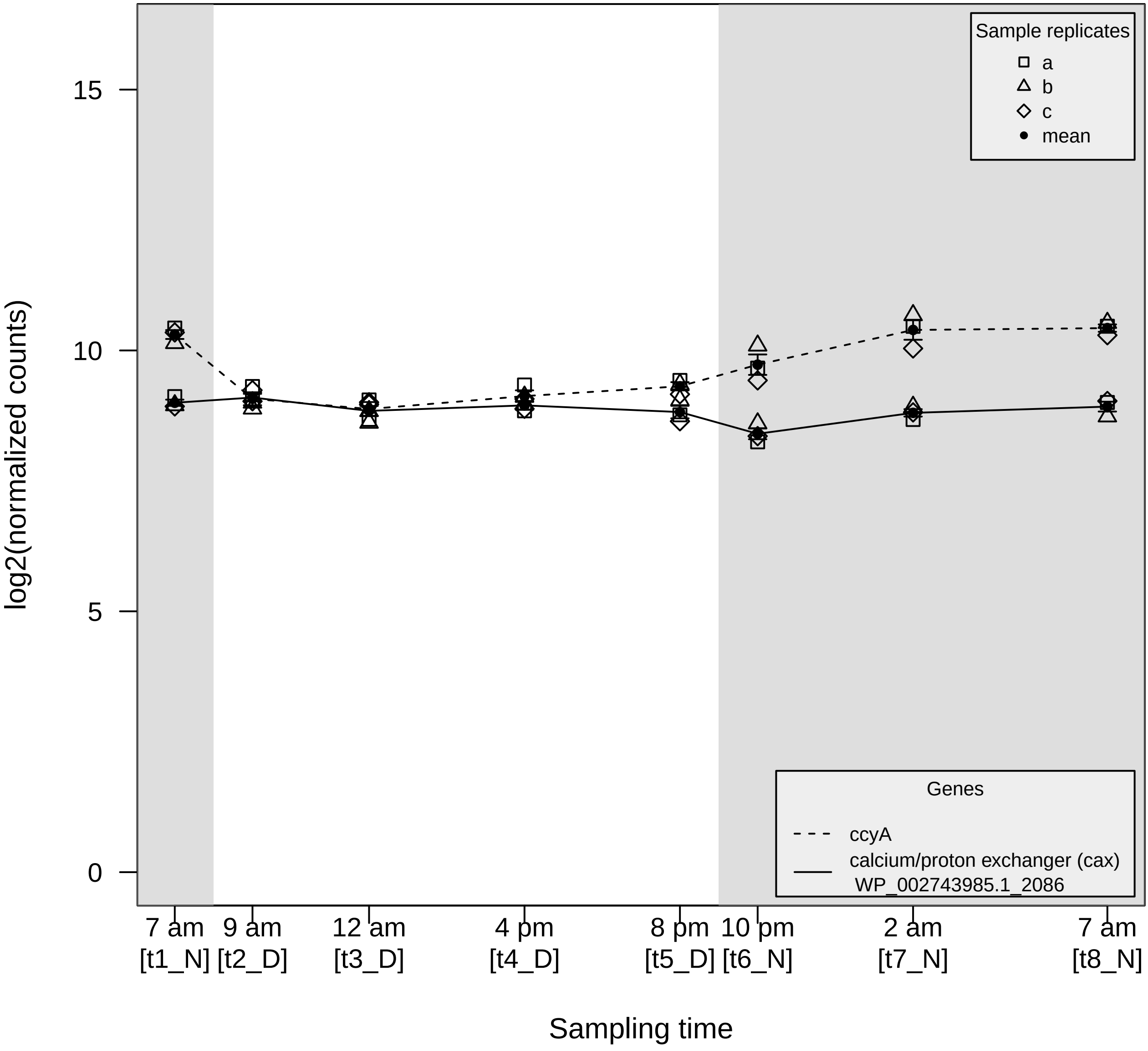

log2(normalized counts)
15
10
5
0
Sample replicates
a
b
c
mean
Genes
ccyA
calcium/proton exchanger (cax)
WP_002743985.1_2086
7 am
[t1_N]
9 am
[t2_D]
12 am
[t3_D]
4 pm
[t4_D]
8 pm
[t5_D]
10 pm
[t6_N]
2 am
[t7_N]
7 am
[t8_N]
Sampling time

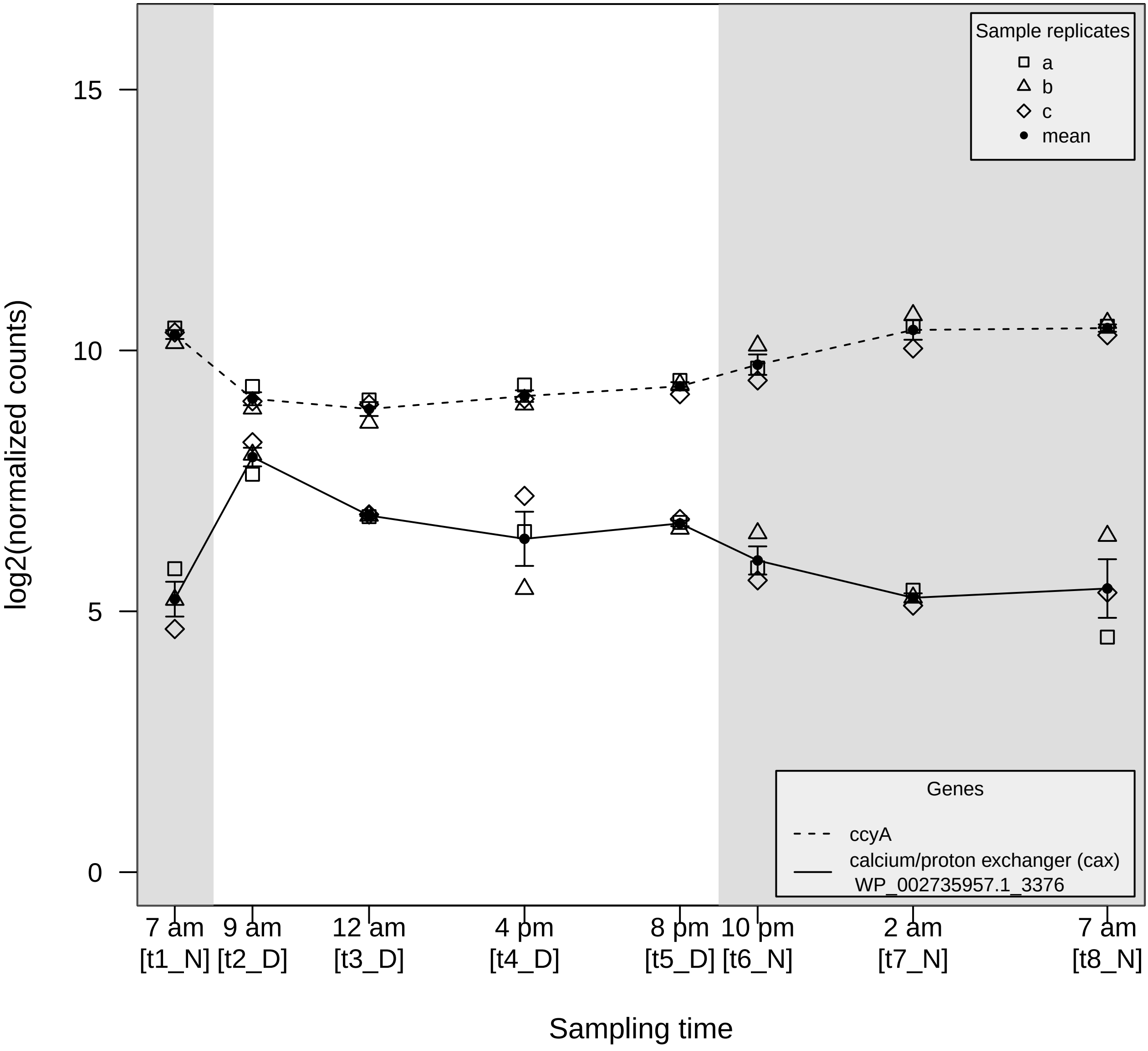

Sample replicates
a
b
c
mean
Genes
ccyA
calcium/proton exchanger (cax)
WP_002735957.1_3376
log2(normalized counts)
0
5
10
15
7 am [t1_N]
9 am [t2_D]
12 am [t3_D]
4 pm [t4_D]
8 pm [t5_D]
10 pm [t6_N]
2 am [t7_N]
7 am [t8_N]
Sampling time

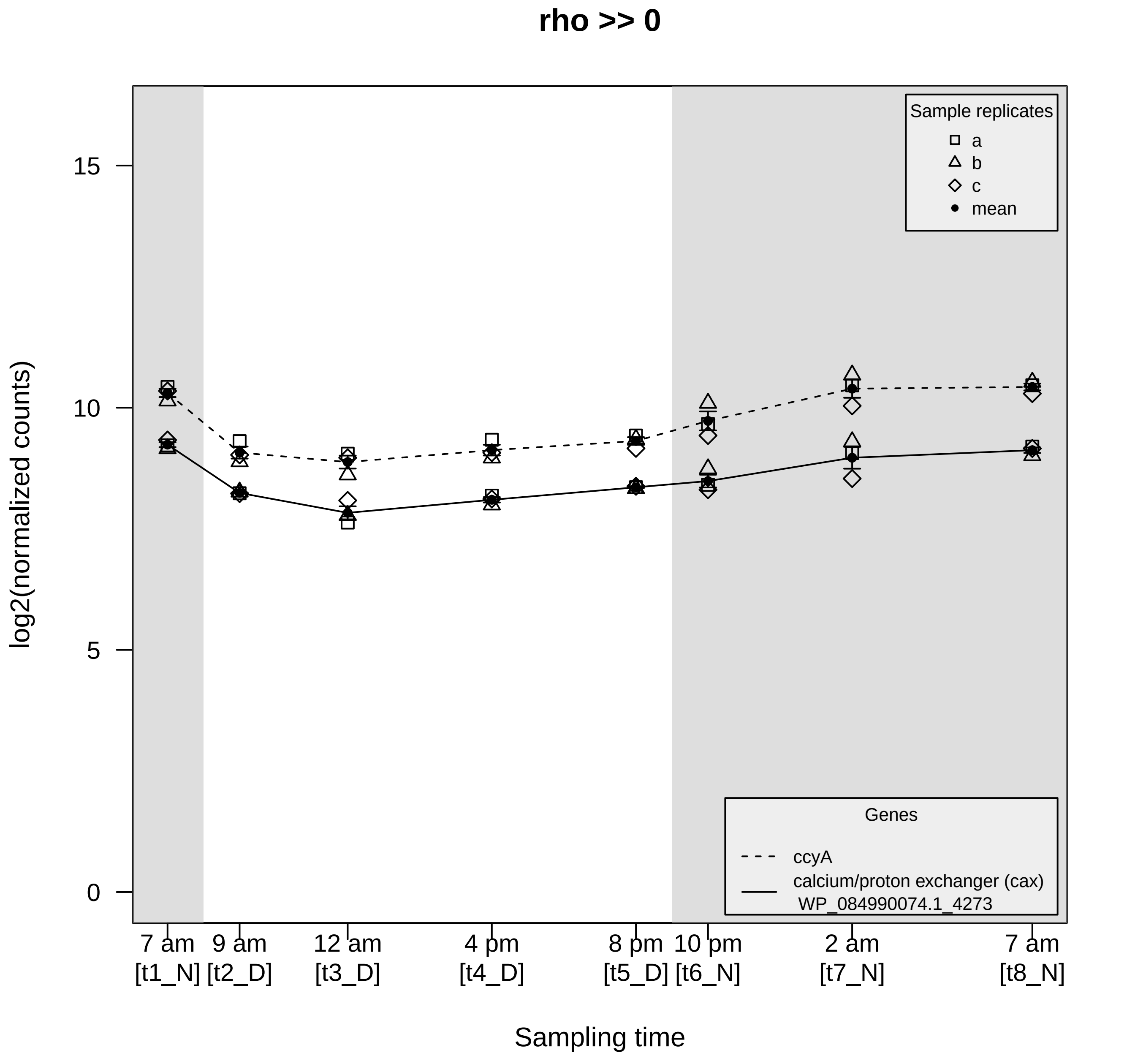
rho >> 0
Sample replicates
a
b
c
mean
Genes
ccyA
calcium/proton exchanger (cax)
WP_084990074.1_4273
log2(normalized counts)
15
10
5
0
7 am [t1_N]
9 am [t2_D]
12 am [t3_D]
4 pm [t4_D]
8 pm [t5_D]
10 pm [t6_N]
2 am [t7_N]
7 am [t8_N]
Sampling time

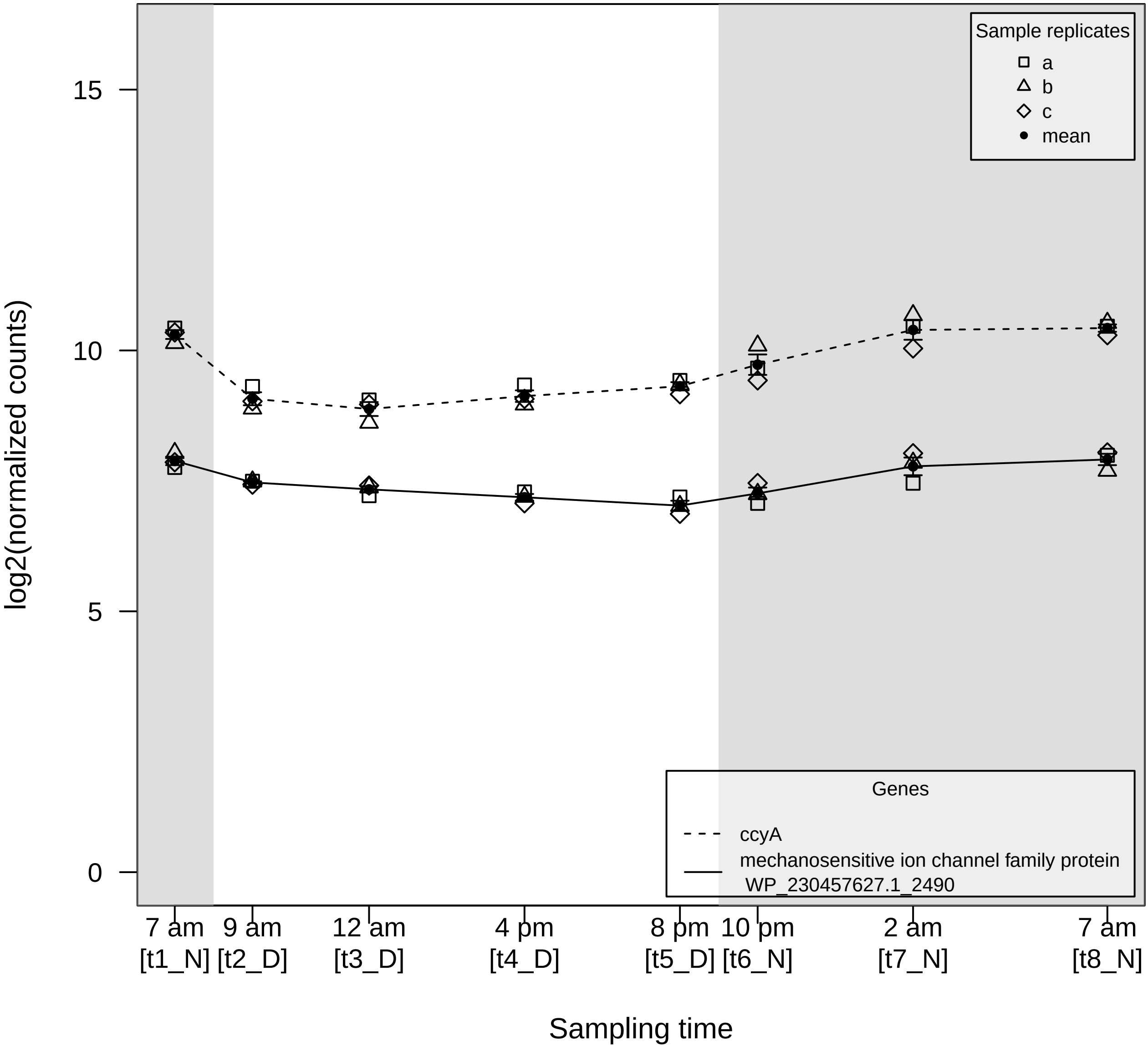

Sample replicates
a
b
c
mean
Genes
ccyA
mechanosensitive ion channel family protein
WP_230457627.1_2490
log2(normalized counts)
15
10
5
0
7 am [t1_N]
9 am [t2_D]
12 am [t3_D]
4 pm [t4_D]
8 pm [t5_D]
10 pm [t6_N]
2 am [t7_N]
7 am [t8_N]
Sampling time

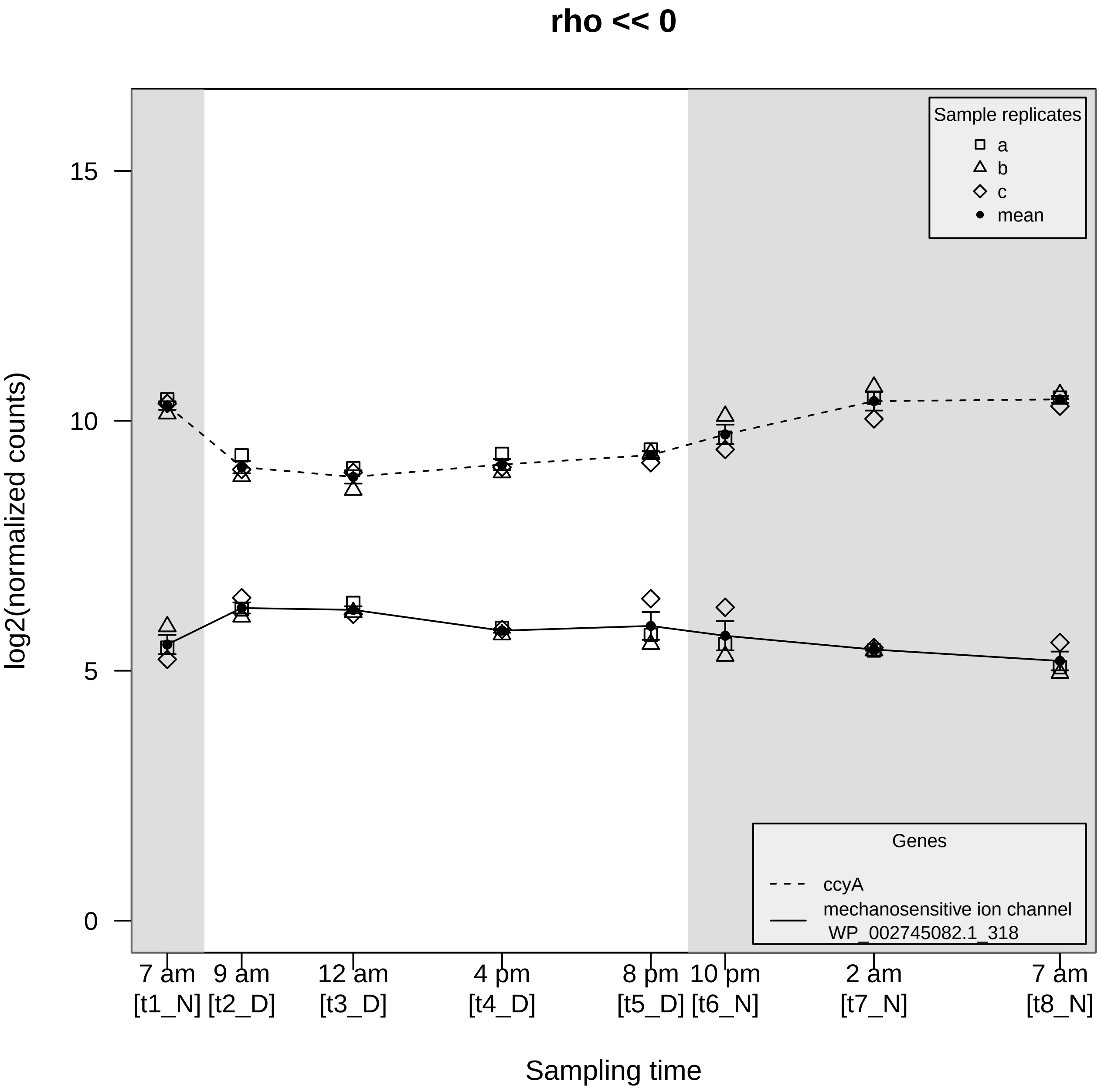

rho << 0
log2(normalized counts)
0
5
10
15
Sample replicates
a
b
c
mean
Genes
ccyA
mechanosensitive ion channel
WP_002745082.1_318
7 am
[t1_N]
9 am
[t2_D]
12 am
[t3_D]
4 pm
[t4_D]
8 pm
[t5_D]
10 pm
[t6_N]
2 am
[t7_N]
7 am
[t8_N]
Sampling time

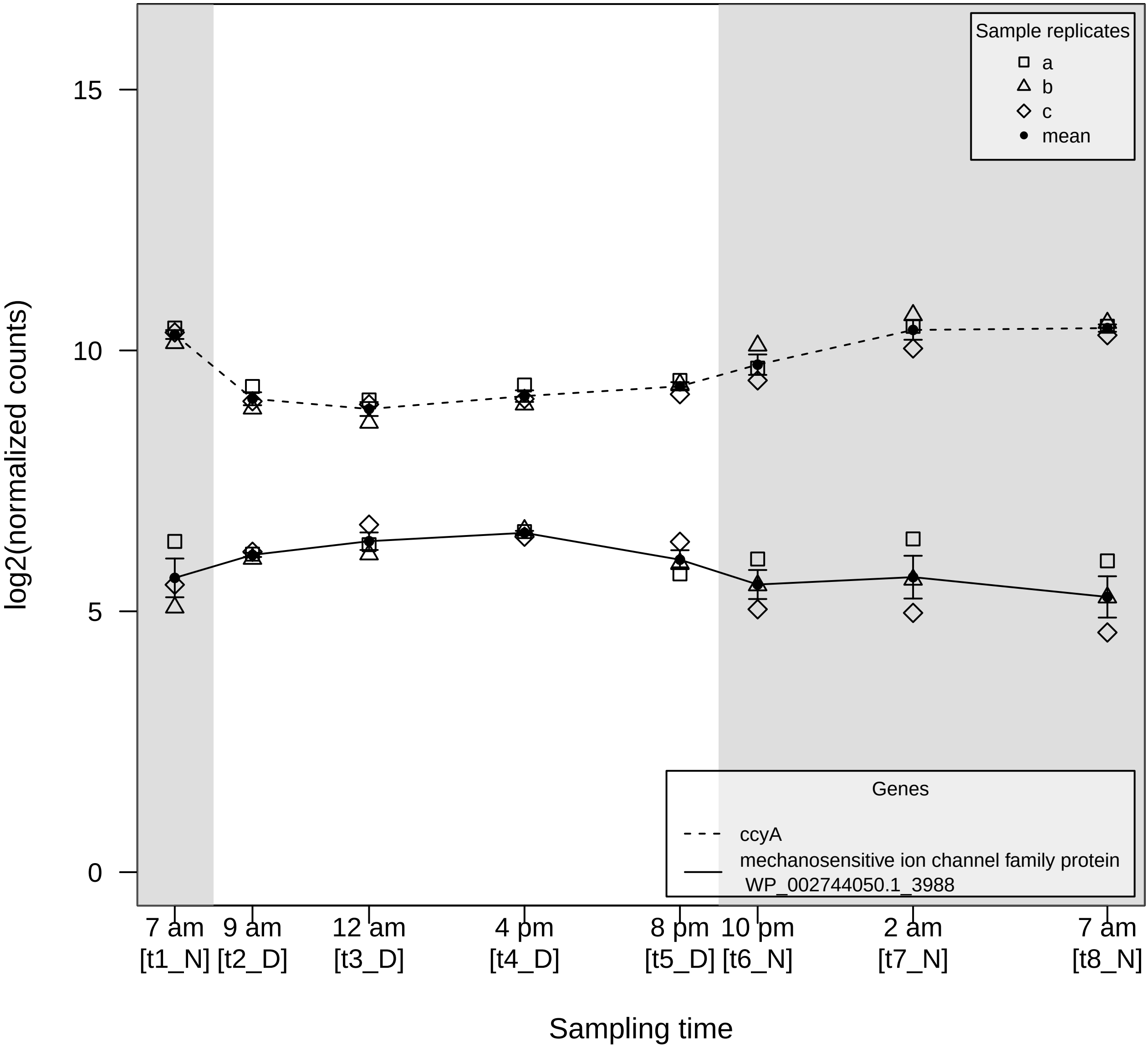

Sample replicates
a
b
c
mean
Genes
ccyA
mechanosensitive ion channel family protein
WP_002744050.1_3988
log2(normalized counts)
15
10
5
0
7 am
[t1_N]
9 am
[t2_D]
12 am
[t3_D]
4 pm
[t4_D]
8 pm
[t5_D]
10 pm
[t6_N]
2 am
[t7_N]
7 am
[t8_N]
Sampling time

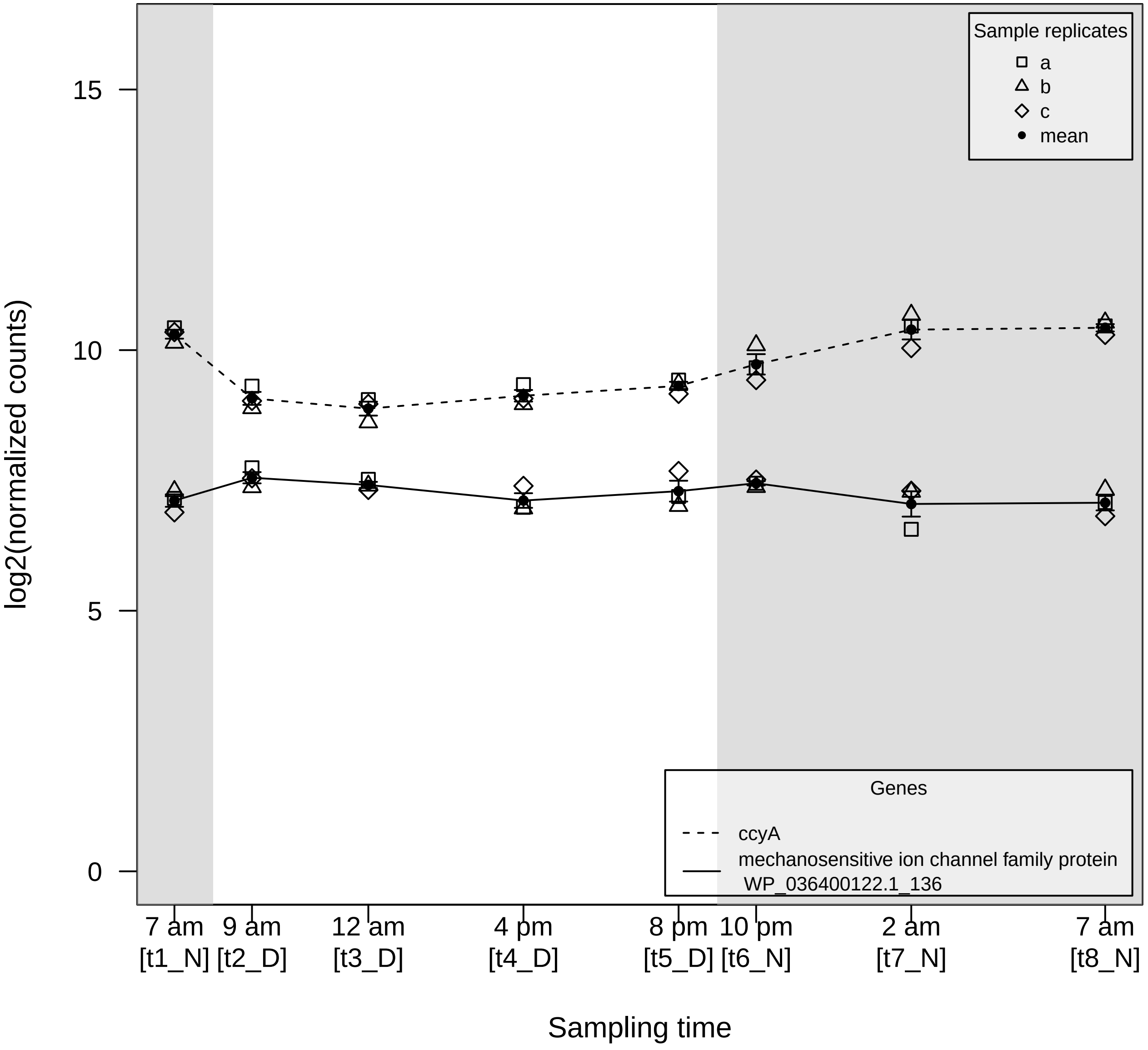

Sample replicates
a
b
c
mean
15
10
5
0
log2(normalized counts)
Genes
ccyA
mechanosensitive ion channel family protein
WP_036400122.1_136
7 am [t1_N]
9 am [t2_D]
12 am [t3_D]
4 pm [t4_D]
8 pm [t5_D]
10 pm [t6_N]
2 am [t7_N]
7 am [t8_N]
Sampling time

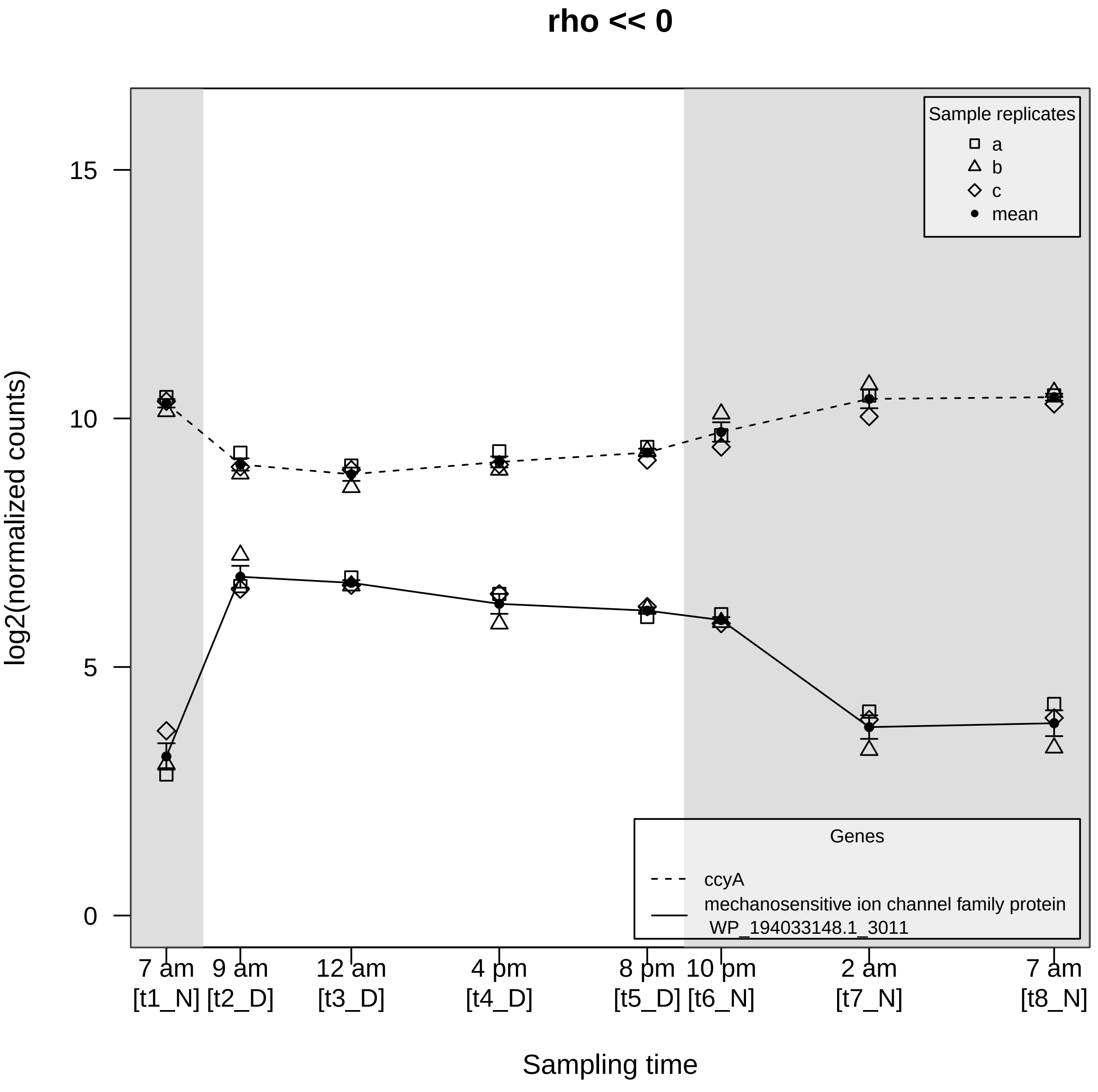

rho << 0
Sample replicates
a
b
c
mean
Genes
ccyA
mechanosensitive ion channel family protein
WP_194033148.1_3011
log2(normalized counts)
15
10
5
0
7 am [t1_N]
9 am [t2_D]
12 am [t3_D]
4 pm [t4_D]
8 pm [t5_D]
10 pm [t6_N]
2 am [t7_N]
7 am [t8_N]
Sampling time

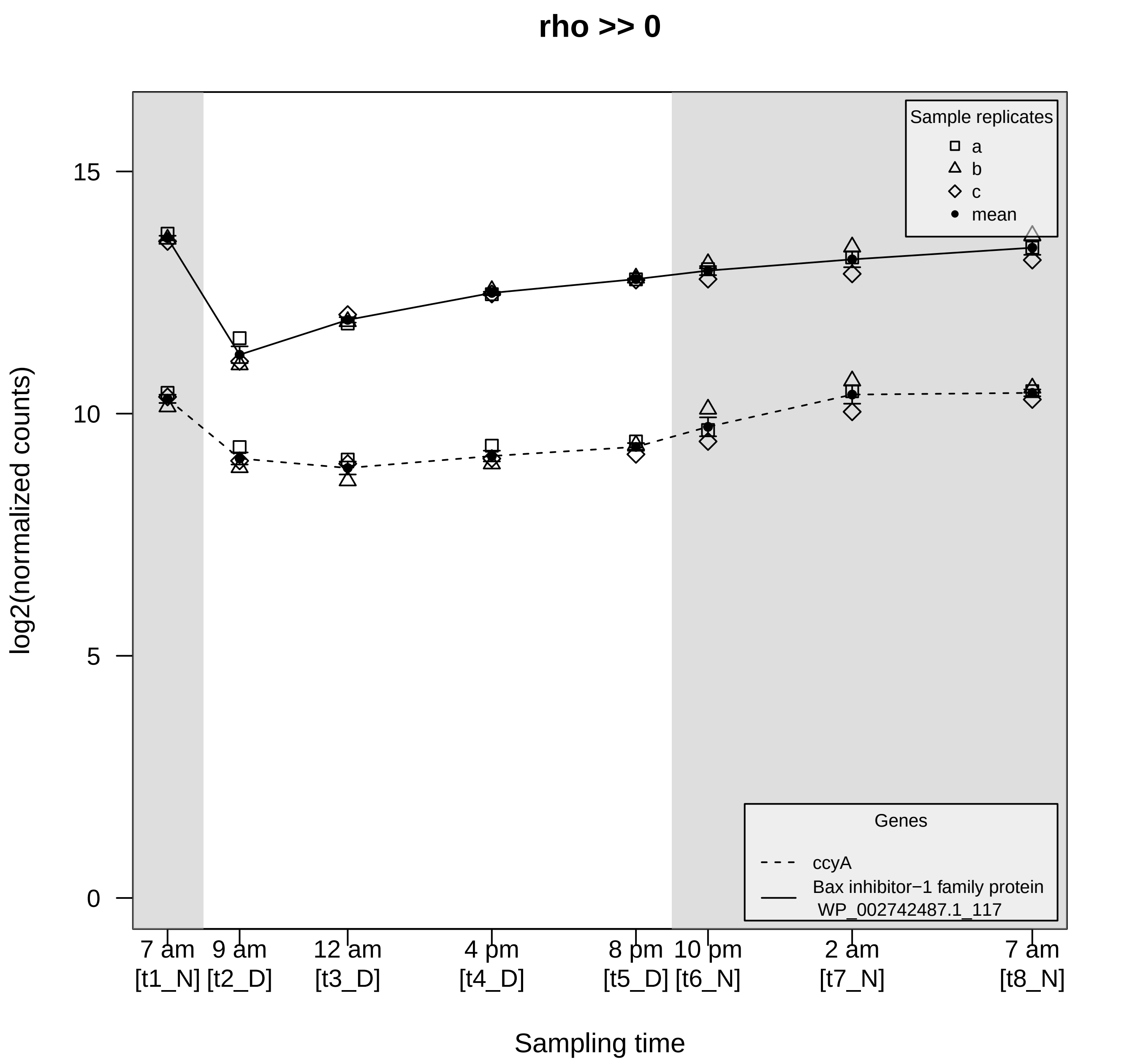

rho >> 0
Sample replicates
a
b
c
mean
Genes
ccyA
Bax inhibitor−1 family protein
WP_002742487.1_117
log2(normalized counts)
15
10
5
0
7 am [t1_N]
9 am [t2_D]
12 am [t3_D]
4 pm [t4_D]
8 pm [t5_D]
10 pm [t6_N]
2 am [t7_N]
7 am [t8_N]
Sampling time
